# Assignment of coronavirus spike protein site-specific glycosylation using GlycReSoft

**DOI:** 10.1101/2020.05.31.125302

**Authors:** Joshua A. Klein, Joseph Zaia

## Abstract

Widely-available LC-MS instruments and methods allow users to acquire glycoproteomics data. Complex glycans, however, add a dimension of complexity to the data analysis workflow. In a sense, complex glycans are post-translationally modified post-translational modifications, reflecting a series of biosynthetic reactions in the secretory pathway that are spatially and temporally regulated. One problem is that complex glycan is micro-heterogeneous, multiplying the complexity of the proteome. Another is that glycopeptide glycans undergo dissociation during tandem MS that must be considered for tandem MS interpretation algorithms and quantitative tools. Fortunately, there are a number of algorithmic tools available for analysis of glycoproteomics LC-MS data. We summarize the principles for glycopeptide data analysis and show use of our GlycReSoft tool to analyze SARS-CoV-2 spike protein site-specific glycosylation.

## Introduction

The analysis of glycopeptides from glycoprotein digests using liquid chromatography-mass spectrometry (LC-MS) is well established [1-9]. As with many protein post-translational modifications, the depth and sensitivity of glycopeptide analysis is highest when an enrichment step is used [5, 6, 10-22]. Glycopeptide LC-MS methods provide maximal dynamic range but require specialized processing steps (Figure 1) to account for glycopeptide heterogeneity and glycosidic bond dissociation [23, 24]. In this review, we summarize bioinformatics methods for processing glycopeptide LC-MS data.

**Figure 1.**
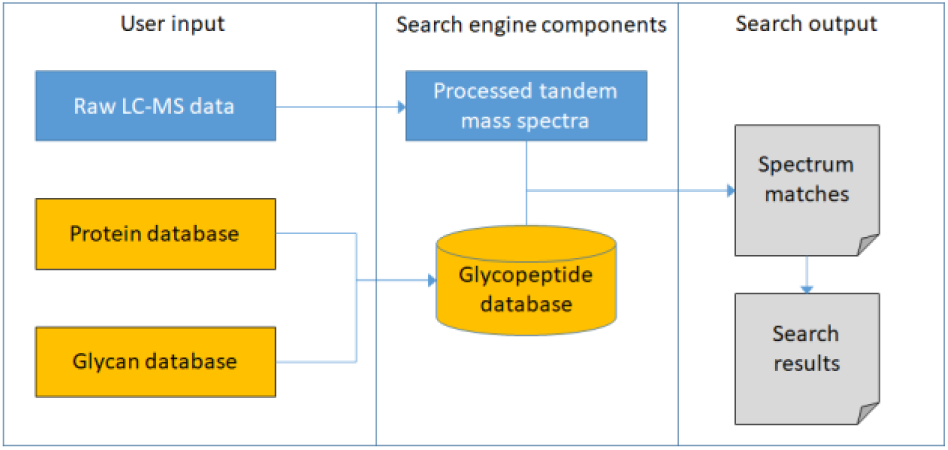
Diagram of a glycoproteomics search engine including inputs, search engine components and outputs.

### Glycopeptide deconvolution

In proteomics, in order to assign the neutral mass of a molecule, it is necessary to convert the raw data from the *m/z* space to the neutral mass space. For unmodified peptides, the elemental composition is approximated using an average amino acid (averagine) to allow estimation of the protein composition [25]. For glycopeptides, it is necessary to adjust the averagine value to include glycosylation. Tryptic glycopeptides tend to be observed over a larger *m/z* and charge state range (2+ to 9+) than typical tryptic peptides (2+ to 4+). In addition, as shown in Figure 2, glycosylation skews the isotopic distribution relative to unmodified peptides. Therefore, specialized deconvolution algorithms are required for glycoproteomics data. SweetNET, a bioinformatics workflow for glycopeptide tandem mass spectral analysis [26] used the MS-DeconV algorithm for spectral deconvolution [27] and the MASCOT [28] for protein identification. The GPQuest glycopeptide spectral library search algorithm [29] used undisclosed isotope pattern fitting and spectral averaging methods for precursor mass calculation. The pGlyco pipeline for identification of glycopeptides from tandem mass spectral data [30] used the pParse algorithm [31], developed by the same group, for deconvolution of precursor and product ions. The GlycoPAT glycoproteomics analysis toolbox [32, 33] deconvolves precursor ions but not product ions. The glyXtool(MS) open*-*source pipeline for semi-automated analysis of glycopeptide mass spectral data [34] uses an OpenMS Feature Finder [35] to calculate precursor ion masses. The GlycReSoft suite of tools for glycomics and glycoproteomics uses an LC-scale deisotoping and charge state deconvolution algorithm for precursor and product ions [36].

**Figure 2.**
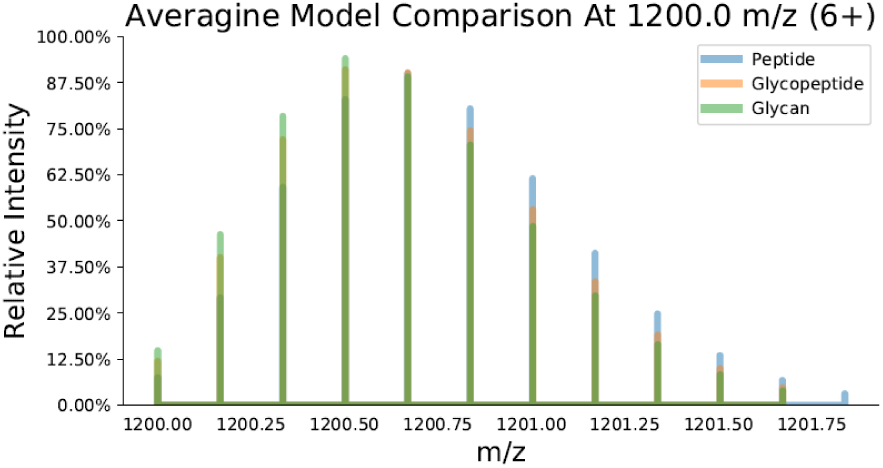
A comparison of three averagine models at *m/z* = 1200 and z = 6+. Note the Glycan model is front-heavy, and the Peptide model is back-heavy, while the Glycopeptide model is balanced between them, as desired.

### Glycopeptide database searching

Glycopeptide identification algorithms use peptide-centric, glycan*-*centric or complete approaches. The peptide-centric method focusses on identifying the peptide backbone sequence, may use peptide + Y ions, but do not control for the false discovery rate of the glycan [37]. By contrast, glycan*-*centric methods [38, 39] identify the attached glycan but do not use peptide backbone dissociation to assign the peptide sequence. Combined methods [40-42] employ a single score that includes both peptide and glycan components and controls the total uncertainty but not the uncertainty of the separate components. Complete methods [43-45] control the uncertainty of glycan and peptide components separately and combined. Some methods use oxonium ions to constrain the range of glycopeptide glycans in a manner that complements use of peptide + Y ions for assigning glycan composition. These approaches assume that there is no ion co-isolation of more than one glycopeptide ion.

A glycoproteomics database search engine includes functions for (i) search space construction, (ii) mass spectrum pre-processing, (iii) a scoring model that evaluates the match between a spectrum and a search space structure, and (iv) a model that evaluates the identification uncertainty for estimation of false discovery rates of glycopeptide sequence matches. The search space uses an input protein list to calculate proteolytic peptides with a list of constant and variable modification rules that include glycosylation. The input protein list may be derived from a FASTA file, an annotated protein sequence format, or an exported proteomics search mzIdentML file. The advantage to using a well-annotated proteome is that the extent of combinatorial expansion of the search space due to inclusion of glycosylation is minimized. There is a degree of subjectivity regarding the makeup of the glycan search space used to construct theoretical glycopeptides. The best practice is to use a measured glycome for this purpose, but this is not always practical. While glycan databases such as GlyTouCan [46] can be used, care must be taken to use the subset of glycans appropriate for the biological system in question. Approaches for estimating glycan search spaces have been described using biosynthetic simulation [47, 48], manual curation [43, 49, 50], and combinatorial expansion [41, 51]. The SweetNET algorithm used a small combinatorial glycan list to extrapolate the set of *N-*glycans, *O*-glycans, and GAG linker saccharides using a spectral network to infer monosaccharide gain/loss in networks of spectra [26].

Glycopeptide tandem MS scoring models depend on the dissociation method and glycopeptide size, meaning that there is no one optimal model that applies to all tandem MS data. For collisional dissociation, collision energy strongly influences the appearance and informational value of glycopeptide tandem mass spectra. As shown in Figure 3, glycopeptide tandem mass spectra contain low *m/z* oxonium ions that act as signatures for glycosylation and high *m/z* ions from loss of monosaccharide units from the precursor ion. Peptide backbone product ions are typically observed only for elevated collision energies. Therefore, use of stepped collision energy has become popular for glycopeptide studies [49]. While electron activated dissociation methods generally favor peptide backbone dissociation over glycosidic bond dissociation, the degree to which vibrational excitation is observed is technique and instrument dependent [52-57].

**Figure 3.**
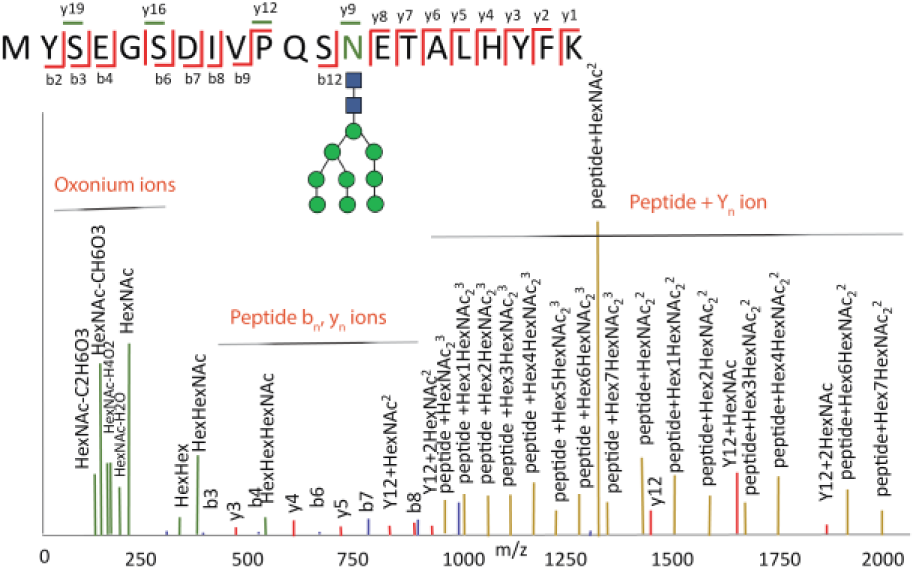
Tandem mass spectrum of an *N-*linked glycopeptide. Peptide backbone product ions are denoted as b_n_ and y_n_. Glycosidic bond cleavage product ions are denoted Y_n_.

As with proteomics of unmodified peptides, empirical models are used to estimate false discovery rate (FDR) for glycopeptides. As in proteomics, glycopeptide data are searched using target decoy analysis [58-60] whereby targets and decoys compete for spectral matching. Some published methods for glycopeptides use structural properties to optimize model performance [30, 33, 44] or employ hierarchical filters [26, 37, 47, 49] to optimize results. For HCD, stepped collision energies most consistently produce peptide+Y ions and peptide b_n_ and y_n_ ions that characterize the glycopeptide glycan and peptide backbone independently [49, 61, 62].

### Glycoproteomics of SARS-CoV-2 spike protein (S)

Whole pathogenic organism vaccines work well against viruses the life cycles of which do not require evasion from the host immune system, including measles, polio, and small pox [63]. By contrast, viruses that have life cycles that depend on the ability to evade the host immune system and have evolved mechanisms that result in suboptimal antibody responses. Immune evasion by molecular mimicry and glycan shielding has been observed and characterized for spike proteins of viruses including HIV-1 envelop protein [64], influenza hemagglutinin [65], Lassa virus glycoprotein complex [66], and corona virus S protein [67].

Glycosylation of the HIV envelope trimer corresponds to about half of its total mass [68]. The dense glycan shield limits the extent of biosynthetic processing, resulting in primarily high mannose *N-*glycans that are thought to interfere with proteolytic processing of envelope peptides for presentation to the major histocompatibility complex [69, 70]. Although studies have identified broadly neutralizing antibodies that recognize the HIV envelope glycan shield, it has not been possible to induces such antibodies in response to vaccine challenge [64]. By contrast, glycosylation of influenza A virus hemagglutinin reflects a balance of immune evasion versus receptor binding. If hemagglutinin glycosylation becomes too dense, it interferes with receptor bniding and/or membrane fusion [71-73].

Four respiratory coronaviruses cause mild, cold-like, symptoms in humans. While most adults have antibodies against these coronaviruses, they have circulated in the human population for centuries [74]. The severe acute respiratory syndrome corona virus (SARS-CoV) zoonotic outbreak in humans was contained within three months after its discovery in 2002. The Middle East respiratory syndrome (MERS) coronavirus has spread zoonotically to humans repeatedly but has so far had limited human*-*to-human spread [75]. By contrast, the SARS-CoV-2 virus jumped from animals to humans in 2019 and caused a global pandemic with incalculable damage to human culture world-wide.

Glycosylation of the SARS-CoV-2 S protein is of interest for development of antiviral strategies that target the virus-angiotensin*-*converting enzyme 2 (ACE2) receptor recognition [76]. The S protein is composed of the amino-terminal receptor binding S1 and carboxy-terminal S2 membrane fusion subunit [77]. Proteolytic cleavage between S1 and S2 is required for receptor binding and membrane fusion [74]. Because antibodies against S1 receptor binding domain have the potential to neutralize the virus, there is interest in using S protein constituents as vaccine candidates [74, 78].

The use of glycan masking and molecular mimicry has been described for human respiratory coronavirus HCoV-NL63 and other coronaviruses [77, 79]. The coronavirus glycan shield was observed to be less dense than that of HIV envelop protein. The most pathogenic coronaviruses (SARS-CoV, MERS and SARS-CoV-2) appear to have S protein trimers able to adopt open and closed conformations [80]. S Protein glycosylation is therefore an important factor to characterize from the point of view of its influences on virus-receptor recognition.

We chose to analyze a published LC-MS data set on SARS-CoV-2 recombinant S protein [81] using our publicly available, open*-*source GlycReSoft program [36]. We show how any biomedical scientist with access to a Windows desktop computer can query publicly available data for S protein site-specific glycosylation.

### Experimental

The site-specific glycosylation of recombinant S protein expressed in human cells was characterized using glycoproteomics liquid-chromatography-mass spectrometry [81]. The authors expressed the pre-fusion S domain with two proline substitutions were used to stabilize the trimer [82]. A “GSAS” substitution at the furin cleavage site and a C-terminal trimerization motif were used to facilitate maintenance of quaternary architecture during glycan processing [83]. They digested separate samples using trypsin, chymotrypsin and alpha-lytic protease, respectively, in order to map glycosylation at all 22 sequons. Size fractionated, reduced, and alkylated S protein was digested with protease and the resulting peptides analyzed using 75 µm internal diameter, 75 cm length reversed phase LC-MS with a 275 min linear gradient. The scan range was 400-1600 and HCD collision energy set to 50%. The instrument was set for top-N data dependent acquisition. A single raw LC-MS data file for each proteolytic enzyme was posted publicly to the MassIVE Database [84].

Glycopeptides were assigned using the GlycReSoft graphical user interface [36] available at http://www.bumc.bu.edu/msr/glycresoft/. Raw files were converted to mzML format using ProteoWizard MSConvert [85] and deconvoluted/deisotoped using the GlycReSoft preprocessing algorithm. A glycan search space was constructed by combining an N-glycan biosynthesis simulation combined with up to one sulfate per glycan composition. A glycopeptide search space was built for each protease using the corresponding mzIdentML or FASTA file and the glycomics search space. Glycopeptides were identified using 0-1 ammonium adducts, with a precursor mass error tolerance of 10 ppm, a product mass error tolerance of 10 ppm. The complete GlycReSoft HTML reports are included as Supplemental Files.

## Results

Total ion chromatograms for the tryptic, chymotryptic and alpha lytic protease digests are shown in Figure 4. The use of a long LC gradient combined with a single HCD collision energy value of 50% maximized the number of glycopeptides that were selected for tandem MS. The proteomics search of the tryptic digest of S protein identified a total of 888 proteins, the top 20 most abundant of which are shown in Figure 5. The S protein was approximately 10-fold more abundant than the next most abundant protein. In order to determine the effect of host proteins on the ability to assign glycopeptides, we compared the GlycReSoft results for a search space constructed using all proteins identified versus that using only SARS-CoV-2 S protein. The results showed that a similar number of glycopeptides were mapped using the complete proteome versus that for the S protein only proteome. This indicated that host proteins did not interfere significantly with the identification of S protein glycosites. We next compared results using no ion adduction versus 0-1 ammonium adduct and 0-1 sodium adduct together. Because the number of glycopeptides identified was similar in both cases, the results demonstrated that there was a low degree ion adduction in the LC-MS runs.

**Figure 4.**
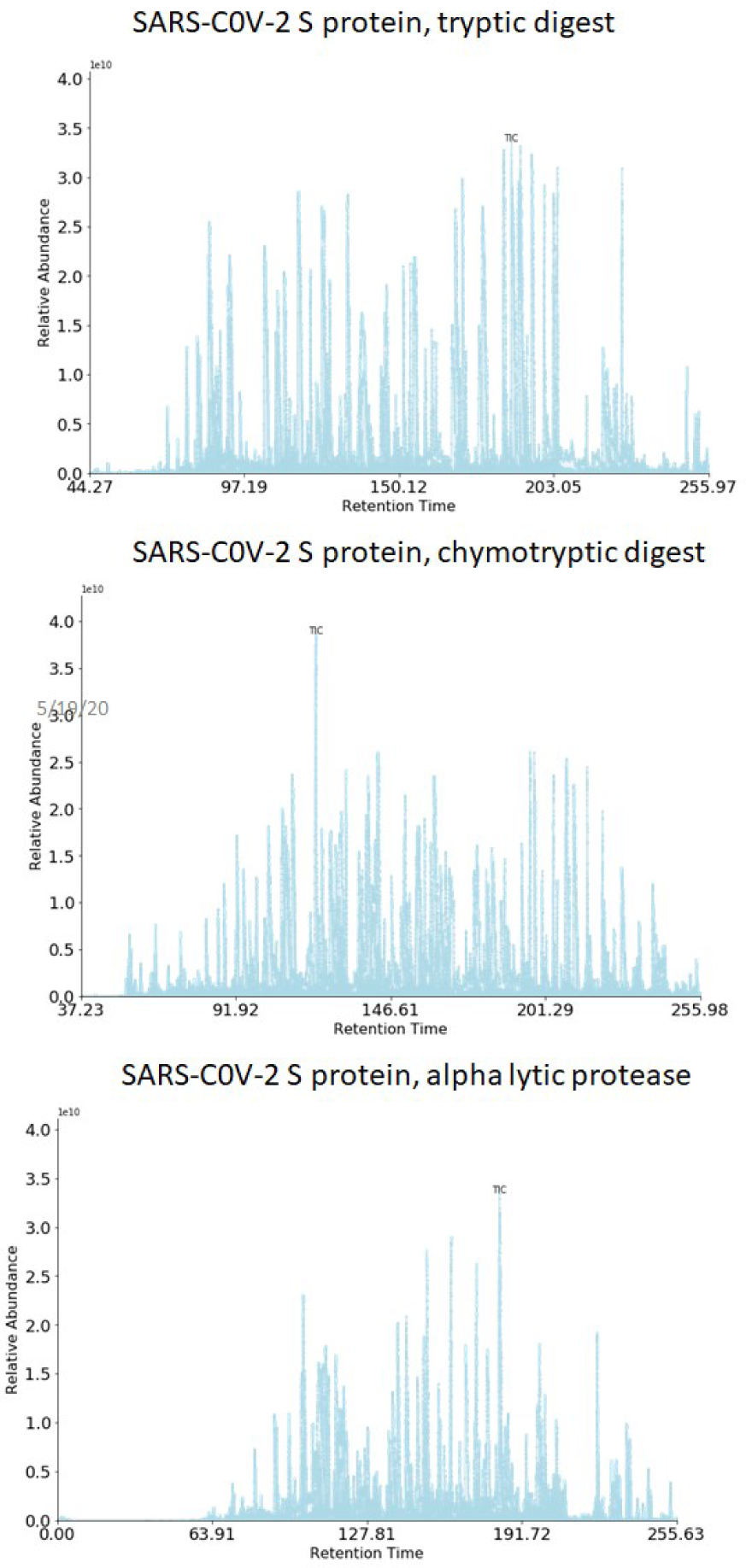
Total ion chromatograms for SARS-CoV-2 S protein proteolytic digests.

**Figure 5.**
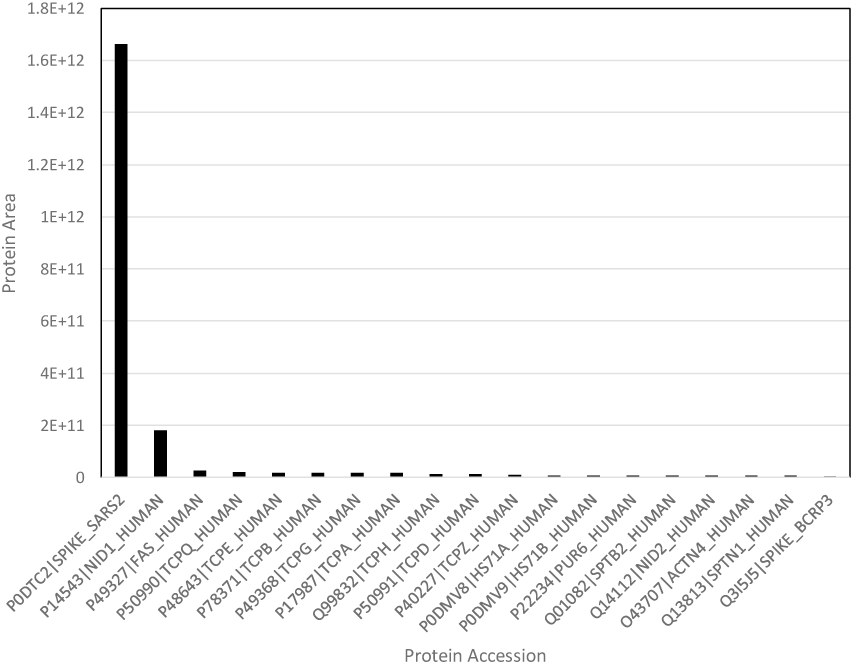
The top 20 most abundant proteins from 888 identified for the SARS-CoV-2 S protein sample from the tryptic digest.

While the glycopeptide tandem mass spectrum shown in Figure 3 was acquired using stepped collision energy, the S protein tandem MS data were acquired using HCD set at 50%. Under these conditions, glycopeptides were extensively fragmented and the abundances of peptide + Y_n_ ions was very low, skewing tandem MS scores to the lower range (see for example Figure 6B). The peptide sequence is identified unambiguously but the lack of peptide+Y_n_ ions limited glycan characterization to intact mass and oxonium ions, leaving core structure unknown. As a balancing factor, it is possible to dissociate more precursor ions when a single collision energy is specified that with stepped collision energy.

**Figure 6.**
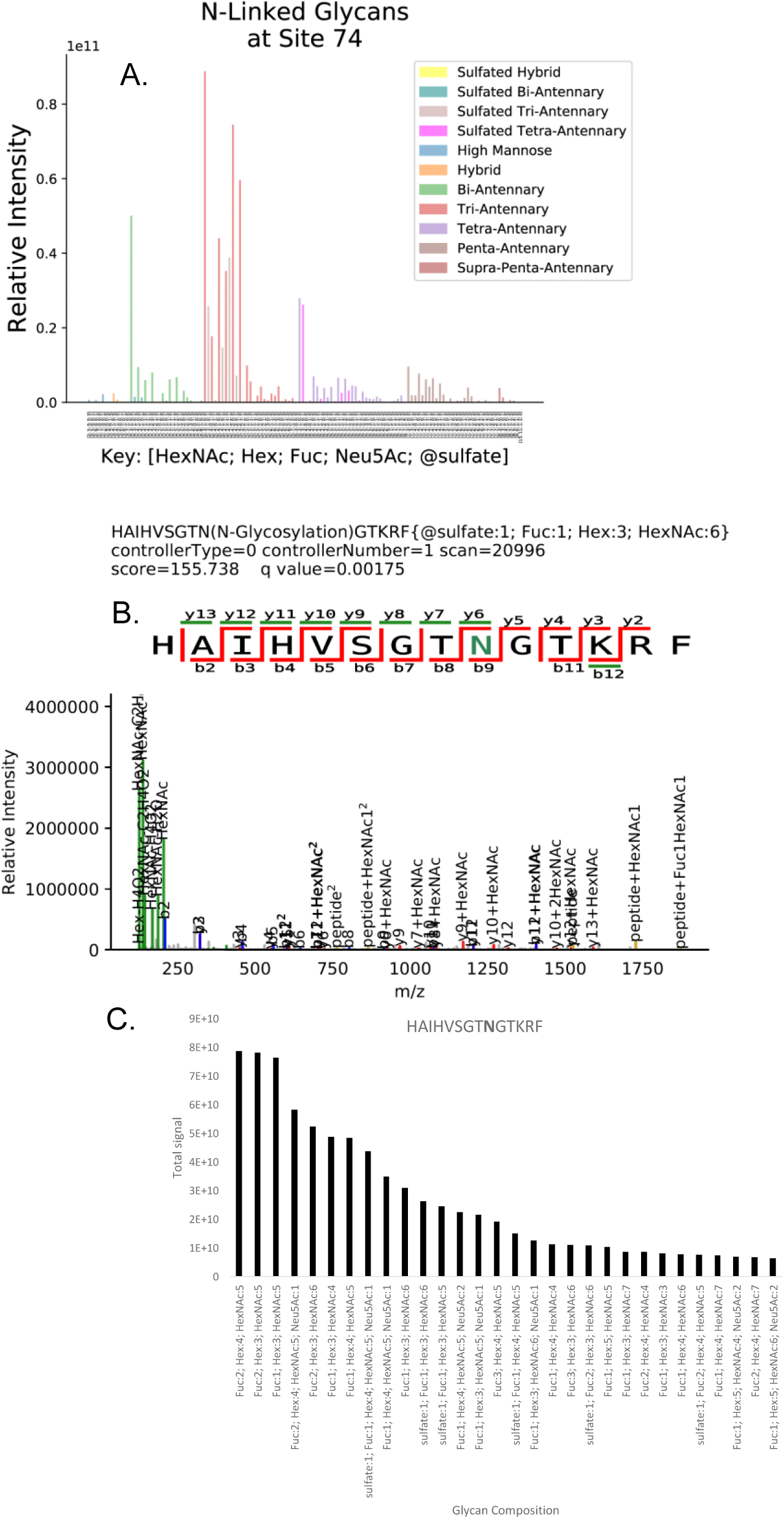
A. Sulfated glycans at position 74 from chymotryptic digest. B. Example annotated tandem mass spectrum. C. Plot of the 30 most abundant glycan compositions at position 74.

We processed LC-MS runs acquired for three proteolytic digests, trypsin, chymotrypsin and alpha lytic protease. The trypsin and alpha-lytic protease search parameters were set to specify one site of glycosylation peptide using a desktop computer using 5 processors. The chymotryptic digest was first considered using only one site of glycosylation per peptide, but the set of identified glycans from that search were used to re-generate the search space allowing up to two sites of glycosylation per peptide for the final reported results, searched on with a shared high performance computing cluster utilizing 16 processors. The glycoforms identified for each glycopeptide are shown Figure 6-Figure 19. The results shown correspond to the enzyme digest that produced the highest glycopeptide abundances for a given glycosite. Overall, the abundances of high mannose, hybrid, and complex *N-*glycan compositions is consistent with those in the original publication [81].

**Figure 7.**
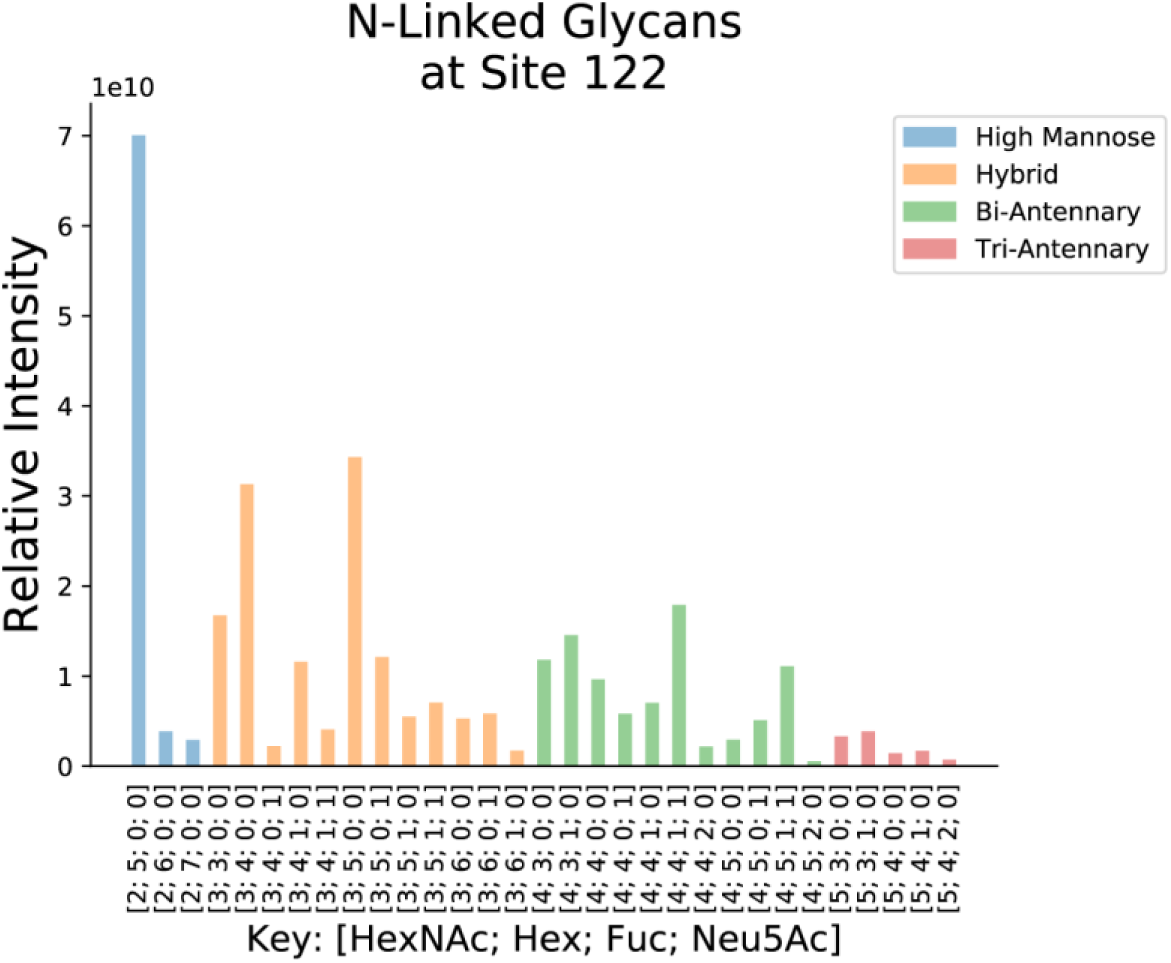
*N-*glycans at site 122 from chymotryptic digest.

**Figure 8.**
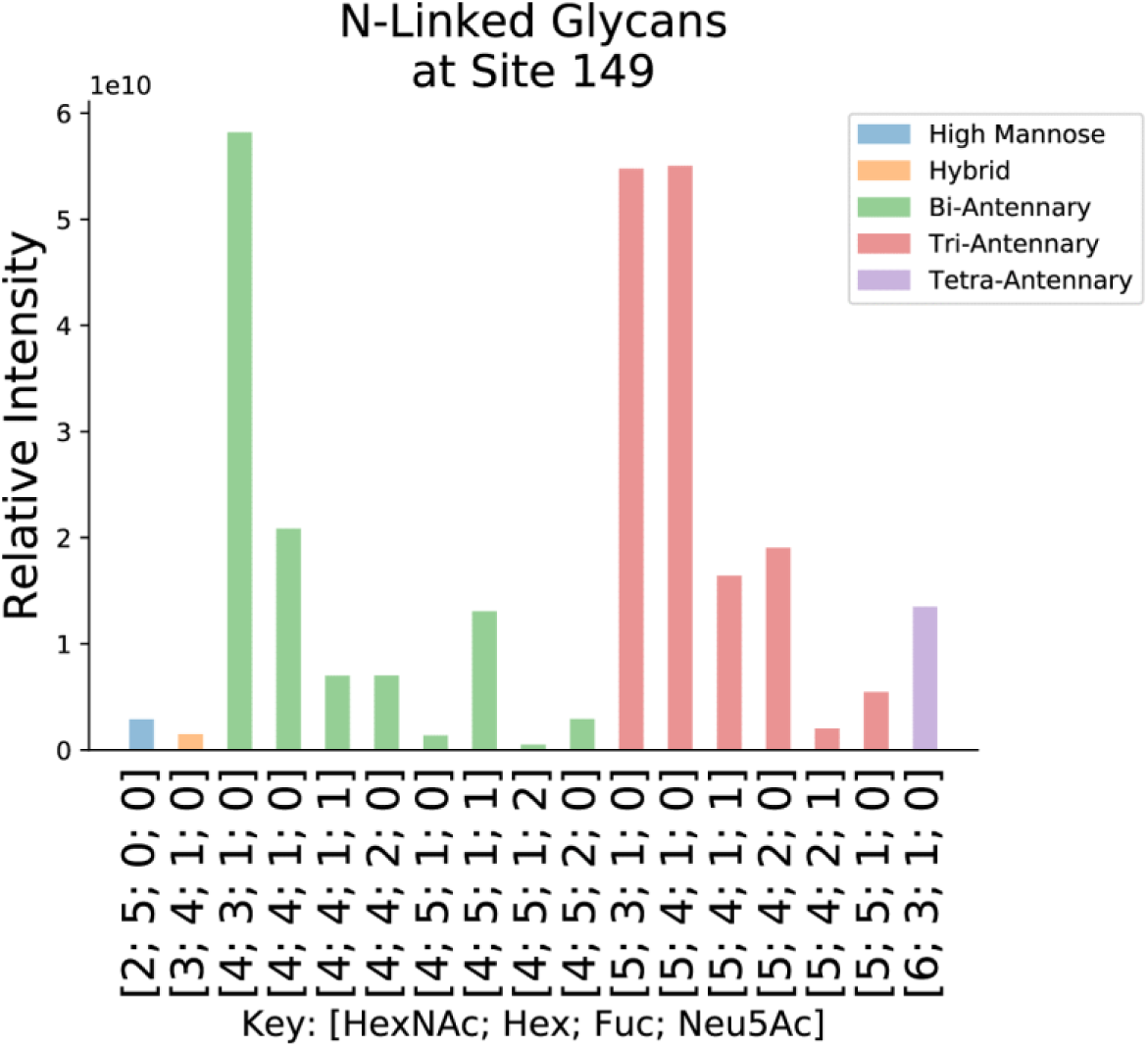
*N-*Glycans at site 149 from chymotryptic digest.

**Figure 9.**
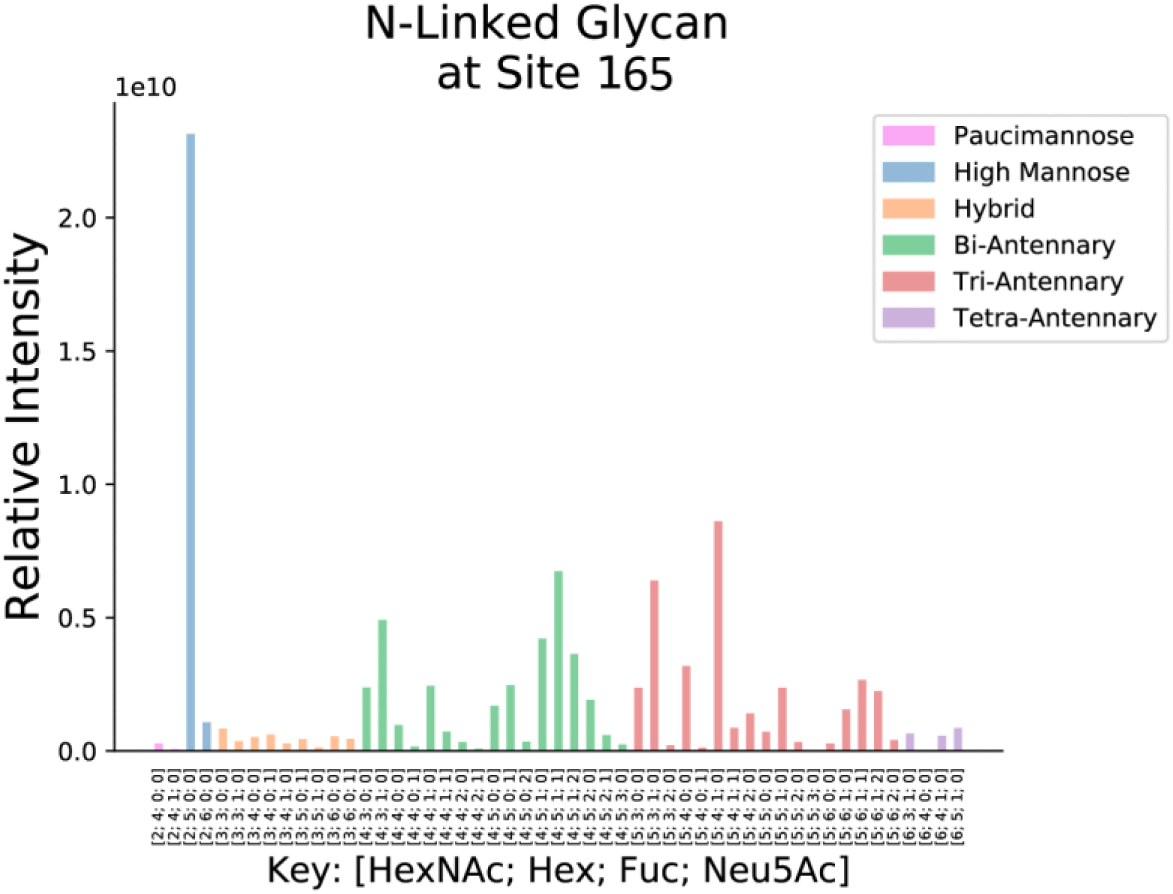
*N-*Glycans at site 165 from tryptic digest.

**Figure 10.**
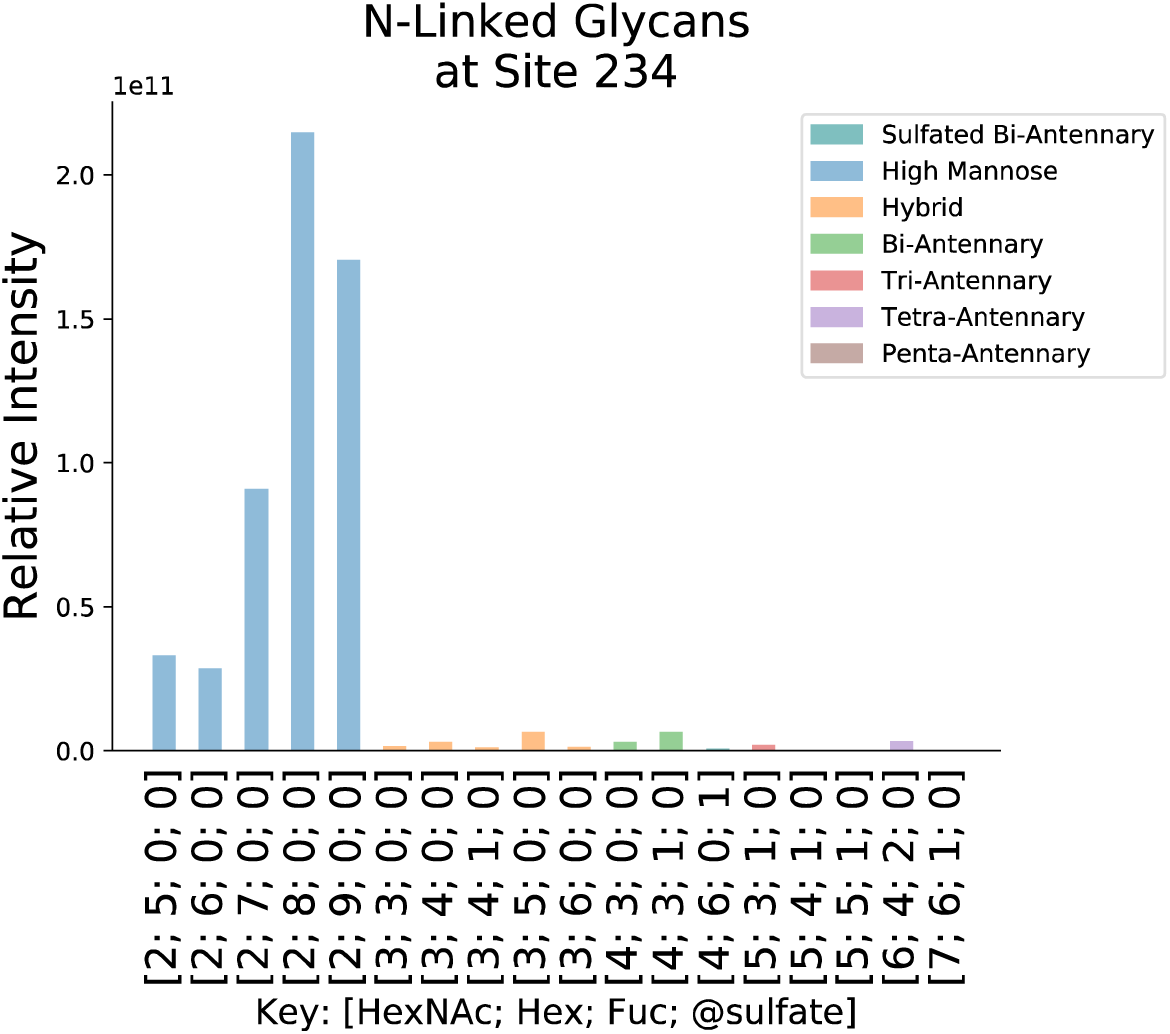
*N-*Glycans at site 234 from chymotryptic digest.

**Figure 11.**
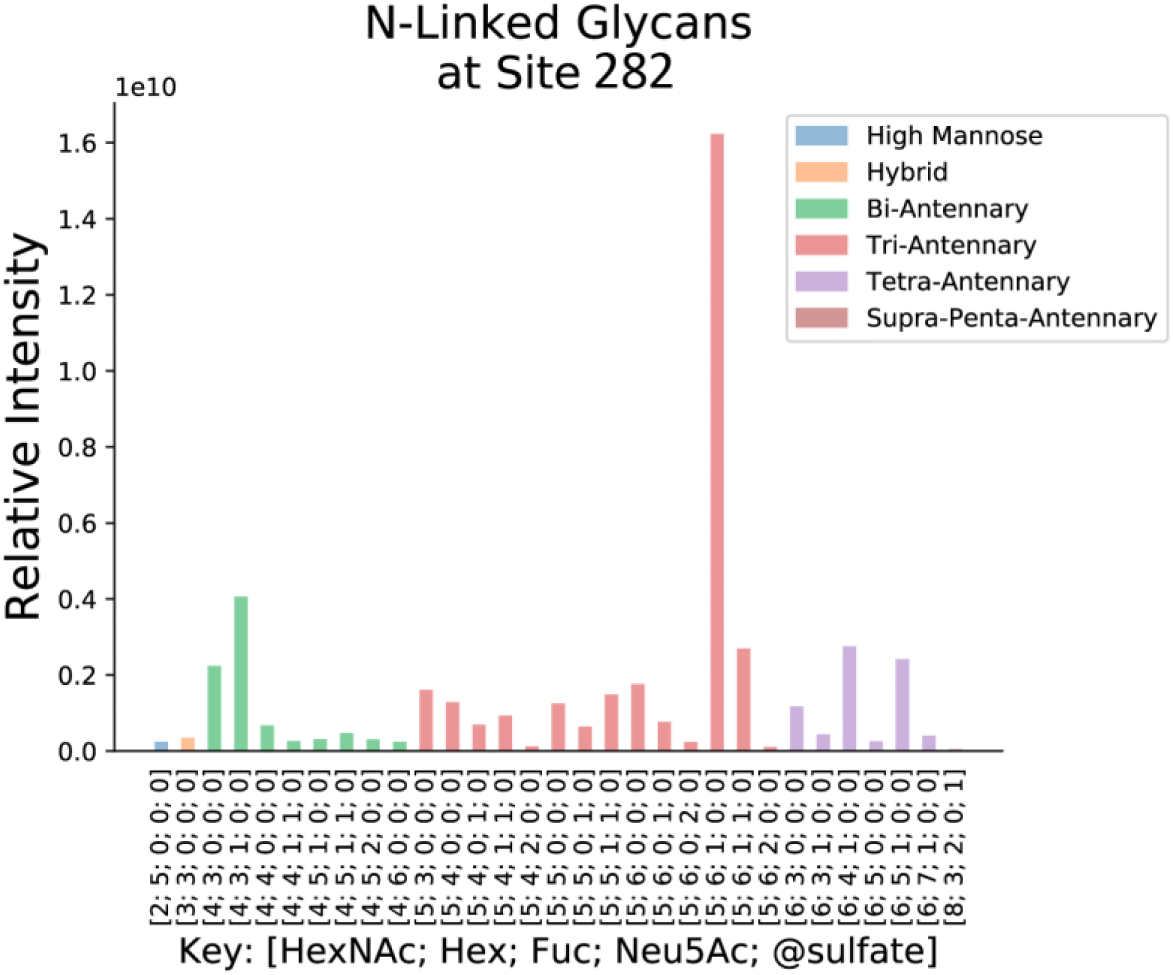
*N-*Glycans at site 282 from tryptic digest.

**Figure 12.**
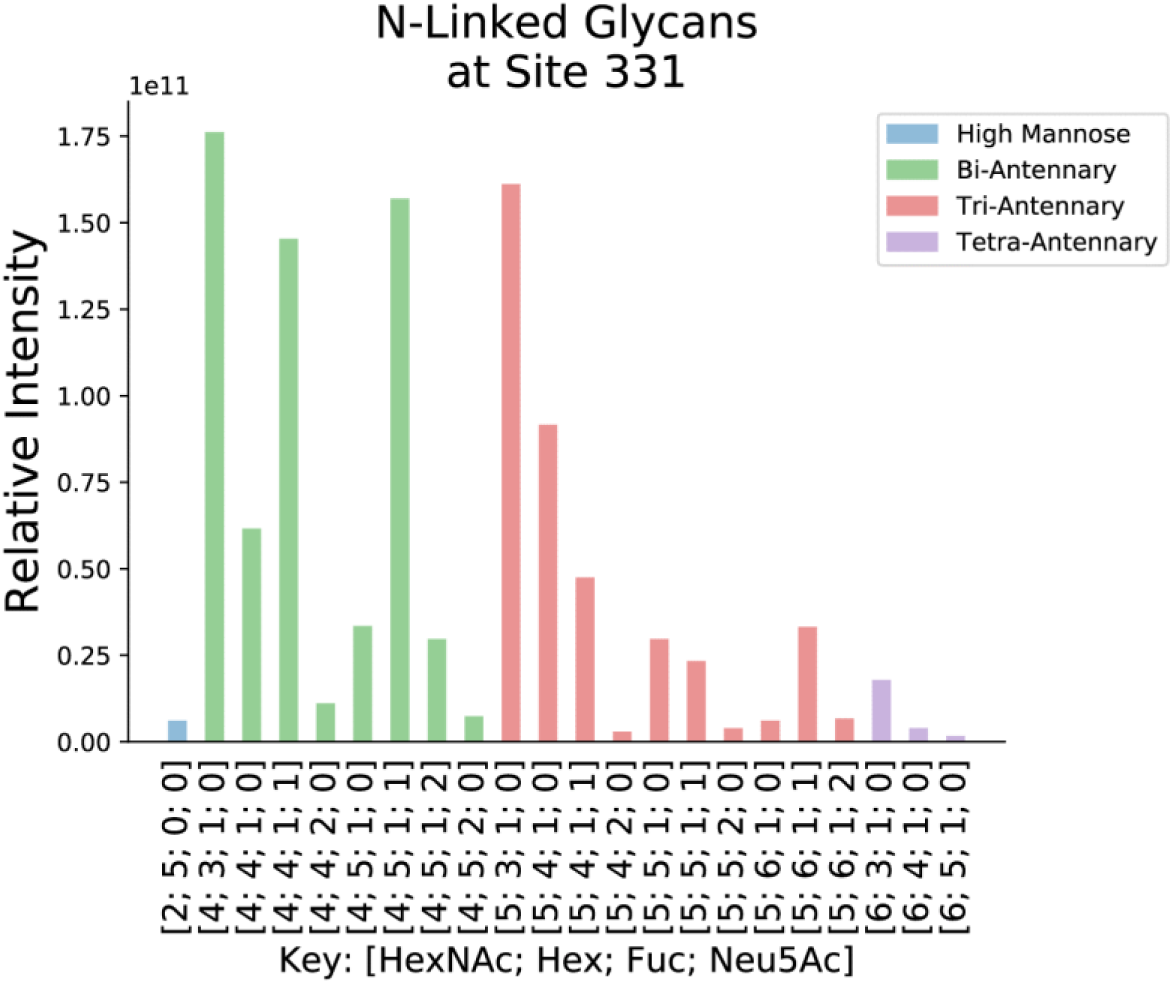
*N-*Glycans at site 331 from chymotryptic digest.

**Figure 13.**
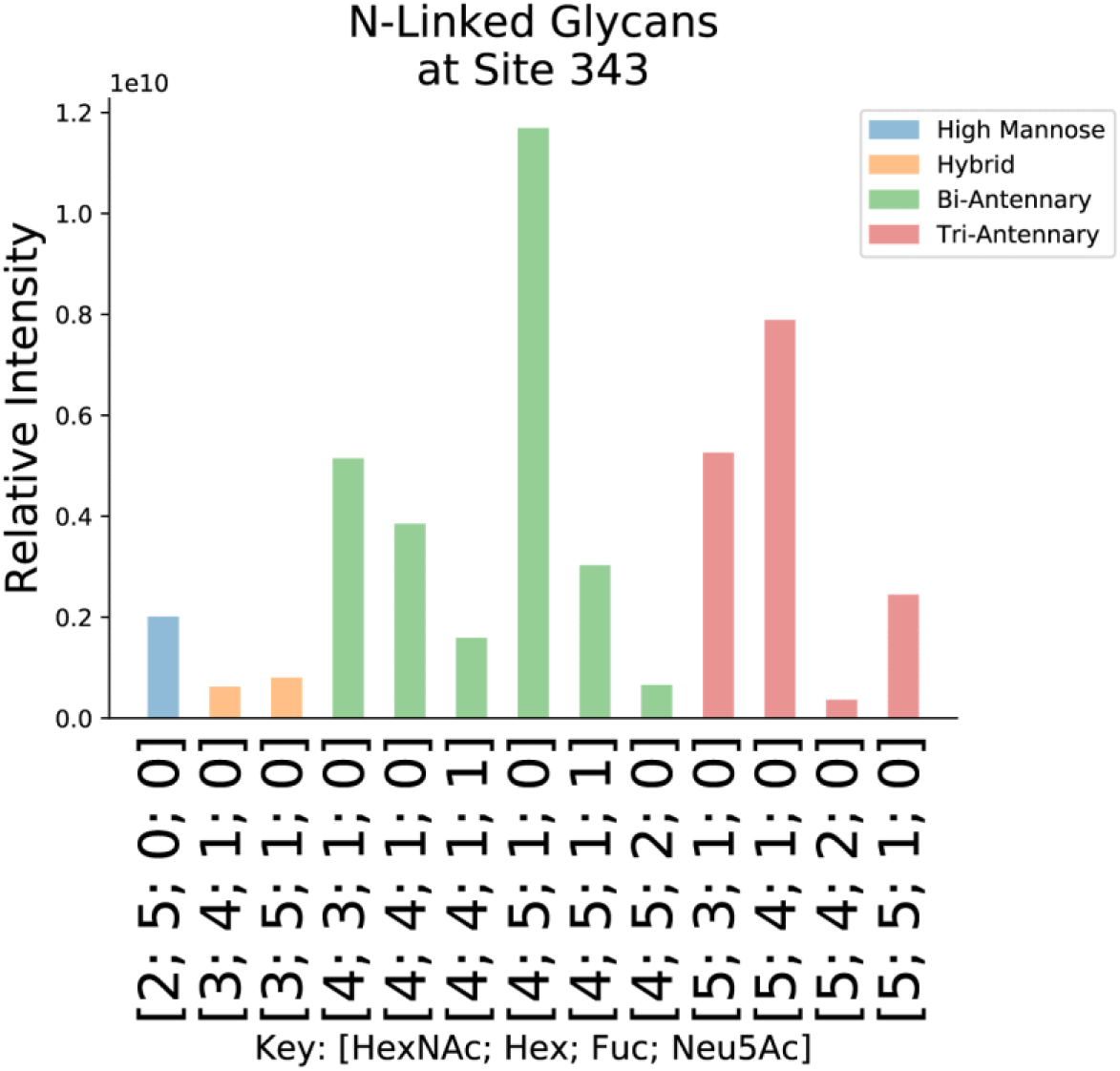
*N-*Glycans at site 343 from chymotryptic search.

**Figure 14.**
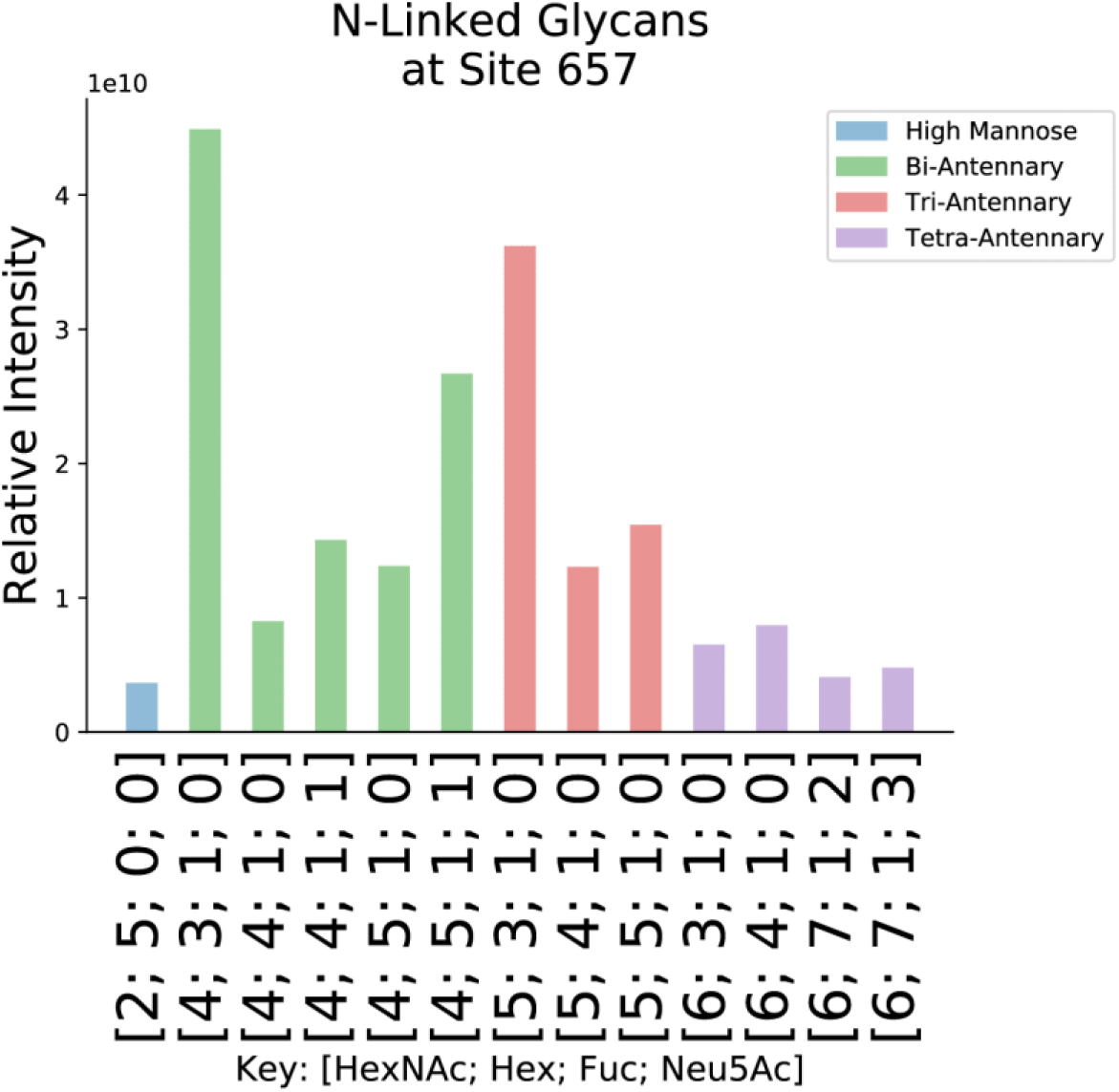
*N-*Glycans at site 657 from chymotryptic digest.

**Figure 15.**
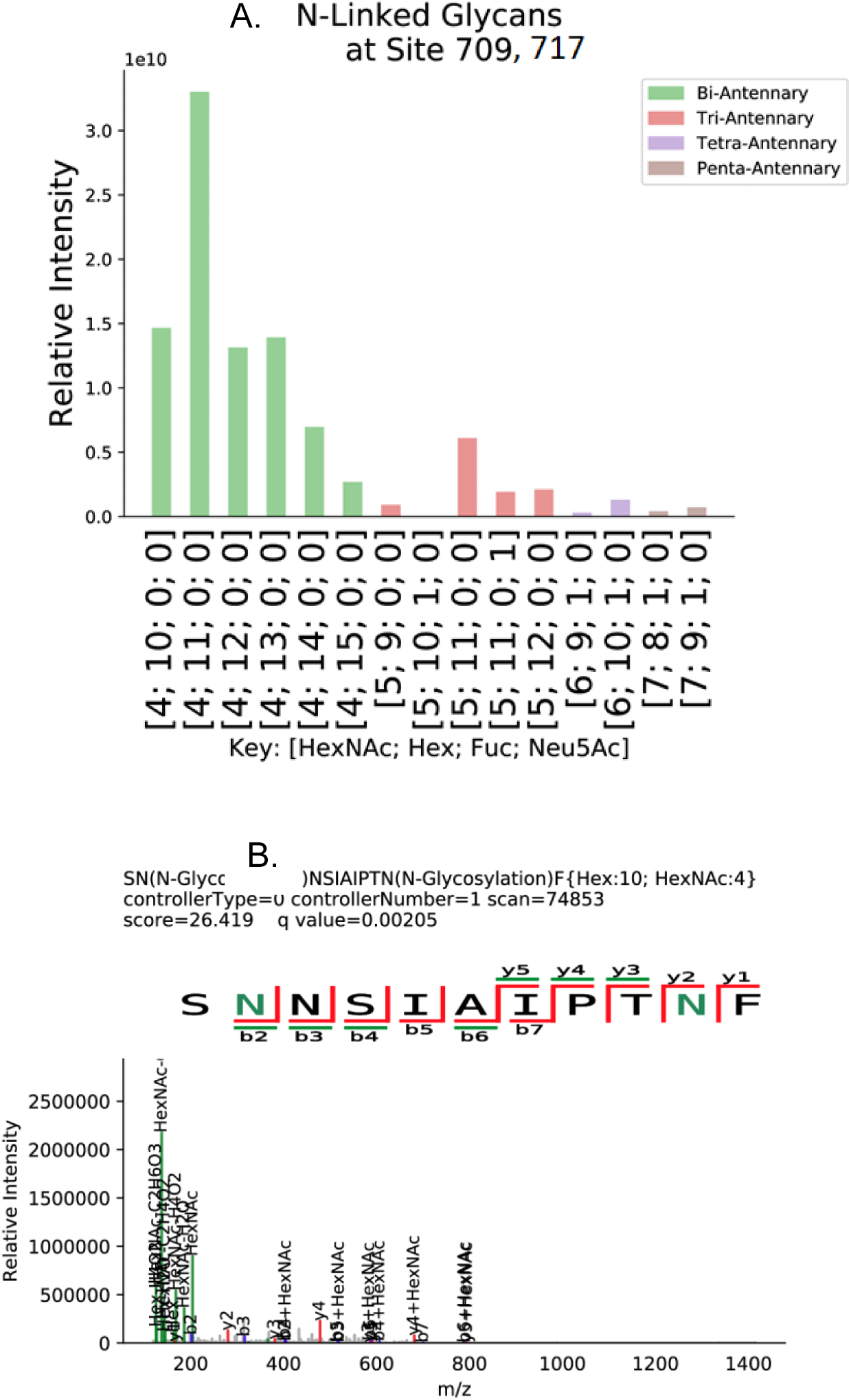
A. *N-*Glycan compositions at sites 709 and 717 from chymotryptic digest. B. Annotated tandem mass spectrum showing two *N*-glycosites.

**Figure 16.**
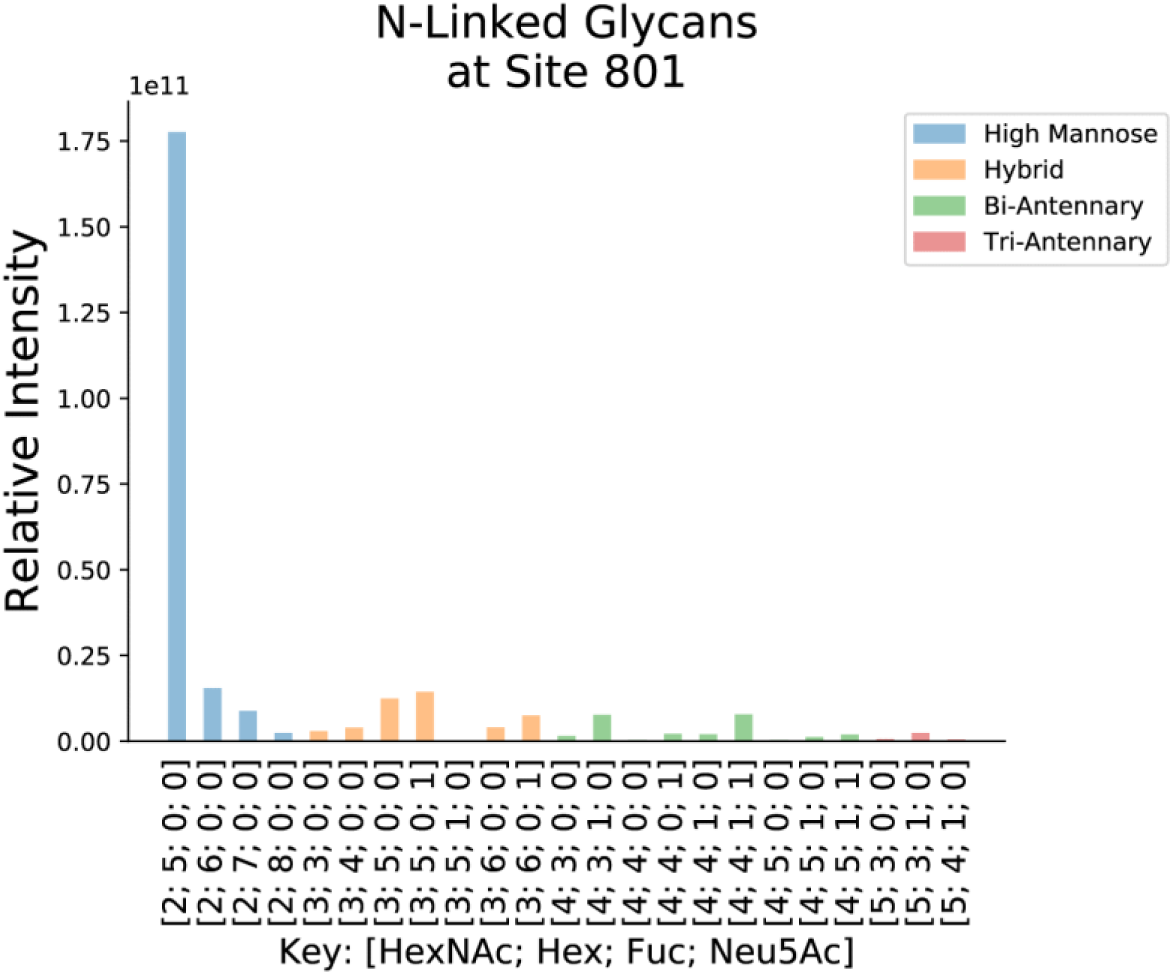
*N-*Glycans at site 801 from chymotryptic digest.

**Figure 17.**
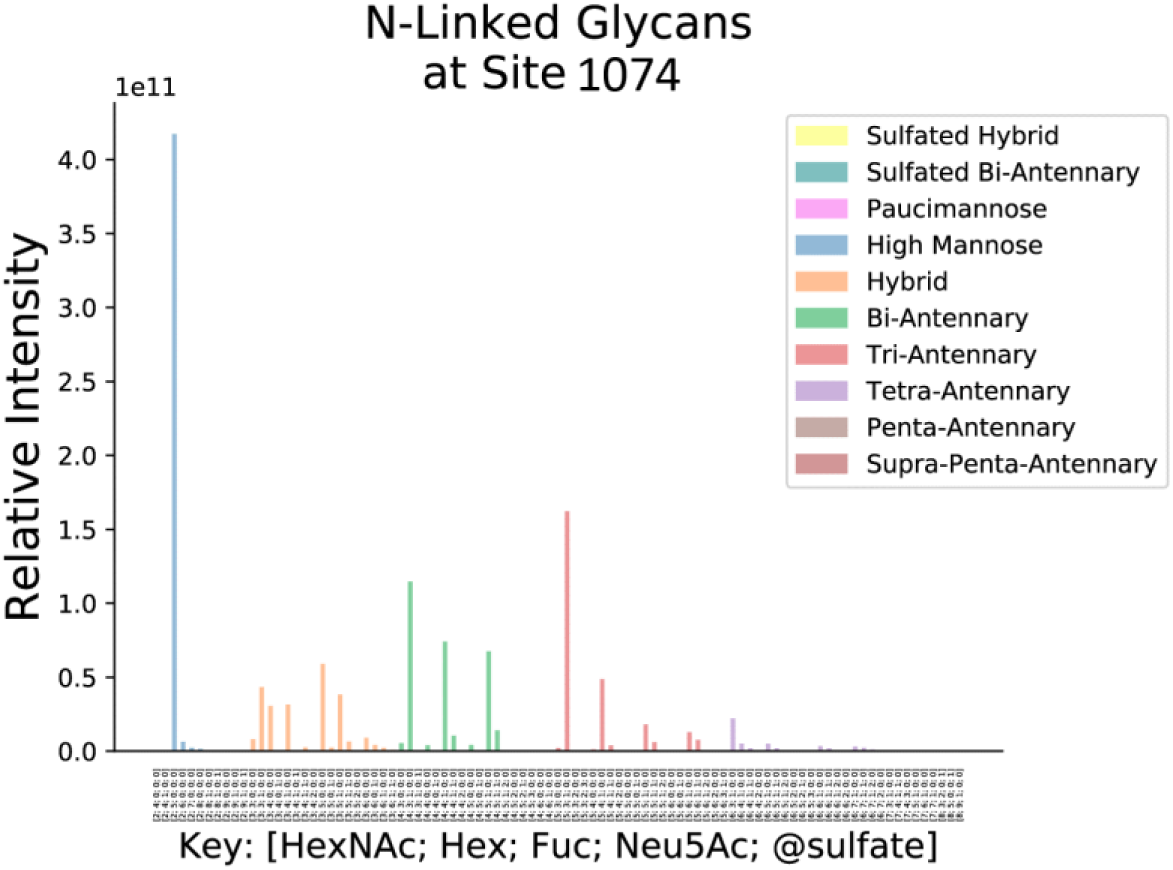
*N-*Glycans at site 1074 from tryptic digest.

**Figure 18.**
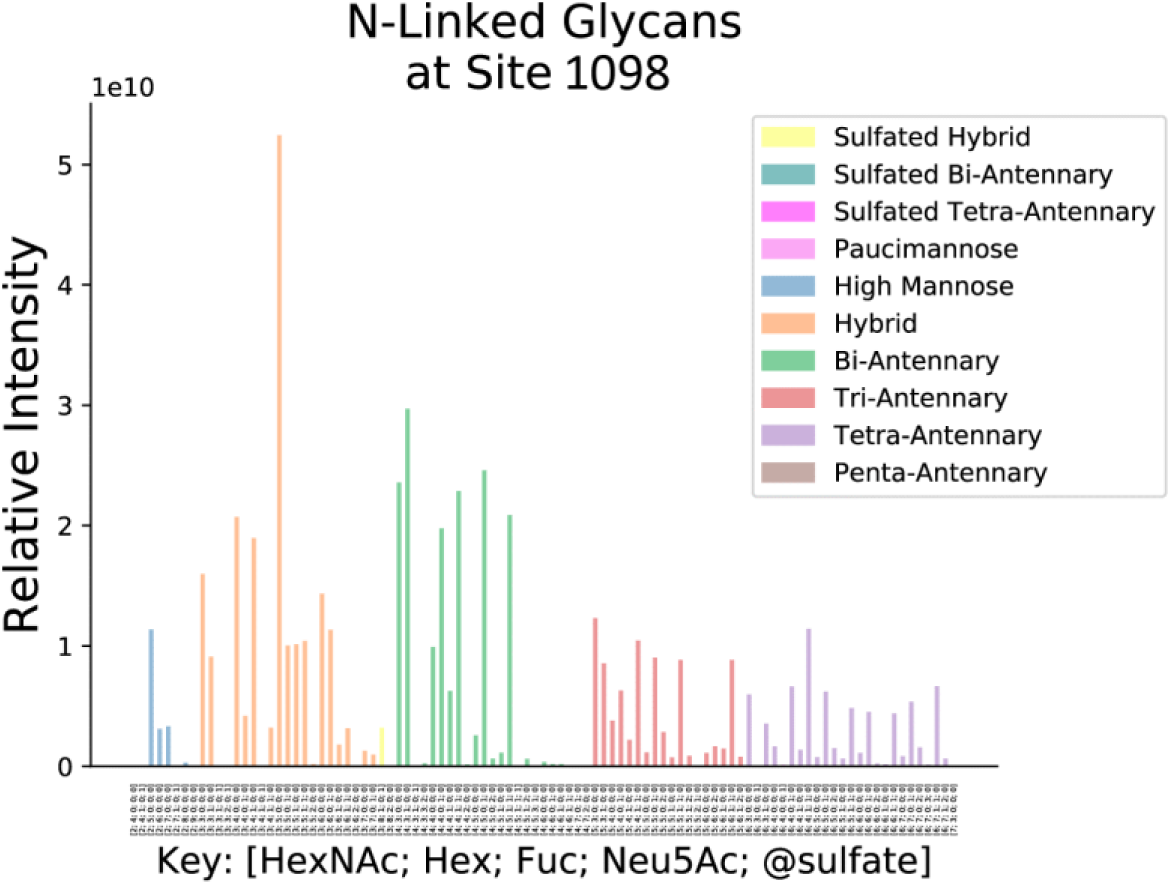
*N-*Glycans at site 1098 from chymotryptic digest.

**Figure 19.**
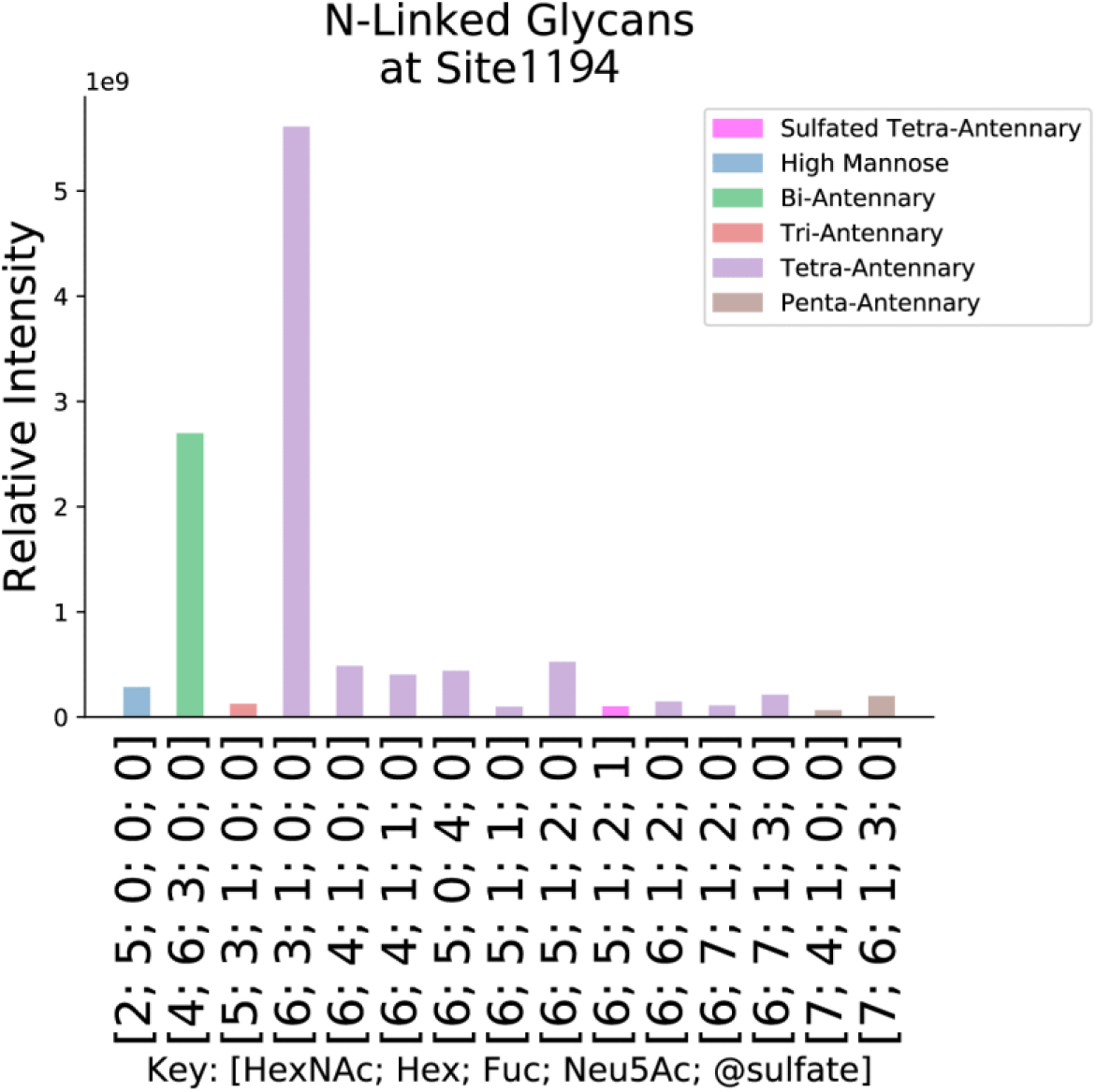
*N-*Glycans at site 1194 from tryptic digest.

*N-*Glycan sulfation is a topic of interest for influenza A virus because this modification influences viral replication, receptor binding, antigenicity and interactions with lectins of the innate immune system [86, 87]. In influenza A virus, the virus neuraminidase enzyme removes all or nearly all of the sialic acid residues from hemagglutinin *N-* glycans. Sulfation has been identified on C-3 of Gal and C-6 of GlcNAc residues of *N-*glycans as a biosynthetic event taking place in the trans-Golgi network [88]. Researchers investigated several influenza vaccine preparations and found sulfation at several *N-*glycan sites for H1N1, H3N2, H5N1, H7N9 and influenza B [89].

In contrast to influenza hemagglutinin, both sialylated and asialo *N-*glycans of S protein are abundant. We therefore included sulfation as a modification to the *N-* glycan search space we used for our analyses. We found position 74 to carry abundant sulfated tri-antennary and tetra-antennary *N-*glycans (Figure 6A). A total of 144 glycan compositions were identified at this position from the chymotryptic digest. An example annotated glycopeptide tandem mass spectrum is shown in Figure 6B. As shown in Figure 6C, 6 of the 30 most abundant glycan compositions at site 74 are sulfated. These abundant sulfated glycans range in composition Fuc_0-2_ Hex_3-4_ HexNAc_5-6_ NeuAc_0-2_, indicating that sulfation is likely placed on a non*-*reducing end HexNAc residue. Sulfation was also detected at trace levels for sites 1074 (Figure 17), 1098 (Figure 18) and 1194 (Figure 19).

As expected, each glycosite reflects a distribution of glycan compositions, consistent with the existence of populations of mature S glycoprotein molecules differing by glycosylation. As shown in Figure 6-Figure 19, glycans at sites 234, 709, 717, 801 are occupied primarily by high mannose *N-*glycans with minimal processing to complex type compositions. Note that sites 709 and 717 were identified in the same chymotryptic peptide (Figure 15) and we assumed one glycan per site. Glycans at sites 122, 165, 801, and 1074 display an abundant Hex_2_HexNAc_5_ composition, indicating processing by mannosidases, along with hybrid, complex biantennary and complex triantennary compositions, indicating that the S protein population undergoes a range from low to high degree of Golgi-mediated biosynthetic processing at these sites. Sites 74, 149, 282, 331, 343, 657, 1098, and 1194 contain extensively processed bi-, tri-, and tetra-antennary compositions, consistent with high degree of accessibility to biosynthetic enzymes at these sites.

## Conclusion

GlycReSoft is an open-source, publicly available software program that can used to analyze glycoproteomics LC-MS data. The program allows the user to specify glycan modifications including sulfation. We show an example of the use of GlycReSoft to assign SARS-CoV-2 S protein glycosylation from a published data set in which we identify sulfated *N*-glycans not identified in the original manuscript.

## Supporting Information

GlycReSoft HTML output summary tiles are provided for the SARS-CoV-2 S protein tryptic and chymotryptic digests, respectively.

