## Supplementary material for "Assignment of coronavirus spike protein site-specific glycosylation using GlycReSoft": 200519 SARS2_S_trypsin.html

### 200519 SARS2\_edited\_tryp\_sulfated\_1\_1\_0.mzid at 210320\_SARS\_CoV\_2\_T (8)

**MS^1^ Mass Accuracy:** 10.0 PPM  
**Grouping Tolerance:** 15.0 PPM  
**MS^n^ Mass Accuracy:** 20.0 PPM  
**Minimum `q`-Value For Clustering:** 0.05

**Glycopeptide Hypothesis:**
200519 SARS2\_edited\_tryp\_sulfated\_1\_1\_0.mzid  
**Sample Searched:**
210320\_SARS\_CoV\_2\_T

| Protein | Matched Glycopeptides |
| --- | --- |
| P0DTC2\|SPIKE\_SARS2 | 650 |

#### P0DTC2|SPIKE\_SARS2

**>P0DTC2|SPIKE\_SARS2**  
 `QCVNLTTRTQLPPAYTNSFTRGVYYPDKVFRSSVLHSTQDLFLPFFSNVTWFHAIHVSGTNGTKRFDNPVLPFNDGVYFA  
STEKSNIIRGWIFGTTLDSKTQSLLIVNNATNVVIKVCEFQFCNDPFLGVYYHKNNKSWMESEFRVYSSANNCTFEYVSQ  
PFLMDLEGKQGNFKNLREFVFKNIDGYFKIYSKHTPINLVRDLPQGFSALEPLVDLPIGINITRFQTLLALHRSYLTPGD  
SSSGWTAGAAAYYVGYLQPRTFLLKYNENGTITDAVDCALDPLSETKCTLKSFTVEKGIYQTSNFRVQPTESIVRFPNIT  
NLCPFGEVFNATRFASVYAWNRKRISNCVADYSVLYNSASFSTFKCYGVSPTKLNDLCFTNVYADSFVIRGDEVRQIAPG  
QTGKIADYNYKLPDDFTGCVIAWNSNNLDSKVGGNYNYLYRLFRKSNLKPFERDISTEIYQAGSTPCNGVEGFNCYFPLQ  
SYGFQPTNGVGYQPYRVVVLSFELLHAPATVCGPKKSTNLVKNKCVNFNFNGLTGTGVLTESNKKFLPFQQFGRDIADTT  
DAVRDPQTLEILDITPCSFGGVSVITPGTNTSNQVAVLYQDVNCTEVPVAIHADQLTPTWRVYSTGSNVFQTRAGCLIGA  
EHVNNSYECDIPIGAGICASYQTQTNSPGSASSVASQSIIAYTMSLGAENSVAYSNNSIAIPTNFTISVTTEILPVSMTK  
TSVDCTMYICGDSTECSNLLLQYGSFCTQLNRALTGIAVEQDKNTQEVFAQVKQIYKTPPIKDFGGFNFSQILPDPSKPS  
KRSFIEDLLFNKVTLADAGFIKQYGDCLGDIAARDLICAQKFNGLTVLPPLLTDEMIAQYTSALLAGTITSGWTFGAGAA  
LQIPFAMQMAYRFNGIGVTQNVLYENQKLIANQFNSAIGKIQDSLSSTASALGKLQDVVNQNAQALNTLVKQLSSNFGAI  
SSVLNDILSRLDPPEAEVQIDRLITGRLQSLQTYVTQQLIRAAEIRASANLAATKMSECVLGQSKRVDFCGKGYHLMSFP  
QSAPHGVVFLHVTYVPAQEKNFTTAPAICHDGKAHFPREGVFVSNGTHWFVTQRNFYEPQIITTDNTFVSGNCDVVIGIV  
NNTVYDPLQPELDSFKEELDKYFKNHTSPDVDLGDISGINASVVNIQKEIDRLNEVAKNLNESLIDLQELGKYEQYIKWP  
WYIWLGFIAGLIAIVMVTIMLCCMTSCCSCLKGCCSCGSCCKFDEDDSEPVLKGVKLHYT`

| Site | Type | Count |
| --- | --- | --- |
| 4 | N-linked | 51 |
| 48 | N-linked | 3 |
| 61 | N-linked | 32 |
| 109 | N-linked | 74 |
| 136 | N-linked | 2 |
| 152 | N-linked | 51 |
| 269 | N-linked | 46 |
| 330 | N-linked | 15 |
| 788 | N-linked | 118 |
| 1061 | N-linked | 120 |
| 1085 | N-linked | 123 |
| 1181 | N-linked | 15 |

| Mass | Sequence | MS_2_ Score | `q`-value | MS_1_ Score | Signal | Protein Coordinates | Tandem Scans | Start Time | Apex Time | End Time |
| --- | --- | --- | --- | --- | --- | --- | --- | --- | --- | --- |
| 3735.5573 | EGVFVSN(N)GTHWFVTQR Hex 6 HexNAc 3 Neu5Ac 1 | 32.896 | 0.001 | 17.440 | 1.135e+10 | 1078 - 1094 | 7 | 162.90 | 163.17 | 165.33 |
| 3817.6102 | EGVFVSN(N)GTHWFVTQR Hex 4 HexNAc 5 Neu5Ac 1 | 30.126 | 0.001 | 17.277 | 6.278e+09 | 1078 - 1094 | 12 | 163.21 | 164.24 | 165.04 |
| 4985.0822 | FPNITNLC(C)PFGEVFN(N)ATR Fuc 2 Hex 6 HexNAc 8 | 29.239 | 0.001 | 18.043 | 9.328e+09 | 315 - 333 | 5 | 207.16 | 207.71 | 208.91 |
| 3573.5051 | EGVFVSN(N)GTHWFVTQR Hex 5 HexNAc 3 Neu5Ac 1 | 28.522 | 0.001 | 17.562 | 1.003e+10 | 1078 - 1094 | 13 | 163.73 | 164.84 | 165.60 |
| 3444.4607 | EGVFVSN(N)GTHWFVTQR Hex 6 HexNAc 3 | 26.529 | 0.001 | 17.657 | 1.435e+10 | 1078 - 1094 | 8 | 148.08 | 148.42 | 149.81 |
| 3963.6672 | EGVFVSN(N)GTHWFVTQR Fuc 1 Hex 4 HexNAc 5 Neu5Ac 1 | 25.824 | 0.001 | 15.013 | 1.046e+10 | 1078 - 1094 | 9 | 162.13 | 163.33 | 163.98 |
| 3282.4079 | EGVFVSN(N)GTHWFVTQR Hex 5 HexNAc 3 | 25.716 | 0.001 | 16.853 | 5.244e+10 | 1078 - 1094 | 14 | 49.80 | 148.97 | 151.25 |
| 3719.5639 | EGVFVSN(N)GTHWFVTQR Fuc 1 Hex 5 HexNAc 3 Neu5Ac 1 | 25.551 | 0.001 | 16.710 | 1.042e+10 | 1078 - 1094 | 13 | 162.78 | 163.05 | 164.67 |
| 3402.3870 | N(N)FTTAPAIC(C)HDGK Fuc 1 Hex 5 HexNAc 5 | 25.129 | 0.001 | 12.409 | 1.797e+10 | 1060 - 1073 | 22 | 83.24 | 83.58 | 86.54 |
| 3133.2527 | N(N)FTTAPAIC(C)HDGK Hex 8 HexNAc 2 | 24.848 | 0.001 | 16.249 | 1.527e+09 | 1060 - 1073 | 7 | 82.74 | 83.16 | 84.13 |
| 4287.8090 | EGVFVSN(N)GTHWFVTQR Fuc 1 Hex 6 HexNAc 5 Neu5Ac 1 | 24.601 | 0.001 | 13.474 | 8.824e+09 | 1078 - 1094 | 13 | 159.74 | 161.35 | 161.98 |
| 3564.4360 | N(N)FTTAPAIC(C)HDGK Fuc 1 Hex 6 HexNAc 5 | 24.523 | 0.001 | 17.483 | 1.276e+10 | 1060 - 1073 | 18 | 82.45 | 83.30 | 84.62 |
| 3614.5316 | EGVFVSN(N)GTHWFVTQR Hex 4 HexNAc 4 Neu5Ac 1 | 24.405 | 0.001 | 17.762 | 1.977e+10 | 1078 - 1094 | 17 | 163.73 | 164.84 | 166.43 |
| 2875.1968 | N(N)FTTAPAIC(C)HDGK Fuc 1 Hex 3 HexNAc 4 | 24.266 | 0.001 | 18.165 | 1.147e+11 | 1060 - 1073 | 34 | 85.34 | 86.85 | 91.36 |
| 3428.4665 | EGVFVSN(N)GTHWFVTQR Fuc 1 Hex 5 HexNAc 3 | 24.177 | 0.001 | 16.715 | 1.013e+10 | 1078 - 1094 | 8 | 147.98 | 148.31 | 149.78 |
| 3328.3436 | N(N)FTTAPAIC(C)HDGK Fuc 1 Hex 4 HexNAc 4 Neu5Ac 1 | 23.763 | 0.001 | 18.056 | 1.040e+10 | 1060 - 1073 | 14 | 98.58 | 99.76 | 101.98 |
| 3776.5830 | EGVFVSN(N)GTHWFVTQR Hex 5 HexNAc 4 Neu5Ac 1 | 23.349 | 0.001 | 16.343 | 2.459e+10 | 1078 - 1094 | 24 | 147.20 | 162.78 | 165.08 |
| 3490.3964 | N(N)FTTAPAIC(C)HDGK Fuc 1 Hex 5 HexNAc 4 Neu5Ac 1 | 23.212 | 0.001 | 17.313 | 1.398e+10 | 1060 - 1073 | 18 | 97.23 | 98.45 | 100.54 |
| 4220.6642 | N(N)FTTAPAIC(C)HDGK Fuc 1 Hex 7 HexNAc 6 Neu5Ac 1 | 23.185 | 0.001 | 14.535 | 2.236e+09 | 1060 - 1073 | 7 | 94.62 | 95.34 | 95.64 |
| 3855.5320 | N(N)FTTAPAIC(C)HDGK Fuc 1 Hex 6 HexNAc 5 Neu5Ac 1 | 22.779 | 0.001 | 13.552 | 7.602e+09 | 1060 - 1073 | 15 | 96.15 | 96.45 | 183.69 |
| 3929.5661 | N(N)FTTAPAIC(C)HDGK Fuc 1 Hex 7 HexNAc 6 | 22.634 | 0.001 | 17.900 | 3.058e+09 | 1060 - 1073 | 7 | 82.36 | 82.57 | 83.17 |
| 3303.3118 | N(N)FTTAPAIC(C)HDGK Hex 6 HexNAc 3 Neu5Ac 1 | 22.528 | 0.001 | 18.348 | 4.092e+09 | 1060 - 1073 | 12 | 97.13 | 98.21 | 99.67 |
| 3287.3172 | N(N)FTTAPAIC(C)HDGK Fuc 1 Hex 5 HexNAc 3 Neu5Ac 1 | 21.982 | 0.001 | 18.841 | 6.443e+09 | 1060 - 1073 | 15 | 98.26 | 99.56 | 100.74 |
| 3161.3829 | EGVFVSN(N)GTHWFVTQR Hex 3 HexNAc 4 | 21.894 | 0.001 | 18.311 | 2.359e+10 | 1078 - 1094 | 9 | 149.54 | 149.87 | 151.31 |
| 3531.4229 | N(N)FTTAPAIC(C)HDGK Fuc 1 Hex 4 HexNAc 5 Neu5Ac 1 | 21.545 | 0.001 | 14.611 | 3.808e+09 | 1060 - 1073 | 9 | 98.39 | 98.94 | 100.16 |
| 2980.2647 | N(N)FTTAPAIC(C)HDGK Fuc 2 Hex 4 HexNAc 3 | 21.320 | 0.001 | -6.783 | 2.816e+08 | 1060 - 1073 | 15 | 57.71 | 85.96 | 87.27 |
| 3693.4759 | N(N)FTTAPAIC(C)HDGK Fuc 1 Hex 5 HexNAc 5 Neu5Ac 1 | 20.696 | 0.001 | 13.773 | 6.116e+09 | 1060 - 1073 | 14 | 96.99 | 97.31 | 99.24 |
| 3240.3566 | N(N)FTTAPAIC(C)HDGK Fuc 1 Hex 4 HexNAc 5 | 20.681 | 0.001 | 11.869 | 4.864e+10 | 1060 - 1073 | 29 | 67.90 | 86.85 | 90.46 |
| 4141.7148 | EGVFVSN(N)GTHWFVTQR Hex 6 HexNAc 5 Neu5Ac 1 | 20.666 | 0.001 | 14.095 | -5.940e+07 | 1078 - 1094 | 8 | 161.23 | 161.50 | 162.55 |
| 3037.2522 | N(N)FTTAPAIC(C)HDGK Fuc 1 Hex 4 HexNAc 4 | 20.347 | 0.001 | 15.735 | 7.425e+10 | 1060 - 1073 | 34 | 84.05 | 86.37 | 91.05 |
| 3760.5885 | EGVFVSN(N)GTHWFVTQR Fuc 1 Hex 4 HexNAc 4 Neu5Ac 1 | 20.315 | 0.001 | 16.007 | 2.289e+10 | 1078 - 1094 | 19 | 162.51 | 163.05 | 165.84 |
| 2834.1721 | N(N)FTTAPAIC(C)HDGK Fuc 1 Hex 4 HexNAc 3 | 19.773 | 0.001 | 13.487 | 3.157e+10 | 1060 - 1073 | 29 | 85.12 | 86.61 | 91.27 |
| 3078.2809 | N(N)FTTAPAIC(C)HDGK Fuc 1 Hex 3 HexNAc 5 | 19.718 | 0.001 | 17.216 | 1.622e+11 | 1060 - 1073 | 49 | 85.60 | 85.85 | 91.03 |
| 3567.5404 | EGVFVSN(N)GTHWFVTQR Hex 3 HexNAc 6 | 19.635 | 0.001 | 17.001 | 5.965e+09 | 1078 - 1094 | 8 | 148.17 | 148.47 | 149.54 |
| 3141.2594 | N(N)FTTAPAIC(C)HDGK Hex 5 HexNAc 3 Neu5Ac 1 | 19.612 | 0.001 | 17.950 | 2.499e+09 | 1060 - 1073 | 10 | 98.14 | 99.34 | 100.65 |
| 3411.4517 | EGVFVSN(N)GTHWFVTQR Hex 4 HexNAc 3 Neu5Ac 1 | 19.422 | 0.001 | 17.379 | 4.175e+09 | 1078 - 1094 | 7 | 164.48 | 165.57 | 166.03 |
| 4652.9035 | EGVFVSN(N)GTHWFVTQR Fuc 1 Hex 7 HexNAc 6 Neu5Ac 1 | 19.018 | 0.001 | 13.448 | 6.631e+09 | 1078 - 1094 | 1 | 159.21 | 159.33 | 160.35 |
| 2971.1901 | N(N)FTTAPAIC(C)HDGK Hex 7 HexNAc 2 | 18.925 | 0.001 | 15.327 | 2.311e+09 | 1060 - 1073 | 4 | 83.38 | 83.64 | 84.62 |
| 3979.6621 | EGVFVSN(N)GTHWFVTQR Hex 5 HexNAc 5 Neu5Ac 1 | 18.925 | 0.001 | 13.263 | 9.020e+09 | 1078 - 1094 | 10 | 161.98 | 162.66 | 164.07 |
| 3079.3324 | EGVFVSN(N)GTHWFVTQR Hex 5 HexNAc 2 | 18.770 | 0.001 | 17.291 | 1.136e+10 | 1078 - 1094 | 15 | 149.26 | 149.67 | 151.21 |
| 3767.5199 | N(N)FTTAPAIC(C)HDGK Fuc 1 Hex 6 HexNAc 6 | 18.727 | 0.001 | 12.146 | 3.293e+09 | 1060 - 1073 | 14 | 82.50 | 82.74 | 84.24 |
| 3605.4755 | N(N)FTTAPAIC(C)HDGK Fuc 1 Hex 5 HexNAc 6 | 18.634 | 0.001 | 9.023 | 4.957e+09 | 1060 - 1073 | 23 | 82.99 | 83.32 | 87.15 |
| 4756.9749 | FPNITNLC(C)PFGEVFN(N)ATR Fuc 1 Hex 8 HexNAc 6 | 18.450 | 0.001 | 14.327 | 3.206e+09 | 315 - 333 | 4 | 208.39 | 208.99 | 209.98 |
| 3012.2169 | N(N)FTTAPAIC(C)HDGK Hex 6 HexNAc 3 | 18.337 | 0.001 | 14.536 | 9.073e+09 | 1060 - 1073 | 10 | 83.94 | 84.55 | 89.39 |
| 3443.5142 | N(N)FTTAPAIC(C)HDGK Fuc 1 Hex 4 HexNAc 6 | 18.240 | 0.001 | 8.048 | 5.051e+09 | 1060 - 1073 | 20 | 65.76 | 84.66 | 87.56 |
| 3781.4914 | N(N)FTTAPAIC(C)HDGK Fuc 1 Hex 5 HexNAc 4 Neu5Ac 2 | 18.206 | 0.001 | 14.664 | 5.967e+08 | 1060 - 1073 | 4 | 113.77 | 113.94 | 114.18 |
| 4506.8477 | EGVFVSN(N)GTHWFVTQR Hex 7 HexNAc 6 Neu5Ac 1 | 18.202 | 0.001 | 16.326 | 5.343e+09 | 1078 - 1094 | 7 | 159.51 | 159.58 | 161.03 |
| 3241.3813 | EGVFVSN(N)GTHWFVTQR Hex 6 HexNAc 2 | 18.102 | 0.001 | 15.803 | 3.113e+09 | 1078 - 1094 | 4 | 148.57 | 148.84 | 149.52 |
| 3557.5096 | EGVFVSN(N)GTHWFVTQR Fuc 1 Hex 4 HexNAc 3 Neu5Ac 1 | 18.068 | 0.001 | 16.636 | 3.208e+09 | 1078 - 1094 | 9 | 163.44 | 163.71 | 165.24 |
| 3344.3387 | N(N)FTTAPAIC(C)HDGK Hex 5 HexNAc 4 Neu5Ac 1 | 17.970 | 0.001 | 16.069 | 7.669e+08 | 1060 - 1073 | 3 | 98.14 | 98.41 | 98.80 |
| 3266.4141 | EGVFVSN(N)GTHWFVTQR Fuc 1 Hex 4 HexNAc 3 | 17.863 | 0.001 | 17.861 | 1.896e+10 | 1078 - 1094 | 16 | 148.73 | 149.06 | 151.00 |
| 4020.7221 | EGVFVSN(N)GTHWFVTQR Hex 4 HexNAc 6 Neu5Ac 1 | 17.407 | 0.001 | 14.467 | 6.608e+09 | 1078 - 1094 | 10 | 162.66 | 163.64 | 164.40 |
| 4182.7419 | EGVFVSN(N)GTHWFVTQR Hex 5 HexNAc 6 Neu5Ac 1 | 17.397 | 0.001 | 13.587 | 6.202e+09 | 1078 - 1094 | 8 | 161.76 | 162.78 | 163.36 |
| 4511.8175 | N(N)FTTAPAIC(C)HDGK Fuc 1 Hex 7 HexNAc 6 Neu5Ac 2 | 17.305 | 0.001 | 12.557 | 1.053e+09 | 1060 - 1073 | 4 | 109.50 | 109.65 | 110.50 |
| 3469.5183 | EGVFVSN(N)GTHWFVTQR Fuc 1 Hex 4 HexNAc 4 | 17.017 | 0.001 | 11.415 | 6.259e+09 | 1078 - 1094 | 15 | 147.79 | 148.42 | 251.23 |
| 2958.3046 | EGVFVSN(N)GTHWFVTQR Hex 3 HexNAc 3 | 16.856 | 0.001 | 19.136 | 1.597e+10 | 1078 - 1094 | 12 | 149.94 | 150.70 | 151.38 |
| 3734.5026 | N(N)FTTAPAIC(C)HDGK Fuc 1 Hex 4 HexNAc 6 Neu5Ac 1 | 16.595 | 0.001 | 14.090 | 1.869e+09 | 1060 - 1073 | 10 | 97.95 | 98.24 | 100.41 |
| 3199.3297 | N(N)FTTAPAIC(C)HDGK Fuc 1 Hex 5 HexNAc 4 | 16.467 | 0.001 | 16.625 | 6.757e+10 | 1060 - 1073 | 33 | 52.04 | 85.14 | 90.13 |
| 3620.5589 | DFGGFN(N)FSQILPDPSKPSK Hex 7 HexNAc 2 | 16.428 | 0.001 | 15.111 | 7.217e+10 | 782 - 801 | 21 | 201.99 | 202.74 | 203.73 |
| 4125.7485 | EGVFVSN(N)GTHWFVTQR Fuc 1 Hex 5 HexNAc 5 Neu5Ac 1 | 16.408 | 0.001 | 10.946 | 8.832e+09 | 1078 - 1094 | 13 | 161.30 | 161.42 | 163.24 |
| 3782.6087 | DFGGFN(N)FSQILPDPSKPSK Hex 8 HexNAc 2 | 16.377 | 0.001 | 15.125 | 6.976e+09 | 782 - 801 | 9 | 201.67 | 202.27 | 202.77 |
| 3104.3756 | EGVFVSN(N)GTHWFVTQR Fuc 1 Hex 3 HexNAc 3 | 16.188 | 0.001 | 16.050 | 9.110e+09 | 1078 - 1094 | 13 | 149.52 | 149.87 | 154.17 |
| 4058.6469 | N(N)FTTAPAIC(C)HDGK Fuc 1 Hex 6 HexNAc 6 Neu5Ac 1 | 16.179 | 0.001 | 10.521 | 1.832e+09 | 1060 - 1073 | 9 | 95.75 | 96.97 | 97.36 |
| 3491.4000 | N(N)FTTAPAIC(C)HDGK Fuc 3 Hex 5 HexNAc 4 | 16.024 | 0.001 | -18.955 | 2.378e+07 | 1060 - 1073 | 2 | 98.23 | 98.23 | 98.23 |
| 3977.7116 | DFGGFN(N)FSQILPDPSKPSK Fuc 1 Hex 4 HexNAc 4 Neu5Ac 1 | 15.862 | 0.001 | 17.544 | 4.320e+10 | 782 - 801 | 17 | 218.16 | 219.87 | 220.65 |
| 3777.3056 | DFGGFN(N)FSQILPDPSKPSKR Hex 7 HexNAc 2 | 15.704 | 0.001 | 10.224 | 3.276e+09 | 782 - 802 | 10 | 112.50 | 185.42 | 185.54 |
| 4919.0260 | FPNITNLC(C)PFGEVFN(N)ATR Fuc 1 Hex 9 HexNAc 6 | 15.613 | 0.001 | 14.214 | 7.881e+08 | 315 - 333 | 2 | 207.38 | 208.29 | 208.73 |
| 3452.5507 | DFGGFN(N)FSQILPDPSKPSKR Hex 5 HexNAc 2 | 15.437 | 0.001 | 15.081 | 2.773e+10 | 782 - 802 | 8 | 186.19 | 186.56 | 187.78 |
| 3307.4384 | EGVFVSN(N)GTHWFVTQR Fuc 1 Hex 3 HexNAc 4 | 15.345 | 0.001 | 17.709 | 2.971e+10 | 1078 - 1094 | 14 | 148.67 | 148.98 | 151.27 |
| 2647.0854 | N(N)FTTAPAIC(C)HDGK Hex 5 HexNAc 2 | 15.187 | 0.001 | 19.402 | 4.173e+11 | 1060 - 1073 | 54 | 83.90 | 85.21 | 128.48 |
| 3922.6408 | EGVFVSN(N)GTHWFVTQR Fuc 1 Hex 5 HexNAc 4 Neu5Ac 1 | 15.151 | 0.001 | 15.385 | 2.087e+10 | 1078 - 1094 | 8 | 161.59 | 162.66 | 163.61 |
| 3526.5140 | EGVFVSN(N)GTHWFVTQR Hex 4 HexNAc 5 | 15.097 | 0.001 | 13.548 | 3.814e+09 | 1078 - 1094 | 10 | 147.92 | 148.60 | 149.60 |
| 3323.4338 | EGVFVSN(N)GTHWFVTQR Hex 4 HexNAc 4 | 14.988 | 0.001 | 14.840 | 9.913e+09 | 1078 - 1094 | 9 | 148.50 | 149.16 | 150.41 |
| 3296.4478 | DFGGFN(N)FSQILPDPSKPSK Hex 5 HexNAc 2 | 14.853 | 0.001 | 18.238 | 7.067e+11 | 782 - 801 | 70 | 187.07 | 204.30 | 215.13 |
| 3848.6730 | DFGGFN(N)FSQILPDPSKPSK Fuc 1 Hex 5 HexNAc 4 | 14.767 | 0.001 | 12.586 | 4.719e+09 | 782 - 801 | 20 | 198.55 | 199.20 | 202.58 |
| 3524.5614 | DFGGFN(N)FSQILPDPSKPSK Fuc 1 Hex 3 HexNAc 4 | 14.684 | 0.001 | 16.974 | 2.119e+10 | 782 - 801 | 11 | 200.81 | 201.46 | 202.58 |
| 3686.6260 | DFGGFN(N)FSQILPDPSKPSK Fuc 1 Hex 4 HexNAc 4 | 14.675 | 0.001 | -4.433 | 1.039e+10 | 782 - 801 | 21 | 134.61 | 200.59 | 202.05 |
| 3458.5125 | DFGGFN(N)FSQILPDPSKPSK Hex 6 HexNAc 2 | 14.646 | 0.001 | 15.430 | 4.311e+10 | 782 - 801 | 22 | 184.18 | 203.44 | 205.03 |
| 4166.7468 | EGVFVSN(N)GTHWFVTQR Fuc 1 Hex 4 HexNAc 6 Neu5Ac 1 | 14.637 | 0.001 | 15.498 | 1.142e+10 | 1078 - 1094 | 5 | 161.76 | 162.74 | 163.40 |
| 3364.4609 | EGVFVSN(N)GTHWFVTQR Hex 3 HexNAc 5 | 14.520 | 0.001 | 17.976 | 1.232e+10 | 1078 - 1094 | 14 | 148.75 | 149.06 | 150.41 |
| 3337.4857 | DFGGFN(N)FSQILPDPSKPSK Hex 4 HexNAc 3 | 14.487 | 0.001 | 15.723 | 2.946e+10 | 782 - 801 | 13 | 204.14 | 204.49 | 206.18 |
| 3485.4875 | EGVFVSN(N)GTHWFVTQR Hex 5 HexNAc 4 | 14.447 | 0.001 | 12.630 | 2.576e+09 | 1078 - 1094 | 11 | 147.90 | 148.17 | 149.60 |
| 4473.8380 | EGVFVSN(N)GTHWFVTQR Hex 5 HexNAc 6 Neu5Ac 2 | 14.299 | 0.001 | 14.718 | 1.504e+09 | 1078 - 1094 | 4 | 180.33 | 180.55 | 181.21 |
| 4328.8949 | EGVFVSN(N)GTHWFVTQR Fuc 1 Hex 5 HexNAc 6 Neu5Ac 1 | 14.255 | 0.001 | 11.445 | 4.827e+09 | 1078 - 1094 | 9 | 160.54 | 161.31 | 162.59 |
| 3145.5013 | TQSLLIVNN(N)ATNVVIK Hex 5 HexNAc 3 | 14.246 | 0.001 | 15.226 | 4.861e+10 | 100 - 116 | 24 | 160.29 | 161.01 | 166.22 |
| 3104.4752 | TQSLLIVNN(N)ATNVVIK Hex 6 HexNAc 2 | 14.170 | 0.001 | 15.680 | 4.374e+09 | 100 - 116 | 5 | 160.43 | 160.95 | 161.29 |
| 4344.8497 | EGVFVSN(N)GTHWFVTQR Hex 6 HexNAc 6 Neu5Ac 1 | 14.144 | 0.001 | 9.210 | 4.501e+09 | 1078 - 1094 | 15 | 160.29 | 161.34 | 233.23 |
| 3785.7363 | TQSLLIVNN(N)ATNVVIK Fuc 1 Hex 5 HexNAc 4 Neu5Ac 1 | 14.030 | 0.001 | 13.116 | 1.424e+10 | 100 - 116 | 12 | 176.75 | 177.22 | 183.92 |
| 2967.4534 | TQSLLIVNN(N)ATNVVIK Fuc 1 Hex 3 HexNAc 3 | 13.919 | 0.001 | 14.232 | 1.165e+10 | 100 - 116 | 7 | 162.24 | 162.64 | 164.32 |
| 6334.7918 | SSVLHSTQDLFLPFFSNVTWFHAIHVSGTN(N)GTK Fuc 1 Hex 8 HexNAc 6 | 13.872 | 0.001 | 14.845 | 7.086e+09 | 31 - 64 | 2 | 208.97 | 209.28 | 210.56 |
| 2809.1446 | N(N)FTTAPAIC(C)HDGK Hex 6 HexNAc 2 | 13.763 | 0.001 | 15.543 | 6.379e+09 | 1060 - 1073 | 10 | 58.13 | 84.47 | 89.35 |
| 2996.2219 | N(N)FTTAPAIC(C)HDGK Fuc 1 Hex 5 HexNAc 3 | 13.757 | 0.001 | 16.374 | 3.829e+10 | 1060 - 1073 | 12 | 84.32 | 86.20 | 90.67 |
| 4139.7644 | DFGGFN(N)FSQILPDPSKPSK Fuc 1 Hex 5 HexNAc 4 Neu5Ac 1 | 13.614 | 0.001 | 17.859 | 9.284e+09 | 782 - 801 | 12 | 216.23 | 218.16 | 219.19 |
| 3499.5441 | DFGGFN(N)FSQILPDPSKPSK Hex 5 HexNAc 3 | 13.600 | 0.001 | 15.502 | 4.679e+10 | 782 - 801 | 19 | 203.04 | 203.51 | 206.98 |
| 3125.2644 | N(N)FTTAPAIC(C)HDGK Fuc 1 Hex 4 HexNAc 3 Neu5Ac 1 | 13.584 | 0.001 | 18.221 | 2.661e+09 | 1060 - 1073 | 6 | 99.45 | 100.52 | 101.26 |
| 3727.6416 | DFGGFN(N)FSQILPDPSKPSK Fuc 1 Hex 3 HexNAc 5 | 13.577 | 0.001 | 16.876 | 1.213e+10 | 782 - 801 | 14 | 200.38 | 201.22 | 202.35 |
| 3053.2442 | N(N)FTTAPAIC(C)HDGK Hex 5 HexNAc 4 | 13.499 | 0.001 | 10.684 | 4.122e+09 | 1060 - 1073 | 6 | 84.05 | 84.84 | 86.41 |
| 3661.6116 | DFGGFN(N)FSQILPDPSKPSK Hex 6 HexNAc 3 | 13.479 | 0.001 | 13.749 | 4.182e+10 | 782 - 801 | 13 | 202.27 | 202.84 | 203.64 |
| 3291.5599 | TQSLLIVNN(N)ATNVVIK Fuc 1 Hex 5 HexNAc 3 | 13.469 | 0.001 | 15.434 | 1.121e+10 | 100 - 116 | 17 | 159.89 | 161.01 | 163.05 |
| 2526.0584 | N(N)FTTAPAIC(C)HDGK Hex 3 HexNAc 3 | 13.462 | 0.001 | 18.004 | 8.044e+09 | 1060 - 1073 | 5 | 86.81 | 87.04 | 87.52 |
| 2688.1118 | N(N)FTTAPAIC(C)HDGK Hex 4 HexNAc 3 | 13.457 | 0.001 | 15.007 | 3.055e+10 | 1060 - 1073 | 13 | 85.85 | 86.08 | 90.80 |
| 3403.4350 | EGVFVSN(N)GTHWFVTQR Hex 7 HexNAc 2 | 13.446 | 0.001 | 17.013 | 3.338e+09 | 1078 - 1094 | 6 | 147.92 | 148.17 | 148.79 |
| 2729.1408 | N(N)FTTAPAIC(C)HDGK Hex 3 HexNAc 4 | 13.437 | 0.001 | 14.133 | 5.478e+09 | 1060 - 1073 | 9 | 86.24 | 86.43 | 125.97 |
| 3952.6833 | DFGGFN(N)FSQILPDPSKPSK Hex 6 HexNAc 3 Neu5Ac 1 | 13.410 | 0.001 | 16.552 | 2.298e+10 | 782 - 801 | 12 | 220.92 | 222.54 | 223.45 |
| 4490.8529 | EGVFVSN(N)GTHWFVTQR Fuc 1 Hex 6 HexNAc 6 Neu5Ac 1 | 13.384 | 0.001 | 8.074 | 4.378e+09 | 1078 - 1094 | 6 | 159.51 | 159.67 | 161.39 |
| 3175.4234 | DFGGFN(N)FSQILPDPSKPSK Hex 3 HexNAc 3 | 13.352 | 0.001 | 17.542 | 5.796e+09 | 782 - 801 | 6 | 205.12 | 205.79 | 206.74 |
| 3623.7163 | TQSLLIVNN(N)ATNVVIK Fuc 1 Hex 4 HexNAc 4 Neu5Ac 1 | 13.106 | 0.001 | 13.320 | 3.015e+10 | 100 - 116 | 24 | 178.19 | 179.51 | 228.63 |
| 3510.5187 | EGVFVSN(N)GTHWFVTQR Fuc 1 Hex 3 HexNAc 5 | 13.064 | 0.001 | 15.098 | 8.552e+09 | 1078 - 1094 | 8 | 147.92 | 148.42 | 149.54 |
| 3307.5547 | TQSLLIVNN(N)ATNVVIK Hex 6 HexNAc 3 | 13.011 | 0.001 | 17.550 | 1.335e+10 | 100 - 116 | 8 | 159.64 | 160.18 | 161.08 |
| 3832.7903 | TPPIKDFGGFN(N)FSQILPDPSKPSK Hex 5 HexNAc 2 | 13.009 | 0.001 | 13.808 | 3.430e+10 | 777 - 801 | 21 | 187.39 | 188.21 | 214.06 |
| 3348.5838 | TQSLLIVNN(N)ATNVVIK Hex 5 HexNAc 4 | 12.846 | 0.001 | 11.089 | 7.817e+09 | 100 - 116 | 12 | 159.58 | 159.77 | 165.53 |
| 2942.4213 | TQSLLIVNN(N)ATNVVIK Hex 5 HexNAc 2 | 12.728 | 0.001 | 18.295 | 1.045e+11 | 100 - 116 | 26 | 159.80 | 161.49 | 166.85 |
| 3420.6043 | TQSLLIVNN(N)ATNVVIK Fuc 1 Hex 4 HexNAc 3 Neu5Ac 1 | 12.693 | 0.001 | -2.514 | 6.072e+09 | 100 - 116 | 13 | 160.62 | 180.24 | 181.37 |
| 3332.5853 | TQSLLIVNN(N)ATNVVIK Fuc 1 Hex 4 HexNAc 4 | 12.653 | 0.001 | 13.013 | 1.643e+10 | 100 - 116 | 13 | 159.61 | 161.01 | 162.51 |
| 2485.0331 | N(N)FTTAPAIC(C)HDGK Hex 4 HexNAc 2 | 12.632 | 0.001 | 11.953 | 3.281e+08 | 1060 - 1073 | 1 | 86.58 | 86.78 | 86.91 |
| 3080.2813 | N(N)FTTAPAIC(C)HDGK Fuc 1 Hex 3 HexNAc 5 | 12.630 | 0.001 | 0.000 | 0.000e+00 | 1060 - 1073 | 5 | - | - | - |
| 3807.9654 | DFGGFN(N)FSQILPDPSKPSK Fuc 1 Hex 6 HexNAc 3 | 12.448 | 0.001 | -5.449 | 1.632e+09 | 782 - 801 | 7 | 201.99 | 202.61 | 214.03 |
| 3993.7083 | DFGGFN(N)FSQILPDPSKPSK Hex 5 HexNAc 4 Neu5Ac 1 | 12.238 | 0.001 | 10.291 | -1.878e+09 | 782 - 801 | 10 | 198.28 | 221.81 | 222.19 |
| 4782.0060 | FPNITNLC(C)PFGEVFN(N)ATR Fuc 2 Hex 6 HexNAc 7 | 12.213 | 0.001 | 14.141 | 1.990e+09 | 315 - 333 | 3 | 208.29 | 208.70 | 209.35 |
| 3224.3035 | N(N)FTTAPAIC(C)HDGK Fuc 2 Hex 3 HexNAc 5 | 12.145 | 0.001 | -9.905 | 3.079e+07 | 1060 - 1073 | 4 | 86.53 | 86.53 | 91.05 |
| 3817.6839 | DFGGFN(N)FSQILPDPSKPSKR Hex 6 HexNAc 3 | 12.088 | 0.001 | -2.978 | 1.063e+08 | 782 - 802 | 1 | 185.32 | 185.42 | 185.42 |
| 2932.2158 | N(N)FTTAPAIC(C)HDGK Hex 3 HexNAc 5 | 12.071 | 0.001 | 10.569 | 2.102e+09 | 1060 - 1073 | 5 | 85.23 | 87.25 | 87.48 |
| 3120.3574 | EGVFVSN(N)GTHWFVTQR Hex 4 HexNAc 3 | 12.040 | 0.001 | 18.617 | 2.073e+10 | 1078 - 1094 | 12 | 149.64 | 149.94 | 151.27 |
| 4156.8882 | TPPIKDFGGFN(N)FSQILPDPSKPSK Hex 7 HexNAc 2 | 11.936 | 0.001 | 14.413 | 2.482e+09 | 777 - 801 | 6 | 186.78 | 187.22 | 187.92 |
| 4270.7659 | EGVFVSN(N)GTHWFVTQR Hex 5 HexNAc 5 Neu5Ac 2 | 11.922 | 0.001 | 12.691 | 2.861e+09 | 1078 - 1094 | 10 | 180.04 | 181.21 | 182.27 |
| 3436.5984 | TQSLLIVNN(N)ATNVVIK Hex 5 HexNAc 3 Neu5Ac 1 | 11.918 | 0.001 | 14.838 | 2.418e+10 | 100 - 116 | 9 | 178.64 | 179.82 | 181.91 |
| 3281.3822 | N(N)FTTAPAIC(C)HDGK Fuc 1 Hex 3 HexNAc 6 | 11.758 | 0.001 | 13.809 | 2.204e+10 | 1060 - 1073 | 13 | 85.25 | 85.50 | 88.21 |
| 2672.1186 | N(N)FTTAPAIC(C)HDGK Fuc 1 Hex 3 HexNAc 3 | 11.682 | 0.001 | 18.236 | 4.331e+10 | 1060 - 1073 | 17 | 86.37 | 87.64 | 90.76 |
| 3345.3591 | N(N)FTTAPAIC(C)HDGK Fuc 2 Hex 5 HexNAc 4 | 11.679 | 0.001 | 16.641 | 6.292e+08 | 1060 - 1073 | 6 | 83.94 | 84.22 | 84.52 |
| 2979.2065 | N(N)FTTAPAIC(C)HDGK Hex 4 HexNAc 3 Neu5Ac 1 | 11.616 | 0.001 | 17.218 | 5.711e+08 | 1060 - 1073 | 3 | 100.16 | 100.39 | 100.88 |
| 3453.6121 | TQSLLIVNN(N)ATNVVIK Fuc 1 Hex 6 HexNAc 3 | 11.528 | 0.001 | 12.358 | 3.695e+09 | 100 - 116 | 8 | 159.44 | 159.55 | 160.78 |
| 3129.5137 | TQSLLIVNN(N)ATNVVIK Fuc 1 Hex 4 HexNAc 3 | 11.491 | 0.001 | 14.488 | 1.638e+10 | 100 - 116 | 13 | 161.03 | 161.67 | 167.83 |
| 3227.5538 | TQSLLIVNN(N)ATNVVIK Hex 3 HexNAc 5 | 11.482 | 0.001 | 14.428 | 5.787e+09 | 100 - 116 | 8 | 160.98 | 161.67 | 162.55 |
| 3672.5721 | EGVFVSN(N)GTHWFVTQR Fuc 1 Hex 4 HexNAc 5 | 11.445 | 0.001 | 13.307 | 2.195e+09 | 1078 - 1094 | 8 | 147.45 | 147.90 | 148.79 |
| 3791.6206 | DFGGFN(N)FSQILPDPSKPSK Fuc 2 Hex 5 HexNAc 3 | 11.419 | 0.001 | 14.075 | -5.759e+08 | 782 - 801 | 4 | 223.21 | 223.57 | 224.32 |
| 3494.6379 | TQSLLIVNN(N)ATNVVIK Fuc 1 Hex 5 HexNAc 4 | 11.406 | 0.001 | 13.622 | 5.268e+09 | 100 - 116 | 11 | 159.38 | 159.67 | 161.18 |
| 3535.6922 | TQSLLIVNN(N)ATNVVIK Fuc 1 Hex 4 HexNAc 5 | 11.340 | 0.001 | -5.732 | 4.238e+09 | 100 - 116 | 10 | 159.42 | 160.93 | 183.66 |
| 3697.7175 | TQSLLIVNN(N)ATNVVIK Fuc 1 Hex 5 HexNAc 5 | 11.055 | 0.001 | 13.470 | 8.398e+08 | 100 - 116 | 2 | 159.47 | 159.61 | 160.41 |
| 3702.6330 | DFGGFN(N)FSQILPDPSKPSK Hex 5 HexNAc 4 | 11.049 | 0.001 | 9.357 | 3.607e+09 | 782 - 801 | 17 | 201.80 | 202.27 | 203.98 |
| 3378.5015 | DFGGFN(N)FSQILPDPSKPSK Hex 3 HexNAc 4 | 10.681 | 0.001 | -3.076 | 1.432e+10 | 782 - 801 | 8 | 142.89 | 204.44 | 205.28 |
| 3994.8559 | TPPIKDFGGFN(N)FSQILPDPSKPSK Hex 6 HexNAc 2 | 10.540 | 0.001 | -2.837 | 1.539e+09 | 777 - 801 | 13 | 102.26 | 187.82 | 193.13 |
| 2983.4482 | TQSLLIVNN(N)ATNVVIK Hex 4 HexNAc 3 | 10.423 | 0.001 | 16.696 | 1.957e+10 | 100 - 116 | 19 | 161.52 | 161.88 | 165.39 |
| 3437.6166 | TQSLLIVNN(N)ATNVVIK Fuc 2 Hex 5 HexNAc 3 | 10.418 | 0.001 | 13.897 | 2.220e+09 | 100 - 116 | 9 | 159.46 | 160.05 | 161.01 |
| 4797.9457 | EGVFVSN(N)GTHWFVTQR Hex 7 HexNAc 6 Neu5Ac 2 | 10.405 | 0.001 | 9.160 | 1.565e+09 | 1078 - 1094 | 3 | 176.84 | 178.23 | 178.38 |
| 3158.2761 | N(N)FTTAPAIC(C)HDGK Fuc 1 Hex 6 HexNAc 3 | 10.388 | 0.001 | 9.878 | 2.318e+09 | 1060 - 1073 | 7 | 41.26 | 84.91 | 85.06 |
| 3519.6708 | TQSLLIVNN(N)ATNVVIK Fuc 2 Hex 3 HexNAc 5 | 10.378 | 0.001 | 15.932 | 1.795e+09 | 100 - 116 | 4 | 159.92 | 160.43 | 160.93 |
| 3628.5736 | DFGGFN(N)FSQILPDPSKPSK Hex 4 HexNAc 3 Neu5Ac 1 | 10.329 | 0.001 | 14.960 | 1.534e+09 | 782 - 801 | 5 | 224.59 | 224.97 | 225.61 |
| 3478.6442 | TQSLLIVNN(N)ATNVVIK Fuc 2 Hex 4 HexNAc 4 | 10.296 | 0.001 | 15.259 | 3.968e+09 | 100 - 116 | 5 | 159.64 | 160.05 | 161.08 |
| 3182.2863 | N(N)FTTAPAIC(C)HDGK Hex 4 HexNAc 4 Neu5Ac 1 | 10.279 | 0.001 | 16.294 | 4.864e+08 | 1060 - 1073 | 4 | 99.41 | 99.66 | 100.43 |
| 3186.5276 | TQSLLIVNN(N)ATNVVIK Hex 4 HexNAc 4 | 10.268 | 0.001 | 14.449 | 1.730e+10 | 100 - 116 | 11 | 160.32 | 161.24 | 162.51 |
| 3170.5342 | TQSLLIVNN(N)ATNVVIK Fuc 1 Hex 3 HexNAc 4 | 10.234 | 0.001 | 15.986 | 2.083e+10 | 100 - 116 | 15 | 161.35 | 161.67 | 167.20 |
| 3445.4148 | N(N)FTTAPAIC(C)HDGK Fuc 1 Hex 4 HexNAc 6 | 10.232 | 0.001 | 0.000 | 0.000e+00 | 1060 - 1073 | 1 | - | - | - |
| 3435.5251 | NLN(N)ESLIDLQELGK Fuc 1 Hex 3 HexNAc 6 | 10.228 | 0.001 | 16.382 | 5.611e+09 | 1178 - 1192 | 9 | 173.32 | 174.01 | 176.00 |
| 2898.1855 | N(N)FTTAPAIC(C)HDGK Fuc 1 Hex 3 HexNAc 4 | 10.140 | 0.001 | 0.000 | 0.000e+00 | 1060 - 1073 | 2 | - | - | - |
| 3681.7262 | TQSLLIVNN(N)ATNVVIK Fuc 2 Hex 4 HexNAc 5 | 10.093 | 0.001 | 14.642 | 2.551e+09 | 100 - 116 | 11 | 159.37 | 159.44 | 160.72 |
| 4396.8161 | YNEN(N)GTITDAVDC(C)ALDPLSETK Fuc 1 Hex 5 HexNAc 5 | 10.068 | 0.001 | 14.003 | -1.309e+09 | 265 - 287 | 10 | 187.17 | 187.58 | 188.30 |
| 3831.6487 | DFGGFN(N)FSQILPDPSKPSK Hex 4 HexNAc 4 Neu5Ac 1 | 9.872 | 0.001 | 14.797 | 7.821e+09 | 782 - 801 | 8 | 223.03 | 223.63 | 224.18 |
| 3790.6216 | DFGGFN(N)FSQILPDPSKPSK Hex 5 HexNAc 3 Neu5Ac 1 | 9.832 | 0.001 | 14.171 | 1.394e+10 | 782 - 801 | 2 | 223.40 | 223.70 | 224.00 |
| 3783.6120 | DFGGFN(N)FSQILPDPSKPSK Hex 8 HexNAc 2 | 9.763 | 0.001 | 0.000 | 0.000e+00 | 782 - 801 | 1 | - | - | - |
| 3631.5463 | EGVFVSN(N)GTHWFVTQR Fuc 1 Hex 5 HexNAc 4 | 9.700 | 0.001 | 12.889 | 1.127e+09 | 1078 - 1094 | 8 | 147.36 | 147.48 | 148.69 |
| 3483.5362 | DFGGFN(N)FSQILPDPSKPSK Fuc 1 Hex 4 HexNAc 3 | 9.694 | 0.001 | 14.465 | 1.314e+09 | 782 - 801 | 4 | 201.02 | 201.46 | 202.05 |
| 3896.5556 | N(N)FTTAPAIC(C)HDGK Fuc 1 Hex 5 HexNAc 6 Neu5Ac 1 | 9.644 | 0.001 | 9.614 | 2.023e+09 | 1060 - 1073 | 17 | 96.49 | 97.80 | 99.72 |
| 3321.4824 | DFGGFN(N)FSQILPDPSKPSK Fuc 1 Hex 3 HexNAc 3 | 9.572 | 0.001 | 13.585 | 8.950e+08 | 782 - 801 | 5 | 201.15 | 202.27 | 202.66 |
| 3484.4343 | N(N)FTTAPAIC(C)HDGK Fuc 1 Hex 3 HexNAc 7 | 9.316 | 0.001 | -5.158 | 1.494e+08 | 1060 - 1073 | 2 | 85.74 | 85.89 | 85.92 |
| 3274.5448 | TQSLLIVNN(N)ATNVVIK Hex 4 HexNAc 3 Neu5Ac 1 | 9.268 | 0.001 | 13.397 | 4.914e+09 | 100 - 116 | 10 | 179.64 | 180.70 | 181.91 |
| 3897.6101 | EGVFVSN(N)GTHWFVTQR Hex 7 HexNAc 3 Neu5Ac 1 | 9.225 | 0.001 | 12.499 | 9.838e+08 | 1078 - 1094 | 7 | 162.21 | 163.33 | 163.64 |
| 3606.5229 | EGVFVSN(N)GTHWFVTQR Hex 7 HexNAc 3 | 9.168 | 0.001 | 14.174 | 1.288e+09 | 1078 - 1094 | 8 | 147.67 | 147.90 | 165.16 |
| 3373.6122 | TQSLLIVNN(N)ATNVVIK Fuc 1 Hex 3 HexNAc 5 | 9.120 | 0.001 | 13.474 | 9.427e+09 | 100 - 116 | 9 | 160.45 | 161.37 | 166.29 |
| 3875.6513 | EGVFVSN(N)GTHWFVTQR Fuc 1 Hex 4 HexNAc 6 | 9.103 | 0.001 | 11.868 | 1.375e+09 | 1078 - 1094 | 5 | 146.88 | 147.50 | 148.17 |
| 3266.5285 | TQSLLIVNN(N)ATNVVIK Hex 7 HexNAc 2 | 9.103 | 0.001 | 16.461 | 6.074e+09 | 100 - 116 | 6 | 159.58 | 159.85 | 160.98 |
| 2850.1914 | N(N)FTTAPAIC(C)HDGK Hex 5 HexNAc 3 | 9.078 | 0.001 | 16.584 | 5.918e+10 | 1060 - 1073 | 19 | 84.55 | 84.86 | 90.57 |
| 4341.8266 | NLN(N)ESLIDLQELGK Fuc 1 Hex 5 HexNAc 6 Neu5Ac 2 | 9.075 | 0.001 | 14.369 | 5.234e+08 | 1178 - 1192 | 2 | 214.67 | 214.97 | 215.28 |
| 3598.6531 | TQSLLIVNN(N)ATNVVIK Hex 6 HexNAc 3 Neu5Ac 1 | 9.074 | 0.001 | 11.173 | 9.020e+09 | 100 - 116 | 14 | 177.67 | 177.91 | 184.87 |
| 4581.8850 | EGVFVSN(N)GTHWFVTQR Fuc 1 Hex 6 HexNAc 5 Neu5Ac 2 | 9.069 | 0.001 | 0.000 | 0.000e+00 | 1078 - 1094 | 2 | - | - | - |
| 3729.5951 | EGVFVSN(N)GTHWFVTQR Hex 4 HexNAc 6 | 9.038 | 0.001 | 13.115 | 1.643e+09 | 1078 - 1094 | 7 | 147.58 | 148.04 | 148.81 |
| 4857.9921 | VYSSANN(N)C(C)TFEYVSQPFLMDLEGK Fuc 1 Hex 5 HexNAc 4 Neu5Ac 1 | 9.030 | 0.001 | 14.936 | 6.745e+09 | 145 - 169 | 9 | 230.02 | 231.36 | 232.92 |
| 3094.2700 | N(N)FTTAPAIC(C)HDGK Hex 4 HexNAc 5 | 9.005 | 0.001 | 9.876 | 1.241e+09 | 1060 - 1073 | 7 | 83.94 | 86.54 | 86.72 |
| 3713.5986 | EGVFVSN(N)GTHWFVTQR Fuc 1 Hex 3 HexNAc 6 | 9.000 | 0.001 | 17.209 | 3.551e+09 | 1078 - 1094 | 7 | 147.67 | 147.87 | 148.69 |
| 3582.7003 | TQSLLIVNN(N)ATNVVIK Fuc 1 Hex 5 HexNAc 3 Neu5Ac 1 | 8.893 | 0.001 | 11.883 | 1.211e+10 | 100 - 116 | 9 | 178.16 | 179.45 | 179.91 |
| 3024.4743 | TQSLLIVNN(N)ATNVVIK Hex 3 HexNAc 4 | 8.873 | 0.001 | 14.990 | 9.150e+09 | 100 - 116 | 12 | 161.60 | 161.95 | 163.81 |
| 6992.0394 | SSVLHSTQDLFLPFFSNVTWFHAIHVSGTN(N)GTK Fuc 3 Hex 9 HexNAc 7 | 8.859 | 0.001 | 12.079 | 1.619e+09 | 31 - 64 | 4 | 207.00 | 207.21 | 207.55 |
| 3449.3705 | N(N)FTTAPAIC(C)HDGK Fuc 1 Hex 6 HexNAc 3 Neu5Ac 1 | 8.858 | 0.001 | 16.597 | 7.835e+08 | 1060 - 1073 | 5 | 98.07 | 98.37 | 98.75 |
| 3850.6208 | EGVFVSN(N)GTHWFVTQR Hex 6 HexNAc 5 | 8.836 | 0.001 | 17.037 | 1.123e+09 | 1078 - 1094 | 5 | 146.80 | 147.06 | 147.27 |
| 2821.3949 | TQSLLIVNN(N)ATNVVIK Hex 3 HexNAc 3 | 8.819 | 0.001 | 14.951 | 1.279e+10 | 100 - 116 | 6 | 162.41 | 162.88 | 163.61 |
| 4432.9784 | EGVFVSN(N)GTHWFVTQR Hex 6 HexNAc 5 Neu5Ac 2 | 8.799 | 0.001 | 9.877 | 1.669e+09 | 1078 - 1094 | 7 | 179.71 | 179.85 | 179.99 |
| 3283.3588 | N(N)FTTAPAIC(C)HDGK Fuc 1 Hex 3 HexNAc 6 | 8.753 | 0.001 | 0.000 | 0.000e+00 | 1060 - 1073 | 1 | - | - | - |
| 4503.8806 | NLN(N)ESLIDLQELGK Fuc 1 Hex 6 HexNAc 6 Neu5Ac 2 | 8.681 | 0.001 | 9.122 | 1.479e+08 | 1178 - 1192 | 3 | 213.39 | 213.57 | 214.03 |
| 3889.7942 | DFGGFN(N)FSQILPDPSKPSK Fuc 1 Hex 4 HexNAc 5 | 8.641 | 0.001 | 11.646 | 4.125e+09 | 782 - 801 | 16 | 198.90 | 199.95 | 201.02 |
| 5210.1204 | FPNITNLC(C)PFGEVFN(N)ATR Fuc 1 Hex 9 HexNAc 6 Neu5Ac 1 | 8.635 | 0.001 | 14.076 | 6.933e+08 | 315 - 333 | 1 | 226.73 | 227.17 | 227.61 |
| 4051.3071 | NLN(N)ESLIDLQELGK Fuc 1 Hex 5 HexNAc 6 Neu5Ac 1 | 8.598 | 0.001 | -5.982 | 9.736e+07 | 1178 - 1192 | 5 | 109.73 | 109.81 | 199.28 |
| 3988.9064 | TPPIKDFGGFN(N)FSQILPDPSKPSKR Hex 5 HexNAc 2 | 8.540 | 0.001 | 13.133 | 1.386e+09 | 777 - 802 | 10 | 175.30 | 175.79 | 176.69 |
| 3134.4234 | DFGGFN(N)FSQILPDPSKPSK Hex 4 HexNAc 2 | 8.532 | 0.001 | 14.160 | 2.405e+10 | 782 - 801 | 20 | 203.18 | 204.27 | 205.64 |
| 3540.5815 | DFGGFN(N)FSQILPDPSKPSK Hex 4 HexNAc 4 | 8.506 | 0.001 | 10.768 | 4.588e+09 | 782 - 801 | 12 | 203.27 | 203.53 | 204.83 |
| 3936.6852 | DFGGFN(N)FSQILPDPSKPSK Fuc 1 Hex 5 HexNAc 3 Neu5Ac 1 | 8.430 | 0.001 | 12.392 | 3.964e+09 | 782 - 801 | 6 | 219.39 | 219.95 | 220.30 |
| 6537.8719 | SSVLHSTQDLFLPFFSN(N)VTWFHAIHVSGTNGTK Fuc 1 Hex 8 HexNAc 7 | 8.363 | 0.001 | 13.880 | 9.937e+09 | 31 - 64 | 2 | 208.37 | 208.87 | 210.10 |
| 6537.8719 | SSVLHSTQDLFLPFFSNVTWFHAIHVSGTN(N)GTK Fuc 1 Hex 8 HexNAc 7 | 8.363 | 0.001 | 13.880 | 9.937e+09 | 31 - 64 | 2 | 208.37 | 208.87 | 210.10 |
| 4412.8111 | YNEN(N)GTITDAVDC(C)ALDPLSETK Hex 6 HexNAc 5 | 8.289 | 0.001 | 15.818 | 1.762e+09 | 265 - 287 | 5 | 187.07 | 187.39 | 187.75 |
| 6683.9297 | SSVLHSTQDLFLPFFSNVTWFHAIHVSGTN(N)GTK Fuc 2 Hex 8 HexNAc 7 | 8.231 | 0.001 | 13.922 | 8.107e+09 | 31 - 64 | 2 | 207.68 | 208.46 | 209.20 |
| 4035.8633 | TPPIKDFGGFN(N)FSQILPDPSKPSK Hex 5 HexNAc 3 | 8.206 | 0.001 | 12.703 | 1.786e+09 | 777 - 801 | 7 | 187.26 | 187.82 | 188.32 |
| 3994.7259 | DFGGFN(N)FSQILPDPSKPSK Fuc 2 Hex 5 HexNAc 4 | 8.180 | 0.001 | 14.054 | 4.828e+08 | 782 - 801 | 6 | 198.24 | 198.59 | 199.24 |
| 2891.1911 | N(N)FTTAPAIC(C)HDGK Hex 4 HexNAc 4 | 8.180 | 0.001 | 10.926 | 4.014e+09 | 1060 - 1073 | 6 | 85.36 | 85.97 | 87.28 |
| 4187.6503 | N(N)FTTAPAIC(C)HDGK Fuc 1 Hex 5 HexNAc 6 Neu5Ac 2 | 8.120 | 0.001 | 11.981 | 1.645e+08 | 1060 - 1073 | 1 | 112.11 | 112.25 | 112.44 |
| 4558.8672 | YNEN(N)GTITDAVDC(C)ALDPLSETK Fuc 1 Hex 6 HexNAc 5 | 8.110 | 0.001 | 16.187 | 1.623e+10 | 265 - 287 | 5 | 186.61 | 186.85 | 187.63 |
| 3389.6118 | TQSLLIVNN(N)ATNVVIK Hex 4 HexNAc 5 | 8.055 | 0.001 | -4.973 | 3.141e+09 | 100 - 116 | 13 | 159.70 | 161.01 | 243.64 |
| 4213.7383 | EGVFVSN(N)GTHWFVTQR Fuc 1 Hex 5 HexNAc 4 Neu5Ac 2 | 8.015 | 0.001 | 13.328 | 6.261e+08 | 1078 - 1094 | 5 | 180.47 | 180.70 | 181.91 |
| 3938.7098 | DFGGFN(N)FSQILPDPSKPSKR Hex 8 HexNAc 2 | 7.993 | 0.001 | 13.579 | 5.060e+08 | 782 - 802 | 2 | 184.50 | 185.06 | 185.36 |
| 3418.3757 | N(N)FTTAPAIC(C)HDGK Hex 6 HexNAc 5 | 7.978 | 0.001 | 15.663 | 5.269e+08 | 1060 - 1073 | 6 | 82.85 | 83.08 | 83.44 |
| 3978.7111 | DFGGFN(N)FSQILPDPSKPSK Fuc 3 Hex 4 HexNAc 4 | 7.953 | 0.001 | -18.010 | 1.098e+07 | 782 - 801 | 2 | 219.22 | 219.22 | 219.22 |
| 3275.5646 | TQSLLIVNN(N)ATNVVIK Fuc 2 Hex 4 HexNAc 3 | 7.949 | 0.001 | 10.738 | 9.464e+08 | 100 - 116 | 6 | 160.82 | 161.01 | 161.69 |
| 3640.6945 | TQSLLIVNN(N)ATNVVIK Fuc 2 Hex 5 HexNAc 4 | 7.949 | 0.001 | 13.215 | 2.085e+09 | 100 - 116 | 3 | 159.29 | 159.39 | 160.01 |
| 3883.7429 | DFGGFN(N)FSQILPDPSKPSKR Fuc 1 Hex 3 HexNAc 5 | 7.924 | 0.001 | 13.171 | 3.130e+08 | 782 - 802 | 2 | 183.49 | 184.01 | 184.38 |
| 3331.5272 | DFGGFN(N)FSQILPDPSKPSKR Hex 3 HexNAc 3 | 7.904 | 0.001 | 12.409 | 1.794e+08 | 782 - 802 | 1 | 187.58 | 187.96 | 188.06 |
| 4133.8137 | DFGGFN(N)FSQILPDPSKPSKR Fuc 1 Hex 4 HexNAc 4 Neu5Ac 1 | 7.892 | 0.001 | 13.119 | 5.791e+08 | 782 - 802 | 2 | 200.25 | 200.57 | 201.07 |
| 7070.9771 | SSVLHSTQDLFLPFFSNVTWFHAIHVSGTN(N)GTK @sulfate 1 Fuc 1 Hex 9 HexNAc 7 Neu5Ac 1 | 7.881 | 0.001 | 14.239 | 3.111e+09 | 31 - 64 | 2 | 231.94 | 232.62 | 233.10 |
| 3256.3227 | N(N)FTTAPAIC(C)HDGK Hex 5 HexNAc 5 | 7.880 | 0.001 | 10.088 | 5.237e+08 | 1060 - 1073 | 9 | 83.08 | 83.30 | 84.50 |
| 4794.9777 | NLN(N)ESLIDLQELGK Hex 5 HexNAc 6 Neu5Ac 4 | 7.829 | 0.001 | 9.605 | 4.402e+08 | 1178 - 1192 | 4 | 238.01 | 240.73 | 240.88 |
| 4553.8937 | FPNITNLC(C)PFGEVFN(N)ATR Fuc 1 Hex 8 HexNAc 5 | 7.821 | 0.001 | 13.582 | 6.631e+08 | 315 - 333 | 4 | 209.58 | 210.03 | 210.38 |
| 3680.6635 | DFGGFN(N)FSQILPDPSKPSKR Fuc 1 Hex 3 HexNAc 4 | 7.806 | 0.001 | 14.974 | 6.143e+08 | 782 - 802 | 5 | 183.72 | 184.18 | 184.67 |
| 4566.8965 | VYSSANN(N)C(C)TFEYVSQPFLMDLEGK Fuc 1 Hex 5 HexNAc 4 | 7.791 | 0.001 | 17.610 | 4.209e+09 | 145 - 169 | 7 | 211.04 | 211.50 | 212.06 |
| 3477.6340 | TQSLLIVNN(N)ATNVVIK Hex 4 HexNAc 4 Neu5Ac 1 | 7.755 | 0.001 | 13.356 | 1.289e+10 | 100 - 116 | 12 | 178.85 | 179.95 | 181.94 |
| 2801.2599 | NLN(N)ESLIDLQELGK Hex 5 HexNAc 2 | 7.744 | 0.001 | 12.979 | 2.847e+08 | 1178 - 1192 | 4 | 176.79 | 177.09 | 177.17 |
| 3688.5860 | EGVFVSN(N)GTHWFVTQR Hex 5 HexNAc 5 | 7.744 | 0.001 | 13.566 | 1.151e+09 | 1078 - 1094 | 7 | 147.45 | 147.87 | 148.67 |
| 4461.8660 | VYSSANN(N)C(C)TFEYVSQPFLMDLEGK Hex 4 HexNAc 5 | 7.728 | 0.001 | 15.650 | 3.193e+09 | 145 - 169 | 7 | 211.83 | 212.70 | 213.36 |
| 4607.9232 | VYSSANN(N)C(C)TFEYVSQPFLMDLEGK Fuc 1 Hex 4 HexNAc 5 | 7.669 | 0.001 | 16.369 | 8.616e+09 | 145 - 169 | 11 | 211.21 | 212.23 | 213.16 |
| 6845.9798 | SSVLHSTQDLFLPFFSNVTWFHAIHVSGTN(N)GTK Fuc 2 Hex 9 HexNAc 7 | 7.638 | 0.001 | 14.046 | 6.113e+09 | 31 - 64 | 6 | 207.10 | 207.51 | 208.83 |
| 3201.2975 | N(N)FTTAPAIC(C)HDGK Fuc 1 Hex 5 HexNAc 4 | 7.580 | 0.001 | 0.000 | 0.000e+00 | 1060 - 1073 | 1 | - | - | - |
| 3477.5546 | DFGGFN(N)FSQILPDPSKPSKR Fuc 1 Hex 3 HexNAc 3 | 7.547 | 0.001 | 8.804 | 1.107e+08 | 782 - 802 | 2 | 186.70 | 186.76 | 186.81 |
| 4944.0567 | FPNITNLC(C)PFGEVFN(N)ATR Fuc 2 Hex 7 HexNAc 7 | 7.543 | 0.001 | 13.909 | 7.550e+08 | 315 - 333 | 2 | 207.59 | 208.06 | 208.39 |
| 3881.6147 | EGVFVSN(N)GTHWFVTQR Fuc 1 Hex 6 HexNAc 3 Neu5Ac 1 | 7.522 | 0.001 | 16.417 | 3.179e+09 | 1078 - 1094 | 7 | 161.98 | 162.21 | 163.49 |
| 2933.2188 | N(N)FTTAPAIC(C)HDGK Hex 3 HexNAc 5 | 7.470 | 0.001 | 0.000 | 0.000e+00 | 1060 - 1073 | 3 | - | - | - |
| 4430.8580 | DFGGFN(N)FSQILPDPSKPSK Fuc 1 Hex 5 HexNAc 4 Neu5Ac 2 | 7.459 | 0.001 | 14.417 | 4.514e+08 | 782 - 801 | 2 | 239.53 | 239.94 | 240.33 |
| 3482.4941 | DFGGFN(N)FSQILPDPSKPSK Hex 6 HexNAc 2 | 7.387 | 0.001 | 0.000 | 0.000e+00 | 782 - 801 | 6 | - | - | - |
| 4849.9626 | YNEN(N)GTITDAVDC(C)ALDPLSETK Fuc 1 Hex 6 HexNAc 5 Neu5Ac 1 | 7.334 | 0.001 | 16.197 | 2.700e+09 | 265 - 287 | 2 | 204.77 | 205.44 | 206.00 |
| 3996.9803 | EGVFVSN(N)GTHWFVTQR Fuc 1 Hex 6 HexNAc 5 | 7.229 | 0.001 | -4.592 | 1.473e+09 | 1078 - 1094 | 14 | 111.52 | 146.45 | 147.03 |
| 4072.7105 | YNEN(N)GTITDAVDC(C)ALDPLSETK Fuc 1 Hex 3 HexNAc 5 | 7.219 | 0.001 | 15.337 | -4.581e+08 | 265 - 287 | 12 | 189.38 | 189.57 | 190.54 |
| 3869.6333 | YNEN(N)GTITDAVDC(C)ALDPLSETK Fuc 1 Hex 3 HexNAc 4 | 7.191 | 0.001 | -1.403 | 4.063e+09 | 265 - 287 | 7 | 152.36 | 190.39 | 190.66 |
| 3183.3127 | N(N)FTTAPAIC(C)HDGK Fuc 2 Hex 4 HexNAc 4 | 7.027 | 0.001 | -3.413 | 2.294e+08 | 1060 - 1073 | 6 | 57.56 | 86.60 | 86.72 |
| 3485.4380 | N(N)FTTAPAIC(C)HDGK Fuc 1 Hex 3 HexNAc 7 | 7.003 | 0.001 | 0.000 | 0.000e+00 | 1060 - 1073 | 1 | - | - | - |
| 4108.8146 | DFGGFN(N)FSQILPDPSKPSKR Hex 6 HexNAc 3 Neu5Ac 1 | 6.877 | 0.001 | -4.058 | 2.219e+08 | 782 - 802 | 4 | 100.16 | 202.71 | 202.71 |
| 3998.6804 | EGVFVSN(N)GTHWFVTQR Fuc 1 Hex 6 HexNAc 5 | 6.873 | 0.001 | 0.000 | 0.000e+00 | 1078 - 1094 | 2 | - | - | - |
| 4782.9445 | EGVFVSN(N)GTHWFVTQR Fuc 1 Hex 6 HexNAc 6 Neu5Ac 2 | 6.870 | 0.001 | 0.000 | 0.000e+00 | 1078 - 1094 | 1 | - | - | - |
| 3493.5780 | DFGGFN(N)FSQILPDPSKPSKR Hex 4 HexNAc 3 | 6.839 | 0.001 | 10.771 | 6.890e+08 | 782 - 802 | 3 | 186.65 | 186.86 | 187.58 |
| 3842.7166 | DFGGFN(N)FSQILPDPSKPSKR Fuc 1 Hex 4 HexNAc 4 | 6.827 | 0.001 | 11.197 | 2.518e+08 | 782 - 802 | 5 | 182.88 | 183.59 | 184.10 |
| 4014.6793 | VYSSANN(N)C(C)TFEYVSQPFLMDLEGK Hex 5 HexNAc 2 | 6.782 | 0.001 | 18.324 | 2.314e+10 | 145 - 169 | 9 | 213.39 | 213.77 | 215.69 |
| 4445.8701 | VYSSANN(N)C(C)TFEYVSQPFLMDLEGK Fuc 1 Hex 3 HexNAc 5 | 6.771 | 0.001 | 16.020 | 6.395e+09 | 145 - 169 | 8 | 212.38 | 213.28 | 213.75 |
| 3639.6962 | TQSLLIVNN(N)ATNVVIK Hex 5 HexNAc 4 Neu5Ac 1 | 6.728 | 0.001 | 13.786 | 6.191e+09 | 100 - 116 | 11 | 177.44 | 177.63 | 178.69 |
| 4474.8451 | EGVFVSN(N)GTHWFVTQR Hex 5 HexNAc 6 Neu5Ac 2 | 6.695 | 0.001 | 0.000 | 0.000e+00 | 1078 - 1094 | 3 | - | - | - |
| 4769.9767 | VYSSANN(N)C(C)TFEYVSQPFLMDLEGK Fuc 1 Hex 5 HexNAc 5 | 6.681 | 0.001 | 15.615 | 2.375e+09 | 145 - 169 | 5 | 210.38 | 211.50 | 212.00 |
| 7137.0780 | SSVLHSTQDLFLPFFSNVTWFHAIHVSGTN(N)GTK Fuc 2 Hex 9 HexNAc 7 Neu5Ac 1 | 6.676 | 0.001 | 14.919 | 4.492e+09 | 31 - 64 | 4 | 217.29 | 218.28 | 219.31 |
| 7137.0780 | SSVLHSTQDLFLPFFSN(N)VTWFHAIHVSGTNGTK Fuc 2 Hex 9 HexNAc 7 Neu5Ac 1 | 6.676 | 0.001 | 14.919 | 4.492e+09 | 31 - 64 | 4 | 217.29 | 218.28 | 219.31 |
| 3665.2358 | TQSLLIVNN(N)ATNVVIK Fuc 1 Hex 3 HexNAc 5 Neu5Ac 1 | 6.658 | 0.001 | 9.186 | 3.227e+07 | 100 - 116 | 3 | 204.73 | 204.81 | 204.98 |
| 3632.5446 | EGVFVSN(N)GTHWFVTQR Fuc 1 Hex 5 HexNAc 4 | 6.652 | 0.001 | 0.000 | 0.000e+00 | 1078 - 1094 | 2 | - | - | - |
| 4711.9353 | VYSSANN(N)C(C)TFEYVSQPFLMDLEGK Hex 5 HexNAc 4 Neu5Ac 1 | 6.649 | 0.001 | 13.685 | 2.472e+09 | 145 - 169 | 6 | 230.60 | 232.04 | 233.38 |
| 4489.0120 | TPPIKDFGGFN(N)FSQILPDPSKPSK Hex 6 HexNAc 3 Neu5Ac 1 | 6.627 | 0.001 | 12.891 | 7.899e+08 | 777 - 801 | 2 | 201.69 | 202.27 | 202.54 |
| 4146.6242 | N(N)FTTAPAIC(C)HDGK Fuc 1 Hex 6 HexNAc 5 Neu5Ac 2 | 6.620 | 0.001 | 11.861 | 3.756e+08 | 1060 - 1073 | 2 | 111.58 | 111.73 | 112.01 |
| 4434.8190 | EGVFVSN(N)GTHWFVTQR Hex 6 HexNAc 5 Neu5Ac 2 | 6.616 | 0.001 | 0.000 | 0.000e+00 | 1078 - 1094 | 4 | - | - | - |
| 3590.5192 | EGVFVSN(N)GTHWFVTQR Fuc 1 Hex 6 HexNAc 3 | 6.474 | 0.001 | 15.229 | 1.810e+09 | 1078 - 1094 | 7 | 147.61 | 147.79 | 148.44 |
| 3574.5206 | EGVFVSN(N)GTHWFVTQR Fuc 2 Hex 5 HexNAc 3 | 6.424 | 0.001 | 10.587 | 1.767e+08 | 1078 - 1094 | 4 | 147.63 | 147.63 | 148.39 |
| 4998.0580 | NLN(N)ESLIDLQELGK Fuc 1 Hex 6 HexNAc 7 Neu5Ac 3 | 6.328 | 0.001 | 12.317 | 2.004e+08 | 1178 - 1192 | 2 | 240.58 | 240.73 | 240.94 |
| 3864.6632 | DFGGFN(N)FSQILPDPSKPSK Hex 6 HexNAc 4 | 6.287 | 0.001 | 8.043 | 8.773e+07 | 782 - 801 | 3 | 202.23 | 202.54 | 202.61 |
| 4484.8338 | YNEN(N)GTITDAVDC(C)ALDPLSETK Fuc 1 Hex 5 HexNAc 4 Neu5Ac 1 | 6.252 | 0.001 | 12.706 | 4.763e+08 | 265 - 287 | 2 | 206.51 | 206.82 | 207.51 |
| 3581.5807 | DFGGFN(N)FSQILPDPSKPSK Hex 3 HexNAc 5 | 6.251 | 0.001 | 12.979 | 6.473e+09 | 782 - 801 | 2 | 203.95 | 204.31 | 204.77 |
| 3955.6950 | DFGGFN(N)FSQILPDPSKPSK Hex 6 HexNAc 3 Neu5Ac 1 | 6.230 | 0.001 | 0.000 | 0.000e+00 | 782 - 801 | 2 | - | - | - |
| 3897.5577 | N(N)FTTAPAIC(C)HDGK Fuc 3 Hex 5 HexNAc 6 | 6.225 | 0.001 | -20.042 | 9.966e+06 | 1060 - 1073 | 1 | 97.01 | 97.01 | 97.09 |
| 3704.6142 | DFGGFN(N)FSQILPDPSKPSK Hex 5 HexNAc 4 | 6.223 | 0.001 | 0.000 | 0.000e+00 | 782 - 801 | 4 | - | - | - |
| 3947.7275 | DFGGFN(N)FSQILPDPSKPSKR Fuc 2 Hex 5 HexNAc 3 | 6.216 | 0.001 | -21.823 | 1.785e+07 | 782 - 802 | 1 | 203.79 | 203.79 | 203.79 |
| 5438.2331 | FPNITNLC(C)PFGEVFN(N)ATR Fuc 2 Hex 7 HexNAc 8 Neu5Ac 1 | 6.114 | 0.001 | 14.144 | -6.742e+08 | 315 - 333 | 5 | 224.53 | 225.89 | 226.13 |
| 5002.2966 | DFGGFN(N)FSQILPDPSKPSKRSFIEDLLFNK Hex 7 HexNAc 2 | 6.102 | 0.001 | 0.000 | 0.000e+00 | 782 - 812 | 1 | - | - | - |
| 2435.0245 | QC(C)VN(N)LTTR Fuc 1 Hex 3 HexNAc 4 | 5.999 | 0.011 | 13.974 | 1.738e+08 | 0 - 8 | 2 | 113.22 | 113.43 | 113.58 |
| 3946.7247 | DFGGFN(N)FSQILPDPSKPSKR Hex 5 HexNAc 3 Neu5Ac 1 | 5.962 | 0.002 | 9.240 | 5.108e+08 | 782 - 802 | 2 | 203.40 | 204.05 | 204.05 |
| 3809.5927 | EGVFVSN(N)GTHWFVTQR @sulfate 1 Hex 4 HexNAc 6 | 5.951 | 0.002 | 12.279 | 1.013e+08 | 1078 - 1094 | 2 | 147.16 | 147.27 | 147.38 |
| 4932.0297 | VYSSANN(N)C(C)TFEYVSQPFLMDLEGK Fuc 1 Hex 6 HexNAc 5 | 5.904 | 0.002 | 16.792 | 1.554e+09 | 145 - 169 | 2 | 209.81 | 210.06 | 210.52 |
| 5180.2863 | TPPIKDFGGFN(N)FSQILPDPSKPSK Fuc 3 Hex 8 HexNAc 4 | 5.898 | 0.001 | 13.786 | 8.950e+08 | 777 - 801 | 9 | 203.84 | 204.21 | 204.53 |
| 4712.9559 | VYSSANN(N)C(C)TFEYVSQPFLMDLEGK Fuc 2 Hex 5 HexNAc 4 | 5.827 | 0.002 | 14.507 | 1.923e+09 | 145 - 169 | 4 | 210.56 | 210.85 | 211.46 |
| 4676.0959 | TPPIKDFGGFN(N)FSQILPDPSKPSK Fuc 1 Hex 5 HexNAc 4 Neu5Ac 1 | 5.817 | 0.002 | 13.525 | 1.118e+09 | 777 - 801 | 6 | 198.52 | 198.94 | 199.40 |
| 3938.6371 | EGVFVSN(N)GTHWFVTQR Hex 6 HexNAc 4 Neu5Ac 1 | 5.676 | 0.002 | 11.189 | 1.866e+08 | 1078 - 1094 | 4 | 163.05 | 163.40 | 163.61 |
| 4197.9152 | TPPIKDFGGFN(N)FSQILPDPSKPSK Hex 6 HexNAc 3 | 5.668 | 0.002 | 13.470 | 8.389e+08 | 777 - 801 | 3 | 187.05 | 187.49 | 187.86 |
| 4053.7635 | EGVFVSN(N)GTHWFVTQR Hex 6 HexNAc 6 | 5.665 | 0.002 | 13.468 | 1.116e+09 | 1078 - 1094 | 9 | 146.45 | 146.88 | 147.27 |
| 4722.1979 | GWIFGTTLDSKTQSLLIVNN(N)ATNVVIK @sulfate 1 Hex 5 HexNAc 3 Neu5Ac 1 | 5.652 | 0.002 | 0.000 | 0.000e+00 | 89 - 116 | 1 | - | - | - |
| 4067.6808 | EGVFVSN(N)GTHWFVTQR Hex 5 HexNAc 4 Neu5Ac 2 | 5.646 | 0.002 | 11.314 | 6.373e+08 | 1078 - 1094 | 7 | 180.82 | 182.80 | 183.11 |
| 4326.9992 | TPPIKDFGGFN(N)FSQILPDPSKPSK Hex 5 HexNAc 3 Neu5Ac 1 | 5.642 | 0.002 | -5.196 | 2.385e+08 | 777 - 801 | 5 | 202.27 | 202.74 | 208.25 |
| 3970.5927 | N(N)FTTAPAIC(C)HDGK Fuc 1 Hex 6 HexNAc 7 | 5.625 | 0.002 | 14.802 | 4.635e+08 | 1060 - 1073 | 3 | 82.77 | 83.03 | 83.21 |
| 4687.9116 | YNEN(N)GTITDAVDC(C)ALDPLSETK Fuc 1 Hex 5 HexNAc 5 Neu5Ac 1 | 5.622 | 0.002 | 13.483 | 1.491e+09 | 265 - 287 | 4 | 205.82 | 206.14 | 206.77 |
| 4271.7625 | EGVFVSN(N)GTHWFVTQR Fuc 2 Hex 5 HexNAc 5 Neu5Ac 1 | 5.620 | 0.002 | -19.406 | 1.528e+07 | 1078 - 1094 | 3 | 181.01 | 181.01 | 181.01 |
| 4648.9486 | VYSSANN(N)C(C)TFEYVSQPFLMDLEGK Fuc 1 Hex 3 HexNAc 6 | 5.612 | 0.002 | 12.586 | 6.476e+08 | 145 - 169 | 2 | 211.76 | 212.23 | 212.77 |
| 3242.3325 | N(N)FTTAPAIC(C)HDGK Fuc 1 Hex 4 HexNAc 5 | 5.611 | 0.002 | 0.000 | 0.000e+00 | 1060 - 1073 | 5 | - | - | - |
| 2836.1781 | N(N)FTTAPAIC(C)HDGK Fuc 1 Hex 4 HexNAc 3 | 5.608 | 0.002 | 0.000 | 0.000e+00 | 1060 - 1073 | 2 | - | - | - |
| 2719.1145 | QC(C)VN(N)LTTR Fuc 1 Hex 6 HexNAc 3 | 5.572 | 0.011 | 10.159 | 5.600e+08 | 0 - 8 | 3 | 110.86 | 111.05 | 111.41 |
| 4638.9083 | EGVFVSN(N)GTHWFVTQR Hex 6 HexNAc 6 Neu5Ac 2 | 5.566 | 0.002 | 0.000 | 0.000e+00 | 1078 - 1094 | 2 | - | - | - |
| 3926.6524 | YNEN(N)GTITDAVDC(C)ALDPLSETK Hex 3 HexNAc 5 | 5.557 | 0.002 | 16.476 | 1.615e+09 | 265 - 287 | 6 | 189.99 | 190.24 | 190.57 |
| 5239.3956 | SSVLHSTQDLFLPFFSNVTWFHAIHVSGTN(N)GTK Fuc 1 Hex 5 HexNAc 3 | 5.554 | 0.002 | 12.955 | 3.453e+08 | 31 - 64 | 1 | 213.19 | 213.47 | 213.67 |
| 3645.6233 | DFGGFN(N)FSQILPDPSKPSK Fuc 1 Hex 5 HexNAc 3 | 5.501 | 0.002 | 10.210 | 1.986e+09 | 782 - 801 | 15 | 200.32 | 203.25 | 203.69 |
| 3891.6467 | EGVFVSN(N)GTHWFVTQR Hex 5 HexNAc 6 | 5.487 | 0.002 | 12.547 | 7.584e+08 | 1078 - 1094 | 6 | 146.80 | 147.48 | 148.04 |
| 3216.2990 | N(N)FTTAPAIC(C)HDGK Hex 6 HexNAc 4 | 5.479 | 0.002 | 0.000 | 0.000e+00 | 1060 - 1073 | 1 | - | - | - |
| 4037.7050 | EGVFVSN(N)GTHWFVTQR Fuc 1 Hex 5 HexNAc 6 | 5.419 | 0.002 | 13.258 | 6.434e+08 | 1078 - 1094 | 4 | 146.57 | 146.94 | 147.36 |
| 4665.9337 | NLN(N)ESLIDLQELGK Fuc 1 Hex 7 HexNAc 6 Neu5Ac 2 | 5.402 | 0.002 | 11.082 | 1.085e+08 | 1178 - 1192 | 1 | 212.50 | 212.88 | 213.05 |
| 4180.7896 | DFGGFN(N)FSQILPDPSKPSK Fuc 1 Hex 4 HexNAc 5 Neu5Ac 1 | 5.396 | 0.002 | 11.137 | 2.119e+08 | 782 - 801 | 4 | 218.64 | 219.22 | 219.53 |
| 4051.7473 | DFGGFN(N)FSQILPDPSKPSK Fuc 1 Hex 5 HexNAc 5 | 5.324 | 0.002 | 10.533 | 1.641e+08 | 782 - 801 | 3 | 198.69 | 198.94 | 199.35 |
| 2631.0834 | N(N)FTTAPAIC(C)HDGK Fuc 1 Hex 4 HexNAc 2 | 5.253 | 0.002 | -21.275 | 3.971e+07 | 1060 - 1073 | 4 | 86.14 | 86.15 | 94.37 |
| 5441.2453 | FPNITNLC(C)PFGEVFN(N)ATR Fuc 2 Hex 7 HexNAc 8 Neu5Ac 1 | 5.236 | 0.002 | 0.000 | 0.000e+00 | 315 - 333 | 1 | - | - | - |
| 6642.9042 | SSVLHSTQDLFLPFFSN(N)VTWFHAIHVSGTNGTK Fuc 2 Hex 9 HexNAc 6 | 5.222 | 0.002 | 12.611 | 1.334e+09 | 31 - 64 | 1 | 207.87 | 208.46 | 208.78 |
| 6642.9042 | SSVLHSTQDLFLPFFSNVTWFHAIHVSGTN(N)GTK Fuc 2 Hex 9 HexNAc 6 | 5.222 | 0.002 | 12.611 | 1.334e+09 | 31 - 64 | 1 | 207.87 | 208.46 | 208.78 |
| 6887.0058 | SSVLHSTQDLFLPFFSNVTWFHAIHVSGTN(N)GTK Fuc 2 Hex 8 HexNAc 8 | 5.221 | 0.002 | 13.025 | 4.050e+09 | 31 - 64 | 5 | 207.13 | 207.51 | 208.83 |
| 2917.2763 | EGVFVSN(N)GTHWFVTQR Hex 4 HexNAc 2 | 5.200 | 0.002 | 10.478 | 5.520e+07 | 1078 - 1094 | 1 | 150.56 | 150.66 | 150.85 |
| 4242.7918 | VYSSANN(N)C(C)TFEYVSQPFLMDLEGK Fuc 1 Hex 3 HexNAc 4 | 5.190 | 0.002 | 17.598 | 4.912e+09 | 145 - 169 | 5 | 213.05 | 213.44 | 214.06 |
| 6828.9648 | SSVLHSTQDLFLPFFSNVTWFHAIHVSGTN(N)GTK Fuc 1 Hex 8 HexNAc 7 Neu5Ac 1 | 5.189 | 0.002 | 15.231 | 3.133e+09 | 31 - 64 | 3 | 218.28 | 218.68 | 219.42 |
| 4299.8122 | VYSSANN(N)C(C)TFEYVSQPFLMDLEGK Hex 3 HexNAc 5 | 5.152 | 0.002 | 15.459 | 2.378e+09 | 145 - 169 | 4 | 213.05 | 213.67 | 214.06 |
| 5149.0876 | VYSSANN(N)C(C)TFEYVSQPFLMDLEGK Fuc 1 Hex 5 HexNAc 4 Neu5Ac 2 | 5.120 | 0.002 | 15.916 | 3.647e+09 | 145 - 169 | 6 | 253.38 | 255.64 | 255.97 |
| 4404.8446 | VYSSANN(N)C(C)TFEYVSQPFLMDLEGK Fuc 1 Hex 4 HexNAc 4 | 5.118 | 0.002 | 15.898 | 2.445e+09 | 145 - 169 | 6 | 211.96 | 212.60 | 212.91 |
| 6095.9254 | SNIIRGWIFGTTLDSKTQSLLIVNN(N)ATNVVIK Fuc 1 Hex 5 HexNAc 8 | 5.031 | 0.002 | 11.923 | 2.004e+09 | 84 - 116 | 2 | 161.49 | 161.67 | 161.95 |
| 5183.2943 | TPPIKDFGGFN(N)FSQILPDPSKPSK Fuc 3 Hex 8 HexNAc 4 | 4.988 | 0.002 | 0.000 | 0.000e+00 | 777 - 801 | 1 | - | - | - |
| 4623.9203 | VYSSANN(N)C(C)TFEYVSQPFLMDLEGK Hex 5 HexNAc 5 | 4.975 | 0.002 | 12.457 | 7.166e+08 | 145 - 169 | 4 | 210.81 | 211.96 | 212.56 |
| 4132.6460 | N(N)FTTAPAIC(C)HDGK Fuc 1 Hex 7 HexNAc 7 | 4.937 | 0.002 | 14.523 | 2.013e+08 | 1060 - 1073 | 3 | 82.57 | 82.74 | 83.01 |
| 3441.3308 | N(N)FTTAPAIC(C)HDGK Fuc 1 Hex 9 HexNAc 2 | 4.910 | 0.002 | -20.707 | 7.653e+06 | 1060 - 1073 | 1 | 83.46 | 83.58 | 83.58 |
| 4318.9403 | TPPIKDFGGFN(N)FSQILPDPSKPSK Hex 8 HexNAc 2 | 4.876 | 0.002 | 11.362 | 9.470e+08 | 777 - 801 | 3 | 186.76 | 186.84 | 187.52 |
| 4222.9927 | TPPIKDFGGFN(N)FSQILPDPSKPSK Fuc 1 Hex 4 HexNAc 4 | 4.800 | 0.002 | 11.488 | 4.554e+08 | 777 - 801 | 6 | 184.73 | 185.46 | 185.48 |
| 4060.6247 | N(N)FTTAPAIC(C)HDGK Hex 5 HexNAc 6 Neu5Ac 2 | 4.764 | 0.002 | 0.000 | 0.000e+00 | 1060 - 1073 | 1 | - | - | - |
| 3808.5601 | N(N)FTTAPAIC(C)HDGK Fuc 1 Hex 5 HexNAc 7 | 4.763 | 0.002 | 12.175 | 2.905e+08 | 1060 - 1073 | 6 | 83.24 | 84.34 | 84.50 |
| 2889.1838 | QC(C)VN(N)LTTR Fuc 3 Hex 4 HexNAc 4 | 4.748 | 0.011 | 15.711 | 1.375e+10 | 0 - 8 | 7 | 133.02 | 133.41 | 134.57 |
| 4525.8584 | YNEN(N)GTITDAVDC(C)ALDPLSETK Fuc 1 Hex 4 HexNAc 5 Neu5Ac 1 | 4.746 | 0.002 | 14.549 | 9.321e+08 | 265 - 287 | 2 | 207.13 | 207.38 | 207.84 |
| 3834.7869 | EGVFVSN(N)GTHWFVTQR Fuc 1 Hex 5 HexNAc 5 | 4.702 | 0.004 | 11.699 | 7.346e+08 | 1078 - 1094 | 10 | 142.20 | 147.27 | 147.74 |
| 4272.7644 | EGVFVSN(N)GTHWFVTQR Hex 5 HexNAc 5 Neu5Ac 2 | 4.688 | 0.005 | 0.000 | 0.000e+00 | 1078 - 1094 | 1 | - | - | - |
| 4418.8269 | EGVFVSN(N)GTHWFVTQR Fuc 1 Hex 5 HexNAc 5 Neu5Ac 2 | 4.681 | 0.005 | 0.000 | 0.000e+00 | 1078 - 1094 | 1 | - | - | - |
| 3404.3867 | N(N)FTTAPAIC(C)HDGK Fuc 1 Hex 5 HexNAc 5 | 4.674 | 0.005 | 0.000 | 0.000e+00 | 1060 - 1073 | 2 | - | - | - |
| 5044.3259 | TPPIKDFGGFN(N)FSQILPDPSKPSK Fuc 4 Hex 5 HexNAc 5 | 4.671 | 0.005 | 0.000 | 0.000e+00 | 777 - 801 | 1 | - | - | - |
| 3055.2520 | N(N)FTTAPAIC(C)HDGK Hex 5 HexNAc 4 | 4.661 | 0.005 | 0.000 | 0.000e+00 | 1060 - 1073 | 1 | - | - | - |
| 4197.7868 | QC(C)VN(N)LTTRTQLPPAYTNSFTR @sulfate 1 Fuc 1 Hex 3 HexNAc 5 | 4.657 | 0.005 | 0.000 | 0.000e+00 | 0 - 21 | 1 | - | - | - |
| 3532.4284 | N(N)FTTAPAIC(C)HDGK Fuc 1 Hex 4 HexNAc 5 Neu5Ac 1 | 4.641 | 0.006 | 0.000 | 0.000e+00 | 1060 - 1073 | 2 | - | - | - |
| 4183.7448 | EGVFVSN(N)GTHWFVTQR Fuc 3 Hex 3 HexNAc 4 Neu5Ac 2 | 4.636 | 0.006 | 6.179 | 2.398e+08 | 1078 - 1094 | 2 | 162.38 | 162.66 | 162.66 |
| 4437.8433 | YNEN(N)GTITDAVDC(C)ALDPLSETK Fuc 1 Hex 4 HexNAc 6 | 4.631 | 0.006 | 11.617 | 2.751e+09 | 265 - 287 | 4 | 187.63 | 188.24 | 188.38 |
| 4923.9994 | YNEN(N)GTITDAVDC(C)ALDPLSETK Fuc 1 Hex 7 HexNAc 6 | 4.608 | 0.006 | 12.301 | 4.062e+08 | 265 - 287 | 1 | 185.26 | 185.42 | 185.50 |
| 3615.5503 | EGVFVSN(N)GTHWFVTQR Fuc 2 Hex 4 HexNAc 4 | 4.604 | 0.006 | 12.034 | 1.489e+08 | 1078 - 1094 | 2 | 147.72 | 147.76 | 148.34 |
| 4416.8749 | EGVFVSN(N)GTHWFVTQR Fuc 1 Hex 5 HexNAc 5 Neu5Ac 2 | 4.595 | 0.006 | 11.931 | 8.743e+08 | 1078 - 1094 | 6 | 180.13 | 180.25 | 181.24 |
| 5147.1372 | FPNITNLC(C)PFGEVFN(N)ATR Fuc 2 Hex 7 HexNAc 8 | 4.579 | 0.006 | 12.807 | 2.559e+09 | 315 - 333 | 2 | 206.23 | 207.03 | 208.25 |
| 4703.9063 | YNEN(N)GTITDAVDC(C)ALDPLSETK Hex 6 HexNAc 5 Neu5Ac 1 | 4.562 | 0.006 | 12.887 | 7.659e+08 | 265 - 287 | 2 | 205.48 | 205.72 | 206.18 |
| 3551.6610 | TQSLLIVNN(N)ATNVVIK Hex 5 HexNAc 5 | 4.540 | 0.006 | 13.522 | 6.632e+08 | 100 - 116 | 5 | 159.51 | 159.74 | 160.59 |
| 4031.7868 | YNEN(N)GTITDAVDC(C)ALDPLSETK Fuc 1 Hex 4 HexNAc 4 | 4.533 | 0.006 | 10.116 | -1.041e+09 | 265 - 287 | 10 | 189.11 | 189.52 | 189.62 |
| 3276.5652 | TQSLLIVNN(N)ATNVVIK Fuc 2 Hex 4 HexNAc 3 | 4.528 | 0.006 | 0.000 | 0.000e+00 | 100 - 116 | 3 | - | - | - |
| 3326.2685 | N(N)FTTAPAIC(C)HDGK @sulfate 1 Hex 3 HexNAc 5 Neu5Ac 1 | 4.501 | 0.006 | 0.000 | 0.000e+00 | 1060 - 1073 | 1 | - | - | - |
| 4250.8334 | YNEN(N)GTITDAVDC(C)ALDPLSETK Hex 5 HexNAc 5 | 4.499 | 0.006 | 11.626 | 1.252e+09 | 265 - 287 | 8 | 187.82 | 188.21 | 188.35 |
| 3984.5718 | N(N)FTTAPAIC(C)HDGK Fuc 1 Hex 5 HexNAc 5 Neu5Ac 2 | 4.489 | 0.006 | 11.827 | 3.145e+08 | 1060 - 1073 | 2 | 112.88 | 113.01 | 113.38 |
| 4578.8699 | EGVFVSN(N)GTHWFVTQR Fuc 1 Hex 6 HexNAc 5 Neu5Ac 2 | 4.489 | 0.006 | 12.560 | 8.037e+08 | 1078 - 1094 | 3 | 177.84 | 177.95 | 178.18 |
| 4214.7422 | EGVFVSN(N)GTHWFVTQR Fuc 3 Hex 5 HexNAc 4 Neu5Ac 1 | 4.482 | 0.006 | -5.045 | 1.231e+07 | 1078 - 1094 | 1 | 180.94 | 181.01 | 181.17 |
| 6699.9310 | SSVLHSTQDLFLPFFSNVTWFHAIHVSGTN(N)GTK Fuc 1 Hex 9 HexNAc 7 | 4.470 | 0.006 | 12.390 | 5.332e+09 | 31 - 64 | 6 | 207.62 | 208.46 | 209.28 |
| 2955.1667 | N(N)FTTAPAIC(C)HDGK @sulfate 1 Fuc 1 Hex 3 HexNAc 4 | 4.466 | 0.006 | -19.707 | 4.728e+06 | 1060 - 1073 | 1 | 84.59 | 84.59 | 84.59 |
| 3565.4874 | EGVFVSN(N)GTHWFVTQR Hex 8 HexNAc 2 | 4.460 | 0.006 | 12.795 | 3.145e+08 | 1078 - 1094 | 2 | 147.65 | 147.79 | 148.14 |
| 3709.5852 | N(N)FTTAPAIC(C)HDGK Hex 6 HexNAc 5 Neu5Ac 1 | 4.435 | 0.006 | 10.562 | 9.570e+07 | 1060 - 1073 | 4 | 96.23 | 96.39 | 96.52 |
| 3743.6530 | DFGGFN(N)FSQILPDPSKPSK Hex 4 HexNAc 5 | 4.424 | 0.006 | -6.275 | 1.823e+08 | 782 - 801 | 11 | 202.20 | 203.56 | 215.72 |
| 3040.2603 | N(N)FTTAPAIC(C)HDGK Fuc 1 Hex 4 HexNAc 4 | 4.416 | 0.006 | 0.000 | 0.000e+00 | 1060 - 1073 | 1 | - | - | - |
| 4753.9801 | VYSSANN(N)C(C)TFEYVSQPFLMDLEGK Fuc 2 Hex 4 HexNAc 5 | 4.405 | 0.006 | 11.035 | 1.410e+09 | 145 - 169 | 3 | 210.81 | 210.96 | 212.20 |
| 3589.4346 | N(N)FTTAPAIC(C)HDGK Fuc 2 Hex 4 HexNAc 6 | 4.395 | 0.006 | -20.078 | 3.363e+06 | 1060 - 1073 | 1 | 83.05 | 83.05 | 83.05 |
| 4420.8401 | VYSSANN(N)C(C)TFEYVSQPFLMDLEGK Hex 5 HexNAc 4 | 4.393 | 0.006 | 14.675 | 1.699e+09 | 145 - 169 | 5 | 211.69 | 211.96 | 213.36 |
| 2914.1470 | N(N)FTTAPAIC(C)HDGK @sulfate 1 Fuc 1 Hex 4 HexNAc 3 | 4.385 | 0.006 | -6.549 | 2.044e+07 | 1060 - 1073 | 1 | 86.78 | 86.78 | 87.07 |
| 2915.1492 | N(N)FTTAPAIC(C)HDGK @sulfate 1 Fuc 2 Hex 3 HexNAc 3 | 4.363 | 0.006 | 0.000 | 0.000e+00 | 1060 - 1073 | 1 | - | - | - |
| 3815.5112 | EGVFVSN(N)GTHWFVTQR @sulfate 1 Fuc 1 Hex 8 HexNAc 2 | 4.359 | 0.006 | 0.000 | 0.000e+00 | 1078 - 1094 | 1 | - | - | - |
| 3322.4870 | DFGGFN(N)FSQILPDPSKPSK Fuc 1 Hex 3 HexNAc 3 | 4.352 | 0.006 | 0.000 | 0.000e+00 | 782 - 801 | 2 | - | - | - |
| 3819.6865 | DFGGFN(N)FSQILPDPSKPSKR Hex 6 HexNAc 3 | 4.324 | 0.007 | 0.000 | 0.000e+00 | 782 - 802 | 1 | - | - | - |
| 3816.8634 | DFGGFN(N)FSQILPDPSKPSK Fuc 3 Hex 3 HexNAc 4 | 4.298 | 0.006 | -5.218 | 1.362e+08 | 782 - 801 | 8 | 172.68 | 200.13 | 200.38 |
| 4421.7843 | NLN(N)ESLIDLQELGK @sulfate 1 Fuc 1 Hex 5 HexNAc 6 Neu5Ac 2 | 4.290 | 0.007 | 11.600 | 1.035e+08 | 1178 - 1192 | 1 | 247.51 | 247.67 | 247.95 |
| 3597.5822 | NLN(N)ESLIDLQELGK Fuc 1 Hex 4 HexNAc 6 | 4.288 | 0.007 | 13.173 | 4.851e+08 | 1178 - 1192 | 4 | 173.73 | 174.01 | 174.74 |
| 4295.8648 | DFGGFN(N)FSQILPDPSKPSKR Fuc 1 Hex 5 HexNAc 4 Neu5Ac 1 | 4.278 | 0.007 | 11.772 | 4.024e+08 | 782 - 802 | 2 | 198.69 | 199.20 | 199.40 |
| 4096.7339 | VYSSANN(N)C(C)TFEYVSQPFLMDLEGK Hex 3 HexNAc 4 | 4.274 | 0.007 | 14.949 | 2.390e+09 | 145 - 169 | 4 | 213.63 | 214.10 | 214.83 |
| 3359.2719 | N(N)FTTAPAIC(C)HDGK @sulfate 1 Fuc 1 Hex 8 HexNAc 2 | 4.268 | 0.007 | -4.711 | 1.204e+07 | 1060 - 1073 | 1 | 99.72 | 99.72 | 99.92 |
| 4695.9406 | VYSSANN(N)C(C)TFEYVSQPFLMDLEGK Fuc 1 Hex 4 HexNAc 4 Neu5Ac 1 | 4.198 | 0.007 | 13.813 | 7.246e+08 | 145 - 169 | 3 | 232.45 | 232.69 | 233.58 |
| 4258.7873 | VYSSANN(N)C(C)TFEYVSQPFLMDLEGK Hex 4 HexNAc 4 | 4.161 | 0.007 | 11.173 | 9.705e+08 | 145 - 169 | 6 | 212.50 | 212.84 | 214.06 |
| 4176.7333 | VYSSANN(N)C(C)TFEYVSQPFLMDLEGK Hex 6 HexNAc 2 | 4.150 | 0.007 | 16.208 | 1.079e+09 | 145 - 169 | 3 | 212.70 | 213.01 | 213.39 |
| 4514.0452 | TPPIKDFGGFN(N)FSQILPDPSKPSK Fuc 1 Hex 4 HexNAc 4 Neu5Ac 1 | 4.134 | 0.007 | 10.801 | 1.733e+09 | 777 - 801 | 2 | 199.90 | 199.97 | 200.51 |
| 4899.0192 | VYSSANN(N)C(C)TFEYVSQPFLMDLEGK Fuc 1 Hex 4 HexNAc 5 Neu5Ac 1 | 4.134 | 0.007 | 10.632 | 8.644e+08 | 145 - 169 | 4 | 230.35 | 232.34 | 233.94 |
| 5223.1210 | VYSSANN(N)C(C)TFEYVSQPFLMDLEGK Fuc 1 Hex 6 HexNAc 5 Neu5Ac 1 | 4.130 | 0.007 | 12.607 | 2.668e+09 | 145 - 169 | 5 | 229.41 | 229.63 | 230.99 |
| 6106.6810 | SSVLHSTQDLFLPFFSNVTWFHAIHVSGTN(N)GTK @sulfate 1 Hex 7 HexNAc 6 | 4.114 | 0.007 | 11.398 | 2.492e+08 | 31 - 64 | 2 | 209.53 | 209.85 | 210.19 |
| 3035.2064 | QC(C)VN(N)LTTR Fuc 1 Hex 6 HexNAc 3 Neu5Ac 1 | 4.023 | 0.011 | 0.000 | 0.000e+00 | 0 - 8 | 1 | - | - | - |
| 2930.1928 | QC(C)VN(N)LTTR Fuc 2 Hex 5 HexNAc 4 | 4.013 | 0.011 | 0.000 | 0.000e+00 | 0 - 8 | 1 | - | - | - |
| 2580.0525 | QC(C)VN(N)LTTR Hex 3 HexNAc 4 Neu5Ac 1 | 3.957 | 0.011 | 16.612 | 8.138e+09 | 0 - 8 | 3 | 111.78 | 112.44 | 114.02 |
| 4379.8001 | YNEN(N)GTITDAVDC(C)ALDPLSETK Hex 4 HexNAc 5 Neu5Ac 1 | 3.877 | 0.007 | 13.495 | 6.966e+08 | 265 - 287 | 4 | 207.93 | 208.22 | 208.52 |
| 4599.8939 | YNEN(N)GTITDAVDC(C)ALDPLSETK Fuc 1 Hex 5 HexNAc 6 | 3.868 | 0.007 | 13.363 | 2.418e+09 | 265 - 287 | 6 | 186.83 | 187.20 | 187.98 |
| 4276.7980 | YNEN(N)GTITDAVDC(C)ALDPLSETK Fuc 1 Hex 3 HexNAc 6 | 3.867 | 0.007 | 0.000 | 0.000e+00 | 265 - 287 | 3 | - | - | - |
| 4129.8828 | YNEN(N)GTITDAVDC(C)ALDPLSETK Hex 3 HexNAc 6 | 3.842 | 0.007 | 13.062 | 1.174e+09 | 265 - 287 | 3 | 189.84 | 189.99 | 190.21 |
| 4322.7782 | YNEN(N)GTITDAVDC(C)ALDPLSETK Fuc 1 Hex 4 HexNAc 4 Neu5Ac 1 | 3.807 | 0.008 | 12.259 | 2.628e+08 | 265 - 287 | 2 | 207.96 | 208.06 | 208.43 |
| 4199.8361 | EGVFVSN(N)GTHWFVTQR Fuc 1 Hex 6 HexNAc 6 | 3.751 | 0.009 | 10.016 | 1.631e+08 | 1078 - 1094 | 5 | 146.29 | 146.69 | 147.09 |
| 3647.6126 | EGVFVSN(N)GTHWFVTQR Hex 6 HexNAc 4 | 3.744 | 0.009 | -5.615 | 3.695e+08 | 1078 - 1094 | 6 | 147.65 | 147.92 | 184.42 |
| 3710.4919 | N(N)FTTAPAIC(C)HDGK Fuc 2 Hex 6 HexNAc 5 | 3.742 | 0.009 | 11.232 | 3.612e+08 | 1060 - 1073 | 6 | 82.39 | 83.35 | 83.53 |
| 4223.6779 | N(N)FTTAPAIC(C)HDGK Hex 6 HexNAc 6 Neu5Ac 2 | 3.740 | 0.009 | 0.000 | 0.000e+00 | 1060 - 1073 | 1 | - | - | - |
| 3891.7011 | DFGGFN(N)FSQILPDPSKPSK Fuc 1 Hex 4 HexNAc 5 | 3.733 | 0.010 | 0.000 | 0.000e+00 | 782 - 801 | 1 | - | - | - |
| 4148.6281 | N(N)FTTAPAIC(C)HDGK Fuc 1 Hex 6 HexNAc 5 Neu5Ac 2 | 3.725 | 0.010 | 0.000 | 0.000e+00 | 1060 - 1073 | 1 | - | - | - |
| 3714.5991 | EGVFVSN(N)GTHWFVTQR Fuc 1 Hex 3 HexNAc 6 | 3.722 | 0.010 | 0.000 | 0.000e+00 | 1078 - 1094 | 1 | - | - | - |
| 3142.4075 | N(N)FTTAPAIC(C)HDGK Fuc 2 Hex 5 HexNAc 3 | 3.721 | 0.010 | -22.357 | 7.386e+07 | 1060 - 1073 | 2 | 58.27 | 86.31 | 86.39 |
| 3428.5799 | TQSLLIVNN(N)ATNVVIK Hex 8 HexNAc 2 | 3.719 | 0.010 | 11.551 | 5.310e+08 | 100 - 116 | 2 | 159.40 | 159.51 | 159.85 |
| 3832.6742 | DFGGFN(N)FSQILPDPSKPSK Fuc 2 Hex 4 HexNAc 4 | 3.713 | 0.011 | 9.680 | 5.285e+08 | 782 - 801 | 3 | 199.98 | 200.25 | 200.65 |
| 6722.9142 | SSVLHSTQDLFLPFFSNVTWFHAIHVSGTN(N)GTK @sulfate 1 Fuc 2 Hex 9 HexNAc 6 | 3.707 | 0.011 | -19.279 | 2.845e+07 | 31 - 64 | 1 | 207.89 | 207.89 | 207.89 |
| 3347.3626 | N(N)FTTAPAIC(C)HDGK Fuc 2 Hex 5 HexNAc 4 | 3.707 | 0.011 | 0.000 | 0.000e+00 | 1060 - 1073 | 1 | - | - | - |
| 5309.1898 | FPNITNLC(C)PFGEVFN(N)ATR Fuc 2 Hex 8 HexNAc 8 | 3.705 | 0.011 | 14.117 | 1.470e+09 | 315 - 333 | 2 | 205.28 | 206.55 | 207.48 |
| 3429.5824 | TQSLLIVNN(N)ATNVVIK Hex 8 HexNAc 2 | 3.697 | 0.011 | 0.000 | 0.000e+00 | 100 - 116 | 1 | - | - | - |
| 3637.4770 | N(N)FTTAPAIC(C)HDGK Fuc 2 Hex 5 HexNAc 4 Neu5Ac 1 | 3.681 | 0.011 | 8.089 | 2.481e+06 | 1060 - 1073 | 1 | 67.74 | 67.84 | 67.87 |
| 3903.6030 | DFGGFN(N)FSQILPDPSKPSK @sulfate 1 Hex 7 HexNAc 3 | 3.677 | 0.012 | 10.782 | 1.190e+08 | 782 - 801 | 3 | 224.97 | 225.27 | 225.55 |
| 3727.5534 | EGVFVSN(N)GTHWFVTQR Hex 9 HexNAc 2 | 3.660 | 0.012 | -5.284 | 3.144e+07 | 1078 - 1094 | 1 | 147.38 | 147.56 | 147.56 |
| 3226.4300 | TQSLLIVNN(N)ATNVVIK @sulfate 1 Hex 5 HexNAc 3 | 3.659 | 0.012 | -20.996 | 4.370e+06 | 100 - 116 | 1 | 152.60 | 152.75 | 152.75 |
| 5043.3293 | DFGGFN(N)FSQILPDPSKPSKRSFIEDLLFNK Hex 6 HexNAc 3 | 3.656 | 0.012 | 0.000 | 0.000e+00 | 782 - 812 | 1 | - | - | - |
| 4619.8957 | EGVFVSN(N)GTHWFVTQR Fuc 1 Hex 5 HexNAc 6 Neu5Ac 2 | 3.654 | 0.012 | 0.000 | 0.000e+00 | 1078 - 1094 | 1 | - | - | - |
| 3680.7044 | TQSLLIVNN(N)ATNVVIK Hex 4 HexNAc 5 Neu5Ac 1 | 3.645 | 0.013 | 9.690 | 3.457e+08 | 100 - 116 | 5 | 177.95 | 178.14 | 178.31 |
| 4147.6242 | N(N)FTTAPAIC(C)HDGK Fuc 3 Hex 6 HexNAc 5 Neu5Ac 1 | 3.645 | 0.013 | -5.616 | 3.969e+07 | 1060 - 1073 | 3 | 111.78 | 111.78 | 112.02 |
| 3295.2960 | N(N)FTTAPAIC(C)HDGK Hex 9 HexNAc 2 | 3.643 | 0.013 | 14.188 | 1.428e+08 | 1060 - 1073 | 3 | 82.79 | 82.94 | 83.19 |
| 3522.5136 | DFGGFN(N)FSQILPDPSKPSK Hex 5 HexNAc 3 | 3.642 | 0.013 | 0.000 | 0.000e+00 | 782 - 801 | 1 | - | - | - |
| 3629.5805 | DFGGFN(N)FSQILPDPSKPSK Hex 4 HexNAc 3 Neu5Ac 1 | 3.642 | 0.013 | 0.000 | 0.000e+00 | 782 - 801 | 1 | - | - | - |
| 3479.6299 | TQSLLIVNN(N)ATNVVIK Hex 4 HexNAc 4 Neu5Ac 1 | 3.641 | 0.013 | 0.000 | 0.000e+00 | 100 - 116 | 2 | - | - | - |
| 4635.8930 | EGVFVSN(N)GTHWFVTQR Hex 6 HexNAc 6 Neu5Ac 2 | 3.639 | 0.013 | 8.153 | 2.193e+08 | 1078 - 1094 | 1 | 178.05 | 178.18 | 178.35 |
| 3510.6372 | TQSLLIVNN(N)ATNVVIK Hex 6 HexNAc 4 | 3.625 | 0.013 | 9.347 | 1.811e+08 | 100 - 116 | 4 | 159.64 | 160.05 | 160.93 |
| 4085.6953 | EGVFVSN(N)GTHWFVTQR Fuc 1 Hex 6 HexNAc 4 Neu5Ac 1 | 3.620 | 0.013 | -3.788 | 2.489e+07 | 1078 - 1094 | 3 | 162.38 | 162.55 | 162.55 |
| 3898.6113 | EGVFVSN(N)GTHWFVTQR Hex 7 HexNAc 3 Neu5Ac 1 | 3.620 | 0.013 | 0.000 | 0.000e+00 | 1078 - 1094 | 1 | - | - | - |
| 3320.4370 | DFGGFN(N)FSQILPDPSKPSK Hex 5 HexNAc 2 | 3.609 | 0.014 | 0.000 | 0.000e+00 | 782 - 801 | 2 | - | - | - |
| 3203.5585 | TQSLLIVNN(N)ATNVVIK Hex 4 HexNAc 4 | 3.594 | 0.015 | 0.000 | 0.000e+00 | 100 - 116 | 1 | - | - | - |
| 2781.1383 | TQSLLIVNN(N)ATNVVIK Hex 4 HexNAc 2 | 3.583 | 0.015 | 9.142 | 3.300e+08 | 100 - 116 | 5 | 161.33 | 161.47 | 163.07 |
| 4263.9723 | TPPIKDFGGFN(N)FSQILPDPSKPSK Fuc 1 Hex 3 HexNAc 5 | 3.574 | 0.016 | 9.384 | 1.747e+08 | 777 - 801 | 3 | 185.18 | 185.50 | 185.50 |
| 5078.0883 | VYSSANN(N)C(C)TFEYVSQPFLMDLEGK Fuc 2 Hex 6 HexNAc 5 | 3.569 | 0.016 | 13.462 | 4.174e+08 | 145 - 169 | 1 | 209.43 | 209.64 | 210.03 |
| 4960.0536 | FPNITNLC(C)PFGEVFN(N)ATR Fuc 1 Hex 8 HexNAc 7 | 3.536 | 0.016 | 13.743 | 9.002e+08 | 315 - 333 | 4 | 207.81 | 208.73 | 209.10 |
| 3521.3241 | N(N)FTTAPAIC(C)HDGK @sulfate 1 Fuc 1 Hex 9 HexNAc 2 | 3.522 | 0.016 | 4.493 | 1.460e+07 | 1060 - 1073 | 2 | 98.31 | 98.35 | 98.55 |
| 3873.8108 | TPPIKDFGGFN(N)FSQILPDPSKPSK Hex 4 HexNAc 3 | 3.522 | 0.016 | 10.567 | 8.851e+08 | 777 - 801 | 3 | 187.92 | 188.24 | 188.35 |
| 3674.5830 | EGVFVSN(N)GTHWFVTQR Fuc 1 Hex 4 HexNAc 5 | 3.521 | 0.016 | 0.000 | 0.000e+00 | 1078 - 1094 | 2 | - | - | - |
| 3279.2792 | N(N)FTTAPAIC(C)HDGK Fuc 1 Hex 8 HexNAc 2 | 3.514 | 0.016 | -21.336 | 2.427e+06 | 1060 - 1073 | 2 | 84.29 | 84.32 | 84.32 |
| 4379.8138 | VYSSANN(N)C(C)TFEYVSQPFLMDLEGK Hex 6 HexNAc 3 | 3.511 | 0.016 | 14.886 | 5.426e+08 | 145 - 169 | 2 | 212.10 | 212.32 | 212.56 |
| 4670.9090 | VYSSANN(N)C(C)TFEYVSQPFLMDLEGK Hex 6 HexNAc 3 Neu5Ac 1 | 3.511 | 0.016 | 14.879 | 4.541e+08 | 145 - 169 | 2 | 232.17 | 232.34 | 232.62 |
| 3332.4014 | EGVFVSN(N)GTHWFVTQR @sulfate 1 Fuc 2 Hex 3 HexNAc 3 | 3.490 | 0.016 | 0.000 | 0.000e+00 | 1078 - 1094 | 1 | - | - | - |
| 4385.0000 | TPPIKDFGGFN(N)FSQILPDPSKPSK Fuc 1 Hex 5 HexNAc 4 | 3.479 | 0.016 | 10.594 | 2.719e+08 | 777 - 801 | 3 | 184.57 | 184.57 | 184.99 |
| 4055.7078 | VYSSANN(N)C(C)TFEYVSQPFLMDLEGK Hex 4 HexNAc 3 | 3.477 | 0.016 | 12.652 | 5.245e+08 | 145 - 169 | 4 | 213.83 | 214.10 | 214.46 |
| 3823.5180 | NN(N)KSWMESEFR Hex 6 HexNAc 7 | 3.464 | 0.016 | 0.000 | 0.000e+00 | 134 - 145 | 1 | - | - | - |
| 3398.3337 | QC(C)VN(N)LTTR Hex 5 HexNAc 5 Neu5Ac 2 | 3.458 | 0.011 | 12.938 | 1.478e+09 | 0 - 8 | 4 | 129.82 | 130.11 | 131.14 |
| 3875.8146 | TPPIKDFGGFN(N)FSQILPDPSKPSK Hex 4 HexNAc 3 | 3.453 | 0.016 | 0.000 | 0.000e+00 | 777 - 801 | 1 | - | - | - |
| 3113.4822 | TQSLLIVNN(N)ATNVVIK Fuc 2 Hex 3 HexNAc 3 | 3.449 | 0.016 | -20.985 | 1.291e+08 | 100 - 116 | 2 | 161.10 | 161.29 | 161.29 |
| 2946.2096 | QC(C)VN(N)LTTR Fuc 2 Hex 4 HexNAc 5 | 3.446 | 0.011 | 14.194 | 5.876e+08 | 0 - 8 | 5 | 109.62 | 110.12 | 111.81 |
| 4157.8884 | TPPIKDFGGFN(N)FSQILPDPSKPSK Hex 7 HexNAc 2 | 3.445 | 0.016 | 0.000 | 0.000e+00 | 777 - 801 | 4 | - | - | - |
| 3439.3797 | QC(C)VN(N)LTTR Hex 4 HexNAc 6 Neu5Ac 2 | 3.444 | 0.011 | 12.996 | 5.737e+08 | 0 - 8 | 3 | 130.39 | 130.85 | 131.35 |
| 4957.0289 | NLN(N)ESLIDLQELGK Fuc 1 Hex 7 HexNAc 6 Neu5Ac 3 | 3.437 | 0.016 | 12.667 | 2.134e+08 | 1178 - 1192 | 1 | 236.36 | 236.99 | 237.41 |
| 2871.1706 | QC(C)VN(N)LTTR Hex 3 HexNAc 4 Neu5Ac 2 | 3.436 | 0.011 | 15.426 | 9.818e+09 | 0 - 8 | 19 | 132.55 | 134.44 | 135.73 |
| 4752.9620 | VYSSANN(N)C(C)TFEYVSQPFLMDLEGK Hex 4 HexNAc 5 Neu5Ac 1 | 3.422 | 0.016 | 10.994 | 1.239e+08 | 145 - 169 | 1 | 232.84 | 233.08 | 233.26 |
| 2703.1197 | QC(C)VN(N)LTTR Fuc 2 Hex 5 HexNAc 3 | 3.422 | 0.011 | 12.109 | -8.442e+08 | 0 - 8 | 1 | 111.01 | 111.19 | 111.52 |
| 2527.0620 | N(N)FTTAPAIC(C)HDGK Hex 3 HexNAc 3 | 3.421 | 0.016 | 0.000 | 0.000e+00 | 1060 - 1073 | 1 | - | - | - |
| 2873.1600 | QC(C)VN(N)LTTR Hex 3 HexNAc 4 Neu5Ac 2 | 3.414 | 0.011 | 0.000 | 0.000e+00 | 0 - 8 | 1 | - | - | - |
| 3236.5161 | QC(C)VN(N)LTTR Fuc 1 Hex 6 HexNAc 4 Neu5Ac 1 | 3.414 | 0.011 | -4.671 | 2.951e+08 | 0 - 8 | 10 | 109.75 | 132.04 | 160.98 |
| 3033.2436 | QC(C)VN(N)LTTR Hex 4 HexNAc 4 Neu5Ac 2 | 3.412 | 0.011 | 12.824 | 1.265e+10 | 0 - 8 | 16 | 130.44 | 132.67 | 134.31 |
| 2864.0894 | QC(C)VN(N)LTTR @sulfate 1 Fuc 1 Hex 5 HexNAc 4 | 3.411 | 0.011 | 0.000 | 0.000e+00 | 0 - 8 | 2 | - | - | - |
| 2930.2038 | QC(C)VN(N)LTTR Fuc 1 Hex 3 HexNAc 5 Neu5Ac 1 | 3.410 | 0.011 | 0.000 | 0.000e+00 | 0 - 8 | 1 | - | - | - |
| 3074.2266 | QC(C)VN(N)LTTR Hex 3 HexNAc 5 Neu5Ac 2 | 3.395 | 0.011 | 12.780 | 8.572e+09 | 0 - 8 | 9 | 130.77 | 131.29 | 135.20 |
| 2801.0843 | QC(C)VN(N)LTTR Fuc 1 Hex 4 HexNAc 5 | 3.394 | 0.011 | -9.700 | 3.300e+07 | 0 - 8 | 7 | 47.55 | 130.20 | 130.31 |
| 2926.3994 | TQSLLIVNN(N)ATNVVIK Fuc 1 Hex 4 HexNAc 2 | 3.383 | 0.016 | -19.338 | 8.649e+06 | 100 - 116 | 1 | 164.77 | 164.77 | 164.77 |
| 2888.2454 | QC(C)VN(N)LTTR Fuc 1 Hex 4 HexNAc 4 Neu5Ac 1 | 3.381 | 0.011 | -3.105 | 3.136e+08 | 0 - 8 | 3 | 94.01 | 134.40 | 134.76 |
| 3052.2594 | QC(C)VN(N)LTTR Fuc 3 Hex 5 HexNAc 4 | 3.378 | 0.011 | 10.653 | 1.592e+09 | 0 - 8 | 3 | 108.92 | 109.04 | 109.22 |
| 2906.1229 | QC(C)VN(N)LTTR Fuc 2 Hex 5 HexNAc 4 | 3.371 | 0.011 | 9.434 | 1.296e+09 | 0 - 8 | 5 | 109.73 | 112.62 | 112.93 |
| 4201.7652 | VYSSANN(N)C(C)TFEYVSQPFLMDLEGK Fuc 1 Hex 4 HexNAc 3 | 3.360 | 0.016 | 12.643 | 2.840e+08 | 145 - 169 | 4 | 213.01 | 213.36 | 213.67 |
| 2638.1040 | QC(C)VN(N)LTTR Fuc 1 Hex 3 HexNAc 5 | 3.356 | 0.011 | 13.104 | 3.087e+08 | 0 - 8 | 4 | 112.34 | 113.75 | 114.02 |
| 3893.6539 | VYSSANN(N)C(C)TFEYVSQPFLMDLEGK Hex 3 HexNAc 3 | 3.342 | 0.016 | 16.360 | 8.432e+08 | 145 - 169 | 4 | 214.72 | 214.97 | 215.45 |
| 4060.7029 | EGVFVSN(N)GTHWFVTQR Fuc 2 Hex 8 HexNAc 3 | 3.338 | 0.016 | -20.511 | 7.995e+06 | 1078 - 1094 | 1 | 161.95 | 161.95 | 161.95 |
| 3888.6796 | NLN(N)ESLIDLQELGK Fuc 1 Hex 4 HexNAc 6 Neu5Ac 1 | 3.331 | 0.016 | 10.399 | 4.037e+08 | 1178 - 1192 | 4 | 193.46 | 193.62 | 193.84 |
| 2598.0835 | QC(C)VN(N)LTTR Fuc 1 Hex 4 HexNAc 4 | 3.329 | 0.011 | 0.000 | 0.000e+00 | 0 - 8 | 1 | - | - | - |
| 3852.6282 | VYSSANN(N)C(C)TFEYVSQPFLMDLEGK Hex 4 HexNAc 2 | 3.329 | 0.016 | 14.676 | 2.821e+08 | 145 - 169 | 1 | 214.76 | 215.00 | 215.24 |
| 3179.2578 | QC(C)VN(N)LTTR Fuc 1 Hex 4 HexNAc 4 Neu5Ac 2 | 3.321 | 0.011 | 14.454 | 4.181e+08 | 0 - 8 | 2 | 131.01 | 131.27 | 131.49 |
| 2418.9996 | QC(C)VN(N)LTTR Fuc 1 Hex 4 HexNAc 3 | 3.307 | 0.011 | 0.000 | 0.000e+00 | 0 - 8 | 1 | - | - | - |
| 3220.2840 | QC(C)VN(N)LTTR Fuc 1 Hex 3 HexNAc 5 Neu5Ac 2 | 3.295 | 0.011 | 13.528 | 6.839e+08 | 0 - 8 | 2 | 131.74 | 131.96 | 132.12 |
| 5081.0803 | FPNITNLC(C)PFGEVFN(N)ATR @sulfate 1 Fuc 1 Hex 7 HexNAc 8 | 3.295 | 0.016 | 10.740 | 1.506e+08 | 315 - 333 | 1 | 206.68 | 206.98 | 207.16 |
| 2573.0636 | QC(C)VN(N)LTTR Hex 6 HexNAc 3 | 3.277 | 0.011 | 8.210 | 6.111e+07 | 0 - 8 | 2 | 112.13 | 112.41 | 112.41 |
| 3723.5736 | YNEN(N)GTITDAVDC(C)ALDPLSETK Hex 3 HexNAc 4 | 3.273 | 0.016 | -6.816 | 2.239e+09 | 265 - 287 | 2 | 98.03 | 191.29 | 191.44 |
| 4764.2849 | GWIFGTTLDSKTQSLLIVNN(N)ATNVVIK Fuc 1 Hex 4 HexNAc 5 | 3.270 | 0.016 | 0.000 | 0.000e+00 | 89 - 116 | 1 | - | - | - |
| 4101.7285 | EGVFVSN(N)GTHWFVTQR Fuc 2 Hex 7 HexNAc 4 | 3.265 | 0.016 | -21.456 | 4.088e+07 | 1078 - 1094 | 1 | 162.70 | 162.70 | 162.70 |
| 4089.1348 | YNEN(N)GTITDAVDC(C)ALDPLSETK Hex 4 HexNAc 5 | 3.265 | 0.016 | -6.857 | 1.286e+09 | 265 - 287 | 10 | 102.60 | 189.40 | 190.39 |
| 4033.6821 | YNEN(N)GTITDAVDC(C)ALDPLSETK Fuc 1 Hex 4 HexNAc 4 | 3.261 | 0.016 | 0.000 | 0.000e+00 | 265 - 287 | 4 | - | - | - |
| 4995.0011 | YNEN(N)GTITDAVDC(C)ALDPLSETK Hex 6 HexNAc 5 Neu5Ac 2 | 3.250 | 0.016 | 14.759 | 2.432e+08 | 265 - 287 | 2 | 226.86 | 227.11 | 227.38 |
| 4542.2903 | YNEN(N)GTITDAVDC(C)ALDPLSETK Hex 5 HexNAc 5 Neu5Ac 1 | 3.240 | 0.016 | 6.616 | 6.465e+08 | 265 - 287 | 5 | 187.01 | 206.74 | 207.25 |
| 3725.5790 | YNEN(N)GTITDAVDC(C)ALDPLSETK Hex 3 HexNAc 4 | 3.233 | 0.016 | 0.000 | 0.000e+00 | 265 - 287 | 1 | - | - | - |
| 3520.4944 | YNEN(N)GTITDAVDC(C)ALDPLSETK Hex 3 HexNAc 3 | 3.220 | 0.016 | 11.950 | 3.487e+08 | 265 - 287 | 4 | 192.10 | 192.23 | 192.51 |
| 4235.0079 | YNEN(N)GTITDAVDC(C)ALDPLSETK Fuc 1 Hex 4 HexNAc 5 | 3.219 | 0.010 | 9.048 | -1.760e+08 | 265 - 287 | 19 | 188.26 | 189.25 | 189.55 |
| 4907.9150 | YNEN(N)GTITDAVDC(C)ALDPLSETK @sulfate 1 Fuc 2 Hex 3 HexNAc 8 | 3.216 | 0.016 | 13.341 | 4.787e+07 | 265 - 287 | 1 | 73.56 | 73.76 | 73.95 |
| 4193.8821 | YNEN(N)GTITDAVDC(C)ALDPLSETK Fuc 1 Hex 5 HexNAc 4 | 3.216 | 0.016 | 13.025 | 3.140e+08 | 265 - 287 | 6 | 188.13 | 188.29 | 188.58 |
| 2191.9295 | QC(C)VN(N)LTTR Fuc 1 Hex 4 HexNAc 2 | 3.211 | 0.011 | 14.630 | 5.167e+09 | 0 - 8 | 3 | 112.34 | 114.14 | 115.13 |
| 2555.0214 | QC(C)VN(N)LTTR Hex 7 HexNAc 2 | 3.208 | 0.011 | 0.000 | 0.000e+00 | 0 - 8 | 1 | - | - | - |
| 3641.7183 | YNEN(N)GTITDAVDC(C)ALDPLSETK Hex 5 HexNAc 2 | 3.196 | 0.016 | -12.682 | 2.462e+08 | 265 - 287 | 3 | 190.54 | 190.78 | 202.61 |
| 2391.9752 | QC(C)VN(N)LTTR Hex 6 HexNAc 2 | 3.186 | 0.011 | 0.000 | 0.000e+00 | 0 - 8 | 2 | - | - | - |
| 4090.7096 | YNEN(N)GTITDAVDC(C)ALDPLSETK Hex 4 HexNAc 5 | 3.185 | 0.016 | 0.000 | 0.000e+00 | 265 - 287 | 3 | - | - | - |
| 3399.3345 | QC(C)VN(N)LTTR Fuc 2 Hex 5 HexNAc 5 Neu5Ac 1 | 3.179 | 0.011 | -19.511 | 3.920e+07 | 0 - 8 | 2 | 130.18 | 130.18 | 130.18 |
| 3885.6272 | YNEN(N)GTITDAVDC(C)ALDPLSETK Hex 4 HexNAc 4 | 3.172 | 0.016 | 13.291 | 6.740e+08 | 265 - 287 | 4 | 190.00 | 190.18 | 190.46 |
| 3927.6476 | YNEN(N)GTITDAVDC(C)ALDPLSETK Hex 3 HexNAc 5 | 3.154 | 0.016 | 0.000 | 0.000e+00 | 265 - 287 | 1 | - | - | - |
| 3075.5936 | QC(C)VN(N)LTTR Fuc 4 Hex 3 HexNAc 5 | 3.102 | 0.011 | 9.180 | 2.098e+08 | 0 - 8 | 8 | 109.02 | 109.02 | 110.21 |
| 2369.9781 | QC(C)VN(N)LTTR Hex 6 HexNAc 2 | 3.083 | 0.011 | 10.445 | 2.553e+07 | 0 - 8 | 2 | 112.89 | 112.93 | 113.01 |
| 3953.6767 | DFGGFN(N)FSQILPDPSKPSK Fuc 2 Hex 6 HexNAc 3 | 3.082 | 0.017 | -21.433 | 9.808e+06 | 782 - 801 | 2 | 222.36 | 222.36 | 222.52 |
| 4841.9520 | VYSSANN(N)C(C)TFEYVSQPFLMDLEGK Fuc 2 Hex 4 HexNAc 4 Neu5Ac 1 | 3.066 | 0.017 | 11.389 | 7.805e+07 | 145 - 169 | 3 | 238.87 | 239.14 | 239.34 |
| 4181.8073 | DFGGFN(N)FSQILPDPSKPSK Hex 3 HexNAc 5 Neu5Ac 2 | 3.039 | 0.017 | 0.000 | 0.000e+00 | 782 - 801 | 1 | - | - | - |
| 2972.4988 | DFGGFN(N)FSQILPDPSKPSK Hex 3 HexNAc 2 | 3.026 | 0.017 | 10.975 | 4.016e+09 | 782 - 801 | 6 | 203.71 | 204.29 | 206.27 |
| 7033.0684 | SSVLHSTQDLFLPFFSNVTWFHAIHVSGTN(N)GTK Fuc 3 Hex 8 HexNAc 8 | 2.996 | 0.017 | 11.825 | 7.878e+08 | 31 - 64 | 1 | 206.91 | 207.19 | 207.74 |
| 3647.5911 | DFGGFN(N)FSQILPDPSKPSK Fuc 1 Hex 5 HexNAc 3 | 2.989 | 0.017 | 0.000 | 0.000e+00 | 782 - 801 | 1 | - | - | - |
| 6991.0184 | SSVLHSTQDLFLPFFSNVTWFHAIHVSGTN(N)GTK Fuc 1 Hex 9 HexNAc 7 Neu5Ac 1 | 2.987 | 0.017 | 13.222 | 4.551e+09 | 31 - 64 | 2 | 217.69 | 218.12 | 219.60 |
| 4973.0558 | VYSSANN(N)C(C)TFEYVSQPFLMDLEGK Fuc 1 Hex 5 HexNAc 6 | 2.986 | 0.017 | 13.928 | 8.593e+08 | 145 - 169 | 3 | 209.88 | 210.19 | 211.11 |
| 6480.8528 | SSVLHSTQDLFLPFFSNVTWFHAIHVSGTN(N)GTK Fuc 2 Hex 8 HexNAc 6 | 2.975 | 0.017 | 10.168 | 2.649e+08 | 31 - 64 | 1 | 208.62 | 208.87 | 209.07 |
| 3599.6713 | TQSLLIVNN(N)ATNVVIK Fuc 2 Hex 6 HexNAc 3 | 2.972 | 0.018 | 9.673 | 2.042e+08 | 100 - 116 | 2 | 159.44 | 159.51 | 159.67 |
| 4811.0032 | VYSSANN(N)C(C)TFEYVSQPFLMDLEGK Fuc 1 Hex 4 HexNAc 6 | 2.963 | 0.018 | 11.819 | 5.704e+08 | 145 - 169 | 2 | 210.71 | 211.26 | 211.96 |
| 3988.8146 | TQSLLIVNN(N)ATNVVIK Fuc 1 Hex 5 HexNAc 5 Neu5Ac 1 | 2.952 | 0.019 | 12.105 | 3.158e+08 | 100 - 116 | 3 | 176.41 | 176.79 | 177.15 |
| 4946.0080 | EGVFVSN(N)GTHWFVTQR Fuc 1 Hex 7 HexNAc 6 Neu5Ac 2 | 2.948 | 0.019 | 0.000 | 0.000e+00 | 1078 - 1094 | 3 | - | - | - |
| 4785.9736 | VYSSANN(N)C(C)TFEYVSQPFLMDLEGK Hex 6 HexNAc 5 | 2.946 | 0.019 | 12.701 | 2.800e+08 | 145 - 169 | 1 | 210.26 | 210.52 | 210.91 |
| 6573.9127 | QIYKTPPIKDFGGFN(N)FSQILPDPSKPSK @sulfate 1 Fuc 4 Hex 7 HexNAc 8 | 2.943 | 0.019 | 5.316 | 7.879e+07 | 773 - 801 | 1 | 233.74 | 233.93 | 234.28 |
| 4550.9022 | VYSSANN(N)C(C)TFEYVSQPFLMDLEGK Fuc 2 Hex 4 HexNAc 4 | 2.943 | 0.019 | 12.826 | 3.367e+08 | 145 - 169 | 2 | 211.79 | 211.96 | 212.43 |
| 4591.9294 | VYSSANN(N)C(C)TFEYVSQPFLMDLEGK Fuc 2 Hex 3 HexNAc 5 | 2.942 | 0.019 | 12.478 | 2.136e+08 | 145 - 169 | 1 | 211.93 | 212.06 | 212.26 |
| 6643.9065 | SSVLHSTQDLFLPFFSNVTWFHAIHVSGTN(N)GTK Fuc 2 Hex 9 HexNAc 6 | 2.941 | 0.019 | 7.694 | 6.708e+07 | 31 - 64 | 1 | 208.22 | 208.62 | 208.87 |
| 3629.4625 | EGVFVSN(N)GTHWFVTQR @sulfate 1 Fuc 1 Hex 7 HexNAc 2 | 2.936 | 0.019 | 10.601 | 8.671e+06 | 1078 - 1094 | 2 | 98.54 | 98.59 | 98.73 |
| 6131.7124 | SSVLHSTQDLFLPFFSNVTWFHAIHVSGTN(N)GTK Fuc 1 Hex 8 HexNAc 5 | 2.934 | 0.019 | 12.539 | 4.150e+08 | 31 - 64 | 1 | 209.70 | 210.06 | 210.32 |
| 4216.8404 | EGVFVSN(N)GTHWFVTQR Hex 7 HexNAc 6 | 2.917 | 0.007 | -7.541 | 8.328e+08 | 1078 - 1094 | 7 | 145.55 | 145.76 | 213.44 |
| 5514.2197 | VYSSANN(N)C(C)TFEYVSQPFLMDLEGK Fuc 1 Hex 6 HexNAc 5 Neu5Ac 2 | 2.913 | 0.020 | 10.864 | 2.246e+09 | 145 - 169 | 5 | 252.61 | 252.89 | 254.92 |
| 4349.7033 | N(N)FTTAPAIC(C)HDGK Fuc 1 Hex 6 HexNAc 6 Neu5Ac 2 | 2.909 | 0.020 | 7.434 | 5.944e+07 | 1060 - 1073 | 1 | 110.32 | 110.53 | 110.56 |
| 3429.4733 | EGVFVSN(N)GTHWFVTQR Fuc 1 Hex 5 HexNAc 3 | 2.903 | 0.020 | 0.000 | 0.000e+00 | 1078 - 1094 | 1 | - | - | - |
| 4944.0018 | EGVFVSN(N)GTHWFVTQR Fuc 1 Hex 7 HexNAc 6 Neu5Ac 2 | 2.899 | 0.020 | 13.743 | 6.414e+08 | 1078 - 1094 | 2 | 176.21 | 176.37 | 177.15 |
| 3827.7644 | TQSLLIVNN(N)ATNVVIK Fuc 1 Hex 4 HexNAc 5 Neu5Ac 1 | 2.892 | 0.020 | 0.000 | 0.000e+00 | 100 - 116 | 1 | - | - | - |
| 2960.1171 | QC(C)VN(N)LTTR @sulfate 1 Fuc 1 Hex 7 HexNAc 3 | 2.891 | 0.011 | 11.463 | 4.544e+07 | 0 - 8 | 1 | 109.88 | 110.16 | 110.27 |
| 3826.7607 | TQSLLIVNN(N)ATNVVIK Fuc 1 Hex 4 HexNAc 5 Neu5Ac 1 | 2.885 | 0.021 | 10.313 | 1.377e+09 | 100 - 116 | 3 | 177.55 | 177.74 | 178.31 |
| 3937.5823 | N(N)FTTAPAIC(C)HDGK Fuc 1 Hex 4 HexNAc 7 Neu5Ac 1 | 2.875 | 0.022 | 10.287 | 7.201e+07 | 1060 - 1073 | 2 | 97.95 | 98.14 | 98.35 |
| 3770.6198 | EGVFVSN(N)GTHWFVTQR Hex 3 HexNAc 7 | 2.875 | 0.022 | 8.476 | 4.262e+07 | 1078 - 1094 | 4 | 148.04 | 148.14 | 148.22 |
| 4100.6884 | EGVFVSN(N)GTHWFVTQR Hex 7 HexNAc 4 Neu5Ac 1 | 2.871 | 0.022 | -3.747 | 2.178e+07 | 1078 - 1094 | 2 | 162.44 | 162.51 | 162.51 |
| 4916.0352 | VYSSANN(N)C(C)TFEYVSQPFLMDLEGK Fuc 2 Hex 5 HexNAc 5 | 2.870 | 0.022 | 11.605 | 3.343e+08 | 145 - 169 | 3 | 210.12 | 211.21 | 211.53 |
| 4210.7712 | YNEN(N)GTITDAVDC(C)ALDPLSETK Hex 6 HexNAc 4 | 2.869 | 0.022 | -7.510 | 2.502e+08 | 265 - 287 | 2 | 188.69 | 188.86 | 188.88 |
| 4117.9169 | TPPIKDFGGFN(N)FSQILPDPSKPSK Hex 3 HexNAc 5 | 2.868 | 0.022 | 10.038 | 1.656e+08 | 777 - 801 | 2 | 187.70 | 188.18 | 188.26 |
| 3783.5085 | N(N)FTTAPAIC(C)HDGK Hex 7 HexNAc 6 | 2.866 | 0.022 | 0.000 | 0.000e+00 | 1060 - 1073 | 1 | - | - | - |
| 4508.8554 | VYSSANN(N)C(C)TFEYVSQPFLMDLEGK Hex 5 HexNAc 3 Neu5Ac 1 | 2.863 | 0.022 | 13.872 | 4.477e+08 | 145 - 169 | 2 | 233.04 | 233.19 | 233.74 |
| 2783.1416 | QC(C)VN(N)LTTR Hex 3 HexNAc 5 Neu5Ac 1 | 2.862 | 0.011 | 14.434 | 2.373e+10 | 0 - 8 | 5 | 111.09 | 111.31 | 114.07 |
| 5004.0517 | VYSSANN(N)C(C)TFEYVSQPFLMDLEGK Fuc 2 Hex 5 HexNAc 4 Neu5Ac 1 | 2.858 | 0.023 | 12.534 | 6.011e+08 | 145 - 169 | 6 | 230.29 | 230.63 | 232.19 |
| 5003.0297 | VYSSANN(N)C(C)TFEYVSQPFLMDLEGK Hex 5 HexNAc 4 Neu5Ac 2 | 2.856 | 0.023 | 14.379 | 3.535e+08 | 145 - 169 | 1 | 254.37 | 254.83 | 255.05 |
| 3361.3540 | N(N)FTTAPAIC(C)HDGK Fuc 1 Hex 6 HexNAc 4 | 2.854 | 0.023 | 12.305 | 5.208e+07 | 1060 - 1073 | 1 | 83.87 | 83.97 | 84.18 |
| 3616.5514 | EGVFVSN(N)GTHWFVTQR Fuc 2 Hex 4 HexNAc 4 | 2.854 | 0.023 | 0.000 | 0.000e+00 | 1078 - 1094 | 1 | - | - | - |
| 4076.8902 | TPPIKDFGGFN(N)FSQILPDPSKPSK Hex 4 HexNAc 4 | 2.851 | 0.024 | 12.495 | 1.325e+08 | 777 - 801 | 2 | 187.60 | 187.82 | 188.09 |
| 4149.8067 | DFGGFN(N)FSQILPDPSKPSKR Hex 5 HexNAc 4 Neu5Ac 1 | 2.848 | 0.024 | 12.436 | 7.628e+07 | 782 - 802 | 3 | 201.83 | 202.02 | 202.42 |
| 4664.9453 | VYSSANN(N)C(C)TFEYVSQPFLMDLEGK Hex 4 HexNAc 6 | 2.840 | 0.026 | 6.379 | 3.795e+07 | 145 - 169 | 3 | 211.65 | 211.65 | 211.93 |
| 3419.5651 | TQSLLIVNN(N)ATNVVIK @sulfate 1 Fuc 2 Hex 3 HexNAc 4 | 2.819 | 0.026 | 0.000 | 0.000e+00 | 100 - 116 | 1 | - | - | - |
| 3387.3998 | EGVFVSN(N)GTHWFVTQR @sulfate 1 Fuc 1 Hex 3 HexNAc 4 | 2.811 | 0.026 | -5.269 | 7.844e+06 | 1078 - 1094 | 1 | 167.59 | 167.62 | 167.77 |
| 3938.5894 | N(N)FTTAPAIC(C)HDGK Fuc 3 Hex 4 HexNAc 7 | 2.809 | 0.026 | 10.954 | 4.736e+08 | 1060 - 1073 | 2 | 170.64 | 170.79 | 171.53 |
| 5182.2785 | QIYKTPPIKDFGGFN(N)FSQILPDPSKPSK @sulfate 1 Fuc 1 Hex 6 HexNAc 4 | 2.807 | 0.026 | 0.000 | 0.000e+00 | 773 - 801 | 1 | - | - | - |
| 4549.8817 | VYSSANN(N)C(C)TFEYVSQPFLMDLEGK Hex 4 HexNAc 4 Neu5Ac 1 | 2.790 | 0.027 | 12.964 | 1.722e+08 | 145 - 169 | 2 | 233.04 | 233.30 | 233.66 |
| 3193.2606 | N(N)FTTAPAIC(C)HDGK @sulfate 1 Hex 4 HexNAc 5 | 2.773 | 0.028 | 0.000 | 0.000e+00 | 1060 - 1073 | 1 | - | - | - |
| 3494.4494 | EGVFVSN(N)GTHWFVTQR @sulfate 1 Fuc 2 Hex 4 HexNAc 3 | 2.767 | 0.028 | 0.000 | 0.000e+00 | 1078 - 1094 | 1 | - | - | - |
| 4150.8596 | TQSLLIVNN(N)ATNVVIK Fuc 1 Hex 6 HexNAc 5 Neu5Ac 1 | 2.762 | 0.029 | 12.207 | 4.651e+08 | 100 - 116 | 3 | 175.13 | 175.28 | 176.00 |
| 3913.5414 | N(N)FTTAPAIC(C)HDGK @sulfate 1 Fuc 2 Hex 3 HexNAc 8 | 2.758 | 0.030 | -19.039 | 9.101e+06 | 1060 - 1073 | 3 | 82.39 | 82.39 | 82.39 |
| 4039.7128 | VYSSANN(N)C(C)TFEYVSQPFLMDLEGK Fuc 1 Hex 3 HexNAc 3 | 2.738 | 0.030 | 15.338 | 3.693e+08 | 145 - 169 | 2 | 214.24 | 214.40 | 214.69 |
| 4312.9890 | TPPIKDFGGFN(N)FSQILPDPSKPSKR Hex 7 HexNAc 2 | 2.733 | 0.030 | 12.173 | 9.076e+07 | 777 - 802 | 1 | 174.69 | 175.04 | 175.19 |
| 2745.1423 | QC(C)VN(N)LTTR Fuc 2 Hex 4 HexNAc 4 | 2.728 | 0.011 | -4.648 | 1.411e+07 | 0 - 8 | 1 | 115.07 | 115.24 | 115.24 |
| 4150.9375 | TPPIKDFGGFN(N)FSQILPDPSKPSKR Hex 6 HexNAc 2 | 2.727 | 0.031 | 2.931 | 2.609e+07 | 777 - 802 | 2 | 175.36 | 175.44 | 175.64 |
| 2945.1841 | QC(C)VN(N)LTTR Hex 4 HexNAc 5 Neu5Ac 1 | 2.720 | 0.011 | 16.431 | 3.587e+09 | 0 - 8 | 2 | 111.50 | 111.71 | 112.37 |
| 2376.9731 | QC(C)VN(N)LTTR Hex 3 HexNAc 3 Neu5Ac 1 | 2.715 | 0.011 | 15.367 | 6.965e+08 | 0 - 8 | 2 | 112.89 | 113.09 | 113.43 |
| 4100.6375 | N(N)FTTAPAIC(C)HDGK Fuc 2 Hex 3 HexNAc 5 Neu5Ac 3 | 2.711 | 0.031 | 5.760 | 1.691e+07 | 1060 - 1073 | 2 | 97.01 | 97.01 | 97.27 |
| 4049.6848 | YNEN(N)GTITDAVDC(C)ALDPLSETK Hex 5 HexNAc 4 | 2.702 | 0.032 | 0.000 | 0.000e+00 | 265 - 287 | 1 | - | - | - |
| 3887.6317 | YNEN(N)GTITDAVDC(C)ALDPLSETK Hex 4 HexNAc 4 | 2.689 | 0.032 | 0.000 | 0.000e+00 | 265 - 287 | 1 | - | - | - |
| 4454.2187 | YNEN(N)GTITDAVDC(C)ALDPLSETK Hex 5 HexNAc 6 | 2.678 | 0.032 | -4.570 | 2.570e+08 | 265 - 287 | 3 | 184.18 | 187.70 | 187.82 |
| 4236.7731 | YNEN(N)GTITDAVDC(C)ALDPLSETK Fuc 1 Hex 4 HexNAc 5 | 2.661 | 0.032 | 0.000 | 0.000e+00 | 265 - 287 | 2 | - | - | - |
| 3753.5318 | N(N)FTTAPAIC(C)HDGK Fuc 2 Hex 5 HexNAc 6 | 2.655 | 0.032 | 8.672 | 8.614e+07 | 1060 - 1073 | 2 | 128.32 | 128.41 | 128.58 |
| 5143.0632 | YNEN(N)GTITDAVDC(C)ALDPLSETK Fuc 1 Hex 6 HexNAc 5 Neu5Ac 2 | 2.639 | 0.032 | 0.000 | 0.000e+00 | 265 - 287 | 1 | - | - | - |
| 4339.7952 | YNEN(N)GTITDAVDC(C)ALDPLSETK Fuc 2 Hex 5 HexNAc 4 | 2.623 | 0.032 | 13.944 | 3.111e+08 | 265 - 287 | 3 | 187.55 | 187.82 | 188.13 |
| 6613.9362 | QIYKTPPIKDFGGFN(N)FSQILPDPSKPSK Fuc 4 Hex 9 HexNAc 7 | 2.619 | 0.032 | 8.526 | 6.820e+07 | 773 - 801 | 1 | 232.69 | 233.08 | 233.55 |
| 4042.5835 | N(N)FTTAPAIC(C)HDGK Fuc 2 Hex 5 HexNAc 6 Neu5Ac 1 | 2.609 | 0.032 | -3.704 | 8.434e+06 | 1060 - 1073 | 2 | 98.50 | 98.55 | 98.55 |
| 6496.8468 | SSVLHSTQDLFLPFFSNVTWFHAIHVSGTN(N)GTK Fuc 1 Hex 9 HexNAc 6 | 2.600 | 0.032 | 12.816 | 2.188e+09 | 31 - 64 | 2 | 208.43 | 208.87 | 209.43 |
| 4252.7683 | YNEN(N)GTITDAVDC(C)ALDPLSETK Hex 5 HexNAc 5 | 2.594 | 0.032 | 0.000 | 0.000e+00 | 265 - 287 | 1 | - | - | - |
| 7282.1084 | SSVLHSTQDLFLPFFSNVTWFHAIHVSGTN(N)GTK Fuc 1 Hex 9 HexNAc 7 Neu5Ac 2 | 2.577 | 0.032 | 14.936 | 3.288e+09 | 31 - 64 | 2 | 229.71 | 229.91 | 230.78 |
| 3341.4362 | EGVFVSN(N)GTHWFVTQR Hex 4 HexNAc 4 | 2.573 | 0.032 | 0.000 | 0.000e+00 | 1078 - 1094 | 1 | - | - | - |
| 5600.2849 | FPNITNLC(C)PFGEVFN(N)ATR Fuc 2 Hex 8 HexNAc 8 Neu5Ac 1 | 2.505 | 0.033 | 12.523 | 3.776e+09 | 315 - 333 | 4 | 222.93 | 224.77 | 225.75 |
| 6318.7696 | SSVLHSTQDLFLPFFSNVTWFHAIHVSGTN(N)GTK Fuc 2 Hex 7 HexNAc 6 | 2.491 | 0.033 | 10.785 | 2.695e+08 | 31 - 64 | 3 | 209.02 | 209.20 | 210.06 |
| 5355.1480 | FPNITNLC(C)PFGEVFN(N)ATR Hex 9 HexNAc 6 Neu5Ac 2 | 2.490 | 0.033 | 12.902 | 4.659e+08 | 315 - 333 | 1 | 225.34 | 225.61 | 225.94 |
| 6026.7050 | QIYKTPPIKDFGGFN(N)FSQILPDPSKPSK Hex 4 HexNAc 8 Neu5Ac 2 | 2.471 | 0.033 | 0.000 | 0.000e+00 | 773 - 801 | 1 | - | - | - |
| 6601.8063 | SSVLHSTQDLFLPFFSNVTWFHAIHVSGTN(N)GTK @sulfate 1 Fuc 2 Hex 7 HexNAc 7 | 2.465 | 0.033 | 11.532 | 8.286e+08 | 31 - 64 | 2 | 220.86 | 221.16 | 221.53 |
| 3979.7194 | DFGGFN(N)FSQILPDPSKPSK Fuc 1 Hex 4 HexNAc 4 Neu5Ac 1 | 2.442 | 0.034 | 0.000 | 0.000e+00 | 782 - 801 | 1 | - | - | - |
| 4162.7477 | DFGGFN(N)FSQILPDPSKPSK Fuc 1 Hex 5 HexNAc 4 Neu5Ac 1 | 2.432 | 0.034 | 0.000 | 0.000e+00 | 782 - 801 | 1 | - | - | - |
| 5360.4411 | SSVLHSTQDLFLPFFSNVTWFHAIHVSGTN(N)GTK Fuc 1 Hex 7 HexNAc 2 | 2.432 | 0.034 | -5.577 | 2.262e+07 | 31 - 64 | 1 | 223.98 | 223.98 | 224.18 |
| 4344.8493 | DFGGFN(N)FSQILPDPSKPSK Fuc 1 Hex 5 HexNAc 5 Neu5Ac 1 | 2.417 | 0.034 | 0.000 | 0.000e+00 | 782 - 801 | 1 | - | - | - |
| 3914.8373 | TPPIKDFGGFN(N)FSQILPDPSKPSK Hex 3 HexNAc 4 | 2.404 | 0.034 | 9.131 | 2.682e+08 | 777 - 801 | 2 | 187.82 | 188.30 | 188.30 |
| 4343.9725 | TPPIKDFGGFN(N)FSQILPDPSKPSK Fuc 1 Hex 6 HexNAc 3 | 2.382 | 0.035 | 8.424 | 4.437e+07 | 777 - 801 | 2 | 187.11 | 187.22 | 187.36 |
| 4346.8035 | VYSSANN(N)C(C)TFEYVSQPFLMDLEGK Hex 4 HexNAc 3 Neu5Ac 1 | 2.366 | 0.035 | 14.736 | 6.169e+08 | 145 - 169 | 4 | 233.99 | 234.28 | 234.61 |
| 4516.0406 | TPPIKDFGGFN(N)FSQILPDPSKPSK Fuc 1 Hex 4 HexNAc 4 Neu5Ac 1 | 2.355 | 0.035 | 0.000 | 0.000e+00 | 777 - 801 | 1 | - | - | - |
| 4198.9133 | TPPIKDFGGFN(N)FSQILPDPSKPSK Hex 6 HexNAc 3 | 2.345 | 0.035 | 0.000 | 0.000e+00 | 777 - 801 | 1 | - | - | - |
| 2963.2146 | QC(C)VN(N)LTTR Fuc 1 Hex 5 HexNAc 5 | 2.341 | 0.011 | 0.000 | 0.000e+00 | 0 - 8 | 1 | - | - | - |
| 4060.8947 | TPPIKDFGGFN(N)FSQILPDPSKPSK Fuc 1 Hex 3 HexNAc 4 | 2.339 | 0.035 | -5.695 | 1.373e+08 | 777 - 801 | 2 | 185.38 | 185.67 | 185.67 |
| 2718.0620 | QC(C)VN(N)LTTR @sulfate 1 Fuc 1 Hex 3 HexNAc 5 | 2.339 | 0.011 | 0.000 | 0.000e+00 | 0 - 8 | 1 | - | - | - |
| 2842.1874 | QC(C)VN(N)LTTR Fuc 1 Hex 3 HexNAc 6 | 2.332 | 0.011 | 0.000 | 0.000e+00 | 0 - 8 | 1 | - | - | - |
| 4859.0136 | VYSSANN(N)C(C)TFEYVSQPFLMDLEGK Fuc 3 Hex 5 HexNAc 4 | 2.306 | 0.035 | 13.827 | 2.440e+08 | 145 - 169 | 1 | 210.12 | 210.59 | 210.91 |
| 4363.8182 | VYSSANN(N)C(C)TFEYVSQPFLMDLEGK Fuc 1 Hex 5 HexNAc 3 | 2.298 | 0.035 | 11.591 | 1.361e+08 | 145 - 169 | 3 | 212.35 | 212.60 | 212.95 |
| 3125.3465 | QC(C)VN(N)LTTR Fuc 1 Hex 6 HexNAc 5 | 2.294 | 0.019 | 11.252 | 1.245e+09 | 0 - 8 | 4 | 109.12 | 109.28 | 110.50 |
| 2962.2146 | QC(C)VN(N)LTTR Fuc 1 Hex 5 HexNAc 5 | 2.289 | 0.019 | -20.628 | 1.399e+07 | 0 - 8 | 1 | 111.73 | 111.76 | 111.76 |
| 3965.6731 | EGVFVSN(N)GTHWFVTQR Fuc 1 Hex 4 HexNAc 5 Neu5Ac 1 | 2.270 | 0.035 | 0.000 | 0.000e+00 | 1078 - 1094 | 1 | - | - | - |
| 3601.4113 | QC(C)VN(N)LTTR Hex 5 HexNAc 6 Neu5Ac 2 | 2.269 | 0.019 | 13.444 | 3.040e+08 | 0 - 8 | 2 | 131.04 | 131.26 | 131.49 |
| 3591.6381 | TQSLLIVNN(N)ATNVVIK Hex 9 HexNAc 2 | 2.263 | 0.035 | -21.461 | 1.110e+06 | 100 - 116 | 1 | 135.31 | 135.31 | 135.31 |
| 3640.6633 | DFGGFN(N)FSQILPDPSKPSKR Fuc 1 Hex 4 HexNAc 3 | 2.255 | 0.035 | 8.716 | 9.873e+06 | 782 - 802 | 1 | 184.23 | 184.38 | 184.42 |
| 4216.9944 | TPPIKDFGGFN(N)FSQILPDPSKPSKR Fuc 1 Hex 3 HexNAc 4 | 2.254 | 0.035 | 4.398 | 2.952e+07 | 777 - 802 | 1 | 173.42 | 173.60 | 173.84 |
| 3624.7027 | TQSLLIVNN(N)ATNVVIK Fuc 3 Hex 4 HexNAc 4 | 2.247 | 0.035 | 13.233 | 1.202e+08 | 100 - 116 | 3 | 159.64 | 159.77 | 160.01 |
| 2395.0095 | QC(C)VN(N)LTTR Fuc 1 Hex 4 HexNAc 3 | 2.246 | 0.019 | 14.478 | 1.201e+10 | 0 - 8 | 2 | 112.90 | 113.38 | 115.24 |
| 3750.5531 | EGVFVSN(N)GTHWFVTQR Hex 9 HexNAc 2 | 2.243 | 0.035 | 0.000 | 0.000e+00 | 1078 - 1094 | 1 | - | - | - |
| 2597.0878 | QC(C)VN(N)LTTR Fuc 1 Hex 4 HexNAc 4 | 2.243 | 0.019 | -19.678 | 4.786e+06 | 0 - 8 | 1 | 115.21 | 115.21 | 115.21 |
| 3432.3306 | QC(C)VN(N)LTTR @sulfate 1 Fuc 2 Hex 4 HexNAc 7 | 2.237 | 0.025 | 11.505 | 2.235e+07 | 0 - 8 | 2 | 63.20 | 63.34 | 63.65 |
| 3462.6319 | TQSLLIVNN(N)ATNVVIK Fuc 1 Hex 3 HexNAc 4 Neu5Ac 1 | 2.235 | 0.035 | 0.000 | 0.000e+00 | 100 - 116 | 1 | - | - | - |
| 5089.0409 | EGVFVSN(N)GTHWFVTQR Hex 7 HexNAc 6 Neu5Ac 3 | 2.235 | 0.035 | 13.261 | 1.304e+08 | 1078 - 1094 | 2 | 198.39 | 198.62 | 198.84 |
| 2865.1731 | QC(C)VN(N)LTTR Fuc 2 Hex 6 HexNAc 3 | 2.233 | 0.025 | 13.320 | 2.570e+09 | 0 - 8 | 2 | 109.64 | 109.99 | 110.53 |
| 3615.6241 | TQSLLIVNN(N)ATNVVIK @sulfate 1 Fuc 1 Hex 4 HexNAc 5 | 2.224 | 0.036 | 14.552 | 2.154e+08 | 100 - 116 | 4 | 181.69 | 181.94 | 182.18 |
| 3700.7199 | TQSLLIVNN(N)ATNVVIK Fuc 1 Hex 5 HexNAc 5 | 2.219 | 0.036 | 0.000 | 0.000e+00 | 100 - 116 | 1 | - | - | - |
| 3794.6247 | EGVFVSN(N)GTHWFVTQR Fuc 1 Hex 6 HexNAc 4 | 2.216 | 0.036 | 6.344 | 1.936e+08 | 1078 - 1094 | 1 | 223.72 | 223.73 | 223.98 |
| 3607.5183 | EGVFVSN(N)GTHWFVTQR Hex 7 HexNAc 3 | 2.210 | 0.036 | 0.000 | 0.000e+00 | 1078 - 1094 | 1 | - | - | - |
| 2929.1978 | QC(C)VN(N)LTTR Fuc 1 Hex 3 HexNAc 5 Neu5Ac 1 | 2.208 | 0.025 | 6.432 | 5.505e+07 | 0 - 8 | 2 | 133.43 | 133.44 | 133.63 |
| 5062.0666 | VYSSANN(N)C(C)TFEYVSQPFLMDLEGK Fuc 3 Hex 5 HexNAc 5 | 2.205 | 0.036 | -20.470 | 1.996e+07 | 145 - 169 | 1 | 209.95 | 209.95 | 210.10 |
| 3412.5449 | TQSLLIVNN(N)ATNVVIK @sulfate 1 Fuc 1 Hex 4 HexNAc 4 | 2.203 | 0.036 | 11.786 | 5.580e+07 | 100 - 116 | 2 | 183.15 | 183.22 | 183.53 |
| 3276.5513 | TQSLLIVNN(N)ATNVVIK Hex 4 HexNAc 3 Neu5Ac 1 | 2.191 | 0.036 | 0.000 | 0.000e+00 | 100 - 116 | 2 | - | - | - |
| 3736.5784 | EGVFVSN(N)GTHWFVTQR Fuc 2 Hex 6 HexNAc 3 | 2.188 | 0.036 | -4.919 | 3.797e+07 | 1078 - 1094 | 1 | 147.16 | 147.27 | 147.27 |
| 3476.5944 | TQSLLIVNN(N)ATNVVIK Fuc 1 Hex 6 HexNAc 3 | 2.182 | 0.036 | 0.000 | 0.000e+00 | 100 - 116 | 1 | - | - | - |
| 3430.6335 | TQSLLIVNN(N)ATNVVIK Hex 3 HexNAc 6 | 2.181 | 0.036 | 8.624 | 6.447e+07 | 100 - 116 | 3 | 160.12 | 161.08 | 161.08 |
| 4705.9295 | YNEN(N)GTITDAVDC(C)ALDPLSETK Fuc 2 Hex 6 HexNAc 5 | 2.166 | 0.037 | -5.404 | 1.042e+08 | 265 - 287 | 2 | 186.68 | 186.68 | 186.78 |
| 4003.6368 | EGVFVSN(N)GTHWFVTQR @sulfate 1 Fuc 1 Hex 5 HexNAc 4 Neu5Ac 1 | 2.154 | 0.037 | -21.527 | 2.760e+05 | 1078 - 1094 | 1 | 44.64 | 44.64 | 44.64 |
| 6538.8747 | SSVLHSTQDLFLPFFSNVTWFHAIHVSGTN(N)GTK Fuc 1 Hex 8 HexNAc 7 | 2.153 | 0.037 | -4.382 | 6.961e+07 | 31 - 64 | 1 | 208.56 | 208.73 | 208.73 |
| 6787.9385 | SSVLHSTQDLFLPFFSNVTWFHAIHVSGTN(N)GTK Fuc 1 Hex 9 HexNAc 6 Neu5Ac 1 | 2.144 | 0.037 | 12.679 | 1.256e+09 | 31 - 64 | 2 | 218.28 | 219.06 | 219.42 |
| 6686.9413 | SSVLHSTQDLFLPFFSNVTWFHAIHVSGTN(N)GTK Fuc 2 Hex 8 HexNAc 7 | 2.143 | 0.037 | 0.000 | 0.000e+00 | 31 - 64 | 1 | - | - | - |
| 6416.8450 | SSVLHSTQDLFLPFFSNVTWFHAIHVSGTN(N)GTK Fuc 1 Hex 6 HexNAc 8 | 2.136 | 0.038 | 11.967 | 1.239e+09 | 31 - 64 | 3 | 209.20 | 209.61 | 210.56 |
| 5271.1829 | N(N)FTTAPAIC(C)HDGKAHFPR Fuc 1 Hex 9 HexNAc 8 | 2.129 | 0.038 | 6.068 | 6.201e+08 | 1060 - 1078 | 1 | 85.21 | 85.27 | 85.33 |
| 6171.9698 | SNIIRGWIFGTTLDSKTQSLLIVNN(N)ATNVVIK Fuc 4 Hex 4 HexNAc 7 | 2.126 | 0.038 | 0.000 | 0.000e+00 | 84 - 116 | 1 | - | - | - |
| 3682.5426 | YNEN(N)GTITDAVDC(C)ALDPLSETK Hex 4 HexNAc 3 | 2.126 | 0.038 | 0.000 | 0.000e+00 | 265 - 287 | 1 | - | - | - |
| 3346.3879 | EGVFVSN(N)GTHWFVTQR @sulfate 1 Fuc 1 Hex 4 HexNAc 3 | 2.123 | 0.038 | -6.123 | 1.932e+07 | 1078 - 1094 | 2 | 148.86 | 149.03 | 149.10 |
| 6724.9558 | SSVLHSTQDLFLPFFSNVTWFHAIHVSGTN(N)GTK Fuc 2 Hex 7 HexNAc 8 | 2.118 | 0.039 | 11.063 | 7.238e+08 | 31 - 64 | 1 | 208.22 | 208.68 | 209.20 |
| 4740.2585 | GWIFGTTLDSKTQSLLIVNN(N)ATNVVIK Hex 6 HexNAc 4 | 2.095 | 0.040 | 0.000 | 0.000e+00 | 89 - 116 | 1 | - | - | - |
| 6779.8848 | SSVLHSTQDLFLPFFSNVTWFHAIHVSGTN(N)GTK @sulfate 1 Fuc 1 Hex 9 HexNAc 7 | 2.095 | 0.040 | 12.178 | 1.762e+09 | 31 - 64 | 1 | 220.24 | 220.49 | 220.80 |
| 2045.8721 | QC(C)VN(N)LTTR Hex 4 HexNAc 2 | 2.093 | 0.025 | 10.898 | 7.061e+07 | 0 - 8 | 1 | 115.06 | 115.27 | 115.30 |
| 6073.9315 | SNIIRGWIFGTTLDSKTQSLLIVNN(N)ATNVVIK Fuc 1 Hex 6 HexNAc 7 | 2.090 | 0.040 | 0.000 | 0.000e+00 | 84 - 116 | 1 | - | - | - |
| 4075.6251 | N(N)FTTAPAIC(C)HDGK Fuc 2 Hex 7 HexNAc 6 | 2.089 | 0.040 | 11.410 | 9.476e+07 | 1060 - 1073 | 4 | 82.54 | 82.68 | 82.90 |
| 3998.6581 | VYSSANN(N)C(C)TFEYVSQPFLMDLEGK Fuc 1 Hex 4 HexNAc 2 | 2.088 | 0.040 | -5.198 | 6.193e+07 | 145 - 169 | 2 | 214.20 | 214.38 | 214.38 |
| 5967.8921 | SNIIRGWIFGTTLDSKTQSLLIVNN(N)ATNVVIK Hex 5 HexNAc 8 | 2.083 | 0.040 | 0.000 | 0.000e+00 | 84 - 116 | 1 | - | - | - |
| 3547.4185 | N(N)FTTAPAIC(C)HDGK Hex 5 HexNAc 5 Neu5Ac 1 | 2.083 | 0.040 | 12.394 | 4.657e+07 | 1060 - 1073 | 1 | 97.19 | 97.31 | 97.45 |
| 3330.3765 | EGVFVSN(N)GTHWFVTQR @sulfate 1 Fuc 2 Hex 3 HexNAc 3 | 2.077 | 0.040 | 12.463 | 4.722e+06 | 1078 - 1094 | 1 | 48.55 | 48.62 | 48.82 |
| 3914.5781 | N(N)FTTAPAIC(C)HDGK Fuc 2 Hex 6 HexNAc 6 | 2.073 | 0.040 | -6.511 | 4.145e+06 | 1060 - 1073 | 1 | 82.81 | 82.81 | 82.99 |
| 6055.9079 | SNIIRGWIFGTTLDSKTQSLLIVNN(N)ATNVVIK Fuc 1 Hex 6 HexNAc 7 | 2.066 | 0.040 | -5.778 | 1.830e+08 | 84 - 116 | 1 | 161.49 | 161.62 | 161.62 |
| 4380.8196 | YNEN(N)GTITDAVDC(C)ALDPLSETK Fuc 2 Hex 4 HexNAc 5 | 2.065 | 0.040 | 8.309 | 1.268e+08 | 265 - 287 | 3 | 187.67 | 188.24 | 188.24 |
| 6227.9871 | SNIIRGWIFGTTLDSKTQSLLIVNN(N)ATNVVIK Fuc 3 Hex 4 HexNAc 8 | 2.063 | 0.040 | -22.935 | 1.754e+07 | 84 - 116 | 1 | 161.03 | 161.03 | 161.03 |
| 3619.3609 | N(N)FTTAPAIC(C)HDGK @sulfate 1 Hex 8 HexNAc 4 | 2.057 | 0.041 | 0.000 | 0.000e+00 | 1060 - 1073 | 1 | - | - | - |
| 3945.5256 | N(N)FTTAPAIC(C)HDGK @sulfate 1 Hex 5 HexNAc 8 | 2.056 | 0.041 | -21.217 | 1.517e+06 | 1060 - 1073 | 1 | 82.64 | 82.64 | 82.64 |
| 6903.9648 | SSVLHSTQDLFLPFFSNVTWFHAIHVSGTN(N)GTK Fuc 1 Hex 9 HexNAc 8 | 2.055 | 0.041 | -7.327 | 3.844e+07 | 31 - 64 | 2 | 220.56 | 220.92 | 220.92 |
| 4275.7904 | YNEN(N)GTITDAVDC(C)ALDPLSETK Fuc 1 Hex 3 HexNAc 6 | 2.054 | 0.041 | 12.239 | 4.364e+08 | 265 - 287 | 3 | 189.34 | 189.44 | 189.49 |
| 3490.3579 | N(N)FTTAPAIC(C)HDGK Hex 7 HexNAc 3 Neu5Ac 1 | 2.046 | 0.041 | 0.000 | 0.000e+00 | 1060 - 1073 | 1 | - | - | - |
| 3386.3843 | N(N)FTTAPAIC(C)HDGK Fuc 2 Hex 4 HexNAc 5 | 2.046 | 0.041 | 10.408 | 2.690e+07 | 1060 - 1073 | 2 | 84.34 | 84.50 | 84.62 |
| 7054.9525 | SSVLHSTQDLFLPFFSNVTWFHAIHVSGTN(N)GTK @sulfate 1 Fuc 2 Hex 8 HexNAc 7 Neu5Ac 1 | 2.032 | 0.041 | -20.458 | 2.649e+07 | 31 - 64 | 1 | 232.45 | 232.49 | 232.49 |
| 4191.9645 | TPPIKDFGGFN(N)FSQILPDPSKPSKR Hex 5 HexNAc 3 | 2.015 | 0.042 | 8.574 | 5.640e+07 | 777 - 802 | 1 | 175.13 | 175.44 | 175.75 |
| 3159.4242 | DFGGFN(N)FSQILPDPSKPSK Hex 4 HexNAc 2 | 2.007 | 0.043 | 0.000 | 0.000e+00 | 782 - 801 | 1 | - | - | - |
| 2852.1687 | N(N)FTTAPAIC(C)HDGK Hex 5 HexNAc 3 | 2.005 | 0.043 | 0.000 | 0.000e+00 | 1060 - 1073 | 1 | - | - | - |
| 6613.9297 | QIYKTPPIKDFGGFN(N)FSQILPDPSKPSK Fuc 2 Hex 9 HexNAc 7 Neu5Ac 1 | 1.996 | 0.044 | 6.513 | 5.375e+08 | 773 - 801 | 4 | 221.82 | 222.11 | 223.21 |
| 3658.4691 | NN(N)KSWMESEFR Hex 5 HexNAc 7 | 1.989 | 0.045 | 11.128 | 5.147e+08 | 134 - 145 | 1 | 162.51 | 162.74 | 163.05 |
| 6610.9637 | GYHLMSFPQSAPHGVVFLHVTYVPAQEKN(N)FTTAPAIC(C)HDGK Fuc 1 Hex 5 HexNAc 4 Neu5Ac 1 | 1.988 | 0.045 | 9.900 | 1.618e+08 | 1032 - 1073 | 2 | 167.20 | 167.47 | 167.90 |
| 3807.6722 | NLN(N)ESLIDLQELGK Fuc 3 Hex 6 HexNAc 4 | 1.987 | 0.045 | 17.925 | 2.698e+09 | 1178 - 1192 | 2 | 244.95 | 245.24 | 246.01 |
| 3143.3154 | EGVFVSN(N)GTHWFVTQR @sulfate 1 Fuc 1 Hex 4 HexNAc 2 | 1.975 | 0.046 | 9.090 | 3.539e+07 | 1078 - 1094 | 1 | 101.17 | 101.24 | 101.85 |
| 3994.6049 | EGVFVSN(N)GTHWFVTQR @sulfate 1 Fuc 1 Hex 8 HexNAc 3 | 1.955 | 0.047 | 13.740 | 3.219e+09 | 1078 - 1094 | 7 | 166.22 | 166.50 | 167.94 |
| 3648.5030 | EGVFVSN(N)GTHWFVTQR @sulfate 1 Hex 3 HexNAc 6 | 1.955 | 0.047 | 9.145 | 2.242e+07 | 1078 - 1094 | 1 | 246.50 | 246.69 | 246.94 |
| 3215.2959 | N(N)FTTAPAIC(C)HDGK Hex 6 HexNAc 4 | 1.949 | 0.048 | 10.886 | 5.237e+07 | 1060 - 1073 | 2 | 83.94 | 84.62 | 84.62 |
| 3381.4619 | EGVFVSN(N)GTHWFVTQR Hex 3 HexNAc 5 | 1.935 | 0.048 | 0.000 | 0.000e+00 | 1078 - 1094 | 1 | - | - | - |
| 3228.3729 | EGVFVSN(N)GTHWFVTQR Fuc 1 Hex 5 HexNAc 2 | 1.925 | 0.049 | 0.000 | 0.000e+00 | 1078 - 1094 | 1 | - | - | - |
| 3184.3370 | EGVFVSN(N)GTHWFVTQR @sulfate 1 Fuc 1 Hex 3 HexNAc 3 | 1.923 | 0.049 | -5.742 | 8.403e+06 | 1078 - 1094 | 1 | 150.00 | 150.00 | 150.18 |
| 4426.0254 | TPPIKDFGGFN(N)FSQILPDPSKPSK Fuc 1 Hex 4 HexNAc 5 | 1.921 | 0.049 | 10.395 | 2.051e+08 | 777 - 801 | 3 | 184.50 | 184.93 | 185.42 |
| 4530.0776 | TPPIKDFGGFN(N)FSQILPDPSKPSK Hex 5 HexNAc 4 Neu5Ac 1 | 1.912 | 0.049 | 15.468 | -9.361e+08 | 777 - 801 | 1 | 199.46 | 199.80 | 200.12 |
| 5182.2873 | TPPIKDFGGFN(N)FSQILPDPSKPSK Fuc 1 Hex 6 HexNAc 7 | 1.911 | 0.049 | 0.000 | 0.000e+00 | 777 - 801 | 1 | - | - | - |
| 3800.6606 | NLN(N)ESLIDLQELGK Fuc 1 Hex 4 HexNAc 7 | 1.909 | 0.049 | 9.106 | 6.522e+07 | 1178 - 1192 | 1 | 173.65 | 173.73 | 174.09 |
| 3232.4501 | NLN(N)ESLIDLQELGK Fuc 1 Hex 3 HexNAc 5 | 1.900 | 0.049 | 13.067 | 1.268e+08 | 1178 - 1192 | 2 | 176.28 | 176.43 | 176.69 |
| 4320.9475 | TPPIKDFGGFN(N)FSQILPDPSKPSK Hex 8 HexNAc 2 | 1.899 | 0.049 | 0.000 | 0.000e+00 | 777 - 801 | 1 | - | - | - |

#### EGVFVSN(N)GTHWFVTQR Hex 6 HexNAc 3 Neu5Ac 1

**Mass:**  3735.5566

(2.760e-07)

**Charge States:** 3, 4

**Start Position**: 1079

**End Position**: 1095

| Profile Score | Tandem Score | TDA q-value | Volume | Spectrum Matches |
| --- | --- | --- | --- | --- |
| 17.4405 | 32.8965 | 0.0010 | 1.1347e+10 | 7 |

**Adduct Signal Fractions**

- **Unmodified**: 1.1347e+10

#### EGVFVSN(N)GTHWFVTQR Hex 4 HexNAc 5 Neu5Ac 1

**Mass:**  3817.6099

(8.810e-08)

**Charge States:** 3, 4

**Start Position**: 1079

**End Position**: 1095

| Profile Score | Tandem Score | TDA q-value | Volume | Spectrum Matches |
| --- | --- | --- | --- | --- |
| 17.2772 | 30.1258 | 0.0010 | 6.2777e+09 | 12 |

**Adduct Signal Fractions**

- **Unmodified**: 6.2777e+09

#### FPNITNLC(C)PFGEVFN(N)ATR Fuc 2 Hex 6 HexNAc 8

**Mass:**  4985.0800

(-7.784e-07)

**Charge States:** 4

**Start Position**: 316

**End Position**: 334

| Profile Score | Tandem Score | TDA q-value | Volume | Spectrum Matches |
| --- | --- | --- | --- | --- |
| 18.0427 | 29.2392 | 0.0010 | 9.3279e+09 | 5 |

**Adduct Signal Fractions**

- **Unmodified**: 9.3279e+09

#### EGVFVSN(N)GTHWFVTQR Hex 5 HexNAc 3 Neu5Ac 1

**Mass:**  3573.5051

(6.968e-07)

**Charge States:** 3, 4

**Start Position**: 1079

**End Position**: 1095

| Profile Score | Tandem Score | TDA q-value | Volume | Spectrum Matches |
| --- | --- | --- | --- | --- |
| 17.5619 | 28.5222 | 0.0010 | 1.0034e+10 | 13 |

**Adduct Signal Fractions**

- **Unmodified**: 1.0034e+10

#### EGVFVSN(N)GTHWFVTQR Hex 6 HexNAc 3

**Mass:**  3444.4597

(-3.417e-07)

**Charge States:** 3, 4

**Start Position**: 1079

**End Position**: 1095

| Profile Score | Tandem Score | TDA q-value | Volume | Spectrum Matches |
| --- | --- | --- | --- | --- |
| 17.6571 | 26.5286 | 0.0010 | 1.4351e+10 | 8 |

**Adduct Signal Fractions**

- **Unmodified**: 1.4351e+10

#### EGVFVSN(N)GTHWFVTQR Fuc 1 Hex 4 HexNAc 5 Neu5Ac 1

**Mass:**  3963.6659

(1.222e-07)

**Charge States:** 3, 4

**Start Position**: 1079

**End Position**: 1095

| Profile Score | Tandem Score | TDA q-value | Volume | Spectrum Matches |
| --- | --- | --- | --- | --- |
| 15.0135 | 25.8235 | 0.0010 | 1.0462e+10 | 9 |

**Adduct Signal Fractions**

- **Unmodified**: 1.0462e+10

#### EGVFVSN(N)GTHWFVTQR Hex 5 HexNAc 3

**Mass:**  3282.4075

(-8.400e-08)

**Charge States:** 3, 4

**Start Position**: 1079

**End Position**: 1095

| Profile Score | Tandem Score | TDA q-value | Volume | Spectrum Matches |
| --- | --- | --- | --- | --- |
| 16.8526 | 25.7156 | 0.0010 | 5.2438e+10 | 14 |

**Adduct Signal Fractions**

- **NH3**: 1.4316e+07
- **Unmodified**: 5.2424e+10

#### EGVFVSN(N)GTHWFVTQR Fuc 1 Hex 5 HexNAc 3 Neu5Ac 1

**Mass:**  3719.5616

(-7.243e-09)

**Charge States:** 3, 4

**Start Position**: 1079

**End Position**: 1095

| Profile Score | Tandem Score | TDA q-value | Volume | Spectrum Matches |
| --- | --- | --- | --- | --- |
| 16.7095 | 25.5514 | 0.0010 | 1.0419e+10 | 13 |

**Adduct Signal Fractions**

- **Unmodified**: 1.0419e+10

#### N(N)FTTAPAIC(C)HDGK Fuc 1 Hex 5 HexNAc 5

**Mass:**  3402.3797

(1.448e-07)

**Charge States:** 3, 4

**Start Position**: 1061

**End Position**: 1074

| Profile Score | Tandem Score | TDA q-value | Volume | Spectrum Matches |
| --- | --- | --- | --- | --- |
| 12.4089 | 25.1288 | 0.0010 | 1.7969e+10 | 22 |

**Adduct Signal Fractions**

- **Unmodified**: 1.7969e+10

#### N(N)FTTAPAIC(C)HDGK Hex 8 HexNAc 2

**Mass:**  3133.2432

(-1.774e-07)

**Charge States:** 2, 3, 4

**Start Position**: 1061

**End Position**: 1074

| Profile Score | Tandem Score | TDA q-value | Volume | Spectrum Matches |
| --- | --- | --- | --- | --- |
| 16.2486 | 24.8476 | 0.0010 | 1.5275e+09 | 7 |

**Adduct Signal Fractions**

- **Unmodified**: 1.5275e+09

#### EGVFVSN(N)GTHWFVTQR Fuc 1 Hex 6 HexNAc 5 Neu5Ac 1

**Mass:**  4287.7741

(-8.017e-08)

**Charge States:** 3, 4

**Start Position**: 1079

**End Position**: 1095

| Profile Score | Tandem Score | TDA q-value | Volume | Spectrum Matches |
| --- | --- | --- | --- | --- |
| 13.4736 | 24.6009 | 0.0010 | 8.8243e+09 | 13 |

**Adduct Signal Fractions**

- **Unmodified**: 8.8243e+09

#### N(N)FTTAPAIC(C)HDGK Fuc 1 Hex 6 HexNAc 5

**Mass:**  3564.4330

(7.713e-08)

**Charge States:** 3, 4, 5

**Start Position**: 1061

**End Position**: 1074

| Profile Score | Tandem Score | TDA q-value | Volume | Spectrum Matches |
| --- | --- | --- | --- | --- |
| 17.4832 | 24.5235 | 0.0010 | 1.2765e+10 | 18 |

**Adduct Signal Fractions**

- **Unmodified**: 1.2765e+10

#### EGVFVSN(N)GTHWFVTQR Hex 4 HexNAc 4 Neu5Ac 1

**Mass:**  3614.5316

(5.595e-07)

**Charge States:** 3, 4

**Start Position**: 1079

**End Position**: 1095

| Profile Score | Tandem Score | TDA q-value | Volume | Spectrum Matches |
| --- | --- | --- | --- | --- |
| 17.7620 | 24.4054 | 0.0010 | 1.9770e+10 | 17 |

**Adduct Signal Fractions**

- **Unmodified**: 1.9770e+10

#### N(N)FTTAPAIC(C)HDGK Fuc 1 Hex 3 HexNAc 4

**Mass:**  2875.1958

(7.427e-07)

**Charge States:** 2, 3, 4

**Start Position**: 1061

**End Position**: 1074

| Profile Score | Tandem Score | TDA q-value | Volume | Spectrum Matches |
| --- | --- | --- | --- | --- |
| 18.1652 | 24.2656 | 0.0010 | 1.1468e+11 | 34 |

**Adduct Signal Fractions**

- **NH3**: 1.0282e+08
- **Unmodified**: 1.1458e+11

#### EGVFVSN(N)GTHWFVTQR Fuc 1 Hex 5 HexNAc 3

**Mass:**  3428.4670

(-3.858e-07)

**Charge States:** 3, 4

**Start Position**: 1079

**End Position**: 1095

| Profile Score | Tandem Score | TDA q-value | Volume | Spectrum Matches |
| --- | --- | --- | --- | --- |
| 16.7149 | 24.1767 | 0.0010 | 1.0131e+10 | 8 |

**Adduct Signal Fractions**

- **Unmodified**: 1.0131e+10

#### N(N)FTTAPAIC(C)HDGK Fuc 1 Hex 4 HexNAc 4 Neu5Ac 1

**Mass:**  3328.3438

(-2.214e-07)

**Charge States:** 3, 4

**Start Position**: 1061

**End Position**: 1074

| Profile Score | Tandem Score | TDA q-value | Volume | Spectrum Matches |
| --- | --- | --- | --- | --- |
| 18.0557 | 23.7630 | 0.0010 | 1.0396e+10 | 14 |

**Adduct Signal Fractions**

- **Na**: 9.1771e+06
- **Unmodified**: 1.0387e+10

#### EGVFVSN(N)GTHWFVTQR Hex 5 HexNAc 4 Neu5Ac 1

**Mass:**  3776.5825

(-3.331e-07)

**Charge States:** 3, 4

**Start Position**: 1079

**End Position**: 1095

| Profile Score | Tandem Score | TDA q-value | Volume | Spectrum Matches |
| --- | --- | --- | --- | --- |
| 16.3426 | 23.3488 | 0.0010 | 2.4590e+10 | 24 |

**Adduct Signal Fractions**

- **NH3**: 2.6018e+07
- **Unmodified**: 2.4564e+10

#### N(N)FTTAPAIC(C)HDGK Fuc 1 Hex 5 HexNAc 4 Neu5Ac 1

**Mass:**  3490.3967

(-3.315e-08)

**Charge States:** 3, 4, 5

**Start Position**: 1061

**End Position**: 1074

| Profile Score | Tandem Score | TDA q-value | Volume | Spectrum Matches |
| --- | --- | --- | --- | --- |
| 17.3130 | 23.2117 | 0.0010 | 1.3975e+10 | 18 |

**Adduct Signal Fractions**

- **Unmodified**: 1.3975e+10

#### N(N)FTTAPAIC(C)HDGK Fuc 1 Hex 7 HexNAc 6 Neu5Ac 1

**Mass:**  4220.6612

(1.189e-07)

**Charge States:** 3, 4

**Start Position**: 1061

**End Position**: 1074

| Profile Score | Tandem Score | TDA q-value | Volume | Spectrum Matches |
| --- | --- | --- | --- | --- |
| 14.5349 | 23.1849 | 0.0010 | 2.2362e+09 | 7 |

**Adduct Signal Fractions**

- **Unmodified**: 2.2362e+09

#### N(N)FTTAPAIC(C)HDGK Fuc 1 Hex 6 HexNAc 5 Neu5Ac 1

**Mass:**  3855.5255

(7.817e-08)

**Charge States:** 3, 4, 5

**Start Position**: 1061

**End Position**: 1074

| Profile Score | Tandem Score | TDA q-value | Volume | Spectrum Matches |
| --- | --- | --- | --- | --- |
| 13.5515 | 22.7792 | 0.0010 | 7.6024e+09 | 15 |

**Adduct Signal Fractions**

- **Unmodified**: 7.6024e+09

#### N(N)FTTAPAIC(C)HDGK Fuc 1 Hex 7 HexNAc 6

**Mass:**  3929.5671

(4.314e-07)

**Charge States:** 3, 4, 5

**Start Position**: 1061

**End Position**: 1074

| Profile Score | Tandem Score | TDA q-value | Volume | Spectrum Matches |
| --- | --- | --- | --- | --- |
| 17.9002 | 22.6342 | 0.0010 | 3.0580e+09 | 7 |

**Adduct Signal Fractions**

- **Unmodified**: 3.0580e+09

#### N(N)FTTAPAIC(C)HDGK Hex 6 HexNAc 3 Neu5Ac 1

**Mass:**  3303.3121

(2.099e-07)

**Charge States:** 3, 4

**Start Position**: 1061

**End Position**: 1074

| Profile Score | Tandem Score | TDA q-value | Volume | Spectrum Matches |
| --- | --- | --- | --- | --- |
| 18.3484 | 22.5283 | 0.0010 | 4.0921e+09 | 12 |

**Adduct Signal Fractions**

- **Unmodified**: 4.0921e+09

#### N(N)FTTAPAIC(C)HDGK Fuc 1 Hex 5 HexNAc 3 Neu5Ac 1

**Mass:**  3287.3155

(9.345e-08)

**Charge States:** 3, 4

**Start Position**: 1061

**End Position**: 1074

| Profile Score | Tandem Score | TDA q-value | Volume | Spectrum Matches |
| --- | --- | --- | --- | --- |
| 18.8406 | 21.9823 | 0.0010 | 6.4428e+09 | 15 |

**Adduct Signal Fractions**

- **Unmodified**: 6.4428e+09

#### EGVFVSN(N)GTHWFVTQR Hex 3 HexNAc 4

**Mass:**  3161.3829

(4.921e-07)

**Charge States:** 2, 3, 4

**Start Position**: 1079

**End Position**: 1095

| Profile Score | Tandem Score | TDA q-value | Volume | Spectrum Matches |
| --- | --- | --- | --- | --- |
| 18.3105 | 21.8943 | 0.0010 | 2.3595e+10 | 9 |

**Adduct Signal Fractions**

- **Unmodified**: 2.3595e+10

#### N(N)FTTAPAIC(C)HDGK Fuc 1 Hex 4 HexNAc 5 Neu5Ac 1

**Mass:**  3531.4224

(1.666e-07)

**Charge States:** 3, 4

**Start Position**: 1061

**End Position**: 1074

| Profile Score | Tandem Score | TDA q-value | Volume | Spectrum Matches |
| --- | --- | --- | --- | --- |
| 14.6107 | 21.5453 | 0.0010 | 3.8079e+09 | 9 |

**Adduct Signal Fractions**

- **Unmodified**: 3.8079e+09

#### N(N)FTTAPAIC(C)HDGK Fuc 2 Hex 4 HexNAc 3

**Mass:**  2980.2261

(5.703e-03)

**Charge States:** 3, 4

**Start Position**: 1061

**End Position**: 1074

| Profile Score | Tandem Score | TDA q-value | Volume | Spectrum Matches |
| --- | --- | --- | --- | --- |
| -6.7833 | 21.3199 | 0.0010 | 2.8161e+08 | 15 |

**Adduct Signal Fractions**

- **Unmodified**: 6.4500e+07
- **NH3**: 2.1711e+08

#### N(N)FTTAPAIC(C)HDGK Fuc 1 Hex 5 HexNAc 5 Neu5Ac 1

**Mass:**  3693.4738

(2.857e-07)

**Charge States:** 3, 4

**Start Position**: 1061

**End Position**: 1074

| Profile Score | Tandem Score | TDA q-value | Volume | Spectrum Matches |
| --- | --- | --- | --- | --- |
| 13.7733 | 20.6964 | 0.0010 | 6.1161e+09 | 14 |

**Adduct Signal Fractions**

- **Unmodified**: 6.1161e+09

#### N(N)FTTAPAIC(C)HDGK Fuc 1 Hex 4 HexNAc 5

**Mass:**  3240.3275

(2.587e-07)

**Charge States:** 3, 4

**Start Position**: 1061

**End Position**: 1074

| Profile Score | Tandem Score | TDA q-value | Volume | Spectrum Matches |
| --- | --- | --- | --- | --- |
| 11.8695 | 20.6810 | 0.0010 | 4.8638e+10 | 29 |

**Adduct Signal Fractions**

- **Na**: 2.2068e+06
- **Unmodified**: 4.8636e+10

#### EGVFVSN(N)GTHWFVTQR Hex 6 HexNAc 5 Neu5Ac 1

**Mass:**  4141.7139

(5.669e-07)

**Charge States:** 3, 4

**Start Position**: 1079

**End Position**: 1095

| Profile Score | Tandem Score | TDA q-value | Volume | Spectrum Matches |
| --- | --- | --- | --- | --- |
| 14.0949 | 20.6661 | 0.0010 | -5.9404e+07 | 8 |

**Adduct Signal Fractions**

- **Unmodified**: 8.5305e+09

#### N(N)FTTAPAIC(C)HDGK Fuc 1 Hex 4 HexNAc 4

**Mass:**  3037.2481

(2.341e-07)

**Charge States:** 2, 3, 4

**Start Position**: 1061

**End Position**: 1074

| Profile Score | Tandem Score | TDA q-value | Volume | Spectrum Matches |
| --- | --- | --- | --- | --- |
| 15.7354 | 20.3466 | 0.0010 | 7.4253e+10 | 34 |

**Adduct Signal Fractions**

- **NH3**: 6.1982e+08
- **Unmodified**: 7.3634e+10

#### EGVFVSN(N)GTHWFVTQR Fuc 1 Hex 4 HexNAc 4 Neu5Ac 1

**Mass:**  3760.5899

(-3.356e-07)

**Charge States:** 3, 4

**Start Position**: 1079

**End Position**: 1095

| Profile Score | Tandem Score | TDA q-value | Volume | Spectrum Matches |
| --- | --- | --- | --- | --- |
| 16.0066 | 20.3154 | 0.0010 | 2.2886e+10 | 19 |

**Adduct Signal Fractions**

- **NH3**: 4.6127e+06
- **Unmodified**: 2.2881e+10

#### N(N)FTTAPAIC(C)HDGK Fuc 1 Hex 4 HexNAc 3

**Mass:**  2834.1690

(2.346e-07)

**Charge States:** 2, 3, 4

**Start Position**: 1061

**End Position**: 1074

| Profile Score | Tandem Score | TDA q-value | Volume | Spectrum Matches |
| --- | --- | --- | --- | --- |
| 13.4867 | 19.7732 | 0.0010 | 3.1575e+10 | 29 |

**Adduct Signal Fractions**

- **Unmodified**: 3.1575e+10

#### N(N)FTTAPAIC(C)HDGK Fuc 1 Hex 3 HexNAc 5

**Mass:**  3078.2757

(-1.629e-08)

**Charge States:** 2, 3, 4

**Start Position**: 1061

**End Position**: 1074

| Profile Score | Tandem Score | TDA q-value | Volume | Spectrum Matches |
| --- | --- | --- | --- | --- |
| 17.2157 | 19.7183 | 0.0010 | 1.6217e+11 | 49 |

**Adduct Signal Fractions**

- **NH3**: 8.2945e+06
- **Unmodified**: 1.6216e+11

#### EGVFVSN(N)GTHWFVTQR Hex 3 HexNAc 6

**Mass:**  3567.5387

(-3.650e-07)

**Charge States:** 3, 4

**Start Position**: 1079

**End Position**: 1095

| Profile Score | Tandem Score | TDA q-value | Volume | Spectrum Matches |
| --- | --- | --- | --- | --- |
| 17.0007 | 19.6346 | 0.0010 | 5.9652e+09 | 8 |

**Adduct Signal Fractions**

- **Unmodified**: 5.9652e+09

#### N(N)FTTAPAIC(C)HDGK Hex 5 HexNAc 3 Neu5Ac 1

**Mass:**  3141.2584

(6.964e-07)

**Charge States:** 2, 3, 4

**Start Position**: 1061

**End Position**: 1074

| Profile Score | Tandem Score | TDA q-value | Volume | Spectrum Matches |
| --- | --- | --- | --- | --- |
| 17.9498 | 19.6121 | 0.0010 | 2.4989e+09 | 10 |

**Adduct Signal Fractions**

- **Unmodified**: 2.4989e+09

#### EGVFVSN(N)GTHWFVTQR Hex 4 HexNAc 3 Neu5Ac 1

**Mass:**  3411.4520

(-1.206e-07)

**Charge States:** 3, 4

**Start Position**: 1079

**End Position**: 1095

| Profile Score | Tandem Score | TDA q-value | Volume | Spectrum Matches |
| --- | --- | --- | --- | --- |
| 17.3792 | 19.4215 | 0.0010 | 4.1746e+09 | 7 |

**Adduct Signal Fractions**

- **Unmodified**: 4.1746e+09

#### EGVFVSN(N)GTHWFVTQR Fuc 1 Hex 7 HexNAc 6 Neu5Ac 1

**Mass:**  4652.9009

(-2.297e-07)

**Charge States:** 3, 4

**Start Position**: 1079

**End Position**: 1095

| Profile Score | Tandem Score | TDA q-value | Volume | Spectrum Matches |
| --- | --- | --- | --- | --- |
| 13.4478 | 19.0179 | 0.0010 | 6.6314e+09 | 1 |

**Adduct Signal Fractions**

- **Unmodified**: 6.6314e+09

#### N(N)FTTAPAIC(C)HDGK Hex 7 HexNAc 2

**Mass:**  2971.1900

(3.174e-07)

**Charge States:** 2, 3

**Start Position**: 1061

**End Position**: 1074

| Profile Score | Tandem Score | TDA q-value | Volume | Spectrum Matches |
| --- | --- | --- | --- | --- |
| 15.3269 | 18.9250 | 0.0010 | 2.3108e+09 | 4 |

**Adduct Signal Fractions**

- **Unmodified**: 2.3108e+09

#### EGVFVSN(N)GTHWFVTQR Hex 5 HexNAc 5 Neu5Ac 1

**Mass:**  3979.6615

(1.191e-07)

**Charge States:** 3, 4

**Start Position**: 1079

**End Position**: 1095

| Profile Score | Tandem Score | TDA q-value | Volume | Spectrum Matches |
| --- | --- | --- | --- | --- |
| 13.2630 | 18.9248 | 0.0010 | 9.0196e+09 | 10 |

**Adduct Signal Fractions**

- **Unmodified**: 9.0196e+09

#### EGVFVSN(N)GTHWFVTQR Hex 5 HexNAc 2

**Mass:**  3079.3313

(5.303e-07)

**Charge States:** 2, 3, 4

**Start Position**: 1079

**End Position**: 1095

| Profile Score | Tandem Score | TDA q-value | Volume | Spectrum Matches |
| --- | --- | --- | --- | --- |
| 17.2907 | 18.7697 | 0.0010 | 1.1364e+10 | 15 |

**Adduct Signal Fractions**

- **Unmodified**: 1.1364e+10

#### N(N)FTTAPAIC(C)HDGK Fuc 1 Hex 6 HexNAc 6

**Mass:**  3767.5136

(2.486e-07)

**Charge States:** 3, 4

**Start Position**: 1061

**End Position**: 1074

| Profile Score | Tandem Score | TDA q-value | Volume | Spectrum Matches |
| --- | --- | --- | --- | --- |
| 12.1465 | 18.7266 | 0.0010 | 3.2935e+09 | 14 |

**Adduct Signal Fractions**

- **Unmodified**: 3.2935e+09

#### N(N)FTTAPAIC(C)HDGK Fuc 1 Hex 5 HexNAc 6

**Mass:**  3605.4590

(7.639e-07)

**Charge States:** 3, 4

**Start Position**: 1061

**End Position**: 1074

| Profile Score | Tandem Score | TDA q-value | Volume | Spectrum Matches |
| --- | --- | --- | --- | --- |
| 9.0232 | 18.6343 | 0.0010 | 4.9574e+09 | 23 |

**Adduct Signal Fractions**

- **Unmodified**: 4.9574e+09

#### FPNITNLC(C)PFGEVFN(N)ATR Fuc 1 Hex 8 HexNAc 6

**Mass:**  4756.9758

(6.262e-07)

**Charge States:** 3, 4

**Start Position**: 316

**End Position**: 334

| Profile Score | Tandem Score | TDA q-value | Volume | Spectrum Matches |
| --- | --- | --- | --- | --- |
| 14.3274 | 18.4500 | 0.0010 | 3.2059e+09 | 4 |

**Adduct Signal Fractions**

- **Unmodified**: 3.2059e+09

#### N(N)FTTAPAIC(C)HDGK Hex 6 HexNAc 3

**Mass:**  3012.2163

(1.580e-07)

**Charge States:** 2, 3, 4

**Start Position**: 1061

**End Position**: 1074

| Profile Score | Tandem Score | TDA q-value | Volume | Spectrum Matches |
| --- | --- | --- | --- | --- |
| 14.5362 | 18.3373 | 0.0010 | 9.0733e+09 | 10 |

**Adduct Signal Fractions**

- **Unmodified**: 9.0733e+09

#### N(N)FTTAPAIC(C)HDGK Fuc 1 Hex 4 HexNAc 6

**Mass:**  3443.4065

(3.843e-07)

**Charge States:** 3, 4

**Start Position**: 1061

**End Position**: 1074

| Profile Score | Tandem Score | TDA q-value | Volume | Spectrum Matches |
| --- | --- | --- | --- | --- |
| 8.0483 | 18.2397 | 0.0010 | 5.0506e+09 | 20 |

**Adduct Signal Fractions**

- **Unmodified**: 5.0506e+09

#### N(N)FTTAPAIC(C)HDGK Fuc 1 Hex 5 HexNAc 4 Neu5Ac 2

**Mass:**  3781.4918

(1.102e-07)

**Charge States:** 3, 4

**Start Position**: 1061

**End Position**: 1074

| Profile Score | Tandem Score | TDA q-value | Volume | Spectrum Matches |
| --- | --- | --- | --- | --- |
| 14.6643 | 18.2060 | 0.0010 | 5.9666e+08 | 4 |

**Adduct Signal Fractions**

- **Unmodified**: 5.9666e+08

#### EGVFVSN(N)GTHWFVTQR Hex 7 HexNAc 6 Neu5Ac 1

**Mass:**  4506.8470

(-3.285e-07)

**Charge States:** 3, 4

**Start Position**: 1079

**End Position**: 1095

| Profile Score | Tandem Score | TDA q-value | Volume | Spectrum Matches |
| --- | --- | --- | --- | --- |
| 16.3262 | 18.2024 | 0.0010 | 5.3431e+09 | 7 |

**Adduct Signal Fractions**

- **Unmodified**: 5.3431e+09

#### EGVFVSN(N)GTHWFVTQR Hex 6 HexNAc 2

**Mass:**  3241.3812

(-2.205e-07)

**Charge States:** 3

**Start Position**: 1079

**End Position**: 1095

| Profile Score | Tandem Score | TDA q-value | Volume | Spectrum Matches |
| --- | --- | --- | --- | --- |
| 15.8028 | 18.1020 | 0.0010 | 3.1130e+09 | 4 |

**Adduct Signal Fractions**

- **Unmodified**: 3.1130e+09

#### EGVFVSN(N)GTHWFVTQR Fuc 1 Hex 4 HexNAc 3 Neu5Ac 1

**Mass:**  3557.5084

(8.226e-07)

**Charge States:** 3, 4

**Start Position**: 1079

**End Position**: 1095

| Profile Score | Tandem Score | TDA q-value | Volume | Spectrum Matches |
| --- | --- | --- | --- | --- |
| 16.6362 | 18.0678 | 0.0010 | 3.2076e+09 | 9 |

**Adduct Signal Fractions**

- **Unmodified**: 3.2076e+09

#### N(N)FTTAPAIC(C)HDGK Hex 5 HexNAc 4 Neu5Ac 1

**Mass:**  3344.3377

(9.857e-08)

**Charge States:** 3, 4

**Start Position**: 1061

**End Position**: 1074

| Profile Score | Tandem Score | TDA q-value | Volume | Spectrum Matches |
| --- | --- | --- | --- | --- |
| 16.0694 | 17.9701 | 0.0010 | 7.6686e+08 | 3 |

**Adduct Signal Fractions**

- **Unmodified**: 7.6686e+08

#### EGVFVSN(N)GTHWFVTQR Fuc 1 Hex 4 HexNAc 3

**Mass:**  3266.4149

(-8.263e-08)

**Charge States:** 3, 4

**Start Position**: 1079

**End Position**: 1095

| Profile Score | Tandem Score | TDA q-value | Volume | Spectrum Matches |
| --- | --- | --- | --- | --- |
| 17.8607 | 17.8633 | 0.0010 | 1.8960e+10 | 16 |

**Adduct Signal Fractions**

- **NH3**: 2.0250e+07
- **Unmodified**: 1.8940e+10

#### EGVFVSN(N)GTHWFVTQR Hex 4 HexNAc 6 Neu5Ac 1

**Mass:**  4020.6899

(2.465e-07)

**Charge States:** 3, 4

**Start Position**: 1079

**End Position**: 1095

| Profile Score | Tandem Score | TDA q-value | Volume | Spectrum Matches |
| --- | --- | --- | --- | --- |
| 14.4671 | 17.4069 | 0.0010 | 6.6080e+09 | 10 |

**Adduct Signal Fractions**

- **Unmodified**: 6.6080e+09

#### EGVFVSN(N)GTHWFVTQR Hex 5 HexNAc 6 Neu5Ac 1

**Mass:**  4182.7416

(6.871e-08)

**Charge States:** 3, 4

**Start Position**: 1079

**End Position**: 1095

| Profile Score | Tandem Score | TDA q-value | Volume | Spectrum Matches |
| --- | --- | --- | --- | --- |
| 13.5874 | 17.3967 | 0.0010 | 6.2022e+09 | 8 |

**Adduct Signal Fractions**

- **Unmodified**: 6.2022e+09

#### N(N)FTTAPAIC(C)HDGK Fuc 1 Hex 7 HexNAc 6 Neu5Ac 2

**Mass:**  4511.7582

(5.453e-07)

**Charge States:** 3, 4

**Start Position**: 1061

**End Position**: 1074

| Profile Score | Tandem Score | TDA q-value | Volume | Spectrum Matches |
| --- | --- | --- | --- | --- |
| 12.5567 | 17.3053 | 0.0010 | 1.0527e+09 | 4 |

**Adduct Signal Fractions**

- **Unmodified**: 1.0527e+09

#### EGVFVSN(N)GTHWFVTQR Fuc 1 Hex 4 HexNAc 4

**Mass:**  3469.4909

(-3.397e-09)

**Charge States:** 3, 4

**Start Position**: 1079

**End Position**: 1095

| Profile Score | Tandem Score | TDA q-value | Volume | Spectrum Matches |
| --- | --- | --- | --- | --- |
| 11.4150 | 17.0175 | 0.0010 | 6.2594e+09 | 15 |

**Adduct Signal Fractions**

- **Unmodified**: 6.2594e+09

#### EGVFVSN(N)GTHWFVTQR Hex 3 HexNAc 3

**Mass:**  2958.3053

(6.168e-07)

**Charge States:** 2, 3

**Start Position**: 1079

**End Position**: 1095

| Profile Score | Tandem Score | TDA q-value | Volume | Spectrum Matches |
| --- | --- | --- | --- | --- |
| 19.1361 | 16.8563 | 0.0010 | 1.5971e+10 | 12 |

**Adduct Signal Fractions**

- **Unmodified**: 1.5971e+10

#### N(N)FTTAPAIC(C)HDGK Fuc 1 Hex 4 HexNAc 6 Neu5Ac 1

**Mass:**  3734.5024

(2.672e-07)

**Charge States:** 3, 4

**Start Position**: 1061

**End Position**: 1074

| Profile Score | Tandem Score | TDA q-value | Volume | Spectrum Matches |
| --- | --- | --- | --- | --- |
| 14.0902 | 16.5949 | 0.0010 | 1.8695e+09 | 10 |

**Adduct Signal Fractions**

- **Unmodified**: 1.8695e+09

#### N(N)FTTAPAIC(C)HDGK Fuc 1 Hex 5 HexNAc 4

**Mass:**  3199.3012

(3.140e-04)

**Charge States:** 3, 4

**Start Position**: 1061

**End Position**: 1074

| Profile Score | Tandem Score | TDA q-value | Volume | Spectrum Matches |
| --- | --- | --- | --- | --- |
| 16.6248 | 16.4674 | 0.0010 | 6.7570e+10 | 33 |

**Adduct Signal Fractions**

- **Unmodified**: 6.7570e+10

#### DFGGFN(N)FSQILPDPSKPSK Hex 7 HexNAc 2

**Mass:**  3620.5561

(3.675e-07)

**Charge States:** 3, 4

**Start Position**: 783

**End Position**: 802

| Profile Score | Tandem Score | TDA q-value | Volume | Spectrum Matches |
| --- | --- | --- | --- | --- |
| 15.1112 | 16.4277 | 0.0010 | 7.2170e+10 | 21 |

**Adduct Signal Fractions**

- **Unmodified**: 7.2170e+10

#### EGVFVSN(N)GTHWFVTQR Fuc 1 Hex 5 HexNAc 5 Neu5Ac 1

**Mass:**  4125.7144

(-2.029e-07)

**Charge States:** 3, 4

**Start Position**: 1079

**End Position**: 1095

| Profile Score | Tandem Score | TDA q-value | Volume | Spectrum Matches |
| --- | --- | --- | --- | --- |
| 10.9464 | 16.4078 | 0.0010 | 8.8319e+09 | 13 |

**Adduct Signal Fractions**

- **Unmodified**: 8.8319e+09

#### DFGGFN(N)FSQILPDPSKPSK Hex 8 HexNAc 2

**Mass:**  3782.6080

(3.159e-07)

**Charge States:** 3, 4

**Start Position**: 783

**End Position**: 802

| Profile Score | Tandem Score | TDA q-value | Volume | Spectrum Matches |
| --- | --- | --- | --- | --- |
| 15.1247 | 16.3772 | 0.0010 | 6.9763e+09 | 9 |

**Adduct Signal Fractions**

- **Unmodified**: 6.9763e+09

#### EGVFVSN(N)GTHWFVTQR Fuc 1 Hex 3 HexNAc 3

**Mass:**  3104.3631

(-1.023e-07)

**Charge States:** 2, 3

**Start Position**: 1079

**End Position**: 1095

| Profile Score | Tandem Score | TDA q-value | Volume | Spectrum Matches |
| --- | --- | --- | --- | --- |
| 16.0496 | 16.1878 | 0.0010 | 9.1100e+09 | 13 |

**Adduct Signal Fractions**

- **NH3**: 1.0969e+08
- **Unmodified**: 9.0003e+09

#### N(N)FTTAPAIC(C)HDGK Fuc 1 Hex 6 HexNAc 6 Neu5Ac 1

**Mass:**  4058.6099

(2.879e-07)

**Charge States:** 3, 4

**Start Position**: 1061

**End Position**: 1074

| Profile Score | Tandem Score | TDA q-value | Volume | Spectrum Matches |
| --- | --- | --- | --- | --- |
| 10.5210 | 16.1786 | 0.0010 | 1.8322e+09 | 9 |

**Adduct Signal Fractions**

- **Unmodified**: 1.8322e+09

#### N(N)FTTAPAIC(C)HDGK Fuc 3 Hex 5 HexNAc 4

**Mass:**  3491.4000

(-4.741e-06)

**Charge States:** 4

**Start Position**: 1061

**End Position**: 1074

| Profile Score | Tandem Score | TDA q-value | Volume | Spectrum Matches |
| --- | --- | --- | --- | --- |
| -18.9545 | 16.0236 | 0.0010 | 2.3776e+07 | 2 |

**Adduct Signal Fractions**

- **Unmodified**: 2.3776e+07

#### DFGGFN(N)FSQILPDPSKPSK Fuc 1 Hex 4 HexNAc 4 Neu5Ac 1

**Mass:**  3977.7112

(1.080e-06)

**Charge States:** 3, 4

**Start Position**: 783

**End Position**: 802

| Profile Score | Tandem Score | TDA q-value | Volume | Spectrum Matches |
| --- | --- | --- | --- | --- |
| 17.5445 | 15.8621 | 0.0010 | 4.3200e+10 | 17 |

**Adduct Signal Fractions**

- **Unmodified**: 4.3200e+10

#### DFGGFN(N)FSQILPDPSKPSKR Hex 7 HexNAc 2

**Mass:**  3776.6566

(5.083e-09)

**Charge States:** 3, 4

**Start Position**: 783

**End Position**: 803

| Profile Score | Tandem Score | TDA q-value | Volume | Spectrum Matches |
| --- | --- | --- | --- | --- |
| 10.2244 | 15.7035 | 0.0010 | 3.2765e+09 | 10 |

**Adduct Signal Fractions**

- **Unmodified**: 3.2765e+09

#### FPNITNLC(C)PFGEVFN(N)ATR Fuc 1 Hex 9 HexNAc 6

**Mass:**  4919.0214

(-1.528e-07)

**Charge States:** 4

**Start Position**: 316

**End Position**: 334

| Profile Score | Tandem Score | TDA q-value | Volume | Spectrum Matches |
| --- | --- | --- | --- | --- |
| 14.2145 | 15.6127 | 0.0010 | 7.8807e+08 | 2 |

**Adduct Signal Fractions**

- **Unmodified**: 7.8807e+08

#### DFGGFN(N)FSQILPDPSKPSKR Hex 5 HexNAc 2

**Mass:**  3452.5507

(-5.601e-07)

**Charge States:** 3, 4

**Start Position**: 783

**End Position**: 803

| Profile Score | Tandem Score | TDA q-value | Volume | Spectrum Matches |
| --- | --- | --- | --- | --- |
| 15.0814 | 15.4374 | 0.0010 | 2.7733e+10 | 8 |

**Adduct Signal Fractions**

- **Unmodified**: 2.7733e+10

#### EGVFVSN(N)GTHWFVTQR Fuc 1 Hex 3 HexNAc 4

**Mass:**  3307.4382

(-1.474e-07)

**Charge States:** 3, 4

**Start Position**: 1079

**End Position**: 1095

| Profile Score | Tandem Score | TDA q-value | Volume | Spectrum Matches |
| --- | --- | --- | --- | --- |
| 17.7094 | 15.3449 | 0.0010 | 2.9708e+10 | 14 |

**Adduct Signal Fractions**

- **NH3**: 7.0239e+06
- **Unmodified**: 2.9701e+10

#### N(N)FTTAPAIC(C)HDGK Hex 5 HexNAc 2

**Mass:**  2647.0844

(-2.058e-07)

**Charge States:** 2, 3, 4

**Start Position**: 1061

**End Position**: 1074

| Profile Score | Tandem Score | TDA q-value | Volume | Spectrum Matches |
| --- | --- | --- | --- | --- |
| 19.4019 | 15.1871 | 0.0010 | 4.1732e+11 | 54 |

**Adduct Signal Fractions**

- **Unmodified**: 4.1732e+11

#### EGVFVSN(N)GTHWFVTQR Fuc 1 Hex 5 HexNAc 4 Neu5Ac 1

**Mass:**  3922.6400

(-6.300e-07)

**Charge States:** 3, 4

**Start Position**: 1079

**End Position**: 1095

| Profile Score | Tandem Score | TDA q-value | Volume | Spectrum Matches |
| --- | --- | --- | --- | --- |
| 15.3853 | 15.1510 | 0.0010 | 2.0872e+10 | 8 |

**Adduct Signal Fractions**

- **Unmodified**: 2.0872e+10

#### EGVFVSN(N)GTHWFVTQR Hex 4 HexNAc 5

**Mass:**  3526.5142

(1.935e-07)

**Charge States:** 3, 4

**Start Position**: 1079

**End Position**: 1095

| Profile Score | Tandem Score | TDA q-value | Volume | Spectrum Matches |
| --- | --- | --- | --- | --- |
| 13.5484 | 15.0969 | 0.0010 | 3.8136e+09 | 10 |

**Adduct Signal Fractions**

- **Unmodified**: 3.8136e+09

#### EGVFVSN(N)GTHWFVTQR Hex 4 HexNAc 4

**Mass:**  3323.4329

(-4.921e-07)

**Charge States:** 3, 4

**Start Position**: 1079

**End Position**: 1095

| Profile Score | Tandem Score | TDA q-value | Volume | Spectrum Matches |
| --- | --- | --- | --- | --- |
| 14.8401 | 14.9885 | 0.0010 | 9.9134e+09 | 9 |

**Adduct Signal Fractions**

- **Unmodified**: 9.9134e+09

#### DFGGFN(N)FSQILPDPSKPSK Hex 5 HexNAc 2

**Mass:**  3296.4475

(1.264e-07)

**Charge States:** 3, 4

**Start Position**: 783

**End Position**: 802

| Profile Score | Tandem Score | TDA q-value | Volume | Spectrum Matches |
| --- | --- | --- | --- | --- |
| 18.2377 | 14.8533 | 0.0010 | 7.0669e+11 | 70 |

**Adduct Signal Fractions**

- **Na**: 1.5734e+08
- **Unmodified**: 7.0653e+11

#### DFGGFN(N)FSQILPDPSKPSK Fuc 1 Hex 5 HexNAc 4

**Mass:**  3848.6648

(1.972e-07)

**Charge States:** 3, 4

**Start Position**: 783

**End Position**: 802

| Profile Score | Tandem Score | TDA q-value | Volume | Spectrum Matches |
| --- | --- | --- | --- | --- |
| 12.5858 | 14.7671 | 0.0010 | 4.7187e+09 | 20 |

**Adduct Signal Fractions**

- **NH3**: 1.0044e+07
- **Unmodified**: 4.7086e+09

#### DFGGFN(N)FSQILPDPSKPSK Fuc 1 Hex 3 HexNAc 4

**Mass:**  3524.5645

(-1.316e-07)

**Charge States:** 3, 4

**Start Position**: 783

**End Position**: 802

| Profile Score | Tandem Score | TDA q-value | Volume | Spectrum Matches |
| --- | --- | --- | --- | --- |
| 16.9742 | 14.6843 | 0.0010 | 2.1191e+10 | 11 |

**Adduct Signal Fractions**

- **Unmodified**: 2.1191e+10

#### DFGGFN(N)FSQILPDPSKPSK Fuc 1 Hex 4 HexNAc 4

**Mass:**  3686.6158

(1.120e-06)

**Charge States:** 3, 4

**Start Position**: 783

**End Position**: 802

| Profile Score | Tandem Score | TDA q-value | Volume | Spectrum Matches |
| --- | --- | --- | --- | --- |
| -4.4335 | 14.6750 | 0.0010 | 1.0392e+10 | 21 |

**Adduct Signal Fractions**

- **Unmodified**: 1.0392e+10

#### DFGGFN(N)FSQILPDPSKPSK Hex 6 HexNAc 2

**Mass:**  3458.5014

(-4.665e-07)

**Charge States:** 3, 4

**Start Position**: 783

**End Position**: 802

| Profile Score | Tandem Score | TDA q-value | Volume | Spectrum Matches |
| --- | --- | --- | --- | --- |
| 15.4304 | 14.6460 | 0.0010 | 4.3112e+10 | 22 |

**Adduct Signal Fractions**

- **Na**: 4.1023e+07
- **Unmodified**: 4.3071e+10

#### EGVFVSN(N)GTHWFVTQR Fuc 1 Hex 4 HexNAc 6 Neu5Ac 1

**Mass:**  4166.7465

(-1.141e-07)

**Charge States:** 3, 4

**Start Position**: 1079

**End Position**: 1095

| Profile Score | Tandem Score | TDA q-value | Volume | Spectrum Matches |
| --- | --- | --- | --- | --- |
| 15.4984 | 14.6365 | 0.0010 | 1.1424e+10 | 5 |

**Adduct Signal Fractions**

- **Unmodified**: 1.1424e+10

#### EGVFVSN(N)GTHWFVTQR Hex 3 HexNAc 5

**Mass:**  3364.4609

(-2.177e-07)

**Charge States:** 3, 4

**Start Position**: 1079

**End Position**: 1095

| Profile Score | Tandem Score | TDA q-value | Volume | Spectrum Matches |
| --- | --- | --- | --- | --- |
| 17.9758 | 14.5203 | 0.0010 | 1.2317e+10 | 14 |

**Adduct Signal Fractions**

- **Unmodified**: 1.2317e+10

#### DFGGFN(N)FSQILPDPSKPSK Hex 4 HexNAc 3

**Mass:**  3337.4750

(5.526e-07)

**Charge States:** 3, 4

**Start Position**: 783

**End Position**: 802

| Profile Score | Tandem Score | TDA q-value | Volume | Spectrum Matches |
| --- | --- | --- | --- | --- |
| 15.7229 | 14.4868 | 0.0010 | 2.9459e+10 | 13 |

**Adduct Signal Fractions**

- **Unmodified**: 2.9459e+10

#### EGVFVSN(N)GTHWFVTQR Hex 5 HexNAc 4

**Mass:**  3485.4874

(-9.397e-08)

**Charge States:** 3, 4

**Start Position**: 1079

**End Position**: 1095

| Profile Score | Tandem Score | TDA q-value | Volume | Spectrum Matches |
| --- | --- | --- | --- | --- |
| 12.6297 | 14.4472 | 0.0010 | 2.5763e+09 | 11 |

**Adduct Signal Fractions**

- **Unmodified**: 2.5763e+09

#### EGVFVSN(N)GTHWFVTQR Hex 5 HexNAc 6 Neu5Ac 2

**Mass:**  4473.8375

(-2.092e-07)

**Charge States:** 3, 4

**Start Position**: 1079

**End Position**: 1095

| Profile Score | Tandem Score | TDA q-value | Volume | Spectrum Matches |
| --- | --- | --- | --- | --- |
| 14.7179 | 14.2987 | 0.0010 | 1.5038e+09 | 4 |

**Adduct Signal Fractions**

- **Unmodified**: 1.5038e+09

#### EGVFVSN(N)GTHWFVTQR Fuc 1 Hex 5 HexNAc 6 Neu5Ac 1

**Mass:**  4328.7982

(-5.592e-08)

**Charge States:** 3, 4

**Start Position**: 1079

**End Position**: 1095

| Profile Score | Tandem Score | TDA q-value | Volume | Spectrum Matches |
| --- | --- | --- | --- | --- |
| 11.4446 | 14.2553 | 0.0010 | 4.8272e+09 | 9 |

**Adduct Signal Fractions**

- **Unmodified**: 4.8272e+09

#### TQSLLIVNN(N)ATNVVIK Hex 5 HexNAc 3

**Mass:**  3145.5011

(1.574e-07)

**Charge States:** 2, 3, 4

**Start Position**: 101

**End Position**: 117

| Profile Score | Tandem Score | TDA q-value | Volume | Spectrum Matches |
| --- | --- | --- | --- | --- |
| 15.2262 | 14.2463 | 0.0010 | 4.8615e+10 | 24 |

**Adduct Signal Fractions**

- **Unmodified**: 4.8615e+10

#### TQSLLIVNN(N)ATNVVIK Hex 6 HexNAc 2

**Mass:**  3104.4757

(5.003e-07)

**Charge States:** 2, 3

**Start Position**: 101

**End Position**: 117

| Profile Score | Tandem Score | TDA q-value | Volume | Spectrum Matches |
| --- | --- | --- | --- | --- |
| 15.6798 | 14.1695 | 0.0010 | 4.3736e+09 | 5 |

**Adduct Signal Fractions**

- **Unmodified**: 4.3736e+09

#### EGVFVSN(N)GTHWFVTQR Hex 6 HexNAc 6 Neu5Ac 1

**Mass:**  4344.7947

(5.990e-07)

**Charge States:** 3, 4

**Start Position**: 1079

**End Position**: 1095

| Profile Score | Tandem Score | TDA q-value | Volume | Spectrum Matches |
| --- | --- | --- | --- | --- |
| 9.2097 | 14.1439 | 0.0010 | 4.5012e+09 | 15 |

**Adduct Signal Fractions**

- **Unmodified**: 4.5012e+09

#### TQSLLIVNN(N)ATNVVIK Fuc 1 Hex 5 HexNAc 4 Neu5Ac 1

**Mass:**  3785.7378

(6.944e-07)

**Charge States:** 3, 4

**Start Position**: 101

**End Position**: 117

| Profile Score | Tandem Score | TDA q-value | Volume | Spectrum Matches |
| --- | --- | --- | --- | --- |
| 13.1160 | 14.0301 | 0.0010 | 1.4243e+10 | 12 |

**Adduct Signal Fractions**

- **Unmodified**: 1.4243e+10

#### TQSLLIVNN(N)ATNVVIK Fuc 1 Hex 3 HexNAc 3

**Mass:**  2967.4548

(6.481e-08)

**Charge States:** 2, 3

**Start Position**: 101

**End Position**: 117

| Profile Score | Tandem Score | TDA q-value | Volume | Spectrum Matches |
| --- | --- | --- | --- | --- |
| 14.2320 | 13.9192 | 0.0010 | 1.1655e+10 | 7 |

**Adduct Signal Fractions**

- **Unmodified**: 1.1655e+10

#### SSVLHSTQDLFLPFFSNVTWFHAIHVSGTN(N)GTK Fuc 1 Hex 8 HexNAc 6

**Mass:**  6334.7920

(8.198e-07)

**Charge States:** 4, 5, 6

**Start Position**: 32

**End Position**: 65

| Profile Score | Tandem Score | TDA q-value | Volume | Spectrum Matches |
| --- | --- | --- | --- | --- |
| 14.8453 | 13.8715 | 0.0010 | 7.0861e+09 | 2 |

**Adduct Signal Fractions**

- **Unmodified**: 7.0861e+09

#### N(N)FTTAPAIC(C)HDGK Hex 6 HexNAc 2

**Mass:**  2809.1373

(-8.523e-08)

**Charge States:** 2, 3

**Start Position**: 1061

**End Position**: 1074

| Profile Score | Tandem Score | TDA q-value | Volume | Spectrum Matches |
| --- | --- | --- | --- | --- |
| 15.5434 | 13.7629 | 0.0010 | 6.3794e+09 | 10 |

**Adduct Signal Fractions**

- **Unmodified**: 6.3794e+09

#### N(N)FTTAPAIC(C)HDGK Fuc 1 Hex 5 HexNAc 3

**Mass:**  2996.2211

(-5.093e-07)

**Charge States:** 2, 3, 4

**Start Position**: 1061

**End Position**: 1074

| Profile Score | Tandem Score | TDA q-value | Volume | Spectrum Matches |
| --- | --- | --- | --- | --- |
| 16.3742 | 13.7569 | 0.0010 | 3.8295e+10 | 12 |

**Adduct Signal Fractions**

- **Unmodified**: 3.8295e+10

#### DFGGFN(N)FSQILPDPSKPSK Fuc 1 Hex 5 HexNAc 4 Neu5Ac 1

**Mass:**  4139.7630

(5.072e-07)

**Charge States:** 3, 4, 5

**Start Position**: 783

**End Position**: 802

| Profile Score | Tandem Score | TDA q-value | Volume | Spectrum Matches |
| --- | --- | --- | --- | --- |
| 17.8594 | 13.6145 | 0.0010 | 9.2840e+09 | 12 |

**Adduct Signal Fractions**

- **Unmodified**: 9.2840e+09

#### DFGGFN(N)FSQILPDPSKPSK Hex 5 HexNAc 3

**Mass:**  3499.5287

(-1.941e-07)

**Charge States:** 3, 4

**Start Position**: 783

**End Position**: 802

| Profile Score | Tandem Score | TDA q-value | Volume | Spectrum Matches |
| --- | --- | --- | --- | --- |
| 15.5024 | 13.5998 | 0.0010 | 4.6795e+10 | 19 |

**Adduct Signal Fractions**

- **Unmodified**: 4.6795e+10

#### N(N)FTTAPAIC(C)HDGK Fuc 1 Hex 4 HexNAc 3 Neu5Ac 1

**Mass:**  3125.2644

(1.949e-07)

**Charge States:** 2, 3, 4

**Start Position**: 1061

**End Position**: 1074

| Profile Score | Tandem Score | TDA q-value | Volume | Spectrum Matches |
| --- | --- | --- | --- | --- |
| 18.2207 | 13.5836 | 0.0010 | 2.6611e+09 | 6 |

**Adduct Signal Fractions**

- **Unmodified**: 2.6611e+09

#### DFGGFN(N)FSQILPDPSKPSK Fuc 1 Hex 3 HexNAc 5

**Mass:**  3727.6396

(3.341e-07)

**Charge States:** 3, 4, 5

**Start Position**: 783

**End Position**: 802

| Profile Score | Tandem Score | TDA q-value | Volume | Spectrum Matches |
| --- | --- | --- | --- | --- |
| 16.8761 | 13.5772 | 0.0010 | 1.2129e+10 | 14 |

**Adduct Signal Fractions**

- **Unmodified**: 1.2129e+10

#### N(N)FTTAPAIC(C)HDGK Hex 5 HexNAc 4

**Mass:**  3053.2443

(2.110e-07)

**Charge States:** 3

**Start Position**: 1061

**End Position**: 1074

| Profile Score | Tandem Score | TDA q-value | Volume | Spectrum Matches |
| --- | --- | --- | --- | --- |
| 10.6836 | 13.4986 | 0.0010 | 4.1224e+09 | 6 |

**Adduct Signal Fractions**

- **Unmodified**: 4.1224e+09

#### DFGGFN(N)FSQILPDPSKPSK Hex 6 HexNAc 3

**Mass:**  3661.5812

(-8.349e-08)

**Charge States:** 3, 4

**Start Position**: 783

**End Position**: 802

| Profile Score | Tandem Score | TDA q-value | Volume | Spectrum Matches |
| --- | --- | --- | --- | --- |
| 13.7487 | 13.4791 | 0.0010 | 4.1818e+10 | 13 |

**Adduct Signal Fractions**

- **Unmodified**: 4.1818e+10

#### TQSLLIVNN(N)ATNVVIK Fuc 1 Hex 5 HexNAc 3

**Mass:**  3291.5583

(-9.123e-09)

**Charge States:** 3, 4

**Start Position**: 101

**End Position**: 117

| Profile Score | Tandem Score | TDA q-value | Volume | Spectrum Matches |
| --- | --- | --- | --- | --- |
| 15.4341 | 13.4691 | 0.0010 | 1.1211e+10 | 17 |

**Adduct Signal Fractions**

- **Unmodified**: 1.1211e+10

#### N(N)FTTAPAIC(C)HDGK Hex 3 HexNAc 3

**Mass:**  2526.0589

(8.853e-08)

**Charge States:** 2, 3

**Start Position**: 1061

**End Position**: 1074

| Profile Score | Tandem Score | TDA q-value | Volume | Spectrum Matches |
| --- | --- | --- | --- | --- |
| 18.0044 | 13.4623 | 0.0010 | 8.0435e+09 | 5 |

**Adduct Signal Fractions**

- **Unmodified**: 8.0435e+09

#### N(N)FTTAPAIC(C)HDGK Hex 4 HexNAc 3

**Mass:**  2688.1109

(-1.192e-07)

**Charge States:** 2, 3

**Start Position**: 1061

**End Position**: 1074

| Profile Score | Tandem Score | TDA q-value | Volume | Spectrum Matches |
| --- | --- | --- | --- | --- |
| 15.0068 | 13.4566 | 0.0010 | 3.0555e+10 | 13 |

**Adduct Signal Fractions**

- **Unmodified**: 3.0555e+10

#### EGVFVSN(N)GTHWFVTQR Hex 7 HexNAc 2

**Mass:**  3403.4348

(2.047e-07)

**Charge States:** 3

**Start Position**: 1079

**End Position**: 1095

| Profile Score | Tandem Score | TDA q-value | Volume | Spectrum Matches |
| --- | --- | --- | --- | --- |
| 17.0129 | 13.4462 | 0.0010 | 3.3379e+09 | 6 |

**Adduct Signal Fractions**

- **Unmodified**: 3.3379e+09

#### N(N)FTTAPAIC(C)HDGK Hex 3 HexNAc 4

**Mass:**  2729.1378

(1.353e-07)

**Charge States:** 2, 3, 4

**Start Position**: 1061

**End Position**: 1074

| Profile Score | Tandem Score | TDA q-value | Volume | Spectrum Matches |
| --- | --- | --- | --- | --- |
| 14.1326 | 13.4375 | 0.0010 | 5.4784e+09 | 9 |

**Adduct Signal Fractions**

- **Unmodified**: 5.4784e+09

#### DFGGFN(N)FSQILPDPSKPSK Hex 6 HexNAc 3 Neu5Ac 1

**Mass:**  3952.6807

(7.945e-07)

**Charge States:** 3, 4

**Start Position**: 783

**End Position**: 802

| Profile Score | Tandem Score | TDA q-value | Volume | Spectrum Matches |
| --- | --- | --- | --- | --- |
| 16.5520 | 13.4105 | 0.0010 | 2.2975e+10 | 12 |

**Adduct Signal Fractions**

- **Unmodified**: 2.2975e+10

#### EGVFVSN(N)GTHWFVTQR Fuc 1 Hex 6 HexNAc 6 Neu5Ac 1

**Mass:**  4490.8513

(5.043e-07)

**Charge States:** 3, 4

**Start Position**: 1079

**End Position**: 1095

| Profile Score | Tandem Score | TDA q-value | Volume | Spectrum Matches |
| --- | --- | --- | --- | --- |
| 8.0744 | 13.3836 | 0.0010 | 4.3785e+09 | 6 |

**Adduct Signal Fractions**

- **Unmodified**: 4.3785e+09

#### DFGGFN(N)FSQILPDPSKPSK Hex 3 HexNAc 3

**Mass:**  3175.4239

(-3.906e-07)

**Charge States:** 3, 4

**Start Position**: 783

**End Position**: 802

| Profile Score | Tandem Score | TDA q-value | Volume | Spectrum Matches |
| --- | --- | --- | --- | --- |
| 17.5418 | 13.3515 | 0.0010 | 5.7961e+09 | 6 |

**Adduct Signal Fractions**

- **Unmodified**: 5.7961e+09

#### TQSLLIVNN(N)ATNVVIK Fuc 1 Hex 4 HexNAc 4 Neu5Ac 1

**Mass:**  3623.6835

(6.341e-07)

**Charge States:** 3, 4

**Start Position**: 101

**End Position**: 117

| Profile Score | Tandem Score | TDA q-value | Volume | Spectrum Matches |
| --- | --- | --- | --- | --- |
| 13.3204 | 13.1065 | 0.0010 | 3.0152e+10 | 24 |

**Adduct Signal Fractions**

- **Unmodified**: 3.0152e+10

#### EGVFVSN(N)GTHWFVTQR Fuc 1 Hex 3 HexNAc 5

**Mass:**  3510.5178

(-1.958e-07)

**Charge States:** 3, 4

**Start Position**: 1079

**End Position**: 1095

| Profile Score | Tandem Score | TDA q-value | Volume | Spectrum Matches |
| --- | --- | --- | --- | --- |
| 15.0976 | 13.0645 | 0.0010 | 8.5517e+09 | 8 |

**Adduct Signal Fractions**

- **Unmodified**: 8.5517e+09

#### TQSLLIVNN(N)ATNVVIK Hex 6 HexNAc 3

**Mass:**  3307.5549

(2.474e-07)

**Charge States:** 3, 4

**Start Position**: 101

**End Position**: 117

| Profile Score | Tandem Score | TDA q-value | Volume | Spectrum Matches |
| --- | --- | --- | --- | --- |
| 17.5495 | 13.0113 | 0.0010 | 1.3351e+10 | 8 |

**Adduct Signal Fractions**

- **Unmodified**: 1.3351e+10

#### TPPIKDFGGFN(N)FSQILPDPSKPSK Hex 5 HexNAc 2

**Mass:**  3832.7840

(5.258e-07)

**Charge States:** 3, 4, 5

**Start Position**: 778

**End Position**: 802

| Profile Score | Tandem Score | TDA q-value | Volume | Spectrum Matches |
| --- | --- | --- | --- | --- |
| 13.8082 | 13.0089 | 0.0010 | 3.4302e+10 | 21 |

**Adduct Signal Fractions**

- **Na**: 2.9675e+06
- **Unmodified**: 3.4299e+10

#### TQSLLIVNN(N)ATNVVIK Hex 5 HexNAc 4

**Mass:**  3348.5804

(5.991e-07)

**Charge States:** 3

**Start Position**: 101

**End Position**: 117

| Profile Score | Tandem Score | TDA q-value | Volume | Spectrum Matches |
| --- | --- | --- | --- | --- |
| 11.0891 | 12.8461 | 0.0010 | 7.8174e+09 | 12 |

**Adduct Signal Fractions**

- **Unmodified**: 7.8174e+09

#### TQSLLIVNN(N)ATNVVIK Hex 5 HexNAc 2

**Mass:**  2942.4203

(-6.368e-08)

**Charge States:** 2, 3

**Start Position**: 101

**End Position**: 117

| Profile Score | Tandem Score | TDA q-value | Volume | Spectrum Matches |
| --- | --- | --- | --- | --- |
| 18.2947 | 12.7278 | 0.0010 | 1.0451e+11 | 26 |

**Adduct Signal Fractions**

- **Unmodified**: 1.0451e+11

#### TQSLLIVNN(N)ATNVVIK Fuc 1 Hex 4 HexNAc 3 Neu5Ac 1

**Mass:**  3420.6015

(3.724e-07)

**Charge States:** 3

**Start Position**: 101

**End Position**: 117

| Profile Score | Tandem Score | TDA q-value | Volume | Spectrum Matches |
| --- | --- | --- | --- | --- |
| -2.5136 | 12.6928 | 0.0010 | 6.0720e+09 | 13 |

**Adduct Signal Fractions**

- **NH3**: 5.3594e+08
- **Unmodified**: 5.5361e+09

#### TQSLLIVNN(N)ATNVVIK Fuc 1 Hex 4 HexNAc 4

**Mass:**  3332.5848

(-2.864e-07)

**Charge States:** 3, 4

**Start Position**: 101

**End Position**: 117

| Profile Score | Tandem Score | TDA q-value | Volume | Spectrum Matches |
| --- | --- | --- | --- | --- |
| 13.0135 | 12.6535 | 0.0010 | 1.6426e+10 | 13 |

**Adduct Signal Fractions**

- **Unmodified**: 1.6426e+10

#### N(N)FTTAPAIC(C)HDGK Hex 4 HexNAc 2

**Mass:**  2485.0343

(1.061e-06)

**Charge States:** 3

**Start Position**: 1061

**End Position**: 1074

| Profile Score | Tandem Score | TDA q-value | Volume | Spectrum Matches |
| --- | --- | --- | --- | --- |
| 11.9526 | 12.6320 | 0.0010 | 3.2806e+08 | 1 |

**Adduct Signal Fractions**

- **Unmodified**: 3.2806e+08

#### N(N)FTTAPAIC(C)HDGK Fuc 1 Hex 3 HexNAc 5

**Mass:**  3080.2813

(6.517e-04)

**Start Position**: 1061

**End Position**: 1074

| Profile Score | Tandem Score | TDA q-value | Volume | Spectrum Matches |
| --- | --- | --- | --- | --- |
| 0.0000 | 12.6300 | 0.0010 | 0.0000e+00 | 5 |

Adduct:

No Chromatogram Found

#### DFGGFN(N)FSQILPDPSKPSK Fuc 1 Hex 6 HexNAc 3

**Mass:**  3807.6378

(5.065e-08)

**Charge States:** 3, 4

**Start Position**: 783

**End Position**: 802

| Profile Score | Tandem Score | TDA q-value | Volume | Spectrum Matches |
| --- | --- | --- | --- | --- |
| -5.4495 | 12.4483 | 0.0010 | 1.6317e+09 | 7 |

**Adduct Signal Fractions**

- **Unmodified**: 1.6317e+09

#### DFGGFN(N)FSQILPDPSKPSK Hex 5 HexNAc 4 Neu5Ac 1

**Mass:**  3993.7089

(8.908e-07)

**Charge States:** 3, 4

**Start Position**: 783

**End Position**: 802

| Profile Score | Tandem Score | TDA q-value | Volume | Spectrum Matches |
| --- | --- | --- | --- | --- |
| 10.2912 | 12.2385 | 0.0010 | -1.8776e+09 | 10 |

**Adduct Signal Fractions**

- **Unmodified**: 6.7123e+09

#### FPNITNLC(C)PFGEVFN(N)ATR Fuc 2 Hex 6 HexNAc 7

**Mass:**  4782.0106

(3.565e-07)

**Charge States:** 3, 4

**Start Position**: 316

**End Position**: 334

| Profile Score | Tandem Score | TDA q-value | Volume | Spectrum Matches |
| --- | --- | --- | --- | --- |
| 14.1413 | 12.2127 | 0.0010 | 1.9898e+09 | 3 |

**Adduct Signal Fractions**

- **Unmodified**: 1.9898e+09

#### N(N)FTTAPAIC(C)HDGK Fuc 2 Hex 3 HexNAc 5

**Mass:**  3224.3024

(5.272e-03)

**Charge States:** 3, 4

**Start Position**: 1061

**End Position**: 1074

| Profile Score | Tandem Score | TDA q-value | Volume | Spectrum Matches |
| --- | --- | --- | --- | --- |
| -9.9054 | 12.1448 | 0.0010 | 3.0792e+07 | 4 |

**Adduct Signal Fractions**

- **NH3**: 3.0792e+07

#### DFGGFN(N)FSQILPDPSKPSKR Hex 6 HexNAc 3

**Mass:**  3817.6840

(3.373e-07)

**Charge States:** 4

**Start Position**: 783

**End Position**: 803

| Profile Score | Tandem Score | TDA q-value | Volume | Spectrum Matches |
| --- | --- | --- | --- | --- |
| -2.9775 | 12.0878 | 0.0010 | 1.0629e+08 | 1 |

**Adduct Signal Fractions**

- **Unmodified**: 1.0629e+08

#### N(N)FTTAPAIC(C)HDGK Hex 3 HexNAc 5

**Mass:**  2932.2157

(-2.599e-07)

**Charge States:** 3

**Start Position**: 1061

**End Position**: 1074

| Profile Score | Tandem Score | TDA q-value | Volume | Spectrum Matches |
| --- | --- | --- | --- | --- |
| 10.5688 | 12.0710 | 0.0010 | 2.1021e+09 | 5 |

**Adduct Signal Fractions**

- **Unmodified**: 2.1021e+09

#### EGVFVSN(N)GTHWFVTQR Hex 4 HexNAc 3

**Mass:**  3120.3578

(4.419e-07)

**Charge States:** 2, 3, 4

**Start Position**: 1079

**End Position**: 1095

| Profile Score | Tandem Score | TDA q-value | Volume | Spectrum Matches |
| --- | --- | --- | --- | --- |
| 18.6171 | 12.0402 | 0.0010 | 2.0728e+10 | 12 |

**Adduct Signal Fractions**

- **Unmodified**: 2.0728e+10

#### TPPIKDFGGFN(N)FSQILPDPSKPSK Hex 7 HexNAc 2

**Mass:**  4156.8888

(-1.130e-07)

**Charge States:** 3, 4, 5

**Start Position**: 778

**End Position**: 802

| Profile Score | Tandem Score | TDA q-value | Volume | Spectrum Matches |
| --- | --- | --- | --- | --- |
| 14.4133 | 11.9357 | 0.0010 | 2.4822e+09 | 6 |

**Adduct Signal Fractions**

- **Unmodified**: 2.4822e+09

#### EGVFVSN(N)GTHWFVTQR Hex 5 HexNAc 5 Neu5Ac 2

**Mass:**  4270.7591

(5.555e-07)

**Charge States:** 3, 4

**Start Position**: 1079

**End Position**: 1095

| Profile Score | Tandem Score | TDA q-value | Volume | Spectrum Matches |
| --- | --- | --- | --- | --- |
| 12.6912 | 11.9223 | 0.0010 | 2.8607e+09 | 10 |

**Adduct Signal Fractions**

- **Unmodified**: 2.8607e+09

#### TQSLLIVNN(N)ATNVVIK Hex 5 HexNAc 3 Neu5Ac 1

**Mass:**  3436.5982

(5.781e-07)

**Charge States:** 3, 4

**Start Position**: 101

**End Position**: 117

| Profile Score | Tandem Score | TDA q-value | Volume | Spectrum Matches |
| --- | --- | --- | --- | --- |
| 14.8385 | 11.9177 | 0.0010 | 2.4181e+10 | 9 |

**Adduct Signal Fractions**

- **Unmodified**: 2.4181e+10

#### N(N)FTTAPAIC(C)HDGK Fuc 1 Hex 3 HexNAc 6

**Mass:**  3281.3541

(5.843e-07)

**Charge States:** 3, 4

**Start Position**: 1061

**End Position**: 1074

| Profile Score | Tandem Score | TDA q-value | Volume | Spectrum Matches |
| --- | --- | --- | --- | --- |
| 13.8086 | 11.7583 | 0.0010 | 2.2037e+10 | 13 |

**Adduct Signal Fractions**

- **Unmodified**: 2.2037e+10

#### N(N)FTTAPAIC(C)HDGK Fuc 1 Hex 3 HexNAc 3

**Mass:**  2672.1147

(-2.612e-07)

**Charge States:** 2, 3

**Start Position**: 1061

**End Position**: 1074

| Profile Score | Tandem Score | TDA q-value | Volume | Spectrum Matches |
| --- | --- | --- | --- | --- |
| 18.2361 | 11.6820 | 0.0010 | 4.3310e+10 | 17 |

**Adduct Signal Fractions**

- **Unmodified**: 4.3310e+10

#### N(N)FTTAPAIC(C)HDGK Fuc 2 Hex 5 HexNAc 4

**Mass:**  3345.3584

(1.740e-07)

**Charge States:** 3, 4

**Start Position**: 1061

**End Position**: 1074

| Profile Score | Tandem Score | TDA q-value | Volume | Spectrum Matches |
| --- | --- | --- | --- | --- |
| 16.6414 | 11.6786 | 0.0010 | 6.2916e+08 | 6 |

**Adduct Signal Fractions**

- **Unmodified**: 6.2916e+08

#### N(N)FTTAPAIC(C)HDGK Hex 4 HexNAc 3 Neu5Ac 1

**Mass:**  2979.2064

(7.153e-08)

**Charge States:** 3, 4

**Start Position**: 1061

**End Position**: 1074

| Profile Score | Tandem Score | TDA q-value | Volume | Spectrum Matches |
| --- | --- | --- | --- | --- |
| 17.2178 | 11.6157 | 0.0010 | 5.7107e+08 | 3 |

**Adduct Signal Fractions**

- **Unmodified**: 5.7107e+08

#### TQSLLIVNN(N)ATNVVIK Fuc 1 Hex 6 HexNAc 3

**Mass:**  3453.6111

(6.749e-07)

**Charge States:** 3, 4

**Start Position**: 101

**End Position**: 117

| Profile Score | Tandem Score | TDA q-value | Volume | Spectrum Matches |
| --- | --- | --- | --- | --- |
| 12.3575 | 11.5276 | 0.0010 | 3.6951e+09 | 8 |

**Adduct Signal Fractions**

- **Unmodified**: 3.6951e+09

#### TQSLLIVNN(N)ATNVVIK Fuc 1 Hex 4 HexNAc 3

**Mass:**  3129.5063

(1.623e-08)

**Charge States:** 2, 3

**Start Position**: 101

**End Position**: 117

| Profile Score | Tandem Score | TDA q-value | Volume | Spectrum Matches |
| --- | --- | --- | --- | --- |
| 14.4885 | 11.4914 | 0.0010 | 1.6377e+10 | 13 |

**Adduct Signal Fractions**

- **Unmodified**: 1.6377e+10

#### TQSLLIVNN(N)ATNVVIK Hex 3 HexNAc 5

**Mass:**  3227.5517

(-1.115e-07)

**Charge States:** 3, 4

**Start Position**: 101

**End Position**: 117

| Profile Score | Tandem Score | TDA q-value | Volume | Spectrum Matches |
| --- | --- | --- | --- | --- |
| 14.4277 | 11.4821 | 0.0010 | 5.7869e+09 | 8 |

**Adduct Signal Fractions**

- **Unmodified**: 5.7869e+09

#### EGVFVSN(N)GTHWFVTQR Fuc 1 Hex 4 HexNAc 5

**Mass:**  3672.5723

(1.582e-08)

**Charge States:** 3, 4

**Start Position**: 1079

**End Position**: 1095

| Profile Score | Tandem Score | TDA q-value | Volume | Spectrum Matches |
| --- | --- | --- | --- | --- |
| 13.3071 | 11.4448 | 0.0010 | 2.1948e+09 | 8 |

**Adduct Signal Fractions**

- **Unmodified**: 2.1948e+09

#### DFGGFN(N)FSQILPDPSKPSK Fuc 2 Hex 5 HexNAc 3

**Mass:**  3791.6220

(-5.724e-06)

**Charge States:** 3, 4

**Start Position**: 783

**End Position**: 802

| Profile Score | Tandem Score | TDA q-value | Volume | Spectrum Matches |
| --- | --- | --- | --- | --- |
| 14.0749 | 11.4194 | 0.0010 | -5.7592e+08 | 4 |

**Adduct Signal Fractions**

- **Unmodified**: 2.5194e+10

#### TQSLLIVNN(N)ATNVVIK Fuc 1 Hex 5 HexNAc 4

**Mass:**  3494.6371

(1.086e-07)

**Charge States:** 3, 4

**Start Position**: 101

**End Position**: 117

| Profile Score | Tandem Score | TDA q-value | Volume | Spectrum Matches |
| --- | --- | --- | --- | --- |
| 13.6224 | 11.4060 | 0.0010 | 5.2683e+09 | 11 |

**Adduct Signal Fractions**

- **NH3**: 2.6711e+07
- **Unmodified**: 5.2415e+09

#### TQSLLIVNN(N)ATNVVIK Fuc 1 Hex 4 HexNAc 5

**Mass:**  3535.6660

(9.095e-08)

**Charge States:** 3, 4

**Start Position**: 101

**End Position**: 117

| Profile Score | Tandem Score | TDA q-value | Volume | Spectrum Matches |
| --- | --- | --- | --- | --- |
| -5.7323 | 11.3397 | 0.0010 | 4.2381e+09 | 10 |

**Adduct Signal Fractions**

- **Unmodified**: 4.2381e+09

#### TQSLLIVNN(N)ATNVVIK Fuc 1 Hex 5 HexNAc 5

**Mass:**  3697.7176

(-1.175e-07)

**Charge States:** 3, 4

**Start Position**: 101

**End Position**: 117

| Profile Score | Tandem Score | TDA q-value | Volume | Spectrum Matches |
| --- | --- | --- | --- | --- |
| 13.4696 | 11.0549 | 0.0010 | 8.3983e+08 | 2 |

**Adduct Signal Fractions**

- **Unmodified**: 8.3983e+08

#### DFGGFN(N)FSQILPDPSKPSK Hex 5 HexNAc 4

**Mass:**  3702.6091

(4.838e-07)

**Charge States:** 3, 4

**Start Position**: 783

**End Position**: 802

| Profile Score | Tandem Score | TDA q-value | Volume | Spectrum Matches |
| --- | --- | --- | --- | --- |
| 9.3565 | 11.0489 | 0.0010 | 3.6075e+09 | 17 |

**Adduct Signal Fractions**

- **Unmodified**: 3.6075e+09

#### DFGGFN(N)FSQILPDPSKPSK Hex 3 HexNAc 4

**Mass:**  3378.5010

(-1.690e-07)

**Charge States:** 3, 4

**Start Position**: 783

**End Position**: 802

| Profile Score | Tandem Score | TDA q-value | Volume | Spectrum Matches |
| --- | --- | --- | --- | --- |
| -3.0764 | 10.6811 | 0.0010 | 1.4316e+10 | 8 |

**Adduct Signal Fractions**

- **Unmodified**: 1.4316e+10

#### TPPIKDFGGFN(N)FSQILPDPSKPSK Hex 6 HexNAc 2

**Mass:**  3994.8336

(4.970e-07)

**Charge States:** 3, 4, 5

**Start Position**: 778

**End Position**: 802

| Profile Score | Tandem Score | TDA q-value | Volume | Spectrum Matches |
| --- | --- | --- | --- | --- |
| -2.8367 | 10.5397 | 0.0010 | 1.5393e+09 | 13 |

**Adduct Signal Fractions**

- **Unmodified**: 1.5393e+09

#### TQSLLIVNN(N)ATNVVIK Hex 4 HexNAc 3

**Mass:**  2983.4484

(-9.320e-07)

**Charge States:** 2, 3, 4

**Start Position**: 101

**End Position**: 117

| Profile Score | Tandem Score | TDA q-value | Volume | Spectrum Matches |
| --- | --- | --- | --- | --- |
| 16.6959 | 10.4234 | 0.0010 | 1.9566e+10 | 19 |

**Adduct Signal Fractions**

- **NH3**: 1.4092e+08
- **Unmodified**: 1.9425e+10

#### TQSLLIVNN(N)ATNVVIK Fuc 2 Hex 5 HexNAc 3

**Mass:**  3437.6160

(4.898e-07)

**Charge States:** 3

**Start Position**: 101

**End Position**: 117

| Profile Score | Tandem Score | TDA q-value | Volume | Spectrum Matches |
| --- | --- | --- | --- | --- |
| 13.8974 | 10.4181 | 0.0010 | 2.2205e+09 | 9 |

**Adduct Signal Fractions**

- **NH3**: 8.6674e+07
- **Unmodified**: 2.1338e+09

#### EGVFVSN(N)GTHWFVTQR Hex 7 HexNAc 6 Neu5Ac 2

**Mass:**  4797.9469

(1.390e-07)

**Charge States:** 4

**Start Position**: 1079

**End Position**: 1095

| Profile Score | Tandem Score | TDA q-value | Volume | Spectrum Matches |
| --- | --- | --- | --- | --- |
| 9.1602 | 10.4054 | 0.0010 | 1.5648e+09 | 3 |

**Adduct Signal Fractions**

- **Unmodified**: 1.5648e+09

#### N(N)FTTAPAIC(C)HDGK Fuc 1 Hex 6 HexNAc 3

**Mass:**  3158.2756

(3.188e-04)

**Charge States:** 3

**Start Position**: 1061

**End Position**: 1074

| Profile Score | Tandem Score | TDA q-value | Volume | Spectrum Matches |
| --- | --- | --- | --- | --- |
| 9.8778 | 10.3879 | 0.0010 | 2.3184e+09 | 7 |

**Adduct Signal Fractions**

- **Unmodified**: 2.3184e+09

#### TQSLLIVNN(N)ATNVVIK Fuc 2 Hex 3 HexNAc 5

**Mass:**  3519.6716

(-4.564e-07)

**Charge States:** 3, 4

**Start Position**: 101

**End Position**: 117

| Profile Score | Tandem Score | TDA q-value | Volume | Spectrum Matches |
| --- | --- | --- | --- | --- |
| 15.9317 | 10.3776 | 0.0010 | 1.7949e+09 | 4 |

**Adduct Signal Fractions**

- **Unmodified**: 1.7949e+09

#### DFGGFN(N)FSQILPDPSKPSK Hex 4 HexNAc 3 Neu5Ac 1

**Mass:**  3628.5741

(-5.543e-08)

**Charge States:** 3, 4

**Start Position**: 783

**End Position**: 802

| Profile Score | Tandem Score | TDA q-value | Volume | Spectrum Matches |
| --- | --- | --- | --- | --- |
| 14.9596 | 10.3291 | 0.0010 | 1.5336e+09 | 5 |

**Adduct Signal Fractions**

- **Unmodified**: 1.5336e+09

#### TQSLLIVNN(N)ATNVVIK Fuc 2 Hex 4 HexNAc 4

**Mass:**  3478.6429

(-3.406e-08)

**Charge States:** 3, 4

**Start Position**: 101

**End Position**: 117

| Profile Score | Tandem Score | TDA q-value | Volume | Spectrum Matches |
| --- | --- | --- | --- | --- |
| 15.2592 | 10.2957 | 0.0010 | 3.9676e+09 | 5 |

**Adduct Signal Fractions**

- **Unmodified**: 3.9676e+09

#### N(N)FTTAPAIC(C)HDGK Hex 4 HexNAc 4 Neu5Ac 1

**Mass:**  3182.2882

(-2.568e-07)

**Charge States:** 3, 4

**Start Position**: 1061

**End Position**: 1074

| Profile Score | Tandem Score | TDA q-value | Volume | Spectrum Matches |
| --- | --- | --- | --- | --- |
| 16.2938 | 10.2791 | 0.0010 | 4.8639e+08 | 4 |

**Adduct Signal Fractions**

- **Unmodified**: 4.8639e+08

#### TQSLLIVNN(N)ATNVVIK Hex 4 HexNAc 4

**Mass:**  3186.5268

(1.900e-07)

**Charge States:** 2, 3

**Start Position**: 101

**End Position**: 117

| Profile Score | Tandem Score | TDA q-value | Volume | Spectrum Matches |
| --- | --- | --- | --- | --- |
| 14.4495 | 10.2681 | 0.0010 | 1.7296e+10 | 11 |

**Adduct Signal Fractions**

- **Unmodified**: 1.7296e+10

#### TQSLLIVNN(N)ATNVVIK Fuc 1 Hex 3 HexNAc 4

**Mass:**  3170.5322

(-7.656e-08)

**Charge States:** 2, 3, 4

**Start Position**: 101

**End Position**: 117

| Profile Score | Tandem Score | TDA q-value | Volume | Spectrum Matches |
| --- | --- | --- | --- | --- |
| 15.9858 | 10.2339 | 0.0010 | 2.0830e+10 | 15 |

**Adduct Signal Fractions**

- **Unmodified**: 2.0830e+10

#### N(N)FTTAPAIC(C)HDGK Fuc 1 Hex 4 HexNAc 6

**Mass:**  3445.4148

(5.832e-04)

**Start Position**: 1061

**End Position**: 1074

| Profile Score | Tandem Score | TDA q-value | Volume | Spectrum Matches |
| --- | --- | --- | --- | --- |
| 0.0000 | 10.2319 | 0.0010 | 0.0000e+00 | 1 |

Adduct:

No Chromatogram Found

#### NLN(N)ESLIDLQELGK Fuc 1 Hex 3 HexNAc 6

**Mass:**  3435.5309

(7.306e-08)

**Charge States:** 3, 4

**Start Position**: 1179

**End Position**: 1193

| Profile Score | Tandem Score | TDA q-value | Volume | Spectrum Matches |
| --- | --- | --- | --- | --- |
| 16.3820 | 10.2282 | 0.0010 | 5.6114e+09 | 9 |

**Adduct Signal Fractions**

- **Unmodified**: 8.2054e+08
- **NH3**: 4.7908e+09

#### N(N)FTTAPAIC(C)HDGK Fuc 1 Hex 3 HexNAc 4

**Mass:**  2898.1855

(6.966e-04)

**Start Position**: 1061

**End Position**: 1074

| Profile Score | Tandem Score | TDA q-value | Volume | Spectrum Matches |
| --- | --- | --- | --- | --- |
| 0.0000 | 10.1395 | 0.0010 | 0.0000e+00 | 2 |

Adduct:

No Chromatogram Found

#### TQSLLIVNN(N)ATNVVIK Fuc 2 Hex 4 HexNAc 5

**Mass:**  3681.7217

(-7.911e-07)

**Charge States:** 3, 4

**Start Position**: 101

**End Position**: 117

| Profile Score | Tandem Score | TDA q-value | Volume | Spectrum Matches |
| --- | --- | --- | --- | --- |
| 14.6415 | 10.0933 | 0.0010 | 2.5509e+09 | 11 |

**Adduct Signal Fractions**

- **Unmodified**: 2.5509e+09

#### YNEN(N)GTITDAVDC(C)ALDPLSETK Fuc 1 Hex 5 HexNAc 5

**Mass:**  4396.8187

(2.811e-07)

**Charge States:** 3, 4

**Start Position**: 266

**End Position**: 288

| Profile Score | Tandem Score | TDA q-value | Volume | Spectrum Matches |
| --- | --- | --- | --- | --- |
| 14.0029 | 10.0680 | 0.0010 | -1.3094e+09 | 10 |

**Adduct Signal Fractions**

- **NH3**: 1.2325e+07
- **Unmodified**: 7.2682e+09

#### DFGGFN(N)FSQILPDPSKPSK Hex 4 HexNAc 4 Neu5Ac 1

**Mass:**  3831.6493

(-2.223e-07)

**Charge States:** 3, 4

**Start Position**: 783

**End Position**: 802

| Profile Score | Tandem Score | TDA q-value | Volume | Spectrum Matches |
| --- | --- | --- | --- | --- |
| 14.7974 | 9.8720 | 0.0010 | 7.8210e+09 | 8 |

**Adduct Signal Fractions**

- **Unmodified**: 7.8210e+09

#### DFGGFN(N)FSQILPDPSKPSK Hex 5 HexNAc 3 Neu5Ac 1

**Mass:**  3790.6200

(-1.102e-06)

**Charge States:** 4

**Start Position**: 783

**End Position**: 802

| Profile Score | Tandem Score | TDA q-value | Volume | Spectrum Matches |
| --- | --- | --- | --- | --- |
| 14.1710 | 9.8316 | 0.0010 | 1.3936e+10 | 2 |

**Adduct Signal Fractions**

- **Unmodified**: 1.3936e+10

#### DFGGFN(N)FSQILPDPSKPSK Hex 8 HexNAc 2

**Mass:**  3783.6120

(2.655e-04)

**Start Position**: 783

**End Position**: 802

| Profile Score | Tandem Score | TDA q-value | Volume | Spectrum Matches |
| --- | --- | --- | --- | --- |
| 0.0000 | 9.7628 | 0.0010 | 0.0000e+00 | 1 |

Adduct:

No Chromatogram Found

#### EGVFVSN(N)GTHWFVTQR Fuc 1 Hex 5 HexNAc 4

**Mass:**  3631.5457

(9.088e-08)

**Charge States:** 3, 4

**Start Position**: 1079

**End Position**: 1095

| Profile Score | Tandem Score | TDA q-value | Volume | Spectrum Matches |
| --- | --- | --- | --- | --- |
| 12.8891 | 9.6996 | 0.0010 | 1.1272e+09 | 8 |

**Adduct Signal Fractions**

- **Unmodified**: 1.1272e+09

#### DFGGFN(N)FSQILPDPSKPSK Fuc 1 Hex 4 HexNAc 3

**Mass:**  3483.5364

(2.495e-07)

**Charge States:** 3, 4

**Start Position**: 783

**End Position**: 802

| Profile Score | Tandem Score | TDA q-value | Volume | Spectrum Matches |
| --- | --- | --- | --- | --- |
| 14.4647 | 9.6944 | 0.0010 | 1.3140e+09 | 4 |

**Adduct Signal Fractions**

- **Unmodified**: 1.3140e+09

#### N(N)FTTAPAIC(C)HDGK Fuc 1 Hex 5 HexNAc 6 Neu5Ac 1

**Mass:**  3896.5557

(6.117e-07)

**Charge States:** 3, 4

**Start Position**: 1061

**End Position**: 1074

| Profile Score | Tandem Score | TDA q-value | Volume | Spectrum Matches |
| --- | --- | --- | --- | --- |
| 9.6135 | 9.6440 | 0.0010 | 2.0227e+09 | 17 |

**Adduct Signal Fractions**

- **Unmodified**: 2.0227e+09

#### DFGGFN(N)FSQILPDPSKPSK Fuc 1 Hex 3 HexNAc 3

**Mass:**  3321.4823

(4.261e-07)

**Charge States:** 3, 4

**Start Position**: 783

**End Position**: 802

| Profile Score | Tandem Score | TDA q-value | Volume | Spectrum Matches |
| --- | --- | --- | --- | --- |
| 13.5847 | 9.5716 | 0.0010 | 8.9499e+08 | 5 |

**Adduct Signal Fractions**

- **Unmodified**: 8.9499e+08

#### N(N)FTTAPAIC(C)HDGK Fuc 1 Hex 3 HexNAc 7

**Mass:**  3484.4383

(-8.939e-07)

**Charge States:** 3

**Start Position**: 1061

**End Position**: 1074

| Profile Score | Tandem Score | TDA q-value | Volume | Spectrum Matches |
| --- | --- | --- | --- | --- |
| -5.1581 | 9.3159 | 0.0010 | 1.4942e+08 | 2 |

**Adduct Signal Fractions**

- **Unmodified**: 1.4942e+08

#### TQSLLIVNN(N)ATNVVIK Hex 4 HexNAc 3 Neu5Ac 1

**Mass:**  3274.5443

(-4.326e-07)

**Charge States:** 3, 4

**Start Position**: 101

**End Position**: 117

| Profile Score | Tandem Score | TDA q-value | Volume | Spectrum Matches |
| --- | --- | --- | --- | --- |
| 13.3972 | 9.2679 | 0.0010 | 4.9139e+09 | 10 |

**Adduct Signal Fractions**

- **NH3**: 1.4242e+07
- **Unmodified**: 4.8996e+09

#### EGVFVSN(N)GTHWFVTQR Hex 7 HexNAc 3 Neu5Ac 1

**Mass:**  3897.6083

(2.947e-09)

**Charge States:** 3, 4

**Start Position**: 1079

**End Position**: 1095

| Profile Score | Tandem Score | TDA q-value | Volume | Spectrum Matches |
| --- | --- | --- | --- | --- |
| 12.4989 | 9.2246 | 0.0010 | 9.8377e+08 | 7 |

**Adduct Signal Fractions**

- **Unmodified**: 9.8377e+08

#### EGVFVSN(N)GTHWFVTQR Hex 7 HexNAc 3

**Mass:**  3606.5143

(-3.430e-07)

**Charge States:** 3, 4

**Start Position**: 1079

**End Position**: 1095

| Profile Score | Tandem Score | TDA q-value | Volume | Spectrum Matches |
| --- | --- | --- | --- | --- |
| 14.1741 | 9.1680 | 0.0010 | 1.2878e+09 | 8 |

**Adduct Signal Fractions**

- **Na**: 1.1204e+07
- **Unmodified**: 1.2766e+09

#### TQSLLIVNN(N)ATNVVIK Fuc 1 Hex 3 HexNAc 5

**Mass:**  3373.6110

(-1.356e-07)

**Charge States:** 3, 4

**Start Position**: 101

**End Position**: 117

| Profile Score | Tandem Score | TDA q-value | Volume | Spectrum Matches |
| --- | --- | --- | --- | --- |
| 13.4738 | 9.1200 | 0.0010 | 9.4273e+09 | 9 |

**Adduct Signal Fractions**

- **Unmodified**: 9.4273e+09

#### EGVFVSN(N)GTHWFVTQR Fuc 1 Hex 4 HexNAc 6

**Mass:**  3875.6510

(-4.898e-08)

**Charge States:** 3, 4

**Start Position**: 1079

**End Position**: 1095

| Profile Score | Tandem Score | TDA q-value | Volume | Spectrum Matches |
| --- | --- | --- | --- | --- |
| 11.8679 | 9.1033 | 0.0010 | 1.3750e+09 | 5 |

**Adduct Signal Fractions**

- **NH3**: 5.0218e+07
- **Unmodified**: 1.3248e+09

#### TQSLLIVNN(N)ATNVVIK Hex 7 HexNAc 2

**Mass:**  3266.5301

(6.196e-07)

**Charge States:** 3

**Start Position**: 101

**End Position**: 117

| Profile Score | Tandem Score | TDA q-value | Volume | Spectrum Matches |
| --- | --- | --- | --- | --- |
| 16.4611 | 9.1027 | 0.0010 | 6.0737e+09 | 6 |

**Adduct Signal Fractions**

- **Unmodified**: 6.0737e+09

#### N(N)FTTAPAIC(C)HDGK Hex 5 HexNAc 3

**Mass:**  2850.1651

(3.528e-04)

**Charge States:** 2, 3

**Start Position**: 1061

**End Position**: 1074

| Profile Score | Tandem Score | TDA q-value | Volume | Spectrum Matches |
| --- | --- | --- | --- | --- |
| 16.5840 | 9.0779 | 0.0010 | 5.9177e+10 | 19 |

**Adduct Signal Fractions**

- **Unmodified**: 5.9177e+10

#### NLN(N)ESLIDLQELGK Fuc 1 Hex 5 HexNAc 6 Neu5Ac 2

**Mass:**  4341.8260

(2.790e-07)

**Charge States:** 3, 4

**Start Position**: 1179

**End Position**: 1193

| Profile Score | Tandem Score | TDA q-value | Volume | Spectrum Matches |
| --- | --- | --- | --- | --- |
| 14.3686 | 9.0747 | 0.0010 | 5.2342e+08 | 2 |

**Adduct Signal Fractions**

- **Unmodified**: 5.2342e+08

#### TQSLLIVNN(N)ATNVVIK Hex 6 HexNAc 3 Neu5Ac 1

**Mass:**  3598.6497

(-2.795e-07)

**Charge States:** 3, 4

**Start Position**: 101

**End Position**: 117

| Profile Score | Tandem Score | TDA q-value | Volume | Spectrum Matches |
| --- | --- | --- | --- | --- |
| 11.1730 | 9.0737 | 0.0010 | 9.0196e+09 | 14 |

**Adduct Signal Fractions**

- **Unmodified**: 9.0196e+09

#### EGVFVSN(N)GTHWFVTQR Fuc 1 Hex 6 HexNAc 5 Neu5Ac 2

**Mass:**  4581.8850

(6.587e-04)

**Start Position**: 1079

**End Position**: 1095

| Profile Score | Tandem Score | TDA q-value | Volume | Spectrum Matches |
| --- | --- | --- | --- | --- |
| 0.0000 | 9.0686 | 0.0010 | 0.0000e+00 | 2 |

Adduct:

No Chromatogram Found

#### EGVFVSN(N)GTHWFVTQR Hex 4 HexNAc 6

**Mass:**  3729.5940

(3.321e-07)

**Charge States:** 3, 4

**Start Position**: 1079

**End Position**: 1095

| Profile Score | Tandem Score | TDA q-value | Volume | Spectrum Matches |
| --- | --- | --- | --- | --- |
| 13.1149 | 9.0381 | 0.0010 | 1.6430e+09 | 7 |

**Adduct Signal Fractions**

- **Unmodified**: 1.6430e+09

#### VYSSANN(N)C(C)TFEYVSQPFLMDLEGK Fuc 1 Hex 5 HexNAc 4 Neu5Ac 1

**Mass:**  4857.9955

(3.709e-07)

**Charge States:** 4, 5

**Start Position**: 146

**End Position**: 170

| Profile Score | Tandem Score | TDA q-value | Volume | Spectrum Matches |
| --- | --- | --- | --- | --- |
| 14.9361 | 9.0298 | 0.0010 | 6.7447e+09 | 9 |

**Adduct Signal Fractions**

- **NH3**: 1.4853e+08
- **Unmodified**: 6.5962e+09

#### N(N)FTTAPAIC(C)HDGK Hex 4 HexNAc 5

**Mass:**  3094.2702

(2.931e-07)

**Charge States:** 3, 4

**Start Position**: 1061

**End Position**: 1074

| Profile Score | Tandem Score | TDA q-value | Volume | Spectrum Matches |
| --- | --- | --- | --- | --- |
| 9.8756 | 9.0046 | 0.0010 | 1.2407e+09 | 7 |

**Adduct Signal Fractions**

- **Unmodified**: 1.2407e+09

#### EGVFVSN(N)GTHWFVTQR Fuc 1 Hex 3 HexNAc 6

**Mass:**  3713.5993

(-6.376e-08)

**Charge States:** 3, 4

**Start Position**: 1079

**End Position**: 1095

| Profile Score | Tandem Score | TDA q-value | Volume | Spectrum Matches |
| --- | --- | --- | --- | --- |
| 17.2089 | 8.9997 | 0.0010 | 3.5506e+09 | 7 |

**Adduct Signal Fractions**

- **Unmodified**: 3.5506e+09

#### TQSLLIVNN(N)ATNVVIK Fuc 1 Hex 5 HexNAc 3 Neu5Ac 1

**Mass:**  3582.6569

(3.366e-07)

**Charge States:** 3, 4

**Start Position**: 101

**End Position**: 117

| Profile Score | Tandem Score | TDA q-value | Volume | Spectrum Matches |
| --- | --- | --- | --- | --- |
| 11.8827 | 8.8930 | 0.0010 | 1.2109e+10 | 9 |

**Adduct Signal Fractions**

- **Unmodified**: 1.2109e+10

#### TQSLLIVNN(N)ATNVVIK Hex 3 HexNAc 4

**Mass:**  3024.4738

(-2.667e-07)

**Charge States:** 2, 3, 4

**Start Position**: 101

**End Position**: 117

| Profile Score | Tandem Score | TDA q-value | Volume | Spectrum Matches |
| --- | --- | --- | --- | --- |
| 14.9901 | 8.8729 | 0.0010 | 9.1501e+09 | 12 |

**Adduct Signal Fractions**

- **Unmodified**: 9.1501e+09

#### SSVLHSTQDLFLPFFSNVTWFHAIHVSGTN(N)GTK Fuc 3 Hex 9 HexNAc 7

**Mass:**  6992.0399

(1.818e-07)

**Charge States:** 5, 6

**Start Position**: 32

**End Position**: 65

| Profile Score | Tandem Score | TDA q-value | Volume | Spectrum Matches |
| --- | --- | --- | --- | --- |
| 12.0789 | 8.8594 | 0.0010 | 1.6191e+09 | 4 |

**Adduct Signal Fractions**

- **Unmodified**: 1.6191e+09

#### N(N)FTTAPAIC(C)HDGK Fuc 1 Hex 6 HexNAc 3 Neu5Ac 1

**Mass:**  3449.3707

(2.486e-07)

**Charge States:** 3, 4

**Start Position**: 1061

**End Position**: 1074

| Profile Score | Tandem Score | TDA q-value | Volume | Spectrum Matches |
| --- | --- | --- | --- | --- |
| 16.5968 | 8.8579 | 0.0010 | 7.8346e+08 | 5 |

**Adduct Signal Fractions**

- **Unmodified**: 7.8346e+08

#### EGVFVSN(N)GTHWFVTQR Hex 6 HexNAc 5

**Mass:**  3850.6212

(9.236e-08)

**Charge States:** 3, 4

**Start Position**: 1079

**End Position**: 1095

| Profile Score | Tandem Score | TDA q-value | Volume | Spectrum Matches |
| --- | --- | --- | --- | --- |
| 17.0368 | 8.8359 | 0.0010 | 1.1226e+09 | 5 |

**Adduct Signal Fractions**

- **Unmodified**: 1.1226e+09

#### TQSLLIVNN(N)ATNVVIK Hex 3 HexNAc 3

**Mass:**  2821.3947

(-2.912e-07)

**Charge States:** 2, 3

**Start Position**: 101

**End Position**: 117

| Profile Score | Tandem Score | TDA q-value | Volume | Spectrum Matches |
| --- | --- | --- | --- | --- |
| 14.9506 | 8.8193 | 0.0010 | 1.2795e+10 | 6 |

**Adduct Signal Fractions**

- **Unmodified**: 1.2795e+10

#### EGVFVSN(N)GTHWFVTQR Hex 6 HexNAc 5 Neu5Ac 2

**Mass:**  4432.8141

(1.479e-06)

**Charge States:** 3, 4

**Start Position**: 1079

**End Position**: 1095

| Profile Score | Tandem Score | TDA q-value | Volume | Spectrum Matches |
| --- | --- | --- | --- | --- |
| 9.8771 | 8.7992 | 0.0010 | 1.6690e+09 | 7 |

**Adduct Signal Fractions**

- **Unmodified**: 1.6690e+09

#### N(N)FTTAPAIC(C)HDGK Fuc 1 Hex 3 HexNAc 6

**Mass:**  3283.3588

(6.110e-04)

**Start Position**: 1061

**End Position**: 1074

| Profile Score | Tandem Score | TDA q-value | Volume | Spectrum Matches |
| --- | --- | --- | --- | --- |
| 0.0000 | 8.7528 | 0.0010 | 0.0000e+00 | 1 |

Adduct:

No Chromatogram Found

#### NLN(N)ESLIDLQELGK Fuc 1 Hex 6 HexNAc 6 Neu5Ac 2

**Mass:**  4503.8811

(1.013e-06)

**Charge States:** 3, 4

**Start Position**: 1179

**End Position**: 1193

| Profile Score | Tandem Score | TDA q-value | Volume | Spectrum Matches |
| --- | --- | --- | --- | --- |
| 9.1216 | 8.6807 | 0.0010 | 1.4786e+08 | 3 |

**Adduct Signal Fractions**

- **Unmodified**: 1.4786e+08

#### DFGGFN(N)FSQILPDPSKPSK Fuc 1 Hex 4 HexNAc 5

**Mass:**  3890.6969

(9.595e-07)

**Charge States:** 3, 4

**Start Position**: 783

**End Position**: 802

| Profile Score | Tandem Score | TDA q-value | Volume | Spectrum Matches |
| --- | --- | --- | --- | --- |
| 11.6464 | 8.6412 | 0.0010 | 4.1250e+09 | 16 |

**Adduct Signal Fractions**

- **Unmodified**: 4.1250e+09

#### FPNITNLC(C)PFGEVFN(N)ATR Fuc 1 Hex 9 HexNAc 6 Neu5Ac 1

**Mass:**  5210.1205

(2.995e-07)

**Charge States:** 4

**Start Position**: 316

**End Position**: 334

| Profile Score | Tandem Score | TDA q-value | Volume | Spectrum Matches |
| --- | --- | --- | --- | --- |
| 14.0765 | 8.6348 | 0.0010 | 6.9329e+08 | 1 |

**Adduct Signal Fractions**

- **Unmodified**: 6.9329e+08

#### NLN(N)ESLIDLQELGK Fuc 1 Hex 5 HexNAc 6 Neu5Ac 1

**Mass:**  4050.7616

(1.822e-06)

**Charge States:** 3, 4

**Start Position**: 1179

**End Position**: 1193

| Profile Score | Tandem Score | TDA q-value | Volume | Spectrum Matches |
| --- | --- | --- | --- | --- |
| -5.9822 | 8.5979 | 0.0010 | 9.7364e+07 | 5 |

**Adduct Signal Fractions**

- **Unmodified**: 9.7364e+07

#### TPPIKDFGGFN(N)FSQILPDPSKPSKR Hex 5 HexNAc 2

**Mass:**  3988.8841

(2.928e-07)

**Charge States:** 4, 5

**Start Position**: 778

**End Position**: 803

| Profile Score | Tandem Score | TDA q-value | Volume | Spectrum Matches |
| --- | --- | --- | --- | --- |
| 13.1332 | 8.5400 | 0.0010 | 1.3862e+09 | 10 |

**Adduct Signal Fractions**

- **Unmodified**: 1.3862e+09

#### DFGGFN(N)FSQILPDPSKPSK Hex 4 HexNAc 2

**Mass:**  3134.3947

(3.191e-07)

**Charge States:** 3, 4

**Start Position**: 783

**End Position**: 802

| Profile Score | Tandem Score | TDA q-value | Volume | Spectrum Matches |
| --- | --- | --- | --- | --- |
| 14.1603 | 8.5316 | 0.0010 | 2.4054e+10 | 20 |

**Adduct Signal Fractions**

- **Unmodified**: 2.4054e+10

#### DFGGFN(N)FSQILPDPSKPSK Hex 4 HexNAc 4

**Mass:**  3540.5552

(-1.271e-08)

**Charge States:** 3, 4

**Start Position**: 783

**End Position**: 802

| Profile Score | Tandem Score | TDA q-value | Volume | Spectrum Matches |
| --- | --- | --- | --- | --- |
| 10.7679 | 8.5057 | 0.0010 | 4.5884e+09 | 12 |

**Adduct Signal Fractions**

- **Unmodified**: 4.5884e+09

#### DFGGFN(N)FSQILPDPSKPSK Fuc 1 Hex 5 HexNAc 3 Neu5Ac 1

**Mass:**  3936.6847

(7.295e-07)

**Charge States:** 3, 4

**Start Position**: 783

**End Position**: 802

| Profile Score | Tandem Score | TDA q-value | Volume | Spectrum Matches |
| --- | --- | --- | --- | --- |
| 12.3923 | 8.4304 | 0.0010 | 3.9638e+09 | 6 |

**Adduct Signal Fractions**

- **Unmodified**: 3.9638e+09

#### SSVLHSTQDLFLPFFSN(N)VTWFHAIHVSGTNGTK Fuc 1 Hex 8 HexNAc 7

**Mass:**  6537.8745

(8.808e-07)

**Charge States:** 5, 6, 7

**Start Position**: 32

**End Position**: 65

| Profile Score | Tandem Score | TDA q-value | Volume | Spectrum Matches |
| --- | --- | --- | --- | --- |
| 13.8804 | 8.3629 | 0.0010 | 9.9367e+09 | 2 |

**Adduct Signal Fractions**

- **Unmodified**: 9.9367e+09

#### SSVLHSTQDLFLPFFSNVTWFHAIHVSGTN(N)GTK Fuc 1 Hex 8 HexNAc 7

**Mass:**  6537.8745

(8.808e-07)

**Charge States:** 5, 6, 7

**Start Position**: 32

**End Position**: 65

| Profile Score | Tandem Score | TDA q-value | Volume | Spectrum Matches |
| --- | --- | --- | --- | --- |
| 13.8804 | 8.3629 | 0.0010 | 9.9367e+09 | 2 |

**Adduct Signal Fractions**

- **Unmodified**: 9.9367e+09

#### YNEN(N)GTITDAVDC(C)ALDPLSETK Hex 6 HexNAc 5

**Mass:**  4412.8102

(8.040e-07)

**Charge States:** 3, 4

**Start Position**: 266

**End Position**: 288

| Profile Score | Tandem Score | TDA q-value | Volume | Spectrum Matches |
| --- | --- | --- | --- | --- |
| 15.8184 | 8.2893 | 0.0010 | 1.7617e+09 | 5 |

**Adduct Signal Fractions**

- **Unmodified**: 1.7617e+09

#### SSVLHSTQDLFLPFFSNVTWFHAIHVSGTN(N)GTK Fuc 2 Hex 8 HexNAc 7

**Mass:**  6683.9386

(6.942e-07)

**Charge States:** 5, 6, 7

**Start Position**: 32

**End Position**: 65

| Profile Score | Tandem Score | TDA q-value | Volume | Spectrum Matches |
| --- | --- | --- | --- | --- |
| 13.9220 | 8.2311 | 0.0010 | 8.1072e+09 | 2 |

**Adduct Signal Fractions**

- **Unmodified**: 8.1072e+09

#### TPPIKDFGGFN(N)FSQILPDPSKPSK Hex 5 HexNAc 3

**Mass:**  4035.8605

(6.057e-07)

**Charge States:** 3, 4, 5

**Start Position**: 778

**End Position**: 802

| Profile Score | Tandem Score | TDA q-value | Volume | Spectrum Matches |
| --- | --- | --- | --- | --- |
| 12.7027 | 8.2062 | 0.0010 | 1.7864e+09 | 7 |

**Adduct Signal Fractions**

- **Unmodified**: 1.7864e+09

#### DFGGFN(N)FSQILPDPSKPSK Fuc 2 Hex 5 HexNAc 4

**Mass:**  3994.7263

(5.379e-07)

**Charge States:** 3, 4

**Start Position**: 783

**End Position**: 802

| Profile Score | Tandem Score | TDA q-value | Volume | Spectrum Matches |
| --- | --- | --- | --- | --- |
| 14.0544 | 8.1797 | 0.0010 | 4.8282e+08 | 6 |

**Adduct Signal Fractions**

- **Unmodified**: 4.8282e+08

#### N(N)FTTAPAIC(C)HDGK Hex 4 HexNAc 4

**Mass:**  2891.1909

(1.611e-06)

**Charge States:** 3

**Start Position**: 1061

**End Position**: 1074

| Profile Score | Tandem Score | TDA q-value | Volume | Spectrum Matches |
| --- | --- | --- | --- | --- |
| 10.9258 | 8.1797 | 0.0010 | 4.0138e+09 | 6 |

**Adduct Signal Fractions**

- **Unmodified**: 4.0138e+09

#### N(N)FTTAPAIC(C)HDGK Fuc 1 Hex 5 HexNAc 6 Neu5Ac 2

**Mass:**  4187.6521

(3.452e-07)

**Charge States:** 3, 4

**Start Position**: 1061

**End Position**: 1074

| Profile Score | Tandem Score | TDA q-value | Volume | Spectrum Matches |
| --- | --- | --- | --- | --- |
| 11.9808 | 8.1196 | 0.0010 | 1.6449e+08 | 1 |

**Adduct Signal Fractions**

- **Unmodified**: 1.6449e+08

#### YNEN(N)GTITDAVDC(C)ALDPLSETK Fuc 1 Hex 6 HexNAc 5

**Mass:**  4558.8673

(-1.501e-08)

**Charge States:** 3, 4

**Start Position**: 266

**End Position**: 288

| Profile Score | Tandem Score | TDA q-value | Volume | Spectrum Matches |
| --- | --- | --- | --- | --- |
| 16.1872 | 8.1101 | 0.0010 | 1.6231e+10 | 5 |

**Adduct Signal Fractions**

- **Unmodified**: 1.6231e+10

#### TQSLLIVNN(N)ATNVVIK Hex 4 HexNAc 5

**Mass:**  3389.6071

(3.385e-07)

**Charge States:** 3, 4

**Start Position**: 101

**End Position**: 117

| Profile Score | Tandem Score | TDA q-value | Volume | Spectrum Matches |
| --- | --- | --- | --- | --- |
| -4.9726 | 8.0553 | 0.0010 | 3.1406e+09 | 13 |

**Adduct Signal Fractions**

- **Unmodified**: 3.1406e+09

#### EGVFVSN(N)GTHWFVTQR Fuc 1 Hex 5 HexNAc 4 Neu5Ac 2

**Mass:**  4213.7375

(2.968e-07)

**Charge States:** 3, 4

**Start Position**: 1079

**End Position**: 1095

| Profile Score | Tandem Score | TDA q-value | Volume | Spectrum Matches |
| --- | --- | --- | --- | --- |
| 13.3284 | 8.0154 | 0.0010 | 6.2607e+08 | 5 |

**Adduct Signal Fractions**

- **Unmodified**: 6.2607e+08

#### DFGGFN(N)FSQILPDPSKPSKR Hex 8 HexNAc 2

**Mass:**  3938.7104

(4.964e-08)

**Charge States:** 4

**Start Position**: 783

**End Position**: 803

| Profile Score | Tandem Score | TDA q-value | Volume | Spectrum Matches |
| --- | --- | --- | --- | --- |
| 13.5795 | 7.9926 | 0.0010 | 5.0604e+08 | 2 |

**Adduct Signal Fractions**

- **Unmodified**: 5.0604e+08

#### N(N)FTTAPAIC(C)HDGK Hex 6 HexNAc 5

**Mass:**  3418.3750

(8.252e-08)

**Charge States:** 3, 4

**Start Position**: 1061

**End Position**: 1074

| Profile Score | Tandem Score | TDA q-value | Volume | Spectrum Matches |
| --- | --- | --- | --- | --- |
| 15.6630 | 7.9782 | 0.0010 | 5.2693e+08 | 6 |

**Adduct Signal Fractions**

- **Unmodified**: 5.2693e+08

#### DFGGFN(N)FSQILPDPSKPSK Fuc 3 Hex 4 HexNAc 4

**Mass:**  3978.7111

(-4.512e-06)

**Charge States:** 4

**Start Position**: 783

**End Position**: 802

| Profile Score | Tandem Score | TDA q-value | Volume | Spectrum Matches |
| --- | --- | --- | --- | --- |
| -18.0099 | 7.9527 | 0.0010 | 1.0975e+07 | 2 |

**Adduct Signal Fractions**

- **Unmodified**: 1.0975e+07

#### TQSLLIVNN(N)ATNVVIK Fuc 2 Hex 4 HexNAc 3

**Mass:**  3275.5642

(3.711e-08)

**Charge States:** 3

**Start Position**: 101

**End Position**: 117

| Profile Score | Tandem Score | TDA q-value | Volume | Spectrum Matches |
| --- | --- | --- | --- | --- |
| 10.7376 | 7.9493 | 0.0010 | 9.4636e+08 | 6 |

**Adduct Signal Fractions**

- **Unmodified**: 9.4636e+08

#### TQSLLIVNN(N)ATNVVIK Fuc 2 Hex 5 HexNAc 4

**Mass:**  3640.6928

(-4.269e-07)

**Charge States:** 3

**Start Position**: 101

**End Position**: 117

| Profile Score | Tandem Score | TDA q-value | Volume | Spectrum Matches |
| --- | --- | --- | --- | --- |
| 13.2149 | 7.9491 | 0.0010 | 2.0848e+09 | 3 |

**Adduct Signal Fractions**

- **Unmodified**: 2.0848e+09

#### DFGGFN(N)FSQILPDPSKPSKR Fuc 1 Hex 3 HexNAc 5

**Mass:**  3883.7423

(8.214e-07)

**Charge States:** 3, 4

**Start Position**: 783

**End Position**: 803

| Profile Score | Tandem Score | TDA q-value | Volume | Spectrum Matches |
| --- | --- | --- | --- | --- |
| 13.1713 | 7.9237 | 0.0010 | 3.1300e+08 | 2 |

**Adduct Signal Fractions**

- **Unmodified**: 3.1300e+08

#### DFGGFN(N)FSQILPDPSKPSKR Hex 3 HexNAc 3

**Mass:**  3331.5290

(6.508e-07)

**Charge States:** 4

**Start Position**: 783

**End Position**: 803

| Profile Score | Tandem Score | TDA q-value | Volume | Spectrum Matches |
| --- | --- | --- | --- | --- |
| 12.4086 | 7.9043 | 0.0010 | 1.7937e+08 | 1 |

**Adduct Signal Fractions**

- **Unmodified**: 1.7937e+08

#### DFGGFN(N)FSQILPDPSKPSKR Fuc 1 Hex 4 HexNAc 4 Neu5Ac 1

**Mass:**  4133.8138

(1.019e-06)

**Charge States:** 4

**Start Position**: 783

**End Position**: 803

| Profile Score | Tandem Score | TDA q-value | Volume | Spectrum Matches |
| --- | --- | --- | --- | --- |
| 13.1189 | 7.8916 | 0.0010 | 5.7908e+08 | 2 |

**Adduct Signal Fractions**

- **Unmodified**: 5.7908e+08

#### SSVLHSTQDLFLPFFSNVTWFHAIHVSGTN(N)GTK @sulfate 1 Fuc 1 Hex 9 HexNAc 7 Neu5Ac 1

**Mass:**  7070.9791

(8.694e-07)

**Charge States:** 5, 6, 7

**Start Position**: 32

**End Position**: 65

| Profile Score | Tandem Score | TDA q-value | Volume | Spectrum Matches |
| --- | --- | --- | --- | --- |
| 14.2392 | 7.8810 | 0.0010 | 3.1107e+09 | 2 |

**Adduct Signal Fractions**

- **Unmodified**: 3.1107e+09

#### N(N)FTTAPAIC(C)HDGK Hex 5 HexNAc 5

**Mass:**  3256.3230

(3.118e-07)

**Charge States:** 3, 4

**Start Position**: 1061

**End Position**: 1074

| Profile Score | Tandem Score | TDA q-value | Volume | Spectrum Matches |
| --- | --- | --- | --- | --- |
| 10.0878 | 7.8801 | 0.0010 | 5.2371e+08 | 9 |

**Adduct Signal Fractions**

- **Unmodified**: 5.2371e+08

#### NLN(N)ESLIDLQELGK Hex 5 HexNAc 6 Neu5Ac 4

**Mass:**  4794.9821

(3.562e-03)

**Charge States:** 4

**Start Position**: 1179

**End Position**: 1193

| Profile Score | Tandem Score | TDA q-value | Volume | Spectrum Matches |
| --- | --- | --- | --- | --- |
| 9.6045 | 7.8291 | 0.0010 | 4.4016e+08 | 4 |

**Adduct Signal Fractions**

- **Unmodified**: 4.4016e+08

#### FPNITNLC(C)PFGEVFN(N)ATR Fuc 1 Hex 8 HexNAc 5

**Mass:**  4553.8963

(5.397e-07)

**Charge States:** 3, 4

**Start Position**: 316

**End Position**: 334

| Profile Score | Tandem Score | TDA q-value | Volume | Spectrum Matches |
| --- | --- | --- | --- | --- |
| 13.5823 | 7.8214 | 0.0010 | 6.6309e+08 | 4 |

**Adduct Signal Fractions**

- **Unmodified**: 6.6309e+08

#### DFGGFN(N)FSQILPDPSKPSKR Fuc 1 Hex 3 HexNAc 4

**Mass:**  3680.6629

(4.002e-07)

**Charge States:** 3, 4

**Start Position**: 783

**End Position**: 803

| Profile Score | Tandem Score | TDA q-value | Volume | Spectrum Matches |
| --- | --- | --- | --- | --- |
| 14.9736 | 7.8063 | 0.0010 | 6.1428e+08 | 5 |

**Adduct Signal Fractions**

- **Unmodified**: 6.1428e+08

#### VYSSANN(N)C(C)TFEYVSQPFLMDLEGK Fuc 1 Hex 5 HexNAc 4

**Mass:**  4566.8974

(-1.227e-07)

**Charge States:** 3, 4

**Start Position**: 146

**End Position**: 170

| Profile Score | Tandem Score | TDA q-value | Volume | Spectrum Matches |
| --- | --- | --- | --- | --- |
| 17.6100 | 7.7911 | 0.0010 | 4.2091e+09 | 7 |

**Adduct Signal Fractions**

- **Unmodified**: 4.2091e+09

#### TQSLLIVNN(N)ATNVVIK Hex 4 HexNAc 4 Neu5Ac 1

**Mass:**  3477.6263

(2.949e-07)

**Charge States:** 3, 4

**Start Position**: 101

**End Position**: 117

| Profile Score | Tandem Score | TDA q-value | Volume | Spectrum Matches |
| --- | --- | --- | --- | --- |
| 13.3562 | 7.7547 | 0.0010 | 1.2892e+10 | 12 |

**Adduct Signal Fractions**

- **Unmodified**: 1.2892e+10

#### NLN(N)ESLIDLQELGK Hex 5 HexNAc 2

**Mass:**  2801.2612

(3.159e-07)

**Charge States:** 2, 3

**Start Position**: 1179

**End Position**: 1193

| Profile Score | Tandem Score | TDA q-value | Volume | Spectrum Matches |
| --- | --- | --- | --- | --- |
| 12.9787 | 7.7444 | 0.0010 | 2.8468e+08 | 4 |

**Adduct Signal Fractions**

- **Unmodified**: 2.8468e+08

#### EGVFVSN(N)GTHWFVTQR Hex 5 HexNAc 5

**Mass:**  3688.5674

(-3.112e-09)

**Charge States:** 3, 4

**Start Position**: 1079

**End Position**: 1095

| Profile Score | Tandem Score | TDA q-value | Volume | Spectrum Matches |
| --- | --- | --- | --- | --- |
| 13.5665 | 7.7442 | 0.0010 | 1.1511e+09 | 7 |

**Adduct Signal Fractions**

- **Unmodified**: 1.1511e+09

#### VYSSANN(N)C(C)TFEYVSQPFLMDLEGK Hex 4 HexNAc 5

**Mass:**  4461.8658

(2.603e-07)

**Charge States:** 3, 4

**Start Position**: 146

**End Position**: 170

| Profile Score | Tandem Score | TDA q-value | Volume | Spectrum Matches |
| --- | --- | --- | --- | --- |
| 15.6499 | 7.7277 | 0.0010 | 3.1926e+09 | 7 |

**Adduct Signal Fractions**

- **Unmodified**: 3.1926e+09

#### VYSSANN(N)C(C)TFEYVSQPFLMDLEGK Fuc 1 Hex 4 HexNAc 5

**Mass:**  4607.9230

(-3.575e-07)

**Charge States:** 3, 4

**Start Position**: 146

**End Position**: 170

| Profile Score | Tandem Score | TDA q-value | Volume | Spectrum Matches |
| --- | --- | --- | --- | --- |
| 16.3694 | 7.6688 | 0.0010 | 8.6161e+09 | 11 |

**Adduct Signal Fractions**

- **NH3**: 1.6585e+08
- **Unmodified**: 8.4502e+09

#### SSVLHSTQDLFLPFFSNVTWFHAIHVSGTN(N)GTK Fuc 2 Hex 9 HexNAc 7

**Mass:**  6845.9834

(3.007e-07)

**Charge States:** 5, 6, 7

**Start Position**: 32

**End Position**: 65

| Profile Score | Tandem Score | TDA q-value | Volume | Spectrum Matches |
| --- | --- | --- | --- | --- |
| 14.0459 | 7.6377 | 0.0010 | 6.1128e+09 | 6 |

**Adduct Signal Fractions**

- **Unmodified**: 6.1128e+09

#### N(N)FTTAPAIC(C)HDGK Fuc 1 Hex 5 HexNAc 4

**Mass:**  3201.2975

(6.241e-04)

**Start Position**: 1061

**End Position**: 1074

| Profile Score | Tandem Score | TDA q-value | Volume | Spectrum Matches |
| --- | --- | --- | --- | --- |
| 0.0000 | 7.5801 | 0.0010 | 0.0000e+00 | 1 |

Adduct:

No Chromatogram Found

#### DFGGFN(N)FSQILPDPSKPSKR Fuc 1 Hex 3 HexNAc 3

**Mass:**  3477.5535

(4.888e-03)

**Charge States:** 4

**Start Position**: 783

**End Position**: 803

| Profile Score | Tandem Score | TDA q-value | Volume | Spectrum Matches |
| --- | --- | --- | --- | --- |
| 8.8038 | 7.5470 | 0.0010 | 1.1074e+08 | 2 |

**Adduct Signal Fractions**

- **NH3**: 1.1074e+08

#### FPNITNLC(C)PFGEVFN(N)ATR Fuc 2 Hex 7 HexNAc 7

**Mass:**  4944.0542

(2.472e-07)

**Charge States:** 4

**Start Position**: 316

**End Position**: 334

| Profile Score | Tandem Score | TDA q-value | Volume | Spectrum Matches |
| --- | --- | --- | --- | --- |
| 13.9090 | 7.5428 | 0.0010 | 7.5503e+08 | 2 |

**Adduct Signal Fractions**

- **Unmodified**: 7.5503e+08

#### EGVFVSN(N)GTHWFVTQR Fuc 1 Hex 6 HexNAc 3 Neu5Ac 1

**Mass:**  3881.6162

(-5.023e-07)

**Charge States:** 3, 4

**Start Position**: 1079

**End Position**: 1095

| Profile Score | Tandem Score | TDA q-value | Volume | Spectrum Matches |
| --- | --- | --- | --- | --- |
| 16.4170 | 7.5221 | 0.0010 | 3.1790e+09 | 7 |

**Adduct Signal Fractions**

- **Unmodified**: 3.1790e+09

#### N(N)FTTAPAIC(C)HDGK Hex 3 HexNAc 5

**Mass:**  2933.2188

(6.842e-04)

**Start Position**: 1061

**End Position**: 1074

| Profile Score | Tandem Score | TDA q-value | Volume | Spectrum Matches |
| --- | --- | --- | --- | --- |
| 0.0000 | 7.4695 | 0.0010 | 0.0000e+00 | 3 |

Adduct:

No Chromatogram Found

#### DFGGFN(N)FSQILPDPSKPSK Fuc 1 Hex 5 HexNAc 4 Neu5Ac 2

**Mass:**  4430.8578

(1.296e-07)

**Charge States:** 3, 4

**Start Position**: 783

**End Position**: 802

| Profile Score | Tandem Score | TDA q-value | Volume | Spectrum Matches |
| --- | --- | --- | --- | --- |
| 14.4174 | 7.4591 | 0.0010 | 4.5137e+08 | 2 |

**Adduct Signal Fractions**

- **Unmodified**: 4.5137e+08

#### DFGGFN(N)FSQILPDPSKPSK Hex 6 HexNAc 2

**Mass:**  3482.4941

(8.697e-04)

**Start Position**: 783

**End Position**: 802

| Profile Score | Tandem Score | TDA q-value | Volume | Spectrum Matches |
| --- | --- | --- | --- | --- |
| 0.0000 | 7.3865 | 0.0010 | 0.0000e+00 | 6 |

Adduct:

No Chromatogram Found

#### YNEN(N)GTITDAVDC(C)ALDPLSETK Fuc 1 Hex 6 HexNAc 5 Neu5Ac 1

**Mass:**  4849.9597

(7.311e-07)

**Charge States:** 4

**Start Position**: 266

**End Position**: 288

| Profile Score | Tandem Score | TDA q-value | Volume | Spectrum Matches |
| --- | --- | --- | --- | --- |
| 16.1969 | 7.3337 | 0.0010 | 2.6996e+09 | 2 |

**Adduct Signal Fractions**

- **Unmodified**: 2.6996e+09

#### EGVFVSN(N)GTHWFVTQR Fuc 1 Hex 6 HexNAc 5

**Mass:**  3996.6744

(7.520e-04)

**Charge States:** 3, 4

**Start Position**: 1079

**End Position**: 1095

| Profile Score | Tandem Score | TDA q-value | Volume | Spectrum Matches |
| --- | --- | --- | --- | --- |
| -4.5921 | 7.2292 | 0.0010 | 1.4731e+09 | 14 |

**Adduct Signal Fractions**

- **Unmodified**: 1.4731e+09

#### YNEN(N)GTITDAVDC(C)ALDPLSETK Fuc 1 Hex 3 HexNAc 5

**Mass:**  4072.7098

(3.910e-07)

**Charge States:** 3, 4

**Start Position**: 266

**End Position**: 288

| Profile Score | Tandem Score | TDA q-value | Volume | Spectrum Matches |
| --- | --- | --- | --- | --- |
| 15.3370 | 7.2186 | 0.0010 | -4.5809e+08 | 12 |

**Adduct Signal Fractions**

- **Unmodified**: 3.8197e+10

#### YNEN(N)GTITDAVDC(C)ALDPLSETK Fuc 1 Hex 3 HexNAc 4

**Mass:**  3869.6297

(1.637e-08)

**Charge States:** 3, 4

**Start Position**: 266

**End Position**: 288

| Profile Score | Tandem Score | TDA q-value | Volume | Spectrum Matches |
| --- | --- | --- | --- | --- |
| -1.4026 | 7.1913 | 0.0010 | 4.0629e+09 | 7 |

**Adduct Signal Fractions**

- **Unmodified**: 4.0629e+09

#### N(N)FTTAPAIC(C)HDGK Fuc 2 Hex 4 HexNAc 4

**Mass:**  3183.3034

(2.707e-07)

**Charge States:** 3

**Start Position**: 1061

**End Position**: 1074

| Profile Score | Tandem Score | TDA q-value | Volume | Spectrum Matches |
| --- | --- | --- | --- | --- |
| -3.4127 | 7.0274 | 0.0010 | 2.2944e+08 | 6 |

**Adduct Signal Fractions**

- **Unmodified**: 2.2944e+08

#### N(N)FTTAPAIC(C)HDGK Fuc 1 Hex 3 HexNAc 7

**Mass:**  3485.4380

(2.884e-04)

**Start Position**: 1061

**End Position**: 1074

| Profile Score | Tandem Score | TDA q-value | Volume | Spectrum Matches |
| --- | --- | --- | --- | --- |
| 0.0000 | 7.0033 | 0.0010 | 0.0000e+00 | 1 |

Adduct:

No Chromatogram Found

#### DFGGFN(N)FSQILPDPSKPSKR Hex 6 HexNAc 3 Neu5Ac 1

**Mass:**  4108.7819

(9.259e-07)

**Charge States:** 4, 5

**Start Position**: 783

**End Position**: 803

| Profile Score | Tandem Score | TDA q-value | Volume | Spectrum Matches |
| --- | --- | --- | --- | --- |
| -4.0585 | 6.8770 | 0.0010 | 2.2188e+08 | 4 |

**Adduct Signal Fractions**

- **Unmodified**: 2.2188e+08

#### EGVFVSN(N)GTHWFVTQR Fuc 1 Hex 6 HexNAc 5

**Mass:**  3998.6804

(5.010e-04)

**Start Position**: 1079

**End Position**: 1095

| Profile Score | Tandem Score | TDA q-value | Volume | Spectrum Matches |
| --- | --- | --- | --- | --- |
| 0.0000 | 6.8728 | 0.0010 | 0.0000e+00 | 2 |

Adduct:

No Chromatogram Found

#### EGVFVSN(N)GTHWFVTQR Fuc 1 Hex 6 HexNAc 6 Neu5Ac 2

**Mass:**  4782.9445

(2.083e-04)

**Start Position**: 1079

**End Position**: 1095

| Profile Score | Tandem Score | TDA q-value | Volume | Spectrum Matches |
| --- | --- | --- | --- | --- |
| 0.0000 | 6.8699 | 0.0010 | 0.0000e+00 | 1 |

Adduct:

No Chromatogram Found

#### DFGGFN(N)FSQILPDPSKPSKR Hex 4 HexNAc 3

**Mass:**  3493.5786

(7.610e-07)

**Charge States:** 4

**Start Position**: 783

**End Position**: 803

| Profile Score | Tandem Score | TDA q-value | Volume | Spectrum Matches |
| --- | --- | --- | --- | --- |
| 10.7705 | 6.8395 | 0.0010 | 6.8901e+08 | 3 |

**Adduct Signal Fractions**

- **Unmodified**: 6.8901e+08

#### DFGGFN(N)FSQILPDPSKPSKR Fuc 1 Hex 4 HexNAc 4

**Mass:**  3842.7159

(2.863e-07)

**Charge States:** 4

**Start Position**: 783

**End Position**: 803

| Profile Score | Tandem Score | TDA q-value | Volume | Spectrum Matches |
| --- | --- | --- | --- | --- |
| 11.1968 | 6.8270 | 0.0010 | 2.5185e+08 | 5 |

**Adduct Signal Fractions**

- **Unmodified**: 2.5185e+08

#### VYSSANN(N)C(C)TFEYVSQPFLMDLEGK Hex 5 HexNAc 2

**Mass:**  4014.6763

(3.793e-07)

**Charge States:** 3, 4

**Start Position**: 146

**End Position**: 170

| Profile Score | Tandem Score | TDA q-value | Volume | Spectrum Matches |
| --- | --- | --- | --- | --- |
| 18.3245 | 6.7820 | 0.0010 | 2.3136e+10 | 9 |

**Adduct Signal Fractions**

- **Unmodified**: 2.3136e+10

#### VYSSANN(N)C(C)TFEYVSQPFLMDLEGK Fuc 1 Hex 3 HexNAc 5

**Mass:**  4445.8699

(-7.838e-08)

**Charge States:** 3, 4

**Start Position**: 146

**End Position**: 170

| Profile Score | Tandem Score | TDA q-value | Volume | Spectrum Matches |
| --- | --- | --- | --- | --- |
| 16.0201 | 6.7714 | 0.0010 | 6.3945e+09 | 8 |

**Adduct Signal Fractions**

- **NH3**: 8.6475e+07
- **Unmodified**: 6.3080e+09

#### TQSLLIVNN(N)ATNVVIK Hex 5 HexNAc 4 Neu5Ac 1

**Mass:**  3639.6761

(4.394e-07)

**Charge States:** 3, 4

**Start Position**: 101

**End Position**: 117

| Profile Score | Tandem Score | TDA q-value | Volume | Spectrum Matches |
| --- | --- | --- | --- | --- |
| 13.7855 | 6.7282 | 0.0010 | 6.1912e+09 | 11 |

**Adduct Signal Fractions**

- **Unmodified**: 6.1912e+09

#### EGVFVSN(N)GTHWFVTQR Hex 5 HexNAc 6 Neu5Ac 2

**Mass:**  4474.8451

(2.252e-04)

**Start Position**: 1079

**End Position**: 1095

| Profile Score | Tandem Score | TDA q-value | Volume | Spectrum Matches |
| --- | --- | --- | --- | --- |
| 0.0000 | 6.6947 | 0.0010 | 0.0000e+00 | 3 |

Adduct:

No Chromatogram Found

#### VYSSANN(N)C(C)TFEYVSQPFLMDLEGK Fuc 1 Hex 5 HexNAc 5

**Mass:**  4769.9768

(1.225e-07)

**Charge States:** 3, 4

**Start Position**: 146

**End Position**: 170

| Profile Score | Tandem Score | TDA q-value | Volume | Spectrum Matches |
| --- | --- | --- | --- | --- |
| 15.6147 | 6.6812 | 0.0010 | 2.3747e+09 | 5 |

**Adduct Signal Fractions**

- **Unmodified**: 2.3747e+09

#### SSVLHSTQDLFLPFFSNVTWFHAIHVSGTN(N)GTK Fuc 2 Hex 9 HexNAc 7 Neu5Ac 1

**Mass:**  7137.0806

(1.252e-07)

**Charge States:** 5, 6, 7

**Start Position**: 32

**End Position**: 65

| Profile Score | Tandem Score | TDA q-value | Volume | Spectrum Matches |
| --- | --- | --- | --- | --- |
| 14.9194 | 6.6758 | 0.0010 | 4.4918e+09 | 4 |

**Adduct Signal Fractions**

- **Unmodified**: 4.4918e+09

#### SSVLHSTQDLFLPFFSN(N)VTWFHAIHVSGTNGTK Fuc 2 Hex 9 HexNAc 7 Neu5Ac 1

**Mass:**  7137.0806

(1.252e-07)

**Charge States:** 5, 6, 7

**Start Position**: 32

**End Position**: 65

| Profile Score | Tandem Score | TDA q-value | Volume | Spectrum Matches |
| --- | --- | --- | --- | --- |
| 14.9194 | 6.6758 | 0.0010 | 4.4918e+09 | 4 |

**Adduct Signal Fractions**

- **Unmodified**: 4.4918e+09

#### TQSLLIVNN(N)ATNVVIK Fuc 1 Hex 3 HexNAc 5 Neu5Ac 1

**Mass:**  3664.6854

(2.762e-04)

**Charge States:** 3

**Start Position**: 101

**End Position**: 117

| Profile Score | Tandem Score | TDA q-value | Volume | Spectrum Matches |
| --- | --- | --- | --- | --- |
| 9.1865 | 6.6581 | 0.0010 | 3.2266e+07 | 3 |

**Adduct Signal Fractions**

- **Unmodified**: 3.2266e+07

#### EGVFVSN(N)GTHWFVTQR Fuc 1 Hex 5 HexNAc 4

**Mass:**  3632.5446

(2.750e-04)

**Start Position**: 1079

**End Position**: 1095

| Profile Score | Tandem Score | TDA q-value | Volume | Spectrum Matches |
| --- | --- | --- | --- | --- |
| 0.0000 | 6.6520 | 0.0010 | 0.0000e+00 | 2 |

Adduct:

No Chromatogram Found

#### VYSSANN(N)C(C)TFEYVSQPFLMDLEGK Hex 5 HexNAc 4 Neu5Ac 1

**Mass:**  4711.9349

(-3.397e-07)

**Charge States:** 3, 4

**Start Position**: 146

**End Position**: 170

| Profile Score | Tandem Score | TDA q-value | Volume | Spectrum Matches |
| --- | --- | --- | --- | --- |
| 13.6845 | 6.6491 | 0.0010 | 2.4723e+09 | 6 |

**Adduct Signal Fractions**

- **Unmodified**: 2.4723e+09

#### TPPIKDFGGFN(N)FSQILPDPSKPSK Hex 6 HexNAc 3 Neu5Ac 1

**Mass:**  4489.0103

(2.332e-07)

**Charge States:** 4, 5

**Start Position**: 778

**End Position**: 802

| Profile Score | Tandem Score | TDA q-value | Volume | Spectrum Matches |
| --- | --- | --- | --- | --- |
| 12.8911 | 6.6267 | 0.0010 | 7.8991e+08 | 2 |

**Adduct Signal Fractions**

- **Unmodified**: 7.8991e+08

#### N(N)FTTAPAIC(C)HDGK Fuc 1 Hex 6 HexNAc 5 Neu5Ac 2

**Mass:**  4146.6254

(-1.340e-07)

**Charge States:** 4

**Start Position**: 1061

**End Position**: 1074

| Profile Score | Tandem Score | TDA q-value | Volume | Spectrum Matches |
| --- | --- | --- | --- | --- |
| 11.8608 | 6.6195 | 0.0010 | 3.7556e+08 | 2 |

**Adduct Signal Fractions**

- **Unmodified**: 3.7556e+08

#### EGVFVSN(N)GTHWFVTQR Hex 6 HexNAc 5 Neu5Ac 2

**Mass:**  4434.8190

(4.527e-04)

**Start Position**: 1079

**End Position**: 1095

| Profile Score | Tandem Score | TDA q-value | Volume | Spectrum Matches |
| --- | --- | --- | --- | --- |
| 0.0000 | 6.6158 | 0.0010 | 0.0000e+00 | 4 |

Adduct:

No Chromatogram Found

#### EGVFVSN(N)GTHWFVTQR Fuc 1 Hex 6 HexNAc 3

**Mass:**  3590.5179

(2.625e-07)

**Charge States:** 3, 4

**Start Position**: 1079

**End Position**: 1095

| Profile Score | Tandem Score | TDA q-value | Volume | Spectrum Matches |
| --- | --- | --- | --- | --- |
| 15.2290 | 6.4738 | 0.0010 | 1.8099e+09 | 7 |

**Adduct Signal Fractions**

- **Unmodified**: 1.8099e+09

#### EGVFVSN(N)GTHWFVTQR Fuc 2 Hex 5 HexNAc 3

**Mass:**  3574.5212

(4.756e-03)

**Charge States:** 3

**Start Position**: 1079

**End Position**: 1095

| Profile Score | Tandem Score | TDA q-value | Volume | Spectrum Matches |
| --- | --- | --- | --- | --- |
| 10.5872 | 6.4237 | 0.0010 | 1.7670e+08 | 4 |

**Adduct Signal Fractions**

- **NH3**: 2.0569e+07
- **Unmodified**: 1.5613e+08

#### NLN(N)ESLIDLQELGK Fuc 1 Hex 6 HexNAc 7 Neu5Ac 3

**Mass:**  4998.0601

(8.938e-07)

**Charge States:** 4

**Start Position**: 1179

**End Position**: 1193

| Profile Score | Tandem Score | TDA q-value | Volume | Spectrum Matches |
| --- | --- | --- | --- | --- |
| 12.3168 | 6.3279 | 0.0010 | 2.0044e+08 | 2 |

**Adduct Signal Fractions**

- **Unmodified**: 2.0044e+08

#### DFGGFN(N)FSQILPDPSKPSK Hex 6 HexNAc 4

**Mass:**  3864.6645

(2.699e-07)

**Charge States:** 3, 4

**Start Position**: 783

**End Position**: 802

| Profile Score | Tandem Score | TDA q-value | Volume | Spectrum Matches |
| --- | --- | --- | --- | --- |
| 8.0430 | 6.2871 | 0.0010 | 8.7730e+07 | 3 |

**Adduct Signal Fractions**

- **Unmodified**: 8.7730e+07

#### YNEN(N)GTITDAVDC(C)ALDPLSETK Fuc 1 Hex 5 HexNAc 4 Neu5Ac 1

**Mass:**  4484.8352

(5.733e-07)

**Charge States:** 3, 4

**Start Position**: 266

**End Position**: 288

| Profile Score | Tandem Score | TDA q-value | Volume | Spectrum Matches |
| --- | --- | --- | --- | --- |
| 12.7056 | 6.2523 | 0.0010 | 4.7630e+08 | 2 |

**Adduct Signal Fractions**

- **Unmodified**: 4.7630e+08

#### DFGGFN(N)FSQILPDPSKPSK Hex 3 HexNAc 5

**Mass:**  3581.5784

(2.006e-07)

**Charge States:** 3, 4

**Start Position**: 783

**End Position**: 802

| Profile Score | Tandem Score | TDA q-value | Volume | Spectrum Matches |
| --- | --- | --- | --- | --- |
| 12.9788 | 6.2511 | 0.0010 | 6.4728e+09 | 2 |

**Adduct Signal Fractions**

- **Unmodified**: 6.4728e+09

#### DFGGFN(N)FSQILPDPSKPSK Hex 6 HexNAc 3 Neu5Ac 1

**Mass:**  3955.6950

(5.085e-04)

**Start Position**: 783

**End Position**: 802

| Profile Score | Tandem Score | TDA q-value | Volume | Spectrum Matches |
| --- | --- | --- | --- | --- |
| 0.0000 | 6.2300 | 0.0010 | 0.0000e+00 | 2 |

Adduct:

No Chromatogram Found

#### N(N)FTTAPAIC(C)HDGK Fuc 3 Hex 5 HexNAc 6

**Mass:**  3897.5580

(-4.444e-06)

**Charge States:** 3, 4

**Start Position**: 1061

**End Position**: 1074

| Profile Score | Tandem Score | TDA q-value | Volume | Spectrum Matches |
| --- | --- | --- | --- | --- |
| -20.0421 | 6.2245 | 0.0010 | 9.9663e+06 | 1 |

**Adduct Signal Fractions**

- **Unmodified**: 9.9663e+06

#### DFGGFN(N)FSQILPDPSKPSK Hex 5 HexNAc 4

**Mass:**  3704.6142

(5.428e-04)

**Start Position**: 783

**End Position**: 802

| Profile Score | Tandem Score | TDA q-value | Volume | Spectrum Matches |
| --- | --- | --- | --- | --- |
| 0.0000 | 6.2228 | 0.0010 | 0.0000e+00 | 4 |

Adduct:

No Chromatogram Found

#### DFGGFN(N)FSQILPDPSKPSKR Fuc 2 Hex 5 HexNAc 3

**Mass:**  3947.7275

(-4.625e-06)

**Charge States:** 4

**Start Position**: 783

**End Position**: 803

| Profile Score | Tandem Score | TDA q-value | Volume | Spectrum Matches |
| --- | --- | --- | --- | --- |
| -21.8225 | 6.2163 | 0.0010 | 1.7853e+07 | 1 |

**Adduct Signal Fractions**

- **Unmodified**: 1.7853e+07

#### FPNITNLC(C)PFGEVFN(N)ATR Fuc 2 Hex 7 HexNAc 8 Neu5Ac 1

**Mass:**  5438.2331

(6.970e-07)

**Charge States:** 4

**Start Position**: 316

**End Position**: 334

| Profile Score | Tandem Score | TDA q-value | Volume | Spectrum Matches |
| --- | --- | --- | --- | --- |
| 14.1436 | 6.1137 | 0.0010 | -6.7417e+08 | 5 |

**Adduct Signal Fractions**

- **Unmodified**: 3.6208e+09

#### DFGGFN(N)FSQILPDPSKPSKRSFIEDLLFNK Hex 7 HexNAc 2

**Mass:**  5002.2966

(3.815e-03)

**Start Position**: 783

**End Position**: 813

| Profile Score | Tandem Score | TDA q-value | Volume | Spectrum Matches |
| --- | --- | --- | --- | --- |
| 0.0000 | 6.1017 | 0.0010 | 0.0000e+00 | 1 |

Adduct:

No Chromatogram Found

#### QC(C)VN(N)LTTR Fuc 1 Hex 3 HexNAc 4

**Mass:**  2435.0241

(-7.326e-07)

**Charge States:** 2, 3

**Start Position**: 1

**End Position**: 9

| Profile Score | Tandem Score | TDA q-value | Volume | Spectrum Matches |
| --- | --- | --- | --- | --- |
| 13.9743 | 5.9995 | 0.0110 | 1.7379e+08 | 2 |

**Adduct Signal Fractions**

- **Unmodified**: 1.7379e+08

#### DFGGFN(N)FSQILPDPSKPSKR Hex 5 HexNAc 3 Neu5Ac 1

**Mass:**  3946.7228

(6.731e-07)

**Charge States:** 4

**Start Position**: 783

**End Position**: 803

| Profile Score | Tandem Score | TDA q-value | Volume | Spectrum Matches |
| --- | --- | --- | --- | --- |
| 9.2398 | 5.9617 | 0.0016 | 5.1084e+08 | 2 |

**Adduct Signal Fractions**

- **Unmodified**: 5.1084e+08

#### EGVFVSN(N)GTHWFVTQR @sulfate 1 Hex 4 HexNAc 6

**Mass:**  3809.5933

(5.512e-06)

**Charge States:** 3

**Start Position**: 1079

**End Position**: 1095

| Profile Score | Tandem Score | TDA q-value | Volume | Spectrum Matches |
| --- | --- | --- | --- | --- |
| 12.2793 | 5.9513 | 0.0016 | 1.0129e+08 | 2 |

**Adduct Signal Fractions**

- **Unmodified**: 1.0129e+08

#### VYSSANN(N)C(C)TFEYVSQPFLMDLEGK Fuc 1 Hex 6 HexNAc 5

**Mass:**  4932.0279

(2.243e-07)

**Charge States:** 4

**Start Position**: 146

**End Position**: 170

| Profile Score | Tandem Score | TDA q-value | Volume | Spectrum Matches |
| --- | --- | --- | --- | --- |
| 16.7925 | 5.9040 | 0.0016 | 1.5542e+09 | 2 |

**Adduct Signal Fractions**

- **Unmodified**: 1.5542e+09

#### TPPIKDFGGFN(N)FSQILPDPSKPSK Fuc 3 Hex 8 HexNAc 4

**Mass:**  5180.2787

(3.488e-03)

**Charge States:** 4, 5

**Start Position**: 778

**End Position**: 802

| Profile Score | Tandem Score | TDA q-value | Volume | Spectrum Matches |
| --- | --- | --- | --- | --- |
| 13.7856 | 5.8976 | 0.0010 | 8.9504e+08 | 9 |

**Adduct Signal Fractions**

- **Unmodified**: 3.0960e+10

#### VYSSANN(N)C(C)TFEYVSQPFLMDLEGK Fuc 2 Hex 5 HexNAc 4

**Mass:**  4712.9551

(4.575e-07)

**Charge States:** 3, 4

**Start Position**: 146

**End Position**: 170

| Profile Score | Tandem Score | TDA q-value | Volume | Spectrum Matches |
| --- | --- | --- | --- | --- |
| 14.5068 | 5.8269 | 0.0016 | 1.9226e+09 | 4 |

**Adduct Signal Fractions**

- **Unmodified**: 1.9226e+09

#### TPPIKDFGGFN(N)FSQILPDPSKPSK Fuc 1 Hex 5 HexNAc 4 Neu5Ac 1

**Mass:**  4676.0938

(3.998e-07)

**Charge States:** 4, 5

**Start Position**: 778

**End Position**: 802

| Profile Score | Tandem Score | TDA q-value | Volume | Spectrum Matches |
| --- | --- | --- | --- | --- |
| 13.5249 | 5.8173 | 0.0016 | 1.1176e+09 | 6 |

**Adduct Signal Fractions**

- **Unmodified**: 1.1176e+09

#### EGVFVSN(N)GTHWFVTQR Hex 6 HexNAc 4 Neu5Ac 1

**Mass:**  3938.6357

(1.044e-06)

**Charge States:** 3, 4

**Start Position**: 1079

**End Position**: 1095

| Profile Score | Tandem Score | TDA q-value | Volume | Spectrum Matches |
| --- | --- | --- | --- | --- |
| 11.1888 | 5.6759 | 0.0016 | 1.8663e+08 | 4 |

**Adduct Signal Fractions**

- **Unmodified**: 1.8663e+08

#### TPPIKDFGGFN(N)FSQILPDPSKPSK Hex 6 HexNAc 3

**Mass:**  4197.9175

(9.484e-07)

**Charge States:** 3, 4, 5

**Start Position**: 778

**End Position**: 802

| Profile Score | Tandem Score | TDA q-value | Volume | Spectrum Matches |
| --- | --- | --- | --- | --- |
| 13.4703 | 5.6682 | 0.0016 | 8.3887e+08 | 3 |

**Adduct Signal Fractions**

- **Unmodified**: 8.3887e+08

#### EGVFVSN(N)GTHWFVTQR Hex 6 HexNAc 6

**Mass:**  4053.6985

(7.504e-07)

**Charge States:** 3, 4

**Start Position**: 1079

**End Position**: 1095

| Profile Score | Tandem Score | TDA q-value | Volume | Spectrum Matches |
| --- | --- | --- | --- | --- |
| 13.4679 | 5.6655 | 0.0016 | 1.1161e+09 | 9 |

**Adduct Signal Fractions**

- **Unmodified**: 1.1161e+09

#### GWIFGTTLDSKTQSLLIVNN(N)ATNVVIK @sulfate 1 Hex 5 HexNAc 3 Neu5Ac 1

**Mass:**  4722.1979

(7.734e-06)

**Start Position**: 90

**End Position**: 117

| Profile Score | Tandem Score | TDA q-value | Volume | Spectrum Matches |
| --- | --- | --- | --- | --- |
| 0.0000 | 5.6517 | 0.0016 | 0.0000e+00 | 1 |

Adduct:

No Chromatogram Found

#### EGVFVSN(N)GTHWFVTQR Hex 5 HexNAc 4 Neu5Ac 2

**Mass:**  4067.6817

(6.677e-07)

**Charge States:** 3, 4

**Start Position**: 1079

**End Position**: 1095

| Profile Score | Tandem Score | TDA q-value | Volume | Spectrum Matches |
| --- | --- | --- | --- | --- |
| 11.3144 | 5.6461 | 0.0016 | 6.3725e+08 | 7 |

**Adduct Signal Fractions**

- **Unmodified**: 6.3725e+08

#### TPPIKDFGGFN(N)FSQILPDPSKPSK Hex 5 HexNAc 3 Neu5Ac 1

**Mass:**  4326.9578

(3.096e-07)

**Charge States:** 4

**Start Position**: 778

**End Position**: 802

| Profile Score | Tandem Score | TDA q-value | Volume | Spectrum Matches |
| --- | --- | --- | --- | --- |
| -5.1963 | 5.6423 | 0.0016 | 2.3850e+08 | 5 |

**Adduct Signal Fractions**

- **Unmodified**: 2.3850e+08

#### N(N)FTTAPAIC(C)HDGK Fuc 1 Hex 6 HexNAc 7

**Mass:**  3970.5931

(8.004e-07)

**Charge States:** 3, 4

**Start Position**: 1061

**End Position**: 1074

| Profile Score | Tandem Score | TDA q-value | Volume | Spectrum Matches |
| --- | --- | --- | --- | --- |
| 14.8021 | 5.6247 | 0.0016 | 4.6353e+08 | 3 |

**Adduct Signal Fractions**

- **Unmodified**: 4.6353e+08

#### YNEN(N)GTITDAVDC(C)ALDPLSETK Fuc 1 Hex 5 HexNAc 5 Neu5Ac 1

**Mass:**  4687.9135

(1.344e-07)

**Charge States:** 3, 4

**Start Position**: 266

**End Position**: 288

| Profile Score | Tandem Score | TDA q-value | Volume | Spectrum Matches |
| --- | --- | --- | --- | --- |
| 13.4832 | 5.6222 | 0.0016 | 1.4910e+09 | 4 |

**Adduct Signal Fractions**

- **Unmodified**: 1.4910e+09

#### EGVFVSN(N)GTHWFVTQR Fuc 2 Hex 5 HexNAc 5 Neu5Ac 1

**Mass:**  4271.7625

(-3.106e-06)

**Charge States:** 3

**Start Position**: 1079

**End Position**: 1095

| Profile Score | Tandem Score | TDA q-value | Volume | Spectrum Matches |
| --- | --- | --- | --- | --- |
| -19.4064 | 5.6196 | 0.0016 | 1.5279e+07 | 3 |

**Adduct Signal Fractions**

- **Unmodified**: 1.5279e+07

#### VYSSANN(N)C(C)TFEYVSQPFLMDLEGK Fuc 1 Hex 3 HexNAc 6

**Mass:**  4648.9477

(-4.914e-07)

**Charge States:** 3, 4

**Start Position**: 146

**End Position**: 170

| Profile Score | Tandem Score | TDA q-value | Volume | Spectrum Matches |
| --- | --- | --- | --- | --- |
| 12.5856 | 5.6116 | 0.0016 | 6.4759e+08 | 2 |

**Adduct Signal Fractions**

- **Unmodified**: 6.4759e+08

#### N(N)FTTAPAIC(C)HDGK Fuc 1 Hex 4 HexNAc 5

**Mass:**  3242.3325

(9.279e-04)

**Start Position**: 1061

**End Position**: 1074

| Profile Score | Tandem Score | TDA q-value | Volume | Spectrum Matches |
| --- | --- | --- | --- | --- |
| 0.0000 | 5.6112 | 0.0016 | 0.0000e+00 | 5 |

Adduct:

No Chromatogram Found

#### N(N)FTTAPAIC(C)HDGK Fuc 1 Hex 4 HexNAc 3

**Mass:**  2836.1781

(7.090e-04)

**Start Position**: 1061

**End Position**: 1074

| Profile Score | Tandem Score | TDA q-value | Volume | Spectrum Matches |
| --- | --- | --- | --- | --- |
| 0.0000 | 5.6085 | 0.0016 | 0.0000e+00 | 2 |

Adduct:

No Chromatogram Found

#### QC(C)VN(N)LTTR Fuc 1 Hex 6 HexNAc 3

**Mass:**  2719.1160

(8.826e-03)

**Charge States:** 2, 3

**Start Position**: 1

**End Position**: 9

| Profile Score | Tandem Score | TDA q-value | Volume | Spectrum Matches |
| --- | --- | --- | --- | --- |
| 10.1594 | 5.5721 | 0.0110 | 5.6002e+08 | 3 |

**Adduct Signal Fractions**

- **Na**: 4.8550e+09

#### EGVFVSN(N)GTHWFVTQR Hex 6 HexNAc 6 Neu5Ac 2

**Mass:**  4638.9083

(4.325e-04)

**Start Position**: 1079

**End Position**: 1095

| Profile Score | Tandem Score | TDA q-value | Volume | Spectrum Matches |
| --- | --- | --- | --- | --- |
| 0.0000 | 5.5663 | 0.0016 | 0.0000e+00 | 2 |

Adduct:

No Chromatogram Found

#### YNEN(N)GTITDAVDC(C)ALDPLSETK Hex 3 HexNAc 5

**Mass:**  3926.6504

(4.110e-07)

**Charge States:** 3, 4

**Start Position**: 266

**End Position**: 288

| Profile Score | Tandem Score | TDA q-value | Volume | Spectrum Matches |
| --- | --- | --- | --- | --- |
| 16.4755 | 5.5569 | 0.0016 | 1.6145e+09 | 6 |

**Adduct Signal Fractions**

- **Unmodified**: 5.9095e+09

#### SSVLHSTQDLFLPFFSNVTWFHAIHVSGTN(N)GTK Fuc 1 Hex 5 HexNAc 3

**Mass:**  5239.3983

(7.616e-07)

**Charge States:** 4, 5

**Start Position**: 32

**End Position**: 65

| Profile Score | Tandem Score | TDA q-value | Volume | Spectrum Matches |
| --- | --- | --- | --- | --- |
| 12.9545 | 5.5540 | 0.0016 | 3.4533e+08 | 1 |

**Adduct Signal Fractions**

- **Unmodified**: 3.4533e+08

#### DFGGFN(N)FSQILPDPSKPSK Fuc 1 Hex 5 HexNAc 3

**Mass:**  3645.5870

(8.778e-08)

**Charge States:** 3, 4

**Start Position**: 783

**End Position**: 802

| Profile Score | Tandem Score | TDA q-value | Volume | Spectrum Matches |
| --- | --- | --- | --- | --- |
| 10.2095 | 5.5009 | 0.0016 | 1.9857e+09 | 15 |

**Adduct Signal Fractions**

- **Unmodified**: 1.9857e+09

#### EGVFVSN(N)GTHWFVTQR Hex 5 HexNAc 6

**Mass:**  3891.6460

(2.490e-07)

**Charge States:** 3, 4

**Start Position**: 1079

**End Position**: 1095

| Profile Score | Tandem Score | TDA q-value | Volume | Spectrum Matches |
| --- | --- | --- | --- | --- |
| 12.5468 | 5.4868 | 0.0016 | 7.5836e+08 | 6 |

**Adduct Signal Fractions**

- **Unmodified**: 7.5836e+08

#### N(N)FTTAPAIC(C)HDGK Hex 6 HexNAc 4

**Mass:**  3216.2990

(3.120e-04)

**Start Position**: 1061

**End Position**: 1074

| Profile Score | Tandem Score | TDA q-value | Volume | Spectrum Matches |
| --- | --- | --- | --- | --- |
| 0.0000 | 5.4788 | 0.0016 | 0.0000e+00 | 1 |

Adduct:

No Chromatogram Found

#### EGVFVSN(N)GTHWFVTQR Fuc 1 Hex 5 HexNAc 6

**Mass:**  4037.7042

(6.440e-07)

**Charge States:** 3, 4

**Start Position**: 1079

**End Position**: 1095

| Profile Score | Tandem Score | TDA q-value | Volume | Spectrum Matches |
| --- | --- | --- | --- | --- |
| 13.2583 | 5.4187 | 0.0016 | 6.4339e+08 | 4 |

**Adduct Signal Fractions**

- **Unmodified**: 6.4339e+08

#### NLN(N)ESLIDLQELGK Fuc 1 Hex 7 HexNAc 6 Neu5Ac 2

**Mass:**  4665.9340

(1.000e-06)

**Charge States:** 4

**Start Position**: 1179

**End Position**: 1193

| Profile Score | Tandem Score | TDA q-value | Volume | Spectrum Matches |
| --- | --- | --- | --- | --- |
| 11.0817 | 5.4025 | 0.0016 | 1.0854e+08 | 1 |

**Adduct Signal Fractions**

- **Unmodified**: 1.0854e+08

#### DFGGFN(N)FSQILPDPSKPSK Fuc 1 Hex 4 HexNAc 5 Neu5Ac 1

**Mass:**  4180.7912

(1.719e-07)

**Charge States:** 3, 4

**Start Position**: 783

**End Position**: 802

| Profile Score | Tandem Score | TDA q-value | Volume | Spectrum Matches |
| --- | --- | --- | --- | --- |
| 11.1373 | 5.3961 | 0.0016 | 2.1190e+08 | 4 |

**Adduct Signal Fractions**

- **Unmodified**: 2.1190e+08

#### DFGGFN(N)FSQILPDPSKPSK Fuc 1 Hex 5 HexNAc 5

**Mass:**  4051.7471

(1.652e-07)

**Charge States:** 3, 4

**Start Position**: 783

**End Position**: 802

| Profile Score | Tandem Score | TDA q-value | Volume | Spectrum Matches |
| --- | --- | --- | --- | --- |
| 10.5329 | 5.3244 | 0.0016 | 1.6414e+08 | 3 |

**Adduct Signal Fractions**

- **Unmodified**: 1.6414e+08

#### N(N)FTTAPAIC(C)HDGK Fuc 1 Hex 4 HexNAc 2

**Mass:**  2631.0939

(6.461e-03)

**Charge States:** 2

**Start Position**: 1061

**End Position**: 1074

| Profile Score | Tandem Score | TDA q-value | Volume | Spectrum Matches |
| --- | --- | --- | --- | --- |
| -21.2745 | 5.2528 | 0.0016 | 3.9705e+07 | 4 |

**Adduct Signal Fractions**

- **NH3**: 6.9215e+06
- **Unmodified**: 3.2784e+07

#### FPNITNLC(C)PFGEVFN(N)ATR Fuc 2 Hex 7 HexNAc 8 Neu5Ac 1

**Mass:**  5441.2453

(5.543e-04)

**Start Position**: 316

**End Position**: 334

| Profile Score | Tandem Score | TDA q-value | Volume | Spectrum Matches |
| --- | --- | --- | --- | --- |
| 0.0000 | 5.2358 | 0.0016 | 0.0000e+00 | 1 |

Adduct:

No Chromatogram Found

#### SSVLHSTQDLFLPFFSN(N)VTWFHAIHVSGTNGTK Fuc 2 Hex 9 HexNAc 6

**Mass:**  6642.9117

(-4.198e-07)

**Charge States:** 5, 6

**Start Position**: 32

**End Position**: 65

| Profile Score | Tandem Score | TDA q-value | Volume | Spectrum Matches |
| --- | --- | --- | --- | --- |
| 12.6111 | 5.2220 | 0.0016 | 1.3338e+09 | 1 |

**Adduct Signal Fractions**

- **Unmodified**: 1.3338e+09

#### SSVLHSTQDLFLPFFSNVTWFHAIHVSGTN(N)GTK Fuc 2 Hex 9 HexNAc 6

**Mass:**  6642.9117

(-4.198e-07)

**Charge States:** 5, 6

**Start Position**: 32

**End Position**: 65

| Profile Score | Tandem Score | TDA q-value | Volume | Spectrum Matches |
| --- | --- | --- | --- | --- |
| 12.6111 | 5.2220 | 0.0016 | 1.3338e+09 | 1 |

**Adduct Signal Fractions**

- **Unmodified**: 1.3338e+09

#### SSVLHSTQDLFLPFFSNVTWFHAIHVSGTN(N)GTK Fuc 2 Hex 8 HexNAc 8

**Mass:**  6887.0092

(7.244e-07)

**Charge States:** 5, 6, 7

**Start Position**: 32

**End Position**: 65

| Profile Score | Tandem Score | TDA q-value | Volume | Spectrum Matches |
| --- | --- | --- | --- | --- |
| 13.0253 | 5.2209 | 0.0016 | 4.0500e+09 | 5 |

**Adduct Signal Fractions**

- **Unmodified**: 4.0500e+09

#### EGVFVSN(N)GTHWFVTQR Hex 4 HexNAc 2

**Mass:**  2917.2794

(1.935e-06)

**Charge States:** 3

**Start Position**: 1079

**End Position**: 1095

| Profile Score | Tandem Score | TDA q-value | Volume | Spectrum Matches |
| --- | --- | --- | --- | --- |
| 10.4783 | 5.1999 | 0.0016 | 5.5197e+07 | 1 |

**Adduct Signal Fractions**

- **Unmodified**: 5.5197e+07

#### VYSSANN(N)C(C)TFEYVSQPFLMDLEGK Fuc 1 Hex 3 HexNAc 4

**Mass:**  4242.7937

(3.025e-07)

**Charge States:** 3, 4

**Start Position**: 146

**End Position**: 170

| Profile Score | Tandem Score | TDA q-value | Volume | Spectrum Matches |
| --- | --- | --- | --- | --- |
| 17.5982 | 5.1898 | 0.0016 | 4.9117e+09 | 5 |

**Adduct Signal Fractions**

- **Unmodified**: 4.9117e+09

#### SSVLHSTQDLFLPFFSNVTWFHAIHVSGTN(N)GTK Fuc 1 Hex 8 HexNAc 7 Neu5Ac 1

**Mass:**  6828.9641

(3.064e-07)

**Charge States:** 5, 6, 7

**Start Position**: 32

**End Position**: 65

| Profile Score | Tandem Score | TDA q-value | Volume | Spectrum Matches |
| --- | --- | --- | --- | --- |
| 15.2308 | 5.1894 | 0.0016 | 3.1334e+09 | 3 |

**Adduct Signal Fractions**

- **Unmodified**: 3.1334e+09

#### VYSSANN(N)C(C)TFEYVSQPFLMDLEGK Hex 3 HexNAc 5

**Mass:**  4299.8114

(-5.207e-07)

**Charge States:** 3, 4

**Start Position**: 146

**End Position**: 170

| Profile Score | Tandem Score | TDA q-value | Volume | Spectrum Matches |
| --- | --- | --- | --- | --- |
| 15.4586 | 5.1518 | 0.0016 | 2.3781e+09 | 4 |

**Adduct Signal Fractions**

- **Unmodified**: 2.3781e+09

#### VYSSANN(N)C(C)TFEYVSQPFLMDLEGK Fuc 1 Hex 5 HexNAc 4 Neu5Ac 2

**Mass:**  5149.0875

(1.608e-07)

**Charge States:** 4, 5

**Start Position**: 146

**End Position**: 170

| Profile Score | Tandem Score | TDA q-value | Volume | Spectrum Matches |
| --- | --- | --- | --- | --- |
| 15.9160 | 5.1195 | 0.0016 | 3.6471e+09 | 6 |

**Adduct Signal Fractions**

- **Unmodified**: 3.6471e+09

#### VYSSANN(N)C(C)TFEYVSQPFLMDLEGK Fuc 1 Hex 4 HexNAc 4

**Mass:**  4404.8455

(3.236e-08)

**Charge States:** 3, 4

**Start Position**: 146

**End Position**: 170

| Profile Score | Tandem Score | TDA q-value | Volume | Spectrum Matches |
| --- | --- | --- | --- | --- |
| 15.8985 | 5.1183 | 0.0016 | 2.4450e+09 | 6 |

**Adduct Signal Fractions**

- **Unmodified**: 2.4450e+09

#### SNIIRGWIFGTTLDSKTQSLLIVNN(N)ATNVVIK Fuc 1 Hex 5 HexNAc 8

**Mass:**  6095.9266

(2.797e-03)

**Charge States:** 4, 5

**Start Position**: 85

**End Position**: 117

| Profile Score | Tandem Score | TDA q-value | Volume | Spectrum Matches |
| --- | --- | --- | --- | --- |
| 11.9235 | 5.0309 | 0.0016 | 2.0040e+09 | 2 |

**Adduct Signal Fractions**

- **NH3**: 2.0040e+09

#### TPPIKDFGGFN(N)FSQILPDPSKPSK Fuc 3 Hex 8 HexNAc 4

**Mass:**  5183.2943

(3.878e-03)

**Start Position**: 778

**End Position**: 802

| Profile Score | Tandem Score | TDA q-value | Volume | Spectrum Matches |
| --- | --- | --- | --- | --- |
| 0.0000 | 4.9876 | 0.0016 | 0.0000e+00 | 1 |

Adduct:

No Chromatogram Found

#### VYSSANN(N)C(C)TFEYVSQPFLMDLEGK Hex 5 HexNAc 5

**Mass:**  4623.9210

(6.800e-07)

**Charge States:** 3, 4

**Start Position**: 146

**End Position**: 170

| Profile Score | Tandem Score | TDA q-value | Volume | Spectrum Matches |
| --- | --- | --- | --- | --- |
| 12.4566 | 4.9749 | 0.0016 | 7.1662e+08 | 4 |

**Adduct Signal Fractions**

- **Unmodified**: 7.1662e+08

#### N(N)FTTAPAIC(C)HDGK Fuc 1 Hex 7 HexNAc 7

**Mass:**  4132.6478

(6.315e-07)

**Charge States:** 3, 4

**Start Position**: 1061

**End Position**: 1074

| Profile Score | Tandem Score | TDA q-value | Volume | Spectrum Matches |
| --- | --- | --- | --- | --- |
| 14.5226 | 4.9370 | 0.0016 | 2.0126e+08 | 3 |

**Adduct Signal Fractions**

- **Unmodified**: 2.0126e+08

#### N(N)FTTAPAIC(C)HDGK Fuc 1 Hex 9 HexNAc 2

**Mass:**  3441.3311

(-5.476e-06)

**Charge States:** 3, 4

**Start Position**: 1061

**End Position**: 1074

| Profile Score | Tandem Score | TDA q-value | Volume | Spectrum Matches |
| --- | --- | --- | --- | --- |
| -20.7065 | 4.9104 | 0.0016 | 7.6530e+06 | 1 |

**Adduct Signal Fractions**

- **Unmodified**: 7.6530e+06

#### TPPIKDFGGFN(N)FSQILPDPSKPSK Hex 8 HexNAc 2

**Mass:**  4318.9382

(-3.600e-07)

**Charge States:** 4

**Start Position**: 778

**End Position**: 802

| Profile Score | Tandem Score | TDA q-value | Volume | Spectrum Matches |
| --- | --- | --- | --- | --- |
| 11.3619 | 4.8757 | 0.0016 | 9.4700e+08 | 3 |

**Adduct Signal Fractions**

- **Unmodified**: 9.4700e+08

#### TPPIKDFGGFN(N)FSQILPDPSKPSK Fuc 1 Hex 4 HexNAc 4

**Mass:**  4222.9465

(2.382e-04)

**Charge States:** 4

**Start Position**: 778

**End Position**: 802

| Profile Score | Tandem Score | TDA q-value | Volume | Spectrum Matches |
| --- | --- | --- | --- | --- |
| 11.4882 | 4.7997 | 0.0024 | 4.5542e+08 | 6 |

**Adduct Signal Fractions**

- **Unmodified**: 4.5542e+08

#### N(N)FTTAPAIC(C)HDGK Hex 5 HexNAc 6 Neu5Ac 2

**Mass:**  4060.6247

(4.709e-03)

**Start Position**: 1061

**End Position**: 1074

| Profile Score | Tandem Score | TDA q-value | Volume | Spectrum Matches |
| --- | --- | --- | --- | --- |
| 0.0000 | 4.7636 | 0.0024 | 0.0000e+00 | 1 |

Adduct:

No Chromatogram Found

#### N(N)FTTAPAIC(C)HDGK Fuc 1 Hex 5 HexNAc 7

**Mass:**  3808.5387

(3.336e-08)

**Charge States:** 3, 4

**Start Position**: 1061

**End Position**: 1074

| Profile Score | Tandem Score | TDA q-value | Volume | Spectrum Matches |
| --- | --- | --- | --- | --- |
| 12.1749 | 4.7626 | 0.0024 | 2.9049e+08 | 6 |

**Adduct Signal Fractions**

- **Unmodified**: 2.9049e+08

#### QC(C)VN(N)LTTR Fuc 3 Hex 4 HexNAc 4

**Mass:**  2889.1834

(7.954e-03)

**Charge States:** 2, 3

**Start Position**: 1

**End Position**: 9

| Profile Score | Tandem Score | TDA q-value | Volume | Spectrum Matches |
| --- | --- | --- | --- | --- |
| 15.7106 | 4.7475 | 0.0110 | 1.3749e+10 | 7 |

**Adduct Signal Fractions**

- **Na**: 1.3749e+10

#### YNEN(N)GTITDAVDC(C)ALDPLSETK Fuc 1 Hex 4 HexNAc 5 Neu5Ac 1

**Mass:**  4525.8584

(2.216e-07)

**Charge States:** 3, 4

**Start Position**: 266

**End Position**: 288

| Profile Score | Tandem Score | TDA q-value | Volume | Spectrum Matches |
| --- | --- | --- | --- | --- |
| 14.5490 | 4.7464 | 0.0024 | 9.3211e+08 | 2 |

**Adduct Signal Fractions**

- **Unmodified**: 9.3211e+08

#### EGVFVSN(N)GTHWFVTQR Fuc 1 Hex 5 HexNAc 5

**Mass:**  3834.6242

(2.607e-04)

**Charge States:** 3, 4

**Start Position**: 1079

**End Position**: 1095

| Profile Score | Tandem Score | TDA q-value | Volume | Spectrum Matches |
| --- | --- | --- | --- | --- |
| 11.6995 | 4.7018 | 0.0041 | 7.3458e+08 | 10 |

**Adduct Signal Fractions**

- **Unmodified**: 7.3458e+08

#### EGVFVSN(N)GTHWFVTQR Hex 5 HexNAc 5 Neu5Ac 2

**Mass:**  4272.7644

(4.697e-04)

**Start Position**: 1079

**End Position**: 1095

| Profile Score | Tandem Score | TDA q-value | Volume | Spectrum Matches |
| --- | --- | --- | --- | --- |
| 0.0000 | 4.6880 | 0.0048 | 0.0000e+00 | 1 |

Adduct:

No Chromatogram Found

#### EGVFVSN(N)GTHWFVTQR Fuc 1 Hex 5 HexNAc 5 Neu5Ac 2

**Mass:**  4418.8269

(4.553e-04)

**Start Position**: 1079

**End Position**: 1095

| Profile Score | Tandem Score | TDA q-value | Volume | Spectrum Matches |
| --- | --- | --- | --- | --- |
| 0.0000 | 4.6813 | 0.0048 | 0.0000e+00 | 1 |

Adduct:

No Chromatogram Found

#### N(N)FTTAPAIC(C)HDGK Fuc 1 Hex 5 HexNAc 5

**Mass:**  3404.3867

(8.834e-04)

**Start Position**: 1061

**End Position**: 1074

| Profile Score | Tandem Score | TDA q-value | Volume | Spectrum Matches |
| --- | --- | --- | --- | --- |
| 0.0000 | 4.6742 | 0.0048 | 0.0000e+00 | 2 |

Adduct:

No Chromatogram Found

#### TPPIKDFGGFN(N)FSQILPDPSKPSK Fuc 4 Hex 5 HexNAc 5

**Mass:**  5044.3259

(3.596e-03)

**Start Position**: 778

**End Position**: 802

| Profile Score | Tandem Score | TDA q-value | Volume | Spectrum Matches |
| --- | --- | --- | --- | --- |
| 0.0000 | 4.6713 | 0.0048 | 0.0000e+00 | 1 |

Adduct:

No Chromatogram Found

#### N(N)FTTAPAIC(C)HDGK Hex 5 HexNAc 4

**Mass:**  3055.2520

(6.580e-04)

**Start Position**: 1061

**End Position**: 1074

| Profile Score | Tandem Score | TDA q-value | Volume | Spectrum Matches |
| --- | --- | --- | --- | --- |
| 0.0000 | 4.6614 | 0.0048 | 0.0000e+00 | 1 |

Adduct:

No Chromatogram Found

#### QC(C)VN(N)LTTRTQLPPAYTNSFTR @sulfate 1 Fuc 1 Hex 3 HexNAc 5

**Mass:**  4197.7868

(7.125e-04)

**Start Position**: 1

**End Position**: 22

| Profile Score | Tandem Score | TDA q-value | Volume | Spectrum Matches |
| --- | --- | --- | --- | --- |
| 0.0000 | 4.6573 | 0.0048 | 0.0000e+00 | 1 |

Adduct:

No Chromatogram Found

#### N(N)FTTAPAIC(C)HDGK Fuc 1 Hex 4 HexNAc 5 Neu5Ac 1

**Mass:**  3532.4284

(2.848e-04)

**Start Position**: 1061

**End Position**: 1074

| Profile Score | Tandem Score | TDA q-value | Volume | Spectrum Matches |
| --- | --- | --- | --- | --- |
| 0.0000 | 4.6407 | 0.0057 | 0.0000e+00 | 2 |

Adduct:

No Chromatogram Found

#### EGVFVSN(N)GTHWFVTQR Fuc 3 Hex 3 HexNAc 4 Neu5Ac 2

**Mass:**  4183.7450

(4.787e-04)

**Charge States:** 3

**Start Position**: 1079

**End Position**: 1095

| Profile Score | Tandem Score | TDA q-value | Volume | Spectrum Matches |
| --- | --- | --- | --- | --- |
| 6.1787 | 4.6355 | 0.0057 | 2.3982e+08 | 2 |

**Adduct Signal Fractions**

- **Unmodified**: 2.3982e+08

#### YNEN(N)GTITDAVDC(C)ALDPLSETK Fuc 1 Hex 4 HexNAc 6

**Mass:**  4437.8449

(1.888e-07)

**Charge States:** 3, 4

**Start Position**: 266

**End Position**: 288

| Profile Score | Tandem Score | TDA q-value | Volume | Spectrum Matches |
| --- | --- | --- | --- | --- |
| 11.6173 | 4.6307 | 0.0057 | 2.7511e+09 | 4 |

**Adduct Signal Fractions**

- **Unmodified**: 2.7511e+09

#### YNEN(N)GTITDAVDC(C)ALDPLSETK Fuc 1 Hex 7 HexNAc 6

**Mass:**  4923.9985

(6.357e-08)

**Charge States:** 4

**Start Position**: 266

**End Position**: 288

| Profile Score | Tandem Score | TDA q-value | Volume | Spectrum Matches |
| --- | --- | --- | --- | --- |
| 12.3015 | 4.6081 | 0.0057 | 4.0624e+08 | 1 |

**Adduct Signal Fractions**

- **Unmodified**: 4.0624e+08

#### EGVFVSN(N)GTHWFVTQR Fuc 2 Hex 4 HexNAc 4

**Mass:**  3615.5484

(1.172e-07)

**Charge States:** 3

**Start Position**: 1079

**End Position**: 1095

| Profile Score | Tandem Score | TDA q-value | Volume | Spectrum Matches |
| --- | --- | --- | --- | --- |
| 12.0336 | 4.6038 | 0.0057 | 1.4888e+08 | 2 |

**Adduct Signal Fractions**

- **Unmodified**: 1.4888e+08

#### EGVFVSN(N)GTHWFVTQR Fuc 1 Hex 5 HexNAc 5 Neu5Ac 2

**Mass:**  4416.8187

(3.331e-07)

**Charge States:** 3, 4

**Start Position**: 1079

**End Position**: 1095

| Profile Score | Tandem Score | TDA q-value | Volume | Spectrum Matches |
| --- | --- | --- | --- | --- |
| 11.9308 | 4.5954 | 0.0057 | 8.7431e+08 | 6 |

**Adduct Signal Fractions**

- **Unmodified**: 8.7431e+08

#### FPNITNLC(C)PFGEVFN(N)ATR Fuc 2 Hex 7 HexNAc 8

**Mass:**  5147.1353

(6.871e-08)

**Charge States:** 4

**Start Position**: 316

**End Position**: 334

| Profile Score | Tandem Score | TDA q-value | Volume | Spectrum Matches |
| --- | --- | --- | --- | --- |
| 12.8067 | 4.5791 | 0.0057 | 2.5586e+09 | 2 |

**Adduct Signal Fractions**

- **Unmodified**: 2.5586e+09

#### YNEN(N)GTITDAVDC(C)ALDPLSETK Hex 6 HexNAc 5 Neu5Ac 1

**Mass:**  4703.9062

(6.162e-07)

**Charge States:** 3, 4

**Start Position**: 266

**End Position**: 288

| Profile Score | Tandem Score | TDA q-value | Volume | Spectrum Matches |
| --- | --- | --- | --- | --- |
| 12.8867 | 4.5622 | 0.0057 | 7.6590e+08 | 2 |

**Adduct Signal Fractions**

- **Unmodified**: 7.6590e+08

#### TQSLLIVNN(N)ATNVVIK Hex 5 HexNAc 5

**Mass:**  3551.6605

(6.435e-07)

**Charge States:** 3, 4

**Start Position**: 101

**End Position**: 117

| Profile Score | Tandem Score | TDA q-value | Volume | Spectrum Matches |
| --- | --- | --- | --- | --- |
| 13.5222 | 4.5396 | 0.0057 | 6.6320e+08 | 5 |

**Adduct Signal Fractions**

- **Unmodified**: 6.6320e+08

#### YNEN(N)GTITDAVDC(C)ALDPLSETK Fuc 1 Hex 4 HexNAc 4

**Mass:**  4031.6827

(2.497e-04)

**Charge States:** 3, 4

**Start Position**: 266

**End Position**: 288

| Profile Score | Tandem Score | TDA q-value | Volume | Spectrum Matches |
| --- | --- | --- | --- | --- |
| 10.1162 | 4.5329 | 0.0057 | -1.0405e+09 | 10 |

**Adduct Signal Fractions**

- **Unmodified**: 3.2545e+09

#### TQSLLIVNN(N)ATNVVIK Fuc 2 Hex 4 HexNAc 3

**Mass:**  3276.5652

(3.058e-04)

**Start Position**: 101

**End Position**: 117

| Profile Score | Tandem Score | TDA q-value | Volume | Spectrum Matches |
| --- | --- | --- | --- | --- |
| 0.0000 | 4.5280 | 0.0057 | 0.0000e+00 | 3 |

Adduct:

No Chromatogram Found

#### N(N)FTTAPAIC(C)HDGK @sulfate 1 Hex 3 HexNAc 5 Neu5Ac 1

**Mass:**  3326.2685

(6.963e-03)

**Start Position**: 1061

**End Position**: 1074

| Profile Score | Tandem Score | TDA q-value | Volume | Spectrum Matches |
| --- | --- | --- | --- | --- |
| 0.0000 | 4.5008 | 0.0057 | 0.0000e+00 | 1 |

Adduct:

No Chromatogram Found

#### YNEN(N)GTITDAVDC(C)ALDPLSETK Hex 5 HexNAc 5

**Mass:**  4250.7610

(1.163e-06)

**Charge States:** 3, 4

**Start Position**: 266

**End Position**: 288

| Profile Score | Tandem Score | TDA q-value | Volume | Spectrum Matches |
| --- | --- | --- | --- | --- |
| 11.6256 | 4.4991 | 0.0057 | 1.2517e+09 | 8 |

**Adduct Signal Fractions**

- **Unmodified**: 1.2517e+09

#### N(N)FTTAPAIC(C)HDGK Fuc 1 Hex 5 HexNAc 5 Neu5Ac 2

**Mass:**  3984.5730

(5.014e-07)

**Charge States:** 3, 4

**Start Position**: 1061

**End Position**: 1074

| Profile Score | Tandem Score | TDA q-value | Volume | Spectrum Matches |
| --- | --- | --- | --- | --- |
| 11.8273 | 4.4891 | 0.0057 | 3.1445e+08 | 2 |

**Adduct Signal Fractions**

- **Unmodified**: 3.1445e+08

#### EGVFVSN(N)GTHWFVTQR Fuc 1 Hex 6 HexNAc 5 Neu5Ac 2

**Mass:**  4578.8663

(1.322e-06)

**Charge States:** 3, 4

**Start Position**: 1079

**End Position**: 1095

| Profile Score | Tandem Score | TDA q-value | Volume | Spectrum Matches |
| --- | --- | --- | --- | --- |
| 12.5601 | 4.4886 | 0.0057 | 8.0371e+08 | 3 |

**Adduct Signal Fractions**

- **Unmodified**: 8.0371e+08

#### EGVFVSN(N)GTHWFVTQR Fuc 3 Hex 5 HexNAc 4 Neu5Ac 1

**Mass:**  4214.7423

(-3.521e-06)

**Charge States:** 4

**Start Position**: 1079

**End Position**: 1095

| Profile Score | Tandem Score | TDA q-value | Volume | Spectrum Matches |
| --- | --- | --- | --- | --- |
| -5.0454 | 4.4818 | 0.0057 | 1.2310e+07 | 1 |

**Adduct Signal Fractions**

- **Unmodified**: 1.2310e+07

#### SSVLHSTQDLFLPFFSNVTWFHAIHVSGTN(N)GTK Fuc 1 Hex 9 HexNAc 7

**Mass:**  6699.9262

(1.092e-06)

**Charge States:** 5, 6, 7

**Start Position**: 32

**End Position**: 65

| Profile Score | Tandem Score | TDA q-value | Volume | Spectrum Matches |
| --- | --- | --- | --- | --- |
| 12.3904 | 4.4702 | 0.0057 | 5.3321e+09 | 6 |

**Adduct Signal Fractions**

- **Unmodified**: 5.3321e+09

#### N(N)FTTAPAIC(C)HDGK @sulfate 1 Fuc 1 Hex 3 HexNAc 4

**Mass:**  2955.1667

(5.767e-03)

**Charge States:** 3

**Start Position**: 1061

**End Position**: 1074

| Profile Score | Tandem Score | TDA q-value | Volume | Spectrum Matches |
| --- | --- | --- | --- | --- |
| -19.7072 | 4.4661 | 0.0057 | 4.7277e+06 | 1 |

**Adduct Signal Fractions**

- **NH3**: 4.7277e+06

#### EGVFVSN(N)GTHWFVTQR Hex 8 HexNAc 2

**Mass:**  3565.4870

(-3.330e-07)

**Charge States:** 3

**Start Position**: 1079

**End Position**: 1095

| Profile Score | Tandem Score | TDA q-value | Volume | Spectrum Matches |
| --- | --- | --- | --- | --- |
| 12.7945 | 4.4603 | 0.0057 | 3.1449e+08 | 2 |

**Adduct Signal Fractions**

- **Unmodified**: 3.1449e+08

#### N(N)FTTAPAIC(C)HDGK Hex 6 HexNAc 5 Neu5Ac 1

**Mass:**  3709.4731

(2.700e-04)

**Charge States:** 3, 4

**Start Position**: 1061

**End Position**: 1074

| Profile Score | Tandem Score | TDA q-value | Volume | Spectrum Matches |
| --- | --- | --- | --- | --- |
| 10.5617 | 4.4347 | 0.0063 | 9.5703e+07 | 4 |

**Adduct Signal Fractions**

- **Unmodified**: 9.5703e+07

#### DFGGFN(N)FSQILPDPSKPSK Hex 4 HexNAc 5

**Mass:**  3743.6346

(4.135e-07)

**Charge States:** 3, 4

**Start Position**: 783

**End Position**: 802

| Profile Score | Tandem Score | TDA q-value | Volume | Spectrum Matches |
| --- | --- | --- | --- | --- |
| -6.2752 | 4.4244 | 0.0063 | 1.8233e+08 | 11 |

**Adduct Signal Fractions**

- **Unmodified**: 1.8233e+08

#### N(N)FTTAPAIC(C)HDGK Fuc 1 Hex 4 HexNAc 4

**Mass:**  3040.2603

(9.918e-04)

**Start Position**: 1061

**End Position**: 1074

| Profile Score | Tandem Score | TDA q-value | Volume | Spectrum Matches |
| --- | --- | --- | --- | --- |
| 0.0000 | 4.4164 | 0.0063 | 0.0000e+00 | 1 |

Adduct:

No Chromatogram Found

#### VYSSANN(N)C(C)TFEYVSQPFLMDLEGK Fuc 2 Hex 4 HexNAc 5

**Mass:**  4753.9816

(-2.238e-08)

**Charge States:** 3, 4

**Start Position**: 146

**End Position**: 170

| Profile Score | Tandem Score | TDA q-value | Volume | Spectrum Matches |
| --- | --- | --- | --- | --- |
| 11.0351 | 4.4049 | 0.0063 | 1.4103e+09 | 3 |

**Adduct Signal Fractions**

- **NH3**: 7.8099e+07
- **Unmodified**: 1.3322e+09

#### N(N)FTTAPAIC(C)HDGK Fuc 2 Hex 4 HexNAc 6

**Mass:**  3589.4346

(4.735e-03)

**Charge States:** 4

**Start Position**: 1061

**End Position**: 1074

| Profile Score | Tandem Score | TDA q-value | Volume | Spectrum Matches |
| --- | --- | --- | --- | --- |
| -20.0777 | 4.3952 | 0.0063 | 3.3631e+06 | 1 |

**Adduct Signal Fractions**

- **NH3**: 3.3631e+06

#### VYSSANN(N)C(C)TFEYVSQPFLMDLEGK Hex 5 HexNAc 4

**Mass:**  4420.8406

(6.299e-07)

**Charge States:** 3, 4

**Start Position**: 146

**End Position**: 170

| Profile Score | Tandem Score | TDA q-value | Volume | Spectrum Matches |
| --- | --- | --- | --- | --- |
| 14.6755 | 4.3933 | 0.0063 | 1.6988e+09 | 5 |

**Adduct Signal Fractions**

- **Unmodified**: 1.6988e+09

#### N(N)FTTAPAIC(C)HDGK @sulfate 1 Fuc 1 Hex 4 HexNAc 3

**Mass:**  2914.1471

(7.450e-06)

**Charge States:** 4, 5

**Start Position**: 1061

**End Position**: 1074

| Profile Score | Tandem Score | TDA q-value | Volume | Spectrum Matches |
| --- | --- | --- | --- | --- |
| -6.5492 | 4.3854 | 0.0063 | 2.0437e+07 | 1 |

**Adduct Signal Fractions**

- **Unmodified**: 2.0437e+07

#### N(N)FTTAPAIC(C)HDGK @sulfate 1 Fuc 2 Hex 3 HexNAc 3

**Mass:**  2915.1492

(5.872e-03)

**Start Position**: 1061

**End Position**: 1074

| Profile Score | Tandem Score | TDA q-value | Volume | Spectrum Matches |
| --- | --- | --- | --- | --- |
| 0.0000 | 4.3634 | 0.0063 | 0.0000e+00 | 1 |

Adduct:

No Chromatogram Found

#### EGVFVSN(N)GTHWFVTQR @sulfate 1 Fuc 1 Hex 8 HexNAc 2

**Mass:**  3815.5112

(6.332e-03)

**Start Position**: 1079

**End Position**: 1095

| Profile Score | Tandem Score | TDA q-value | Volume | Spectrum Matches |
| --- | --- | --- | --- | --- |
| 0.0000 | 4.3587 | 0.0063 | 0.0000e+00 | 1 |

Adduct:

No Chromatogram Found

#### DFGGFN(N)FSQILPDPSKPSK Fuc 1 Hex 3 HexNAc 3

**Mass:**  3322.4870

(3.028e-04)

**Start Position**: 783

**End Position**: 802

| Profile Score | Tandem Score | TDA q-value | Volume | Spectrum Matches |
| --- | --- | --- | --- | --- |
| 0.0000 | 4.3522 | 0.0063 | 0.0000e+00 | 2 |

Adduct:

No Chromatogram Found

#### DFGGFN(N)FSQILPDPSKPSKR Hex 6 HexNAc 3

**Mass:**  3819.6865

(5.249e-04)

**Start Position**: 783

**End Position**: 803

| Profile Score | Tandem Score | TDA q-value | Volume | Spectrum Matches |
| --- | --- | --- | --- | --- |
| 0.0000 | 4.3236 | 0.0068 | 0.0000e+00 | 1 |

Adduct:

No Chromatogram Found

#### DFGGFN(N)FSQILPDPSKPSK Fuc 3 Hex 3 HexNAc 4

**Mass:**  3817.6812

(4.455e-03)

**Charge States:** 3, 4

**Start Position**: 783

**End Position**: 802

| Profile Score | Tandem Score | TDA q-value | Volume | Spectrum Matches |
| --- | --- | --- | --- | --- |
| -5.2181 | 4.2976 | 0.0057 | 1.3616e+08 | 8 |

**Adduct Signal Fractions**

- **Unmodified**: 2.8200e+07
- **NH3**: 1.0796e+08

#### NLN(N)ESLIDLQELGK @sulfate 1 Fuc 1 Hex 5 HexNAc 6 Neu5Ac 2

**Mass:**  4421.7834

(4.614e-07)

**Charge States:** 3, 4

**Start Position**: 1179

**End Position**: 1193

| Profile Score | Tandem Score | TDA q-value | Volume | Spectrum Matches |
| --- | --- | --- | --- | --- |
| 11.5998 | 4.2896 | 0.0068 | 1.0345e+08 | 1 |

**Adduct Signal Fractions**

- **Unmodified**: 1.0345e+08

#### NLN(N)ESLIDLQELGK Fuc 1 Hex 4 HexNAc 6

**Mass:**  3597.5836

(3.662e-07)

**Charge States:** 3, 4

**Start Position**: 1179

**End Position**: 1193

| Profile Score | Tandem Score | TDA q-value | Volume | Spectrum Matches |
| --- | --- | --- | --- | --- |
| 13.1729 | 4.2876 | 0.0068 | 4.8511e+08 | 4 |

**Adduct Signal Fractions**

- **Unmodified**: 4.8511e+08

#### DFGGFN(N)FSQILPDPSKPSKR Fuc 1 Hex 5 HexNAc 4 Neu5Ac 1

**Mass:**  4295.8633

(6.588e-07)

**Charge States:** 4

**Start Position**: 783

**End Position**: 803

| Profile Score | Tandem Score | TDA q-value | Volume | Spectrum Matches |
| --- | --- | --- | --- | --- |
| 11.7718 | 4.2784 | 0.0068 | 4.0236e+08 | 2 |

**Adduct Signal Fractions**

- **Unmodified**: 4.0236e+08

#### VYSSANN(N)C(C)TFEYVSQPFLMDLEGK Hex 3 HexNAc 4

**Mass:**  4096.7314

(7.934e-08)

**Charge States:** 3, 4

**Start Position**: 146

**End Position**: 170

| Profile Score | Tandem Score | TDA q-value | Volume | Spectrum Matches |
| --- | --- | --- | --- | --- |
| 14.9489 | 4.2736 | 0.0068 | 2.3896e+09 | 4 |

**Adduct Signal Fractions**

- **Unmodified**: 2.3896e+09

#### N(N)FTTAPAIC(C)HDGK @sulfate 1 Fuc 1 Hex 8 HexNAc 2

**Mass:**  3359.2696

(6.848e-03)

**Charge States:** 4

**Start Position**: 1061

**End Position**: 1074

| Profile Score | Tandem Score | TDA q-value | Volume | Spectrum Matches |
| --- | --- | --- | --- | --- |
| -4.7111 | 4.2677 | 0.0068 | 1.2043e+07 | 1 |

**Adduct Signal Fractions**

- **Na**: 1.2043e+07

#### VYSSANN(N)C(C)TFEYVSQPFLMDLEGK Fuc 1 Hex 4 HexNAc 4 Neu5Ac 1

**Mass:**  4695.9386

(6.568e-07)

**Charge States:** 3, 4

**Start Position**: 146

**End Position**: 170

| Profile Score | Tandem Score | TDA q-value | Volume | Spectrum Matches |
| --- | --- | --- | --- | --- |
| 13.8132 | 4.1984 | 0.0068 | 7.2459e+08 | 3 |

**Adduct Signal Fractions**

- **Unmodified**: 7.2459e+08

#### VYSSANN(N)C(C)TFEYVSQPFLMDLEGK Hex 4 HexNAc 4

**Mass:**  4258.7880

(9.718e-08)

**Charge States:** 3, 4

**Start Position**: 146

**End Position**: 170

| Profile Score | Tandem Score | TDA q-value | Volume | Spectrum Matches |
| --- | --- | --- | --- | --- |
| 11.1730 | 4.1605 | 0.0068 | 9.7051e+08 | 6 |

**Adduct Signal Fractions**

- **Unmodified**: 9.7051e+08

#### VYSSANN(N)C(C)TFEYVSQPFLMDLEGK Hex 6 HexNAc 2

**Mass:**  4176.7340

(6.702e-07)

**Charge States:** 3, 4

**Start Position**: 146

**End Position**: 170

| Profile Score | Tandem Score | TDA q-value | Volume | Spectrum Matches |
| --- | --- | --- | --- | --- |
| 16.2081 | 4.1501 | 0.0068 | 1.0793e+09 | 3 |

**Adduct Signal Fractions**

- **Unmodified**: 1.0793e+09

#### TPPIKDFGGFN(N)FSQILPDPSKPSK Fuc 1 Hex 4 HexNAc 4 Neu5Ac 1

**Mass:**  4514.0427

(3.499e-07)

**Charge States:** 4

**Start Position**: 778

**End Position**: 802

| Profile Score | Tandem Score | TDA q-value | Volume | Spectrum Matches |
| --- | --- | --- | --- | --- |
| 10.8007 | 4.1344 | 0.0068 | 1.7332e+09 | 2 |

**Adduct Signal Fractions**

- **Unmodified**: 1.7332e+09

#### VYSSANN(N)C(C)TFEYVSQPFLMDLEGK Fuc 1 Hex 4 HexNAc 5 Neu5Ac 1

**Mass:**  4899.0190

(1.447e-07)

**Charge States:** 4

**Start Position**: 146

**End Position**: 170

| Profile Score | Tandem Score | TDA q-value | Volume | Spectrum Matches |
| --- | --- | --- | --- | --- |
| 10.6319 | 4.1337 | 0.0068 | 8.6441e+08 | 4 |

**Adduct Signal Fractions**

- **Unmodified**: 8.6441e+08

#### VYSSANN(N)C(C)TFEYVSQPFLMDLEGK Fuc 1 Hex 6 HexNAc 5 Neu5Ac 1

**Mass:**  5223.1201

(-9.529e-07)

**Charge States:** 4

**Start Position**: 146

**End Position**: 170

| Profile Score | Tandem Score | TDA q-value | Volume | Spectrum Matches |
| --- | --- | --- | --- | --- |
| 12.6068 | 4.1300 | 0.0068 | 2.6681e+09 | 5 |

**Adduct Signal Fractions**

- **Unmodified**: 2.6681e+09

#### SSVLHSTQDLFLPFFSNVTWFHAIHVSGTN(N)GTK @sulfate 1 Hex 7 HexNAc 6

**Mass:**  6106.6824

(7.231e-06)

**Charge States:** 5

**Start Position**: 32

**End Position**: 65

| Profile Score | Tandem Score | TDA q-value | Volume | Spectrum Matches |
| --- | --- | --- | --- | --- |
| 11.3981 | 4.1142 | 0.0068 | 2.4923e+08 | 2 |

**Adduct Signal Fractions**

- **Unmodified**: 2.4923e+08

#### QC(C)VN(N)LTTR Fuc 1 Hex 6 HexNAc 3 Neu5Ac 1

**Mass:**  3035.2064

(8.642e-03)

**Start Position**: 1

**End Position**: 9

| Profile Score | Tandem Score | TDA q-value | Volume | Spectrum Matches |
| --- | --- | --- | --- | --- |
| 0.0000 | 4.0232 | 0.0110 | 0.0000e+00 | 1 |

Adduct:

No Chromatogram Found

#### QC(C)VN(N)LTTR Fuc 2 Hex 5 HexNAc 4

**Mass:**  2930.1928

(8.607e-03)

**Start Position**: 1

**End Position**: 9

| Profile Score | Tandem Score | TDA q-value | Volume | Spectrum Matches |
| --- | --- | --- | --- | --- |
| 0.0000 | 4.0130 | 0.0110 | 0.0000e+00 | 1 |

Adduct:

No Chromatogram Found

#### QC(C)VN(N)LTTR Hex 3 HexNAc 4 Neu5Ac 1

**Mass:**  2580.0524

(-3.796e-06)

**Charge States:** 2, 3

**Start Position**: 1

**End Position**: 9

| Profile Score | Tandem Score | TDA q-value | Volume | Spectrum Matches |
| --- | --- | --- | --- | --- |
| 16.6118 | 3.9575 | 0.0110 | 8.1376e+09 | 3 |

**Adduct Signal Fractions**

- **Unmodified**: 8.1376e+09

#### YNEN(N)GTITDAVDC(C)ALDPLSETK Hex 4 HexNAc 5 Neu5Ac 1

**Mass:**  4379.8008

(-3.067e-07)

**Charge States:** 3, 4

**Start Position**: 266

**End Position**: 288

| Profile Score | Tandem Score | TDA q-value | Volume | Spectrum Matches |
| --- | --- | --- | --- | --- |
| 13.4954 | 3.8767 | 0.0068 | 6.9656e+08 | 4 |

**Adduct Signal Fractions**

- **Unmodified**: 6.9656e+08

#### YNEN(N)GTITDAVDC(C)ALDPLSETK Fuc 1 Hex 5 HexNAc 6

**Mass:**  4599.8940

(-2.236e-07)

**Charge States:** 3, 4

**Start Position**: 266

**End Position**: 288

| Profile Score | Tandem Score | TDA q-value | Volume | Spectrum Matches |
| --- | --- | --- | --- | --- |
| 13.3635 | 3.8679 | 0.0068 | 2.4176e+09 | 6 |

**Adduct Signal Fractions**

- **Unmodified**: 2.4176e+09

#### YNEN(N)GTITDAVDC(C)ALDPLSETK Fuc 1 Hex 3 HexNAc 6

**Mass:**  4276.7980

(2.362e-04)

**Start Position**: 266

**End Position**: 288

| Profile Score | Tandem Score | TDA q-value | Volume | Spectrum Matches |
| --- | --- | --- | --- | --- |
| 0.0000 | 3.8672 | 0.0068 | 0.0000e+00 | 3 |

Adduct:

No Chromatogram Found

#### YNEN(N)GTITDAVDC(C)ALDPLSETK Hex 3 HexNAc 6

**Mass:**  4129.7329

(9.523e-07)

**Charge States:** 3, 4

**Start Position**: 266

**End Position**: 288

| Profile Score | Tandem Score | TDA q-value | Volume | Spectrum Matches |
| --- | --- | --- | --- | --- |
| 13.0618 | 3.8415 | 0.0068 | 1.1739e+09 | 3 |

**Adduct Signal Fractions**

- **Unmodified**: 1.1739e+09

#### YNEN(N)GTITDAVDC(C)ALDPLSETK Fuc 1 Hex 4 HexNAc 4 Neu5Ac 1

**Mass:**  4322.7754

(-2.717e-07)

**Charge States:** 3, 4

**Start Position**: 266

**End Position**: 288

| Profile Score | Tandem Score | TDA q-value | Volume | Spectrum Matches |
| --- | --- | --- | --- | --- |
| 12.2587 | 3.8073 | 0.0081 | 2.6282e+08 | 2 |

**Adduct Signal Fractions**

- **Unmodified**: 2.6282e+08

#### EGVFVSN(N)GTHWFVTQR Fuc 1 Hex 6 HexNAc 6

**Mass:**  4200.7697

(-5.390e-08)

**Charge States:** 3, 4

**Start Position**: 1079

**End Position**: 1095

| Profile Score | Tandem Score | TDA q-value | Volume | Spectrum Matches |
| --- | --- | --- | --- | --- |
| 10.0164 | 3.7507 | 0.0093 | 1.6309e+08 | 5 |

**Adduct Signal Fractions**

- **Unmodified**: 1.6309e+08

#### EGVFVSN(N)GTHWFVTQR Hex 6 HexNAc 4

**Mass:**  3647.5405

(1.570e-08)

**Charge States:** 3

**Start Position**: 1079

**End Position**: 1095

| Profile Score | Tandem Score | TDA q-value | Volume | Spectrum Matches |
| --- | --- | --- | --- | --- |
| -5.6154 | 3.7439 | 0.0093 | 3.6950e+08 | 6 |

**Adduct Signal Fractions**

- **Unmodified**: 3.6950e+08

#### N(N)FTTAPAIC(C)HDGK Fuc 2 Hex 6 HexNAc 5

**Mass:**  3710.4916

(-8.571e-08)

**Charge States:** 3, 4

**Start Position**: 1061

**End Position**: 1074

| Profile Score | Tandem Score | TDA q-value | Volume | Spectrum Matches |
| --- | --- | --- | --- | --- |
| 11.2321 | 3.7422 | 0.0093 | 3.6121e+08 | 6 |

**Adduct Signal Fractions**

- **Unmodified**: 3.6121e+08

#### N(N)FTTAPAIC(C)HDGK Hex 6 HexNAc 6 Neu5Ac 2

**Mass:**  4223.6779

(4.766e-03)

**Start Position**: 1061

**End Position**: 1074

| Profile Score | Tandem Score | TDA q-value | Volume | Spectrum Matches |
| --- | --- | --- | --- | --- |
| 0.0000 | 3.7400 | 0.0093 | 0.0000e+00 | 1 |

Adduct:

No Chromatogram Found

#### DFGGFN(N)FSQILPDPSKPSK Fuc 1 Hex 4 HexNAc 5

**Mass:**  3891.7011

(5.164e-04)

**Start Position**: 783

**End Position**: 802

| Profile Score | Tandem Score | TDA q-value | Volume | Spectrum Matches |
| --- | --- | --- | --- | --- |
| 0.0000 | 3.7335 | 0.0099 | 0.0000e+00 | 1 |

Adduct:

No Chromatogram Found

#### N(N)FTTAPAIC(C)HDGK Fuc 1 Hex 6 HexNAc 5 Neu5Ac 2

**Mass:**  4148.6281

(4.834e-04)

**Start Position**: 1061

**End Position**: 1074

| Profile Score | Tandem Score | TDA q-value | Volume | Spectrum Matches |
| --- | --- | --- | --- | --- |
| 0.0000 | 3.7255 | 0.0104 | 0.0000e+00 | 1 |

Adduct:

No Chromatogram Found

#### EGVFVSN(N)GTHWFVTQR Fuc 1 Hex 3 HexNAc 6

**Mass:**  3714.5991

(2.693e-04)

**Start Position**: 1079

**End Position**: 1095

| Profile Score | Tandem Score | TDA q-value | Volume | Spectrum Matches |
| --- | --- | --- | --- | --- |
| 0.0000 | 3.7217 | 0.0104 | 0.0000e+00 | 1 |

Adduct:

No Chromatogram Found

#### N(N)FTTAPAIC(C)HDGK Fuc 2 Hex 5 HexNAc 3

**Mass:**  3142.2835

(3.188e-04)

**Charge States:** 3

**Start Position**: 1061

**End Position**: 1074

| Profile Score | Tandem Score | TDA q-value | Volume | Spectrum Matches |
| --- | --- | --- | --- | --- |
| -22.3571 | 3.7208 | 0.0104 | 7.3855e+07 | 2 |

**Adduct Signal Fractions**

- **Unmodified**: 7.3855e+07

#### TQSLLIVNN(N)ATNVVIK Hex 8 HexNAc 2

**Mass:**  3428.5791

(-3.423e-08)

**Charge States:** 3

**Start Position**: 101

**End Position**: 117

| Profile Score | Tandem Score | TDA q-value | Volume | Spectrum Matches |
| --- | --- | --- | --- | --- |
| 11.5515 | 3.7186 | 0.0104 | 5.3099e+08 | 2 |

**Adduct Signal Fractions**

- **Unmodified**: 5.3099e+08

#### DFGGFN(N)FSQILPDPSKPSK Fuc 2 Hex 4 HexNAc 4

**Mass:**  3832.6742

(1.574e-06)

**Charge States:** 3, 4

**Start Position**: 783

**End Position**: 802

| Profile Score | Tandem Score | TDA q-value | Volume | Spectrum Matches |
| --- | --- | --- | --- | --- |
| 9.6799 | 3.7134 | 0.0109 | 5.2850e+08 | 3 |

**Adduct Signal Fractions**

- **Unmodified**: 5.2850e+08

#### SSVLHSTQDLFLPFFSNVTWFHAIHVSGTN(N)GTK @sulfate 1 Fuc 2 Hex 9 HexNAc 6

**Mass:**  6722.9142

(2.541e-03)

**Charge States:** 5

**Start Position**: 32

**End Position**: 65

| Profile Score | Tandem Score | TDA q-value | Volume | Spectrum Matches |
| --- | --- | --- | --- | --- |
| -19.2793 | 3.7073 | 0.0109 | 2.8449e+07 | 1 |

**Adduct Signal Fractions**

- **NH3**: 2.8449e+07

#### N(N)FTTAPAIC(C)HDGK Fuc 2 Hex 5 HexNAc 4

**Mass:**  3347.3626

(5.990e-04)

**Start Position**: 1061

**End Position**: 1074

| Profile Score | Tandem Score | TDA q-value | Volume | Spectrum Matches |
| --- | --- | --- | --- | --- |
| 0.0000 | 3.7069 | 0.0109 | 0.0000e+00 | 1 |

Adduct:

No Chromatogram Found

#### FPNITNLC(C)PFGEVFN(N)ATR Fuc 2 Hex 8 HexNAc 8

**Mass:**  5309.1924

(4.399e-07)

**Charge States:** 4

**Start Position**: 316

**End Position**: 334

| Profile Score | Tandem Score | TDA q-value | Volume | Spectrum Matches |
| --- | --- | --- | --- | --- |
| 14.1174 | 3.7053 | 0.0109 | 1.4696e+09 | 2 |

**Adduct Signal Fractions**

- **Unmodified**: 1.4696e+09

#### TQSLLIVNN(N)ATNVVIK Hex 8 HexNAc 2

**Mass:**  3429.5824

(2.923e-04)

**Start Position**: 101

**End Position**: 117

| Profile Score | Tandem Score | TDA q-value | Volume | Spectrum Matches |
| --- | --- | --- | --- | --- |
| 0.0000 | 3.6968 | 0.0109 | 0.0000e+00 | 1 |

Adduct:

No Chromatogram Found

#### N(N)FTTAPAIC(C)HDGK Fuc 2 Hex 5 HexNAc 4 Neu5Ac 1

**Mass:**  3637.4774

(2.761e-04)

**Charge States:** 3

**Start Position**: 1061

**End Position**: 1074

| Profile Score | Tandem Score | TDA q-value | Volume | Spectrum Matches |
| --- | --- | --- | --- | --- |
| 8.0892 | 3.6805 | 0.0115 | 2.4808e+06 | 1 |

**Adduct Signal Fractions**

- **Unmodified**: 2.4808e+06

#### DFGGFN(N)FSQILPDPSKPSK @sulfate 1 Hex 7 HexNAc 3

**Mass:**  3903.6033

(3.128e-06)

**Charge States:** 3, 4

**Start Position**: 783

**End Position**: 802

| Profile Score | Tandem Score | TDA q-value | Volume | Spectrum Matches |
| --- | --- | --- | --- | --- |
| 10.7818 | 3.6770 | 0.0121 | 1.1898e+08 | 3 |

**Adduct Signal Fractions**

- **Unmodified**: 1.1898e+08

#### EGVFVSN(N)GTHWFVTQR Hex 9 HexNAc 2

**Mass:**  3727.5724

(-8.155e-07)

**Charge States:** 3

**Start Position**: 1079

**End Position**: 1095

| Profile Score | Tandem Score | TDA q-value | Volume | Spectrum Matches |
| --- | --- | --- | --- | --- |
| -5.2837 | 3.6600 | 0.0125 | 3.1443e+07 | 1 |

**Adduct Signal Fractions**

- **Unmodified**: 3.1443e+07

#### TQSLLIVNN(N)ATNVVIK @sulfate 1 Hex 5 HexNAc 3

**Mass:**  3226.4286

(3.101e-04)

**Charge States:** 3

**Start Position**: 101

**End Position**: 117

| Profile Score | Tandem Score | TDA q-value | Volume | Spectrum Matches |
| --- | --- | --- | --- | --- |
| -20.9956 | 3.6590 | 0.0125 | 4.3700e+06 | 1 |

**Adduct Signal Fractions**

- **Unmodified**: 4.3700e+06

#### DFGGFN(N)FSQILPDPSKPSKRSFIEDLLFNK Hex 6 HexNAc 3

**Mass:**  5043.3293

(3.785e-03)

**Start Position**: 783

**End Position**: 813

| Profile Score | Tandem Score | TDA q-value | Volume | Spectrum Matches |
| --- | --- | --- | --- | --- |
| 0.0000 | 3.6557 | 0.0125 | 0.0000e+00 | 1 |

Adduct:

No Chromatogram Found

#### EGVFVSN(N)GTHWFVTQR Fuc 1 Hex 5 HexNAc 6 Neu5Ac 2

**Mass:**  4619.8957

(4.846e-08)

**Start Position**: 1079

**End Position**: 1095

| Profile Score | Tandem Score | TDA q-value | Volume | Spectrum Matches |
| --- | --- | --- | --- | --- |
| 0.0000 | 3.6536 | 0.0125 | 0.0000e+00 | 1 |

Adduct:

No Chromatogram Found

#### TQSLLIVNN(N)ATNVVIK Hex 4 HexNAc 5 Neu5Ac 1

**Mass:**  3680.7046

(5.934e-07)

**Charge States:** 3, 4

**Start Position**: 101

**End Position**: 117

| Profile Score | Tandem Score | TDA q-value | Volume | Spectrum Matches |
| --- | --- | --- | --- | --- |
| 9.6897 | 3.6451 | 0.0133 | 3.4567e+08 | 5 |

**Adduct Signal Fractions**

- **Unmodified**: 3.4567e+08

#### N(N)FTTAPAIC(C)HDGK Fuc 3 Hex 6 HexNAc 5 Neu5Ac 1

**Mass:**  4147.6241

(-4.618e-06)

**Charge States:** 4

**Start Position**: 1061

**End Position**: 1074

| Profile Score | Tandem Score | TDA q-value | Volume | Spectrum Matches |
| --- | --- | --- | --- | --- |
| -5.6164 | 3.6447 | 0.0133 | 3.9691e+07 | 3 |

**Adduct Signal Fractions**

- **Unmodified**: 3.9691e+07

#### N(N)FTTAPAIC(C)HDGK Hex 9 HexNAc 2

**Mass:**  3295.2967

(2.518e-07)

**Charge States:** 3

**Start Position**: 1061

**End Position**: 1074

| Profile Score | Tandem Score | TDA q-value | Volume | Spectrum Matches |
| --- | --- | --- | --- | --- |
| 14.1882 | 3.6434 | 0.0133 | 1.4284e+08 | 3 |

**Adduct Signal Fractions**

- **Unmodified**: 1.4284e+08

#### DFGGFN(N)FSQILPDPSKPSK Hex 5 HexNAc 3

**Mass:**  3522.5136

(6.568e-03)

**Start Position**: 783

**End Position**: 802

| Profile Score | Tandem Score | TDA q-value | Volume | Spectrum Matches |
| --- | --- | --- | --- | --- |
| 0.0000 | 3.6419 | 0.0133 | 0.0000e+00 | 1 |

Adduct:

No Chromatogram Found
