## Supplementary material for "Assignment of coronavirus spike protein site-specific glycosylation using GlycReSoft": 210320_SARS2_S_chymotrypsin.html

### 210320\_SARS\_CoV\_2\_C @ GlycopeptideHypothesis-de2712e9621044c8ae9c543abd7c44a3

**MS^1^ Mass Accuracy:** 10.0 PPM  
**Grouping Tolerance:** 15.0 PPM  
**MS^n^ Mass Accuracy:** 10.0 PPM  
**Minimum `q`-Value For Clustering:** 0.05

**Glycopeptide Hypothesis:**
GlycopeptideHypothesis-de2712e9621044c8ae9c543abd7c44a3  
**Sample Searched:**
210320\_SARS\_CoV\_2\_C

| Protein | Matched Glycopeptides |
| --- | --- |
| sp\|P0DTC2\|SPIKE\_SARS2 Spike glycoprotein OS=Severe acute respiratory syndrome coronavirus 2 OX=2697049 GN=S PE=1 SV=1 | 477 |

#### sp|P0DTC2|SPIKE\_SARS2 Spike glycoprotein OS=Severe acute respiratory syndrome coronavirus 2 OX=2697049 GN=S PE=1 SV=1

**>sp|P0DTC2|SPIKE\_SARS2 Spike glycoprotein OS=Severe acute respiratory syndrome coronavirus 2 OX=2697049 GN=S PE=1 SV=1**  
 `MFVFLVLLPLVSSQCVNLTTRTQLPPAYTNSFTRGVYYPDKVFRSSVLHSTQDLFLPFFSNVTWFHAIHVSGTNGTKRFD  
NPVLPFNDGVYFASTEKSNIIRGWIFGTTLDSKTQSLLIVNNATNVVIKVCEFQFCNDPFLGVYYHKNNKSWMESEFRVY  
SSANNCTFEYVSQPFLMDLEGKQGNFKNLREFVFKNIDGYFKIYSKHTPINLVRDLPQGFSALEPLVDLPIGINITRFQT  
LLALHRSYLTPGDSSSGWTAGAAAYYVGYLQPRTFLLKYNENGTITDAVDCALDPLSETKCTLKSFTVEKGIYQTSNFRV  
QPTESIVRFPNITNLCPFGEVFNATRFASVYAWNRKRISNCVADYSVLYNSASFSTFKCYGVSPTKLNDLCFTNVYADSF  
VIRGDEVRQIAPGQTGKIADYNYKLPDDFTGCVIAWNSNNLDSKVGGNYNYLYRLFRKSNLKPFERDISTEIYQAGSTPC  
NGVEGFNCYFPLQSYGFQPTNGVGYQPYRVVVLSFELLHAPATVCGPKKSTNLVKNKCVNFNFNGLTGTGVLTESNKKFL  
PFQQFGRDIADTTDAVRDPQTLEILDITPCSFGGVSVITPGTNTSNQVAVLYQDVNCTEVPVAIHADQLTPTWRVYSTGS  
NVFQTRAGCLIGAEHVNNSYECDIPIGAGICASYQTQTNSPGSASSVASQSIIAYTMSLGAENSVAYSNNSIAIPTNFTI  
SVTTEILPVSMTKTSVDCTMYICGDSTECSNLLLQYGSFCTQLNRALTGIAVEQDKNTQEVFAQVKQIYKTPPIKDFGGF  
NFSQILPDPSKPSKRSFIEDLLFNKVTLADAGFIKQYGDCLGDIAARDLICAQKFNGLTVLPPLLTDEMIAQYTSALLAG  
TITSGWTFGAGAALQIPFAMQMAYRFNGIGVTQNVLYENQKLIANQFNSAIGKIQDSLSSTASALGKLQDVVNQNAQALN  
TLVKQLSSNFGAISSVLNDILSRLDPPEAEVQIDRLITGRLQSLQTYVTQQLIRAAEIRASANLAATKMSECVLGQSKRV  
DFCGKGYHLMSFPQSAPHGVVFLHVTYVPAQEKNFTTAPAICHDGKAHFPREGVFVSNGTHWFVTQRNFYEPQIITTDNT  
FVSGNCDVVIGIVNNTVYDPLQPELDSFKEELDKYFKNHTSPDVDLGDISGINASVVNIQKEIDRLNEVAKNLNESLIDL  
QELGKYEQYIKWPWYIWLGFIAGLIAIVMVTIMLCCMTSCCSCLKGCCSCGSCCKFDEDDSEPVLKGVKLHYT`

| Site | Type | Count |
| --- | --- | --- |
| 61 | N-linked | 1 |
| 74 | N-linked | 144 |
| 122 | N-linked | 47 |
| 149 | N-linked | 28 |
| 165 | N-linked | 1 |
| 234 | N-linked | 25 |
| 331 | N-linked | 58 |
| 343 | N-linked | 13 |
| 657 | N-linked | 18 |
| 709 | N-linked | 16 |
| 717 | N-linked | 18 |
| 801 | N-linked | 41 |
| 1074 | N-linked | 54 |
| 1098 | N-linked | 29 |
| 1134 | N-linked | 1 |

| Mass | Sequence | MS_2_ Score | `q`-value | MS_1_ Score | Signal | Protein Coordinates | Tandem Scans | Start Time | Apex Time | End Time |
| --- | --- | --- | --- | --- | --- | --- | --- | --- | --- | --- |
| 3333.4595 | HAIHVSGTN(N)GTKRF Fuc 1 Hex 4 HexNAc 5 | 165.934 | 0.002 | 13.716 | 4.356e+10 | 65 - 79 | 66 | 52.26 | 54.65 | 63.19 |
| 3495.5188 | HAIHVSGTN(N)GTKRF Fuc 1 Hex 5 HexNAc 5 | 161.396 | 0.002 | 16.129 | 9.884e+09 | 65 - 79 | 46 | 52.70 | 55.88 | 57.96 |
| 3276.4460 | HAIHVSGTN(N)GTKRF Fuc 2 Hex 4 HexNAc 4 | 160.342 | 0.002 | 16.242 | 7.950e+09 | 65 - 79 | 23 | 52.54 | 53.83 | 57.82 |
| 2968.3328 | HAIHVSGTN(N)GTKRF Fuc 1 Hex 3 HexNAc 4 | 158.254 | 0.002 | 17.792 | 4.959e+10 | 65 - 79 | 58 | 52.98 | 54.81 | 63.30 |
| 3413.4148 | HAIHVSGTN(N)GTKRF @sulfate 1 Fuc 1 Hex 4 HexNAc 5 | 157.377 | 0.002 | 16.458 | 1.446e+10 | 65 - 79 | 21 | 62.36 | 62.95 | 67.64 |
| 3770.6127 | HAIHVSGTN(N)GTKRF Fuc 2 Hex 4 HexNAc 5 Neu5Ac 1 | 157.183 | 0.002 | 16.738 | 5.940e+10 | 65 - 79 | 55 | 60.53 | 62.97 | 67.71 |
| 3704.5155 | HAIHVSGTN(N)GTKRF @sulfate 1 Fuc 1 Hex 4 HexNAc 5 Neu5Ac 1 | 155.558 | 0.002 | 17.966 | 3.811e+10 | 65 - 79 | 26 | 72.26 | 73.59 | 78.07 |
| 3454.4471 | HAIHVSGTN(N)GTKRF @sulfate 1 Fuc 1 Hex 3 HexNAc 6 | 154.558 | 0.002 | 16.563 | 2.603e+10 | 65 - 79 | 25 | 62.43 | 63.35 | 67.60 |
| 3479.5201 | HAIHVSGTN(N)GTKRF Fuc 2 Hex 4 HexNAc 5 | 153.875 | 0.002 | 17.118 | 7.365e+10 | 65 - 79 | 67 | 52.09 | 53.83 | 63.21 |
| 3682.8576 | HAIHVSGTN(N)GTKRF Fuc 2 Hex 4 HexNAc 6 | 151.105 | 0.002 | 6.666 | 3.861e+09 | 65 - 79 | 37 | 52.51 | 53.09 | 170.29 |
| 3624.5606 | HAIHVSGTN(N)GTKRF Fuc 1 Hex 4 HexNAc 5 Neu5Ac 1 | 146.832 | 0.002 | 14.868 | 3.373e+10 | 65 - 79 | 53 | 60.59 | 61.66 | 67.75 |
| 3171.4114 | HAIHVSGTN(N)GTKRF Fuc 1 Hex 3 HexNAc 5 | 146.773 | 0.002 | 14.769 | 8.776e+10 | 65 - 79 | 68 | 53.09 | 54.90 | 62.87 |
| 3130.3775 | HAIHVSGTN(N)GTKRF Fuc 1 Hex 4 HexNAc 4 | 146.556 | 0.002 | 15.364 | 9.342e+09 | 65 - 79 | 35 | 52.87 | 54.80 | 63.04 |
| 3844.9121 | HAIHVSGTN(N)GTKRF Fuc 2 Hex 5 HexNAc 6 | 144.406 | 0.002 | -5.968 | 4.465e+09 | 65 - 79 | 24 | 52.03 | 52.75 | 169.24 |
| 3251.3676 | HAIHVSGTN(N)GTKRF @sulfate 1 Fuc 1 Hex 3 HexNAc 5 | 143.811 | 0.002 | 18.266 | 2.535e+10 | 65 - 79 | 26 | 62.77 | 63.49 | 67.68 |
| 3438.4928 | HAIHVSGTN(N)GTKRF Fuc 2 Hex 5 HexNAc 4 | 141.152 | 0.002 | 12.785 | 3.092e+09 | 65 - 79 | 21 | 52.13 | 53.14 | 55.95 |
| 3221.4126 | VDLPIGIN(N)ITRF Hex 9 HexNAc 2 | 140.583 | 0.002 | 17.613 | 8.530e+08 | 226 - 238 | 2 | 202.81 | 203.18 | 203.66 |
| 2740.2128 | HAIHVSGTN(N)GTKRF Hex 5 HexNAc 2 | 140.451 | 0.002 | 16.908 | 6.511e+08 | 65 - 79 | 3 | 52.28 | 53.14 | 53.94 |
| 3536.5411 | HAIHVSGTN(N)GTKRF Fuc 1 Hex 4 HexNAc 6 | 139.578 | 0.002 | 13.720 | 6.951e+09 | 65 - 79 | 37 | 52.42 | 53.72 | 57.99 |
| 3874.6226 | HAIHVSGTN(N)GTKRF Fuc 1 Hex 5 HexNAc 4 Neu5Ac 2 | 139.529 | 0.002 | 14.510 | 6.704e+09 | 65 - 79 | 16 | 62.70 | 72.16 | 73.40 |
| 3210.3360 | HAIHVSGTN(N)GTKRF @sulfate 1 Fuc 1 Hex 4 HexNAc 4 | 139.252 | 0.002 | 14.313 | 1.301e+09 | 65 - 79 | 4 | 62.74 | 63.21 | 64.23 |
| 3739.6166 | HAIHVSGTN(N)GTKRF Fuc 1 Hex 4 HexNAc 7 | 137.971 | 0.002 | 14.441 | 7.625e+09 | 65 - 79 | 36 | 52.77 | 53.68 | 57.79 |
| 3374.4890 | HAIHVSGTN(N)GTKRF Fuc 1 Hex 3 HexNAc 6 | 137.529 | 0.002 | 13.795 | 2.752e+10 | 65 - 79 | 57 | 52.80 | 54.93 | 61.98 |
| 3372.3974 | HAIHVSGTN(N)GTKRF @sulfate 1 Fuc 1 Hex 5 HexNAc 4 | 137.358 | 0.002 | 14.594 | 4.433e+08 | 65 - 79 | 5 | 62.19 | 62.48 | 63.13 |
| 2484.0441 | GEVFN(N)ATRF Fuc 1 Hex 3 HexNAc 4 | 135.956 | 0.008 | 17.393 | 5.146e+09 | 338 - 347 | 4 | 124.20 | 124.47 | 124.81 |
| 3559.4805 | HAIHVSGTN(N)GTKRF @sulfate 1 Fuc 2 Hex 4 HexNAc 5 | 135.942 | 0.002 | 14.229 | 7.097e+09 | 65 - 79 | 23 | 61.77 | 62.05 | 66.58 |
| 2982.2453 | HAIHVSGTN(N)GTKRF @sulfate 1 Hex 6 HexNAc 2 | 133.753 | 0.002 | 15.660 | 5.706e+08 | 65 - 79 | 3 | 62.13 | 62.55 | 63.01 |
| 3885.6795 | HAIHVSGTN(N)GTKRF Fuc 2 Hex 4 HexNAc 7 | 132.565 | 0.002 | 14.555 | 6.377e+09 | 65 - 79 | 38 | 52.39 | 53.05 | 57.60 |
| 3048.2894 | HAIHVSGTN(N)GTKRF @sulfate 1 Fuc 1 Hex 3 HexNAc 4 | 130.044 | 0.002 | 15.031 | 1.463e+09 | 65 - 79 | 8 | 63.42 | 63.65 | 65.09 |
| 3185.3232 | HAIHVSGTN(N)GTKRF @sulfate 1 Hex 6 HexNAc 3 | 129.754 | 0.002 | 16.269 | 4.322e+08 | 65 - 79 | 2 | 61.71 | 61.88 | 62.67 |
| 3421.4804 | HAIHVSGTN(N)GTKRF Fuc 1 Hex 4 HexNAc 4 Neu5Ac 1 | 129.587 | 0.002 | 13.261 | 5.916e+09 | 65 - 79 | 27 | 60.93 | 62.48 | 65.32 |
| 3948.7288 | HAIHVSGTN(N)GTKRF Fuc 1 Hex 6 HexNAc 5 Neu5Ac 1 | 129.334 | 0.002 | 12.952 | 2.351e+09 | 65 - 79 | 23 | 51.84 | 60.84 | 62.74 |
| 2927.2979 | HAIHVSGTN(N)GTKRF Fuc 1 Hex 4 HexNAc 3 | 128.156 | 0.002 | 15.118 | 2.392e+09 | 65 - 79 | 11 | 53.47 | 53.81 | 56.24 |
| 3099.2461 | GEVFN(N)ATRF Fuc 1 Hex 5 HexNAc 4 Neu5Ac 1 | 127.348 | 0.008 | 17.963 | 3.025e+09 | 338 - 347 | 8 | 143.39 | 143.78 | 145.62 |
| 2147.9002 | VPAQEKN(N)F Hex 5 HexNAc 2 | 126.457 | 0.008 | 19.916 | 6.033e+10 | 1067 - 1075 | 23 | 66.45 | 67.87 | 71.99 |
| 2309.9523 | VPAQEKN(N)F Hex 6 HexNAc 2 | 126.426 | 0.008 | 16.804 | 1.862e+09 | 1067 - 1075 | 10 | 66.42 | 66.85 | 68.56 |
| 3989.6921 | HAIHVSGTN(N)GTKRF Fuc 1 Hex 5 HexNAc 6 Neu5Ac 1 | 126.353 | 0.002 | 12.507 | 6.594e+09 | 65 - 79 | 21 | 60.39 | 61.12 | 65.99 |
| 3965.6398 | HAIHVSGTN(N)GTKRF @sulfate 1 Fuc 2 Hex 4 HexNAc 7 | 125.111 | 0.002 | 14.938 | 1.077e+09 | 65 - 79 | 6 | 61.46 | 62.72 | 63.19 |
| 3059.3581 | VDLPIGIN(N)ITRF Hex 8 HexNAc 2 | 123.508 | 0.002 | 16.319 | 1.623e+09 | 226 - 238 | 3 | 203.06 | 203.63 | 203.95 |
| 3729.5933 | HAIHVSGTN(N)GTKRF Fuc 2 Hex 5 HexNAc 4 Neu5Ac 1 | 121.193 | 0.002 | 11.605 | 1.364e+09 | 65 - 79 | 12 | 60.45 | 61.20 | 63.12 |
| 3698.6006 | HAIHVSGTN(N)GTKRF Fuc 1 Hex 5 HexNAc 6 | 119.536 | 0.002 | 14.059 | 4.178e+09 | 65 - 79 | 26 | 52.49 | 53.47 | 57.84 |
| 4135.7524 | HAIHVSGTN(N)GTKRF Fuc 2 Hex 5 HexNAc 6 Neu5Ac 1 | 119.361 | 0.002 | 12.976 | 4.277e+09 | 65 - 79 | 25 | 60.06 | 61.65 | 64.47 |
| 3292.4406 | HAIHVSGTN(N)GTKRF Fuc 1 Hex 5 HexNAc 4 | 119.167 | 0.002 | 14.056 | 2.492e+09 | 65 - 79 | 33 | 52.06 | 54.51 | 62.55 |
| 3641.5703 | HAIHVSGTN(N)GTKRF Fuc 2 Hex 5 HexNAc 5 | 117.143 | 0.002 | 14.557 | 4.239e+09 | 65 - 79 | 33 | 52.39 | 56.23 | 57.75 |
| 3657.5262 | HAIHVSGTN(N)GTKRF @sulfate 1 Fuc 1 Hex 3 HexNAc 7 | 116.229 | 0.002 | 12.194 | 1.856e+09 | 65 - 79 | 14 | 62.31 | 63.80 | 65.32 |
| 3577.5666 | HAIHVSGTN(N)GTKRF Fuc 1 Hex 3 HexNAc 7 | 115.985 | 0.002 | 13.333 | 9.557e+09 | 65 - 79 | 45 | 53.26 | 54.49 | 57.90 |
| 3860.6514 | HAIHVSGTN(N)GTKRF Fuc 1 Hex 6 HexNAc 6 | 115.446 | 0.002 | 13.565 | 8.901e+08 | 65 - 79 | 6 | 53.20 | 53.66 | 57.60 |
| 3786.6103 | HAIHVSGTN(N)GTKRF Fuc 1 Hex 5 HexNAc 5 Neu5Ac 1 | 115.393 | 0.002 | 11.591 | 5.587e+09 | 65 - 79 | 30 | 60.72 | 62.94 | 66.01 |
| 2350.9790 | VPAQEKN(N)F Hex 5 HexNAc 3 | 114.707 | 0.008 | 17.059 | 5.201e+09 | 1067 - 1075 | 15 | 67.05 | 67.50 | 70.59 |
| 2513.0305 | VPAQEKN(N)F Hex 6 HexNAc 3 | 114.700 | 0.008 | 17.572 | 2.684e+09 | 1067 - 1075 | 8 | 66.85 | 67.31 | 68.91 |
| 2334.9836 | VPAQEKN(N)F Fuc 1 Hex 4 HexNAc 3 | 114.532 | 0.008 | 13.644 | 3.556e+09 | 1067 - 1075 | 13 | 69.41 | 70.55 | 71.90 |
| 4176.7707 | HAIHVSGTN(N)GTKRF Fuc 2 Hex 4 HexNAc 7 Neu5Ac 1 | 113.549 | 0.002 | 15.083 | 4.991e+09 | 65 - 79 | 19 | 60.18 | 61.96 | 63.12 |
| 2808.1490 | GEVFN(N)ATRF Fuc 1 Hex 5 HexNAc 4 | 113.090 | 0.008 | 16.495 | 1.169e+10 | 338 - 347 | 10 | 122.33 | 122.57 | 124.41 |
| 2954.2082 | GEVFN(N)ATRF Fuc 2 Hex 5 HexNAc 4 | 106.943 | 0.008 | 14.840 | 6.543e+08 | 338 - 347 | 3 | 121.31 | 121.71 | 121.95 |
| 2697.0427 | VSN(N)GTHW Fuc 1 Hex 4 HexNAc 4 Neu5Ac 1 | 106.508 | 0.008 | 15.181 | 3.210e+09 | 1095 - 1102 | 11 | 83.25 | 83.54 | 85.80 |
| 2646.0960 | GEVFN(N)ATRF Fuc 1 Hex 4 HexNAc 4 | 105.506 | 0.008 | 15.985 | 3.847e+09 | 338 - 347 | 10 | 122.98 | 123.97 | 124.47 |
| 2849.1781 | GEVFN(N)ATRF Fuc 1 Hex 4 HexNAc 5 | 104.620 | 0.008 | 14.976 | 7.886e+09 | 338 - 347 | 11 | 122.56 | 123.67 | 124.77 |
| 2700.1167 | VPAQEKN(N)F Fuc 1 Hex 5 HexNAc 4 | 103.650 | 0.008 | 17.247 | 4.362e+09 | 1067 - 1075 | 13 | 68.79 | 69.67 | 71.90 |
| 2188.9265 | VPAQEKN(N)F Hex 4 HexNAc 3 | 103.541 | 0.008 | 15.236 | 2.955e+09 | 1067 - 1075 | 9 | 68.36 | 68.81 | 70.53 |
| 2405.9490 | VSN(N)GTHW Fuc 1 Hex 4 HexNAc 4 | 103.485 | 0.008 | 12.944 | 7.952e+08 | 1095 - 1102 | 10 | 68.62 | 69.24 | 69.85 |
| 3780.6700 | HAIHVSGTN(N)GTKRF Fuc 1 Hex 3 HexNAc 8 | 102.334 | 0.002 | 13.990 | 3.846e+09 | 65 - 79 | 22 | 53.14 | 53.97 | 56.05 |
| 2991.2097 | VPAQEKN(N)F Fuc 1 Hex 5 HexNAc 4 Neu5Ac 1 | 101.580 | 0.008 | 18.281 | 3.168e+09 | 1067 - 1075 | 9 | 82.67 | 83.09 | 85.07 |
| 2687.1231 | GEVFN(N)ATRF Fuc 1 Hex 3 HexNAc 5 | 100.540 | 0.008 | 16.783 | 5.260e+09 | 338 - 347 | 7 | 123.67 | 124.56 | 124.84 |
| 2446.9766 | VSN(N)GTHW Fuc 1 Hex 3 HexNAc 5 | 97.590 | 0.008 | 15.574 | 2.917e+08 | 1095 - 1102 | 4 | 69.50 | 69.82 | 70.10 |
| 2300.9185 | VSN(N)GTHW Hex 3 HexNAc 5 | 97.292 | 0.008 | 16.436 | 4.283e+08 | 1095 - 1102 | 4 | 68.20 | 68.45 | 68.62 |
| 2835.0630 | VSN(N)GTHW Hex 7 HexNAc 3 Neu5Ac 1 | 95.121 | 0.008 | 9.626 | 5.791e+06 | 1095 - 1102 | 1 | 66.88 | 66.99 | 67.09 |
| 2859.0958 | VSN(N)GTHW Fuc 1 Hex 5 HexNAc 4 Neu5Ac 1 | 95.119 | 0.008 | 17.739 | 2.924e+09 | 1095 - 1102 | 10 | 81.94 | 82.28 | 84.88 |
| 2900.1235 | VSN(N)GTHW Fuc 1 Hex 4 HexNAc 5 Neu5Ac 1 | 94.976 | 0.008 | 18.439 | 4.556e+08 | 1095 - 1102 | 4 | 84.71 | 85.01 | 85.32 |
| 2788.1333 | VPAQEKN(N)F Fuc 1 Hex 5 HexNAc 3 Neu5Ac 1 | 94.923 | 0.008 | 17.102 | 1.829e+09 | 1067 - 1075 | 6 | 83.57 | 84.06 | 84.92 |
| 3065.2487 | VPAQEKN(N)F Fuc 1 Hex 6 HexNAc 5 | 94.917 | 0.008 | 17.029 | 1.940e+09 | 1067 - 1075 | 10 | 66.99 | 67.41 | 69.89 |
| 2497.0358 | VPAQEKN(N)F Fuc 1 Hex 5 HexNAc 3 | 94.753 | 0.008 | 18.073 | 2.403e+09 | 1067 - 1075 | 12 | 68.79 | 70.16 | 71.49 |
| 2579.0891 | VPAQEKN(N)F Fuc 1 Hex 3 HexNAc 5 | 94.632 | 0.008 | 15.990 | 1.332e+10 | 1067 - 1075 | 32 | 69.46 | 69.92 | 75.21 |
| 2026.8728 | VPAQEKN(N)F Hex 3 HexNAc 3 | 94.463 | 0.008 | 17.057 | 1.135e+09 | 1067 - 1075 | 9 | 69.29 | 69.71 | 70.70 |
| 2656.0171 | VSN(N)GTHW Fuc 1 Hex 5 HexNAc 3 Neu5Ac 1 | 94.411 | 0.008 | 17.864 | 1.012e+09 | 1095 - 1102 | 8 | 82.85 | 83.16 | 85.07 |
| 2509.9596 | VSN(N)GTHW Hex 5 HexNAc 3 Neu5Ac 1 | 91.533 | 0.008 | 16.243 | 1.571e+09 | 1095 - 1102 | 7 | 82.11 | 84.19 | 84.50 |
| 2650.0563 | VSN(N)GTHW Fuc 1 Hex 3 HexNAc 6 | 91.047 | 0.008 | 15.750 | 6.934e+08 | 1095 - 1102 | 6 | 67.88 | 68.14 | 68.42 |
| 2376.0107 | VPAQEKN(N)F Fuc 1 Hex 3 HexNAc 4 | 90.924 | 0.008 | 16.998 | 1.451e+10 | 1067 - 1075 | 18 | 70.63 | 71.08 | 73.48 |
| 3006.1665 | VSN(N)GTHWF Fuc 1 Hex 5 HexNAc 4 Neu5Ac 1 | 90.741 | 0.008 | 13.606 | 5.382e+09 | 1095 - 1103 | 10 | 135.96 | 136.29 | 138.34 |
| 2829.1586 | VPAQEKN(N)F Fuc 1 Hex 4 HexNAc 4 Neu5Ac 1 | 88.678 | 0.008 | 15.982 | 3.158e+09 | 1067 - 1075 | 6 | 84.00 | 84.44 | 85.32 |
| 2642.0735 | VPAQEKN(N)F Hex 5 HexNAc 3 Neu5Ac 1 | 88.412 | 0.008 | 17.022 | 1.273e+09 | 1067 - 1075 | 4 | 82.28 | 82.63 | 83.29 |
| 2365.9313 | VSN(N)GTHWF Hex 5 HexNAc 3 | 88.156 | 0.008 | 13.519 | 3.595e+09 | 1095 - 1103 | 9 | 116.07 | 116.44 | 117.36 |
| 3032.2392 | VPAQEKN(N)F Fuc 1 Hex 4 HexNAc 5 Neu5Ac 1 | 87.700 | 0.008 | 15.816 | 1.027e+09 | 1067 - 1075 | 6 | 82.96 | 83.50 | 84.94 |
| 2097.8385 | VSN(N)GTHW Hex 3 HexNAc 4 | 87.127 | 0.008 | 17.276 | 9.201e+08 | 1095 - 1102 | 5 | 68.20 | 68.49 | 69.03 |
| 2844.1138 | VSN(N)GTHWF Fuc 1 Hex 4 HexNAc 4 Neu5Ac 1 | 86.188 | 0.008 | 13.926 | 8.286e+09 | 1095 - 1103 | 9 | 138.04 | 138.33 | 140.04 |
| 2957.1455 | VSN(N)GTHW Hex 4 HexNAc 6 Neu5Ac 1 | 85.918 | 0.008 | 16.683 | 3.715e+08 | 1095 - 1102 | 3 | 82.57 | 83.05 | 83.37 |
| 2903.1978 | VPAQEKN(N)F Fuc 1 Hex 5 HexNAc 5 | 85.538 | 0.008 | 12.864 | 2.674e+09 | 1067 - 1075 | 14 | 67.18 | 67.57 | 70.87 |
| 2553.0158 | VSN(N)GTHWF Fuc 1 Hex 4 HexNAc 4 | 85.028 | 0.008 | 16.674 | 9.007e+08 | 1095 - 1103 | 7 | 115.55 | 116.29 | 116.80 |
| 2538.0658 | VPAQEKN(N)F Fuc 1 Hex 4 HexNAc 4 | 84.671 | 0.008 | 16.509 | 5.805e+09 | 1067 - 1075 | 15 | 69.31 | 70.73 | 72.09 |
| 2804.1265 | VPAQEKN(N)F Hex 6 HexNAc 3 Neu5Ac 1 | 84.307 | 0.008 | 15.223 | 3.410e+09 | 1067 - 1075 | 6 | 81.04 | 81.43 | 82.34 |
| 2713.0371 | VSN(N)GTHW Hex 5 HexNAc 4 Neu5Ac 1 | 84.001 | 0.008 | 16.552 | 1.254e+09 | 1095 - 1102 | 6 | 81.52 | 81.77 | 82.60 |
| 1985.8470 | VPAQEKN(N)F Hex 4 HexNAc 2 | 83.627 | 0.008 | 15.444 | 6.049e+09 | 1067 - 1075 | 16 | 66.48 | 67.89 | 69.96 |
| 2550.9852 | VSN(N)GTHW Hex 4 HexNAc 4 Neu5Ac 1 | 83.342 | 0.008 | 18.530 | 2.098e+09 | 1095 - 1102 | 10 | 82.67 | 83.02 | 85.27 |
| 2243.8957 | VSN(N)GTHW Fuc 1 Hex 3 HexNAc 4 | 83.321 | 0.008 | 17.417 | 1.348e+09 | 1095 - 1102 | 8 | 69.43 | 69.75 | 70.43 |
| 2672.0106 | VSN(N)GTHW Hex 6 HexNAc 3 Neu5Ac 1 | 83.242 | 0.008 | 16.719 | 1.104e+09 | 1095 - 1102 | 6 | 82.28 | 82.63 | 82.92 |
| 2741.1421 | VPAQEKN(N)F Fuc 1 Hex 4 HexNAc 5 | 83.191 | 0.008 | 14.803 | 1.140e+10 | 1067 - 1075 | 22 | 68.16 | 72.11 | 74.09 |
| 2659.0898 | VPAQEKN(N)F Fuc 1 Hex 6 HexNAc 3 | 80.374 | 0.008 | 16.491 | 2.674e+08 | 1067 - 1075 | 3 | 68.59 | 68.89 | 69.29 |
| 2626.0786 | VPAQEKN(N)F Fuc 1 Hex 4 HexNAc 3 Neu5Ac 1 | 77.924 | 0.008 | 12.966 | 8.513e+08 | 1067 - 1075 | 4 | 84.78 | 85.16 | 85.42 |
| 2229.9510 | VPAQEKN(N)F Hex 3 HexNAc 4 | 72.200 | 0.008 | 16.258 | 7.669e+08 | 1067 - 1075 | 8 | 69.17 | 69.53 | 71.26 |
| 1823.7980 | VPAQEKN(N)F Hex 3 HexNAc 2 | 71.590 | 0.008 | 15.561 | 1.700e+09 | 1067 - 1075 | 7 | 67.54 | 67.89 | 70.31 |
| 4030.7166 | HAIHVSGTN(N)GTKRF Fuc 1 Hex 4 HexNAc 7 Neu5Ac 1 | 64.845 | 0.002 | 13.800 | 6.102e+09 | 65 - 79 | 17 | 60.77 | 62.81 | 63.28 |
| 3827.6403 | HAIHVSGTN(N)GTKRF Fuc 1 Hex 4 HexNAc 6 Neu5Ac 1 | 61.881 | 0.002 | 11.824 | 4.294e+09 | 65 - 79 | 29 | 60.63 | 61.22 | 66.18 |
| 3819.5750 | HAIHVSGTN(N)GTKRF @sulfate 1 Fuc 1 Hex 4 HexNAc 7 | 60.852 | 0.002 | 12.758 | 1.913e+09 | 65 - 79 | 15 | 61.79 | 62.00 | 65.26 |
| 3669.6899 | SALEPLVDLPIGIN(N)ITRF Hex 8 HexNAc 2 | 54.908 | 0.002 | 16.682 | 2.130e+11 | 220 - 238 | 47 | 239.23 | 240.26 | 252.11 |
| 4280.7826 | HAIHVSGTN(N)GTKRF Fuc 1 Hex 5 HexNAc 6 Neu5Ac 2 | 52.434 | 0.002 | 11.376 | 6.326e+09 | 65 - 79 | 23 | 69.89 | 70.89 | 73.43 |
| 2573.2006 | VDLPIGIN(N)ITRF Hex 5 HexNAc 2 | 51.798 | 0.002 | 15.897 | 2.213e+08 | 226 - 238 | 1 | 205.60 | 205.85 | 206.27 |
| 3583.5286 | HAIHVSGTN(N)GTKRF Fuc 1 Hex 5 HexNAc 4 Neu5Ac 1 | 51.017 | 0.002 | 13.982 | 6.065e+09 | 65 - 79 | 27 | 60.49 | 61.88 | 65.30 |
| 4110.6746 | HAIHVSGTN(N)GTKRF @sulfate 1 Fuc 1 Hex 4 HexNAc 7 Neu5Ac 1 | 48.760 | 0.002 | 11.893 | 4.241e+09 | 65 - 79 | 14 | 71.85 | 72.71 | 74.97 |
| 4401.7710 | HAIHVSGTN(N)GTKRF Fuc 1 Hex 7 HexNAc 5 Neu5Ac 2 | 48.737 | 0.002 | 15.500 | 1.117e+09 | 65 - 79 | 3 | 83.37 | 83.77 | 84.28 |
| 3625.5736 | FHAIHVSGTN(N)GTKRF Hex 4 HexNAc 5 Neu5Ac 1 | 47.010 | 0.002 | 16.321 | 1.773e+10 | 64 - 79 | 16 | 52.03 | 52.93 | 57.82 |
| 3218.4048 | HAIHVSGTN(N)GTKRF Fuc 1 Hex 4 HexNAc 3 Neu5Ac 1 | 45.303 | 0.002 | 12.974 | 6.860e+08 | 65 - 79 | 3 | 62.33 | 62.97 | 64.44 |
| 3584.5535 | HAIHVSGTN(N)GTKRF Fuc 3 Hex 5 HexNAc 4 | 44.978 | 0.002 | -3.334 | 5.221e+08 | 65 - 79 | 6 | 52.00 | 52.83 | 63.53 |
| 4239.7616 | HAIHVSGTN(N)GTKRF Fuc 1 Hex 6 HexNAc 5 Neu5Ac 2 | 41.727 | 0.002 | 14.114 | 1.789e+09 | 65 - 79 | 5 | 69.63 | 70.95 | 71.48 |
| 3831.7439 | SALEPLVDLPIGIN(N)ITRF Hex 9 HexNAc 2 | 41.424 | 0.002 | 18.110 | 1.697e+11 | 220 - 238 | 25 | 239.08 | 239.47 | 243.24 |
| 3345.5853 | SALEPLVDLPIGIN(N)ITRF Hex 6 HexNAc 2 | 41.238 | 0.002 | 17.812 | 2.837e+10 | 220 - 238 | 22 | 239.34 | 241.96 | 247.62 |
| 3507.6395 | SALEPLVDLPIGIN(N)ITRF Hex 7 HexNAc 2 | 39.498 | 0.002 | 18.851 | 9.023e+10 | 220 - 238 | 33 | 239.21 | 241.20 | 248.79 |
| 3787.6309 | HAIHVSGTN(N)GTKRF Fuc 3 Hex 5 HexNAc 5 | 38.635 | 0.002 | 15.608 | 5.048e+08 | 65 - 79 | 8 | 52.20 | 53.22 | 53.66 |
| 4360.7403 | HAIHVSGTN(N)GTKRF @sulfate 1 Fuc 1 Hex 5 HexNAc 6 Neu5Ac 2 | 37.124 | 0.002 | 15.225 | 3.150e+09 | 65 - 79 | 8 | 83.09 | 83.54 | 85.07 |
| 3183.5338 | SALEPLVDLPIGIN(N)ITRF Hex 5 HexNAc 2 | 36.866 | 0.002 | 17.224 | 3.287e+10 | 220 - 238 | 22 | 239.38 | 242.94 | 247.58 |
| 3011.2294 | GEVFN(N)ATRF Fuc 1 Hex 5 HexNAc 5 | 36.457 | 0.008 | 17.026 | 2.446e+09 | 338 - 347 | 3 | 122.58 | 122.83 | 123.41 |
| 3907.5964 | HAIHVSGTN(N)GTKRF @sulfate 1 Fuc 1 Hex 4 HexNAc 6 Neu5Ac 1 | 36.444 | 0.002 | 12.057 | 9.662e+08 | 65 - 79 | 6 | 72.22 | 72.99 | 74.85 |
| 2255.9323 | GEVFN(N)ATRF Hex 5 HexNAc 2 | 36.237 | 0.008 | 16.437 | 2.008e+09 | 338 - 347 | 2 | 126.04 | 126.30 | 126.82 |
| 2897.3067 | VDLPIGIN(N)ITRF Hex 7 HexNAc 2 | 35.975 | 0.002 | 13.154 | 8.419e+08 | 226 - 238 | 4 | 204.29 | 204.47 | 204.75 |
| 2443.0186 | GEVFN(N)ATRF Fuc 1 Hex 4 HexNAc 3 | 34.965 | 0.008 | 17.229 | 6.205e+08 | 338 - 347 | 4 | 124.24 | 124.47 | 124.81 |
| 2937.1907 | GEVFN(N)ATRF Fuc 1 Hex 4 HexNAc 4 Neu5Ac 1 | 34.404 | 0.008 | 17.504 | 1.588e+09 | 338 - 347 | 6 | 145.07 | 145.41 | 146.55 |
| 3778.5547 | HAIHVSGTN(N)GTKRF @sulfate 1 Fuc 1 Hex 5 HexNAc 6 | 33.150 | 0.002 | 15.513 | 2.845e+08 | 65 - 79 | 1 | 61.54 | 61.71 | 62.02 |
| 2605.0711 | GEVFN(N)ATRF Fuc 1 Hex 5 HexNAc 3 | 32.308 | 0.008 | 12.406 | 8.022e+08 | 338 - 347 | 9 | 123.30 | 123.91 | 124.20 |
| 4895.9861 | HAIHVSGTN(N)GTKRF Fuc 1 Hex 7 HexNAc 6 Neu5Ac 3 | 32.135 | 0.002 | 12.497 | 1.859e+09 | 65 - 79 | 6 | 79.07 | 80.41 | 81.06 |
| 3771.6366 | FHAIHVSGTN(N)GTKRF Fuc 1 Hex 4 HexNAc 5 Neu5Ac 1 | 31.763 | 0.002 | 15.177 | 1.473e+09 | 64 - 79 | 9 | 52.89 | 53.16 | 84.12 |
| 3074.3681 | HAIHVSGTN(N)GTKRF Hex 4 HexNAc 3 Neu5Ac 1 | 31.668 | 0.002 | -6.429 | 4.105e+06 | 65 - 79 | 1 | 54.94 | 54.95 | 55.04 |
| 3626.5893 | FHAIHVSGTN(N)GTKRF Fuc 2 Hex 4 HexNAc 5 | 31.515 | 0.002 | 15.308 | 7.896e+08 | 64 - 79 | 9 | 74.46 | 74.68 | 75.75 |
| 3803.6219 | HAIHVSGTN(N)GTKRF Fuc 2 Hex 6 HexNAc 5 | 30.968 | 0.002 | 13.734 | 3.213e+08 | 65 - 79 | 10 | 52.17 | 52.60 | 54.04 |
| 3115.3990 | FHAIHVSGTN(N)GTKRF Fuc 1 Hex 3 HexNAc 4 | 30.610 | 0.002 | 14.804 | 4.226e+08 | 64 - 79 | 7 | 75.20 | 75.84 | 76.46 |
| 4077.7259 | HAIHVSGTN(N)GTKRF Fuc 1 Hex 5 HexNAc 5 Neu5Ac 2 | 30.408 | 0.002 | 10.593 | 1.778e+09 | 65 - 79 | 12 | 70.87 | 71.50 | 73.37 |
| 4069.6464 | HAIHVSGTN(N)GTKRF @sulfate 1 Fuc 1 Hex 5 HexNAc 6 Neu5Ac 1 | 29.961 | 0.002 | 12.939 | 2.540e+09 | 65 - 79 | 12 | 71.59 | 72.43 | 74.41 |
| 4530.8525 | HAIHVSGTN(N)GTKRF Fuc 1 Hex 6 HexNAc 5 Neu5Ac 3 | 29.819 | 0.002 | 13.067 | 4.267e+09 | 65 - 79 | 7 | 80.82 | 81.65 | 82.15 |
| 4516.8767 | HAIHVSGTN(N)GTKRF Fuc 1 Hex 7 HexNAc 7 Neu5Ac 1 | 29.652 | 0.002 | 15.588 | 6.275e+08 | 65 - 79 | 3 | 61.57 | 61.73 | 62.43 |
| 4354.8195 | HAIHVSGTN(N)GTKRF Fuc 1 Hex 6 HexNAc 7 Neu5Ac 1 | 29.533 | 0.002 | 11.450 | 1.187e+09 | 65 - 79 | 9 | 60.04 | 61.30 | 63.13 |
| 3375.4978 | FHAIHVSGTN(N)GTKRF Hex 3 HexNAc 6 | 28.908 | 0.002 | -4.232 | 1.023e+07 | 64 - 79 | 3 | 56.23 | 56.31 | 56.31 |
| 2735.2534 | VDLPIGIN(N)ITRF Hex 6 HexNAc 2 | 27.117 | 0.002 | 13.634 | 1.916e+08 | 226 - 238 | 2 | 204.86 | 205.01 | 205.20 |
| 3917.6865 | FHAIHVSGTN(N)GTKRF Fuc 2 Hex 4 HexNAc 5 Neu5Ac 1 | 26.886 | 0.002 | 10.020 | 1.669e+08 | 64 - 79 | 4 | 82.89 | 85.07 | 85.20 |
| 3609.4268 | SN(N)NSIAIPTN(N)F Hex 10 HexNAc 4 | 26.419 | 0.002 | 17.216 | 1.466e+10 | 707 - 718 | 7 | 156.68 | 156.98 | 158.20 |
| 2594.0429 | VSN(N)GTHWF Fuc 1 Hex 3 HexNAc 5 | 26.181 | 0.008 | 15.285 | 1.227e+09 | 1095 - 1103 | 5 | 115.88 | 116.21 | 116.71 |
| 3521.5582 | FHAIHVSGTN(N)GTKRF Fuc 1 Hex 3 HexNAc 6 | 25.870 | 0.002 | -5.376 | 3.475e+08 | 64 - 79 | 6 | 54.86 | 75.47 | 75.87 |
| 4022.7031 | HAIHVSGTN(N)GTKRF Fuc 1 Hex 7 HexNAc 6 | 25.691 | 0.002 | 16.245 | 1.862e+08 | 65 - 79 | 2 | 53.16 | 53.32 | 53.61 |
| 3560.4861 | FHAIHVSGTN(N)GTKRF @sulfate 1 Fuc 1 Hex 4 HexNAc 5 | 25.303 | 0.002 | 14.575 | 1.885e+08 | 64 - 79 | 2 | 85.01 | 85.23 | 85.32 |
| 3990.7116 | HAIHVSGTN(N)GTKRF Fuc 3 Hex 5 HexNAc 6 | 25.283 | 0.002 | 14.290 | 1.291e+09 | 65 - 79 | 10 | 51.82 | 53.09 | 53.70 |
| 4151.7445 | HAIHVSGTN(N)GTKRF Fuc 1 Hex 6 HexNAc 6 Neu5Ac 1 | 25.216 | 0.002 | 12.144 | 1.540e+09 | 65 - 79 | 21 | 59.98 | 64.21 | 65.65 |
| 2782.1705 | VPAQEKN(N)F Fuc 1 Hex 3 HexNAc 6 | 25.006 | 0.008 | 16.562 | 1.374e+09 | 1067 - 1075 | 5 | 69.41 | 69.78 | 70.92 |
| 4256.7306 | HAIHVSGTN(N)GTKRF @sulfate 1 Fuc 2 Hex 4 HexNAc 7 Neu5Ac 1 | 24.950 | 0.002 | 15.535 | 2.068e+09 | 65 - 79 | 10 | 72.24 | 73.18 | 74.32 |
| 4063.7238 | HAIHVSGTN(N)GTKRF Fuc 1 Hex 6 HexNAc 7 | 24.377 | 0.002 | -5.617 | 4.325e+08 | 65 - 79 | 12 | 53.24 | 53.49 | 96.29 |
| 4604.8944 | HAIHVSGTN(N)GTKRF Fuc 1 Hex 7 HexNAc 6 Neu5Ac 2 | 24.231 | 0.002 | 13.350 | 9.793e+08 | 65 - 79 | 9 | 68.68 | 69.22 | 71.24 |
| 2162.8513 | VSN(N)GTHWF Hex 5 HexNAc 2 | 23.909 | 0.008 | 18.265 | 1.453e+09 | 1095 - 1103 | 2 | 116.94 | 117.23 | 117.91 |
| 4662.9318 | HAIHVSGTN(N)GTKRF Fuc 2 Hex 7 HexNAc 7 Neu5Ac 1 | 23.834 | 0.002 | 12.675 | 1.477e+08 | 65 - 79 | 2 | 61.46 | 61.63 | 61.77 |
| 2511.9901 | VSN(N)GTHWF Fuc 1 Hex 5 HexNAc 3 | 23.677 | 0.008 | 17.713 | 1.499e+09 | 1095 - 1103 | 6 | 115.18 | 115.82 | 116.64 |
| 3103.2040 | VSN(N)GTHW Fuc 1 Hex 4 HexNAc 6 Neu5Ac 1 | 23.631 | 0.008 | 16.971 | 4.628e+08 | 1095 - 1102 | 3 | 83.16 | 83.54 | 84.06 |
| 2996.2108 | GEVFN(N)ATRF Fuc 2 Hex 4 HexNAc 5 | 23.530 | 0.008 | 9.100 | 3.610e+08 | 338 - 347 | 4 | 92.44 | 92.55 | 93.52 |
| 2390.9629 | VSN(N)GTHWF Fuc 1 Hex 3 HexNAc 4 | 23.024 | 0.008 | 16.403 | 2.553e+09 | 1095 - 1103 | 5 | 116.94 | 117.17 | 117.64 |
| 2349.9384 | VSN(N)GTHWF Fuc 1 Hex 4 HexNAc 3 | 22.982 | 0.008 | 16.567 | 1.810e+09 | 1095 - 1103 | 5 | 116.97 | 117.28 | 117.67 |
| 3047.1955 | VSN(N)GTHWF Fuc 1 Hex 4 HexNAc 5 Neu5Ac 1 | 22.602 | 0.008 | 13.210 | 7.013e+08 | 1095 - 1103 | 2 | 136.99 | 137.36 | 137.56 |
| 2527.9823 | VSN(N)GTHWF Hex 6 HexNAc 3 | 22.568 | 0.008 | 17.453 | 2.312e+09 | 1095 - 1103 | 7 | 114.32 | 115.49 | 116.00 |
| 3860.6058 | HAIHVSGTN(N)GTKRF @sulfate 1 Fuc 1 Hex 3 HexNAc 8 | 22.451 | 0.002 | 11.710 | 1.311e+09 | 65 - 79 | 13 | 62.13 | 62.39 | 65.30 |
| 2698.0575 | VSN(N)GTHWF Hex 4 HexNAc 4 Neu5Ac 1 | 22.367 | 0.008 | 16.230 | 3.420e+09 | 1095 - 1103 | 5 | 139.14 | 139.32 | 139.60 |
| 3268.3276 | VPAQEKN(N)F Fuc 1 Hex 6 HexNAc 6 | 21.950 | 0.023 | 7.517 | 6.346e+07 | 1067 - 1075 | 1 | 72.40 | 72.40 | 72.74 |
| 3183.2951 | HKNN(N)KSWMESEF Fuc 1 Hex 3 HexNAc 5 | 21.535 | 0.002 | 13.912 | 2.675e+10 | 145 - 157 | 15 | 91.45 | 92.05 | 94.28 |
| 3973.6971 | HAIHVSGTN(N)GTKRF Fuc 2 Hex 4 HexNAc 6 Neu5Ac 1 | 21.372 | 0.002 | 13.406 | 1.365e+09 | 65 - 79 | 17 | 60.39 | 62.10 | 63.17 |
| 4094.7212 | HAIHVSGTN(N)GTKRF Fuc 2 Hex 6 HexNAc 5 Neu5Ac 1 | 21.084 | 0.002 | 11.785 | 8.012e+08 | 65 - 79 | 16 | 59.94 | 61.60 | 62.81 |
| 3771.4766 | SN(N)NSIAIPTN(N)F Hex 11 HexNAc 4 | 20.669 | 0.002 | 17.127 | 3.301e+10 | 707 - 718 | 7 | 154.48 | 155.68 | 156.24 |
| 4095.5815 | SN(N)NSIAIPTN(N)F Hex 13 HexNAc 4 | 20.555 | 0.002 | 17.695 | 1.393e+10 | 707 - 718 | 5 | 153.45 | 154.14 | 155.16 |
| 4297.8040 | HAIHVSGTN(N)GTKRF Fuc 2 Hex 6 HexNAc 6 Neu5Ac 1 | 20.527 | 0.002 | 15.961 | 1.576e+08 | 65 - 79 | 2 | 63.75 | 63.95 | 64.16 |
| 4042.8254 | RVQPTESIVRFPN(N)ITNL Fuc 1 Hex 5 HexNAc 4 Neu5Ac 1 | 20.411 | 0.002 | 15.370 | 1.269e+10 | 318 - 335 | 7 | 183.69 | 184.52 | 185.05 |
| 3224.5603 | SALEPLVDLPIGIN(N)ITRF Hex 4 HexNAc 3 | 20.338 | 0.002 | 16.013 | 3.038e+09 | 220 - 238 | 3 | 242.79 | 243.20 | 244.86 |
| 3487.5873 | N(N)FSQILPDPSKPSKRSF Hex 7 HexNAc 2 | 20.183 | 0.002 | 13.184 | 4.444e+09 | 800 - 817 | 8 | 123.85 | 124.53 | 159.31 |
| 3204.4770 | N(N)FSQILPDPSKPSKRSF Hex 4 HexNAc 3 | 19.394 | 0.002 | 12.738 | 3.615e+09 | 800 - 817 | 6 | 125.79 | 126.36 | 126.92 |
| 3199.5132 | RVQPTESIVRFPN(N)ITNL Hex 5 HexNAc 2 | 19.107 | 0.002 | 17.409 | 6.809e+08 | 318 - 335 | 4 | 170.89 | 171.36 | 172.02 |
| 3595.4204 | HKNN(N)KSWMESEF Fuc 1 Hex 5 HexNAc 4 Neu5Ac 1 | 18.527 | 0.004 | 16.910 | 6.982e+09 | 145 - 157 | 10 | 104.37 | 104.63 | 105.97 |
| 3751.7308 | RVQPTESIVRFPN(N)ITNL Fuc 1 Hex 5 HexNAc 4 | 18.175 | 0.004 | 14.730 | 9.264e+09 | 318 - 335 | 6 | 166.85 | 167.17 | 167.75 |
| 4426.8468 | HAIHVSGTN(N)GTKRF Fuc 2 Hex 5 HexNAc 6 Neu5Ac 2 | 18.108 | 0.004 | 12.803 | 2.870e+09 | 65 - 79 | 14 | 69.22 | 71.48 | 72.02 |
| 4427.8512 | FHAIHVSGTN(N)GTKRF Fuc 1 Hex 5 HexNAc 6 Neu5Ac 2 | 18.068 | 0.004 | -5.096 | 5.432e+06 | 64 - 79 | 1 | 69.75 | 69.75 | 69.92 |
| 3331.3727 | HAIHVSGTN(N)GTKRF @sulfate 1 Fuc 1 Hex 6 HexNAc 3 | 18.060 | 0.004 | 12.640 | 4.581e+07 | 65 - 79 | 1 | 62.13 | 62.19 | 62.55 |
| 3933.5295 | SN(N)NSIAIPTN(N)F Hex 12 HexNAc 4 | 17.793 | 0.004 | 15.734 | 1.313e+10 | 707 - 718 | 5 | 153.73 | 154.77 | 155.48 |
| 3163.4523 | N(N)FSQILPDPSKPSKRSF Hex 5 HexNAc 2 | 17.583 | 0.004 | 18.678 | 1.215e+11 | 800 - 817 | 25 | 125.03 | 125.99 | 128.34 |
| 4807.9683 | HAIHVSGTN(N)GTKRF Fuc 1 Hex 7 HexNAc 7 Neu5Ac 2 | 17.551 | 0.004 | 15.595 | 2.100e+08 | 65 - 79 | 4 | 70.34 | 70.53 | 70.87 |
| 4257.6354 | SN(N)NSIAIPTN(N)F Hex 14 HexNAc 4 | 17.508 | 0.004 | 13.760 | 6.963e+09 | 707 - 718 | 7 | 152.56 | 153.11 | 154.67 |
| 4442.8423 | HAIHVSGTN(N)GTKRF Fuc 1 Hex 6 HexNAc 6 Neu5Ac 2 | 17.282 | 0.006 | 11.830 | 1.052e+09 | 65 - 79 | 17 | 69.29 | 71.59 | 72.05 |
| 4245.9061 | RVQPTESIVRFPN(N)ITNL Fuc 1 Hex 5 HexNAc 5 Neu5Ac 1 | 17.163 | 0.006 | 11.688 | 1.759e+09 | 318 - 335 | 3 | 183.50 | 184.55 | 185.23 |
| 3318.4795 | FHAIHVSGTN(N)GTKRF Fuc 1 Hex 3 HexNAc 5 | 16.975 | 0.006 | 8.895 | 1.037e+09 | 64 - 79 | 15 | 55.98 | 76.38 | 76.87 |
| 4395.8528 | HAIHVSGTN(N)GTKRF Fuc 1 Hex 5 HexNAc 8 Neu5Ac 1 | 16.841 | 0.007 | 14.212 | 6.126e+08 | 65 - 79 | 6 | 60.53 | 61.57 | 61.91 |
| 3345.3484 | HKNN(N)KSWMESEF Fuc 1 Hex 4 HexNAc 5 | 16.465 | 0.007 | 15.318 | 2.141e+10 | 145 - 157 | 18 | 90.97 | 92.05 | 93.65 |
| 3325.4977 | N(N)FSQILPDPSKPSKRSF Hex 6 HexNAc 2 | 16.462 | 0.007 | 12.985 | 1.416e+10 | 800 - 817 | 15 | 124.39 | 125.12 | 125.89 |
| 4699.0537 | RVQPTESIVRFPN(N)ITNL Fuc 1 Hex 6 HexNAc 5 Neu5Ac 2 | 16.419 | 0.007 | 13.729 | 2.641e+09 | 318 - 335 | 6 | 201.59 | 202.98 | 203.93 |
| 4083.8511 | RVQPTESIVRFPN(N)ITNL Fuc 1 Hex 4 HexNAc 5 Neu5Ac 1 | 16.138 | 0.007 | 13.200 | 5.214e+09 | 318 - 335 | 12 | 183.89 | 184.29 | 186.27 |
| 3880.7719 | RVQPTESIVRFPN(N)ITNL Fuc 1 Hex 4 HexNAc 4 Neu5Ac 1 | 16.118 | 0.007 | 17.834 | 1.713e+10 | 318 - 335 | 9 | 184.55 | 185.75 | 186.66 |
| 3427.6265 | RVQPTESIVRFPN(N)ITNL Fuc 1 Hex 3 HexNAc 4 | 16.039 | 0.007 | 16.109 | 2.767e+10 | 318 - 335 | 11 | 168.16 | 168.78 | 170.71 |
| 3265.5863 | SALEPLVDLPIGIN(N)ITRF Hex 3 HexNAc 4 | 15.723 | 0.007 | 16.467 | 3.211e+09 | 220 - 238 | 6 | 242.75 | 243.19 | 244.32 |
| 3391.5609 | N(N)FSQILPDPSKPSKRSF Fuc 1 Hex 3 HexNAc 4 | 15.705 | 0.007 | 13.034 | 5.482e+09 | 800 - 817 | 9 | 124.24 | 124.93 | 125.37 |
| 3974.5626 | SN(N)NSIAIPTN(N)F Hex 11 HexNAc 5 | 15.562 | 0.007 | -1.984 | 6.102e+09 | 707 - 718 | 4 | 123.06 | 155.29 | 155.57 |
| 3649.6068 | N(N)FSQILPDPSKPSKRSF Hex 8 HexNAc 2 | 15.156 | 0.008 | 16.933 | 1.659e+09 | 800 - 817 | 6 | 123.71 | 124.16 | 124.73 |
| 4001.5779 | SN(N)NSIAIPTN(N)F Fuc 1 Hex 9 HexNAc 6 | 15.093 | 0.008 | 11.907 | 3.053e+08 | 707 - 718 | 4 | 105.44 | 105.76 | 105.92 |
| 3553.6108 | N(N)FSQILPDPSKPSKRSF Fuc 1 Hex 4 HexNAc 4 | 15.059 | 0.008 | 15.850 | 1.245e+09 | 800 - 817 | 6 | 123.70 | 124.47 | 124.81 |
| 4812.1103 | RVQPTESIVRFPN(N)ITNLC(C)PF Fuc 1 Hex 6 HexNAc 5 Neu5Ac 1 | 14.895 | 0.008 | 17.081 | 7.110e+09 | 318 - 338 | 5 | 206.41 | 207.47 | 208.06 |
| 3501.4393 | HAIHVSGTN(N)GTKRF @sulfate 1 Fuc 1 Hex 4 HexNAc 4 Neu5Ac 1 | 14.711 | 0.008 | 14.408 | 6.956e+07 | 65 - 79 | 2 | 71.77 | 71.93 | 72.05 |
| 3366.5594 | N(N)FSQILPDPSKPSKRSF Hex 5 HexNAc 3 | 14.605 | 0.008 | 12.043 | 1.212e+10 | 800 - 817 | 20 | 124.49 | 125.53 | 126.47 |
| 3917.6854 | HAIHVSGTN(N)GTKRF Hex 6 HexNAc 7 | 14.539 | 0.008 | 17.434 | 4.941e+08 | 65 - 79 | 1 | 84.78 | 84.92 | 85.11 |
| 3411.6434 | SALEPLVDLPIGIN(N)ITRF Fuc 1 Hex 3 HexNAc 4 | 14.455 | 0.008 | 14.977 | 6.505e+09 | 220 - 238 | 8 | 239.61 | 240.24 | 241.49 |
| 2820.1779 | HAIHVSGTN(N)GTKRF @sulfate 1 Hex 5 HexNAc 2 | 14.398 | 0.008 | 12.684 | 1.983e+07 | 65 - 79 | 1 | 62.97 | 63.07 | 63.15 |
| 3782.5010 | HKNN(N)KSWMESEF Fuc 2 Hex 4 HexNAc 5 Neu5Ac 1 | 14.109 | 0.010 | 16.027 | 2.039e+09 | 145 - 157 | 4 | 106.05 | 106.35 | 106.84 |
| 3433.3672 | HKNN(N)KSWMESEF Fuc 1 Hex 4 HexNAc 4 Neu5Ac 1 | 14.047 | 0.010 | 16.125 | 4.333e+09 | 145 - 157 | 7 | 105.41 | 106.72 | 107.32 |
| 3480.5292 | FHAIHVSGTN(N)GTKRF Fuc 1 Hex 4 HexNAc 5 | 14.033 | 0.010 | 11.029 | 4.750e+08 | 64 - 79 | 12 | 74.57 | 75.27 | 76.29 |
| 3386.6127 | SALEPLVDLPIGIN(N)ITRF Hex 5 HexNAc 3 | 13.843 | 0.010 | 18.025 | 6.480e+09 | 220 - 238 | 2 | 241.62 | 242.12 | 242.85 |
| 3142.2702 | HKNN(N)KSWMESEF Fuc 1 Hex 4 HexNAc 4 | 13.831 | 0.010 | 14.174 | 9.152e+09 | 145 - 157 | 5 | 91.34 | 92.01 | 92.25 |
| 3613.4441 | YHKNN(N)KSWMESEF Fuc 2 Hex 5 HexNAc 4 | 13.779 | 0.010 | 16.856 | 1.680e+09 | 144 - 157 | 4 | 93.30 | 93.57 | 94.29 |
| 3062.5082 | SALEPLVDLPIGIN(N)ITRF Hex 3 HexNAc 3 | 13.770 | 0.010 | 18.309 | 1.531e+09 | 220 - 238 | 2 | 243.95 | 244.32 | 245.28 |
| 4527.0453 | VPAQEKN(N)FTTAPAIC(C)HDGKAHFPREGVF Fuc 1 Hex 4 HexNAc 3 | 13.712 | 0.010 | 16.219 | 2.214e+10 | 1067 - 1095 | 10 | 117.14 | 118.04 | 119.55 |
| 4006.7059 | HAIHVSGTN(N)GTKRF Fuc 2 Hex 6 HexNAc 6 | 13.659 | 0.010 | 10.419 | 2.539e+08 | 65 - 79 | 9 | 54.35 | 56.67 | 57.75 |
| 3450.3808 | HKNN(N)KSWMESEF Fuc 2 Hex 5 HexNAc 4 | 13.571 | 0.010 | 16.077 | 1.248e+09 | 145 - 157 | 4 | 90.65 | 91.07 | 91.62 |
| 3851.5812 | FHAIHVSGTN(N)GTKRF @sulfate 1 Fuc 1 Hex 4 HexNAc 5 Neu5Ac 1 | 13.538 | 0.010 | 15.591 | 7.345e+08 | 64 - 79 | 4 | 96.90 | 97.09 | 97.33 |
| 3586.6105 | GGFN(N)FSQILPDPSKPSKRSF Hex 6 HexNAc 2 | 13.448 | 0.010 | 11.847 | 3.623e+08 | 797 - 817 | 3 | 165.14 | 165.45 | 165.79 |
| 4136.6083 | SN(N)NSIAIPTN(N)F Hex 12 HexNAc 5 | 13.307 | 0.010 | 15.643 | 2.116e+09 | 707 - 718 | 4 | 153.90 | 154.33 | 155.13 |
| 3386.3753 | HKNN(N)KSWMESEF Fuc 1 Hex 3 HexNAc 6 | 13.222 | 0.010 | 11.952 | 4.432e+09 | 145 - 157 | 9 | 91.32 | 91.74 | 94.06 |
| 3424.5610 | GGFN(N)FSQILPDPSKPSKRSF Hex 5 HexNAc 2 | 13.192 | 0.010 | 14.746 | 1.697e+10 | 797 - 817 | 12 | 165.00 | 166.11 | 166.97 |
| 4771.1521 | VPAQEKN(N)FTTAPAIC(C)HDGKAHFPREGVF Fuc 1 Hex 3 HexNAc 5 | 13.126 | 0.010 | 13.449 | 5.001e+10 | 1067 - 1095 | 21 | 117.09 | 117.79 | 119.59 |
| 3223.4861 | VDLPIGIN(N)ITRF Hex 4 HexNAc 6 | 13.106 | 0.010 | 0.000 | 0.000e+00 | 226 - 238 | 1 | - | - | - |
| 4358.9627 | RVQPTESIVRFPN(N)ITNLC(C)PF Fuc 1 Hex 5 HexNAc 5 | 13.103 | 0.010 | 15.268 | 6.885e+09 | 318 - 338 | 7 | 190.64 | 191.78 | 192.68 |
| 4035.8660 | RVQPTESIVRFPN(N)ITNLC(C)PFGEVF Hex 5 HexNAc 2 | 13.085 | 0.010 | 17.920 | 3.046e+09 | 318 - 342 | 4 | 219.84 | 220.15 | 221.17 |
| 3636.4487 | HKNN(N)KSWMESEF Fuc 1 Hex 4 HexNAc 5 Neu5Ac 1 | 13.072 | 0.010 | 13.379 | 8.888e+09 | 145 - 157 | 17 | 104.64 | 104.98 | 107.66 |
| 3245.4972 | N(N)FSQILPDPSKPSKRSF Hex 3 HexNAc 4 | 12.996 | 0.010 | 12.595 | 1.553e+09 | 800 - 817 | 6 | 125.99 | 126.36 | 126.93 |
| 4689.0989 | VPAQEKN(N)FTTAPAIC(C)HDGKAHFPREGVF Fuc 1 Hex 5 HexNAc 3 | 12.936 | 0.010 | 16.523 | 1.357e+10 | 1067 - 1095 | 13 | 117.36 | 117.79 | 119.27 |
| 3748.6605 | GGFN(N)FSQILPDPSKPSKRSF Hex 7 HexNAc 2 | 12.901 | 0.010 | 13.725 | 2.996e+09 | 797 - 817 | 7 | 164.47 | 165.01 | 165.45 |
| 4879.1750 | RVQPTESIVRFPN(N)ITNLC(C)PFGEVF Fuc 1 Hex 5 HexNAc 4 Neu5Ac 1 | 12.804 | 0.010 | 18.005 | 5.809e+10 | 318 - 342 | 12 | 232.00 | 233.39 | 236.40 |
| 3235.4939 | LIVNN(N)ATNVVIKVC(C)EF Fuc 1 Hex 4 HexNAc 3 | 12.777 | 0.010 | 16.574 | 2.324e+09 | 117 - 133 | 4 | 179.00 | 179.31 | 180.18 |
| 4196.9091 | RVQPTESIVRFPN(N)ITNLC(C)PF Fuc 1 Hex 4 HexNAc 5 | 12.628 | 0.010 | 13.200 | 1.935e+10 | 318 - 338 | 18 | 191.19 | 192.68 | 194.67 |
| 4840.0693 | QTRAGC(C)LIGAEHVN(N)NSYEC(C)DIPIGAGIC(C)ASY Fuc 1 Hex 3 HexNAc 4 | 12.565 | 0.010 | 16.847 | 4.211e+10 | 643 - 674 | 9 | 165.97 | 166.28 | 167.69 |
| 3451.3901 | YHKNN(N)KSWMESEF Fuc 2 Hex 4 HexNAc 4 | 12.441 | 0.011 | 15.202 | 4.965e+09 | 144 - 157 | 7 | 93.67 | 94.28 | 94.82 |
| 3657.6250 | N(N)FSQILPDPSKPSKRSF Hex 5 HexNAc 3 Neu5Ac 1 | 12.334 | 0.011 | -2.919 | 1.335e+10 | 800 - 817 | 12 | 82.80 | 140.11 | 140.68 |
| 4281.8079 | HAIHVSGTN(N)GTKRF Fuc 3 Hex 5 HexNAc 6 Neu5Ac 1 | 12.332 | 0.011 | 11.648 | 1.012e+09 | 65 - 79 | 16 | 59.85 | 61.77 | 62.70 |
| 3705.5334 | FHAIHVSGTN(N)GTKRF @sulfate 1 Hex 4 HexNAc 5 Neu5Ac 1 | 12.311 | 0.011 | 14.462 | 6.062e+08 | 64 - 79 | 2 | 61.91 | 62.16 | 62.72 |
| 3491.4064 | HKNN(N)KSWMESEF Fuc 2 Hex 4 HexNAc 5 | 12.099 | 0.012 | 16.650 | 9.459e+09 | 145 - 157 | 6 | 90.81 | 91.15 | 91.74 |
| 5017.9704 | HAIHVSGTN(N)GTKRF @sulfate 1 Fuc 3 Hex 6 HexNAc 7 Neu5Ac 2 | 11.938 | 0.012 | -4.504 | 6.687e+07 | 65 - 79 | 1 | 94.66 | 94.68 | 94.87 |
| 3591.5538 | IVNN(N)ATNVVIKVC(C)EF Hex 6 HexNAc 3 Neu5Ac 1 | 11.886 | 0.012 | 15.937 | 5.844e+09 | 118 - 133 | 8 | 177.91 | 179.14 | 179.72 |
| 4521.0131 | RVQPTESIVRFPN(N)ITNLC(C)PF Fuc 1 Hex 6 HexNAc 5 | 11.782 | 0.013 | 16.759 | 2.128e+09 | 318 - 338 | 5 | 190.37 | 190.72 | 192.04 |
| 3226.3777 | HAIHVSGTN(N)GTKRF Hex 8 HexNAc 2 | 11.773 | 0.013 | 15.004 | 2.941e+07 | 65 - 79 | 2 | 51.57 | 51.74 | 52.34 |
| 3284.4655 | IVNN(N)ATNVVIKVC(C)EF Fuc 1 Hex 5 HexNAc 3 | 11.624 | 0.013 | 14.064 | 4.295e+09 | 118 - 133 | 5 | 159.26 | 159.55 | 160.49 |
| 3429.4996 | IVNN(N)ATNVVIKVC(C)EF Hex 5 HexNAc 3 Neu5Ac 1 | 11.522 | 0.013 | 14.960 | 8.168e+09 | 118 - 133 | 10 | 178.75 | 180.18 | 180.88 |
| 3616.5844 | IVNN(N)ATNVVIKVC(C)EF Fuc 1 Hex 4 HexNAc 4 Neu5Ac 1 | 11.521 | 0.013 | 15.944 | 1.157e+10 | 118 - 133 | 11 | 177.62 | 177.91 | 180.14 |
| 3507.4032 | HKNN(N)KSWMESEF Fuc 1 Hex 5 HexNAc 5 | 11.449 | 0.013 | 15.819 | 5.447e+09 | 145 - 157 | 10 | 90.29 | 91.72 | 92.01 |
| 3288.3283 | HKNN(N)KSWMESEF Fuc 2 Hex 4 HexNAc 4 | 11.420 | 0.013 | 15.185 | 2.037e+09 | 145 - 157 | 6 | 91.15 | 91.53 | 91.99 |
| 3776.6740 | VDLPIGIN(N)ITRF Fuc 1 Hex 4 HexNAc 8 | 11.407 | 0.013 | 0.000 | 0.000e+00 | 226 - 238 | 1 | - | - | - |
| 4892.1752 | VPAQEKN(N)FTTAPAIC(C)HDGKAHFPREGVF Fuc 1 Hex 5 HexNAc 4 | 11.277 | 0.013 | 16.092 | 2.060e+10 | 1067 - 1095 | 9 | 116.64 | 117.33 | 119.39 |
| 3413.5170 | IVNN(N)ATNVVIKVC(C)EF Fuc 1 Hex 4 HexNAc 3 Neu5Ac 1 | 11.270 | 0.013 | 13.914 | 3.569e+09 | 118 - 133 | 9 | 178.38 | 178.96 | 180.18 |
| 3163.4377 | IVNN(N)ATNVVIKVC(C)EF Fuc 1 Hex 3 HexNAc 4 | 11.159 | 0.013 | 16.343 | 1.206e+10 | 118 - 133 | 12 | 160.01 | 160.23 | 161.60 |
| 4933.2018 | VPAQEKN(N)FTTAPAIC(C)HDGKAHFPREGVF Fuc 1 Hex 4 HexNAc 5 | 10.965 | 0.014 | 13.804 | 2.137e+10 | 1067 - 1095 | 15 | 116.94 | 117.69 | 119.59 |
| 4568.0695 | VPAQEKN(N)FTTAPAIC(C)HDGKAHFPREGVF Fuc 1 Hex 3 HexNAc 4 | 10.953 | 0.014 | 18.091 | 6.336e+10 | 1067 - 1095 | 12 | 117.11 | 118.07 | 119.61 |
| 3932.6678 | HAIHVSGTN(N)GTKRF Fuc 2 Hex 5 HexNAc 5 Neu5Ac 1 | 10.823 | 0.014 | 11.011 | 8.861e+08 | 65 - 79 | 21 | 60.25 | 61.61 | 65.72 |
| 4040.6151 | SN(N)NSIAIPTN(N)F Fuc 1 Hex 8 HexNAc 7 | 10.674 | 0.013 | 11.693 | 4.177e+08 | 707 - 718 | 3 | 155.95 | 156.06 | 156.19 |
| 4047.7290 | HAIHVSGTN(N)GTKRF Fuc 2 Hex 5 HexNAc 7 | 10.670 | 0.014 | 11.806 | 1.259e+08 | 65 - 79 | 9 | 52.60 | 52.83 | 53.55 |
| 3650.4544 | SN(N)NSIAIPTN(N)F Hex 9 HexNAc 5 | 10.594 | 0.013 | 16.488 | 9.037e+08 | 707 - 718 | 1 | 157.50 | 157.73 | 157.93 |
| 4006.7901 | N(N)FSQILPDPSKPSKRSF Fuc 1 Hex 5 HexNAc 4 Neu5Ac 1 | 10.585 | 0.014 | -2.920 | 1.956e+09 | 800 - 817 | 6 | 136.95 | 137.47 | 235.84 |
| 3259.4324 | IVNN(N)ATNVVIKVC(C)EF Hex 7 HexNAc 2 | 10.585 | 0.014 | 17.570 | 2.300e+09 | 118 - 133 | 3 | 159.31 | 159.58 | 160.04 |
| 5293.2127 | QTRAGC(C)LIGAEHVN(N)NSYEC(C)DIPIGAGIC(C)ASY Fuc 1 Hex 4 HexNAc 4 Neu5Ac 1 | 10.566 | 0.014 | 15.789 | 1.431e+10 | 643 - 674 | 10 | 177.58 | 178.36 | 180.14 |
| 3398.4337 | FHAIHVSGTN(N)GTKRF @sulfate 1 Fuc 1 Hex 3 HexNAc 5 | 10.465 | 0.014 | 13.007 | 3.629e+08 | 64 - 79 | 2 | 85.77 | 85.82 | 86.12 |
| 3614.7224 | SALEPLVDLPIGIN(N)ITRF Fuc 1 Hex 3 HexNAc 5 | 10.379 | 0.016 | 14.722 | 2.038e+09 | 220 - 238 | 5 | 239.06 | 239.18 | 239.53 |
| 3300.4598 | IVNN(N)ATNVVIKVC(C)EF Hex 6 HexNAc 3 | 10.264 | 0.016 | 17.277 | 5.284e+09 | 118 - 133 | 3 | 159.58 | 159.87 | 160.41 |
| 3850.5727 | HAIHVSGTN(N)GTKRF @sulfate 1 Fuc 2 Hex 4 HexNAc 5 Neu5Ac 1 | 10.209 | 0.017 | 9.996 | 2.122e+08 | 65 - 79 | 3 | 72.60 | 76.69 | 76.87 |
| 3366.5183 | IVNN(N)ATNVVIKVC(C)EF Fuc 1 Hex 3 HexNAc 5 | 10.193 | 0.017 | 15.819 | 3.864e+09 | 118 - 133 | 4 | 159.41 | 160.11 | 160.49 |
| 5095.2551 | VPAQEKN(N)FTTAPAIC(C)HDGKAHFPREGVF Fuc 1 Hex 5 HexNAc 5 | 10.135 | 0.017 | 16.299 | 1.332e+10 | 1067 - 1095 | 11 | 116.36 | 116.97 | 117.93 |
| 3325.4890 | IVNN(N)ATNVVIKVC(C)EF Fuc 1 Hex 4 HexNAc 4 | 10.135 | 0.017 | 15.070 | 5.618e+09 | 118 - 133 | 6 | 159.09 | 159.58 | 160.55 |
| 4686.9474 | HAIHVSGTN(N)GTKRF Fuc 1 Hex 5 HexNAc 8 Neu5Ac 2 | 10.098 | 0.017 | 14.590 | 4.624e+08 | 65 - 79 | 6 | 71.41 | 71.57 | 71.90 |
| 3575.5593 | IVNN(N)ATNVVIKVC(C)EF Fuc 1 Hex 5 HexNAc 3 Neu5Ac 1 | 10.091 | 0.017 | 15.837 | 4.488e+09 | 118 - 133 | 5 | 177.56 | 178.82 | 179.93 |
| 3446.5178 | IVNN(N)ATNVVIKVC(C)EF Fuc 1 Hex 6 HexNAc 3 | 10.045 | 0.017 | 15.613 | 1.717e+09 | 118 - 133 | 5 | 158.92 | 159.23 | 160.01 |
| 2935.3291 | IVNN(N)ATNVVIKVC(C)EF Hex 5 HexNAc 2 | 10.006 | 0.017 | 17.309 | 6.077e+10 | 118 - 133 | 19 | 160.87 | 161.13 | 163.84 |
| 3305.3316 | YHKNN(N)KSWMESEF Fuc 1 Hex 4 HexNAc 4 | 9.950 | 0.017 | 14.647 | 1.168e+10 | 144 - 157 | 9 | 94.69 | 95.34 | 96.54 |
| 3652.6663 | GGFN(N)FSQILPDPSKPSKRSF Fuc 1 Hex 3 HexNAc 4 | 9.669 | 0.017 | 13.488 | 2.248e+09 | 797 - 817 | 6 | 163.44 | 163.96 | 164.36 |
| 3960.5514 | SN(N)NSIAIPTN(N)F Fuc 1 Hex 10 HexNAc 5 | 9.667 | 0.017 | -4.946 | 5.745e+07 | 707 - 718 | 1 | 102.94 | 103.24 | 103.24 |
| 5205.2027 | QTRAGC(C)LIGAEHVN(N)NSYEC(C)DIPIGAGIC(C)ASY Fuc 1 Hex 4 HexNAc 5 | 9.562 | 0.020 | 15.838 | 1.126e+10 | 643 - 674 | 7 | 164.85 | 165.41 | 166.07 |
| 4034.8544 | RVQPTESIVRFPN(N)ITNLC(C)PF Fuc 1 Hex 3 HexNAc 5 | 9.555 | 0.020 | 16.283 | 4.794e+10 | 318 - 338 | 16 | 191.93 | 192.74 | 195.36 |
| 3993.8287 | RVQPTESIVRFPN(N)ITNLC(C)PF Fuc 1 Hex 4 HexNAc 4 | 9.542 | 0.020 | 13.989 | 1.638e+10 | 318 - 338 | 9 | 191.41 | 192.72 | 195.17 |
| 3251.4887 | LIVNN(N)ATNVVIKVC(C)EF Hex 5 HexNAc 3 | 9.513 | 0.020 | 18.016 | 5.543e+09 | 117 - 133 | 3 | 179.22 | 179.47 | 180.18 |
| 3789.6895 | GGFN(N)FSQILPDPSKPSKRSF Hex 6 HexNAc 3 | 9.493 | 0.020 | 13.114 | 1.123e+09 | 797 - 817 | 5 | 164.77 | 165.21 | 165.45 |
| 4611.9575 | QTRAGC(C)LIGAEHVN(N)NSYEC(C)DIPIGAGIC(C)ASY Hex 5 HexNAc 2 | 9.465 | 0.020 | 16.780 | 3.655e+09 | 643 - 674 | 3 | 166.53 | 166.70 | 167.01 |
| 3438.5752 | LIVNN(N)ATNVVIKVC(C)EF Fuc 1 Hex 4 HexNAc 4 | 9.456 | 0.020 | 12.087 | 1.417e+09 | 117 - 133 | 7 | 178.03 | 178.58 | 179.82 |
| 3831.7753 | RVQPTESIVRFPN(N)ITNLC(C)PF Fuc 1 Hex 3 HexNAc 4 | 9.445 | 0.020 | 17.265 | 4.475e+10 | 318 - 338 | 12 | 192.47 | 193.17 | 195.33 |
| 4155.8821 | RVQPTESIVRFPN(N)ITNLC(C)PF Fuc 1 Hex 5 HexNAc 4 | 9.428 | 0.020 | 17.927 | 7.511e+09 | 318 - 338 | 9 | 191.15 | 191.58 | 192.68 |
| 4238.6763 | EPLVDLPIGIN(N)ITRF Fuc 1 Hex 6 HexNAc 7 | 9.410 | 0.020 | -7.092 | 2.356e+08 | 223 - 238 | 2 | 219.54 | 219.81 | 239.53 |
| 4467.0546 | RVQPTESIVRFPN(N)ITNLC(C)PFGEVF Fuc 1 Hex 3 HexNAc 5 | 9.396 | 0.020 | 17.638 | 9.033e+10 | 318 - 342 | 17 | 215.94 | 216.85 | 221.07 |
| 5002.1225 | QTRAGC(C)LIGAEHVN(N)NSYEC(C)DIPIGAGIC(C)ASY Fuc 1 Hex 4 HexNAc 4 | 9.378 | 0.020 | 12.786 | 7.977e+09 | 643 - 674 | 6 | 165.29 | 165.71 | 167.04 |
| 3594.6414 | N(N)FSQILPDPSKPSKRSF Fuc 1 Hex 3 HexNAc 5 | 9.342 | 0.020 | 12.909 | 1.415e+09 | 800 - 817 | 5 | 124.16 | 124.77 | 125.19 |
| 4192.7677 | HAIHVSGTN(N)GTKRF Fuc 1 Hex 5 HexNAc 7 Neu5Ac 1 | 9.267 | 0.021 | 10.569 | 1.047e+09 | 65 - 79 | 21 | 60.34 | 62.94 | 63.17 |
| 3528.6732 | N(N)FSQILPDPSKPSKRSF Hex 6 HexNAc 3 | 9.253 | 0.021 | -4.681 | 2.916e+09 | 800 - 817 | 5 | 124.13 | 124.73 | 241.68 |
| 3828.6569 | FHAIHVSGTN(N)GTKRF Hex 4 HexNAc 6 Neu5Ac 1 | 9.247 | 0.021 | 12.302 | 4.425e+08 | 64 - 79 | 2 | 52.39 | 53.68 | 53.83 |
| 4426.0311 | RVQPTESIVRFPN(N)ITNLC(C)PFGEVF Fuc 1 Hex 4 HexNAc 4 | 9.231 | 0.021 | 16.395 | 3.642e+10 | 318 - 342 | 11 | 215.33 | 216.50 | 219.54 |
| 2980.2164 | HKNN(N)KSWMESEF Fuc 1 Hex 3 HexNAc 4 | 9.212 | 0.021 | 16.615 | 3.791e+10 | 145 - 157 | 17 | 91.28 | 92.42 | 94.19 |
| 4000.5889 | SN(N)NSIAIPTN(N)F Fuc 1 Hex 9 HexNAc 6 | 9.200 | 0.021 | 0.000 | 0.000e+00 | 707 - 718 | 2 | - | - | - |
| 3819.6782 | N(N)FSQILPDPSKPSKRSF Hex 6 HexNAc 3 Neu5Ac 1 | 9.195 | 0.021 | -3.173 | 6.164e+09 | 800 - 817 | 7 | 138.98 | 139.42 | 179.76 |
| 3627.6416 | GGFN(N)FSQILPDPSKPSKRSF Hex 5 HexNAc 3 | 9.152 | 0.021 | 12.233 | 3.627e+08 | 797 - 817 | 4 | 165.05 | 165.33 | 165.64 |
| 3954.5830 | HAIHVSGTN(N)GTKRF @sulfate 1 Fuc 1 Hex 5 HexNAc 4 Neu5Ac 2 | 9.094 | 0.021 | 12.800 | 1.483e+08 | 65 - 79 | 3 | 82.85 | 83.02 | 83.50 |
| 3776.7761 | SALEPLVDLPIGIN(N)ITRF Fuc 1 Hex 4 HexNAc 5 | 9.072 | 0.021 | 15.677 | 3.682e+08 | 220 - 238 | 4 | 238.07 | 238.68 | 238.95 |
| 5043.1477 | QTRAGC(C)LIGAEHVN(N)NSYEC(C)DIPIGAGIC(C)ASY Fuc 1 Hex 3 HexNAc 5 | 9.053 | 0.021 | 13.287 | 3.338e+10 | 643 - 674 | 12 | 165.54 | 166.16 | 167.72 |
| 4284.9239 | RVQPTESIVRFPN(N)ITNLC(C)PF Fuc 1 Hex 4 HexNAc 4 Neu5Ac 1 | 9.000 | 0.022 | 16.098 | 5.238e+10 | 318 - 338 | 16 | 209.04 | 209.86 | 210.94 |
| 4974.2272 | VPAQEKN(N)FTTAPAIC(C)HDGKAHFPREGVF Fuc 1 Hex 3 HexNAc 6 | 8.958 | 0.022 | 16.385 | 7.738e+09 | 1067 - 1095 | 9 | 117.17 | 117.51 | 118.37 |
| 2976.3531 | IVNN(N)ATNVVIKVC(C)EF Hex 4 HexNAc 3 | 8.937 | 0.022 | 15.787 | 2.304e+10 | 118 - 133 | 10 | 161.13 | 161.60 | 162.29 |
| 5913.5400 | VPAQEKN(N)FTTAPAIC(C)HDGKAHFPREGVF Fuc 1 Hex 7 HexNAc 6 Neu5Ac 1 | 8.874 | 0.022 | 16.154 | 1.591e+09 | 1067 - 1095 | 4 | 123.80 | 124.24 | 124.66 |
| 4588.0847 | RVQPTESIVRFPN(N)ITNLC(C)PFGEVF Fuc 1 Hex 5 HexNAc 4 | 8.847 | 0.022 | 18.122 | 1.669e+10 | 318 - 342 | 8 | 214.38 | 215.48 | 215.91 |
| 4730.1205 | VPAQEKN(N)FTTAPAIC(C)HDGKAHFPREGVF Fuc 1 Hex 4 HexNAc 4 | 8.807 | 0.022 | 13.956 | 2.527e+10 | 1067 - 1095 | 16 | 117.20 | 117.85 | 119.59 |
| 3448.5766 | N(N)FSQILPDPSKPSKRSF Hex 3 HexNAc 5 | 8.794 | 0.022 | 12.095 | 5.992e+08 | 800 - 817 | 7 | 126.04 | 126.41 | 126.76 |
| 3138.4056 | IVNN(N)ATNVVIKVC(C)EF Hex 5 HexNAc 3 | 8.751 | 0.022 | 16.037 | 2.878e+10 | 118 - 133 | 8 | 159.79 | 160.49 | 161.64 |
| 3630.7037 | RVQPTESIVRFPN(N)ITNL Fuc 1 Hex 3 HexNAc 5 | 8.714 | 0.022 | 15.535 | 2.283e+10 | 318 - 335 | 11 | 167.59 | 168.54 | 170.39 |
| 4105.8137 | GGFN(N)FSQILPDPSKPSKRSF Fuc 1 Hex 4 HexNAc 4 Neu5Ac 1 | 8.682 | 0.022 | 15.422 | 1.071e+09 | 797 - 817 | 5 | 178.08 | 178.71 | 179.21 |
| 4629.1091 | RVQPTESIVRFPN(N)ITNLC(C)PFGEVF Fuc 1 Hex 4 HexNAc 5 | 8.677 | 0.022 | 14.371 | 5.697e+10 | 318 - 342 | 18 | 215.11 | 216.08 | 219.30 |
| 4791.1638 | RVQPTESIVRFPN(N)ITNLC(C)PFGEVF Fuc 1 Hex 5 HexNAc 5 | 8.661 | 0.022 | 13.528 | 1.513e+10 | 318 - 342 | 8 | 214.16 | 215.60 | 215.91 |
| 3537.5507 | FHAIHVSGTN(N)GTKRF Hex 4 HexNAc 6 | 8.602 | 0.022 | 6.292 | 4.885e+07 | 64 - 79 | 3 | 54.59 | 54.59 | 54.88 |
| 3596.4285 | YHKNN(N)KSWMESEF Fuc 1 Hex 4 HexNAc 4 Neu5Ac 1 | 8.600 | 0.022 | 15.459 | 2.649e+09 | 144 - 157 | 3 | 107.45 | 107.73 | 107.96 |
| 4851.1529 | VPAQEKN(N)FTTAPAIC(C)HDGKAHFPREGVF Fuc 1 Hex 6 HexNAc 3 | 8.568 | 0.022 | 15.549 | 1.464e+09 | 1067 - 1095 | 6 | 117.17 | 117.44 | 117.67 |
| 3715.6629 | N(N)FSQILPDPSKPSKRSF Fuc 1 Hex 5 HexNAc 4 | 8.555 | 0.022 | 13.970 | 1.250e+09 | 800 - 817 | 4 | 123.08 | 123.55 | 124.49 |
| 3569.6081 | N(N)FSQILPDPSKPSKRSF Hex 5 HexNAc 4 | 8.500 | 0.023 | 13.554 | 3.401e+08 | 800 - 817 | 4 | 124.20 | 124.60 | 124.87 |
| 3470.5283 | IVNN(N)ATNVVIKVC(C)EF Hex 4 HexNAc 4 Neu5Ac 1 | 8.497 | 0.023 | 14.495 | 5.837e+09 | 118 - 133 | 10 | 178.92 | 180.25 | 180.83 |
| 5455.2667 | QTRAGC(C)LIGAEHVN(N)NSYEC(C)DIPIGAGIC(C)ASY Fuc 1 Hex 5 HexNAc 4 Neu5Ac 1 | 8.496 | 0.023 | 15.835 | 2.562e+10 | 643 - 674 | 12 | 176.72 | 177.55 | 180.18 |
| 5164.1722 | QTRAGC(C)LIGAEHVN(N)NSYEC(C)DIPIGAGIC(C)ASY Fuc 1 Hex 5 HexNAc 4 | 8.485 | 0.023 | 17.669 | 1.237e+10 | 643 - 674 | 3 | 164.74 | 165.00 | 165.41 |
| 3814.7251 | GGFN(N)FSQILPDPSKPSKRSF Fuc 1 Hex 4 HexNAc 4 | 8.400 | 0.023 | 12.712 | 7.729e+08 | 797 - 817 | 4 | 163.10 | 163.55 | 163.92 |
| 2761.2191 | LHVTYVPAQEKN(N)F Hex 5 HexNAc 2 | 8.310 | 0.023 | 13.810 | 8.226e+08 | 1062 - 1075 | 1 | 111.42 | 111.53 | 111.61 |
| 4734.1407 | RVQPTESIVRFPN(N)ITNLC(C)PFGEVF Fuc 2 Hex 5 HexNAc 4 | 8.304 | 0.023 | 17.576 | 3.851e+09 | 318 - 342 | 4 | 214.62 | 215.17 | 215.83 |
| 3601.5412 | HAIHVSGTN(N)GTKRF Hex 6 HexNAc 4 Neu5Ac 1 | 8.289 | 0.023 | 0.921 | 1.930e+07 | 65 - 79 | 3 | 63.41 | 63.41 | 63.81 |
| 3542.5860 | LIVNN(N)ATNVVIKVC(C)EF Hex 5 HexNAc 3 Neu5Ac 1 | 8.195 | 0.023 | 15.207 | 3.957e+09 | 117 - 133 | 8 | 197.74 | 197.93 | 199.69 |
| 4488.0032 | RVQPTESIVRFPN(N)ITNLC(C)PF Fuc 1 Hex 4 HexNAc 5 Neu5Ac 1 | 8.188 | 0.023 | 12.567 | 2.250e+10 | 318 - 338 | 14 | 208.00 | 208.32 | 210.92 |
| 3789.5131 | SNNSIAIPTN(N)F Fuc 2 Hex 5 HexNAc 6 Neu5Ac 1 | 8.186 | 0.023 | 0.000 | 0.000e+00 | 707 - 718 | 1 | - | - | - |
| 5257.3091 | VPAQEKN(N)FTTAPAIC(C)HDGKAHFPREGVF Fuc 1 Hex 6 HexNAc 5 | 8.148 | 0.023 | 18.035 | 8.684e+09 | 1067 - 1095 | 6 | 116.21 | 116.83 | 117.69 |
| 4572.0873 | RVQPTESIVRFPN(N)ITNLC(C)PFGEVF Fuc 2 Hex 4 HexNAc 4 | 8.121 | 0.023 | 13.662 | 7.873e+09 | 318 - 342 | 5 | 215.17 | 216.24 | 216.85 |
| 5367.2522 | QTRAGC(C)LIGAEHVN(N)NSYEC(C)DIPIGAGIC(C)ASY Fuc 1 Hex 5 HexNAc 5 | 8.052 | 0.024 | 13.596 | 1.542e+10 | 643 - 674 | 4 | 164.33 | 165.02 | 165.54 |
| 3372.5174 | LIVNN(N)ATNVVIKVC(C)EF Hex 7 HexNAc 2 | 8.036 | 0.024 | 17.240 | 6.186e+08 | 117 - 133 | 2 | 178.50 | 178.78 | 179.14 |
| 4237.9331 | RVQPTESIVRFPN(N)ITNLC(C)PF Fuc 1 Hex 3 HexNAc 6 | 8.025 | 0.024 | 15.041 | 3.352e+09 | 318 - 338 | 5 | 191.68 | 192.47 | 192.76 |
| 3276.5208 | LIVNN(N)ATNVVIKVC(C)EF Fuc 1 Hex 3 HexNAc 4 | 7.994 | 0.024 | 15.764 | 2.479e+09 | 117 - 133 | 5 | 178.96 | 179.25 | 180.18 |
| 3589.6783 | RVQPTESIVRFPN(N)ITNL Fuc 1 Hex 4 HexNAc 4 | 7.927 | 0.024 | 15.622 | 8.778e+09 | 318 - 335 | 11 | 167.29 | 168.20 | 169.80 |
| 4032.7436 | FHAIHVSGTN(N)GTKRF Fuc 2 Hex 4 HexNAc 7 | 7.913 | 0.024 | -4.063 | 2.596e+07 | 64 - 79 | 2 | 53.88 | 53.88 | 53.97 |
| 4301.9400 | RVQPTESIVRFPN(N)ITNLC(C)PF Fuc 2 Hex 5 HexNAc 4 | 7.901 | 0.024 | 16.136 | 1.879e+09 | 318 - 338 | 4 | 190.83 | 191.24 | 192.27 |
| 4342.9671 | RVQPTESIVRFPN(N)ITNLC(C)PF Fuc 2 Hex 4 HexNAc 5 | 7.888 | 0.024 | 15.627 | 2.951e+09 | 318 - 338 | 7 | 190.96 | 191.27 | 192.72 |
| 2845.2343 | KTPPIKDFGGFN(N)F Hex 6 HexNAc 2 | 7.878 | 0.024 | 15.352 | 9.608e+08 | 789 - 802 | 4 | 154.58 | 155.09 | 155.41 |
| 4818.1431 | VPAQEKN(N)FTTAPAIC(C)HDGKAHFPREGVF Fuc 1 Hex 4 HexNAc 3 Neu5Ac 1 | 7.852 | 0.024 | 14.726 | 2.046e+09 | 1067 - 1095 | 3 | 126.39 | 126.64 | 126.87 |
| 3512.5813 | N(N)FSQILPDPSKPSKRSF Fuc 1 Hex 5 HexNAc 3 | 7.836 | 0.024 | 12.333 | 1.396e+08 | 800 - 817 | 2 | 124.10 | 124.73 | 124.77 |
| 4419.6882 | SN(N)NSIAIPTN(N)F Hex 15 HexNAc 4 | 7.816 | 0.024 | 15.442 | 2.699e+09 | 707 - 718 | 3 | 152.02 | 152.20 | 152.56 |
| 4533.8537 | HAIHVSGTN(N)GTKRF Fuc 1 Hex 6 HexNAc 5 Neu5Ac 3 | 7.808 | 0.024 | 0.000 | 0.000e+00 | 65 - 79 | 1 | - | - | - |
| 4446.9745 | RVQPTESIVRFPN(N)ITNLC(C)PF Fuc 1 Hex 5 HexNAc 4 Neu5Ac 1 | 7.786 | 0.024 | 15.672 | 8.618e+10 | 318 - 338 | 14 | 207.59 | 208.71 | 209.53 |
| 4937.0117 | HAIHVSGTN(N)GTKRF Fuc 1 Hex 6 HexNAc 7 Neu5Ac 3 | 7.757 | 0.025 | 11.574 | 1.628e+09 | 65 - 79 | 9 | 79.28 | 80.66 | 81.06 |
| 3954.8098 | RVQPTESIVRFPN(N)ITNL Fuc 1 Hex 5 HexNAc 5 | 7.722 | 0.026 | 13.020 | 7.627e+09 | 318 - 335 | 8 | 166.45 | 167.21 | 167.72 |
| 5244.3130 | RVQPTESIVRFPN(N)ITNLC(C)PFGEVF Fuc 1 Hex 6 HexNAc 5 Neu5Ac 1 | 7.708 | 0.026 | 14.533 | 1.778e+10 | 318 - 342 | 6 | 230.65 | 232.15 | 232.73 |
| 3603.6654 | RVQPTESIVRFPN(N)ITNLC(C)PF Hex 5 HexNAc 2 | 7.682 | 0.026 | 14.850 | 2.441e+09 | 318 - 338 | 5 | 195.85 | 196.01 | 196.83 |
| 4738.0750 | RVQPTESIVRFPN(N)ITNLC(C)PF Fuc 1 Hex 5 HexNAc 4 Neu5Ac 2 | 7.659 | 0.026 | 14.547 | 1.153e+10 | 318 - 338 | 6 | 227.22 | 228.73 | 229.07 |
| 3778.5949 | HAIHVSGTN(N)GTKRF Fuc 1 Hex 8 HexNAc 4 | 7.644 | 0.026 | 17.540 | 4.966e+08 | 65 - 79 | 1 | 86.08 | 86.29 | 86.49 |
| 4717.1246 | RVQPTESIVRFPN(N)ITNLC(C)PFGEVF Fuc 1 Hex 4 HexNAc 4 Neu5Ac 1 | 7.588 | 0.026 | 18.144 | 7.583e+10 | 318 - 342 | 19 | 233.11 | 234.60 | 238.31 |
| 4161.6408 | SN(N)NSIAIPTN(N)F Fuc 1 Hex 10 HexNAc 6 | 7.561 | 0.026 | 15.672 | 1.304e+09 | 707 - 718 | 3 | 154.33 | 154.63 | 155.34 |
| 3855.7486 | GGFN(N)FSQILPDPSKPSKRSF Fuc 1 Hex 3 HexNAc 5 | 7.548 | 0.026 | 14.261 | 1.007e+09 | 797 - 817 | 5 | 163.22 | 163.66 | 163.95 |
| 3528.5723 | IVNN(N)ATNVVIKVC(C)EF Fuc 1 Hex 4 HexNAc 5 | 7.538 | 0.026 | 12.821 | 1.714e+09 | 118 - 133 | 4 | 158.75 | 159.44 | 160.04 |
| 4080.7816 | GGFN(N)FSQILPDPSKPSKRSF Hex 6 HexNAc 3 Neu5Ac 1 | 7.525 | 0.026 | 12.704 | 1.400e+09 | 797 - 817 | 6 | 179.79 | 180.47 | 180.90 |
| 3122.4111 | IVNN(N)ATNVVIKVC(C)EF Fuc 1 Hex 4 HexNAc 3 | 7.524 | 0.026 | 16.122 | 9.301e+09 | 118 - 133 | 10 | 159.97 | 160.23 | 161.60 |
| 4562.0415 | RVQPTESIVRFPN(N)ITNLC(C)PF Fuc 1 Hex 5 HexNAc 6 | 7.469 | 0.026 | 14.375 | 1.679e+09 | 318 - 338 | 3 | 190.27 | 191.24 | 192.11 |
| 3267.4481 | IVNN(N)ATNVVIKVC(C)EF Hex 4 HexNAc 3 Neu5Ac 1 | 7.461 | 0.026 | 13.277 | 2.238e+09 | 118 - 133 | 4 | 179.76 | 180.87 | 181.81 |
| 2648.1374 | HVTYVPAQEKN(N)F Hex 5 HexNAc 2 | 7.430 | 0.026 | 17.842 | 1.357e+09 | 1063 - 1075 | 3 | 90.99 | 91.26 | 91.56 |
| 4645.9154 | HAIHVSGTN(N)GTKRF Fuc 1 Hex 6 HexNAc 7 Neu5Ac 2 | 7.420 | 0.026 | 11.224 | 3.967e+09 | 65 - 79 | 22 | 68.83 | 71.79 | 75.29 |
| 3017.3805 | IVNN(N)ATNVVIKVC(C)EF Hex 3 HexNAc 4 | 7.409 | 0.026 | 14.657 | 1.182e+10 | 118 - 133 | 7 | 161.22 | 161.60 | 162.05 |
| 3382.5129 | IVNN(N)ATNVVIKVC(C)EF Hex 4 HexNAc 5 | 7.338 | 0.026 | 11.350 | 1.441e+09 | 118 - 133 | 5 | 159.61 | 160.17 | 160.85 |
| 4031.7371 | FHAIHVSGTN(N)GTKRF Hex 4 HexNAc 7 Neu5Ac 1 | 7.320 | 0.026 | 13.417 | 1.845e+09 | 64 - 79 | 5 | 52.17 | 53.69 | 54.49 |
| 7409.0268 | SN(N)VTWFHAIHVSGTN(N)GTKRF Fuc 1 Hex 11 HexNAc 13 Neu5Ac 2 | 7.319 | 0.026 | 10.784 | 1.465e+08 | 59 - 79 | 3 | 73.51 | 73.59 | 73.74 |
| 3179.4333 | IVNN(N)ATNVVIKVC(C)EF Hex 4 HexNAc 4 | 7.313 | 0.026 | 13.734 | 9.669e+09 | 118 - 133 | 9 | 159.97 | 160.78 | 161.64 |
| 3698.6924 | N(N)FSQILPDPSKPSKRSF Hex 4 HexNAc 4 Neu5Ac 1 | 7.290 | 0.026 | 11.592 | 2.168e+09 | 800 - 817 | 6 | 139.84 | 140.30 | 140.52 |
| 2814.3009 | IVNN(N)ATNVVIKVC(C)EF Hex 3 HexNAc 3 | 7.256 | 0.026 | 16.209 | 1.675e+10 | 118 - 133 | 5 | 162.43 | 162.67 | 163.86 |
| 3220.4579 | IVNN(N)ATNVVIKVC(C)EF Hex 3 HexNAc 5 | 7.245 | 0.026 | 15.745 | 2.820e+09 | 118 - 133 | 4 | 160.80 | 161.41 | 161.69 |
| 3918.7306 | GGFN(N)FSQILPDPSKPSKRSF Hex 5 HexNAc 3 Neu5Ac 1 | 7.242 | 0.026 | 15.782 | 1.105e+09 | 797 - 817 | 6 | 181.11 | 181.51 | 181.85 |
| 3346.3583 | YHKNN(N)KSWMESEF Fuc 1 Hex 3 HexNAc 5 | 7.216 | 0.026 | 14.648 | 2.801e+10 | 144 - 157 | 15 | 94.89 | 96.38 | 97.61 |
| 3097.3792 | IVNN(N)ATNVVIKVC(C)EF Hex 6 HexNAc 2 | 7.211 | 0.026 | 15.894 | 3.040e+09 | 118 - 133 | 4 | 159.44 | 160.27 | 160.87 |
| 3048.4094 | LIVNN(N)ATNVVIKVC(C)EF Hex 5 HexNAc 2 | 7.190 | 0.026 | 15.217 | 9.290e+09 | 117 - 133 | 8 | 179.51 | 180.06 | 180.95 |
| 3407.5520 | N(N)FSQILPDPSKPSKRSF Hex 4 HexNAc 4 | 7.185 | 0.026 | 10.312 | 3.381e+08 | 800 - 817 | 3 | 125.76 | 125.95 | 126.41 |
| 4920.2044 | RVQPTESIVRFPN(N)ITNLC(C)PFGEVF Fuc 1 Hex 4 HexNAc 5 Neu5Ac 1 | 7.185 | 0.026 | 16.153 | 1.973e+10 | 318 - 342 | 9 | 232.53 | 232.88 | 235.22 |
| 3548.6643 | SALEPLVDLPIGIN(N)ITRF Hex 6 HexNAc 3 | 7.183 | 0.026 | 15.520 | 1.282e+09 | 220 - 238 | 4 | 240.82 | 241.25 | 241.73 |
| 3927.7141 | FHAIHVSGTN(N)GTKRF Fuc 1 Hex 3 HexNAc 8 | 7.113 | 0.026 | 6.719 | 3.471e+07 | 64 - 79 | 1 | 53.76 | 53.76 | 54.08 |
| 3942.7033 | HAIHVSGTN(N)GTKRF Fuc 1 Hex 4 HexNAc 8 | 7.108 | 0.026 | 10.068 | 1.039e+08 | 65 - 79 | 3 | 53.58 | 53.95 | 54.07 |
| 3210.4627 | LIVNN(N)ATNVVIKVC(C)EF Hex 6 HexNAc 2 | 7.090 | 0.026 | 16.063 | 8.388e+08 | 117 - 133 | 2 | 179.14 | 179.44 | 180.00 |
| 3042.4200 | N(N)FSQILPDPSKPSKRSF Hex 3 HexNAc 3 | 7.067 | 0.026 | 11.626 | 2.694e+09 | 800 - 817 | 7 | 126.82 | 127.38 | 127.90 |
| 5298.3343 | VPAQEKN(N)FTTAPAIC(C)HDGKAHFPREGVF Fuc 1 Hex 5 HexNAc 6 | 7.066 | 0.026 | 12.613 | 3.516e+09 | 1067 - 1095 | 6 | 116.44 | 116.74 | 117.69 |
| 3143.2810 | YHKNN(N)KSWMESEF Fuc 1 Hex 3 HexNAc 4 | 6.985 | 0.029 | 17.333 | 2.028e+10 | 144 - 157 | 12 | 95.27 | 95.98 | 97.37 |
| 4007.7061 | HAIHVSGTN(N)GTKRF Fuc 2 Hex 6 HexNAc 6 | 6.972 | 0.030 | 0.000 | 0.000e+00 | 65 - 79 | 1 | - | - | - |
| 3465.5796 | GGFN(N)FSQILPDPSKPSKRSF Hex 4 HexNAc 3 | 6.968 | 0.030 | 10.867 | 4.001e+08 | 797 - 817 | 1 | 166.22 | 166.39 | 166.82 |
| 4953.2124 | RVQPTESIVRFPN(N)ITNLC(C)PFGEVF Fuc 1 Hex 6 HexNAc 5 | 6.967 | 0.030 | 17.710 | 4.019e+09 | 318 - 342 | 4 | 213.85 | 214.62 | 215.76 |
| 3007.2855 | KTPPIKDFGGFN(N)F Hex 7 HexNAc 2 | 6.840 | 0.030 | 16.650 | 1.420e+09 | 789 - 802 | 4 | 153.86 | 154.20 | 154.83 |
| 4209.7840 | HAIHVSGTN(N)GTKRF Fuc 2 Hex 6 HexNAc 7 | 6.783 | 0.030 | 12.068 | 1.486e+08 | 65 - 79 | 6 | 52.39 | 53.14 | 53.51 |
| 2887.2872 | FHAIHVSGTN(N)GTKRF Hex 5 HexNAc 2 | 6.767 | 0.030 | 12.972 | 1.461e+07 | 64 - 79 | 2 | 74.88 | 75.08 | 75.20 |
| 2683.1807 | KTPPIKDFGGFN(N)F Hex 5 HexNAc 2 | 6.753 | 0.030 | 15.648 | 3.918e+10 | 789 - 802 | 8 | 155.29 | 155.69 | 156.46 |
| 4263.9755 | RVQPTESIVRFPN(N)ITNLC(C)PFGEVF Fuc 1 Hex 3 HexNAc 4 | 6.664 | 0.030 | 19.672 | 1.037e+11 | 318 - 342 | 14 | 216.20 | 216.99 | 221.04 |
| 3561.5593 | EPLVDLPIGIN(N)ITRF @sulfate 1 Hex 6 HexNAc 4 | 6.573 | 0.030 | 10.338 | 6.927e+08 | 223 - 238 | 1 | 240.72 | 241.33 | 241.73 |
| 4505.0179 | RVQPTESIVRFPN(N)ITNLC(C)PF Fuc 2 Hex 5 HexNAc 5 | 6.532 | 0.031 | 15.687 | 1.172e+09 | 318 - 338 | 6 | 190.72 | 191.47 | 192.15 |
| 5188.0808 | HAIHVSGTN(N)GTKRF Fuc 1 Hex 7 HexNAc 6 Neu5Ac 4 | 6.466 | 0.031 | -3.437 | 1.556e+08 | 65 - 79 | 1 | 92.19 | 92.20 | 92.21 |
| 3897.7880 | RVQPTESIVRFPN(N)ITNL Fuc 2 Hex 5 HexNAc 4 | 6.432 | 0.032 | 13.279 | 1.668e+09 | 318 - 335 | 3 | 166.63 | 166.87 | 167.20 |
| 4557.9033 | HAIHVSGTN(N)GTKRF Fuc 1 Hex 6 HexNAc 8 Neu5Ac 1 | 6.387 | 0.033 | 12.407 | 1.583e+08 | 65 - 79 | 5 | 61.96 | 62.17 | 62.45 |
| 3341.4862 | IVNN(N)ATNVVIKVC(C)EF Hex 5 HexNAc 4 | 6.381 | 0.033 | 15.664 | 2.945e+09 | 118 - 133 | 3 | 159.31 | 159.58 | 160.44 |
| 3529.5814 | N(N)FSQILPDPSKPSKRSF Hex 6 HexNAc 3 | 6.368 | 0.033 | 0.000 | 0.000e+00 | 800 - 817 | 1 | - | - | - |
| 3854.7287 | EPLVDLPIGIN(N)ITRF Fuc 2 Hex 4 HexNAc 6 | 6.347 | 0.034 | 10.849 | 3.245e+09 | 223 - 238 | 2 | 239.22 | 239.53 | 240.18 |
| 3277.4522 | FHAIHVSGTN(N)GTKRF Fuc 1 Hex 4 HexNAc 4 | 6.339 | 0.034 | 10.339 | 9.804e+07 | 64 - 79 | 5 | 74.73 | 75.64 | 75.92 |
| 5246.2274 | QTRAGC(C)LIGAEHVN(N)NSYEC(C)DIPIGAGIC(C)ASY Fuc 1 Hex 3 HexNAc 6 | 6.337 | 0.034 | 14.883 | 6.515e+09 | 643 - 674 | 8 | 165.05 | 165.33 | 166.07 |
| 5622.4401 | VPAQEKN(N)FTTAPAIC(C)HDGKAHFPREGVF Fuc 1 Hex 7 HexNAc 6 | 6.331 | 0.034 | 18.242 | 2.081e+09 | 1067 - 1095 | 2 | 116.16 | 116.44 | 116.92 |
| 5438.2425 | Q(P)TRAGC(C)LIGAEHVN(N)NSYEC(C)DIPIGAGIC(C)ASY Fuc 1 Hex 5 HexNAc 4 Neu5Ac 1 | 6.301 | 0.034 | 16.657 | 1.069e+09 | 643 - 674 | 3 | 198.33 | 199.49 | 199.77 |
| 3632.5793 | IVNN(N)ATNVVIKVC(C)EF Hex 5 HexNAc 4 Neu5Ac 1 | 6.282 | 0.034 | 15.123 | 4.921e+09 | 118 - 133 | 14 | 177.61 | 177.90 | 180.04 |
| 5408.2813 | QTRAGC(C)LIGAEHVN(N)NSYEC(C)DIPIGAGIC(C)ASY Fuc 1 Hex 4 HexNAc 6 | 6.244 | 0.034 | 14.549 | 7.945e+09 | 643 - 674 | 6 | 164.48 | 164.91 | 165.45 |
| 4339.9600 | VPAQEKN(N)FTTAPAIC(C)HDGKAHFPREGVF Hex 5 HexNAc 2 | 6.206 | 0.034 | 16.331 | 6.693e+09 | 1067 - 1095 | 8 | 117.83 | 118.07 | 118.64 |
| 3567.5358 | HAIHVSGTN(N)GTKRF Fuc 2 Hex 4 HexNAc 4 Neu5Ac 1 | 6.203 | 0.034 | 11.303 | 1.816e+07 | 65 - 79 | 1 | 62.81 | 62.92 | 63.15 |
| 3471.5505 | IVNN(N)ATNVVIKVC(C)EF Fuc 2 Hex 4 HexNAc 4 | 6.192 | 0.034 | 13.847 | 2.171e+09 | 118 - 133 | 3 | 158.80 | 159.02 | 159.93 |
| 4333.9244 | RVQPTESIVRFPN(N)ITNL Fuc 1 Hex 5 HexNAc 4 Neu5Ac 2 | 6.187 | 0.034 | 15.008 | 8.135e+09 | 318 - 335 | 7 | 204.07 | 204.29 | 205.07 |
| 3508.4109 | YHKNN(N)KSWMESEF Fuc 1 Hex 4 HexNAc 5 | 6.184 | 0.034 | 15.998 | 3.362e+10 | 144 - 157 | 16 | 93.52 | 95.60 | 96.59 |
| 5082.2560 | RVQPTESIVRFPN(N)ITNLC(C)PFGEVF Fuc 1 Hex 5 HexNAc 5 Neu5Ac 1 | 6.164 | 0.034 | 15.636 | 1.110e+10 | 318 - 342 | 9 | 231.66 | 233.39 | 234.09 |
| 3778.6368 | IVNN(N)ATNVVIKVC(C)EF Fuc 1 Hex 5 HexNAc 4 Neu5Ac 1 | 6.123 | 0.034 | 15.957 | 1.028e+10 | 118 - 133 | 13 | 176.38 | 176.64 | 179.33 |
| 4937.2190 | RVQPTESIVRFPN(N)ITNLC(C)PFGEVF Fuc 2 Hex 5 HexNAc 5 | 6.121 | 0.034 | 16.423 | 2.131e+09 | 318 - 342 | 4 | 214.35 | 215.39 | 215.76 |
| 5170.2757 | RVQPTESIVRFPN(N)ITNLC(C)PFGEVF Fuc 1 Hex 5 HexNAc 4 Neu5Ac 2 | 6.114 | 0.034 | 18.409 | 1.000e+10 | 318 - 342 | 7 | 252.47 | 253.86 | 255.41 |
| 4670.1335 | RVQPTESIVRFPN(N)ITNLC(C)PFGEVF Fuc 1 Hex 3 HexNAc 6 | 6.089 | 0.034 | 13.328 | 1.448e+10 | 318 - 342 | 6 | 215.51 | 216.25 | 217.35 |
| 4500.8809 | HAIHVSGTN(N)GTKRF Fuc 2 Hex 6 HexNAc 7 Neu5Ac 1 | 6.087 | 0.034 | 10.089 | 4.569e+08 | 65 - 79 | 13 | 59.88 | 60.93 | 62.87 |
| 4407.9587 | RVQPTESIVRFPN(N)ITNL Fuc 1 Hex 6 HexNAc 5 Neu5Ac 1 | 6.084 | 0.035 | 14.112 | 8.268e+09 | 318 - 335 | 6 | 182.63 | 183.27 | 184.21 |
| 3926.7070 | HAIHVSGTN(N)GTKRFDNPVLPF @sulfate 1 Hex 7 HexNAc 2 | 6.076 | 0.035 | 13.432 | 2.152e+09 | 65 - 86 | 10 | 52.94 | 53.22 | 54.07 |
| 3901.6733 | HAIHVSGTN(N)GTKRF Fuc 1 Hex 5 HexNAc 7 | 6.065 | 0.035 | 14.952 | 2.033e+08 | 65 - 79 | 3 | 54.95 | 55.21 | 55.79 |
| 3688.6430 | LIVNN(N)ATNVVIKVC(C)EF Fuc 1 Hex 5 HexNAc 3 Neu5Ac 1 | 6.053 | 0.035 | 10.543 | 2.576e+09 | 117 - 133 | 6 | 196.35 | 197.57 | 198.33 |
| 4202.6688 | SN(N)NSIAIPTN(N)F Fuc 1 Hex 9 HexNAc 7 | 5.952 | 0.037 | 13.683 | 7.164e+08 | 707 - 718 | 4 | 154.41 | 154.83 | 155.33 |
| 3663.4891 | HAIHVSGTN(N)GTKRF @sulfate 1 Fuc 1 Hex 5 HexNAc 4 Neu5Ac 1 | 5.945 | 0.038 | 12.590 | 1.100e+08 | 65 - 79 | 2 | 71.15 | 71.30 | 71.79 |
| 3758.4813 | YHKNN(N)KSWMESEF Fuc 1 Hex 5 HexNAc 4 Neu5Ac 1 | 5.886 | 0.042 | 15.591 | 6.081e+09 | 144 - 157 | 6 | 106.63 | 107.59 | 107.86 |
| 3654.4683 | YHKNN(N)KSWMESEF Fuc 2 Hex 4 HexNAc 5 | 5.854 | 0.042 | 16.487 | 9.605e+09 | 144 - 157 | 5 | 93.33 | 93.72 | 94.44 |
| 3102.2533 | YHKNN(N)KSWMESEF Fuc 1 Hex 4 HexNAc 3 | 5.849 | 0.042 | 16.034 | 1.492e+09 | 144 - 157 | 4 | 95.48 | 95.80 | 96.31 |
| 3729.6698 | LIVNN(N)ATNVVIKVC(C)EF Fuc 1 Hex 4 HexNAc 4 Neu5Ac 1 | 5.758 | 0.042 | 13.308 | 6.335e+09 | 117 - 133 | 8 | 196.27 | 197.66 | 198.86 |
| 3792.7586 | RVQPTESIVRFPN(N)ITNL Fuc 1 Hex 4 HexNAc 5 | 5.730 | 0.042 | 12.693 | 1.528e+10 | 318 - 335 | 16 | 166.90 | 167.66 | 169.72 |
| 3413.4593 | HAIHVSGTN(N)GTKRF Fuc 1 Hex 7 HexNAc 3 | 5.723 | 0.042 | 11.802 | 2.321e+07 | 65 - 79 | 4 | 53.00 | 53.11 | 53.40 |
| 3603.5816 | HAIHVSGTN(N)GTKRF Fuc 2 Hex 6 HexNAc 4 | 5.708 | 0.042 | 7.342 | 4.497e+07 | 65 - 79 | 1 | 172.32 | 172.51 | 172.57 |
| 6767.7208 | QTRAGC(C)LIGAEHVN(N)NSYEC(C)DIPIGAGIC(C)ASY Fuc 1 Hex 7 HexNAc 6 Neu5Ac 3 | 5.692 | 0.042 | 13.344 | 4.780e+09 | 643 - 674 | 5 | 205.53 | 206.69 | 207.87 |
| 3526.5902 | LIVNN(N)ATNVVIKVC(C)EF Fuc 1 Hex 4 HexNAc 3 Neu5Ac 1 | 5.674 | 0.042 | 15.702 | 5.098e+08 | 117 - 133 | 3 | 198.12 | 198.37 | 199.12 |
| 4139.8879 | RVQPTESIVRFPN(N)ITNLC(C)PF Fuc 2 Hex 4 HexNAc 4 | 5.673 | 0.042 | 14.763 | 2.115e+09 | 318 - 338 | 7 | 190.93 | 192.35 | 192.74 |
| 6476.6250 | QTRAGC(C)LIGAEHVN(N)NSYEC(C)DIPIGAGIC(C)ASY Fuc 1 Hex 7 HexNAc 6 Neu5Ac 2 | 5.662 | 0.042 | 14.221 | 4.084e+09 | 643 - 674 | 5 | 188.79 | 189.73 | 191.81 |
| 3585.5630 | FHAIHVSGTN(N)GTKRF Fuc 2 Hex 5 HexNAc 4 | 5.625 | 0.042 | 12.858 | 4.677e+07 | 64 - 79 | 3 | 74.35 | 74.46 | 74.68 |
| 3886.6899 | FHAIHVSGTN(N)GTKRF Fuc 1 Hex 4 HexNAc 7 | 5.621 | 0.042 | 12.952 | 9.667e+07 | 64 - 79 | 1 | 74.37 | 74.53 | 74.92 |
| 5103.2054 | RVQPTESIVRFPN(N)ITNLC(C)PF Fuc 1 Hex 6 HexNAc 5 Neu5Ac 2 | 5.591 | 0.042 | 12.734 | 4.110e+09 | 318 - 338 | 8 | 225.84 | 227.46 | 228.58 |
| 3670.6929 | EPLVDLPIGIN(N)ITRF Fuc 1 Hex 5 HexNAc 5 | 5.577 | 0.042 | -7.188 | 1.499e+07 | 223 - 238 | 1 | 241.26 | 241.29 | 241.54 |
| 4053.6876 | IGAEHVN(N)NSYEC(C)DIPIGAGIC(C)ASY Fuc 1 Hex 3 HexNAc 4 | 5.531 | 0.042 | 16.976 | 2.795e+09 | 650 - 674 | 4 | 170.33 | 170.60 | 171.81 |
| 3266.4283 | VSGNC(C)DVVIGIVN(N)NTVY Fuc 1 Hex 3 HexNAc 4 | 5.519 | 0.042 | 16.467 | 3.012e+09 | 1121 - 1138 | 3 | 193.42 | 193.71 | 194.63 |
| 4650.0539 | RVQPTESIVRFPN(N)ITNLC(C)PF Fuc 1 Hex 5 HexNAc 5 Neu5Ac 1 | 5.495 | 0.042 | 11.411 | 1.045e+10 | 318 - 338 | 9 | 207.15 | 208.17 | 208.91 |
| 3303.5310 | GGFN(N)FSQILPDPSKPSKRSF Hex 3 HexNAc 3 | 5.473 | 0.042 | 10.887 | 2.881e+08 | 797 - 817 | 3 | 166.69 | 167.10 | 167.33 |
| 4256.7668 | IGAEHVN(N)NSYEC(C)DIPIGAGIC(C)ASY Fuc 1 Hex 3 HexNAc 5 | 5.447 | 0.042 | 16.120 | 2.828e+09 | 650 - 674 | 6 | 169.69 | 170.51 | 171.85 |
| 5136.2783 | VPAQEKN(N)FTTAPAIC(C)HDGKAHFPREGVF Fuc 1 Hex 4 HexNAc 6 | 5.393 | 0.042 | 14.762 | 3.774e+09 | 1067 - 1095 | 7 | 116.80 | 117.20 | 117.93 |
| 3370.6166 | SALEPLVDLPIGIN(N)ITRF Fuc 1 Hex 4 HexNAc 3 | 5.393 | 0.042 | 12.918 | 1.229e+09 | 220 - 238 | 5 | 239.69 | 240.29 | 240.80 |
| 3756.6914 | N(N)FSQILPDPSKPSKRSF Fuc 1 Hex 4 HexNAc 5 | 5.379 | 0.042 | 13.603 | 4.857e+08 | 800 - 817 | 4 | 123.65 | 124.06 | 124.81 |
| 3844.7052 | N(N)FSQILPDPSKPSKRSF Fuc 1 Hex 4 HexNAc 4 Neu5Ac 1 | 5.371 | 0.042 | 12.147 | 6.807e+09 | 800 - 817 | 9 | 138.18 | 138.72 | 139.42 |
| 4543.0400 | VPAQEKN(N)FTTAPAIC(C)HDGKAHFPREGVF Hex 5 HexNAc 3 | 5.359 | 0.042 | 13.285 | 5.715e+08 | 1067 - 1095 | 4 | 117.67 | 117.85 | 118.33 |
| 4832.1893 | RVQPTESIVRFPN(N)ITNLC(C)PFGEVF Fuc 1 Hex 4 HexNAc 6 | 5.356 | 0.042 | 13.528 | 2.486e+09 | 318 - 342 | 4 | 214.71 | 215.39 | 215.91 |
| 3910.7197 | GGFN(N)FSQILPDPSKPSKRSF Hex 8 HexNAc 2 | 5.340 | 0.042 | 11.421 | 7.622e+08 | 797 - 817 | 3 | 164.34 | 164.64 | 164.99 |
| 2752.1061 | HKNN(N)KSWMESEF Hex 5 HexNAc 2 | 5.325 | 0.043 | 13.778 | 2.905e+09 | 145 - 157 | 3 | 92.38 | 92.64 | 93.42 |
| 3304.3247 | HKNN(N)KSWMESEF Fuc 1 Hex 5 HexNAc 4 | 5.320 | 0.043 | 14.756 | 1.362e+09 | 145 - 157 | 2 | 91.28 | 91.53 | 91.78 |
| 5388.4107 | VPAQEKN(N)FTTAPAIC(C)HDGKAHFPREGVF Fuc 1 Hex 5 HexNAc 5 Neu5Ac 1 | 5.283 | 0.043 | -5.974 | 9.514e+06 | 1067 - 1095 | 2 | 232.99 | 233.08 | 233.08 |
| 3601.5147 | FHAIHVSGTN(N)GTKRF @sulfate 1 Fuc 1 Hex 3 HexNAc 6 | 5.252 | 0.043 | 10.510 | 1.181e+08 | 64 - 79 | 3 | 85.38 | 85.52 | 85.60 |
| 4418.8192 | IGAEHVN(N)NSYEC(C)DIPIGAGIC(C)ASY Fuc 1 Hex 4 HexNAc 5 | 5.235 | 0.043 | 14.514 | 1.048e+09 | 650 - 674 | 3 | 168.89 | 169.72 | 170.36 |
| 4380.9854 | VPAQEKN(N)FTTAPAIC(C)HDGKAHFPREGVF Hex 4 HexNAc 3 | 5.226 | 0.043 | 11.203 | 2.352e+08 | 1067 - 1095 | 3 | 118.23 | 118.37 | 118.50 |
| 2966.1520 | SSANN(N)C(C)TFEYVSQPF Hex 5 HexNAc 2 | 5.218 | 0.043 | 17.731 | 4.646e+09 | 160 - 175 | 6 | 174.06 | 174.52 | 175.55 |
| 3950.5568 | SNNSIAIPTN(N)F Fuc 2 Hex 6 HexNAc 6 Neu5Ac 1 | 5.149 | 0.043 | 13.629 | 5.149e+08 | 707 - 718 | 1 | 154.48 | 154.73 | 155.23 |
| 4215.7019 | HAIHVSGTN(N)GTKRF @sulfate 1 Fuc 2 Hex 5 HexNAc 6 Neu5Ac 1 | 5.139 | 0.043 | 9.628 | 5.056e+08 | 65 - 79 | 5 | 71.83 | 72.09 | 73.93 |
| 3808.5634 | HAIHVSGTN(N)GTKRF @sulfate 1 Hex 5 HexNAc 4 Neu5Ac 2 | 5.127 | 0.043 | 12.113 | 6.756e+07 | 65 - 79 | 1 | 62.77 | 62.89 | 63.17 |
| 4100.8669 | RVQPTESIVRFPN(N)ITNL Fuc 2 Hex 5 HexNAc 5 | 5.126 | 0.043 | 11.586 | 6.716e+08 | 318 - 335 | 3 | 166.70 | 167.12 | 167.24 |
| 2878.2547 | HVTYVPAQEKN(N)F Fuc 1 Hex 3 HexNAc 4 | 5.115 | 0.043 | 0.000 | 0.000e+00 | 1063 - 1075 | 1 | - | - | - |
| 2876.2490 | HVTYVPAQEKN(N)F Fuc 1 Hex 3 HexNAc 4 | 5.095 | 0.043 | 12.047 | 4.778e+08 | 1063 - 1075 | 2 | 92.22 | 92.29 | 92.46 |
| 4313.8014 | HAIHVSGTN(N)GTKRF Fuc 1 Hex 7 HexNAc 6 Neu5Ac 1 | 5.085 | 0.043 | 11.191 | 4.254e+08 | 65 - 79 | 9 | 60.01 | 60.70 | 61.68 |
| 4371.8369 | HAIHVSGTN(N)GTKRF Fuc 2 Hex 7 HexNAc 7 | 5.082 | 0.043 | 10.645 | 4.066e+07 | 65 - 79 | 2 | 54.15 | 54.55 | 54.59 |
| 3064.3283 | HAIHVSGTN(N)GTKRF Hex 7 HexNAc 2 | 5.047 | 0.044 | 10.235 | 8.566e+07 | 65 - 79 | 2 | 51.67 | 52.69 | 53.03 |
| 3549.4392 | YHKNN(N)KSWMESEF Fuc 1 Hex 3 HexNAc 6 | 5.014 | 0.044 | 11.132 | 9.068e+09 | 144 - 157 | 16 | 93.89 | 94.95 | 96.59 |
| 3674.6286 | IVNN(N)ATNVVIKVC(C)EF Fuc 2 Hex 4 HexNAc 5 | 4.999 | 0.045 | 15.996 | 7.208e+08 | 118 - 133 | 2 | 157.91 | 158.12 | 158.32 |
| 4049.5791 | YHKNN(N)KSWMESEF Fuc 1 Hex 5 HexNAc 4 Neu5Ac 2 | 4.977 | 0.045 | 15.928 | 5.211e+08 | 144 - 157 | 1 | 124.06 | 124.24 | 124.60 |
| 3633.6040 | IVNN(N)ATNVVIKVC(C)EF Fuc 2 Hex 5 HexNAc 4 | 4.975 | 0.045 | 12.999 | 5.506e+08 | 118 - 133 | 2 | 157.81 | 157.99 | 158.38 |
| 4265.6517 | SN(N)NSIAIPTN(N)F Hex 11 HexNAc 5 Neu5Ac 1 | 4.960 | 0.045 | 15.873 | 1.905e+09 | 707 - 718 | 4 | 177.66 | 178.00 | 178.40 |
| 3799.5080 | YHKNN(N)KSWMESEF Fuc 1 Hex 4 HexNAc 5 Neu5Ac 1 | 4.946 | 0.046 | 13.765 | 7.536e+09 | 144 - 157 | 6 | 106.84 | 107.22 | 108.00 |
| 4385.8142 | HAIHVSGTN(N)GTKRF Fuc 2 Hex 6 HexNAc 5 Neu5Ac 2 | 4.905 | 0.047 | 11.019 | 4.020e+08 | 65 - 79 | 10 | 69.50 | 70.59 | 71.57 |
| 3866.5678 | HAIHVSGTN(N)GTKRF @sulfate 1 Fuc 1 Hex 5 HexNAc 5 Neu5Ac 1 | 4.898 | 0.047 | 12.501 | 1.033e+08 | 65 - 79 | 2 | 72.58 | 72.77 | 73.27 |
| 3735.7356 | RVQPTESIVRFPN(N)ITNL Fuc 2 Hex 4 HexNAc 4 | 4.883 | 0.047 | 11.002 | 1.049e+09 | 318 - 335 | 6 | 167.45 | 167.66 | 168.16 |
| 3397.5463 | LIVNN(N)ATNVVIKVC(C)EF Fuc 1 Hex 5 HexNAc 3 | 4.868 | 0.047 | 12.959 | 1.215e+09 | 117 - 133 | 3 | 178.36 | 178.69 | 179.82 |
| 4985.0930 | Q(P)TRAGC(C)LIGAEHVN(N)NSYEC(C)DIPIGAGIC(C)ASY Fuc 1 Hex 4 HexNAc 4 | 4.860 | 0.047 | 14.063 | 2.724e+08 | 643 - 674 | 1 | 184.75 | 185.05 | 185.23 |
| 3089.4379 | LIVNN(N)ATNVVIKVC(C)EF Hex 4 HexNAc 3 | 4.846 | 0.048 | 15.531 | 8.261e+09 | 117 - 133 | 4 | 180.00 | 180.36 | 180.81 |
| 3745.6675 | LIVNN(N)ATNVVIKVC(C)EF Hex 5 HexNAc 4 Neu5Ac 1 | 4.838 | 0.049 | 10.189 | 2.015e+08 | 117 - 133 | 3 | 197.42 | 197.46 | 197.60 |
| 3333.5437 | LIVNN(N)ATNVVIKVC(C)EF Hex 3 HexNAc 5 | 4.797 | 0.050 | 14.469 | 4.900e+08 | 117 - 133 | 3 | 179.53 | 179.98 | 180.25 |
| 5184.2758 | VPAQEKN(N)FTTAPAIC(C)HDGKAHFPREGVF Fuc 3 Hex 5 HexNAc 4 | 4.765 | 0.050 | 11.041 | 1.257e+09 | 1067 - 1095 | 4 | 125.16 | 125.45 | 125.95 |
| 3891.7235 | LIVNN(N)ATNVVIKVC(C)EF Fuc 1 Hex 5 HexNAc 4 Neu5Ac 1 | 4.759 | 0.050 | 14.518 | 8.296e+08 | 117 - 133 | 2 | 194.97 | 195.29 | 195.54 |
| 4399.9880 | RVQPTESIVRFPN(N)ITNLC(C)PF Fuc 1 Hex 4 HexNAc 6 | 4.722 | 0.050 | 15.636 | 1.463e+09 | 318 - 338 | 4 | 190.89 | 191.47 | 192.47 |
| 5038.2321 | VPAQEKN(N)FTTAPAIC(C)HDGKAHFPREGVF Fuc 2 Hex 5 HexNAc 4 | 4.654 | 0.050 | 13.786 | 6.580e+08 | 1067 - 1095 | 4 | 116.94 | 117.14 | 117.44 |

#### HAIHVSGTN(N)GTKRF Fuc 1 Hex 4 HexNAc 5

**Mass:**  3333.4590

(-1.571e-06)

**Charge States:** 3, 4, 5

**Start Position**: 66

**End Position**: 80

| Profile Score | Tandem Score | TDA q-value | Volume | Spectrum Matches |
| --- | --- | --- | --- | --- |
| 13.7162 | 165.9341 | 0.0020 | 4.3557e+10 | 66 |

**Adduct Signal Fractions**

- **Unmodified**: 4.3557e+10

#### HAIHVSGTN(N)GTKRF Fuc 1 Hex 5 HexNAc 5

**Mass:**  3495.5164

(-6.129e-07)

**Charge States:** 3, 4, 5

**Start Position**: 66

**End Position**: 80

| Profile Score | Tandem Score | TDA q-value | Volume | Spectrum Matches |
| --- | --- | --- | --- | --- |
| 16.1288 | 161.3963 | 0.0020 | 9.8841e+09 | 46 |

**Adduct Signal Fractions**

- **Unmodified**: 9.8841e+09

#### HAIHVSGTN(N)GTKRF Fuc 2 Hex 4 HexNAc 4

**Mass:**  3276.4466

(1.251e-06)

**Charge States:** 3, 4, 5

**Start Position**: 66

**End Position**: 80

| Profile Score | Tandem Score | TDA q-value | Volume | Spectrum Matches |
| --- | --- | --- | --- | --- |
| 16.2418 | 160.3417 | 0.0020 | 7.9503e+09 | 23 |

**Adduct Signal Fractions**

- **Unmodified**: 7.9503e+09

#### HAIHVSGTN(N)GTKRF Fuc 1 Hex 3 HexNAc 4

**Mass:**  2968.3315

(4.667e-07)

**Charge States:** 2, 3, 4, 5

**Start Position**: 66

**End Position**: 80

| Profile Score | Tandem Score | TDA q-value | Volume | Spectrum Matches |
| --- | --- | --- | --- | --- |
| 17.7919 | 158.2543 | 0.0020 | 4.9589e+10 | 58 |

**Adduct Signal Fractions**

- **Unmodified**: 4.9589e+10

#### HAIHVSGTN(N)GTKRF @sulfate 1 Fuc 1 Hex 4 HexNAc 5

**Mass:**  3413.4131

(-1.136e-06)

**Charge States:** 3, 4

**Start Position**: 66

**End Position**: 80

| Profile Score | Tandem Score | TDA q-value | Volume | Spectrum Matches |
| --- | --- | --- | --- | --- |
| 16.4585 | 157.3770 | 0.0020 | 1.4459e+10 | 21 |

**Adduct Signal Fractions**

- **Unmodified**: 1.4459e+10

#### HAIHVSGTN(N)GTKRF Fuc 2 Hex 4 HexNAc 5 Neu5Ac 1

**Mass:**  3770.6122

(-8.366e-07)

**Charge States:** 3, 4, 5

**Start Position**: 66

**End Position**: 80

| Profile Score | Tandem Score | TDA q-value | Volume | Spectrum Matches |
| --- | --- | --- | --- | --- |
| 16.7382 | 157.1832 | 0.0020 | 5.9402e+10 | 55 |

**Adduct Signal Fractions**

- **Unmodified**: 5.9402e+10

#### HAIHVSGTN(N)GTKRF @sulfate 1 Fuc 1 Hex 4 HexNAc 5 Neu5Ac 1

**Mass:**  3704.5152

(6.185e-07)

**Charge States:** 3, 4, 5

**Start Position**: 66

**End Position**: 80

| Profile Score | Tandem Score | TDA q-value | Volume | Spectrum Matches |
| --- | --- | --- | --- | --- |
| 17.9662 | 155.5578 | 0.0020 | 3.8110e+10 | 26 |

**Adduct Signal Fractions**

- **Unmodified**: 3.8110e+10

#### HAIHVSGTN(N)GTKRF @sulfate 1 Fuc 1 Hex 3 HexNAc 6

**Mass:**  3454.4472

(5.558e-07)

**Charge States:** 3, 4

**Start Position**: 66

**End Position**: 80

| Profile Score | Tandem Score | TDA q-value | Volume | Spectrum Matches |
| --- | --- | --- | --- | --- |
| 16.5633 | 154.5584 | 0.0020 | 2.6032e+10 | 25 |

**Adduct Signal Fractions**

- **Unmodified**: 2.6032e+10

#### HAIHVSGTN(N)GTKRF Fuc 2 Hex 4 HexNAc 5

**Mass:**  3479.5219

(-1.807e-06)

**Charge States:** 3, 4, 5

**Start Position**: 66

**End Position**: 80

| Profile Score | Tandem Score | TDA q-value | Volume | Spectrum Matches |
| --- | --- | --- | --- | --- |
| 17.1176 | 153.8750 | 0.0020 | 7.3646e+10 | 67 |

**Adduct Signal Fractions**

- **Unmodified**: 7.3646e+10

#### HAIHVSGTN(N)GTKRF Fuc 2 Hex 4 HexNAc 6

**Mass:**  3682.6032

(2.071e-06)

**Charge States:** 3, 4, 5

**Start Position**: 66

**End Position**: 80

| Profile Score | Tandem Score | TDA q-value | Volume | Spectrum Matches |
| --- | --- | --- | --- | --- |
| 6.6664 | 151.1049 | 0.0020 | 3.8614e+09 | 37 |

**Adduct Signal Fractions**

- **Unmodified**: 3.8614e+09

#### HAIHVSGTN(N)GTKRF Fuc 1 Hex 4 HexNAc 5 Neu5Ac 1

**Mass:**  3624.5601

(1.252e-06)

**Charge States:** 3, 4, 5

**Start Position**: 66

**End Position**: 80

| Profile Score | Tandem Score | TDA q-value | Volume | Spectrum Matches |
| --- | --- | --- | --- | --- |
| 14.8677 | 146.8319 | 0.0020 | 3.3734e+10 | 53 |

**Adduct Signal Fractions**

- **Unmodified**: 3.3734e+10

#### HAIHVSGTN(N)GTKRF Fuc 1 Hex 3 HexNAc 5

**Mass:**  3171.4119

(1.679e-06)

**Charge States:** 2, 3, 4, 5

**Start Position**: 66

**End Position**: 80

| Profile Score | Tandem Score | TDA q-value | Volume | Spectrum Matches |
| --- | --- | --- | --- | --- |
| 14.7694 | 146.7734 | 0.0020 | 8.7757e+10 | 68 |

**Adduct Signal Fractions**

- **Unmodified**: 8.7757e+10

#### HAIHVSGTN(N)GTKRF Fuc 1 Hex 4 HexNAc 4

**Mass:**  3130.3765

(-1.945e-06)

**Charge States:** 3, 4, 5

**Start Position**: 66

**End Position**: 80

| Profile Score | Tandem Score | TDA q-value | Volume | Spectrum Matches |
| --- | --- | --- | --- | --- |
| 15.3642 | 146.5561 | 0.0020 | 9.3422e+09 | 35 |

**Adduct Signal Fractions**

- **Unmodified**: 9.3422e+09

#### HAIHVSGTN(N)GTKRF Fuc 2 Hex 5 HexNAc 6

**Mass:**  3844.6494

(-5.068e-07)

**Charge States:** 3, 4, 5

**Start Position**: 66

**End Position**: 80

| Profile Score | Tandem Score | TDA q-value | Volume | Spectrum Matches |
| --- | --- | --- | --- | --- |
| -5.9681 | 144.4057 | 0.0020 | 4.4648e+09 | 24 |

**Adduct Signal Fractions**

- **Unmodified**: 4.4648e+09

#### HAIHVSGTN(N)GTKRF @sulfate 1 Fuc 1 Hex 3 HexNAc 5

**Mass:**  3251.3682

(1.413e-06)

**Charge States:** 3, 4

**Start Position**: 66

**End Position**: 80

| Profile Score | Tandem Score | TDA q-value | Volume | Spectrum Matches |
| --- | --- | --- | --- | --- |
| 18.2663 | 143.8109 | 0.0020 | 2.5352e+10 | 26 |

**Adduct Signal Fractions**

- **Unmodified**: 2.5352e+10

#### HAIHVSGTN(N)GTKRF Fuc 2 Hex 5 HexNAc 4

**Mass:**  3438.4905

(-8.244e-07)

**Charge States:** 3, 4, 5

**Start Position**: 66

**End Position**: 80

| Profile Score | Tandem Score | TDA q-value | Volume | Spectrum Matches |
| --- | --- | --- | --- | --- |
| 12.7847 | 141.1517 | 0.0020 | 3.0920e+09 | 21 |

**Adduct Signal Fractions**

- **Unmodified**: 3.0920e+09

#### VDLPIGIN(N)ITRF Hex 9 HexNAc 2

**Mass:**  3221.4131

(6.838e-07)

**Charge States:** 3

**Start Position**: 227

**End Position**: 239

| Profile Score | Tandem Score | TDA q-value | Volume | Spectrum Matches |
| --- | --- | --- | --- | --- |
| 17.6127 | 140.5832 | 0.0020 | 8.5296e+08 | 2 |

**Adduct Signal Fractions**

- **Unmodified**: 8.5296e+08

#### HAIHVSGTN(N)GTKRF Hex 5 HexNAc 2

**Mass:**  2740.2134

(-2.265e-06)

**Charge States:** 3, 4

**Start Position**: 66

**End Position**: 80

| Profile Score | Tandem Score | TDA q-value | Volume | Spectrum Matches |
| --- | --- | --- | --- | --- |
| 16.9080 | 140.4511 | 0.0020 | 6.5111e+08 | 3 |

**Adduct Signal Fractions**

- **Unmodified**: 6.5111e+08

#### HAIHVSGTN(N)GTKRF Fuc 1 Hex 4 HexNAc 6

**Mass:**  3536.5370

(2.548e-06)

**Charge States:** 3, 4, 5

**Start Position**: 66

**End Position**: 80

| Profile Score | Tandem Score | TDA q-value | Volume | Spectrum Matches |
| --- | --- | --- | --- | --- |
| 13.7203 | 139.5777 | 0.0020 | 6.9513e+09 | 37 |

**Adduct Signal Fractions**

- **Unmodified**: 6.9513e+09

#### HAIHVSGTN(N)GTKRF Fuc 1 Hex 5 HexNAc 4 Neu5Ac 2

**Mass:**  3874.6213

(-9.495e-07)

**Charge States:** 3, 4, 5

**Start Position**: 66

**End Position**: 80

| Profile Score | Tandem Score | TDA q-value | Volume | Spectrum Matches |
| --- | --- | --- | --- | --- |
| 14.5101 | 139.5287 | 0.0020 | 6.7035e+09 | 16 |

**Adduct Signal Fractions**

- **Unmodified**: 6.7035e+09

#### HAIHVSGTN(N)GTKRF @sulfate 1 Fuc 1 Hex 4 HexNAc 4

**Mass:**  3210.3359

(-1.283e-06)

**Charge States:** 3, 4

**Start Position**: 66

**End Position**: 80

| Profile Score | Tandem Score | TDA q-value | Volume | Spectrum Matches |
| --- | --- | --- | --- | --- |
| 14.3125 | 139.2517 | 0.0020 | 1.3013e+09 | 4 |

**Adduct Signal Fractions**

- **Unmodified**: 1.3013e+09

#### HAIHVSGTN(N)GTKRF Fuc 1 Hex 4 HexNAc 7

**Mass:**  3739.6148

(-2.079e-06)

**Charge States:** 3, 4, 5

**Start Position**: 66

**End Position**: 80

| Profile Score | Tandem Score | TDA q-value | Volume | Spectrum Matches |
| --- | --- | --- | --- | --- |
| 14.4407 | 137.9707 | 0.0020 | 7.6248e+09 | 36 |

**Adduct Signal Fractions**

- **Unmodified**: 7.6248e+09

#### HAIHVSGTN(N)GTKRF Fuc 1 Hex 3 HexNAc 6

**Mass:**  3374.4881

(2.232e-07)

**Charge States:** 3, 4, 5

**Start Position**: 66

**End Position**: 80

| Profile Score | Tandem Score | TDA q-value | Volume | Spectrum Matches |
| --- | --- | --- | --- | --- |
| 13.7954 | 137.5294 | 0.0020 | 2.7516e+10 | 57 |

**Adduct Signal Fractions**

- **Unmodified**: 2.7516e+10

#### HAIHVSGTN(N)GTKRF @sulfate 1 Fuc 1 Hex 5 HexNAc 4

**Mass:**  3372.3978

(1.741e-06)

**Charge States:** 3, 4

**Start Position**: 66

**End Position**: 80

| Profile Score | Tandem Score | TDA q-value | Volume | Spectrum Matches |
| --- | --- | --- | --- | --- |
| 14.5939 | 137.3578 | 0.0020 | 4.4333e+08 | 5 |

**Adduct Signal Fractions**

- **Unmodified**: 4.4333e+08

#### GEVFN(N)ATRF Fuc 1 Hex 3 HexNAc 4

**Mass:**  2484.0438

(9.303e-07)

**Charge States:** 2, 3

**Start Position**: 339

**End Position**: 348

| Profile Score | Tandem Score | TDA q-value | Volume | Spectrum Matches |
| --- | --- | --- | --- | --- |
| 17.3931 | 135.9559 | 0.0079 | 5.1463e+09 | 4 |

**Adduct Signal Fractions**

- **Unmodified**: 5.1463e+09

#### HAIHVSGTN(N)GTKRF @sulfate 1 Fuc 2 Hex 4 HexNAc 5

**Mass:**  3559.4802

(1.664e-06)

**Charge States:** 3, 4

**Start Position**: 66

**End Position**: 80

| Profile Score | Tandem Score | TDA q-value | Volume | Spectrum Matches |
| --- | --- | --- | --- | --- |
| 14.2291 | 135.9422 | 0.0020 | 7.0970e+09 | 23 |

**Adduct Signal Fractions**

- **Unmodified**: 7.0970e+09

#### HAIHVSGTN(N)GTKRF @sulfate 1 Hex 6 HexNAc 2

**Mass:**  2982.2454

(5.913e-06)

**Charge States:** 3, 4

**Start Position**: 66

**End Position**: 80

| Profile Score | Tandem Score | TDA q-value | Volume | Spectrum Matches |
| --- | --- | --- | --- | --- |
| 15.6603 | 133.7527 | 0.0020 | 5.7060e+08 | 3 |

**Adduct Signal Fractions**

- **Unmodified**: 5.7060e+08

#### HAIHVSGTN(N)GTKRF Fuc 2 Hex 4 HexNAc 7

**Mass:**  3885.6818

(1.047e-06)

**Charge States:** 3, 4, 5

**Start Position**: 66

**End Position**: 80

| Profile Score | Tandem Score | TDA q-value | Volume | Spectrum Matches |
| --- | --- | --- | --- | --- |
| 14.5547 | 132.5647 | 0.0020 | 6.3767e+09 | 38 |

**Adduct Signal Fractions**

- **Unmodified**: 6.3767e+09

#### HAIHVSGTN(N)GTKRF @sulfate 1 Fuc 1 Hex 3 HexNAc 4

**Mass:**  3048.2908

(1.421e-06)

**Charge States:** 3, 4

**Start Position**: 66

**End Position**: 80

| Profile Score | Tandem Score | TDA q-value | Volume | Spectrum Matches |
| --- | --- | --- | --- | --- |
| 15.0309 | 130.0437 | 0.0020 | 1.4627e+09 | 8 |

**Adduct Signal Fractions**

- **Unmodified**: 1.4627e+09

#### HAIHVSGTN(N)GTKRF @sulfate 1 Hex 6 HexNAc 3

**Mass:**  3185.3245

(5.425e-06)

**Charge States:** 3, 4

**Start Position**: 66

**End Position**: 80

| Profile Score | Tandem Score | TDA q-value | Volume | Spectrum Matches |
| --- | --- | --- | --- | --- |
| 16.2692 | 129.7540 | 0.0020 | 4.3221e+08 | 2 |

**Adduct Signal Fractions**

- **Unmodified**: 4.3221e+08

#### HAIHVSGTN(N)GTKRF Fuc 1 Hex 4 HexNAc 4 Neu5Ac 1

**Mass:**  3421.4809

(1.704e-06)

**Charge States:** 3, 4, 5

**Start Position**: 66

**End Position**: 80

| Profile Score | Tandem Score | TDA q-value | Volume | Spectrum Matches |
| --- | --- | --- | --- | --- |
| 13.2605 | 129.5865 | 0.0020 | 5.9155e+09 | 27 |

**Adduct Signal Fractions**

- **Unmodified**: 5.9155e+09

#### HAIHVSGTN(N)GTKRF Fuc 1 Hex 6 HexNAc 5 Neu5Ac 1

**Mass:**  3948.6610

(-4.685e-07)

**Charge States:** 3, 4, 5

**Start Position**: 66

**End Position**: 80

| Profile Score | Tandem Score | TDA q-value | Volume | Spectrum Matches |
| --- | --- | --- | --- | --- |
| 12.9520 | 129.3344 | 0.0020 | 2.3513e+09 | 23 |

**Adduct Signal Fractions**

- **Unmodified**: 2.3513e+09

#### HAIHVSGTN(N)GTKRF Fuc 1 Hex 4 HexNAc 3

**Mass:**  2927.2963

(-1.847e-06)

**Charge States:** 3, 4

**Start Position**: 66

**End Position**: 80

| Profile Score | Tandem Score | TDA q-value | Volume | Spectrum Matches |
| --- | --- | --- | --- | --- |
| 15.1184 | 128.1564 | 0.0020 | 2.3921e+09 | 11 |

**Adduct Signal Fractions**

- **Unmodified**: 2.3921e+09

#### GEVFN(N)ATRF Fuc 1 Hex 5 HexNAc 4 Neu5Ac 1

**Mass:**  3099.2465

(9.961e-07)

**Charge States:** 2, 3

**Start Position**: 339

**End Position**: 348

| Profile Score | Tandem Score | TDA q-value | Volume | Spectrum Matches |
| --- | --- | --- | --- | --- |
| 17.9630 | 127.3482 | 0.0079 | 3.0254e+09 | 8 |

**Adduct Signal Fractions**

- **Unmodified**: 3.0254e+09

#### VPAQEKN(N)F Hex 5 HexNAc 2

**Mass:**  2147.9011

(-2.002e-06)

**Charge States:** 2, 3, 4

**Start Position**: 1068

**End Position**: 1076

| Profile Score | Tandem Score | TDA q-value | Volume | Spectrum Matches |
| --- | --- | --- | --- | --- |
| 19.9164 | 126.4574 | 0.0079 | 6.0333e+10 | 23 |

**Adduct Signal Fractions**

- **Unmodified**: 6.0333e+10

#### VPAQEKN(N)F Hex 6 HexNAc 2

**Mass:**  2309.9510

(4.369e-07)

**Charge States:** 2, 3

**Start Position**: 1068

**End Position**: 1076

| Profile Score | Tandem Score | TDA q-value | Volume | Spectrum Matches |
| --- | --- | --- | --- | --- |
| 16.8036 | 126.4262 | 0.0079 | 1.8623e+09 | 10 |

**Adduct Signal Fractions**

- **Unmodified**: 1.8623e+09

#### HAIHVSGTN(N)GTKRF Fuc 1 Hex 5 HexNAc 6 Neu5Ac 1

**Mass:**  3989.6889

(1.825e-06)

**Charge States:** 3, 4, 5

**Start Position**: 66

**End Position**: 80

| Profile Score | Tandem Score | TDA q-value | Volume | Spectrum Matches |
| --- | --- | --- | --- | --- |
| 12.5072 | 126.3529 | 0.0020 | 6.5940e+09 | 21 |

**Adduct Signal Fractions**

- **Unmodified**: 6.5940e+09

#### HAIHVSGTN(N)GTKRF @sulfate 1 Fuc 2 Hex 4 HexNAc 7

**Mass:**  3965.6406

(1.353e-06)

**Charge States:** 3, 4

**Start Position**: 66

**End Position**: 80

| Profile Score | Tandem Score | TDA q-value | Volume | Spectrum Matches |
| --- | --- | --- | --- | --- |
| 14.9381 | 125.1115 | 0.0020 | 1.0772e+09 | 6 |

**Adduct Signal Fractions**

- **Unmodified**: 1.0772e+09

#### VDLPIGIN(N)ITRF Hex 8 HexNAc 2

**Mass:**  3059.3567

(-4.704e-09)

**Charge States:** 2, 3

**Start Position**: 227

**End Position**: 239

| Profile Score | Tandem Score | TDA q-value | Volume | Spectrum Matches |
| --- | --- | --- | --- | --- |
| 16.3194 | 123.5080 | 0.0020 | 1.6232e+09 | 3 |

**Adduct Signal Fractions**

- **Unmodified**: 1.6232e+09

#### HAIHVSGTN(N)GTKRF Fuc 2 Hex 5 HexNAc 4 Neu5Ac 1

**Mass:**  3729.5908

(1.191e-06)

**Charge States:** 3, 4, 5

**Start Position**: 66

**End Position**: 80

| Profile Score | Tandem Score | TDA q-value | Volume | Spectrum Matches |
| --- | --- | --- | --- | --- |
| 11.6047 | 121.1930 | 0.0020 | 1.3636e+09 | 12 |

**Adduct Signal Fractions**

- **Unmodified**: 1.3636e+09

#### HAIHVSGTN(N)GTKRF Fuc 1 Hex 5 HexNAc 6

**Mass:**  3698.5991

(2.585e-06)

**Charge States:** 3, 4, 5

**Start Position**: 66

**End Position**: 80

| Profile Score | Tandem Score | TDA q-value | Volume | Spectrum Matches |
| --- | --- | --- | --- | --- |
| 14.0593 | 119.5359 | 0.0020 | 4.1781e+09 | 26 |

**Adduct Signal Fractions**

- **Unmodified**: 4.1781e+09

#### HAIHVSGTN(N)GTKRF Fuc 2 Hex 5 HexNAc 6 Neu5Ac 1

**Mass:**  4135.7559

(2.198e-06)

**Charge States:** 3, 4, 5

**Start Position**: 66

**End Position**: 80

| Profile Score | Tandem Score | TDA q-value | Volume | Spectrum Matches |
| --- | --- | --- | --- | --- |
| 12.9761 | 119.3610 | 0.0020 | 4.2774e+09 | 25 |

**Adduct Signal Fractions**

- **Unmodified**: 4.2774e+09

#### HAIHVSGTN(N)GTKRF Fuc 1 Hex 5 HexNAc 4

**Mass:**  3292.4413

(2.225e-06)

**Charge States:** 3, 4, 5

**Start Position**: 66

**End Position**: 80

| Profile Score | Tandem Score | TDA q-value | Volume | Spectrum Matches |
| --- | --- | --- | --- | --- |
| 14.0556 | 119.1673 | 0.0020 | 2.4919e+09 | 33 |

**Adduct Signal Fractions**

- **Unmodified**: 2.4919e+09

#### HAIHVSGTN(N)GTKRF Fuc 2 Hex 5 HexNAc 5

**Mass:**  3641.5702

(-1.158e-07)

**Charge States:** 3, 4, 5

**Start Position**: 66

**End Position**: 80

| Profile Score | Tandem Score | TDA q-value | Volume | Spectrum Matches |
| --- | --- | --- | --- | --- |
| 14.5573 | 117.1430 | 0.0020 | 4.2393e+09 | 33 |

**Adduct Signal Fractions**

- **Unmodified**: 4.2393e+09

#### HAIHVSGTN(N)GTKRF @sulfate 1 Fuc 1 Hex 3 HexNAc 7

**Mass:**  3657.5250

(7.893e-07)

**Charge States:** 3, 4

**Start Position**: 66

**End Position**: 80

| Profile Score | Tandem Score | TDA q-value | Volume | Spectrum Matches |
| --- | --- | --- | --- | --- |
| 12.1942 | 116.2292 | 0.0020 | 1.8563e+09 | 14 |

**Adduct Signal Fractions**

- **Unmodified**: 1.8563e+09

#### HAIHVSGTN(N)GTKRF Fuc 1 Hex 3 HexNAc 7

**Mass:**  3577.5690

(-2.568e-07)

**Charge States:** 3, 4, 5

**Start Position**: 66

**End Position**: 80

| Profile Score | Tandem Score | TDA q-value | Volume | Spectrum Matches |
| --- | --- | --- | --- | --- |
| 13.3325 | 115.9852 | 0.0020 | 9.5565e+09 | 45 |

**Adduct Signal Fractions**

- **Unmodified**: 9.5565e+09

#### HAIHVSGTN(N)GTKRF Fuc 1 Hex 6 HexNAc 6

**Mass:**  3860.6477

(2.089e-06)

**Charge States:** 3, 4, 5

**Start Position**: 66

**End Position**: 80

| Profile Score | Tandem Score | TDA q-value | Volume | Spectrum Matches |
| --- | --- | --- | --- | --- |
| 13.5648 | 115.4459 | 0.0020 | 8.9013e+08 | 6 |

**Adduct Signal Fractions**

- **Unmodified**: 8.9013e+08

#### HAIHVSGTN(N)GTKRF Fuc 1 Hex 5 HexNAc 5 Neu5Ac 1

**Mass:**  3786.6108

(-2.342e-07)

**Charge States:** 3, 4, 5

**Start Position**: 66

**End Position**: 80

| Profile Score | Tandem Score | TDA q-value | Volume | Spectrum Matches |
| --- | --- | --- | --- | --- |
| 11.5912 | 115.3926 | 0.0020 | 5.5868e+09 | 30 |

**Adduct Signal Fractions**

- **Unmodified**: 5.5868e+09

#### VPAQEKN(N)F Hex 5 HexNAc 3

**Mass:**  2350.9777

(-4.265e-07)

**Charge States:** 2, 3

**Start Position**: 1068

**End Position**: 1076

| Profile Score | Tandem Score | TDA q-value | Volume | Spectrum Matches |
| --- | --- | --- | --- | --- |
| 17.0588 | 114.7066 | 0.0079 | 5.2007e+09 | 15 |

**Adduct Signal Fractions**

- **Unmodified**: 5.2007e+09

#### VPAQEKN(N)F Hex 6 HexNAc 3

**Mass:**  2513.0309

(-1.061e-06)

**Charge States:** 2, 3

**Start Position**: 1068

**End Position**: 1076

| Profile Score | Tandem Score | TDA q-value | Volume | Spectrum Matches |
| --- | --- | --- | --- | --- |
| 17.5723 | 114.6996 | 0.0079 | 2.6837e+09 | 8 |

**Adduct Signal Fractions**

- **Unmodified**: 2.6837e+09

#### VPAQEKN(N)F Fuc 1 Hex 4 HexNAc 3

**Mass:**  2334.9835

(-2.636e-07)

**Charge States:** 2, 3

**Start Position**: 1068

**End Position**: 1076

| Profile Score | Tandem Score | TDA q-value | Volume | Spectrum Matches |
| --- | --- | --- | --- | --- |
| 13.6439 | 114.5321 | 0.0079 | 3.5560e+09 | 13 |

**Adduct Signal Fractions**

- **Unmodified**: 3.5560e+09

#### HAIHVSGTN(N)GTKRF Fuc 2 Hex 4 HexNAc 7 Neu5Ac 1

**Mass:**  4176.7673

(-3.340e-07)

**Charge States:** 3, 4, 5

**Start Position**: 66

**End Position**: 80

| Profile Score | Tandem Score | TDA q-value | Volume | Spectrum Matches |
| --- | --- | --- | --- | --- |
| 15.0828 | 113.5492 | 0.0020 | 4.9908e+09 | 19 |

**Adduct Signal Fractions**

- **Unmodified**: 4.9908e+09

#### GEVFN(N)ATRF Fuc 1 Hex 5 HexNAc 4

**Mass:**  2808.1492

(2.236e-07)

**Charge States:** 2, 3

**Start Position**: 339

**End Position**: 348

| Profile Score | Tandem Score | TDA q-value | Volume | Spectrum Matches |
| --- | --- | --- | --- | --- |
| 16.4952 | 113.0899 | 0.0079 | 1.1691e+10 | 10 |

**Adduct Signal Fractions**

- **Unmodified**: 1.1691e+10

#### GEVFN(N)ATRF Fuc 2 Hex 5 HexNAc 4

**Mass:**  2954.2064

(1.786e-06)

**Charge States:** 2, 3

**Start Position**: 339

**End Position**: 348

| Profile Score | Tandem Score | TDA q-value | Volume | Spectrum Matches |
| --- | --- | --- | --- | --- |
| 14.8397 | 106.9433 | 0.0079 | 6.5425e+08 | 3 |

**Adduct Signal Fractions**

- **Unmodified**: 6.5425e+08

#### VSN(N)GTHW Fuc 1 Hex 4 HexNAc 4 Neu5Ac 1

**Mass:**  2697.0424

(-3.593e-07)

**Charge States:** 2, 3

**Start Position**: 1096

**End Position**: 1103

| Profile Score | Tandem Score | TDA q-value | Volume | Spectrum Matches |
| --- | --- | --- | --- | --- |
| 15.1813 | 106.5079 | 0.0079 | 3.2096e+09 | 11 |

**Adduct Signal Fractions**

- **Unmodified**: 3.2096e+09

#### GEVFN(N)ATRF Fuc 1 Hex 4 HexNAc 4

**Mass:**  2646.0960

(2.441e-07)

**Charge States:** 2, 3

**Start Position**: 339

**End Position**: 348

| Profile Score | Tandem Score | TDA q-value | Volume | Spectrum Matches |
| --- | --- | --- | --- | --- |
| 15.9849 | 105.5065 | 0.0079 | 3.8475e+09 | 10 |

**Adduct Signal Fractions**

- **Unmodified**: 3.8475e+09

#### GEVFN(N)ATRF Fuc 1 Hex 4 HexNAc 5

**Mass:**  2849.1804

(1.792e-06)

**Charge States:** 2, 3

**Start Position**: 339

**End Position**: 348

| Profile Score | Tandem Score | TDA q-value | Volume | Spectrum Matches |
| --- | --- | --- | --- | --- |
| 14.9756 | 104.6205 | 0.0079 | 7.8860e+09 | 11 |

**Adduct Signal Fractions**

- **Unmodified**: 7.8860e+09

#### VPAQEKN(N)F Fuc 1 Hex 5 HexNAc 4

**Mass:**  2700.1166

(2.268e-07)

**Charge States:** 2, 3, 4

**Start Position**: 1068

**End Position**: 1076

| Profile Score | Tandem Score | TDA q-value | Volume | Spectrum Matches |
| --- | --- | --- | --- | --- |
| 17.2466 | 103.6499 | 0.0079 | 4.3617e+09 | 13 |

**Adduct Signal Fractions**

- **Unmodified**: 4.3617e+09

#### VPAQEKN(N)F Hex 4 HexNAc 3

**Mass:**  2188.9241

(1.470e-06)

**Charge States:** 2, 3

**Start Position**: 1068

**End Position**: 1076

| Profile Score | Tandem Score | TDA q-value | Volume | Spectrum Matches |
| --- | --- | --- | --- | --- |
| 15.2357 | 103.5411 | 0.0079 | 2.9549e+09 | 9 |

**Adduct Signal Fractions**

- **Unmodified**: 2.9549e+09

#### VSN(N)GTHW Fuc 1 Hex 4 HexNAc 4

**Mass:**  2405.9489

(8.096e-07)

**Charge States:** 2, 3

**Start Position**: 1096

**End Position**: 1103

| Profile Score | Tandem Score | TDA q-value | Volume | Spectrum Matches |
| --- | --- | --- | --- | --- |
| 12.9436 | 103.4855 | 0.0079 | 7.9522e+08 | 10 |

**Adduct Signal Fractions**

- **Unmodified**: 7.9522e+08

#### HAIHVSGTN(N)GTKRF Fuc 1 Hex 3 HexNAc 8

**Mass:**  3780.6554

(2.256e-06)

**Charge States:** 3, 4, 5

**Start Position**: 66

**End Position**: 80

| Profile Score | Tandem Score | TDA q-value | Volume | Spectrum Matches |
| --- | --- | --- | --- | --- |
| 13.9897 | 102.3336 | 0.0020 | 3.8461e+09 | 22 |

**Adduct Signal Fractions**

- **Unmodified**: 3.8461e+09

#### VPAQEKN(N)F Fuc 1 Hex 5 HexNAc 4 Neu5Ac 1

**Mass:**  2991.2095

(-7.271e-07)

**Charge States:** 2, 3

**Start Position**: 1068

**End Position**: 1076

| Profile Score | Tandem Score | TDA q-value | Volume | Spectrum Matches |
| --- | --- | --- | --- | --- |
| 18.2813 | 101.5798 | 0.0079 | 3.1676e+09 | 9 |

**Adduct Signal Fractions**

- **Unmodified**: 3.1676e+09

#### GEVFN(N)ATRF Fuc 1 Hex 3 HexNAc 5

**Mass:**  2687.1231

(4.205e-07)

**Charge States:** 2, 3

**Start Position**: 339

**End Position**: 348

| Profile Score | Tandem Score | TDA q-value | Volume | Spectrum Matches |
| --- | --- | --- | --- | --- |
| 16.7834 | 100.5395 | 0.0079 | 5.2596e+09 | 7 |

**Adduct Signal Fractions**

- **Unmodified**: 5.2596e+09

#### VSN(N)GTHW Fuc 1 Hex 3 HexNAc 5

**Mass:**  2446.9766

(7.689e-07)

**Charge States:** 2, 3

**Start Position**: 1096

**End Position**: 1103

| Profile Score | Tandem Score | TDA q-value | Volume | Spectrum Matches |
| --- | --- | --- | --- | --- |
| 15.5738 | 97.5904 | 0.0079 | 2.9171e+08 | 4 |

**Adduct Signal Fractions**

- **Unmodified**: 2.9171e+08

#### VSN(N)GTHW Hex 3 HexNAc 5

**Mass:**  2300.9174

(1.277e-06)

**Charge States:** 2, 3

**Start Position**: 1096

**End Position**: 1103

| Profile Score | Tandem Score | TDA q-value | Volume | Spectrum Matches |
| --- | --- | --- | --- | --- |
| 16.4363 | 97.2917 | 0.0079 | 4.2826e+08 | 4 |

**Adduct Signal Fractions**

- **Unmodified**: 4.2826e+08

#### VSN(N)GTHW Hex 7 HexNAc 3 Neu5Ac 1

**Mass:**  2835.0611

(3.546e-04)

**Charge States:** 3

**Start Position**: 1096

**End Position**: 1103

| Profile Score | Tandem Score | TDA q-value | Volume | Spectrum Matches |
| --- | --- | --- | --- | --- |
| 9.6263 | 95.1208 | 0.0079 | 5.7908e+06 | 1 |

**Adduct Signal Fractions**

- **Unmodified**: 5.7908e+06

#### VSN(N)GTHW Fuc 1 Hex 5 HexNAc 4 Neu5Ac 1

**Mass:**  2859.0941

(-7.839e-07)

**Charge States:** 2, 3

**Start Position**: 1096

**End Position**: 1103

| Profile Score | Tandem Score | TDA q-value | Volume | Spectrum Matches |
| --- | --- | --- | --- | --- |
| 17.7388 | 95.1189 | 0.0079 | 2.9239e+09 | 10 |

**Adduct Signal Fractions**

- **Unmodified**: 2.9239e+09

#### VSN(N)GTHW Fuc 1 Hex 4 HexNAc 5 Neu5Ac 1

**Mass:**  2900.1233

(3.295e-07)

**Charge States:** 2, 3

**Start Position**: 1096

**End Position**: 1103

| Profile Score | Tandem Score | TDA q-value | Volume | Spectrum Matches |
| --- | --- | --- | --- | --- |
| 18.4388 | 94.9756 | 0.0079 | 4.5559e+08 | 4 |

**Adduct Signal Fractions**

- **Unmodified**: 4.5559e+08

#### VPAQEKN(N)F Fuc 1 Hex 5 HexNAc 3 Neu5Ac 1

**Mass:**  2788.1349

(6.499e-07)

**Charge States:** 2, 3

**Start Position**: 1068

**End Position**: 1076

| Profile Score | Tandem Score | TDA q-value | Volume | Spectrum Matches |
| --- | --- | --- | --- | --- |
| 17.1020 | 94.9227 | 0.0079 | 1.8291e+09 | 6 |

**Adduct Signal Fractions**

- **Unmodified**: 1.8291e+09

#### VPAQEKN(N)F Fuc 1 Hex 6 HexNAc 5

**Mass:**  3065.2477

(-1.270e-07)

**Charge States:** 2, 3, 4

**Start Position**: 1068

**End Position**: 1076

| Profile Score | Tandem Score | TDA q-value | Volume | Spectrum Matches |
| --- | --- | --- | --- | --- |
| 17.0287 | 94.9170 | 0.0079 | 1.9399e+09 | 10 |

**Adduct Signal Fractions**

- **Unmodified**: 1.9399e+09

#### VPAQEKN(N)F Fuc 1 Hex 5 HexNAc 3

**Mass:**  2497.0343

(-8.589e-07)

**Charge States:** 2, 3

**Start Position**: 1068

**End Position**: 1076

| Profile Score | Tandem Score | TDA q-value | Volume | Spectrum Matches |
| --- | --- | --- | --- | --- |
| 18.0725 | 94.7533 | 0.0079 | 2.4032e+09 | 12 |

**Adduct Signal Fractions**

- **Unmodified**: 2.4032e+09

#### VPAQEKN(N)F Fuc 1 Hex 3 HexNAc 5

**Mass:**  2579.0905

(1.765e-07)

**Charge States:** 2, 3, 4

**Start Position**: 1068

**End Position**: 1076

| Profile Score | Tandem Score | TDA q-value | Volume | Spectrum Matches |
| --- | --- | --- | --- | --- |
| 15.9896 | 94.6317 | 0.0079 | 1.3322e+10 | 32 |

**Adduct Signal Fractions**

- **Unmodified**: 1.3322e+10

#### VPAQEKN(N)F Hex 3 HexNAc 3

**Mass:**  2026.8729

(-9.255e-07)

**Charge States:** 2, 3

**Start Position**: 1068

**End Position**: 1076

| Profile Score | Tandem Score | TDA q-value | Volume | Spectrum Matches |
| --- | --- | --- | --- | --- |
| 17.0566 | 94.4630 | 0.0079 | 1.1347e+09 | 9 |

**Adduct Signal Fractions**

- **Unmodified**: 1.1347e+09

#### VSN(N)GTHW Fuc 1 Hex 5 HexNAc 3 Neu5Ac 1

**Mass:**  2656.0190

(9.159e-07)

**Charge States:** 2, 3

**Start Position**: 1096

**End Position**: 1103

| Profile Score | Tandem Score | TDA q-value | Volume | Spectrum Matches |
| --- | --- | --- | --- | --- |
| 17.8636 | 94.4114 | 0.0079 | 1.0123e+09 | 8 |

**Adduct Signal Fractions**

- **Unmodified**: 1.0123e+09

#### VSN(N)GTHW Hex 5 HexNAc 3 Neu5Ac 1

**Mass:**  2509.9596

(2.401e-07)

**Charge States:** 2, 3

**Start Position**: 1096

**End Position**: 1103

| Profile Score | Tandem Score | TDA q-value | Volume | Spectrum Matches |
| --- | --- | --- | --- | --- |
| 16.2434 | 91.5333 | 0.0079 | 1.5712e+09 | 7 |

**Adduct Signal Fractions**

- **Unmodified**: 1.5712e+09

#### VSN(N)GTHW Fuc 1 Hex 3 HexNAc 6

**Mass:**  2650.0565

(1.155e-06)

**Charge States:** 2, 3

**Start Position**: 1096

**End Position**: 1103

| Profile Score | Tandem Score | TDA q-value | Volume | Spectrum Matches |
| --- | --- | --- | --- | --- |
| 15.7501 | 91.0473 | 0.0079 | 6.9344e+08 | 6 |

**Adduct Signal Fractions**

- **Unmodified**: 6.9344e+08

#### VPAQEKN(N)F Fuc 1 Hex 3 HexNAc 4

**Mass:**  2376.0109

(5.650e-07)

**Charge States:** 2, 3, 4

**Start Position**: 1068

**End Position**: 1076

| Profile Score | Tandem Score | TDA q-value | Volume | Spectrum Matches |
| --- | --- | --- | --- | --- |
| 16.9978 | 90.9244 | 0.0079 | 1.4513e+10 | 18 |

**Adduct Signal Fractions**

- **Unmodified**: 1.4513e+10

#### VSN(N)GTHWF Fuc 1 Hex 5 HexNAc 4 Neu5Ac 1

**Mass:**  3006.1663

(4.219e-07)

**Charge States:** 2, 3

**Start Position**: 1096

**End Position**: 1104

| Profile Score | Tandem Score | TDA q-value | Volume | Spectrum Matches |
| --- | --- | --- | --- | --- |
| 13.6057 | 90.7408 | 0.0079 | 5.3820e+09 | 10 |

**Adduct Signal Fractions**

- **Unmodified**: 5.3820e+09

#### VPAQEKN(N)F Fuc 1 Hex 4 HexNAc 4 Neu5Ac 1

**Mass:**  2829.1587

(-2.677e-08)

**Charge States:** 2, 3

**Start Position**: 1068

**End Position**: 1076

| Profile Score | Tandem Score | TDA q-value | Volume | Spectrum Matches |
| --- | --- | --- | --- | --- |
| 15.9820 | 88.6782 | 0.0079 | 3.1580e+09 | 6 |

**Adduct Signal Fractions**

- **Unmodified**: 3.1580e+09

#### VPAQEKN(N)F Hex 5 HexNAc 3 Neu5Ac 1

**Mass:**  2642.0706

(6.501e-07)

**Charge States:** 2, 3

**Start Position**: 1068

**End Position**: 1076

| Profile Score | Tandem Score | TDA q-value | Volume | Spectrum Matches |
| --- | --- | --- | --- | --- |
| 17.0216 | 88.4120 | 0.0079 | 1.2727e+09 | 4 |

**Adduct Signal Fractions**

- **Unmodified**: 1.2727e+09

#### VSN(N)GTHWF Hex 5 HexNAc 3

**Mass:**  2365.9316

(-1.564e-07)

**Charge States:** 2, 3

**Start Position**: 1096

**End Position**: 1104

| Profile Score | Tandem Score | TDA q-value | Volume | Spectrum Matches |
| --- | --- | --- | --- | --- |
| 13.5188 | 88.1556 | 0.0079 | 3.5952e+09 | 9 |

**Adduct Signal Fractions**

- **Unmodified**: 3.5952e+09

#### VPAQEKN(N)F Fuc 1 Hex 4 HexNAc 5 Neu5Ac 1

**Mass:**  3032.2390

(6.424e-07)

**Charge States:** 2, 3, 4

**Start Position**: 1068

**End Position**: 1076

| Profile Score | Tandem Score | TDA q-value | Volume | Spectrum Matches |
| --- | --- | --- | --- | --- |
| 15.8164 | 87.6996 | 0.0079 | 1.0273e+09 | 6 |

**Adduct Signal Fractions**

- **Unmodified**: 1.0273e+09

#### VSN(N)GTHW Hex 3 HexNAc 4

**Mass:**  2097.8390

(-7.818e-08)

**Charge States:** 2, 3

**Start Position**: 1096

**End Position**: 1103

| Profile Score | Tandem Score | TDA q-value | Volume | Spectrum Matches |
| --- | --- | --- | --- | --- |
| 17.2758 | 87.1265 | 0.0079 | 9.2013e+08 | 5 |

**Adduct Signal Fractions**

- **Unmodified**: 9.2013e+08

#### VSN(N)GTHWF Fuc 1 Hex 4 HexNAc 4 Neu5Ac 1

**Mass:**  2844.1121

(-3.755e-08)

**Charge States:** 2, 3

**Start Position**: 1096

**End Position**: 1104

| Profile Score | Tandem Score | TDA q-value | Volume | Spectrum Matches |
| --- | --- | --- | --- | --- |
| 13.9264 | 86.1884 | 0.0079 | 8.2858e+09 | 9 |

**Adduct Signal Fractions**

- **Unmodified**: 8.2858e+09

#### VSN(N)GTHW Hex 4 HexNAc 6 Neu5Ac 1

**Mass:**  2957.1462

(-3.614e-08)

**Charge States:** 2, 3

**Start Position**: 1096

**End Position**: 1103

| Profile Score | Tandem Score | TDA q-value | Volume | Spectrum Matches |
| --- | --- | --- | --- | --- |
| 16.6828 | 85.9183 | 0.0079 | 3.7147e+08 | 3 |

**Adduct Signal Fractions**

- **Unmodified**: 3.7147e+08

#### VPAQEKN(N)F Fuc 1 Hex 5 HexNAc 5

**Mass:**  2903.1981

(1.029e-06)

**Charge States:** 2, 3, 4

**Start Position**: 1068

**End Position**: 1076

| Profile Score | Tandem Score | TDA q-value | Volume | Spectrum Matches |
| --- | --- | --- | --- | --- |
| 12.8644 | 85.5379 | 0.0079 | 2.6735e+09 | 14 |

**Adduct Signal Fractions**

- **Unmodified**: 2.6735e+09

#### VSN(N)GTHWF Fuc 1 Hex 4 HexNAc 4

**Mass:**  2553.0165

(-4.270e-07)

**Charge States:** 2, 3

**Start Position**: 1096

**End Position**: 1104

| Profile Score | Tandem Score | TDA q-value | Volume | Spectrum Matches |
| --- | --- | --- | --- | --- |
| 16.6739 | 85.0278 | 0.0079 | 9.0069e+08 | 7 |

**Adduct Signal Fractions**

- **Unmodified**: 9.0069e+08

#### VPAQEKN(N)F Fuc 1 Hex 4 HexNAc 4

**Mass:**  2538.0661

(1.025e-06)

**Charge States:** 2, 3, 4

**Start Position**: 1068

**End Position**: 1076

| Profile Score | Tandem Score | TDA q-value | Volume | Spectrum Matches |
| --- | --- | --- | --- | --- |
| 16.5085 | 84.6706 | 0.0079 | 5.8050e+09 | 15 |

**Adduct Signal Fractions**

- **Unmodified**: 5.8050e+09

#### VPAQEKN(N)F Hex 6 HexNAc 3 Neu5Ac 1

**Mass:**  2804.1270

(-9.751e-08)

**Charge States:** 2, 3

**Start Position**: 1068

**End Position**: 1076

| Profile Score | Tandem Score | TDA q-value | Volume | Spectrum Matches |
| --- | --- | --- | --- | --- |
| 15.2233 | 84.3067 | 0.0079 | 3.4098e+09 | 6 |

**Adduct Signal Fractions**

- **Unmodified**: 3.4098e+09

#### VSN(N)GTHW Hex 5 HexNAc 4 Neu5Ac 1

**Mass:**  2713.0353

(-1.018e-06)

**Charge States:** 2, 3

**Start Position**: 1096

**End Position**: 1103

| Profile Score | Tandem Score | TDA q-value | Volume | Spectrum Matches |
| --- | --- | --- | --- | --- |
| 16.5522 | 84.0010 | 0.0079 | 1.2538e+09 | 6 |

**Adduct Signal Fractions**

- **Unmodified**: 1.2538e+09

#### VPAQEKN(N)F Hex 4 HexNAc 2

**Mass:**  1985.8474

(1.634e-07)

**Charge States:** 2, 3

**Start Position**: 1068

**End Position**: 1076

| Profile Score | Tandem Score | TDA q-value | Volume | Spectrum Matches |
| --- | --- | --- | --- | --- |
| 15.4442 | 83.6272 | 0.0079 | 6.0489e+09 | 16 |

**Adduct Signal Fractions**

- **Unmodified**: 6.0489e+09

#### VSN(N)GTHW Hex 4 HexNAc 4 Neu5Ac 1

**Mass:**  2550.9847

(-4.791e-07)

**Charge States:** 2, 3

**Start Position**: 1096

**End Position**: 1103

| Profile Score | Tandem Score | TDA q-value | Volume | Spectrum Matches |
| --- | --- | --- | --- | --- |
| 18.5295 | 83.3423 | 0.0079 | 2.0979e+09 | 10 |

**Adduct Signal Fractions**

- **Unmodified**: 2.0979e+09

#### VSN(N)GTHW Fuc 1 Hex 3 HexNAc 4

**Mass:**  2243.8965

(-4.681e-07)

**Charge States:** 2, 3

**Start Position**: 1096

**End Position**: 1103

| Profile Score | Tandem Score | TDA q-value | Volume | Spectrum Matches |
| --- | --- | --- | --- | --- |
| 17.4172 | 83.3212 | 0.0079 | 1.3480e+09 | 8 |

**Adduct Signal Fractions**

- **Unmodified**: 1.3480e+09

#### VSN(N)GTHW Hex 6 HexNAc 3 Neu5Ac 1

**Mass:**  2672.0126

(-6.589e-07)

**Charge States:** 2, 3

**Start Position**: 1096

**End Position**: 1103

| Profile Score | Tandem Score | TDA q-value | Volume | Spectrum Matches |
| --- | --- | --- | --- | --- |
| 16.7191 | 83.2416 | 0.0079 | 1.1044e+09 | 6 |

**Adduct Signal Fractions**

- **Unmodified**: 1.1044e+09

#### VPAQEKN(N)F Fuc 1 Hex 4 HexNAc 5

**Mass:**  2741.1421

(6.948e-08)

**Charge States:** 2, 3, 4

**Start Position**: 1068

**End Position**: 1076

| Profile Score | Tandem Score | TDA q-value | Volume | Spectrum Matches |
| --- | --- | --- | --- | --- |
| 14.8031 | 83.1909 | 0.0079 | 1.1396e+10 | 22 |

**Adduct Signal Fractions**

- **Unmodified**: 1.1396e+10

#### VPAQEKN(N)F Fuc 1 Hex 6 HexNAc 3

**Mass:**  2659.0902

(2.978e-07)

**Charge States:** 2, 3

**Start Position**: 1068

**End Position**: 1076

| Profile Score | Tandem Score | TDA q-value | Volume | Spectrum Matches |
| --- | --- | --- | --- | --- |
| 16.4907 | 80.3735 | 0.0079 | 2.6735e+08 | 3 |

**Adduct Signal Fractions**

- **Unmodified**: 2.6735e+08

#### VPAQEKN(N)F Fuc 1 Hex 4 HexNAc 3 Neu5Ac 1

**Mass:**  2626.0793

(-4.573e-08)

**Charge States:** 2, 3

**Start Position**: 1068

**End Position**: 1076

| Profile Score | Tandem Score | TDA q-value | Volume | Spectrum Matches |
| --- | --- | --- | --- | --- |
| 12.9660 | 77.9238 | 0.0079 | 8.5133e+08 | 4 |

**Adduct Signal Fractions**

- **Unmodified**: 8.5133e+08

#### VPAQEKN(N)F Hex 3 HexNAc 4

**Mass:**  2229.9495

(-1.427e-06)

**Charge States:** 2, 3

**Start Position**: 1068

**End Position**: 1076

| Profile Score | Tandem Score | TDA q-value | Volume | Spectrum Matches |
| --- | --- | --- | --- | --- |
| 16.2582 | 72.1998 | 0.0079 | 7.6689e+08 | 8 |

**Adduct Signal Fractions**

- **Unmodified**: 7.6689e+08

#### VPAQEKN(N)F Hex 3 HexNAc 2

**Mass:**  1823.8004

(3.423e-06)

**Charge States:** 2, 3

**Start Position**: 1068

**End Position**: 1076

| Profile Score | Tandem Score | TDA q-value | Volume | Spectrum Matches |
| --- | --- | --- | --- | --- |
| 15.5610 | 71.5903 | 0.0079 | 1.6998e+09 | 7 |

**Adduct Signal Fractions**

- **Unmodified**: 1.6998e+09

#### HAIHVSGTN(N)GTKRF Fuc 1 Hex 4 HexNAc 7 Neu5Ac 1

**Mass:**  4030.7127

(-6.774e-07)

**Charge States:** 3, 4, 5

**Start Position**: 66

**End Position**: 80

| Profile Score | Tandem Score | TDA q-value | Volume | Spectrum Matches |
| --- | --- | --- | --- | --- |
| 13.8000 | 64.8451 | 0.0020 | 6.1022e+09 | 17 |

**Adduct Signal Fractions**

- **Unmodified**: 6.1022e+09

#### HAIHVSGTN(N)GTKRF Fuc 1 Hex 4 HexNAc 6 Neu5Ac 1

**Mass:**  3827.6406

(1.636e-06)

**Charge States:** 3, 4, 5

**Start Position**: 66

**End Position**: 80

| Profile Score | Tandem Score | TDA q-value | Volume | Spectrum Matches |
| --- | --- | --- | --- | --- |
| 11.8244 | 61.8808 | 0.0020 | 4.2945e+09 | 29 |

**Adduct Signal Fractions**

- **Unmodified**: 4.2945e+09

#### HAIHVSGTN(N)GTKRF @sulfate 1 Fuc 1 Hex 4 HexNAc 7

**Mass:**  3819.5746

(-6.557e-07)

**Charge States:** 3, 4

**Start Position**: 66

**End Position**: 80

| Profile Score | Tandem Score | TDA q-value | Volume | Spectrum Matches |
| --- | --- | --- | --- | --- |
| 12.7580 | 60.8521 | 0.0020 | 1.9127e+09 | 15 |

**Adduct Signal Fractions**

- **Unmodified**: 1.9127e+09

#### SALEPLVDLPIGIN(N)ITRF Hex 8 HexNAc 2

**Mass:**  3669.6889

(-1.488e-07)

**Charge States:** 3, 4

**Start Position**: 221

**End Position**: 239

| Profile Score | Tandem Score | TDA q-value | Volume | Spectrum Matches |
| --- | --- | --- | --- | --- |
| 16.6821 | 54.9085 | 0.0020 | 2.1299e+11 | 47 |

**Adduct Signal Fractions**

- **Unmodified**: 2.1299e+11

#### HAIHVSGTN(N)GTKRF Fuc 1 Hex 5 HexNAc 6 Neu5Ac 2

**Mass:**  4280.7870

(6.094e-07)

**Charge States:** 3, 4, 5

**Start Position**: 66

**End Position**: 80

| Profile Score | Tandem Score | TDA q-value | Volume | Spectrum Matches |
| --- | --- | --- | --- | --- |
| 11.3761 | 52.4338 | 0.0020 | 6.3260e+09 | 23 |

**Adduct Signal Fractions**

- **Unmodified**: 6.3260e+09

#### VDLPIGIN(N)ITRF Hex 5 HexNAc 2

**Mass:**  2573.2009

(2.384e-07)

**Charge States:** 2, 3

**Start Position**: 227

**End Position**: 239

| Profile Score | Tandem Score | TDA q-value | Volume | Spectrum Matches |
| --- | --- | --- | --- | --- |
| 15.8975 | 51.7983 | 0.0020 | 2.2129e+08 | 1 |

**Adduct Signal Fractions**

- **Unmodified**: 2.2129e+08

#### HAIHVSGTN(N)GTKRF Fuc 1 Hex 5 HexNAc 4 Neu5Ac 1

**Mass:**  3583.5291

(-6.244e-07)

**Charge States:** 3, 4, 5

**Start Position**: 66

**End Position**: 80

| Profile Score | Tandem Score | TDA q-value | Volume | Spectrum Matches |
| --- | --- | --- | --- | --- |
| 13.9819 | 51.0166 | 0.0020 | 6.0648e+09 | 27 |

**Adduct Signal Fractions**

- **Unmodified**: 6.0648e+09

#### HAIHVSGTN(N)GTKRF @sulfate 1 Fuc 1 Hex 4 HexNAc 7 Neu5Ac 1

**Mass:**  4110.6751

(5.195e-07)

**Charge States:** 3, 4, 5

**Start Position**: 66

**End Position**: 80

| Profile Score | Tandem Score | TDA q-value | Volume | Spectrum Matches |
| --- | --- | --- | --- | --- |
| 11.8933 | 48.7599 | 0.0020 | 4.2406e+09 | 14 |

**Adduct Signal Fractions**

- **Unmodified**: 4.2406e+09

#### HAIHVSGTN(N)GTKRF Fuc 1 Hex 7 HexNAc 5 Neu5Ac 2

**Mass:**  4401.7731

(-9.409e-06)

**Charge States:** 3, 4, 5

**Start Position**: 66

**End Position**: 80

| Profile Score | Tandem Score | TDA q-value | Volume | Spectrum Matches |
| --- | --- | --- | --- | --- |
| 15.5004 | 48.7373 | 0.0020 | 1.1172e+09 | 3 |

**Adduct Signal Fractions**

- **Unmodified**: 1.1172e+09

#### FHAIHVSGTN(N)GTKRF Hex 4 HexNAc 5 Neu5Ac 1

**Mass:**  3625.5733

(1.204e-06)

**Charge States:** 3, 4, 5

**Start Position**: 65

**End Position**: 80

| Profile Score | Tandem Score | TDA q-value | Volume | Spectrum Matches |
| --- | --- | --- | --- | --- |
| 16.3214 | 47.0101 | 0.0020 | 1.7733e+10 | 16 |

**Adduct Signal Fractions**

- **Unmodified**: 1.7733e+10

#### HAIHVSGTN(N)GTKRF Fuc 1 Hex 4 HexNAc 3 Neu5Ac 1

**Mass:**  3218.4042

(2.145e-06)

**Charge States:** 3, 4

**Start Position**: 66

**End Position**: 80

| Profile Score | Tandem Score | TDA q-value | Volume | Spectrum Matches |
| --- | --- | --- | --- | --- |
| 12.9736 | 45.3031 | 0.0020 | 6.8597e+08 | 3 |

**Adduct Signal Fractions**

- **Unmodified**: 6.8597e+08

#### HAIHVSGTN(N)GTKRF Fuc 3 Hex 5 HexNAc 4

**Mass:**  3584.5325

(1.506e-06)

**Charge States:** 3, 4, 5

**Start Position**: 66

**End Position**: 80

| Profile Score | Tandem Score | TDA q-value | Volume | Spectrum Matches |
| --- | --- | --- | --- | --- |
| -3.3342 | 44.9779 | 0.0020 | 5.2208e+08 | 6 |

**Adduct Signal Fractions**

- **Unmodified**: 5.2208e+08

#### HAIHVSGTN(N)GTKRF Fuc 1 Hex 6 HexNAc 5 Neu5Ac 2

**Mass:**  4239.7640

(8.669e-07)

**Charge States:** 4, 5

**Start Position**: 66

**End Position**: 80

| Profile Score | Tandem Score | TDA q-value | Volume | Spectrum Matches |
| --- | --- | --- | --- | --- |
| 14.1143 | 41.7269 | 0.0020 | 1.7890e+09 | 5 |

**Adduct Signal Fractions**

- **Unmodified**: 1.7890e+09

#### SALEPLVDLPIGIN(N)ITRF Hex 9 HexNAc 2

**Mass:**  3831.7445

(-8.108e-07)

**Charge States:** 3, 4

**Start Position**: 221

**End Position**: 239

| Profile Score | Tandem Score | TDA q-value | Volume | Spectrum Matches |
| --- | --- | --- | --- | --- |
| 18.1097 | 41.4244 | 0.0020 | 1.6966e+11 | 25 |

**Adduct Signal Fractions**

- **Unmodified**: 1.6966e+11

#### SALEPLVDLPIGIN(N)ITRF Hex 6 HexNAc 2

**Mass:**  3345.5859

(-2.816e-08)

**Charge States:** 3, 4

**Start Position**: 221

**End Position**: 239

| Profile Score | Tandem Score | TDA q-value | Volume | Spectrum Matches |
| --- | --- | --- | --- | --- |
| 17.8121 | 41.2378 | 0.0020 | 2.8371e+10 | 22 |

**Adduct Signal Fractions**

- **Unmodified**: 2.8371e+10

#### SALEPLVDLPIGIN(N)ITRF Hex 7 HexNAc 2

**Mass:**  3507.6394

(1.118e-06)

**Charge States:** 3, 4

**Start Position**: 221

**End Position**: 239

| Profile Score | Tandem Score | TDA q-value | Volume | Spectrum Matches |
| --- | --- | --- | --- | --- |
| 18.8507 | 39.4983 | 0.0020 | 9.0231e+10 | 33 |

**Adduct Signal Fractions**

- **Unmodified**: 9.0231e+10

#### HAIHVSGTN(N)GTKRF Fuc 3 Hex 5 HexNAc 5

**Mass:**  3787.6346

(-1.483e-06)

**Charge States:** 3, 4, 5

**Start Position**: 66

**End Position**: 80

| Profile Score | Tandem Score | TDA q-value | Volume | Spectrum Matches |
| --- | --- | --- | --- | --- |
| 15.6081 | 38.6350 | 0.0020 | 5.0482e+08 | 8 |

**Adduct Signal Fractions**

- **Unmodified**: 5.0482e+08

#### HAIHVSGTN(N)GTKRF @sulfate 1 Fuc 1 Hex 5 HexNAc 6 Neu5Ac 2

**Mass:**  4360.7393

(1.346e-07)

**Charge States:** 3, 4, 5

**Start Position**: 66

**End Position**: 80

| Profile Score | Tandem Score | TDA q-value | Volume | Spectrum Matches |
| --- | --- | --- | --- | --- |
| 15.2253 | 37.1237 | 0.0020 | 3.1503e+09 | 8 |

**Adduct Signal Fractions**

- **Unmodified**: 3.1503e+09

#### SALEPLVDLPIGIN(N)ITRF Hex 5 HexNAc 2

**Mass:**  3183.5353

(7.403e-08)

**Charge States:** 2, 3, 4

**Start Position**: 221

**End Position**: 239

| Profile Score | Tandem Score | TDA q-value | Volume | Spectrum Matches |
| --- | --- | --- | --- | --- |
| 17.2238 | 36.8665 | 0.0020 | 3.2867e+10 | 22 |

**Adduct Signal Fractions**

- **Unmodified**: 3.2867e+10

#### GEVFN(N)ATRF Fuc 1 Hex 5 HexNAc 5

**Mass:**  3011.2298

(6.243e-08)

**Charge States:** 2, 3

**Start Position**: 339

**End Position**: 348

| Profile Score | Tandem Score | TDA q-value | Volume | Spectrum Matches |
| --- | --- | --- | --- | --- |
| 17.0261 | 36.4572 | 0.0079 | 2.4461e+09 | 3 |

**Adduct Signal Fractions**

- **Unmodified**: 2.4461e+09

#### HAIHVSGTN(N)GTKRF @sulfate 1 Fuc 1 Hex 4 HexNAc 6 Neu5Ac 1

**Mass:**  3907.5969

(1.024e-06)

**Charge States:** 3, 4

**Start Position**: 66

**End Position**: 80

| Profile Score | Tandem Score | TDA q-value | Volume | Spectrum Matches |
| --- | --- | --- | --- | --- |
| 12.0571 | 36.4438 | 0.0020 | 9.6617e+08 | 6 |

**Adduct Signal Fractions**

- **Unmodified**: 9.6617e+08

#### GEVFN(N)ATRF Hex 5 HexNAc 2

**Mass:**  2255.9314

(2.675e-07)

**Charge States:** 2

**Start Position**: 339

**End Position**: 348

| Profile Score | Tandem Score | TDA q-value | Volume | Spectrum Matches |
| --- | --- | --- | --- | --- |
| 16.4369 | 36.2368 | 0.0079 | 2.0082e+09 | 2 |

**Adduct Signal Fractions**

- **Unmodified**: 2.0082e+09

#### VDLPIGIN(N)ITRF Hex 7 HexNAc 2

**Mass:**  2897.3073

(1.001e-06)

**Charge States:** 2, 3

**Start Position**: 227

**End Position**: 239

| Profile Score | Tandem Score | TDA q-value | Volume | Spectrum Matches |
| --- | --- | --- | --- | --- |
| 13.1540 | 35.9748 | 0.0020 | 8.4186e+08 | 4 |

**Adduct Signal Fractions**

- **Unmodified**: 8.4186e+08

#### GEVFN(N)ATRF Fuc 1 Hex 4 HexNAc 3

**Mass:**  2443.0190

(8.107e-07)

**Charge States:** 2, 3

**Start Position**: 339

**End Position**: 348

| Profile Score | Tandem Score | TDA q-value | Volume | Spectrum Matches |
| --- | --- | --- | --- | --- |
| 17.2294 | 34.9650 | 0.0079 | 6.2049e+08 | 4 |

**Adduct Signal Fractions**

- **Unmodified**: 6.2049e+08

#### GEVFN(N)ATRF Fuc 1 Hex 4 HexNAc 4 Neu5Ac 1

**Mass:**  2937.1877

(1.072e-06)

**Charge States:** 2, 3

**Start Position**: 339

**End Position**: 348

| Profile Score | Tandem Score | TDA q-value | Volume | Spectrum Matches |
| --- | --- | --- | --- | --- |
| 17.5035 | 34.4042 | 0.0079 | 1.5882e+09 | 6 |

**Adduct Signal Fractions**

- **Unmodified**: 1.5882e+09

#### HAIHVSGTN(N)GTKRF @sulfate 1 Fuc 1 Hex 5 HexNAc 6

**Mass:**  3778.5541

(6.678e-07)

**Charge States:** 4

**Start Position**: 66

**End Position**: 80

| Profile Score | Tandem Score | TDA q-value | Volume | Spectrum Matches |
| --- | --- | --- | --- | --- |
| 15.5128 | 33.1502 | 0.0020 | 2.8449e+08 | 1 |

**Adduct Signal Fractions**

- **Unmodified**: 2.8449e+08

#### GEVFN(N)ATRF Fuc 1 Hex 5 HexNAc 3

**Mass:**  2605.0745

(1.085e-06)

**Charge States:** 2, 3

**Start Position**: 339

**End Position**: 348

| Profile Score | Tandem Score | TDA q-value | Volume | Spectrum Matches |
| --- | --- | --- | --- | --- |
| 12.4064 | 32.3083 | 0.0079 | 8.0225e+08 | 9 |

**Adduct Signal Fractions**

- **Unmodified**: 8.0225e+08

#### HAIHVSGTN(N)GTKRF Fuc 1 Hex 7 HexNAc 6 Neu5Ac 3

**Mass:**  4895.9874

(-1.536e-08)

**Charge States:** 4, 5

**Start Position**: 66

**End Position**: 80

| Profile Score | Tandem Score | TDA q-value | Volume | Spectrum Matches |
| --- | --- | --- | --- | --- |
| 12.4967 | 32.1350 | 0.0020 | 1.8586e+09 | 6 |

**Adduct Signal Fractions**

- **Unmodified**: 1.8586e+09

#### FHAIHVSGTN(N)GTKRF Fuc 1 Hex 4 HexNAc 5 Neu5Ac 1

**Mass:**  3771.6404

(3.877e-06)

**Charge States:** 3, 4, 5

**Start Position**: 65

**End Position**: 80

| Profile Score | Tandem Score | TDA q-value | Volume | Spectrum Matches |
| --- | --- | --- | --- | --- |
| 15.1775 | 31.7635 | 0.0020 | 1.4726e+09 | 9 |

**Adduct Signal Fractions**

- **Unmodified**: 1.4726e+09

#### HAIHVSGTN(N)GTKRF Hex 4 HexNAc 3 Neu5Ac 1

**Mass:**  3074.3696

(6.601e-04)

**Charge States:** 4

**Start Position**: 66

**End Position**: 80

| Profile Score | Tandem Score | TDA q-value | Volume | Spectrum Matches |
| --- | --- | --- | --- | --- |
| -6.4294 | 31.6682 | 0.0020 | 4.1046e+06 | 1 |

**Adduct Signal Fractions**

- **Unmodified**: 4.1046e+06

#### FHAIHVSGTN(N)GTKRF Fuc 2 Hex 4 HexNAc 5

**Mass:**  3626.5886

(7.400e-07)

**Charge States:** 3, 4, 5

**Start Position**: 65

**End Position**: 80

| Profile Score | Tandem Score | TDA q-value | Volume | Spectrum Matches |
| --- | --- | --- | --- | --- |
| 15.3085 | 31.5148 | 0.0020 | 7.8955e+08 | 9 |

**Adduct Signal Fractions**

- **Unmodified**: 7.8955e+08

#### HAIHVSGTN(N)GTKRF Fuc 2 Hex 6 HexNAc 5

**Mass:**  3803.6236

(-1.325e-06)

**Charge States:** 4, 5

**Start Position**: 66

**End Position**: 80

| Profile Score | Tandem Score | TDA q-value | Volume | Spectrum Matches |
| --- | --- | --- | --- | --- |
| 13.7344 | 30.9680 | 0.0020 | 3.2135e+08 | 10 |

**Adduct Signal Fractions**

- **Unmodified**: 3.2135e+08

#### FHAIHVSGTN(N)GTKRF Fuc 1 Hex 3 HexNAc 4

**Mass:**  3115.3988

(3.626e-07)

**Charge States:** 3, 4, 5

**Start Position**: 65

**End Position**: 80

| Profile Score | Tandem Score | TDA q-value | Volume | Spectrum Matches |
| --- | --- | --- | --- | --- |
| 14.8038 | 30.6104 | 0.0020 | 4.2261e+08 | 7 |

**Adduct Signal Fractions**

- **Unmodified**: 4.2261e+08

#### HAIHVSGTN(N)GTKRF Fuc 1 Hex 5 HexNAc 5 Neu5Ac 2

**Mass:**  4077.7053

(8.748e-07)

**Charge States:** 4, 5

**Start Position**: 66

**End Position**: 80

| Profile Score | Tandem Score | TDA q-value | Volume | Spectrum Matches |
| --- | --- | --- | --- | --- |
| 10.5928 | 30.4078 | 0.0020 | 1.7778e+09 | 12 |

**Adduct Signal Fractions**

- **Unmodified**: 1.7778e+09

#### HAIHVSGTN(N)GTKRF @sulfate 1 Fuc 1 Hex 5 HexNAc 6 Neu5Ac 1

**Mass:**  4069.6458

(-1.707e-07)

**Charge States:** 3, 4

**Start Position**: 66

**End Position**: 80

| Profile Score | Tandem Score | TDA q-value | Volume | Spectrum Matches |
| --- | --- | --- | --- | --- |
| 12.9385 | 29.9611 | 0.0020 | 2.5405e+09 | 12 |

**Adduct Signal Fractions**

- **Unmodified**: 2.5405e+09

#### HAIHVSGTN(N)GTKRF Fuc 1 Hex 6 HexNAc 5 Neu5Ac 3

**Mass:**  4530.8555

(3.591e-07)

**Charge States:** 4, 5

**Start Position**: 66

**End Position**: 80

| Profile Score | Tandem Score | TDA q-value | Volume | Spectrum Matches |
| --- | --- | --- | --- | --- |
| 13.0666 | 29.8194 | 0.0020 | 4.2667e+09 | 7 |

**Adduct Signal Fractions**

- **Unmodified**: 4.2667e+09

#### HAIHVSGTN(N)GTKRF Fuc 1 Hex 7 HexNAc 7 Neu5Ac 1

**Mass:**  4516.8794

(8.361e-09)

**Charge States:** 4, 5

**Start Position**: 66

**End Position**: 80

| Profile Score | Tandem Score | TDA q-value | Volume | Spectrum Matches |
| --- | --- | --- | --- | --- |
| 15.5884 | 29.6521 | 0.0020 | 6.2747e+08 | 3 |

**Adduct Signal Fractions**

- **Unmodified**: 6.2747e+08

#### HAIHVSGTN(N)GTKRF Fuc 1 Hex 6 HexNAc 7 Neu5Ac 1

**Mass:**  4354.8185

(-7.615e-07)

**Charge States:** 4, 5

**Start Position**: 66

**End Position**: 80

| Profile Score | Tandem Score | TDA q-value | Volume | Spectrum Matches |
| --- | --- | --- | --- | --- |
| 11.4497 | 29.5328 | 0.0020 | 1.1873e+09 | 9 |

**Adduct Signal Fractions**

- **Unmodified**: 1.1873e+09

#### FHAIHVSGTN(N)GTKRF Hex 3 HexNAc 6

**Mass:**  3375.4944

(7.738e-07)

**Charge States:** 5

**Start Position**: 65

**End Position**: 80

| Profile Score | Tandem Score | TDA q-value | Volume | Spectrum Matches |
| --- | --- | --- | --- | --- |
| -4.2324 | 28.9083 | 0.0020 | 1.0225e+07 | 3 |

**Adduct Signal Fractions**

- **Unmodified**: 1.0225e+07

#### VDLPIGIN(N)ITRF Hex 6 HexNAc 2

**Mass:**  2735.2493

(1.241e-06)

**Charge States:** 2, 3

**Start Position**: 227

**End Position**: 239

| Profile Score | Tandem Score | TDA q-value | Volume | Spectrum Matches |
| --- | --- | --- | --- | --- |
| 13.6337 | 27.1170 | 0.0020 | 1.9157e+08 | 2 |

**Adduct Signal Fractions**

- **Unmodified**: 1.9157e+08

#### FHAIHVSGTN(N)GTKRF Fuc 2 Hex 4 HexNAc 5 Neu5Ac 1

**Mass:**  3917.6856

(8.315e-07)

**Charge States:** 4, 5

**Start Position**: 65

**End Position**: 80

| Profile Score | Tandem Score | TDA q-value | Volume | Spectrum Matches |
| --- | --- | --- | --- | --- |
| 10.0202 | 26.8856 | 0.0020 | 1.6689e+08 | 4 |

**Adduct Signal Fractions**

- **Unmodified**: 1.6689e+08

#### SN(N)NSIAIPTN(N)F Hex 10 HexNAc 4

**Mass:**  3609.4279

(1.162e-06)

**Charge States:** 3

**Start Position**: 708

**End Position**: 719

| Profile Score | Tandem Score | TDA q-value | Volume | Spectrum Matches |
| --- | --- | --- | --- | --- |
| 17.2158 | 26.4189 | 0.0020 | 1.4658e+10 | 7 |

**Adduct Signal Fractions**

- **Unmodified**: 1.4658e+10

#### VSN(N)GTHWF Fuc 1 Hex 3 HexNAc 5

**Mass:**  2594.0422

(-2.911e-07)

**Charge States:** 2, 3

**Start Position**: 1096

**End Position**: 1104

| Profile Score | Tandem Score | TDA q-value | Volume | Spectrum Matches |
| --- | --- | --- | --- | --- |
| 15.2847 | 26.1813 | 0.0079 | 1.2267e+09 | 5 |

**Adduct Signal Fractions**

- **Unmodified**: 1.2267e+09

#### FHAIHVSGTN(N)GTKRF Fuc 1 Hex 3 HexNAc 6

**Mass:**  3521.5565

(5.704e-07)

**Charge States:** 3, 4, 5

**Start Position**: 65

**End Position**: 80

| Profile Score | Tandem Score | TDA q-value | Volume | Spectrum Matches |
| --- | --- | --- | --- | --- |
| -5.3761 | 25.8695 | 0.0020 | 3.4750e+08 | 6 |

**Adduct Signal Fractions**

- **Unmodified**: 3.4750e+08

#### HAIHVSGTN(N)GTKRF Fuc 1 Hex 7 HexNAc 6

**Mass:**  4022.7079

(1.515e-06)

**Charge States:** 4, 5

**Start Position**: 66

**End Position**: 80

| Profile Score | Tandem Score | TDA q-value | Volume | Spectrum Matches |
| --- | --- | --- | --- | --- |
| 16.2449 | 25.6914 | 0.0020 | 1.8625e+08 | 2 |

**Adduct Signal Fractions**

- **Unmodified**: 1.8625e+08

#### FHAIHVSGTN(N)GTKRF @sulfate 1 Fuc 1 Hex 4 HexNAc 5

**Mass:**  3560.4856

(-3.997e-07)

**Charge States:** 3, 4

**Start Position**: 65

**End Position**: 80

| Profile Score | Tandem Score | TDA q-value | Volume | Spectrum Matches |
| --- | --- | --- | --- | --- |
| 14.5754 | 25.3031 | 0.0020 | 1.8852e+08 | 2 |

**Adduct Signal Fractions**

- **Unmodified**: 1.8852e+08

#### HAIHVSGTN(N)GTKRF Fuc 3 Hex 5 HexNAc 6

**Mass:**  3990.7111

(6.499e-09)

**Charge States:** 3, 4, 5

**Start Position**: 66

**End Position**: 80

| Profile Score | Tandem Score | TDA q-value | Volume | Spectrum Matches |
| --- | --- | --- | --- | --- |
| 14.2903 | 25.2830 | 0.0020 | 1.2914e+09 | 10 |

**Adduct Signal Fractions**

- **Unmodified**: 1.2914e+09

#### HAIHVSGTN(N)GTKRF Fuc 1 Hex 6 HexNAc 6 Neu5Ac 1

**Mass:**  4151.7401

(1.805e-06)

**Charge States:** 3, 4, 5

**Start Position**: 66

**End Position**: 80

| Profile Score | Tandem Score | TDA q-value | Volume | Spectrum Matches |
| --- | --- | --- | --- | --- |
| 12.1442 | 25.2165 | 0.0020 | 1.5395e+09 | 21 |

**Adduct Signal Fractions**

- **Unmodified**: 1.5395e+09

#### VPAQEKN(N)F Fuc 1 Hex 3 HexNAc 6

**Mass:**  2782.1705

(1.020e-06)

**Charge States:** 2, 3, 4

**Start Position**: 1068

**End Position**: 1076

| Profile Score | Tandem Score | TDA q-value | Volume | Spectrum Matches |
| --- | --- | --- | --- | --- |
| 16.5620 | 25.0062 | 0.0079 | 1.3740e+09 | 5 |

**Adduct Signal Fractions**

- **Unmodified**: 1.3740e+09

#### HAIHVSGTN(N)GTKRF @sulfate 1 Fuc 2 Hex 4 HexNAc 7 Neu5Ac 1

**Mass:**  4256.7291

(-3.667e-07)

**Charge States:** 3, 4, 5

**Start Position**: 66

**End Position**: 80

| Profile Score | Tandem Score | TDA q-value | Volume | Spectrum Matches |
| --- | --- | --- | --- | --- |
| 15.5352 | 24.9497 | 0.0020 | 2.0677e+09 | 10 |

**Adduct Signal Fractions**

- **Unmodified**: 2.0677e+09

#### HAIHVSGTN(N)GTKRF Fuc 1 Hex 6 HexNAc 7

**Mass:**  4063.7211

(-1.017e-06)

**Charge States:** 3, 4, 5

**Start Position**: 66

**End Position**: 80

| Profile Score | Tandem Score | TDA q-value | Volume | Spectrum Matches |
| --- | --- | --- | --- | --- |
| -5.6172 | 24.3773 | 0.0020 | 4.3246e+08 | 12 |

**Adduct Signal Fractions**

- **Unmodified**: 4.3246e+08

#### HAIHVSGTN(N)GTKRF Fuc 1 Hex 7 HexNAc 6 Neu5Ac 2

**Mass:**  4604.8982

(1.111e-06)

**Charge States:** 4, 5

**Start Position**: 66

**End Position**: 80

| Profile Score | Tandem Score | TDA q-value | Volume | Spectrum Matches |
| --- | --- | --- | --- | --- |
| 13.3498 | 24.2305 | 0.0020 | 9.7935e+08 | 9 |

**Adduct Signal Fractions**

- **Unmodified**: 9.7935e+08

#### VSN(N)GTHWF Hex 5 HexNAc 2

**Mass:**  2162.8517

(-6.796e-07)

**Charge States:** 2, 3

**Start Position**: 1096

**End Position**: 1104

| Profile Score | Tandem Score | TDA q-value | Volume | Spectrum Matches |
| --- | --- | --- | --- | --- |
| 18.2650 | 23.9089 | 0.0079 | 1.4528e+09 | 2 |

**Adduct Signal Fractions**

- **Unmodified**: 1.4528e+09

#### HAIHVSGTN(N)GTKRF Fuc 2 Hex 7 HexNAc 7 Neu5Ac 1

**Mass:**  4662.9319

(-7.651e-07)

**Charge States:** 4, 5

**Start Position**: 66

**End Position**: 80

| Profile Score | Tandem Score | TDA q-value | Volume | Spectrum Matches |
| --- | --- | --- | --- | --- |
| 12.6754 | 23.8342 | 0.0020 | 1.4766e+08 | 2 |

**Adduct Signal Fractions**

- **Unmodified**: 1.4766e+08

#### VSN(N)GTHWF Fuc 1 Hex 5 HexNAc 3

**Mass:**  2511.9913

(-7.518e-07)

**Charge States:** 2, 3

**Start Position**: 1096

**End Position**: 1104

| Profile Score | Tandem Score | TDA q-value | Volume | Spectrum Matches |
| --- | --- | --- | --- | --- |
| 17.7129 | 23.6772 | 0.0079 | 1.4991e+09 | 6 |

**Adduct Signal Fractions**

- **Unmodified**: 1.4991e+09

#### VSN(N)GTHW Fuc 1 Hex 4 HexNAc 6 Neu5Ac 1

**Mass:**  3103.2032

(3.455e-07)

**Charge States:** 2, 3

**Start Position**: 1096

**End Position**: 1103

| Profile Score | Tandem Score | TDA q-value | Volume | Spectrum Matches |
| --- | --- | --- | --- | --- |
| 16.9707 | 23.6310 | 0.0079 | 4.6278e+08 | 3 |

**Adduct Signal Fractions**

- **Unmodified**: 4.6278e+08

#### GEVFN(N)ATRF Fuc 2 Hex 4 HexNAc 5

**Mass:**  2996.2113

(3.348e-04)

**Charge States:** 3, 4

**Start Position**: 339

**End Position**: 348

| Profile Score | Tandem Score | TDA q-value | Volume | Spectrum Matches |
| --- | --- | --- | --- | --- |
| 9.1005 | 23.5296 | 0.0079 | 3.6100e+08 | 4 |

**Adduct Signal Fractions**

- **Unmodified**: 3.6100e+08

#### VSN(N)GTHWF Fuc 1 Hex 3 HexNAc 4

**Mass:**  2390.9633

(-1.038e-06)

**Charge States:** 2, 3

**Start Position**: 1096

**End Position**: 1104

| Profile Score | Tandem Score | TDA q-value | Volume | Spectrum Matches |
| --- | --- | --- | --- | --- |
| 16.4025 | 23.0241 | 0.0079 | 2.5526e+09 | 5 |

**Adduct Signal Fractions**

- **Unmodified**: 2.5526e+09

#### VSN(N)GTHWF Fuc 1 Hex 4 HexNAc 3

**Mass:**  2349.9377

(1.373e-06)

**Charge States:** 2, 3

**Start Position**: 1096

**End Position**: 1104

| Profile Score | Tandem Score | TDA q-value | Volume | Spectrum Matches |
| --- | --- | --- | --- | --- |
| 16.5669 | 22.9817 | 0.0079 | 1.8100e+09 | 5 |

**Adduct Signal Fractions**

- **Unmodified**: 1.8100e+09

#### VSN(N)GTHWF Fuc 1 Hex 4 HexNAc 5 Neu5Ac 1

**Mass:**  3047.1922

(3.411e-07)

**Charge States:** 2, 3

**Start Position**: 1096

**End Position**: 1104

| Profile Score | Tandem Score | TDA q-value | Volume | Spectrum Matches |
| --- | --- | --- | --- | --- |
| 13.2099 | 22.6016 | 0.0079 | 7.0126e+08 | 2 |

**Adduct Signal Fractions**

- **Unmodified**: 7.0126e+08

#### VSN(N)GTHWF Hex 6 HexNAc 3

**Mass:**  2527.9809

(-2.561e-07)

**Charge States:** 2, 3

**Start Position**: 1096

**End Position**: 1104

| Profile Score | Tandem Score | TDA q-value | Volume | Spectrum Matches |
| --- | --- | --- | --- | --- |
| 17.4528 | 22.5681 | 0.0079 | 2.3116e+09 | 7 |

**Adduct Signal Fractions**

- **Unmodified**: 2.3116e+09

#### HAIHVSGTN(N)GTKRF @sulfate 1 Fuc 1 Hex 3 HexNAc 8

**Mass:**  3860.6053

(4.420e-07)

**Charge States:** 3, 4

**Start Position**: 66

**End Position**: 80

| Profile Score | Tandem Score | TDA q-value | Volume | Spectrum Matches |
| --- | --- | --- | --- | --- |
| 11.7097 | 22.4510 | 0.0020 | 1.3106e+09 | 13 |

**Adduct Signal Fractions**

- **Unmodified**: 1.3106e+09

#### VSN(N)GTHWF Hex 4 HexNAc 4 Neu5Ac 1

**Mass:**  2698.0588

(2.013e-06)

**Charge States:** 2, 3

**Start Position**: 1096

**End Position**: 1104

| Profile Score | Tandem Score | TDA q-value | Volume | Spectrum Matches |
| --- | --- | --- | --- | --- |
| 16.2303 | 22.3670 | 0.0079 | 3.4198e+09 | 5 |

**Adduct Signal Fractions**

- **Unmodified**: 3.4198e+09

#### VPAQEKN(N)F Fuc 1 Hex 6 HexNAc 6

**Mass:**  3268.3292

(-5.644e-06)

**Charge States:** 3

**Start Position**: 1068

**End Position**: 1076

| Profile Score | Tandem Score | TDA q-value | Volume | Spectrum Matches |
| --- | --- | --- | --- | --- |
| 7.5167 | 21.9501 | 0.0231 | 6.3462e+07 | 1 |

**Adduct Signal Fractions**

- **Unmodified**: 6.3462e+07

#### HKNN(N)KSWMESEF Fuc 1 Hex 3 HexNAc 5

**Mass:**  3183.2965

(-2.687e-07)

**Charge States:** 2, 3, 4

**Start Position**: 146

**End Position**: 158

| Profile Score | Tandem Score | TDA q-value | Volume | Spectrum Matches |
| --- | --- | --- | --- | --- |
| 13.9116 | 21.5345 | 0.0020 | 2.6749e+10 | 15 |

**Adduct Signal Fractions**

- **Unmodified**: 2.6749e+10

#### HAIHVSGTN(N)GTKRF Fuc 2 Hex 4 HexNAc 6 Neu5Ac 1

**Mass:**  3973.6938

(-4.281e-07)

**Charge States:** 4, 5

**Start Position**: 66

**End Position**: 80

| Profile Score | Tandem Score | TDA q-value | Volume | Spectrum Matches |
| --- | --- | --- | --- | --- |
| 13.4062 | 21.3723 | 0.0020 | 1.3647e+09 | 17 |

**Adduct Signal Fractions**

- **Unmodified**: 1.3647e+09

#### HAIHVSGTN(N)GTKRF Fuc 2 Hex 6 HexNAc 5 Neu5Ac 1

**Mass:**  4094.7243

(-6.540e-07)

**Charge States:** 4, 5

**Start Position**: 66

**End Position**: 80

| Profile Score | Tandem Score | TDA q-value | Volume | Spectrum Matches |
| --- | --- | --- | --- | --- |
| 11.7854 | 21.0840 | 0.0020 | 8.0116e+08 | 16 |

**Adduct Signal Fractions**

- **Unmodified**: 8.0116e+08

#### SN(N)NSIAIPTN(N)F Hex 11 HexNAc 4

**Mass:**  3771.4777

(4.433e-07)

**Charge States:** 3

**Start Position**: 708

**End Position**: 719

| Profile Score | Tandem Score | TDA q-value | Volume | Spectrum Matches |
| --- | --- | --- | --- | --- |
| 17.1271 | 20.6693 | 0.0020 | 3.3013e+10 | 7 |

**Adduct Signal Fractions**

- **Unmodified**: 3.3013e+10

#### SN(N)NSIAIPTN(N)F Hex 13 HexNAc 4

**Mass:**  4095.5825

(-2.856e-07)

**Charge States:** 3

**Start Position**: 708

**End Position**: 719

| Profile Score | Tandem Score | TDA q-value | Volume | Spectrum Matches |
| --- | --- | --- | --- | --- |
| 17.6952 | 20.5550 | 0.0020 | 1.3926e+10 | 5 |

**Adduct Signal Fractions**

- **Unmodified**: 1.3926e+10

#### HAIHVSGTN(N)GTKRF Fuc 2 Hex 6 HexNAc 6 Neu5Ac 1

**Mass:**  4297.8014

(1.296e-06)

**Charge States:** 4, 5

**Start Position**: 66

**End Position**: 80

| Profile Score | Tandem Score | TDA q-value | Volume | Spectrum Matches |
| --- | --- | --- | --- | --- |
| 15.9610 | 20.5268 | 0.0020 | 1.5757e+08 | 2 |

**Adduct Signal Fractions**

- **Unmodified**: 1.5757e+08

#### RVQPTESIVRFPN(N)ITNL Fuc 1 Hex 5 HexNAc 4 Neu5Ac 1

**Mass:**  4042.8255

(-1.069e-07)

**Charge States:** 3, 4, 5

**Start Position**: 319

**End Position**: 336

| Profile Score | Tandem Score | TDA q-value | Volume | Spectrum Matches |
| --- | --- | --- | --- | --- |
| 15.3700 | 20.4106 | 0.0020 | 1.2688e+10 | 7 |

**Adduct Signal Fractions**

- **Unmodified**: 1.2688e+10

#### SALEPLVDLPIGIN(N)ITRF Hex 4 HexNAc 3

**Mass:**  3224.5600

(7.417e-07)

**Charge States:** 3, 4

**Start Position**: 221

**End Position**: 239

| Profile Score | Tandem Score | TDA q-value | Volume | Spectrum Matches |
| --- | --- | --- | --- | --- |
| 16.0129 | 20.3384 | 0.0020 | 3.0381e+09 | 3 |

**Adduct Signal Fractions**

- **Unmodified**: 3.0381e+09

#### N(N)FSQILPDPSKPSKRSF Hex 7 HexNAc 2

**Mass:**  3487.5565

(2.226e-06)

**Charge States:** 3, 4

**Start Position**: 801

**End Position**: 818

| Profile Score | Tandem Score | TDA q-value | Volume | Spectrum Matches |
| --- | --- | --- | --- | --- |
| 13.1839 | 20.1833 | 0.0020 | 4.4438e+09 | 8 |

**Adduct Signal Fractions**

- **Unmodified**: 4.4438e+09

#### N(N)FSQILPDPSKPSKRSF Hex 4 HexNAc 3

**Mass:**  3204.4791

(2.571e-06)

**Charge States:** 3, 4

**Start Position**: 801

**End Position**: 818

| Profile Score | Tandem Score | TDA q-value | Volume | Spectrum Matches |
| --- | --- | --- | --- | --- |
| 12.7378 | 19.3937 | 0.0020 | 3.6149e+09 | 6 |

**Adduct Signal Fractions**

- **Unmodified**: 3.6149e+09

#### RVQPTESIVRFPN(N)ITNL Hex 5 HexNAc 2

**Mass:**  3199.5130

(-1.226e-08)

**Charge States:** 3, 4

**Start Position**: 319

**End Position**: 336

| Profile Score | Tandem Score | TDA q-value | Volume | Spectrum Matches |
| --- | --- | --- | --- | --- |
| 17.4094 | 19.1071 | 0.0020 | 6.8091e+08 | 4 |

**Adduct Signal Fractions**

- **Unmodified**: 6.8091e+08

#### HKNN(N)KSWMESEF Fuc 1 Hex 5 HexNAc 4 Neu5Ac 1

**Mass:**  3595.4206

(7.205e-07)

**Charge States:** 3, 4

**Start Position**: 146

**End Position**: 158

| Profile Score | Tandem Score | TDA q-value | Volume | Spectrum Matches |
| --- | --- | --- | --- | --- |
| 16.9104 | 18.5268 | 0.0038 | 6.9818e+09 | 10 |

**Adduct Signal Fractions**

- **Unmodified**: 6.9818e+09

#### RVQPTESIVRFPN(N)ITNL Fuc 1 Hex 5 HexNAc 4

**Mass:**  3751.7295

(1.825e-08)

**Charge States:** 3, 4

**Start Position**: 319

**End Position**: 336

| Profile Score | Tandem Score | TDA q-value | Volume | Spectrum Matches |
| --- | --- | --- | --- | --- |
| 14.7299 | 18.1752 | 0.0038 | 9.2637e+09 | 6 |

**Adduct Signal Fractions**

- **Unmodified**: 9.2637e+09

#### HAIHVSGTN(N)GTKRF Fuc 2 Hex 5 HexNAc 6 Neu5Ac 2

**Mass:**  4426.8483

(1.649e-06)

**Charge States:** 3, 4, 5

**Start Position**: 66

**End Position**: 80

| Profile Score | Tandem Score | TDA q-value | Volume | Spectrum Matches |
| --- | --- | --- | --- | --- |
| 12.8026 | 18.1080 | 0.0038 | 2.8705e+09 | 14 |

**Adduct Signal Fractions**

- **Unmodified**: 2.8705e+09

#### FHAIHVSGTN(N)GTKRF Fuc 1 Hex 5 HexNAc 6 Neu5Ac 2

**Mass:**  4427.8528

(-9.881e-08)

**Charge States:** 4

**Start Position**: 65

**End Position**: 80

| Profile Score | Tandem Score | TDA q-value | Volume | Spectrum Matches |
| --- | --- | --- | --- | --- |
| -5.0960 | 18.0677 | 0.0038 | 5.4318e+06 | 1 |

**Adduct Signal Fractions**

- **Unmodified**: 5.4318e+06

#### HAIHVSGTN(N)GTKRF @sulfate 1 Fuc 1 Hex 6 HexNAc 3

**Mass:**  3331.3726

(2.045e-06)

**Charge States:** 4

**Start Position**: 66

**End Position**: 80

| Profile Score | Tandem Score | TDA q-value | Volume | Spectrum Matches |
| --- | --- | --- | --- | --- |
| 12.6396 | 18.0604 | 0.0038 | 4.5808e+07 | 1 |

**Adduct Signal Fractions**

- **Unmodified**: 4.5808e+07

#### SN(N)NSIAIPTN(N)F Hex 12 HexNAc 4

**Mass:**  3933.5306

(-2.871e-07)

**Charge States:** 3, 4

**Start Position**: 708

**End Position**: 719

| Profile Score | Tandem Score | TDA q-value | Volume | Spectrum Matches |
| --- | --- | --- | --- | --- |
| 15.7344 | 17.7926 | 0.0038 | 1.3130e+10 | 5 |

**Adduct Signal Fractions**

- **Unmodified**: 1.3130e+10

#### N(N)FSQILPDPSKPSKRSF Hex 5 HexNAc 2

**Mass:**  3163.4508

(1.786e-06)

**Charge States:** 2, 3, 4

**Start Position**: 801

**End Position**: 818

| Profile Score | Tandem Score | TDA q-value | Volume | Spectrum Matches |
| --- | --- | --- | --- | --- |
| 18.6776 | 17.5826 | 0.0038 | 1.2146e+11 | 25 |

**Adduct Signal Fractions**

- **Unmodified**: 1.2146e+11

#### HAIHVSGTN(N)GTKRF Fuc 1 Hex 7 HexNAc 7 Neu5Ac 2

**Mass:**  4807.9656

(-8.766e-07)

**Charge States:** 4, 5

**Start Position**: 66

**End Position**: 80

| Profile Score | Tandem Score | TDA q-value | Volume | Spectrum Matches |
| --- | --- | --- | --- | --- |
| 15.5948 | 17.5507 | 0.0038 | 2.1002e+08 | 4 |

**Adduct Signal Fractions**

- **Unmodified**: 2.1002e+08

#### SN(N)NSIAIPTN(N)F Hex 14 HexNAc 4

**Mass:**  4257.6347

(4.207e-08)

**Charge States:** 3

**Start Position**: 708

**End Position**: 719

| Profile Score | Tandem Score | TDA q-value | Volume | Spectrum Matches |
| --- | --- | --- | --- | --- |
| 13.7597 | 17.5077 | 0.0038 | 6.9629e+09 | 7 |

**Adduct Signal Fractions**

- **Unmodified**: 6.9629e+09

#### HAIHVSGTN(N)GTKRF Fuc 1 Hex 6 HexNAc 6 Neu5Ac 2

**Mass:**  4442.8440

(8.736e-07)

**Charge States:** 4, 5

**Start Position**: 66

**End Position**: 80

| Profile Score | Tandem Score | TDA q-value | Volume | Spectrum Matches |
| --- | --- | --- | --- | --- |
| 11.8296 | 17.2821 | 0.0056 | 1.0517e+09 | 17 |

**Adduct Signal Fractions**

- **Unmodified**: 1.0517e+09

#### RVQPTESIVRFPN(N)ITNL Fuc 1 Hex 5 HexNAc 5 Neu5Ac 1

**Mass:**  4245.9115

(1.429e-06)

**Charge States:** 3, 4

**Start Position**: 319

**End Position**: 336

| Profile Score | Tandem Score | TDA q-value | Volume | Spectrum Matches |
| --- | --- | --- | --- | --- |
| 11.6876 | 17.1630 | 0.0056 | 1.7592e+09 | 3 |

**Adduct Signal Fractions**

- **Unmodified**: 1.7592e+09

#### FHAIHVSGTN(N)GTKRF Fuc 1 Hex 3 HexNAc 5

**Mass:**  3318.4798

(2.590e-07)

**Charge States:** 3, 4, 5

**Start Position**: 65

**End Position**: 80

| Profile Score | Tandem Score | TDA q-value | Volume | Spectrum Matches |
| --- | --- | --- | --- | --- |
| 8.8951 | 16.9754 | 0.0056 | 1.0367e+09 | 15 |

**Adduct Signal Fractions**

- **Unmodified**: 1.0367e+09

#### HAIHVSGTN(N)GTKRF Fuc 1 Hex 5 HexNAc 8 Neu5Ac 1

**Mass:**  4395.8543

(1.462e-06)

**Charge States:** 4, 5

**Start Position**: 66

**End Position**: 80

| Profile Score | Tandem Score | TDA q-value | Volume | Spectrum Matches |
| --- | --- | --- | --- | --- |
| 14.2117 | 16.8415 | 0.0069 | 6.1261e+08 | 6 |

**Adduct Signal Fractions**

- **Unmodified**: 6.1261e+08

#### HKNN(N)KSWMESEF Fuc 1 Hex 4 HexNAc 5

**Mass:**  3345.3472

(-6.505e-07)

**Charge States:** 3, 4

**Start Position**: 146

**End Position**: 158

| Profile Score | Tandem Score | TDA q-value | Volume | Spectrum Matches |
| --- | --- | --- | --- | --- |
| 15.3180 | 16.4647 | 0.0069 | 2.1405e+10 | 18 |

**Adduct Signal Fractions**

- **Unmodified**: 2.1405e+10

#### N(N)FSQILPDPSKPSKRSF Hex 6 HexNAc 2

**Mass:**  3325.4972

(-9.033e-08)

**Charge States:** 3, 4

**Start Position**: 801

**End Position**: 818

| Profile Score | Tandem Score | TDA q-value | Volume | Spectrum Matches |
| --- | --- | --- | --- | --- |
| 12.9850 | 16.4617 | 0.0069 | 1.4157e+10 | 15 |

**Adduct Signal Fractions**

- **Unmodified**: 1.4157e+10

#### RVQPTESIVRFPN(N)ITNL Fuc 1 Hex 6 HexNAc 5 Neu5Ac 2

**Mass:**  4699.0543

(2.597e-08)

**Charge States:** 3, 4, 5

**Start Position**: 319

**End Position**: 336

| Profile Score | Tandem Score | TDA q-value | Volume | Spectrum Matches |
| --- | --- | --- | --- | --- |
| 13.7287 | 16.4195 | 0.0069 | 2.6406e+09 | 6 |

**Adduct Signal Fractions**

- **Unmodified**: 2.6406e+09

#### RVQPTESIVRFPN(N)ITNL Fuc 1 Hex 4 HexNAc 5 Neu5Ac 1

**Mass:**  4083.8517

(-1.121e-07)

**Charge States:** 3, 4, 5

**Start Position**: 319

**End Position**: 336

| Profile Score | Tandem Score | TDA q-value | Volume | Spectrum Matches |
| --- | --- | --- | --- | --- |
| 13.2000 | 16.1380 | 0.0069 | 5.2143e+09 | 12 |

**Adduct Signal Fractions**

- **Unmodified**: 5.2143e+09

#### RVQPTESIVRFPN(N)ITNL Fuc 1 Hex 4 HexNAc 4 Neu5Ac 1

**Mass:**  3880.7725

(-8.556e-08)

**Charge States:** 3, 4, 5

**Start Position**: 319

**End Position**: 336

| Profile Score | Tandem Score | TDA q-value | Volume | Spectrum Matches |
| --- | --- | --- | --- | --- |
| 17.8340 | 16.1184 | 0.0069 | 1.7128e+10 | 9 |

**Adduct Signal Fractions**

- **Unmodified**: 1.7128e+10

#### RVQPTESIVRFPN(N)ITNL Fuc 1 Hex 3 HexNAc 4

**Mass:**  3427.6271

(-6.138e-07)

**Charge States:** 3, 4

**Start Position**: 319

**End Position**: 336

| Profile Score | Tandem Score | TDA q-value | Volume | Spectrum Matches |
| --- | --- | --- | --- | --- |
| 16.1088 | 16.0390 | 0.0069 | 2.7674e+10 | 11 |

**Adduct Signal Fractions**

- **Unmodified**: 2.7674e+10

#### SALEPLVDLPIGIN(N)ITRF Hex 3 HexNAc 4

**Mass:**  3265.5860

(2.560e-07)

**Charge States:** 3, 4

**Start Position**: 221

**End Position**: 239

| Profile Score | Tandem Score | TDA q-value | Volume | Spectrum Matches |
| --- | --- | --- | --- | --- |
| 16.4671 | 15.7232 | 0.0069 | 3.2106e+09 | 6 |

**Adduct Signal Fractions**

- **Unmodified**: 3.2106e+09

#### N(N)FSQILPDPSKPSKRSF Fuc 1 Hex 3 HexNAc 4

**Mass:**  3391.5635

(2.367e-06)

**Charge States:** 3, 4

**Start Position**: 801

**End Position**: 818

| Profile Score | Tandem Score | TDA q-value | Volume | Spectrum Matches |
| --- | --- | --- | --- | --- |
| 13.0338 | 15.7052 | 0.0069 | 5.4825e+09 | 9 |

**Adduct Signal Fractions**

- **Unmodified**: 5.4825e+09

#### SN(N)NSIAIPTN(N)F Hex 11 HexNAc 5

**Mass:**  3974.5573

(3.347e-07)

**Charge States:** 3, 4

**Start Position**: 708

**End Position**: 719

| Profile Score | Tandem Score | TDA q-value | Volume | Spectrum Matches |
| --- | --- | --- | --- | --- |
| -1.9841 | 15.5617 | 0.0069 | 6.1017e+09 | 4 |

**Adduct Signal Fractions**

- **Unmodified**: 6.1017e+09

#### N(N)FSQILPDPSKPSKRSF Hex 8 HexNAc 2

**Mass:**  3649.6066

(1.047e-06)

**Charge States:** 3, 4

**Start Position**: 801

**End Position**: 818

| Profile Score | Tandem Score | TDA q-value | Volume | Spectrum Matches |
| --- | --- | --- | --- | --- |
| 16.9327 | 15.1556 | 0.0082 | 1.6589e+09 | 6 |

**Adduct Signal Fractions**

- **Unmodified**: 1.6589e+09

#### SN(N)NSIAIPTN(N)F Fuc 1 Hex 9 HexNAc 6

**Mass:**  4001.5818

(5.024e-04)

**Charge States:** 3, 4

**Start Position**: 708

**End Position**: 719

| Profile Score | Tandem Score | TDA q-value | Volume | Spectrum Matches |
| --- | --- | --- | --- | --- |
| 11.9075 | 15.0931 | 0.0082 | 3.0533e+08 | 4 |

**Adduct Signal Fractions**

- **Unmodified**: 3.0533e+08

#### N(N)FSQILPDPSKPSKRSF Fuc 1 Hex 4 HexNAc 4

**Mass:**  3553.6091

(3.108e-07)

**Charge States:** 3, 4

**Start Position**: 801

**End Position**: 818

| Profile Score | Tandem Score | TDA q-value | Volume | Spectrum Matches |
| --- | --- | --- | --- | --- |
| 15.8504 | 15.0587 | 0.0082 | 1.2449e+09 | 6 |

**Adduct Signal Fractions**

- **Unmodified**: 1.2449e+09

#### RVQPTESIVRFPN(N)ITNLC(C)PF Fuc 1 Hex 6 HexNAc 5 Neu5Ac 1

**Mass:**  4812.1081

(1.824e-07)

**Charge States:** 4, 5

**Start Position**: 319

**End Position**: 339

| Profile Score | Tandem Score | TDA q-value | Volume | Spectrum Matches |
| --- | --- | --- | --- | --- |
| 17.0808 | 14.8949 | 0.0082 | 7.1100e+09 | 5 |

**Adduct Signal Fractions**

- **Unmodified**: 7.1100e+09

#### HAIHVSGTN(N)GTKRF @sulfate 1 Fuc 1 Hex 4 HexNAc 4 Neu5Ac 1

**Mass:**  3501.4380

(1.949e-06)

**Charge States:** 4

**Start Position**: 66

**End Position**: 80

| Profile Score | Tandem Score | TDA q-value | Volume | Spectrum Matches |
| --- | --- | --- | --- | --- |
| 14.4076 | 14.7111 | 0.0082 | 6.9558e+07 | 2 |

**Adduct Signal Fractions**

- **Unmodified**: 6.9558e+07

#### N(N)FSQILPDPSKPSKRSF Hex 5 HexNAc 3

**Mass:**  3366.5307

(1.332e-06)

**Charge States:** 3, 4

**Start Position**: 801

**End Position**: 818

| Profile Score | Tandem Score | TDA q-value | Volume | Spectrum Matches |
| --- | --- | --- | --- | --- |
| 12.0428 | 14.6049 | 0.0082 | 1.2119e+10 | 20 |

**Adduct Signal Fractions**

- **Unmodified**: 1.2119e+10

#### HAIHVSGTN(N)GTKRF Hex 6 HexNAc 7

**Mass:**  3917.6855

(4.409e-06)

**Charge States:** 3, 4, 5

**Start Position**: 66

**End Position**: 80

| Profile Score | Tandem Score | TDA q-value | Volume | Spectrum Matches |
| --- | --- | --- | --- | --- |
| 17.4336 | 14.5389 | 0.0082 | 4.9414e+08 | 1 |

**Adduct Signal Fractions**

- **Unmodified**: 4.9414e+08

#### SALEPLVDLPIGIN(N)ITRF Fuc 1 Hex 3 HexNAc 4

**Mass:**  3411.6422

(-2.509e-07)

**Charge States:** 3, 4

**Start Position**: 221

**End Position**: 239

| Profile Score | Tandem Score | TDA q-value | Volume | Spectrum Matches |
| --- | --- | --- | --- | --- |
| 14.9767 | 14.4553 | 0.0082 | 6.5050e+09 | 8 |

**Adduct Signal Fractions**

- **Unmodified**: 6.5050e+09

#### HAIHVSGTN(N)GTKRF @sulfate 1 Hex 5 HexNAc 2

**Mass:**  2820.1784

(6.371e-07)

**Charge States:** 4

**Start Position**: 66

**End Position**: 80

| Profile Score | Tandem Score | TDA q-value | Volume | Spectrum Matches |
| --- | --- | --- | --- | --- |
| 12.6841 | 14.3978 | 0.0082 | 1.9834e+07 | 1 |

**Adduct Signal Fractions**

- **Unmodified**: 1.9834e+07

#### HKNN(N)KSWMESEF Fuc 2 Hex 4 HexNAc 5 Neu5Ac 1

**Mass:**  3782.4994

(-5.271e-07)

**Charge States:** 3, 4

**Start Position**: 146

**End Position**: 158

| Profile Score | Tandem Score | TDA q-value | Volume | Spectrum Matches |
| --- | --- | --- | --- | --- |
| 16.0266 | 14.1092 | 0.0096 | 2.0387e+09 | 4 |

**Adduct Signal Fractions**

- **Unmodified**: 2.0387e+09

#### HKNN(N)KSWMESEF Fuc 1 Hex 4 HexNAc 4 Neu5Ac 1

**Mass:**  3433.3674

(1.325e-06)

**Charge States:** 3, 4

**Start Position**: 146

**End Position**: 158

| Profile Score | Tandem Score | TDA q-value | Volume | Spectrum Matches |
| --- | --- | --- | --- | --- |
| 16.1254 | 14.0465 | 0.0096 | 4.3335e+09 | 7 |

**Adduct Signal Fractions**

- **Unmodified**: 4.3335e+09

#### FHAIHVSGTN(N)GTKRF Fuc 1 Hex 4 HexNAc 5

**Mass:**  3480.5281

(-2.625e-06)

**Charge States:** 4, 5

**Start Position**: 65

**End Position**: 80

| Profile Score | Tandem Score | TDA q-value | Volume | Spectrum Matches |
| --- | --- | --- | --- | --- |
| 11.0292 | 14.0330 | 0.0096 | 4.7499e+08 | 12 |

**Adduct Signal Fractions**

- **Unmodified**: 4.7499e+08

#### SALEPLVDLPIGIN(N)ITRF Hex 5 HexNAc 3

**Mass:**  3386.6124

(3.527e-07)

**Charge States:** 3, 4

**Start Position**: 221

**End Position**: 239

| Profile Score | Tandem Score | TDA q-value | Volume | Spectrum Matches |
| --- | --- | --- | --- | --- |
| 18.0251 | 13.8428 | 0.0096 | 6.4796e+09 | 2 |

**Adduct Signal Fractions**

- **Unmodified**: 6.4796e+09

#### HKNN(N)KSWMESEF Fuc 1 Hex 4 HexNAc 4

**Mass:**  3142.2714

(1.512e-06)

**Charge States:** 2, 3, 4

**Start Position**: 146

**End Position**: 158

| Profile Score | Tandem Score | TDA q-value | Volume | Spectrum Matches |
| --- | --- | --- | --- | --- |
| 14.1736 | 13.8310 | 0.0096 | 9.1518e+09 | 5 |

**Adduct Signal Fractions**

- **Unmodified**: 9.1518e+09

#### YHKNN(N)KSWMESEF Fuc 2 Hex 5 HexNAc 4

**Mass:**  3613.4451

(3.926e-07)

**Charge States:** 3, 4

**Start Position**: 145

**End Position**: 158

| Profile Score | Tandem Score | TDA q-value | Volume | Spectrum Matches |
| --- | --- | --- | --- | --- |
| 16.8560 | 13.7786 | 0.0101 | 1.6796e+09 | 4 |

**Adduct Signal Fractions**

- **Unmodified**: 1.6796e+09

#### SALEPLVDLPIGIN(N)ITRF Hex 3 HexNAc 3

**Mass:**  3062.5093

(1.175e-06)

**Charge States:** 2, 3, 4

**Start Position**: 221

**End Position**: 239

| Profile Score | Tandem Score | TDA q-value | Volume | Spectrum Matches |
| --- | --- | --- | --- | --- |
| 18.3093 | 13.7701 | 0.0101 | 1.5314e+09 | 2 |

**Adduct Signal Fractions**

- **Unmodified**: 1.5314e+09

#### VPAQEKN(N)FTTAPAIC(C)HDGKAHFPREGVF Fuc 1 Hex 4 HexNAc 3

**Mass:**  4527.0481

(-4.301e-07)

**Charge States:** 3, 4, 5, 6

**Start Position**: 1068

**End Position**: 1096

| Profile Score | Tandem Score | TDA q-value | Volume | Spectrum Matches |
| --- | --- | --- | --- | --- |
| 16.2192 | 13.7119 | 0.0101 | 2.2137e+10 | 10 |

**Adduct Signal Fractions**

- **Unmodified**: 2.2137e+10

#### HAIHVSGTN(N)GTKRF Fuc 2 Hex 6 HexNAc 6

**Mass:**  4006.7019

(8.227e-07)

**Charge States:** 4, 5

**Start Position**: 66

**End Position**: 80

| Profile Score | Tandem Score | TDA q-value | Volume | Spectrum Matches |
| --- | --- | --- | --- | --- |
| 10.4194 | 13.6588 | 0.0101 | 2.5390e+08 | 9 |

**Adduct Signal Fractions**

- **Unmodified**: 2.5390e+08

#### HKNN(N)KSWMESEF Fuc 2 Hex 5 HexNAc 4

**Mass:**  3450.3806

(1.939e-07)

**Charge States:** 3, 4

**Start Position**: 146

**End Position**: 158

| Profile Score | Tandem Score | TDA q-value | Volume | Spectrum Matches |
| --- | --- | --- | --- | --- |
| 16.0773 | 13.5706 | 0.0101 | 1.2476e+09 | 4 |

**Adduct Signal Fractions**

- **Unmodified**: 1.2476e+09

#### FHAIHVSGTN(N)GTKRF @sulfate 1 Fuc 1 Hex 4 HexNAc 5 Neu5Ac 1

**Mass:**  3851.5799

(-5.021e-07)

**Charge States:** 3, 4

**Start Position**: 65

**End Position**: 80

| Profile Score | Tandem Score | TDA q-value | Volume | Spectrum Matches |
| --- | --- | --- | --- | --- |
| 15.5906 | 13.5378 | 0.0101 | 7.3445e+08 | 4 |

**Adduct Signal Fractions**

- **Unmodified**: 7.3445e+08

#### GGFN(N)FSQILPDPSKPSKRSF Hex 6 HexNAc 2

**Mass:**  3586.6111

(6.786e-07)

**Charge States:** 3, 4

**Start Position**: 798

**End Position**: 818

| Profile Score | Tandem Score | TDA q-value | Volume | Spectrum Matches |
| --- | --- | --- | --- | --- |
| 11.8466 | 13.4484 | 0.0101 | 3.6226e+08 | 3 |

**Adduct Signal Fractions**

- **Unmodified**: 3.6226e+08

#### SN(N)NSIAIPTN(N)F Hex 12 HexNAc 5

**Mass:**  4136.6085

(-3.200e-07)

**Charge States:** 3, 4

**Start Position**: 708

**End Position**: 719

| Profile Score | Tandem Score | TDA q-value | Volume | Spectrum Matches |
| --- | --- | --- | --- | --- |
| 15.6434 | 13.3070 | 0.0101 | 2.1157e+09 | 4 |

**Adduct Signal Fractions**

- **Unmodified**: 2.1157e+09

#### HKNN(N)KSWMESEF Fuc 1 Hex 3 HexNAc 6

**Mass:**  3386.3769

(5.192e-07)

**Charge States:** 3, 4

**Start Position**: 146

**End Position**: 158

| Profile Score | Tandem Score | TDA q-value | Volume | Spectrum Matches |
| --- | --- | --- | --- | --- |
| 11.9518 | 13.2219 | 0.0101 | 4.4316e+09 | 9 |

**Adduct Signal Fractions**

- **Unmodified**: 4.4316e+09

#### GGFN(N)FSQILPDPSKPSKRSF Hex 5 HexNAc 2

**Mass:**  3424.5626

(1.857e-06)

**Charge States:** 3, 4

**Start Position**: 798

**End Position**: 818

| Profile Score | Tandem Score | TDA q-value | Volume | Spectrum Matches |
| --- | --- | --- | --- | --- |
| 14.7460 | 13.1918 | 0.0101 | 1.6971e+10 | 12 |

**Adduct Signal Fractions**

- **Unmodified**: 1.6971e+10

#### VPAQEKN(N)FTTAPAIC(C)HDGKAHFPREGVF Fuc 1 Hex 3 HexNAc 5

**Mass:**  4771.1512

(8.171e-07)

**Charge States:** 3, 4, 5, 6

**Start Position**: 1068

**End Position**: 1096

| Profile Score | Tandem Score | TDA q-value | Volume | Spectrum Matches |
| --- | --- | --- | --- | --- |
| 13.4486 | 13.1256 | 0.0101 | 5.0014e+10 | 21 |

**Adduct Signal Fractions**

- **Unmodified**: 5.0014e+10

#### VDLPIGIN(N)ITRF Hex 4 HexNAc 6

**Mass:**  3223.4861

(6.834e-06)

**Start Position**: 227

**End Position**: 239

| Profile Score | Tandem Score | TDA q-value | Volume | Spectrum Matches |
| --- | --- | --- | --- | --- |
| 0.0000 | 13.1062 | 0.0101 | 0.0000e+00 | 1 |

Adduct:

No Chromatogram Found

#### RVQPTESIVRFPN(N)ITNLC(C)PF Fuc 1 Hex 5 HexNAc 5

**Mass:**  4358.9628

(2.103e-08)

**Charge States:** 3, 4, 5

**Start Position**: 319

**End Position**: 339

| Profile Score | Tandem Score | TDA q-value | Volume | Spectrum Matches |
| --- | --- | --- | --- | --- |
| 15.2684 | 13.1034 | 0.0101 | 6.8848e+09 | 7 |

**Adduct Signal Fractions**

- **Unmodified**: 6.8848e+09

#### RVQPTESIVRFPN(N)ITNLC(C)PFGEVF Hex 5 HexNAc 2

**Mass:**  4035.8658

(4.279e-07)

**Charge States:** 3, 4

**Start Position**: 319

**End Position**: 343

| Profile Score | Tandem Score | TDA q-value | Volume | Spectrum Matches |
| --- | --- | --- | --- | --- |
| 17.9198 | 13.0847 | 0.0101 | 3.0461e+09 | 4 |

**Adduct Signal Fractions**

- **Unmodified**: 3.0461e+09

#### HKNN(N)KSWMESEF Fuc 1 Hex 4 HexNAc 5 Neu5Ac 1

**Mass:**  3636.4502

(1.600e-06)

**Charge States:** 3, 4

**Start Position**: 146

**End Position**: 158

| Profile Score | Tandem Score | TDA q-value | Volume | Spectrum Matches |
| --- | --- | --- | --- | --- |
| 13.3785 | 13.0717 | 0.0101 | 8.8881e+09 | 17 |

**Adduct Signal Fractions**

- **Unmodified**: 8.8881e+09

#### N(N)FSQILPDPSKPSKRSF Hex 3 HexNAc 4

**Mass:**  3245.4961

(-3.413e-07)

**Charge States:** 3, 4

**Start Position**: 801

**End Position**: 818

| Profile Score | Tandem Score | TDA q-value | Volume | Spectrum Matches |
| --- | --- | --- | --- | --- |
| 12.5950 | 12.9956 | 0.0101 | 1.5527e+09 | 6 |

**Adduct Signal Fractions**

- **Unmodified**: 1.5527e+09

#### VPAQEKN(N)FTTAPAIC(C)HDGKAHFPREGVF Fuc 1 Hex 5 HexNAc 3

**Mass:**  4689.1009

(1.303e-06)

**Charge States:** 3, 4, 5, 6

**Start Position**: 1068

**End Position**: 1096

| Profile Score | Tandem Score | TDA q-value | Volume | Spectrum Matches |
| --- | --- | --- | --- | --- |
| 16.5228 | 12.9356 | 0.0101 | 1.3571e+10 | 13 |

**Adduct Signal Fractions**

- **Unmodified**: 1.3571e+10

#### GGFN(N)FSQILPDPSKPSKRSF Hex 7 HexNAc 2

**Mass:**  3748.6596

(1.005e-07)

**Charge States:** 3, 4

**Start Position**: 798

**End Position**: 818

| Profile Score | Tandem Score | TDA q-value | Volume | Spectrum Matches |
| --- | --- | --- | --- | --- |
| 13.7248 | 12.9008 | 0.0101 | 2.9964e+09 | 7 |

**Adduct Signal Fractions**

- **Unmodified**: 2.9964e+09

#### RVQPTESIVRFPN(N)ITNLC(C)PFGEVF Fuc 1 Hex 5 HexNAc 4 Neu5Ac 1

**Mass:**  4879.1738

(4.579e-07)

**Charge States:** 4, 5, 6

**Start Position**: 319

**End Position**: 343

| Profile Score | Tandem Score | TDA q-value | Volume | Spectrum Matches |
| --- | --- | --- | --- | --- |
| 18.0053 | 12.8044 | 0.0101 | 5.8093e+10 | 12 |

**Adduct Signal Fractions**

- **Unmodified**: 5.8093e+10

#### LIVNN(N)ATNVVIKVC(C)EF Fuc 1 Hex 4 HexNAc 3

**Mass:**  3235.4934

(-2.551e-07)

**Charge States:** 3

**Start Position**: 118

**End Position**: 134

| Profile Score | Tandem Score | TDA q-value | Volume | Spectrum Matches |
| --- | --- | --- | --- | --- |
| 16.5735 | 12.7766 | 0.0101 | 2.3243e+09 | 4 |

**Adduct Signal Fractions**

- **Unmodified**: 2.3243e+09

#### RVQPTESIVRFPN(N)ITNLC(C)PF Fuc 1 Hex 4 HexNAc 5

**Mass:**  4196.9080

(1.166e-06)

**Charge States:** 3, 4, 5

**Start Position**: 319

**End Position**: 339

| Profile Score | Tandem Score | TDA q-value | Volume | Spectrum Matches |
| --- | --- | --- | --- | --- |
| 13.1999 | 12.6278 | 0.0101 | 1.9350e+10 | 18 |

**Adduct Signal Fractions**

- **Unmodified**: 1.9350e+10

#### QTRAGC(C)LIGAEHVN(N)NSYEC(C)DIPIGAGIC(C)ASY Fuc 1 Hex 3 HexNAc 4

**Mass:**  4840.0697

(7.996e-07)

**Charge States:** 4, 5

**Start Position**: 644

**End Position**: 675

| Profile Score | Tandem Score | TDA q-value | Volume | Spectrum Matches |
| --- | --- | --- | --- | --- |
| 16.8474 | 12.5647 | 0.0101 | 4.2110e+10 | 9 |

**Adduct Signal Fractions**

- **Unmodified**: 4.2110e+10

#### YHKNN(N)KSWMESEF Fuc 2 Hex 4 HexNAc 4

**Mass:**  3451.3905

(-2.632e-07)

**Charge States:** 3, 4

**Start Position**: 145

**End Position**: 158

| Profile Score | Tandem Score | TDA q-value | Volume | Spectrum Matches |
| --- | --- | --- | --- | --- |
| 15.2020 | 12.4408 | 0.0113 | 4.9648e+09 | 7 |

**Adduct Signal Fractions**

- **Unmodified**: 4.9648e+09

#### N(N)FSQILPDPSKPSKRSF Hex 5 HexNAc 3 Neu5Ac 1

**Mass:**  3657.6201

(2.649e-07)

**Charge States:** 3, 4

**Start Position**: 801

**End Position**: 818

| Profile Score | Tandem Score | TDA q-value | Volume | Spectrum Matches |
| --- | --- | --- | --- | --- |
| -2.9195 | 12.3337 | 0.0113 | 1.3350e+10 | 12 |

**Adduct Signal Fractions**

- **Unmodified**: 1.3350e+10

#### HAIHVSGTN(N)GTKRF Fuc 3 Hex 5 HexNAc 6 Neu5Ac 1

**Mass:**  4281.8062

(8.062e-07)

**Charge States:** 4, 5

**Start Position**: 66

**End Position**: 80

| Profile Score | Tandem Score | TDA q-value | Volume | Spectrum Matches |
| --- | --- | --- | --- | --- |
| 11.6483 | 12.3318 | 0.0113 | 1.0115e+09 | 16 |

**Adduct Signal Fractions**

- **Unmodified**: 1.0115e+09

#### FHAIHVSGTN(N)GTKRF @sulfate 1 Hex 4 HexNAc 5 Neu5Ac 1

**Mass:**  3705.5332

(2.074e-06)

**Charge States:** 3, 4

**Start Position**: 65

**End Position**: 80

| Profile Score | Tandem Score | TDA q-value | Volume | Spectrum Matches |
| --- | --- | --- | --- | --- |
| 14.4623 | 12.3106 | 0.0113 | 6.0617e+08 | 2 |

**Adduct Signal Fractions**

- **Unmodified**: 6.0617e+08

#### HKNN(N)KSWMESEF Fuc 2 Hex 4 HexNAc 5

**Mass:**  3491.4058

(-2.535e-07)

**Charge States:** 3, 4

**Start Position**: 146

**End Position**: 158

| Profile Score | Tandem Score | TDA q-value | Volume | Spectrum Matches |
| --- | --- | --- | --- | --- |
| 16.6503 | 12.0992 | 0.0124 | 9.4588e+09 | 6 |

**Adduct Signal Fractions**

- **Unmodified**: 9.4588e+09

#### HAIHVSGTN(N)GTKRF @sulfate 1 Fuc 3 Hex 6 HexNAc 7 Neu5Ac 2

**Mass:**  5017.9696

(-3.993e-06)

**Charge States:** 4

**Start Position**: 66

**End Position**: 80

| Profile Score | Tandem Score | TDA q-value | Volume | Spectrum Matches |
| --- | --- | --- | --- | --- |
| -4.5043 | 11.9380 | 0.0124 | 6.6872e+07 | 1 |

**Adduct Signal Fractions**

- **Unmodified**: 6.6872e+07

#### IVNN(N)ATNVVIKVC(C)EF Hex 6 HexNAc 3 Neu5Ac 1

**Mass:**  3591.5532

(2.357e-07)

**Charge States:** 3, 4

**Start Position**: 119

**End Position**: 134

| Profile Score | Tandem Score | TDA q-value | Volume | Spectrum Matches |
| --- | --- | --- | --- | --- |
| 15.9367 | 11.8858 | 0.0124 | 5.8438e+09 | 8 |

**Adduct Signal Fractions**

- **Unmodified**: 5.8438e+09

#### RVQPTESIVRFPN(N)ITNLC(C)PF Fuc 1 Hex 6 HexNAc 5

**Mass:**  4521.0132

(-5.054e-07)

**Charge States:** 3, 4, 5

**Start Position**: 319

**End Position**: 339

| Profile Score | Tandem Score | TDA q-value | Volume | Spectrum Matches |
| --- | --- | --- | --- | --- |
| 16.7594 | 11.7815 | 0.0131 | 2.1280e+09 | 5 |

**Adduct Signal Fractions**

- **Unmodified**: 2.1280e+09

#### HAIHVSGTN(N)GTKRF Hex 8 HexNAc 2

**Mass:**  3226.3782

(2.139e-07)

**Charge States:** 4

**Start Position**: 66

**End Position**: 80

| Profile Score | Tandem Score | TDA q-value | Volume | Spectrum Matches |
| --- | --- | --- | --- | --- |
| 15.0042 | 11.7734 | 0.0131 | 2.9411e+07 | 2 |

**Adduct Signal Fractions**

- **Unmodified**: 2.9411e+07

#### IVNN(N)ATNVVIKVC(C)EF Fuc 1 Hex 5 HexNAc 3

**Mass:**  3284.4660

(1.049e-06)

**Charge States:** 3

**Start Position**: 119

**End Position**: 134

| Profile Score | Tandem Score | TDA q-value | Volume | Spectrum Matches |
| --- | --- | --- | --- | --- |
| 14.0644 | 11.6237 | 0.0131 | 4.2947e+09 | 5 |

**Adduct Signal Fractions**

- **Unmodified**: 4.2947e+09

#### IVNN(N)ATNVVIKVC(C)EF Hex 5 HexNAc 3 Neu5Ac 1

**Mass:**  3429.4995

(-2.017e-07)

**Charge States:** 3, 4

**Start Position**: 119

**End Position**: 134

| Profile Score | Tandem Score | TDA q-value | Volume | Spectrum Matches |
| --- | --- | --- | --- | --- |
| 14.9598 | 11.5220 | 0.0131 | 8.1678e+09 | 10 |

**Adduct Signal Fractions**

- **Unmodified**: 8.1678e+09

#### IVNN(N)ATNVVIKVC(C)EF Fuc 1 Hex 4 HexNAc 4 Neu5Ac 1

**Mass:**  3616.5825

(1.723e-07)

**Charge States:** 3, 4

**Start Position**: 119

**End Position**: 134

| Profile Score | Tandem Score | TDA q-value | Volume | Spectrum Matches |
| --- | --- | --- | --- | --- |
| 15.9438 | 11.5206 | 0.0131 | 1.1574e+10 | 11 |

**Adduct Signal Fractions**

- **Unmodified**: 1.1574e+10

#### HKNN(N)KSWMESEF Fuc 1 Hex 5 HexNAc 5

**Mass:**  3507.4047

(1.658e-06)

**Charge States:** 3, 4

**Start Position**: 146

**End Position**: 158

| Profile Score | Tandem Score | TDA q-value | Volume | Spectrum Matches |
| --- | --- | --- | --- | --- |
| 15.8192 | 11.4485 | 0.0131 | 5.4468e+09 | 10 |

**Adduct Signal Fractions**

- **Unmodified**: 5.4468e+09

#### HKNN(N)KSWMESEF Fuc 2 Hex 4 HexNAc 4

**Mass:**  3288.3271

(1.400e-08)

**Charge States:** 3, 4

**Start Position**: 146

**End Position**: 158

| Profile Score | Tandem Score | TDA q-value | Volume | Spectrum Matches |
| --- | --- | --- | --- | --- |
| 15.1847 | 11.4195 | 0.0131 | 2.0374e+09 | 6 |

**Adduct Signal Fractions**

- **Unmodified**: 2.0374e+09

#### VDLPIGIN(N)ITRF Fuc 1 Hex 4 HexNAc 8

**Mass:**  3776.6740

(2.631e-04)

**Start Position**: 227

**End Position**: 239

| Profile Score | Tandem Score | TDA q-value | Volume | Spectrum Matches |
| --- | --- | --- | --- | --- |
| 0.0000 | 11.4071 | 0.0131 | 0.0000e+00 | 1 |

Adduct:

No Chromatogram Found

#### VPAQEKN(N)FTTAPAIC(C)HDGKAHFPREGVF Fuc 1 Hex 5 HexNAc 4

**Mass:**  4892.1708

(-1.005e-06)

**Charge States:** 4, 5, 6

**Start Position**: 1068

**End Position**: 1096

| Profile Score | Tandem Score | TDA q-value | Volume | Spectrum Matches |
| --- | --- | --- | --- | --- |
| 16.0919 | 11.2769 | 0.0131 | 2.0598e+10 | 9 |

**Adduct Signal Fractions**

- **Unmodified**: 2.0598e+10

#### IVNN(N)ATNVVIKVC(C)EF Fuc 1 Hex 4 HexNAc 3 Neu5Ac 1

**Mass:**  3413.5377

(-3.222e-07)

**Charge States:** 3, 4

**Start Position**: 119

**End Position**: 134

| Profile Score | Tandem Score | TDA q-value | Volume | Spectrum Matches |
| --- | --- | --- | --- | --- |
| 13.9140 | 11.2697 | 0.0131 | 3.5686e+09 | 9 |

**Adduct Signal Fractions**

- **Unmodified**: 3.5686e+09

#### IVNN(N)ATNVVIKVC(C)EF Fuc 1 Hex 3 HexNAc 4

**Mass:**  3163.4377

(4.316e-07)

**Charge States:** 2, 3

**Start Position**: 119

**End Position**: 134

| Profile Score | Tandem Score | TDA q-value | Volume | Spectrum Matches |
| --- | --- | --- | --- | --- |
| 16.3431 | 11.1591 | 0.0131 | 1.2064e+10 | 12 |

**Adduct Signal Fractions**

- **Unmodified**: 1.2064e+10

#### VPAQEKN(N)FTTAPAIC(C)HDGKAHFPREGVF Fuc 1 Hex 4 HexNAc 5

**Mass:**  4933.1992

(-4.912e-07)

**Charge States:** 4, 5, 6

**Start Position**: 1068

**End Position**: 1096

| Profile Score | Tandem Score | TDA q-value | Volume | Spectrum Matches |
| --- | --- | --- | --- | --- |
| 13.8039 | 10.9646 | 0.0140 | 2.1375e+10 | 15 |

**Adduct Signal Fractions**

- **Unmodified**: 2.1375e+10

#### VPAQEKN(N)FTTAPAIC(C)HDGKAHFPREGVF Fuc 1 Hex 3 HexNAc 4

**Mass:**  4568.0654

(7.683e-07)

**Charge States:** 3, 4, 5, 6

**Start Position**: 1068

**End Position**: 1096

| Profile Score | Tandem Score | TDA q-value | Volume | Spectrum Matches |
| --- | --- | --- | --- | --- |
| 18.0910 | 10.9530 | 0.0140 | 6.3364e+10 | 12 |

**Adduct Signal Fractions**

- **Unmodified**: 6.3364e+10

#### HAIHVSGTN(N)GTKRF Fuc 2 Hex 5 HexNAc 5 Neu5Ac 1

**Mass:**  3932.6685

(5.864e-07)

**Charge States:** 4, 5

**Start Position**: 66

**End Position**: 80

| Profile Score | Tandem Score | TDA q-value | Volume | Spectrum Matches |
| --- | --- | --- | --- | --- |
| 11.0108 | 10.8225 | 0.0140 | 8.8611e+08 | 21 |

**Adduct Signal Fractions**

- **Unmodified**: 8.8611e+08

#### SN(N)NSIAIPTN(N)F Fuc 1 Hex 8 HexNAc 7

**Mass:**  4040.6108

(-6.877e-07)

**Charge States:** 3

**Start Position**: 708

**End Position**: 719

| Profile Score | Tandem Score | TDA q-value | Volume | Spectrum Matches |
| --- | --- | --- | --- | --- |
| 11.6930 | 10.6735 | 0.0131 | 4.1770e+08 | 3 |

**Adduct Signal Fractions**

- **Unmodified**: 4.1770e+08

#### HAIHVSGTN(N)GTKRF Fuc 2 Hex 5 HexNAc 7

**Mass:**  4047.7314

(-9.028e-07)

**Charge States:** 4, 5

**Start Position**: 66

**End Position**: 80

| Profile Score | Tandem Score | TDA q-value | Volume | Spectrum Matches |
| --- | --- | --- | --- | --- |
| 11.8060 | 10.6705 | 0.0140 | 1.2594e+08 | 9 |

**Adduct Signal Fractions**

- **Unmodified**: 1.2594e+08

#### SN(N)NSIAIPTN(N)F Hex 9 HexNAc 5

**Mass:**  3650.4545

(1.324e-06)

**Charge States:** 3

**Start Position**: 708

**End Position**: 719

| Profile Score | Tandem Score | TDA q-value | Volume | Spectrum Matches |
| --- | --- | --- | --- | --- |
| 16.4879 | 10.5941 | 0.0131 | 9.0369e+08 | 1 |

**Adduct Signal Fractions**

- **Unmodified**: 9.0369e+08

#### N(N)FSQILPDPSKPSKRSF Fuc 1 Hex 5 HexNAc 4 Neu5Ac 1

**Mass:**  4006.7632

(1.872e-06)

**Charge States:** 3, 4, 5

**Start Position**: 801

**End Position**: 818

| Profile Score | Tandem Score | TDA q-value | Volume | Spectrum Matches |
| --- | --- | --- | --- | --- |
| -2.9204 | 10.5851 | 0.0140 | 1.9560e+09 | 6 |

**Adduct Signal Fractions**

- **Unmodified**: 1.9560e+09

#### IVNN(N)ATNVVIKVC(C)EF Hex 7 HexNAc 2

**Mass:**  3259.4320

(9.985e-07)

**Charge States:** 3

**Start Position**: 119

**End Position**: 134

| Profile Score | Tandem Score | TDA q-value | Volume | Spectrum Matches |
| --- | --- | --- | --- | --- |
| 17.5705 | 10.5846 | 0.0140 | 2.2997e+09 | 3 |

**Adduct Signal Fractions**

- **Unmodified**: 2.2997e+09

#### QTRAGC(C)LIGAEHVN(N)NSYEC(C)DIPIGAGIC(C)ASY Fuc 1 Hex 4 HexNAc 4 Neu5Ac 1

**Mass:**  5293.2140

(-7.497e-07)

**Charge States:** 4, 5, 6

**Start Position**: 644

**End Position**: 675

| Profile Score | Tandem Score | TDA q-value | Volume | Spectrum Matches |
| --- | --- | --- | --- | --- |
| 15.7887 | 10.5664 | 0.0140 | 1.4308e+10 | 10 |

**Adduct Signal Fractions**

- **Unmodified**: 1.4308e+10

#### FHAIHVSGTN(N)GTKRF @sulfate 1 Fuc 1 Hex 3 HexNAc 5

**Mass:**  3398.4350

(1.016e-07)

**Charge States:** 4

**Start Position**: 65

**End Position**: 80

| Profile Score | Tandem Score | TDA q-value | Volume | Spectrum Matches |
| --- | --- | --- | --- | --- |
| 13.0074 | 10.4649 | 0.0140 | 3.6288e+08 | 2 |

**Adduct Signal Fractions**

- **Unmodified**: 3.6288e+08

#### SALEPLVDLPIGIN(N)ITRF Fuc 1 Hex 3 HexNAc 5

**Mass:**  3614.7201

(3.621e-07)

**Charge States:** 3, 4

**Start Position**: 221

**End Position**: 239

| Profile Score | Tandem Score | TDA q-value | Volume | Spectrum Matches |
| --- | --- | --- | --- | --- |
| 14.7221 | 10.3785 | 0.0164 | 2.0380e+09 | 5 |

**Adduct Signal Fractions**

- **Unmodified**: 2.0380e+09

#### IVNN(N)ATNVVIKVC(C)EF Hex 6 HexNAc 3

**Mass:**  3300.4603

(8.643e-07)

**Charge States:** 3, 4

**Start Position**: 119

**End Position**: 134

| Profile Score | Tandem Score | TDA q-value | Volume | Spectrum Matches |
| --- | --- | --- | --- | --- |
| 17.2769 | 10.2641 | 0.0164 | 5.2842e+09 | 3 |

**Adduct Signal Fractions**

- **Unmodified**: 5.2842e+09

#### HAIHVSGTN(N)GTKRF @sulfate 1 Fuc 2 Hex 4 HexNAc 5 Neu5Ac 1

**Mass:**  3850.5730

(3.109e-07)

**Charge States:** 4

**Start Position**: 66

**End Position**: 80

| Profile Score | Tandem Score | TDA q-value | Volume | Spectrum Matches |
| --- | --- | --- | --- | --- |
| 9.9955 | 10.2086 | 0.0168 | 2.1220e+08 | 3 |

**Adduct Signal Fractions**

- **Unmodified**: 2.1220e+08

#### IVNN(N)ATNVVIKVC(C)EF Fuc 1 Hex 3 HexNAc 5

**Mass:**  3366.5179

(8.657e-07)

**Charge States:** 3, 4

**Start Position**: 119

**End Position**: 134

| Profile Score | Tandem Score | TDA q-value | Volume | Spectrum Matches |
| --- | --- | --- | --- | --- |
| 15.8192 | 10.1935 | 0.0168 | 3.8640e+09 | 4 |

**Adduct Signal Fractions**

- **Unmodified**: 3.8640e+09

#### VPAQEKN(N)FTTAPAIC(C)HDGKAHFPREGVF Fuc 1 Hex 5 HexNAc 5

**Mass:**  5095.2545

(-2.745e-08)

**Charge States:** 4, 5, 6

**Start Position**: 1068

**End Position**: 1096

| Profile Score | Tandem Score | TDA q-value | Volume | Spectrum Matches |
| --- | --- | --- | --- | --- |
| 16.2995 | 10.1352 | 0.0168 | 1.3322e+10 | 11 |

**Adduct Signal Fractions**

- **Unmodified**: 1.3322e+10

#### IVNN(N)ATNVVIKVC(C)EF Fuc 1 Hex 4 HexNAc 4

**Mass:**  3325.4883

(-1.153e-07)

**Charge States:** 3, 4

**Start Position**: 119

**End Position**: 134

| Profile Score | Tandem Score | TDA q-value | Volume | Spectrum Matches |
| --- | --- | --- | --- | --- |
| 15.0702 | 10.1347 | 0.0168 | 5.6178e+09 | 6 |

**Adduct Signal Fractions**

- **Unmodified**: 5.6178e+09

#### HAIHVSGTN(N)GTKRF Fuc 1 Hex 5 HexNAc 8 Neu5Ac 2

**Mass:**  4686.9491

(1.396e-06)

**Charge States:** 4, 5

**Start Position**: 66

**End Position**: 80

| Profile Score | Tandem Score | TDA q-value | Volume | Spectrum Matches |
| --- | --- | --- | --- | --- |
| 14.5902 | 10.0982 | 0.0168 | 4.6242e+08 | 6 |

**Adduct Signal Fractions**

- **Unmodified**: 4.6242e+08

#### IVNN(N)ATNVVIKVC(C)EF Fuc 1 Hex 5 HexNAc 3 Neu5Ac 1

**Mass:**  3575.5594

(2.275e-07)

**Charge States:** 3, 4

**Start Position**: 119

**End Position**: 134

| Profile Score | Tandem Score | TDA q-value | Volume | Spectrum Matches |
| --- | --- | --- | --- | --- |
| 15.8370 | 10.0907 | 0.0168 | 4.4884e+09 | 5 |

**Adduct Signal Fractions**

- **Unmodified**: 4.4884e+09

#### IVNN(N)ATNVVIKVC(C)EF Fuc 1 Hex 6 HexNAc 3

**Mass:**  3446.5171

(6.479e-07)

**Charge States:** 3

**Start Position**: 119

**End Position**: 134

| Profile Score | Tandem Score | TDA q-value | Volume | Spectrum Matches |
| --- | --- | --- | --- | --- |
| 15.6133 | 10.0454 | 0.0168 | 1.7175e+09 | 5 |

**Adduct Signal Fractions**

- **Unmodified**: 1.7175e+09

#### IVNN(N)ATNVVIKVC(C)EF Hex 5 HexNAc 2

**Mass:**  2935.3238

(1.081e-06)

**Charge States:** 2, 3

**Start Position**: 119

**End Position**: 134

| Profile Score | Tandem Score | TDA q-value | Volume | Spectrum Matches |
| --- | --- | --- | --- | --- |
| 17.3091 | 10.0059 | 0.0168 | 6.0765e+10 | 19 |

**Adduct Signal Fractions**

- **Unmodified**: 6.0765e+10

#### YHKNN(N)KSWMESEF Fuc 1 Hex 4 HexNAc 4

**Mass:**  3305.3351

(-2.563e-07)

**Charge States:** 3, 4

**Start Position**: 145

**End Position**: 158

| Profile Score | Tandem Score | TDA q-value | Volume | Spectrum Matches |
| --- | --- | --- | --- | --- |
| 14.6470 | 9.9496 | 0.0168 | 1.1685e+10 | 9 |

**Adduct Signal Fractions**

- **Unmodified**: 1.1685e+10

#### GGFN(N)FSQILPDPSKPSKRSF Fuc 1 Hex 3 HexNAc 4

**Mass:**  3652.6633

(1.046e-06)

**Charge States:** 3, 4

**Start Position**: 798

**End Position**: 818

| Profile Score | Tandem Score | TDA q-value | Volume | Spectrum Matches |
| --- | --- | --- | --- | --- |
| 13.4885 | 9.6690 | 0.0168 | 2.2477e+09 | 6 |

**Adduct Signal Fractions**

- **Unmodified**: 2.2477e+09

#### SN(N)NSIAIPTN(N)F Fuc 1 Hex 10 HexNAc 5

**Mass:**  3960.5496

(5.079e-04)

**Charge States:** 3

**Start Position**: 708

**End Position**: 719

| Profile Score | Tandem Score | TDA q-value | Volume | Spectrum Matches |
| --- | --- | --- | --- | --- |
| -4.9461 | 9.6666 | 0.0168 | 5.7451e+07 | 1 |

**Adduct Signal Fractions**

- **Unmodified**: 5.7451e+07

#### QTRAGC(C)LIGAEHVN(N)NSYEC(C)DIPIGAGIC(C)ASY Fuc 1 Hex 4 HexNAc 5

**Mass:**  5205.2042

(6.466e-07)

**Charge States:** 4, 5

**Start Position**: 644

**End Position**: 675

| Profile Score | Tandem Score | TDA q-value | Volume | Spectrum Matches |
| --- | --- | --- | --- | --- |
| 15.8381 | 9.5619 | 0.0198 | 1.1256e+10 | 7 |

**Adduct Signal Fractions**

- **Unmodified**: 1.1256e+10

#### RVQPTESIVRFPN(N)ITNLC(C)PF Fuc 1 Hex 3 HexNAc 5

**Mass:**  4034.8558

(-8.298e-07)

**Charge States:** 3, 4, 5

**Start Position**: 319

**End Position**: 339

| Profile Score | Tandem Score | TDA q-value | Volume | Spectrum Matches |
| --- | --- | --- | --- | --- |
| 16.2832 | 9.5554 | 0.0198 | 4.7942e+10 | 16 |

**Adduct Signal Fractions**

- **Unmodified**: 4.7942e+10

#### RVQPTESIVRFPN(N)ITNLC(C)PF Fuc 1 Hex 4 HexNAc 4

**Mass:**  3993.8277

(8.584e-07)

**Charge States:** 3, 4, 5

**Start Position**: 319

**End Position**: 339

| Profile Score | Tandem Score | TDA q-value | Volume | Spectrum Matches |
| --- | --- | --- | --- | --- |
| 13.9889 | 9.5422 | 0.0198 | 1.6375e+10 | 9 |

**Adduct Signal Fractions**

- **Unmodified**: 1.6375e+10

#### LIVNN(N)ATNVVIKVC(C)EF Hex 5 HexNAc 3

**Mass:**  3251.4883

(-2.803e-07)

**Charge States:** 3

**Start Position**: 118

**End Position**: 134

| Profile Score | Tandem Score | TDA q-value | Volume | Spectrum Matches |
| --- | --- | --- | --- | --- |
| 18.0160 | 9.5134 | 0.0198 | 5.5431e+09 | 3 |

**Adduct Signal Fractions**

- **Unmodified**: 5.5431e+09

#### GGFN(N)FSQILPDPSKPSKRSF Hex 6 HexNAc 3

**Mass:**  3789.6884

(8.054e-07)

**Charge States:** 4

**Start Position**: 798

**End Position**: 818

| Profile Score | Tandem Score | TDA q-value | Volume | Spectrum Matches |
| --- | --- | --- | --- | --- |
| 13.1142 | 9.4933 | 0.0198 | 1.1225e+09 | 5 |

**Adduct Signal Fractions**

- **Unmodified**: 1.1225e+09

#### QTRAGC(C)LIGAEHVN(N)NSYEC(C)DIPIGAGIC(C)ASY Hex 5 HexNAc 2

**Mass:**  4611.9583

(3.116e-07)

**Charge States:** 3, 4

**Start Position**: 644

**End Position**: 675

| Profile Score | Tandem Score | TDA q-value | Volume | Spectrum Matches |
| --- | --- | --- | --- | --- |
| 16.7801 | 9.4645 | 0.0198 | 3.6553e+09 | 3 |

**Adduct Signal Fractions**

- **Unmodified**: 3.6553e+09

#### LIVNN(N)ATNVVIKVC(C)EF Fuc 1 Hex 4 HexNAc 4

**Mass:**  3438.5764

(8.155e-07)

**Charge States:** 3, 4

**Start Position**: 118

**End Position**: 134

| Profile Score | Tandem Score | TDA q-value | Volume | Spectrum Matches |
| --- | --- | --- | --- | --- |
| 12.0869 | 9.4565 | 0.0198 | 1.4173e+09 | 7 |

**Adduct Signal Fractions**

- **Unmodified**: 1.4173e+09

#### RVQPTESIVRFPN(N)ITNLC(C)PF Fuc 1 Hex 3 HexNAc 4

**Mass:**  3831.7756

(-8.486e-07)

**Charge States:** 3, 4

**Start Position**: 319

**End Position**: 339

| Profile Score | Tandem Score | TDA q-value | Volume | Spectrum Matches |
| --- | --- | --- | --- | --- |
| 17.2652 | 9.4447 | 0.0198 | 4.4746e+10 | 12 |

**Adduct Signal Fractions**

- **Unmodified**: 4.4746e+10

#### RVQPTESIVRFPN(N)ITNLC(C)PF Fuc 1 Hex 5 HexNAc 4

**Mass:**  4155.8804

(2.612e-07)

**Charge States:** 3, 4, 5

**Start Position**: 319

**End Position**: 339

| Profile Score | Tandem Score | TDA q-value | Volume | Spectrum Matches |
| --- | --- | --- | --- | --- |
| 17.9268 | 9.4281 | 0.0198 | 7.5108e+09 | 9 |

**Adduct Signal Fractions**

- **Unmodified**: 7.5108e+09

#### EPLVDLPIGIN(N)ITRF Fuc 1 Hex 6 HexNAc 7

**Mass:**  4238.9472

(4.828e-04)

**Charge States:** 3, 4

**Start Position**: 224

**End Position**: 239

| Profile Score | Tandem Score | TDA q-value | Volume | Spectrum Matches |
| --- | --- | --- | --- | --- |
| -7.0919 | 9.4097 | 0.0198 | 2.3559e+08 | 2 |

**Adduct Signal Fractions**

- **Unmodified**: 2.3559e+08

#### RVQPTESIVRFPN(N)ITNLC(C)PFGEVF Fuc 1 Hex 3 HexNAc 5

**Mass:**  4467.0535

(-9.850e-07)

**Charge States:** 3, 4, 5

**Start Position**: 319

**End Position**: 343

| Profile Score | Tandem Score | TDA q-value | Volume | Spectrum Matches |
| --- | --- | --- | --- | --- |
| 17.6378 | 9.3961 | 0.0198 | 9.0333e+10 | 17 |

**Adduct Signal Fractions**

- **Unmodified**: 9.0333e+10

#### QTRAGC(C)LIGAEHVN(N)NSYEC(C)DIPIGAGIC(C)ASY Fuc 1 Hex 4 HexNAc 4

**Mass:**  5002.1232

(6.947e-07)

**Charge States:** 4, 5

**Start Position**: 644

**End Position**: 675

| Profile Score | Tandem Score | TDA q-value | Volume | Spectrum Matches |
| --- | --- | --- | --- | --- |
| 12.7855 | 9.3775 | 0.0198 | 7.9773e+09 | 6 |

**Adduct Signal Fractions**

- **Unmodified**: 7.9773e+09

#### N(N)FSQILPDPSKPSKRSF Fuc 1 Hex 3 HexNAc 5

**Mass:**  3594.6421

(1.821e-06)

**Charge States:** 3, 4

**Start Position**: 801

**End Position**: 818

| Profile Score | Tandem Score | TDA q-value | Volume | Spectrum Matches |
| --- | --- | --- | --- | --- |
| 12.9087 | 9.3415 | 0.0198 | 1.4147e+09 | 5 |

**Adduct Signal Fractions**

- **Unmodified**: 1.4147e+09

#### HAIHVSGTN(N)GTKRF Fuc 1 Hex 5 HexNAc 7 Neu5Ac 1

**Mass:**  4192.7679

(-7.400e-07)

**Charge States:** 4, 5

**Start Position**: 66

**End Position**: 80

| Profile Score | Tandem Score | TDA q-value | Volume | Spectrum Matches |
| --- | --- | --- | --- | --- |
| 10.5694 | 9.2670 | 0.0205 | 1.0469e+09 | 21 |

**Adduct Signal Fractions**

- **Unmodified**: 1.0469e+09

#### N(N)FSQILPDPSKPSKRSF Hex 6 HexNAc 3

**Mass:**  3528.5775

(-2.325e-07)

**Charge States:** 3, 4

**Start Position**: 801

**End Position**: 818

| Profile Score | Tandem Score | TDA q-value | Volume | Spectrum Matches |
| --- | --- | --- | --- | --- |
| -4.6806 | 9.2529 | 0.0205 | 2.9161e+09 | 5 |

**Adduct Signal Fractions**

- **Unmodified**: 2.9161e+09

#### FHAIHVSGTN(N)GTKRF Hex 4 HexNAc 6 Neu5Ac 1

**Mass:**  3828.6561

(3.390e-06)

**Charge States:** 4, 5

**Start Position**: 65

**End Position**: 80

| Profile Score | Tandem Score | TDA q-value | Volume | Spectrum Matches |
| --- | --- | --- | --- | --- |
| 12.3020 | 9.2468 | 0.0205 | 4.4254e+08 | 2 |

**Adduct Signal Fractions**

- **Unmodified**: 4.4254e+08

#### RVQPTESIVRFPN(N)ITNLC(C)PFGEVF Fuc 1 Hex 4 HexNAc 4

**Mass:**  4426.0343

(1.064e-06)

**Charge States:** 3, 4, 5

**Start Position**: 319

**End Position**: 343

| Profile Score | Tandem Score | TDA q-value | Volume | Spectrum Matches |
| --- | --- | --- | --- | --- |
| 16.3946 | 9.2312 | 0.0205 | 3.6424e+10 | 11 |

**Adduct Signal Fractions**

- **Unmodified**: 3.6424e+10

#### HKNN(N)KSWMESEF Fuc 1 Hex 3 HexNAc 4

**Mass:**  2980.2154

(-7.208e-08)

**Charge States:** 2, 3, 4

**Start Position**: 146

**End Position**: 158

| Profile Score | Tandem Score | TDA q-value | Volume | Spectrum Matches |
| --- | --- | --- | --- | --- |
| 16.6147 | 9.2116 | 0.0205 | 3.7913e+10 | 17 |

**Adduct Signal Fractions**

- **Unmodified**: 3.7913e+10

#### SN(N)NSIAIPTN(N)F Fuc 1 Hex 9 HexNAc 6

**Mass:**  4000.5889

(2.512e-04)

**Start Position**: 708

**End Position**: 719

| Profile Score | Tandem Score | TDA q-value | Volume | Spectrum Matches |
| --- | --- | --- | --- | --- |
| 0.0000 | 9.2004 | 0.0205 | 0.0000e+00 | 2 |

Adduct:

No Chromatogram Found

#### N(N)FSQILPDPSKPSKRSF Hex 6 HexNAc 3 Neu5Ac 1

**Mass:**  3819.6791

(1.152e-06)

**Charge States:** 3, 4

**Start Position**: 801

**End Position**: 818

| Profile Score | Tandem Score | TDA q-value | Volume | Spectrum Matches |
| --- | --- | --- | --- | --- |
| -3.1734 | 9.1955 | 0.0205 | 6.1644e+09 | 7 |

**Adduct Signal Fractions**

- **Unmodified**: 6.1644e+09

#### GGFN(N)FSQILPDPSKPSKRSF Hex 5 HexNAc 3

**Mass:**  3627.6420

(1.926e-06)

**Charge States:** 3, 4

**Start Position**: 798

**End Position**: 818

| Profile Score | Tandem Score | TDA q-value | Volume | Spectrum Matches |
| --- | --- | --- | --- | --- |
| 12.2334 | 9.1517 | 0.0213 | 3.6271e+08 | 4 |

**Adduct Signal Fractions**

- **Unmodified**: 3.6271e+08

#### HAIHVSGTN(N)GTKRF @sulfate 1 Fuc 1 Hex 5 HexNAc 4 Neu5Ac 2

**Mass:**  3954.5831

(1.854e-07)

**Charge States:** 3, 4

**Start Position**: 66

**End Position**: 80

| Profile Score | Tandem Score | TDA q-value | Volume | Spectrum Matches |
| --- | --- | --- | --- | --- |
| 12.8000 | 9.0939 | 0.0213 | 1.4835e+08 | 3 |

**Adduct Signal Fractions**

- **Unmodified**: 1.4835e+08

#### SALEPLVDLPIGIN(N)ITRF Fuc 1 Hex 4 HexNAc 5

**Mass:**  3776.7753

(-1.227e-07)

**Charge States:** 3, 4

**Start Position**: 221

**End Position**: 239

| Profile Score | Tandem Score | TDA q-value | Volume | Spectrum Matches |
| --- | --- | --- | --- | --- |
| 15.6769 | 9.0719 | 0.0213 | 3.6825e+08 | 4 |

**Adduct Signal Fractions**

- **Unmodified**: 3.6825e+08

#### QTRAGC(C)LIGAEHVN(N)NSYEC(C)DIPIGAGIC(C)ASY Fuc 1 Hex 3 HexNAc 5

**Mass:**  5043.1472

(-1.743e-07)

**Charge States:** 4, 5

**Start Position**: 644

**End Position**: 675

| Profile Score | Tandem Score | TDA q-value | Volume | Spectrum Matches |
| --- | --- | --- | --- | --- |
| 13.2875 | 9.0530 | 0.0213 | 3.3379e+10 | 12 |

**Adduct Signal Fractions**

- **Unmodified**: 3.3379e+10

#### RVQPTESIVRFPN(N)ITNLC(C)PF Fuc 1 Hex 4 HexNAc 4 Neu5Ac 1

**Mass:**  4284.9221

(-7.221e-08)

**Charge States:** 3, 4, 5

**Start Position**: 319

**End Position**: 339

| Profile Score | Tandem Score | TDA q-value | Volume | Spectrum Matches |
| --- | --- | --- | --- | --- |
| 16.0981 | 8.9995 | 0.0218 | 5.2384e+10 | 16 |

**Adduct Signal Fractions**

- **Unmodified**: 5.2384e+10

#### VPAQEKN(N)FTTAPAIC(C)HDGKAHFPREGVF Fuc 1 Hex 3 HexNAc 6

**Mass:**  4974.2263

(-4.591e-07)

**Charge States:** 4, 5, 6

**Start Position**: 1068

**End Position**: 1096

| Profile Score | Tandem Score | TDA q-value | Volume | Spectrum Matches |
| --- | --- | --- | --- | --- |
| 16.3854 | 8.9576 | 0.0218 | 7.7381e+09 | 9 |

**Adduct Signal Fractions**

- **Unmodified**: 7.7381e+09

#### IVNN(N)ATNVVIKVC(C)EF Hex 4 HexNAc 3

**Mass:**  2976.3551

(1.047e-06)

**Charge States:** 2, 3

**Start Position**: 119

**End Position**: 134

| Profile Score | Tandem Score | TDA q-value | Volume | Spectrum Matches |
| --- | --- | --- | --- | --- |
| 15.7868 | 8.9373 | 0.0218 | 2.3043e+10 | 10 |

**Adduct Signal Fractions**

- **Unmodified**: 2.3043e+10

#### VPAQEKN(N)FTTAPAIC(C)HDGKAHFPREGVF Fuc 1 Hex 7 HexNAc 6 Neu5Ac 1

**Mass:**  5913.5409

(1.130e-06)

**Charge States:** 4, 5, 6

**Start Position**: 1068

**End Position**: 1096

| Profile Score | Tandem Score | TDA q-value | Volume | Spectrum Matches |
| --- | --- | --- | --- | --- |
| 16.1544 | 8.8739 | 0.0218 | 1.5909e+09 | 4 |

**Adduct Signal Fractions**

- **Unmodified**: 1.5909e+09

#### RVQPTESIVRFPN(N)ITNLC(C)PFGEVF Fuc 1 Hex 5 HexNAc 4

**Mass:**  4588.0853

(2.815e-07)

**Charge States:** 3, 4, 5

**Start Position**: 319

**End Position**: 343

| Profile Score | Tandem Score | TDA q-value | Volume | Spectrum Matches |
| --- | --- | --- | --- | --- |
| 18.1218 | 8.8466 | 0.0218 | 1.6686e+10 | 8 |

**Adduct Signal Fractions**

- **Unmodified**: 1.6686e+10

#### VPAQEKN(N)FTTAPAIC(C)HDGKAHFPREGVF Fuc 1 Hex 4 HexNAc 4

**Mass:**  4730.1182

(-8.439e-07)

**Charge States:** 3, 4, 5, 6

**Start Position**: 1068

**End Position**: 1096

| Profile Score | Tandem Score | TDA q-value | Volume | Spectrum Matches |
| --- | --- | --- | --- | --- |
| 13.9563 | 8.8073 | 0.0218 | 2.5273e+10 | 16 |

**Adduct Signal Fractions**

- **Unmodified**: 2.5273e+10

#### N(N)FSQILPDPSKPSKRSF Hex 3 HexNAc 5

**Mass:**  3448.5759

(-2.211e-07)

**Charge States:** 3, 4

**Start Position**: 801

**End Position**: 818

| Profile Score | Tandem Score | TDA q-value | Volume | Spectrum Matches |
| --- | --- | --- | --- | --- |
| 12.0954 | 8.7944 | 0.0224 | 5.9919e+08 | 7 |

**Adduct Signal Fractions**

- **Unmodified**: 5.9919e+08

#### IVNN(N)ATNVVIKVC(C)EF Hex 5 HexNAc 3

**Mass:**  3138.4052

(3.048e-07)

**Charge States:** 2, 3

**Start Position**: 119

**End Position**: 134

| Profile Score | Tandem Score | TDA q-value | Volume | Spectrum Matches |
| --- | --- | --- | --- | --- |
| 16.0367 | 8.7508 | 0.0224 | 2.8779e+10 | 8 |

**Adduct Signal Fractions**

- **Unmodified**: 2.8779e+10

#### RVQPTESIVRFPN(N)ITNL Fuc 1 Hex 3 HexNAc 5

**Mass:**  3630.7038

(2.335e-07)

**Charge States:** 3, 4

**Start Position**: 319

**End Position**: 336

| Profile Score | Tandem Score | TDA q-value | Volume | Spectrum Matches |
| --- | --- | --- | --- | --- |
| 15.5351 | 8.7145 | 0.0224 | 2.2827e+10 | 11 |

**Adduct Signal Fractions**

- **Unmodified**: 2.2827e+10

#### GGFN(N)FSQILPDPSKPSKRSF Fuc 1 Hex 4 HexNAc 4 Neu5Ac 1

**Mass:**  4105.8137

(-3.113e-07)

**Charge States:** 3, 4, 5

**Start Position**: 798

**End Position**: 818

| Profile Score | Tandem Score | TDA q-value | Volume | Spectrum Matches |
| --- | --- | --- | --- | --- |
| 15.4219 | 8.6820 | 0.0224 | 1.0709e+09 | 5 |

**Adduct Signal Fractions**

- **Unmodified**: 1.0709e+09

#### RVQPTESIVRFPN(N)ITNLC(C)PFGEVF Fuc 1 Hex 4 HexNAc 5

**Mass:**  4629.1089

(2.318e-07)

**Charge States:** 3, 4, 5

**Start Position**: 319

**End Position**: 343

| Profile Score | Tandem Score | TDA q-value | Volume | Spectrum Matches |
| --- | --- | --- | --- | --- |
| 14.3714 | 8.6775 | 0.0224 | 5.6970e+10 | 18 |

**Adduct Signal Fractions**

- **Unmodified**: 5.6970e+10

#### RVQPTESIVRFPN(N)ITNLC(C)PFGEVF Fuc 1 Hex 5 HexNAc 5

**Mass:**  4791.1643

(3.425e-07)

**Charge States:** 3, 4, 5

**Start Position**: 319

**End Position**: 343

| Profile Score | Tandem Score | TDA q-value | Volume | Spectrum Matches |
| --- | --- | --- | --- | --- |
| 13.5277 | 8.6612 | 0.0224 | 1.5127e+10 | 8 |

**Adduct Signal Fractions**

- **Unmodified**: 1.5127e+10

#### FHAIHVSGTN(N)GTKRF Hex 4 HexNAc 6

**Mass:**  3537.5556

(1.128e-06)

**Charge States:** 5

**Start Position**: 65

**End Position**: 80

| Profile Score | Tandem Score | TDA q-value | Volume | Spectrum Matches |
| --- | --- | --- | --- | --- |
| 6.2918 | 8.6016 | 0.0224 | 4.8853e+07 | 3 |

**Adduct Signal Fractions**

- **Unmodified**: 4.8853e+07

#### YHKNN(N)KSWMESEF Fuc 1 Hex 4 HexNAc 4 Neu5Ac 1

**Mass:**  3596.4311

(4.353e-07)

**Charge States:** 3, 4

**Start Position**: 145

**End Position**: 158

| Profile Score | Tandem Score | TDA q-value | Volume | Spectrum Matches |
| --- | --- | --- | --- | --- |
| 15.4589 | 8.6002 | 0.0224 | 2.6488e+09 | 3 |

**Adduct Signal Fractions**

- **Unmodified**: 2.6488e+09

#### VPAQEKN(N)FTTAPAIC(C)HDGKAHFPREGVF Fuc 1 Hex 6 HexNAc 3

**Mass:**  4851.1541

(9.752e-07)

**Charge States:** 4, 5, 6

**Start Position**: 1068

**End Position**: 1096

| Profile Score | Tandem Score | TDA q-value | Volume | Spectrum Matches |
| --- | --- | --- | --- | --- |
| 15.5487 | 8.5676 | 0.0224 | 1.4642e+09 | 6 |

**Adduct Signal Fractions**

- **Unmodified**: 1.4642e+09

#### N(N)FSQILPDPSKPSKRSF Fuc 1 Hex 5 HexNAc 4

**Mass:**  3715.6633

(4.516e-07)

**Charge States:** 3, 4

**Start Position**: 801

**End Position**: 818

| Profile Score | Tandem Score | TDA q-value | Volume | Spectrum Matches |
| --- | --- | --- | --- | --- |
| 13.9701 | 8.5552 | 0.0224 | 1.2504e+09 | 4 |

**Adduct Signal Fractions**

- **Unmodified**: 1.2504e+09

#### N(N)FSQILPDPSKPSKRSF Hex 5 HexNAc 4

**Mass:**  3569.6088

(1.098e-06)

**Charge States:** 3, 4

**Start Position**: 801

**End Position**: 818

| Profile Score | Tandem Score | TDA q-value | Volume | Spectrum Matches |
| --- | --- | --- | --- | --- |
| 13.5545 | 8.5003 | 0.0225 | 3.4009e+08 | 4 |

**Adduct Signal Fractions**

- **Unmodified**: 3.4009e+08

#### IVNN(N)ATNVVIKVC(C)EF Hex 4 HexNAc 4 Neu5Ac 1

**Mass:**  3470.5290

(6.574e-07)

**Charge States:** 3, 4

**Start Position**: 119

**End Position**: 134

| Profile Score | Tandem Score | TDA q-value | Volume | Spectrum Matches |
| --- | --- | --- | --- | --- |
| 14.4949 | 8.4970 | 0.0225 | 5.8375e+09 | 10 |

**Adduct Signal Fractions**

- **Unmodified**: 5.8375e+09

#### QTRAGC(C)LIGAEHVN(N)NSYEC(C)DIPIGAGIC(C)ASY Fuc 1 Hex 5 HexNAc 4 Neu5Ac 1

**Mass:**  5455.2688

(-7.009e-07)

**Charge States:** 4, 5, 6

**Start Position**: 644

**End Position**: 675

| Profile Score | Tandem Score | TDA q-value | Volume | Spectrum Matches |
| --- | --- | --- | --- | --- |
| 15.8354 | 8.4965 | 0.0225 | 2.5619e+10 | 12 |

**Adduct Signal Fractions**

- **Unmodified**: 2.5619e+10

#### QTRAGC(C)LIGAEHVN(N)NSYEC(C)DIPIGAGIC(C)ASY Fuc 1 Hex 5 HexNAc 4

**Mass:**  5164.1717

(-4.537e-08)

**Charge States:** 4, 5

**Start Position**: 644

**End Position**: 675

| Profile Score | Tandem Score | TDA q-value | Volume | Spectrum Matches |
| --- | --- | --- | --- | --- |
| 17.6689 | 8.4849 | 0.0225 | 1.2372e+10 | 3 |

**Adduct Signal Fractions**

- **Unmodified**: 1.2372e+10

#### GGFN(N)FSQILPDPSKPSKRSF Fuc 1 Hex 4 HexNAc 4

**Mass:**  3814.7247

(1.690e-06)

**Charge States:** 3, 4

**Start Position**: 798

**End Position**: 818

| Profile Score | Tandem Score | TDA q-value | Volume | Spectrum Matches |
| --- | --- | --- | --- | --- |
| 12.7121 | 8.4003 | 0.0225 | 7.7294e+08 | 4 |

**Adduct Signal Fractions**

- **Unmodified**: 7.7294e+08

#### LHVTYVPAQEKN(N)F Hex 5 HexNAc 2

**Mass:**  2761.2188

(1.186e-06)

**Charge States:** 3

**Start Position**: 1063

**End Position**: 1076

| Profile Score | Tandem Score | TDA q-value | Volume | Spectrum Matches |
| --- | --- | --- | --- | --- |
| 13.8104 | 8.3104 | 0.0225 | 8.2257e+08 | 1 |

**Adduct Signal Fractions**

- **Unmodified**: 8.2257e+08

#### RVQPTESIVRFPN(N)ITNLC(C)PFGEVF Fuc 2 Hex 5 HexNAc 4

**Mass:**  4734.1393

(6.699e-08)

**Charge States:** 3, 4, 5

**Start Position**: 319

**End Position**: 343

| Profile Score | Tandem Score | TDA q-value | Volume | Spectrum Matches |
| --- | --- | --- | --- | --- |
| 17.5758 | 8.3044 | 0.0225 | 3.8506e+09 | 4 |

**Adduct Signal Fractions**

- **Unmodified**: 3.8506e+09

#### HAIHVSGTN(N)GTKRF Hex 6 HexNAc 4 Neu5Ac 1

**Mass:**  3601.5636

(5.506e-04)

**Charge States:** 4

**Start Position**: 66

**End Position**: 80

| Profile Score | Tandem Score | TDA q-value | Volume | Spectrum Matches |
| --- | --- | --- | --- | --- |
| 0.9210 | 8.2890 | 0.0225 | 1.9303e+07 | 3 |

**Adduct Signal Fractions**

- **Unmodified**: 1.9303e+07

#### LIVNN(N)ATNVVIKVC(C)EF Hex 5 HexNAc 3 Neu5Ac 1

**Mass:**  3542.5881

(4.756e-07)

**Charge States:** 3, 4

**Start Position**: 118

**End Position**: 134

| Profile Score | Tandem Score | TDA q-value | Volume | Spectrum Matches |
| --- | --- | --- | --- | --- |
| 15.2074 | 8.1946 | 0.0225 | 3.9573e+09 | 8 |

**Adduct Signal Fractions**

- **Unmodified**: 3.9573e+09

#### RVQPTESIVRFPN(N)ITNLC(C)PF Fuc 1 Hex 4 HexNAc 5 Neu5Ac 1

**Mass:**  4488.0040

(2.240e-07)

**Charge States:** 3, 4

**Start Position**: 319

**End Position**: 339

| Profile Score | Tandem Score | TDA q-value | Volume | Spectrum Matches |
| --- | --- | --- | --- | --- |
| 12.5670 | 8.1883 | 0.0232 | 2.2504e+10 | 14 |

**Adduct Signal Fractions**

- **Unmodified**: 2.2504e+10

#### SNNSIAIPTN(N)F Fuc 2 Hex 5 HexNAc 6 Neu5Ac 1

**Mass:**  3789.5131

(2.597e-04)

**Start Position**: 708

**End Position**: 719

| Profile Score | Tandem Score | TDA q-value | Volume | Spectrum Matches |
| --- | --- | --- | --- | --- |
| 0.0000 | 8.1858 | 0.0232 | 0.0000e+00 | 1 |

Adduct:

No Chromatogram Found

#### VPAQEKN(N)FTTAPAIC(C)HDGKAHFPREGVF Fuc 1 Hex 6 HexNAc 5

**Mass:**  5257.3107

(4.966e-07)

**Charge States:** 4, 5, 6

**Start Position**: 1068

**End Position**: 1096

| Profile Score | Tandem Score | TDA q-value | Volume | Spectrum Matches |
| --- | --- | --- | --- | --- |
| 18.0350 | 8.1478 | 0.0232 | 8.6838e+09 | 6 |

**Adduct Signal Fractions**

- **Unmodified**: 8.6838e+09

#### RVQPTESIVRFPN(N)ITNLC(C)PFGEVF Fuc 2 Hex 4 HexNAc 4

**Mass:**  4572.0877

(-1.705e-07)

**Charge States:** 3, 4

**Start Position**: 319

**End Position**: 343

| Profile Score | Tandem Score | TDA q-value | Volume | Spectrum Matches |
| --- | --- | --- | --- | --- |
| 13.6617 | 8.1208 | 0.0232 | 7.8733e+09 | 5 |

**Adduct Signal Fractions**

- **Unmodified**: 7.8733e+09

#### QTRAGC(C)LIGAEHVN(N)NSYEC(C)DIPIGAGIC(C)ASY Fuc 1 Hex 5 HexNAc 5

**Mass:**  5367.2503

(-9.395e-08)

**Charge States:** 4, 5

**Start Position**: 644

**End Position**: 675

| Profile Score | Tandem Score | TDA q-value | Volume | Spectrum Matches |
| --- | --- | --- | --- | --- |
| 13.5962 | 8.0516 | 0.0236 | 1.5424e+10 | 4 |

**Adduct Signal Fractions**

- **Unmodified**: 1.5424e+10

#### LIVNN(N)ATNVVIKVC(C)EF Hex 7 HexNAc 2

**Mass:**  3372.5177

(6.487e-07)

**Charge States:** 3

**Start Position**: 118

**End Position**: 134

| Profile Score | Tandem Score | TDA q-value | Volume | Spectrum Matches |
| --- | --- | --- | --- | --- |
| 17.2400 | 8.0365 | 0.0236 | 6.1856e+08 | 2 |

**Adduct Signal Fractions**

- **Unmodified**: 6.1856e+08

#### RVQPTESIVRFPN(N)ITNLC(C)PF Fuc 1 Hex 3 HexNAc 6

**Mass:**  4237.9317

(-5.546e-07)

**Charge States:** 3, 4, 5

**Start Position**: 319

**End Position**: 339

| Profile Score | Tandem Score | TDA q-value | Volume | Spectrum Matches |
| --- | --- | --- | --- | --- |
| 15.0414 | 8.0252 | 0.0236 | 3.3518e+09 | 5 |

**Adduct Signal Fractions**

- **Unmodified**: 3.3518e+09

#### LIVNN(N)ATNVVIKVC(C)EF Fuc 1 Hex 3 HexNAc 4

**Mass:**  3276.5212

(1.490e-07)

**Charge States:** 3

**Start Position**: 118

**End Position**: 134

| Profile Score | Tandem Score | TDA q-value | Volume | Spectrum Matches |
| --- | --- | --- | --- | --- |
| 15.7642 | 7.9942 | 0.0236 | 2.4794e+09 | 5 |

**Adduct Signal Fractions**

- **Unmodified**: 2.4794e+09

#### RVQPTESIVRFPN(N)ITNL Fuc 1 Hex 4 HexNAc 4

**Mass:**  3589.6774

(1.424e-07)

**Charge States:** 3, 4

**Start Position**: 319

**End Position**: 336

| Profile Score | Tandem Score | TDA q-value | Volume | Spectrum Matches |
| --- | --- | --- | --- | --- |
| 15.6218 | 7.9269 | 0.0236 | 8.7777e+09 | 11 |

**Adduct Signal Fractions**

- **Unmodified**: 8.7777e+09

#### FHAIHVSGTN(N)GTKRF Fuc 2 Hex 4 HexNAc 7

**Mass:**  4032.7452

(-4.004e-07)

**Charge States:** 5

**Start Position**: 65

**End Position**: 80

| Profile Score | Tandem Score | TDA q-value | Volume | Spectrum Matches |
| --- | --- | --- | --- | --- |
| -4.0632 | 7.9128 | 0.0236 | 2.5963e+07 | 2 |

**Adduct Signal Fractions**

- **Unmodified**: 2.5963e+07

#### RVQPTESIVRFPN(N)ITNLC(C)PF Fuc 2 Hex 5 HexNAc 4

**Mass:**  4301.9424

(-3.868e-07)

**Charge States:** 3, 4

**Start Position**: 319

**End Position**: 339

| Profile Score | Tandem Score | TDA q-value | Volume | Spectrum Matches |
| --- | --- | --- | --- | --- |
| 16.1360 | 7.9008 | 0.0236 | 1.8795e+09 | 4 |

**Adduct Signal Fractions**

- **Unmodified**: 1.8795e+09

#### RVQPTESIVRFPN(N)ITNLC(C)PF Fuc 2 Hex 4 HexNAc 5

**Mass:**  4342.9669

(6.210e-08)

**Charge States:** 3, 4, 5

**Start Position**: 319

**End Position**: 339

| Profile Score | Tandem Score | TDA q-value | Volume | Spectrum Matches |
| --- | --- | --- | --- | --- |
| 15.6269 | 7.8884 | 0.0242 | 2.9509e+09 | 7 |

**Adduct Signal Fractions**

- **Unmodified**: 2.9509e+09

#### KTPPIKDFGGFN(N)F Hex 6 HexNAc 2

**Mass:**  2845.2356

(1.296e-06)

**Charge States:** 2, 3

**Start Position**: 790

**End Position**: 803

| Profile Score | Tandem Score | TDA q-value | Volume | Spectrum Matches |
| --- | --- | --- | --- | --- |
| 15.3522 | 7.8776 | 0.0242 | 9.6080e+08 | 4 |

**Adduct Signal Fractions**

- **Unmodified**: 9.6080e+08

#### VPAQEKN(N)FTTAPAIC(C)HDGKAHFPREGVF Fuc 1 Hex 4 HexNAc 3 Neu5Ac 1

**Mass:**  4818.1426

(1.370e-06)

**Charge States:** 4, 5, 6

**Start Position**: 1068

**End Position**: 1096

| Profile Score | Tandem Score | TDA q-value | Volume | Spectrum Matches |
| --- | --- | --- | --- | --- |
| 14.7257 | 7.8520 | 0.0242 | 2.0462e+09 | 3 |

**Adduct Signal Fractions**

- **Unmodified**: 2.0462e+09

#### N(N)FSQILPDPSKPSKRSF Fuc 1 Hex 5 HexNAc 3

**Mass:**  3512.5806

(-4.125e-07)

**Charge States:** 4

**Start Position**: 801

**End Position**: 818

| Profile Score | Tandem Score | TDA q-value | Volume | Spectrum Matches |
| --- | --- | --- | --- | --- |
| 12.3329 | 7.8358 | 0.0242 | 1.3955e+08 | 2 |

**Adduct Signal Fractions**

- **Unmodified**: 1.3955e+08

#### SN(N)NSIAIPTN(N)F Hex 15 HexNAc 4

**Mass:**  4419.6875

(3.026e-07)

**Charge States:** 3

**Start Position**: 708

**End Position**: 719

| Profile Score | Tandem Score | TDA q-value | Volume | Spectrum Matches |
| --- | --- | --- | --- | --- |
| 15.4419 | 7.8160 | 0.0242 | 2.6994e+09 | 3 |

**Adduct Signal Fractions**

- **Unmodified**: 2.6994e+09

#### HAIHVSGTN(N)GTKRF Fuc 1 Hex 6 HexNAc 5 Neu5Ac 3

**Mass:**  4533.8537

(6.621e-04)

**Start Position**: 66

**End Position**: 80

| Profile Score | Tandem Score | TDA q-value | Volume | Spectrum Matches |
| --- | --- | --- | --- | --- |
| 0.0000 | 7.8075 | 0.0242 | 0.0000e+00 | 1 |

Adduct:

No Chromatogram Found

#### RVQPTESIVRFPN(N)ITNLC(C)PF Fuc 1 Hex 5 HexNAc 4 Neu5Ac 1

**Mass:**  4446.9773

(-9.800e-07)

**Charge States:** 3, 4, 5

**Start Position**: 319

**End Position**: 339

| Profile Score | Tandem Score | TDA q-value | Volume | Spectrum Matches |
| --- | --- | --- | --- | --- |
| 15.6719 | 7.7861 | 0.0242 | 8.6185e+10 | 14 |

**Adduct Signal Fractions**

- **Unmodified**: 8.6185e+10

#### HAIHVSGTN(N)GTKRF Fuc 1 Hex 6 HexNAc 7 Neu5Ac 3

**Mass:**  4937.0095

(-5.517e-07)

**Charge States:** 4, 5

**Start Position**: 66

**End Position**: 80

| Profile Score | Tandem Score | TDA q-value | Volume | Spectrum Matches |
| --- | --- | --- | --- | --- |
| 11.5744 | 7.7572 | 0.0251 | 1.6281e+09 | 9 |

**Adduct Signal Fractions**

- **Unmodified**: 1.6281e+09

#### RVQPTESIVRFPN(N)ITNL Fuc 1 Hex 5 HexNAc 5

**Mass:**  3954.8086

(2.987e-07)

**Charge States:** 3, 4

**Start Position**: 319

**End Position**: 336

| Profile Score | Tandem Score | TDA q-value | Volume | Spectrum Matches |
| --- | --- | --- | --- | --- |
| 13.0204 | 7.7223 | 0.0258 | 7.6274e+09 | 8 |

**Adduct Signal Fractions**

- **Unmodified**: 7.6274e+09

#### RVQPTESIVRFPN(N)ITNLC(C)PFGEVF Fuc 1 Hex 6 HexNAc 5 Neu5Ac 1

**Mass:**  5244.3145

(8.903e-07)

**Charge States:** 4, 5, 6

**Start Position**: 319

**End Position**: 343

| Profile Score | Tandem Score | TDA q-value | Volume | Spectrum Matches |
| --- | --- | --- | --- | --- |
| 14.5334 | 7.7075 | 0.0258 | 1.7781e+10 | 6 |

**Adduct Signal Fractions**

- **Unmodified**: 1.7781e+10

#### RVQPTESIVRFPN(N)ITNLC(C)PF Hex 5 HexNAc 2

**Mass:**  3603.6637

(-5.020e-07)

**Charge States:** 3, 4

**Start Position**: 319

**End Position**: 339

| Profile Score | Tandem Score | TDA q-value | Volume | Spectrum Matches |
| --- | --- | --- | --- | --- |
| 14.8503 | 7.6816 | 0.0258 | 2.4406e+09 | 5 |

**Adduct Signal Fractions**

- **Unmodified**: 2.4406e+09

#### RVQPTESIVRFPN(N)ITNLC(C)PF Fuc 1 Hex 5 HexNAc 4 Neu5Ac 2

**Mass:**  4738.0769

(3.743e-07)

**Charge States:** 3, 4, 5

**Start Position**: 319

**End Position**: 339

| Profile Score | Tandem Score | TDA q-value | Volume | Spectrum Matches |
| --- | --- | --- | --- | --- |
| 14.5475 | 7.6594 | 0.0258 | 1.1529e+10 | 6 |

**Adduct Signal Fractions**

- **Unmodified**: 1.1529e+10

#### HAIHVSGTN(N)GTKRF Fuc 1 Hex 8 HexNAc 4

**Mass:**  3778.5943

(2.430e-07)

**Charge States:** 3, 4

**Start Position**: 66

**End Position**: 80

| Profile Score | Tandem Score | TDA q-value | Volume | Spectrum Matches |
| --- | --- | --- | --- | --- |
| 17.5401 | 7.6440 | 0.0258 | 4.9663e+08 | 1 |

**Adduct Signal Fractions**

- **Unmodified**: 4.9663e+08

#### RVQPTESIVRFPN(N)ITNLC(C)PFGEVF Fuc 1 Hex 4 HexNAc 4 Neu5Ac 1

**Mass:**  4717.1243

(-1.166e-07)

**Charge States:** 3, 4, 5

**Start Position**: 319

**End Position**: 343

| Profile Score | Tandem Score | TDA q-value | Volume | Spectrum Matches |
| --- | --- | --- | --- | --- |
| 18.1444 | 7.5877 | 0.0262 | 7.5829e+10 | 19 |

**Adduct Signal Fractions**

- **Unmodified**: 7.5829e+10

#### SN(N)NSIAIPTN(N)F Fuc 1 Hex 10 HexNAc 6

**Mass:**  4161.6406

(4.817e-07)

**Charge States:** 3, 4

**Start Position**: 708

**End Position**: 719

| Profile Score | Tandem Score | TDA q-value | Volume | Spectrum Matches |
| --- | --- | --- | --- | --- |
| 15.6719 | 7.5608 | 0.0262 | 1.3041e+09 | 3 |

**Adduct Signal Fractions**

- **Unmodified**: 1.3041e+09

#### GGFN(N)FSQILPDPSKPSKRSF Fuc 1 Hex 3 HexNAc 5

**Mass:**  3855.7496

(1.657e-06)

**Charge States:** 3, 4

**Start Position**: 798

**End Position**: 818

| Profile Score | Tandem Score | TDA q-value | Volume | Spectrum Matches |
| --- | --- | --- | --- | --- |
| 14.2607 | 7.5482 | 0.0262 | 1.0072e+09 | 5 |

**Adduct Signal Fractions**

- **Unmodified**: 1.0072e+09

#### IVNN(N)ATNVVIKVC(C)EF Fuc 1 Hex 4 HexNAc 5

**Mass:**  3528.5725

(1.235e-06)

**Charge States:** 3, 4

**Start Position**: 119

**End Position**: 134

| Profile Score | Tandem Score | TDA q-value | Volume | Spectrum Matches |
| --- | --- | --- | --- | --- |
| 12.8205 | 7.5384 | 0.0262 | 1.7137e+09 | 4 |

**Adduct Signal Fractions**

- **Unmodified**: 1.7137e+09

#### GGFN(N)FSQILPDPSKPSKRSF Hex 6 HexNAc 3 Neu5Ac 1

**Mass:**  4080.7820

(-8.552e-07)

**Charge States:** 4

**Start Position**: 798

**End Position**: 818

| Profile Score | Tandem Score | TDA q-value | Volume | Spectrum Matches |
| --- | --- | --- | --- | --- |
| 12.7043 | 7.5251 | 0.0262 | 1.3997e+09 | 6 |

**Adduct Signal Fractions**

- **Unmodified**: 1.3997e+09

#### IVNN(N)ATNVVIKVC(C)EF Fuc 1 Hex 4 HexNAc 3

**Mass:**  3122.4115

(5.189e-07)

**Charge States:** 2, 3

**Start Position**: 119

**End Position**: 134

| Profile Score | Tandem Score | TDA q-value | Volume | Spectrum Matches |
| --- | --- | --- | --- | --- |
| 16.1216 | 7.5237 | 0.0262 | 9.3013e+09 | 10 |

**Adduct Signal Fractions**

- **Unmodified**: 9.3013e+09

#### RVQPTESIVRFPN(N)ITNLC(C)PF Fuc 1 Hex 5 HexNAc 6

**Mass:**  4562.0439

(7.268e-07)

**Charge States:** 3, 4, 5

**Start Position**: 319

**End Position**: 339

| Profile Score | Tandem Score | TDA q-value | Volume | Spectrum Matches |
| --- | --- | --- | --- | --- |
| 14.3751 | 7.4688 | 0.0262 | 1.6789e+09 | 3 |

**Adduct Signal Fractions**

- **Unmodified**: 1.6789e+09

#### IVNN(N)ATNVVIKVC(C)EF Hex 4 HexNAc 3 Neu5Ac 1

**Mass:**  3267.4478

(1.362e-07)

**Charge States:** 3

**Start Position**: 119

**End Position**: 134

| Profile Score | Tandem Score | TDA q-value | Volume | Spectrum Matches |
| --- | --- | --- | --- | --- |
| 13.2769 | 7.4605 | 0.0262 | 2.2375e+09 | 4 |

**Adduct Signal Fractions**

- **Unmodified**: 2.2375e+09

#### HVTYVPAQEKN(N)F Hex 5 HexNAc 2

**Mass:**  2648.1358

(-3.578e-07)

**Charge States:** 2, 3

**Start Position**: 1064

**End Position**: 1076

| Profile Score | Tandem Score | TDA q-value | Volume | Spectrum Matches |
| --- | --- | --- | --- | --- |
| 17.8419 | 7.4297 | 0.0262 | 1.3574e+09 | 3 |

**Adduct Signal Fractions**

- **Unmodified**: 1.3574e+09

#### HAIHVSGTN(N)GTKRF Fuc 1 Hex 6 HexNAc 7 Neu5Ac 2

**Mass:**  4645.9141

(-5.876e-07)

**Charge States:** 4, 5, 6

**Start Position**: 66

**End Position**: 80

| Profile Score | Tandem Score | TDA q-value | Volume | Spectrum Matches |
| --- | --- | --- | --- | --- |
| 11.2244 | 7.4204 | 0.0262 | 3.9668e+09 | 22 |

**Adduct Signal Fractions**

- **Unmodified**: 3.9668e+09

#### IVNN(N)ATNVVIKVC(C)EF Hex 3 HexNAc 4

**Mass:**  3017.3802

(5.729e-07)

**Charge States:** 2, 3

**Start Position**: 119

**End Position**: 134

| Profile Score | Tandem Score | TDA q-value | Volume | Spectrum Matches |
| --- | --- | --- | --- | --- |
| 14.6575 | 7.4092 | 0.0262 | 1.1818e+10 | 7 |

**Adduct Signal Fractions**

- **Unmodified**: 1.1818e+10

#### IVNN(N)ATNVVIKVC(C)EF Hex 4 HexNAc 5

**Mass:**  3382.5126

(5.622e-07)

**Charge States:** 3

**Start Position**: 119

**End Position**: 134

| Profile Score | Tandem Score | TDA q-value | Volume | Spectrum Matches |
| --- | --- | --- | --- | --- |
| 11.3498 | 7.3377 | 0.0262 | 1.4407e+09 | 5 |

**Adduct Signal Fractions**

- **Unmodified**: 1.4407e+09

#### FHAIHVSGTN(N)GTKRF Hex 4 HexNAc 7 Neu5Ac 1

**Mass:**  4031.7380

(2.788e-06)

**Charge States:** 3, 4, 5

**Start Position**: 65

**End Position**: 80

| Profile Score | Tandem Score | TDA q-value | Volume | Spectrum Matches |
| --- | --- | --- | --- | --- |
| 13.4171 | 7.3203 | 0.0262 | 1.8445e+09 | 5 |

**Adduct Signal Fractions**

- **Unmodified**: 1.8445e+09

#### SN(N)VTWFHAIHVSGTN(N)GTKRF Fuc 1 Hex 11 HexNAc 13 Neu5Ac 2

**Mass:**  7409.0253

(4.058e-06)

**Charge States:** 7

**Start Position**: 60

**End Position**: 80

| Profile Score | Tandem Score | TDA q-value | Volume | Spectrum Matches |
| --- | --- | --- | --- | --- |
| 10.7839 | 7.3188 | 0.0262 | 1.4652e+08 | 3 |

**Adduct Signal Fractions**

- **Unmodified**: 1.4652e+08

#### IVNN(N)ATNVVIKVC(C)EF Hex 4 HexNAc 4

**Mass:**  3179.4339

(1.073e-06)

**Charge States:** 2, 3

**Start Position**: 119

**End Position**: 134

| Profile Score | Tandem Score | TDA q-value | Volume | Spectrum Matches |
| --- | --- | --- | --- | --- |
| 13.7343 | 7.3126 | 0.0262 | 9.6691e+09 | 9 |

**Adduct Signal Fractions**

- **Unmodified**: 9.6691e+09

#### N(N)FSQILPDPSKPSKRSF Hex 4 HexNAc 4 Neu5Ac 1

**Mass:**  3698.6508

(2.726e-04)

**Charge States:** 4

**Start Position**: 801

**End Position**: 818

| Profile Score | Tandem Score | TDA q-value | Volume | Spectrum Matches |
| --- | --- | --- | --- | --- |
| 11.5922 | 7.2897 | 0.0262 | 2.1684e+09 | 6 |

**Adduct Signal Fractions**

- **Unmodified**: 2.1684e+09

#### IVNN(N)ATNVVIKVC(C)EF Hex 3 HexNAc 3

**Mass:**  2814.2998

(3.613e-07)

**Charge States:** 2, 3

**Start Position**: 119

**End Position**: 134

| Profile Score | Tandem Score | TDA q-value | Volume | Spectrum Matches |
| --- | --- | --- | --- | --- |
| 16.2093 | 7.2557 | 0.0264 | 1.6749e+10 | 5 |

**Adduct Signal Fractions**

- **Unmodified**: 1.6749e+10

#### IVNN(N)ATNVVIKVC(C)EF Hex 3 HexNAc 5

**Mass:**  3220.4573

(-1.966e-07)

**Charge States:** 3, 4

**Start Position**: 119

**End Position**: 134

| Profile Score | Tandem Score | TDA q-value | Volume | Spectrum Matches |
| --- | --- | --- | --- | --- |
| 15.7448 | 7.2450 | 0.0264 | 2.8199e+09 | 4 |

**Adduct Signal Fractions**

- **Unmodified**: 2.8199e+09

#### GGFN(N)FSQILPDPSKPSKRSF Hex 5 HexNAc 3 Neu5Ac 1

**Mass:**  3918.7307

(7.213e-08)

**Charge States:** 3, 4

**Start Position**: 798

**End Position**: 818

| Profile Score | Tandem Score | TDA q-value | Volume | Spectrum Matches |
| --- | --- | --- | --- | --- |
| 15.7816 | 7.2417 | 0.0264 | 1.1049e+09 | 6 |

**Adduct Signal Fractions**

- **Unmodified**: 1.1049e+09

#### YHKNN(N)KSWMESEF Fuc 1 Hex 3 HexNAc 5

**Mass:**  3346.3553

(-9.104e-07)

**Charge States:** 3, 4

**Start Position**: 145

**End Position**: 158

| Profile Score | Tandem Score | TDA q-value | Volume | Spectrum Matches |
| --- | --- | --- | --- | --- |
| 14.6481 | 7.2158 | 0.0264 | 2.8007e+10 | 15 |

**Adduct Signal Fractions**

- **Unmodified**: 2.8007e+10

#### IVNN(N)ATNVVIKVC(C)EF Hex 6 HexNAc 2

**Mass:**  3097.3790

(2.067e-07)

**Charge States:** 2, 3

**Start Position**: 119

**End Position**: 134

| Profile Score | Tandem Score | TDA q-value | Volume | Spectrum Matches |
| --- | --- | --- | --- | --- |
| 15.8943 | 7.2113 | 0.0264 | 3.0404e+09 | 4 |

**Adduct Signal Fractions**

- **Unmodified**: 3.0404e+09

#### LIVNN(N)ATNVVIKVC(C)EF Hex 5 HexNAc 2

**Mass:**  3048.4096

(3.343e-07)

**Charge States:** 2, 3

**Start Position**: 118

**End Position**: 134

| Profile Score | Tandem Score | TDA q-value | Volume | Spectrum Matches |
| --- | --- | --- | --- | --- |
| 15.2174 | 7.1898 | 0.0264 | 9.2900e+09 | 8 |

**Adduct Signal Fractions**

- **Unmodified**: 9.2900e+09

#### N(N)FSQILPDPSKPSKRSF Hex 4 HexNAc 4

**Mass:**  3407.5496

(-1.316e-06)

**Charge States:** 4

**Start Position**: 801

**End Position**: 818

| Profile Score | Tandem Score | TDA q-value | Volume | Spectrum Matches |
| --- | --- | --- | --- | --- |
| 10.3117 | 7.1851 | 0.0264 | 3.3807e+08 | 3 |

**Adduct Signal Fractions**

- **Unmodified**: 3.3807e+08

#### RVQPTESIVRFPN(N)ITNLC(C)PFGEVF Fuc 1 Hex 4 HexNAc 5 Neu5Ac 1

**Mass:**  4920.2022

(1.160e-07)

**Charge States:** 4, 5

**Start Position**: 319

**End Position**: 343

| Profile Score | Tandem Score | TDA q-value | Volume | Spectrum Matches |
| --- | --- | --- | --- | --- |
| 16.1527 | 7.1846 | 0.0264 | 1.9734e+10 | 9 |

**Adduct Signal Fractions**

- **Unmodified**: 1.9734e+10

#### SALEPLVDLPIGIN(N)ITRF Hex 6 HexNAc 3

**Mass:**  3548.6642

(-4.290e-08)

**Charge States:** 3

**Start Position**: 221

**End Position**: 239

| Profile Score | Tandem Score | TDA q-value | Volume | Spectrum Matches |
| --- | --- | --- | --- | --- |
| 15.5203 | 7.1828 | 0.0264 | 1.2822e+09 | 4 |

**Adduct Signal Fractions**

- **Unmodified**: 1.2822e+09

#### FHAIHVSGTN(N)GTKRF Fuc 1 Hex 3 HexNAc 8

**Mass:**  3927.7142

(-2.358e-07)

**Charge States:** 5

**Start Position**: 65

**End Position**: 80

| Profile Score | Tandem Score | TDA q-value | Volume | Spectrum Matches |
| --- | --- | --- | --- | --- |
| 6.7190 | 7.1128 | 0.0264 | 3.4706e+07 | 1 |

**Adduct Signal Fractions**

- **Unmodified**: 3.4706e+07

#### HAIHVSGTN(N)GTKRF Fuc 1 Hex 4 HexNAc 8

**Mass:**  3942.7089

(2.404e-06)

**Charge States:** 4, 5

**Start Position**: 66

**End Position**: 80

| Profile Score | Tandem Score | TDA q-value | Volume | Spectrum Matches |
| --- | --- | --- | --- | --- |
| 10.0681 | 7.1082 | 0.0264 | 1.0387e+08 | 3 |

**Adduct Signal Fractions**

- **Unmodified**: 1.0387e+08

#### LIVNN(N)ATNVVIKVC(C)EF Hex 6 HexNAc 2

**Mass:**  3210.4630

(7.966e-08)

**Charge States:** 3

**Start Position**: 118

**End Position**: 134

| Profile Score | Tandem Score | TDA q-value | Volume | Spectrum Matches |
| --- | --- | --- | --- | --- |
| 16.0631 | 7.0904 | 0.0264 | 8.3877e+08 | 2 |

**Adduct Signal Fractions**

- **Unmodified**: 8.3877e+08

#### N(N)FSQILPDPSKPSKRSF Hex 3 HexNAc 3

**Mass:**  3042.4188

(1.717e-06)

**Charge States:** 3, 4

**Start Position**: 801

**End Position**: 818

| Profile Score | Tandem Score | TDA q-value | Volume | Spectrum Matches |
| --- | --- | --- | --- | --- |
| 11.6264 | 7.0674 | 0.0264 | 2.6936e+09 | 7 |

**Adduct Signal Fractions**

- **Unmodified**: 2.6936e+09

#### VPAQEKN(N)FTTAPAIC(C)HDGKAHFPREGVF Fuc 1 Hex 5 HexNAc 6

**Mass:**  5298.3359

(2.721e-07)

**Charge States:** 4, 5, 6

**Start Position**: 1068

**End Position**: 1096

| Profile Score | Tandem Score | TDA q-value | Volume | Spectrum Matches |
| --- | --- | --- | --- | --- |
| 12.6135 | 7.0661 | 0.0264 | 3.5160e+09 | 6 |

**Adduct Signal Fractions**

- **Unmodified**: 3.5160e+09

#### YHKNN(N)KSWMESEF Fuc 1 Hex 3 HexNAc 4

**Mass:**  3143.2825

(7.100e-08)

**Charge States:** 2, 3, 4

**Start Position**: 145

**End Position**: 158

| Profile Score | Tandem Score | TDA q-value | Volume | Spectrum Matches |
| --- | --- | --- | --- | --- |
| 17.3328 | 6.9852 | 0.0288 | 2.0280e+10 | 12 |

**Adduct Signal Fractions**

- **Unmodified**: 2.0280e+10

#### HAIHVSGTN(N)GTKRF Fuc 2 Hex 6 HexNAc 6

**Mass:**  4007.7061

(2.499e-04)

**Start Position**: 66

**End Position**: 80

| Profile Score | Tandem Score | TDA q-value | Volume | Spectrum Matches |
| --- | --- | --- | --- | --- |
| 0.0000 | 6.9719 | 0.0296 | 0.0000e+00 | 1 |

Adduct:

No Chromatogram Found

#### GGFN(N)FSQILPDPSKPSKRSF Hex 4 HexNAc 3

**Mass:**  3465.5793

(-1.458e-06)

**Charge States:** 4

**Start Position**: 798

**End Position**: 818

| Profile Score | Tandem Score | TDA q-value | Volume | Spectrum Matches |
| --- | --- | --- | --- | --- |
| 10.8672 | 6.9681 | 0.0296 | 4.0005e+08 | 1 |

**Adduct Signal Fractions**

- **Unmodified**: 4.0005e+08

#### RVQPTESIVRFPN(N)ITNLC(C)PFGEVF Fuc 1 Hex 6 HexNAc 5

**Mass:**  4953.2122

(-8.469e-07)

**Charge States:** 4, 5

**Start Position**: 319

**End Position**: 343

| Profile Score | Tandem Score | TDA q-value | Volume | Spectrum Matches |
| --- | --- | --- | --- | --- |
| 17.7096 | 6.9667 | 0.0296 | 4.0186e+09 | 4 |

**Adduct Signal Fractions**

- **Unmodified**: 4.0186e+09

#### KTPPIKDFGGFN(N)F Hex 7 HexNAc 2

**Mass:**  3007.2857

(8.432e-08)

**Charge States:** 2, 3

**Start Position**: 790

**End Position**: 803

| Profile Score | Tandem Score | TDA q-value | Volume | Spectrum Matches |
| --- | --- | --- | --- | --- |
| 16.6497 | 6.8400 | 0.0296 | 1.4200e+09 | 4 |

**Adduct Signal Fractions**

- **Unmodified**: 1.4200e+09

#### HAIHVSGTN(N)GTKRF Fuc 2 Hex 6 HexNAc 7

**Mass:**  4209.7845

(-5.616e-07)

**Charge States:** 4, 5

**Start Position**: 66

**End Position**: 80

| Profile Score | Tandem Score | TDA q-value | Volume | Spectrum Matches |
| --- | --- | --- | --- | --- |
| 12.0677 | 6.7825 | 0.0296 | 1.4860e+08 | 6 |

**Adduct Signal Fractions**

- **Unmodified**: 1.4860e+08

#### FHAIHVSGTN(N)GTKRF Hex 5 HexNAc 2

**Mass:**  2887.2871

(3.241e-08)

**Charge States:** 4

**Start Position**: 65

**End Position**: 80

| Profile Score | Tandem Score | TDA q-value | Volume | Spectrum Matches |
| --- | --- | --- | --- | --- |
| 12.9723 | 6.7665 | 0.0296 | 1.4614e+07 | 2 |

**Adduct Signal Fractions**

- **Unmodified**: 1.4614e+07

#### KTPPIKDFGGFN(N)F Hex 5 HexNAc 2

**Mass:**  2683.1805

(7.462e-07)

**Charge States:** 2, 3

**Start Position**: 790

**End Position**: 803

| Profile Score | Tandem Score | TDA q-value | Volume | Spectrum Matches |
| --- | --- | --- | --- | --- |
| 15.6482 | 6.7530 | 0.0296 | 3.9183e+10 | 8 |

**Adduct Signal Fractions**

- **Unmodified**: 3.9183e+10

#### RVQPTESIVRFPN(N)ITNLC(C)PFGEVF Fuc 1 Hex 3 HexNAc 4

**Mass:**  4263.9763

(1.680e-07)

**Charge States:** 3, 4, 5

**Start Position**: 319

**End Position**: 343

| Profile Score | Tandem Score | TDA q-value | Volume | Spectrum Matches |
| --- | --- | --- | --- | --- |
| 19.6717 | 6.6640 | 0.0296 | 1.0374e+11 | 14 |

**Adduct Signal Fractions**

- **Unmodified**: 1.0374e+11

#### EPLVDLPIGIN(N)ITRF @sulfate 1 Hex 6 HexNAc 4

**Mass:**  3561.5603

(2.844e-04)

**Charge States:** 3, 4

**Start Position**: 224

**End Position**: 239

| Profile Score | Tandem Score | TDA q-value | Volume | Spectrum Matches |
| --- | --- | --- | --- | --- |
| 10.3376 | 6.5725 | 0.0296 | 6.9269e+08 | 1 |

**Adduct Signal Fractions**

- **Unmodified**: 6.9269e+08

#### RVQPTESIVRFPN(N)ITNLC(C)PF Fuc 2 Hex 5 HexNAc 5

**Mass:**  4505.0169

(-3.400e-07)

**Charge States:** 3, 4

**Start Position**: 319

**End Position**: 339

| Profile Score | Tandem Score | TDA q-value | Volume | Spectrum Matches |
| --- | --- | --- | --- | --- |
| 15.6869 | 6.5321 | 0.0308 | 1.1719e+09 | 6 |

**Adduct Signal Fractions**

- **Unmodified**: 1.1719e+09

#### HAIHVSGTN(N)GTKRF Fuc 1 Hex 7 HexNAc 6 Neu5Ac 4

**Mass:**  5188.0807

(1.949e-04)

**Charge States:** 4

**Start Position**: 66

**End Position**: 80

| Profile Score | Tandem Score | TDA q-value | Volume | Spectrum Matches |
| --- | --- | --- | --- | --- |
| -3.4370 | 6.4659 | 0.0308 | 1.5560e+08 | 1 |

**Adduct Signal Fractions**

- **Unmodified**: 1.5560e+08

#### RVQPTESIVRFPN(N)ITNL Fuc 2 Hex 5 HexNAc 4

**Mass:**  3897.7888

(1.105e-06)

**Charge States:** 3, 4

**Start Position**: 319

**End Position**: 336

| Profile Score | Tandem Score | TDA q-value | Volume | Spectrum Matches |
| --- | --- | --- | --- | --- |
| 13.2790 | 6.4322 | 0.0315 | 1.6684e+09 | 3 |

**Adduct Signal Fractions**

- **Unmodified**: 1.6684e+09

#### HAIHVSGTN(N)GTKRF Fuc 1 Hex 6 HexNAc 8 Neu5Ac 1

**Mass:**  4557.9059

(9.470e-07)

**Charge States:** 4, 5

**Start Position**: 66

**End Position**: 80

| Profile Score | Tandem Score | TDA q-value | Volume | Spectrum Matches |
| --- | --- | --- | --- | --- |
| 12.4070 | 6.3869 | 0.0329 | 1.5833e+08 | 5 |

**Adduct Signal Fractions**

- **Unmodified**: 1.5833e+08

#### IVNN(N)ATNVVIKVC(C)EF Hex 5 HexNAc 4

**Mass:**  3341.4864

(1.672e-07)

**Charge States:** 3, 4

**Start Position**: 119

**End Position**: 134

| Profile Score | Tandem Score | TDA q-value | Volume | Spectrum Matches |
| --- | --- | --- | --- | --- |
| 15.6638 | 6.3806 | 0.0329 | 2.9455e+09 | 3 |

**Adduct Signal Fractions**

- **Unmodified**: 2.9455e+09

#### N(N)FSQILPDPSKPSKRSF Hex 6 HexNAc 3

**Mass:**  3529.5814

(2.848e-04)

**Start Position**: 801

**End Position**: 818

| Profile Score | Tandem Score | TDA q-value | Volume | Spectrum Matches |
| --- | --- | --- | --- | --- |
| 0.0000 | 6.3682 | 0.0329 | 0.0000e+00 | 1 |

Adduct:

No Chromatogram Found

#### EPLVDLPIGIN(N)ITRF Fuc 2 Hex 4 HexNAc 6

**Mass:**  3854.7275

(-7.266e-06)

**Charge States:** 3, 4

**Start Position**: 224

**End Position**: 239

| Profile Score | Tandem Score | TDA q-value | Volume | Spectrum Matches |
| --- | --- | --- | --- | --- |
| 10.8488 | 6.3469 | 0.0337 | 3.2453e+09 | 2 |

**Adduct Signal Fractions**

- **Unmodified**: 3.2453e+09

#### FHAIHVSGTN(N)GTKRF Fuc 1 Hex 4 HexNAc 4

**Mass:**  3277.4512

(9.861e-07)

**Charge States:** 4, 5

**Start Position**: 65

**End Position**: 80

| Profile Score | Tandem Score | TDA q-value | Volume | Spectrum Matches |
| --- | --- | --- | --- | --- |
| 10.3389 | 6.3393 | 0.0342 | 9.8035e+07 | 5 |

**Adduct Signal Fractions**

- **Unmodified**: 9.8035e+07

#### QTRAGC(C)LIGAEHVN(N)NSYEC(C)DIPIGAGIC(C)ASY Fuc 1 Hex 3 HexNAc 6

**Mass:**  5246.2276

(-4.887e-07)

**Charge States:** 4, 5, 6

**Start Position**: 644

**End Position**: 675

| Profile Score | Tandem Score | TDA q-value | Volume | Spectrum Matches |
| --- | --- | --- | --- | --- |
| 14.8826 | 6.3372 | 0.0342 | 6.5155e+09 | 8 |

**Adduct Signal Fractions**

- **Unmodified**: 6.5155e+09

#### VPAQEKN(N)FTTAPAIC(C)HDGKAHFPREGVF Fuc 1 Hex 7 HexNAc 6

**Mass:**  5622.4409

(2.020e-07)

**Charge States:** 4, 5, 6

**Start Position**: 1068

**End Position**: 1096

| Profile Score | Tandem Score | TDA q-value | Volume | Spectrum Matches |
| --- | --- | --- | --- | --- |
| 18.2417 | 6.3312 | 0.0342 | 2.0808e+09 | 2 |

**Adduct Signal Fractions**

- **Unmodified**: 2.0808e+09

#### Q(P)TRAGC(C)LIGAEHVN(N)NSYEC(C)DIPIGAGIC(C)ASY Fuc 1 Hex 5 HexNAc 4 Neu5Ac 1

**Mass:**  5438.2415

(-2.478e-08)

**Charge States:** 4, 5

**Start Position**: 644

**End Position**: 675

| Profile Score | Tandem Score | TDA q-value | Volume | Spectrum Matches |
| --- | --- | --- | --- | --- |
| 16.6567 | 6.3013 | 0.0342 | 1.0688e+09 | 3 |

**Adduct Signal Fractions**

- **Unmodified**: 1.0688e+09

#### IVNN(N)ATNVVIKVC(C)EF Hex 5 HexNAc 4 Neu5Ac 1

**Mass:**  3632.5780

(-9.144e-08)

**Charge States:** 3, 4

**Start Position**: 119

**End Position**: 134

| Profile Score | Tandem Score | TDA q-value | Volume | Spectrum Matches |
| --- | --- | --- | --- | --- |
| 15.1225 | 6.2817 | 0.0342 | 4.9206e+09 | 14 |

**Adduct Signal Fractions**

- **Unmodified**: 4.9206e+09

#### QTRAGC(C)LIGAEHVN(N)NSYEC(C)DIPIGAGIC(C)ASY Fuc 1 Hex 4 HexNAc 6

**Mass:**  5408.2821

(7.868e-07)

**Charge States:** 4, 5

**Start Position**: 644

**End Position**: 675

| Profile Score | Tandem Score | TDA q-value | Volume | Spectrum Matches |
| --- | --- | --- | --- | --- |
| 14.5491 | 6.2443 | 0.0342 | 7.9450e+09 | 6 |

**Adduct Signal Fractions**

- **Unmodified**: 7.9450e+09

#### VPAQEKN(N)FTTAPAIC(C)HDGKAHFPREGVF Hex 5 HexNAc 2

**Mass:**  4339.9605

(7.404e-11)

**Charge States:** 4, 5, 6

**Start Position**: 1068

**End Position**: 1096

| Profile Score | Tandem Score | TDA q-value | Volume | Spectrum Matches |
| --- | --- | --- | --- | --- |
| 16.3314 | 6.2060 | 0.0345 | 6.6934e+09 | 8 |

**Adduct Signal Fractions**

- **Unmodified**: 6.6934e+09

#### HAIHVSGTN(N)GTKRF Fuc 2 Hex 4 HexNAc 4 Neu5Ac 1

**Mass:**  3567.5324

(4.322e-06)

**Charge States:** 4

**Start Position**: 66

**End Position**: 80

| Profile Score | Tandem Score | TDA q-value | Volume | Spectrum Matches |
| --- | --- | --- | --- | --- |
| 11.3029 | 6.2026 | 0.0345 | 1.8155e+07 | 1 |

**Adduct Signal Fractions**

- **Unmodified**: 1.8155e+07

#### IVNN(N)ATNVVIKVC(C)EF Fuc 2 Hex 4 HexNAc 4

**Mass:**  3471.5508

(7.700e-07)

**Charge States:** 3

**Start Position**: 119

**End Position**: 134

| Profile Score | Tandem Score | TDA q-value | Volume | Spectrum Matches |
| --- | --- | --- | --- | --- |
| 13.8468 | 6.1922 | 0.0345 | 2.1713e+09 | 3 |

**Adduct Signal Fractions**

- **Unmodified**: 2.1713e+09

#### RVQPTESIVRFPN(N)ITNL Fuc 1 Hex 5 HexNAc 4 Neu5Ac 2

**Mass:**  4333.9246

(-5.579e-07)

**Charge States:** 3, 4, 5

**Start Position**: 319

**End Position**: 336

| Profile Score | Tandem Score | TDA q-value | Volume | Spectrum Matches |
| --- | --- | --- | --- | --- |
| 15.0082 | 6.1873 | 0.0345 | 8.1355e+09 | 7 |

**Adduct Signal Fractions**

- **Unmodified**: 8.1355e+09

#### YHKNN(N)KSWMESEF Fuc 1 Hex 4 HexNAc 5

**Mass:**  3508.4081

(1.838e-07)

**Charge States:** 3, 4

**Start Position**: 145

**End Position**: 158

| Profile Score | Tandem Score | TDA q-value | Volume | Spectrum Matches |
| --- | --- | --- | --- | --- |
| 15.9984 | 6.1840 | 0.0345 | 3.3618e+10 | 16 |

**Adduct Signal Fractions**

- **Unmodified**: 3.3618e+10

#### RVQPTESIVRFPN(N)ITNLC(C)PFGEVF Fuc 1 Hex 5 HexNAc 5 Neu5Ac 1

**Mass:**  5082.2536

(1.109e-07)

**Charge States:** 4, 5

**Start Position**: 319

**End Position**: 343

| Profile Score | Tandem Score | TDA q-value | Volume | Spectrum Matches |
| --- | --- | --- | --- | --- |
| 15.6360 | 6.1644 | 0.0345 | 1.1102e+10 | 9 |

**Adduct Signal Fractions**

- **Unmodified**: 1.1102e+10

#### IVNN(N)ATNVVIKVC(C)EF Fuc 1 Hex 5 HexNAc 4 Neu5Ac 1

**Mass:**  3778.6361

(-7.251e-07)

**Charge States:** 3, 4

**Start Position**: 119

**End Position**: 134

| Profile Score | Tandem Score | TDA q-value | Volume | Spectrum Matches |
| --- | --- | --- | --- | --- |
| 15.9573 | 6.1226 | 0.0345 | 1.0280e+10 | 13 |

**Adduct Signal Fractions**

- **Unmodified**: 1.0280e+10

#### RVQPTESIVRFPN(N)ITNLC(C)PFGEVF Fuc 2 Hex 5 HexNAc 5

**Mass:**  4937.2192

(1.063e-07)

**Charge States:** 4, 5

**Start Position**: 319

**End Position**: 343

| Profile Score | Tandem Score | TDA q-value | Volume | Spectrum Matches |
| --- | --- | --- | --- | --- |
| 16.4228 | 6.1213 | 0.0345 | 2.1308e+09 | 4 |

**Adduct Signal Fractions**

- **Unmodified**: 2.1308e+09

#### RVQPTESIVRFPN(N)ITNLC(C)PFGEVF Fuc 1 Hex 5 HexNAc 4 Neu5Ac 2

**Mass:**  5170.2752

(2.659e-07)

**Charge States:** 4, 5, 6

**Start Position**: 319

**End Position**: 343

| Profile Score | Tandem Score | TDA q-value | Volume | Spectrum Matches |
| --- | --- | --- | --- | --- |
| 18.4087 | 6.1144 | 0.0345 | 1.0005e+10 | 7 |

**Adduct Signal Fractions**

- **Unmodified**: 1.0005e+10

#### RVQPTESIVRFPN(N)ITNLC(C)PFGEVF Fuc 1 Hex 3 HexNAc 6

**Mass:**  4670.1328

(-8.556e-07)

**Charge States:** 3, 4

**Start Position**: 319

**End Position**: 343

| Profile Score | Tandem Score | TDA q-value | Volume | Spectrum Matches |
| --- | --- | --- | --- | --- |
| 13.3285 | 6.0892 | 0.0345 | 1.4479e+10 | 6 |

**Adduct Signal Fractions**

- **Unmodified**: 1.4479e+10

#### HAIHVSGTN(N)GTKRF Fuc 2 Hex 6 HexNAc 7 Neu5Ac 1

**Mass:**  4500.8840

(-3.070e-07)

**Charge States:** 4, 5

**Start Position**: 66

**End Position**: 80

| Profile Score | Tandem Score | TDA q-value | Volume | Spectrum Matches |
| --- | --- | --- | --- | --- |
| 10.0889 | 6.0867 | 0.0345 | 4.5691e+08 | 13 |

**Adduct Signal Fractions**

- **Unmodified**: 4.5691e+08

#### RVQPTESIVRFPN(N)ITNL Fuc 1 Hex 6 HexNAc 5 Neu5Ac 1

**Mass:**  4407.9589

(8.360e-07)

**Charge States:** 3, 4

**Start Position**: 319

**End Position**: 336

| Profile Score | Tandem Score | TDA q-value | Volume | Spectrum Matches |
| --- | --- | --- | --- | --- |
| 14.1122 | 6.0837 | 0.0347 | 8.2678e+09 | 6 |

**Adduct Signal Fractions**

- **Unmodified**: 8.2678e+09

#### HAIHVSGTN(N)GTKRFDNPVLPF @sulfate 1 Hex 7 HexNAc 2

**Mass:**  3926.7091

(7.942e-06)

**Charge States:** 3, 4, 5

**Start Position**: 66

**End Position**: 87

| Profile Score | Tandem Score | TDA q-value | Volume | Spectrum Matches |
| --- | --- | --- | --- | --- |
| 13.4321 | 6.0764 | 0.0347 | 2.1525e+09 | 10 |

**Adduct Signal Fractions**

- **Unmodified**: 2.1525e+09

#### HAIHVSGTN(N)GTKRF Fuc 1 Hex 5 HexNAc 7

**Mass:**  3901.6792

(-1.326e-06)

**Charge States:** 4, 5

**Start Position**: 66

**End Position**: 80

| Profile Score | Tandem Score | TDA q-value | Volume | Spectrum Matches |
| --- | --- | --- | --- | --- |
| 14.9518 | 6.0651 | 0.0347 | 2.0328e+08 | 3 |

**Adduct Signal Fractions**

- **Unmodified**: 2.0328e+08

#### LIVNN(N)ATNVVIKVC(C)EF Fuc 1 Hex 5 HexNAc 3 Neu5Ac 1

**Mass:**  3688.6463

(4.775e-07)

**Charge States:** 3, 4

**Start Position**: 118

**End Position**: 134

| Profile Score | Tandem Score | TDA q-value | Volume | Spectrum Matches |
| --- | --- | --- | --- | --- |
| 10.5429 | 6.0529 | 0.0347 | 2.5760e+09 | 6 |

**Adduct Signal Fractions**

- **Unmodified**: 2.5760e+09

#### SN(N)NSIAIPTN(N)F Fuc 1 Hex 9 HexNAc 7

**Mass:**  4202.6685

(8.030e-07)

**Charge States:** 3, 4

**Start Position**: 708

**End Position**: 719

| Profile Score | Tandem Score | TDA q-value | Volume | Spectrum Matches |
| --- | --- | --- | --- | --- |
| 13.6829 | 5.9524 | 0.0373 | 7.1640e+08 | 4 |

**Adduct Signal Fractions**

- **Unmodified**: 7.1640e+08

#### HAIHVSGTN(N)GTKRF @sulfate 1 Fuc 1 Hex 5 HexNAc 4 Neu5Ac 1

**Mass:**  3663.4898

(1.616e-07)

**Charge States:** 4

**Start Position**: 66

**End Position**: 80

| Profile Score | Tandem Score | TDA q-value | Volume | Spectrum Matches |
| --- | --- | --- | --- | --- |
| 12.5895 | 5.9450 | 0.0380 | 1.1005e+08 | 2 |

**Adduct Signal Fractions**

- **Unmodified**: 1.1005e+08

#### YHKNN(N)KSWMESEF Fuc 1 Hex 5 HexNAc 4 Neu5Ac 1

**Mass:**  3758.4807

(3.693e-08)

**Charge States:** 3, 4

**Start Position**: 145

**End Position**: 158

| Profile Score | Tandem Score | TDA q-value | Volume | Spectrum Matches |
| --- | --- | --- | --- | --- |
| 15.5911 | 5.8856 | 0.0419 | 6.0813e+09 | 6 |

**Adduct Signal Fractions**

- **Unmodified**: 6.0813e+09

#### YHKNN(N)KSWMESEF Fuc 2 Hex 4 HexNAc 5

**Mass:**  3654.4673

(-2.938e-08)

**Charge States:** 3, 4

**Start Position**: 145

**End Position**: 158

| Profile Score | Tandem Score | TDA q-value | Volume | Spectrum Matches |
| --- | --- | --- | --- | --- |
| 16.4867 | 5.8537 | 0.0419 | 9.6048e+09 | 5 |

**Adduct Signal Fractions**

- **Unmodified**: 9.6048e+09

#### YHKNN(N)KSWMESEF Fuc 1 Hex 4 HexNAc 3

**Mass:**  3102.2524

(-3.006e-07)

**Charge States:** 2, 3, 4

**Start Position**: 145

**End Position**: 158

| Profile Score | Tandem Score | TDA q-value | Volume | Spectrum Matches |
| --- | --- | --- | --- | --- |
| 16.0336 | 5.8492 | 0.0419 | 1.4920e+09 | 4 |

**Adduct Signal Fractions**

- **Unmodified**: 1.4920e+09

#### LIVNN(N)ATNVVIKVC(C)EF Fuc 1 Hex 4 HexNAc 4 Neu5Ac 1

**Mass:**  3729.6700

(1.284e-06)

**Charge States:** 3, 4

**Start Position**: 118

**End Position**: 134

| Profile Score | Tandem Score | TDA q-value | Volume | Spectrum Matches |
| --- | --- | --- | --- | --- |
| 13.3080 | 5.7578 | 0.0419 | 6.3350e+09 | 8 |

**Adduct Signal Fractions**

- **Unmodified**: 6.3350e+09

#### RVQPTESIVRFPN(N)ITNL Fuc 1 Hex 4 HexNAc 5

**Mass:**  3792.7588

(3.877e-07)

**Charge States:** 3, 4

**Start Position**: 319

**End Position**: 336

| Profile Score | Tandem Score | TDA q-value | Volume | Spectrum Matches |
| --- | --- | --- | --- | --- |
| 12.6926 | 5.7301 | 0.0419 | 1.5278e+10 | 16 |

**Adduct Signal Fractions**

- **Unmodified**: 1.5278e+10

#### HAIHVSGTN(N)GTKRF Fuc 1 Hex 7 HexNAc 3

**Mass:**  3413.4588

(-2.607e-07)

**Charge States:** 4

**Start Position**: 66

**End Position**: 80

| Profile Score | Tandem Score | TDA q-value | Volume | Spectrum Matches |
| --- | --- | --- | --- | --- |
| 11.8024 | 5.7234 | 0.0419 | 2.3208e+07 | 4 |

**Adduct Signal Fractions**

- **Unmodified**: 2.3208e+07

#### HAIHVSGTN(N)GTKRF Fuc 2 Hex 6 HexNAc 4

**Mass:**  3603.5822

(8.389e-04)

**Charge States:** 3

**Start Position**: 66

**End Position**: 80

| Profile Score | Tandem Score | TDA q-value | Volume | Spectrum Matches |
| --- | --- | --- | --- | --- |
| 7.3420 | 5.7079 | 0.0419 | 4.4973e+07 | 1 |

**Adduct Signal Fractions**

- **Unmodified**: 4.4973e+07

#### QTRAGC(C)LIGAEHVN(N)NSYEC(C)DIPIGAGIC(C)ASY Fuc 1 Hex 7 HexNAc 6 Neu5Ac 3

**Mass:**  6767.7204

(-3.565e-07)

**Charge States:** 5, 6

**Start Position**: 644

**End Position**: 675

| Profile Score | Tandem Score | TDA q-value | Volume | Spectrum Matches |
| --- | --- | --- | --- | --- |
| 13.3435 | 5.6918 | 0.0419 | 4.7804e+09 | 5 |

**Adduct Signal Fractions**

- **Unmodified**: 4.7804e+09

#### LIVNN(N)ATNVVIKVC(C)EF Fuc 1 Hex 4 HexNAc 3 Neu5Ac 1

**Mass:**  3526.5911

(4.291e-07)

**Charge States:** 3, 4

**Start Position**: 118

**End Position**: 134

| Profile Score | Tandem Score | TDA q-value | Volume | Spectrum Matches |
| --- | --- | --- | --- | --- |
| 15.7021 | 5.6744 | 0.0419 | 5.0983e+08 | 3 |

**Adduct Signal Fractions**

- **Unmodified**: 5.0983e+08

#### RVQPTESIVRFPN(N)ITNLC(C)PF Fuc 2 Hex 4 HexNAc 4

**Mass:**  4139.8897

(3.953e-07)

**Charge States:** 3, 4

**Start Position**: 319

**End Position**: 339

| Profile Score | Tandem Score | TDA q-value | Volume | Spectrum Matches |
| --- | --- | --- | --- | --- |
| 14.7632 | 5.6729 | 0.0419 | 2.1153e+09 | 7 |

**Adduct Signal Fractions**

- **Unmodified**: 2.1153e+09

#### QTRAGC(C)LIGAEHVN(N)NSYEC(C)DIPIGAGIC(C)ASY Fuc 1 Hex 7 HexNAc 6 Neu5Ac 2

**Mass:**  6476.6248

(-2.935e-07)

**Charge States:** 5, 6

**Start Position**: 644

**End Position**: 675

| Profile Score | Tandem Score | TDA q-value | Volume | Spectrum Matches |
| --- | --- | --- | --- | --- |
| 14.2208 | 5.6620 | 0.0419 | 4.0837e+09 | 5 |

**Adduct Signal Fractions**

- **Unmodified**: 4.0837e+09

#### FHAIHVSGTN(N)GTKRF Fuc 2 Hex 5 HexNAc 4

**Mass:**  3585.5633

(-1.044e-07)

**Charge States:** 4, 5

**Start Position**: 65

**End Position**: 80

| Profile Score | Tandem Score | TDA q-value | Volume | Spectrum Matches |
| --- | --- | --- | --- | --- |
| 12.8579 | 5.6246 | 0.0419 | 4.6774e+07 | 3 |

**Adduct Signal Fractions**

- **Unmodified**: 4.6774e+07

#### FHAIHVSGTN(N)GTKRF Fuc 1 Hex 4 HexNAc 7

**Mass:**  3886.6903

(5.184e-07)

**Charge States:** 4, 5

**Start Position**: 65

**End Position**: 80

| Profile Score | Tandem Score | TDA q-value | Volume | Spectrum Matches |
| --- | --- | --- | --- | --- |
| 12.9520 | 5.6211 | 0.0419 | 9.6669e+07 | 1 |

**Adduct Signal Fractions**

- **Unmodified**: 9.6669e+07

#### RVQPTESIVRFPN(N)ITNLC(C)PF Fuc 1 Hex 6 HexNAc 5 Neu5Ac 2

**Mass:**  5103.2047

(3.562e-08)

**Charge States:** 4, 5

**Start Position**: 319

**End Position**: 339

| Profile Score | Tandem Score | TDA q-value | Volume | Spectrum Matches |
| --- | --- | --- | --- | --- |
| 12.7337 | 5.5910 | 0.0419 | 4.1102e+09 | 8 |

**Adduct Signal Fractions**

- **Unmodified**: 4.1102e+09

#### EPLVDLPIGIN(N)ITRF Fuc 1 Hex 5 HexNAc 5

**Mass:**  3670.6949

(8.249e-04)

**Charge States:** 4

**Start Position**: 224

**End Position**: 239

| Profile Score | Tandem Score | TDA q-value | Volume | Spectrum Matches |
| --- | --- | --- | --- | --- |
| -7.1881 | 5.5768 | 0.0419 | 1.4993e+07 | 1 |

**Adduct Signal Fractions**

- **Unmodified**: 1.4993e+07

#### IGAEHVN(N)NSYEC(C)DIPIGAGIC(C)ASY Fuc 1 Hex 3 HexNAc 4

**Mass:**  4053.6880

(7.158e-08)

**Charge States:** 3, 4

**Start Position**: 651

**End Position**: 675

| Profile Score | Tandem Score | TDA q-value | Volume | Spectrum Matches |
| --- | --- | --- | --- | --- |
| 16.9762 | 5.5313 | 0.0419 | 2.7945e+09 | 4 |

**Adduct Signal Fractions**

- **Unmodified**: 2.7945e+09

#### VSGNC(C)DVVIGIVN(N)NTVY Fuc 1 Hex 3 HexNAc 4

**Mass:**  3266.4286

(2.183e-07)

**Charge States:** 3, 4

**Start Position**: 1122

**End Position**: 1139

| Profile Score | Tandem Score | TDA q-value | Volume | Spectrum Matches |
| --- | --- | --- | --- | --- |
| 16.4675 | 5.5186 | 0.0419 | 3.0117e+09 | 3 |

**Adduct Signal Fractions**

- **Unmodified**: 3.0117e+09

#### RVQPTESIVRFPN(N)ITNLC(C)PF Fuc 1 Hex 5 HexNAc 5 Neu5Ac 1

**Mass:**  4650.0544

(-2.657e-07)

**Charge States:** 3, 4, 5

**Start Position**: 319

**End Position**: 339

| Profile Score | Tandem Score | TDA q-value | Volume | Spectrum Matches |
| --- | --- | --- | --- | --- |
| 11.4109 | 5.4946 | 0.0419 | 1.0454e+10 | 9 |

**Adduct Signal Fractions**

- **Unmodified**: 1.0454e+10

#### GGFN(N)FSQILPDPSKPSKRSF Hex 3 HexNAc 3

**Mass:**  3303.5318

(4.602e-07)

**Charge States:** 4

**Start Position**: 798

**End Position**: 818

| Profile Score | Tandem Score | TDA q-value | Volume | Spectrum Matches |
| --- | --- | --- | --- | --- |
| 10.8868 | 5.4727 | 0.0419 | 2.8808e+08 | 3 |

**Adduct Signal Fractions**

- **Unmodified**: 2.8808e+08

#### IGAEHVN(N)NSYEC(C)DIPIGAGIC(C)ASY Fuc 1 Hex 3 HexNAc 5

**Mass:**  4256.7679

(4.788e-07)

**Charge States:** 3, 4

**Start Position**: 651

**End Position**: 675

| Profile Score | Tandem Score | TDA q-value | Volume | Spectrum Matches |
| --- | --- | --- | --- | --- |
| 16.1199 | 5.4470 | 0.0419 | 2.8277e+09 | 6 |

**Adduct Signal Fractions**

- **Unmodified**: 2.8277e+09

#### VPAQEKN(N)FTTAPAIC(C)HDGKAHFPREGVF Fuc 1 Hex 4 HexNAc 6

**Mass:**  5136.2785

(-9.812e-07)

**Charge States:** 4, 5, 6

**Start Position**: 1068

**End Position**: 1096

| Profile Score | Tandem Score | TDA q-value | Volume | Spectrum Matches |
| --- | --- | --- | --- | --- |
| 14.7619 | 5.3931 | 0.0419 | 3.7744e+09 | 7 |

**Adduct Signal Fractions**

- **Unmodified**: 3.7744e+09

#### SALEPLVDLPIGIN(N)ITRF Fuc 1 Hex 4 HexNAc 3

**Mass:**  3370.6175

(5.045e-08)

**Charge States:** 3

**Start Position**: 221

**End Position**: 239

| Profile Score | Tandem Score | TDA q-value | Volume | Spectrum Matches |
| --- | --- | --- | --- | --- |
| 12.9182 | 5.3928 | 0.0419 | 1.2287e+09 | 5 |

**Adduct Signal Fractions**

- **Unmodified**: 1.2287e+09

#### N(N)FSQILPDPSKPSKRSF Fuc 1 Hex 4 HexNAc 5

**Mass:**  3756.6920

(1.169e-06)

**Charge States:** 3, 4

**Start Position**: 801

**End Position**: 818

| Profile Score | Tandem Score | TDA q-value | Volume | Spectrum Matches |
| --- | --- | --- | --- | --- |
| 13.6033 | 5.3791 | 0.0419 | 4.8570e+08 | 4 |

**Adduct Signal Fractions**

- **Unmodified**: 4.8570e+08

#### N(N)FSQILPDPSKPSKRSF Fuc 1 Hex 4 HexNAc 4 Neu5Ac 1

**Mass:**  3844.7042

(-7.511e-08)

**Charge States:** 3, 4

**Start Position**: 801

**End Position**: 818

| Profile Score | Tandem Score | TDA q-value | Volume | Spectrum Matches |
| --- | --- | --- | --- | --- |
| 12.1466 | 5.3714 | 0.0419 | 6.8070e+09 | 9 |

**Adduct Signal Fractions**

- **Unmodified**: 6.8070e+09

#### VPAQEKN(N)FTTAPAIC(C)HDGKAHFPREGVF Hex 5 HexNAc 3

**Mass:**  4543.0414

(4.693e-07)

**Charge States:** 4, 5, 6

**Start Position**: 1068

**End Position**: 1096

| Profile Score | Tandem Score | TDA q-value | Volume | Spectrum Matches |
| --- | --- | --- | --- | --- |
| 13.2851 | 5.3586 | 0.0419 | 5.7154e+08 | 4 |

**Adduct Signal Fractions**

- **Unmodified**: 5.7154e+08

#### RVQPTESIVRFPN(N)ITNLC(C)PFGEVF Fuc 1 Hex 4 HexNAc 6

**Mass:**  4832.1900

(1.959e-08)

**Charge States:** 4, 5

**Start Position**: 319

**End Position**: 343

| Profile Score | Tandem Score | TDA q-value | Volume | Spectrum Matches |
| --- | --- | --- | --- | --- |
| 13.5276 | 5.3559 | 0.0419 | 2.4860e+09 | 4 |

**Adduct Signal Fractions**

- **Unmodified**: 2.4860e+09

#### GGFN(N)FSQILPDPSKPSKRSF Hex 8 HexNAc 2

**Mass:**  3910.7175

(1.455e-06)

**Charge States:** 4

**Start Position**: 798

**End Position**: 818

| Profile Score | Tandem Score | TDA q-value | Volume | Spectrum Matches |
| --- | --- | --- | --- | --- |
| 11.4207 | 5.3396 | 0.0425 | 7.6216e+08 | 3 |

**Adduct Signal Fractions**

- **Unmodified**: 7.6216e+08

#### HKNN(N)KSWMESEF Hex 5 HexNAc 2

**Mass:**  2752.1071

(5.256e-07)

**Charge States:** 2, 3

**Start Position**: 146

**End Position**: 158

| Profile Score | Tandem Score | TDA q-value | Volume | Spectrum Matches |
| --- | --- | --- | --- | --- |
| 13.7782 | 5.3245 | 0.0426 | 2.9055e+09 | 3 |

**Adduct Signal Fractions**

- **Unmodified**: 2.9055e+09

#### HKNN(N)KSWMESEF Fuc 1 Hex 5 HexNAc 4

**Mass:**  3304.3235

(5.452e-07)

**Charge States:** 3, 4

**Start Position**: 146

**End Position**: 158

| Profile Score | Tandem Score | TDA q-value | Volume | Spectrum Matches |
| --- | --- | --- | --- | --- |
| 14.7564 | 5.3200 | 0.0426 | 1.3624e+09 | 2 |

**Adduct Signal Fractions**

- **Unmodified**: 1.3624e+09

#### VPAQEKN(N)FTTAPAIC(C)HDGKAHFPREGVF Fuc 1 Hex 5 HexNAc 5 Neu5Ac 1

**Mass:**  5388.4193

(3.736e-04)

**Charge States:** 4

**Start Position**: 1068

**End Position**: 1096

| Profile Score | Tandem Score | TDA q-value | Volume | Spectrum Matches |
| --- | --- | --- | --- | --- |
| -5.9740 | 5.2829 | 0.0426 | 9.5140e+06 | 2 |

**Adduct Signal Fractions**

- **Unmodified**: 9.5140e+06

#### FHAIHVSGTN(N)GTKRF @sulfate 1 Fuc 1 Hex 3 HexNAc 6

**Mass:**  3601.5151

(-1.340e-06)

**Charge States:** 4

**Start Position**: 65

**End Position**: 80

| Profile Score | Tandem Score | TDA q-value | Volume | Spectrum Matches |
| --- | --- | --- | --- | --- |
| 10.5097 | 5.2516 | 0.0426 | 1.1814e+08 | 3 |

**Adduct Signal Fractions**

- **Unmodified**: 1.1814e+08

#### IGAEHVN(N)NSYEC(C)DIPIGAGIC(C)ASY Fuc 1 Hex 4 HexNAc 5

**Mass:**  4418.8178

(-1.083e-07)

**Charge States:** 3, 4

**Start Position**: 651

**End Position**: 675

| Profile Score | Tandem Score | TDA q-value | Volume | Spectrum Matches |
| --- | --- | --- | --- | --- |
| 14.5139 | 5.2354 | 0.0426 | 1.0484e+09 | 3 |

**Adduct Signal Fractions**

- **Unmodified**: 1.0484e+09

#### VPAQEKN(N)FTTAPAIC(C)HDGKAHFPREGVF Hex 4 HexNAc 3

**Mass:**  4380.9828

(2.441e-07)

**Charge States:** 4, 5, 6

**Start Position**: 1068

**End Position**: 1096

| Profile Score | Tandem Score | TDA q-value | Volume | Spectrum Matches |
| --- | --- | --- | --- | --- |
| 11.2032 | 5.2260 | 0.0426 | 2.3521e+08 | 3 |

**Adduct Signal Fractions**

- **Unmodified**: 2.3521e+08

#### SSANN(N)C(C)TFEYVSQPF Hex 5 HexNAc 2

**Mass:**  2966.1514

(-6.382e-07)

**Charge States:** 2, 3

**Start Position**: 161

**End Position**: 176

| Profile Score | Tandem Score | TDA q-value | Volume | Spectrum Matches |
| --- | --- | --- | --- | --- |
| 17.7314 | 5.2178 | 0.0426 | 4.6464e+09 | 6 |

**Adduct Signal Fractions**

- **Unmodified**: 4.6464e+09

#### SNNSIAIPTN(N)F Fuc 2 Hex 6 HexNAc 6 Neu5Ac 1

**Mass:**  3950.5557

(-6.161e-06)

**Charge States:** 3

**Start Position**: 708

**End Position**: 719

| Profile Score | Tandem Score | TDA q-value | Volume | Spectrum Matches |
| --- | --- | --- | --- | --- |
| 13.6290 | 5.1492 | 0.0426 | 5.1486e+08 | 1 |

**Adduct Signal Fractions**

- **Unmodified**: 5.1486e+08

#### HAIHVSGTN(N)GTKRF @sulfate 1 Fuc 2 Hex 5 HexNAc 6 Neu5Ac 1

**Mass:**  4215.7010

(-5.980e-07)

**Charge States:** 4

**Start Position**: 66

**End Position**: 80

| Profile Score | Tandem Score | TDA q-value | Volume | Spectrum Matches |
| --- | --- | --- | --- | --- |
| 9.6285 | 5.1389 | 0.0426 | 5.0559e+08 | 5 |

**Adduct Signal Fractions**

- **Unmodified**: 5.0559e+08

#### HAIHVSGTN(N)GTKRF @sulfate 1 Hex 5 HexNAc 4 Neu5Ac 2

**Mass:**  3808.5615

(9.587e-06)

**Charge States:** 5

**Start Position**: 66

**End Position**: 80

| Profile Score | Tandem Score | TDA q-value | Volume | Spectrum Matches |
| --- | --- | --- | --- | --- |
| 12.1132 | 5.1270 | 0.0431 | 6.7562e+07 | 1 |

**Adduct Signal Fractions**

- **Unmodified**: 6.7562e+07

#### RVQPTESIVRFPN(N)ITNL Fuc 2 Hex 5 HexNAc 5

**Mass:**  4100.8671

(-5.645e-07)

**Charge States:** 3, 4

**Start Position**: 319

**End Position**: 336

| Profile Score | Tandem Score | TDA q-value | Volume | Spectrum Matches |
| --- | --- | --- | --- | --- |
| 11.5861 | 5.1258 | 0.0431 | 6.7160e+08 | 3 |

**Adduct Signal Fractions**

- **Unmodified**: 6.7160e+08

#### HVTYVPAQEKN(N)F Fuc 1 Hex 3 HexNAc 4

**Mass:**  2878.2547

(6.975e-04)

**Start Position**: 1064

**End Position**: 1076

| Profile Score | Tandem Score | TDA q-value | Volume | Spectrum Matches |
| --- | --- | --- | --- | --- |
| 0.0000 | 5.1149 | 0.0431 | 0.0000e+00 | 1 |

Adduct:

No Chromatogram Found

#### HVTYVPAQEKN(N)F Fuc 1 Hex 3 HexNAc 4

**Mass:**  2876.2487

(1.304e-08)

**Charge States:** 3

**Start Position**: 1064

**End Position**: 1076

| Profile Score | Tandem Score | TDA q-value | Volume | Spectrum Matches |
| --- | --- | --- | --- | --- |
| 12.0466 | 5.0948 | 0.0433 | 4.7775e+08 | 2 |

**Adduct Signal Fractions**

- **Unmodified**: 4.7775e+08

#### HAIHVSGTN(N)GTKRF Fuc 1 Hex 7 HexNAc 6 Neu5Ac 1

**Mass:**  4313.8029

(1.565e-06)

**Charge States:** 4, 5

**Start Position**: 66

**End Position**: 80

| Profile Score | Tandem Score | TDA q-value | Volume | Spectrum Matches |
| --- | --- | --- | --- | --- |
| 11.1910 | 5.0855 | 0.0433 | 4.2543e+08 | 9 |

**Adduct Signal Fractions**

- **Unmodified**: 4.2543e+08

#### HAIHVSGTN(N)GTKRF Fuc 2 Hex 7 HexNAc 7

**Mass:**  4371.8336

(1.323e-06)

**Charge States:** 4, 5

**Start Position**: 66

**End Position**: 80

| Profile Score | Tandem Score | TDA q-value | Volume | Spectrum Matches |
| --- | --- | --- | --- | --- |
| 10.6451 | 5.0822 | 0.0433 | 4.0663e+07 | 2 |

**Adduct Signal Fractions**

- **Unmodified**: 4.0663e+07

#### HAIHVSGTN(N)GTKRF Hex 7 HexNAc 2

**Mass:**  3064.3304

(5.250e-07)

**Charge States:** 4

**Start Position**: 66

**End Position**: 80

| Profile Score | Tandem Score | TDA q-value | Volume | Spectrum Matches |
| --- | --- | --- | --- | --- |
| 10.2349 | 5.0473 | 0.0439 | 8.5662e+07 | 2 |

**Adduct Signal Fractions**

- **Unmodified**: 8.5662e+07

#### YHKNN(N)KSWMESEF Fuc 1 Hex 3 HexNAc 6

**Mass:**  3549.4382

(-8.974e-07)

**Charge States:** 3, 4

**Start Position**: 145

**End Position**: 158

| Profile Score | Tandem Score | TDA q-value | Volume | Spectrum Matches |
| --- | --- | --- | --- | --- |
| 11.1324 | 5.0141 | 0.0443 | 9.0682e+09 | 16 |

**Adduct Signal Fractions**

- **Unmodified**: 9.0682e+09

#### IVNN(N)ATNVVIKVC(C)EF Fuc 2 Hex 4 HexNAc 5

**Mass:**  3674.6279

(6.018e-07)

**Charge States:** 3

**Start Position**: 119

**End Position**: 134

| Profile Score | Tandem Score | TDA q-value | Volume | Spectrum Matches |
| --- | --- | --- | --- | --- |
| 15.9962 | 4.9985 | 0.0452 | 7.2083e+08 | 2 |

**Adduct Signal Fractions**

- **Unmodified**: 7.2083e+08

#### YHKNN(N)KSWMESEF Fuc 1 Hex 5 HexNAc 4 Neu5Ac 2

**Mass:**  4049.5806

(1.045e-06)

**Charge States:** 3, 4

**Start Position**: 145

**End Position**: 158

| Profile Score | Tandem Score | TDA q-value | Volume | Spectrum Matches |
| --- | --- | --- | --- | --- |
| 15.9284 | 4.9768 | 0.0452 | 5.2109e+08 | 1 |

**Adduct Signal Fractions**

- **Unmodified**: 5.2109e+08

#### IVNN(N)ATNVVIKVC(C)EF Fuc 2 Hex 5 HexNAc 4

**Mass:**  3633.6052

(1.011e-06)

**Charge States:** 3

**Start Position**: 119

**End Position**: 134

| Profile Score | Tandem Score | TDA q-value | Volume | Spectrum Matches |
| --- | --- | --- | --- | --- |
| 12.9994 | 4.9753 | 0.0452 | 5.5057e+08 | 2 |

**Adduct Signal Fractions**

- **Unmodified**: 5.5057e+08

#### SN(N)NSIAIPTN(N)F Hex 11 HexNAc 5 Neu5Ac 1

**Mass:**  4265.6520

(3.210e-07)

**Charge States:** 3, 4

**Start Position**: 708

**End Position**: 719

| Profile Score | Tandem Score | TDA q-value | Volume | Spectrum Matches |
| --- | --- | --- | --- | --- |
| 15.8733 | 4.9604 | 0.0452 | 1.9053e+09 | 4 |

**Adduct Signal Fractions**

- **Unmodified**: 1.9053e+09

#### YHKNN(N)KSWMESEF Fuc 1 Hex 4 HexNAc 5 Neu5Ac 1

**Mass:**  3799.5076

(-9.511e-08)

**Charge States:** 3, 4

**Start Position**: 145

**End Position**: 158

| Profile Score | Tandem Score | TDA q-value | Volume | Spectrum Matches |
| --- | --- | --- | --- | --- |
| 13.7651 | 4.9463 | 0.0456 | 7.5360e+09 | 6 |

**Adduct Signal Fractions**

- **Unmodified**: 7.5360e+09

#### HAIHVSGTN(N)GTKRF Fuc 2 Hex 6 HexNAc 5 Neu5Ac 2

**Mass:**  4385.8132

(-6.507e-07)

**Charge States:** 4, 5

**Start Position**: 66

**End Position**: 80

| Profile Score | Tandem Score | TDA q-value | Volume | Spectrum Matches |
| --- | --- | --- | --- | --- |
| 11.0186 | 4.9050 | 0.0468 | 4.0202e+08 | 10 |

**Adduct Signal Fractions**

- **Unmodified**: 4.0202e+08

#### HAIHVSGTN(N)GTKRF @sulfate 1 Fuc 1 Hex 5 HexNAc 5 Neu5Ac 1

**Mass:**  3866.5667

(-5.689e-08)

**Charge States:** 4

**Start Position**: 66

**End Position**: 80

| Profile Score | Tandem Score | TDA q-value | Volume | Spectrum Matches |
| --- | --- | --- | --- | --- |
| 12.5012 | 4.8977 | 0.0468 | 1.0329e+08 | 2 |

**Adduct Signal Fractions**

- **Unmodified**: 1.0329e+08

#### RVQPTESIVRFPN(N)ITNL Fuc 2 Hex 4 HexNAc 4

**Mass:**  3735.7370

(6.504e-07)

**Charge States:** 3, 4

**Start Position**: 319

**End Position**: 336

| Profile Score | Tandem Score | TDA q-value | Volume | Spectrum Matches |
| --- | --- | --- | --- | --- |
| 11.0021 | 4.8830 | 0.0468 | 1.0493e+09 | 6 |

**Adduct Signal Fractions**

- **Unmodified**: 1.0493e+09

#### LIVNN(N)ATNVVIKVC(C)EF Fuc 1 Hex 5 HexNAc 3

**Mass:**  3397.5463

(5.117e-07)

**Charge States:** 3

**Start Position**: 118

**End Position**: 134

| Profile Score | Tandem Score | TDA q-value | Volume | Spectrum Matches |
| --- | --- | --- | --- | --- |
| 12.9587 | 4.8676 | 0.0468 | 1.2153e+09 | 3 |

**Adduct Signal Fractions**

- **Unmodified**: 1.2153e+09

#### Q(P)TRAGC(C)LIGAEHVN(N)NSYEC(C)DIPIGAGIC(C)ASY Fuc 1 Hex 4 HexNAc 4

**Mass:**  4985.0932

(2.048e-07)

**Charge States:** 4, 5

**Start Position**: 644

**End Position**: 675

| Profile Score | Tandem Score | TDA q-value | Volume | Spectrum Matches |
| --- | --- | --- | --- | --- |
| 14.0627 | 4.8601 | 0.0473 | 2.7238e+08 | 1 |

**Adduct Signal Fractions**

- **Unmodified**: 2.7238e+08

#### LIVNN(N)ATNVVIKVC(C)EF Hex 4 HexNAc 3

**Mass:**  3089.4386

(1.025e-06)

**Charge States:** 2, 3

**Start Position**: 118

**End Position**: 134

| Profile Score | Tandem Score | TDA q-value | Volume | Spectrum Matches |
| --- | --- | --- | --- | --- |
| 15.5315 | 4.8460 | 0.0479 | 8.2613e+09 | 4 |

**Adduct Signal Fractions**

- **Unmodified**: 8.2613e+09

#### LIVNN(N)ATNVVIKVC(C)EF Hex 5 HexNAc 4 Neu5Ac 1

**Mass:**  3745.6695

(4.867e-07)

**Charge States:** 3

**Start Position**: 118

**End Position**: 134

| Profile Score | Tandem Score | TDA q-value | Volume | Spectrum Matches |
| --- | --- | --- | --- | --- |
| 10.1891 | 4.8379 | 0.0491 | 2.0150e+08 | 3 |

**Adduct Signal Fractions**

- **Unmodified**: 2.0150e+08

#### LIVNN(N)ATNVVIKVC(C)EF Hex 3 HexNAc 5

**Mass:**  3333.5437

(6.843e-07)

**Charge States:** 3, 4

**Start Position**: 118

**End Position**: 134

| Profile Score | Tandem Score | TDA q-value | Volume | Spectrum Matches |
| --- | --- | --- | --- | --- |
| 14.4686 | 4.7967 | 0.0499 | 4.9001e+08 | 3 |

**Adduct Signal Fractions**

- **Unmodified**: 4.9001e+08

#### VPAQEKN(N)FTTAPAIC(C)HDGKAHFPREGVF Fuc 3 Hex 5 HexNAc 4

**Mass:**  5184.2732

(-1.979e-06)

**Charge States:** 4, 5

**Start Position**: 1068

**End Position**: 1096

| Profile Score | Tandem Score | TDA q-value | Volume | Spectrum Matches |
| --- | --- | --- | --- | --- |
| 11.0413 | 4.7655 | 0.0499 | 1.2565e+09 | 4 |

**Adduct Signal Fractions**

- **Unmodified**: 1.2565e+09

#### LIVNN(N)ATNVVIKVC(C)EF Fuc 1 Hex 5 HexNAc 4 Neu5Ac 1

**Mass:**  3891.7250

(8.347e-07)

**Charge States:** 3, 4

**Start Position**: 118

**End Position**: 134

| Profile Score | Tandem Score | TDA q-value | Volume | Spectrum Matches |
| --- | --- | --- | --- | --- |
| 14.5181 | 4.7587 | 0.0499 | 8.2960e+08 | 2 |

**Adduct Signal Fractions**

- **Unmodified**: 8.2960e+08

#### RVQPTESIVRFPN(N)ITNLC(C)PF Fuc 1 Hex 4 HexNAc 6

**Mass:**  4399.9878

(-8.070e-08)

**Charge States:** 3, 4

**Start Position**: 319

**End Position**: 339

| Profile Score | Tandem Score | TDA q-value | Volume | Spectrum Matches |
| --- | --- | --- | --- | --- |
| 15.6359 | 4.7215 | 0.0499 | 1.4626e+09 | 4 |

**Adduct Signal Fractions**

- **Unmodified**: 1.4626e+09

#### VPAQEKN(N)FTTAPAIC(C)HDGKAHFPREGVF Fuc 2 Hex 5 HexNAc 4

**Mass:**  5038.2343

(-6.186e-07)

**Charge States:** 4, 5, 6

**Start Position**: 1068

**End Position**: 1096

| Profile Score | Tandem Score | TDA q-value | Volume | Spectrum Matches |
| --- | --- | --- | --- | --- |
| 13.7862 | 4.6540 | 0.0499 | 6.5803e+08 | 4 |

**Adduct Signal Fractions**

- **Unmodified**: 6.5803e+08
